# Engaging Unstabilized Alkyl Radicals with Pyridoxal Radical Biocatalysis: Enantiodivergent Synthesis of Aliphatic Non-Canonical Amino Acids

**DOI:** 10.64898/2026.01.21.700944

**Authors:** Lei Cheng, Jasper Chen, Zhiyu Bo, Xiangyu Zhang, Peng Liu, Yang Yang

## Abstract

Harnessing transient, unstabilized alkyl radical intermediates for the enantioselective construction of value-added chemical entities remains a fundamental challenge in biocatalysis. Through the repurposing and directed evolution of pyridoxal phosphate (PLP)-dependent tryptophan synthases, we advanced an open-shell enzyme platform capable of intercepting transient alkyl radicals for the efficient and enantioselective synthesis of aliphatic non-canonical amino acids. Engineering an orthogonal pair of radical PLP enzymes allowed unstabilized alkyl radicals, generated from diverse aliphatic organoboronates, to undergo dehydroxylative C(sp^3^)–C(sp^3^) coupling with a common L-serine donor, affording either L- or D-amino acids with excellent enantiopurity in an enzyme-controlled fashion. Mechanistic and computational investigations employing radical clock substrates and unusual radical-mediated rearrangement processes revealed that the radical intermediates generated in this system exhibit unexpectedly long lifetimes, highlighting the power of this dual enzyme-photocatalyst platform to engage unactivated alkyl radicals. Collectively, these findings delineate a potentially general strategy for generating and utilizing unstabilized alkyl radicals and underscore the synthetic potential of radical pyridoxal biocatalysts for stereodivergent amino acid construction.

## Introduction

Anaerobic radical enzymes,^1,2^ including radical S-adenosylmethionine (SAM) enzymes,^3^ cobalamin-dependent enzymes,^4^ and glycyl radical enzymes^5^ among others, harness transient radical intermediates for highly challenging transformations, many of which remain inaccessible to state-of-the-art small-molecule catalysts. For example, lysine 2,3-aminomutase,^6^ methylornithine synthase^7,8^ and fatty acid photodecarboxylase^9^ carry out unimolecular free radical reactions using diverse SAM/PLP, SAM and flavin adenine dinucleotide (FAD)-based mechanisms, respectively (Figure 1a). Despite this fascinating radical enzymology, only a very limited subset of natural radical enzymes could leverage unstabilized alkyl radicals as productive intermediates for intermolecular C–C bond formation.^1-9^ A rare example is provided by glycyl radical enzymes such as (1-methylpentyl)succinate synthase, which employ enzymatic thiyl radicals to generate unstabilized and unfunctionalized alkyl radicals from hydrocarbon substrates for enantioselective C–C bond formation.^10,11^ Their remarkable control over radical reactivity arises from generating these open-shell intermediates within the enzyme’s active site in proximity to both the cofactor and the coupling partner, thereby enabling tightly orchestrated transformations. While this spatial and temporal confinement minimizes radical escape, it inherently limits substrate scope and synthetic versatility.

**Figure 1.**
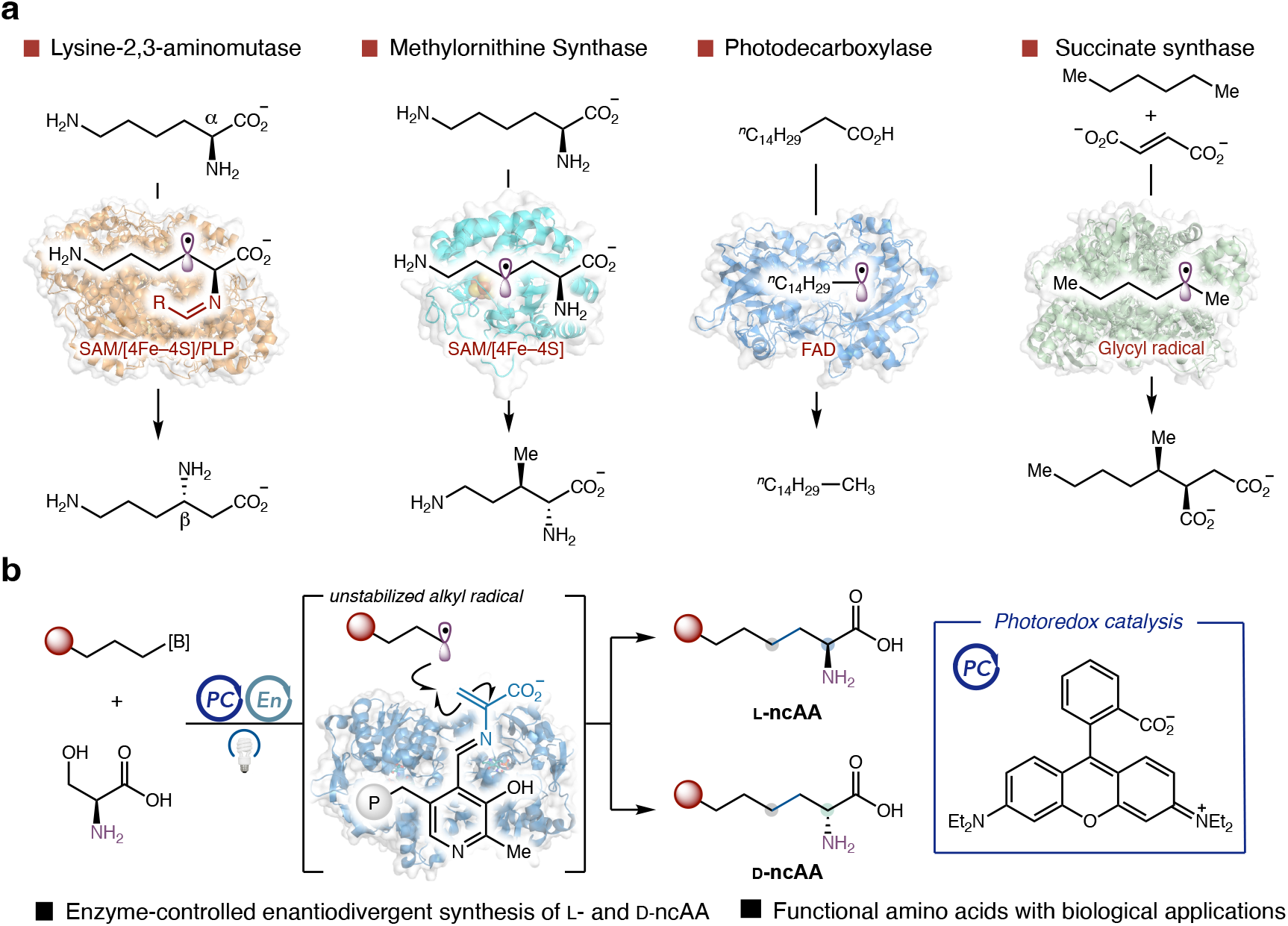
Biocatalytic transformation of unstabilized alkyl radicals. **a**, Representative examples of anaerobic radical enzymes in transforming short-lived alkyl radical species. **b**, Pyridoxal radical biocatalysis engaged short-lived alkyl radicals for aliphatic noncanonical amino acids synthesis.

Over the past decade, efforts to expand the catalytic repertoire of enzymes have led to biocatalytic transformations not found in natural enzymology, including those which are unknown in both organic chemistry and biochemistry.^12-14^ Among these advances, cooperative photobiocatalysis has emerged as a powerful strategy for achieving stereocontrol in otherwise challenging free radical-mediated reactions.^15-18^ Recently, our group and others have demonstrated that organic and metallocofactor-dependent enzymes can be repurposed for intermolecular asymmetric radical transformations involving exogenously generated radical species, through the cooperative activation using a photoredox catalyst and an enzyme—with or without a native redox cofactor.^19-35^ However, to date, the majority of cooperative photobiocatalytic radical reactions remain limited to the use of stabilized carbon-centered radicals such as benzyl radicals^20,25,26,30,31,36^ and *α*-carbonyl radicals^37,38^ for C–C bond formation. Harnessing highly reactive, unstabilized alkyl radicals for cooperative stereoselective photobiocatalysis, particularly in intermolecular C(sp^3^)–C(sp^3^) bond formation, has remained a formidable challenge.^39,40^ Only a handful of examples have been reported, typically with modest catalytic efficiency and/or stereoselectivity.^23,36,41^ The transient nature of these intermediates, together with their limited ability to engage in stabilizing interactions such as hydrogen bonding and salt-bridge interactions within enzyme active sites, often results in diminished activity and stereoselectivity. The lack of efficient and general biocatalysts for transforming unstabilized alkyl radicals under anaerobic conditions is in stark contrast to recent advances in small-molecule catalysis, where highly efficient and enantioselective asymmetric radical transformations have emerged.^42,43^ These limitations underscore the need for enzyme systems capable of generating and transforming such transient unstabilized alkyl radicals with high fidelity and selectivity.

Through the cooperative use of a photosensitizer and an engineered pyridoxal enzyme, our laboratory recently introduced pyridoxal radical biocatalysis as a new strategy for C(sp^3^)–C(sp^3^) coupling between a catalytically formed free radical and an enzymatic covalent intermediate to produce value-added, stereochemically enriched products.^19^ We envisioned that this dual catalyst system could be leveraged to afford an efficient process to generate and utilize highly reactive primary alkyl radicals, taking advantage of the unique active site environment of PLP-dependent tryptophan synthases. In contrast to the semi-solvent exposed active sites of widely studied flavin-dependent “ene” reductases and nicotinamide-dependent ketoreductases,^44,45^ tryptophan synthases feature a narrow substrate tunnel and undergo pronounced conformational changes between open and closed states during catalysis.^46-48^ We reasoned that engineering this tunnel could enable the encapsulation and stabilization of otherwise short-lived primary alkyl radicals, facilitating their coupling with the catalytically formed enzymatic covalent intermediate. Conformational gating might minimize radical escape, thereby mimicking the spatial and temporal control characteristic of native anaerobic radical enzymes. In contrast to radical SAM/PLP-dependent enzymes, our system employs an external photosensitizer to generate radicals under mild and redox-neutral conditions, offering enhanced tunability and operational simplicity.

Herein, we report the successful implementation of this envisioned photobiocatalytic C–C coupling of unstabilized alkyl radical intermediates. In contrast to reactions with stabilized benzyl radicals, only a very limited subset of PLP enzymes showed detectable activity in this demanding transformation. Through multiple rounds of directed evolution, we engineered a complementary set of radical PLP enzymes that deliver either enantiomer of the C–C coupling products with excellent control. These enantiodivergent biocatalysts enabled the convergent synthesis of aliphatic non-canonical amino acids, which are invaluable building blocks for peptide therapeutics and bioactive natural products,^49^ and could be incorporated into functional proteins via genetic codon expansion methods for therapeutic, biochemical, and biocatalysis applications.^50-52^ Notably, the ability to efficiently engage unstabilized alkyl radical intermediates with extensively engineered enzymes highlights the outstanding synthetic potential of radical PLP biocatalysts and expands the frontiers of enzyme repurposing beyond the scope of pyridoxal radical biocatalysis.

## Results and discussion

At the outset of this study, we employed (2-phenylethyl)boronic acid **1a** and L-serine **2a** as model substrates to evaluate the full collection of PLP-dependent tryptophan synthase β-subunit variants generated in our previous studies. Organoboronic acids were used as the radical precursor to allow for overall redox-neutral coupling. Although most enzyme variants showed very low levels of activity, Y301H was found to be a critical activating mutation.^22^ Incorporation of Y301H to L-*Pf*PLP^β^ afforded the first variant with appreciable activity, affording **3a** in 15% yield albeit with low enantioselectivity (52:48 e.r.). This L-*Pf*PLP^β^ Y301H variant was therefore selected as the starting template for directed evolution via iterative site-saturation mutagenesis (SSM) and screening.

Guided by molecular docking using AutoDock and substrate tunnel analysis using CAVER,^53^ our initial engineering efforts targeted active-site residues within 4.0 Å of the PLP covalent intermediate as well as residues lining the substrate access tunnel for site-saturation mutagenesis (Figure 2c). A key challenge at the outset was the lack of a reliable high-throughput screening protocol, since radical PLP enzymes displayed very low initial activity in cell-free lysates. Unlike biocatalytic reactions using stabilized benzyl radicals, use of the unstabilized alkyl radical precursor **1a** provided <1% product **3a** under lysate-based reactions in a 96-position blue LED (440 nm) photoreactor with rhodamine B (RhB) as the photocatalyst. To address this issue, we systematically optimized reaction parameters and found that switching to green LEDs (527 nm) in combination with eosin B in place of RhB afforded **3a** in appreciable yields, enabling robust high-throughput screening (Figure 2d).

**Figure 2.**
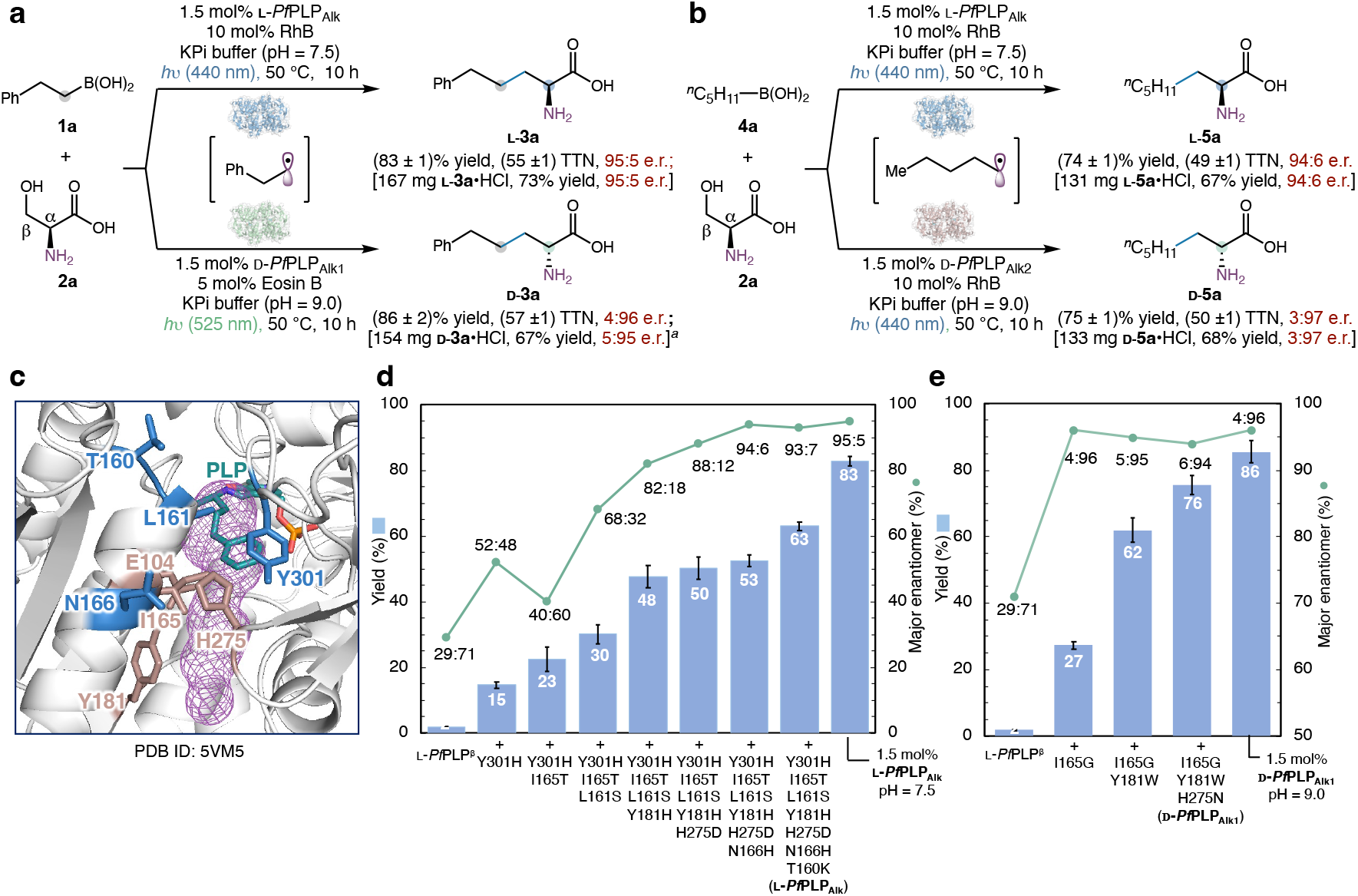
Directed evolution of enantiodivergent pyridoxal radical enzymes to engage completely unstabilized alkyl radicals. **a**, Enantiodivergent photobiocatalytic synthesis of aliphatic non-canonical amino acids via homobenzyl radical: results with final L-*Pf*PLP_Alk_ and D-*Pf*PLP_Alk1_ variants. **b**, Enantiodivergent photobiocatalytic synthesis of aliphatic non-canonical amino acids via 1-pentyl radical: results with final L-*Pf*PLP_Alk_ and D-*Pf*PLP_Alk2_ variants. **c**, Docking results and the substrate tunnel generated by CAVER based on the crystal structure created from PDB ID 5VM5. **d**, Directed evolution of L-*Pf*PLP_Alk_ using **1a** and **2a** as the substrates. e, Directed evolution of D-*Pf*PLP_Alk1_ using **1a** and **2a** as the substrates. For UV-active amino acids (L-**3** or D-**3**), the yields were determined by LC-MS analysis; For UV-inactive amino acids (L-**5** or D-**5**), the products were derivatized with 2,4-dinitrofluorobenzene (DNFB) to generate UV-active derivatives for yield determination; All the reactions were performed in technical triplicates and averaged yields were reported; Error bars indicate standard deviation; Batch-to-batch variation across biological replicates of independently purified enzymes gave the following results: L-**3a**, 82–83%; D-**3a**, 85–87%; L-**5a**, 73–75%; D-**5a**, 75–78%. ^*a*^1.0 mol% D-*Pf*PLP_Alk1_ was used.

Using L-*Pf*PLP^β^ Y301H as the starting template, two rounds of directed evolution led to the identification of beneficial mutations I165T and L161S, increasing the yield to 30% with an enantiomeric ratio (e.r.) of 68:32. Both residues 165 and 161 are located in an α-helix defining the narrow substrate tunnel. In the fourth round of engineering, incorporation of Y181H further enhanced enzyme performance, affording L-**3a** in 48% yield and 82:18 e.r.. The fifth and sixth rounds led to beneficial mutations H275D and N166H, increasing enantioselectivity to 94:6 e.r. with a modest improvement in activity. A final round of engineering revealed T160K as a key mutation, culminating in the variant L-*Pf*PLP^β^ Y301H I165T L161S Y181H H275D N166H T160K, hereafter designated L-*Pf*PLP_Alk_ (radical alkylating PLP enzyme from Pyrococcus furiosus). Increasing the enzyme loading to 1.5 mol% and adjusting the reaction pH from 8.0 to 7.5 further enhanced activity and selectivity, affording L-**3a** in 83% yield and 95:5 e.r.. Notably, L-*Pf*PLP_Alk_ was not limited to the transformation of substrate **1a**: without additional protein engineering, L-*Pf*PLP_Alk_ catalyzed the coupling of aliphatic 1-pentylboronic **4a** to provide L-**5a** in 74% yield and 94:6 e.r., underscoring its ability to transform highly reactive, completely unstabilized primary alkyl radicals (Figure 2b; see supplementary Tables S7–S8 for further details).

A central feature of our tryptophan synthase-catalyzed C(sp^3^)–C(sp^3^) radical coupling is the ability to achieve enantiodivergent synthesis of non-canonical amino acids. To engineer efficient D-amino acid producing enzymes for unstabilized alkyl radicals, we first evaluated *Pf*PLP^β^ variants known to invert α-stereochemistry. While our previously reported 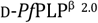 (L-*Pf*PLP^β^ E104P) only provided D-**3a** in 5% yield and 9:91 e.r., the L-*Pf*PLP^β^ I165G variant afforded D-**3a** in 27% yield with a promising enantiomeric ratio of 4:96. Targeted site-saturation mutagenesis of residues surrounding G165 identified Y181W as highly beneficial, increasing the yield of D-**3a** to 62% while maintaining the e.r.. Addition of H275N afforded the final variant L-*Pf*PLP^β^ I165G Y181W H275N (D-*Pf*PLP_Alk1_), which delivered D-**3a** in 76% yield and 6:94 e.r.. Under optimized conditions (1.5 mol% D-*Pf*PLP_Alk1_ pH 9.0), D-**3a** was obtained in 86% yield and 4:96 e.r. (Figure 2e).

Encouraged by this result, we examined aliphatic substrate **4a** with D-*Pf*PLP_Alk1_. Under standard conditions, the desired product D-**5a** was obtained in 55% yield and 34:66 e.r., indicating reduced enantiocontrol over aliphatic radicals. This prompted a parallel engineering campaign starting from 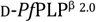. Introduction of Y181K provided a new variant L-*Pf*PLP^β^ E104P Y181K (D-*Pf*PLP_Alk2_), which improved the yield and enantioselectivity for D-**5a** to 47% and 4:96 e.r.. Switching back to RhB (440 nm) further improved the yield of D-**5a** to 64%. By increasing the enzyme loading to 1.5 mol% and adjusting the reaction pH from 8.5 to 9.0, D-**5a** was obtained in 75% yield and 3:97 e.r. (Figure 2b; see supplementary Table S17 for details). Finally, this photobiocatalytic radical C(sp^3^)–C(sp^3^) coupling could be scaled up conveniently. On a 1.0 mmol scale, with 1–1.5 mol% enzyme loading and Kessil LED irradiation, all four standard reactions shown in Figures 2a and 2b afforded over 100 mg amino acid products while maintaining excellent enantioselectivity, demonstrating the potential utility of these PLP biocatalysts for laboratory-scale synthesis (See supplementary Figure S5-S6 for details).

With a set of enantiodivergent PLP enzymes in hand, we next explored the substrate scope of the photobiocatalytic asymmetric dehydroxylative C(sp^3^)–C(sp^3^) coupling. Under the optimized reaction conditions, diverse organoboron reagents, including organoboronic acids, pinacol boronate esters, and trifluoroborates, displayed comparable activity (supplementary Table S6). Accordingly, substrates for this study were selected primarily on the basis of synthetic accessibility. To systematically evaluate the synthetic utility of evolved PLP enzymes, we organized the substrate scope into two categories, including (i) alkyl radical precursors bearing a pendant aromatic substituent (Figure 3), and (ii) radical precursors featuring aliphatic chains (Figure 4).

**Figure 3.**
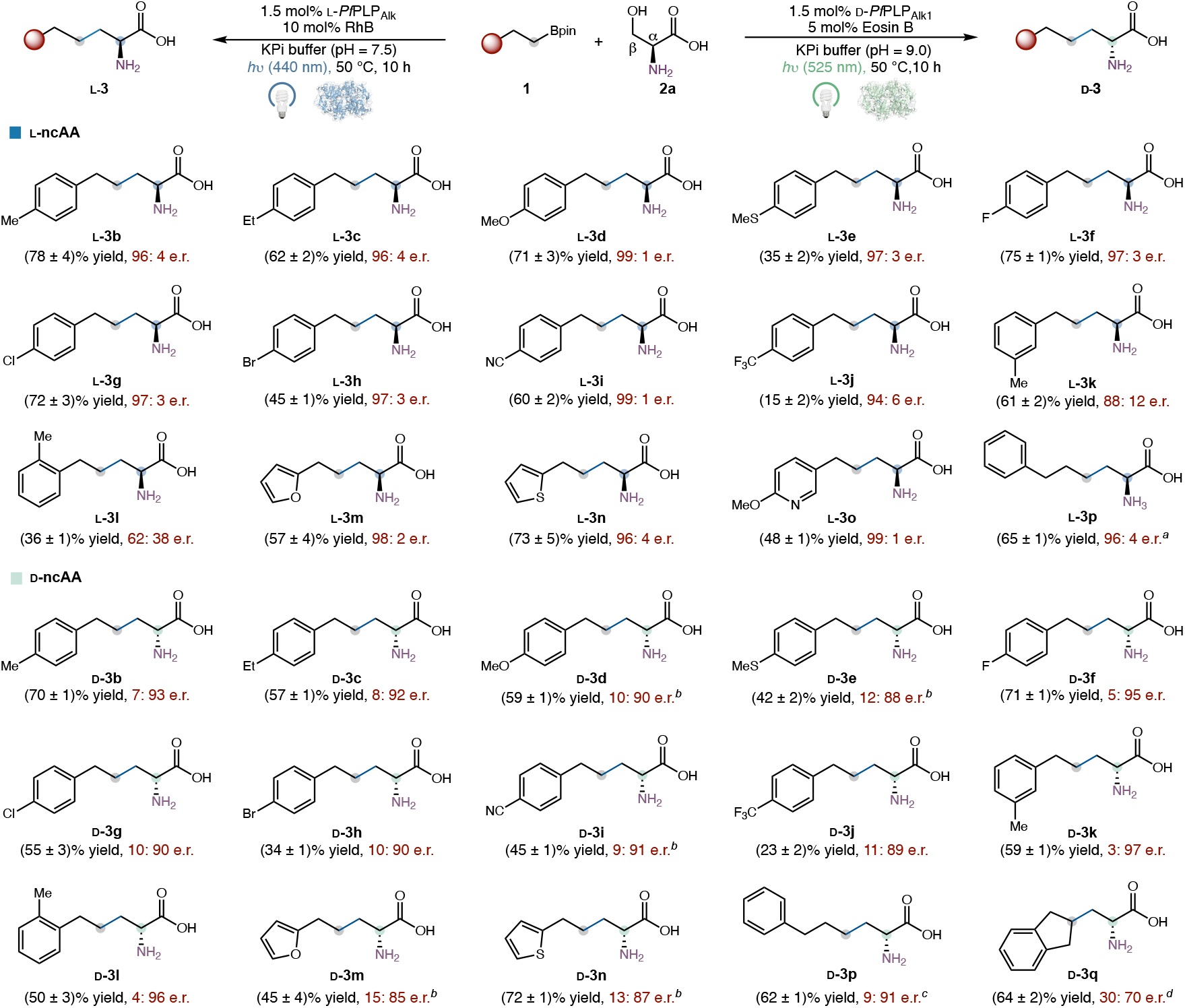
Substrate scope of enantiodivergent photobiocatalytic C(sp^3^)–C(sp^3^) coupling. **a**, L-amino acid synthesis using L-*Pf*PLP_Alk_. Reaction conditions: **1** (4.0 mM), **2a** (20.0 mM), 1.5 mol% L-*Pf*PLP_Alk_, 10 mol% RhB, *h*ν (440 nm, 5.0 W), 200 mM KPi buffer (pH = 7.5), DMSO (8% v/v), 50 °C, 10 h, under N_2_. **b**, D-amino acid synthesis using D-*Pf*PLP_Alk1_. Reaction conditions: **1** (4.0 mM), **2a** (20.0 mM), 1.5 mol% D-*Pf*PLP_Alk1_, 5 mol% Eosin B, *h*ν (525 nm, 3.0 W), 200 mM KPi buffer (pH = 9.0), DMSO (8% v/v), 50 °C, 10 h, under N_2_. All the reactions were performed as technical triplicates and averaged yields were reported. ^*a*^1.5 mol% L-*Pf*PLP^β^ Y301H I165T L161S Y181H H275D N166H was used in lieu of L-*Pf*PLP_Alk_. ^*b*^10 mol% Eosin B, *h*ν (525 nm, 2.0 W), 200 mM KPi buffer (pH = 8.5). ^*c*^1.5 mol% L-*Pf*PLP^β^ I165G Y181W was used in lieu of D-*Pf*PLP_Alk1_. ^*d*^1.5 mol% D-*Pf*PLP_Alk2_ was used in lieu of D-*Pf*PLP_Alk1._

**Figure 4.**
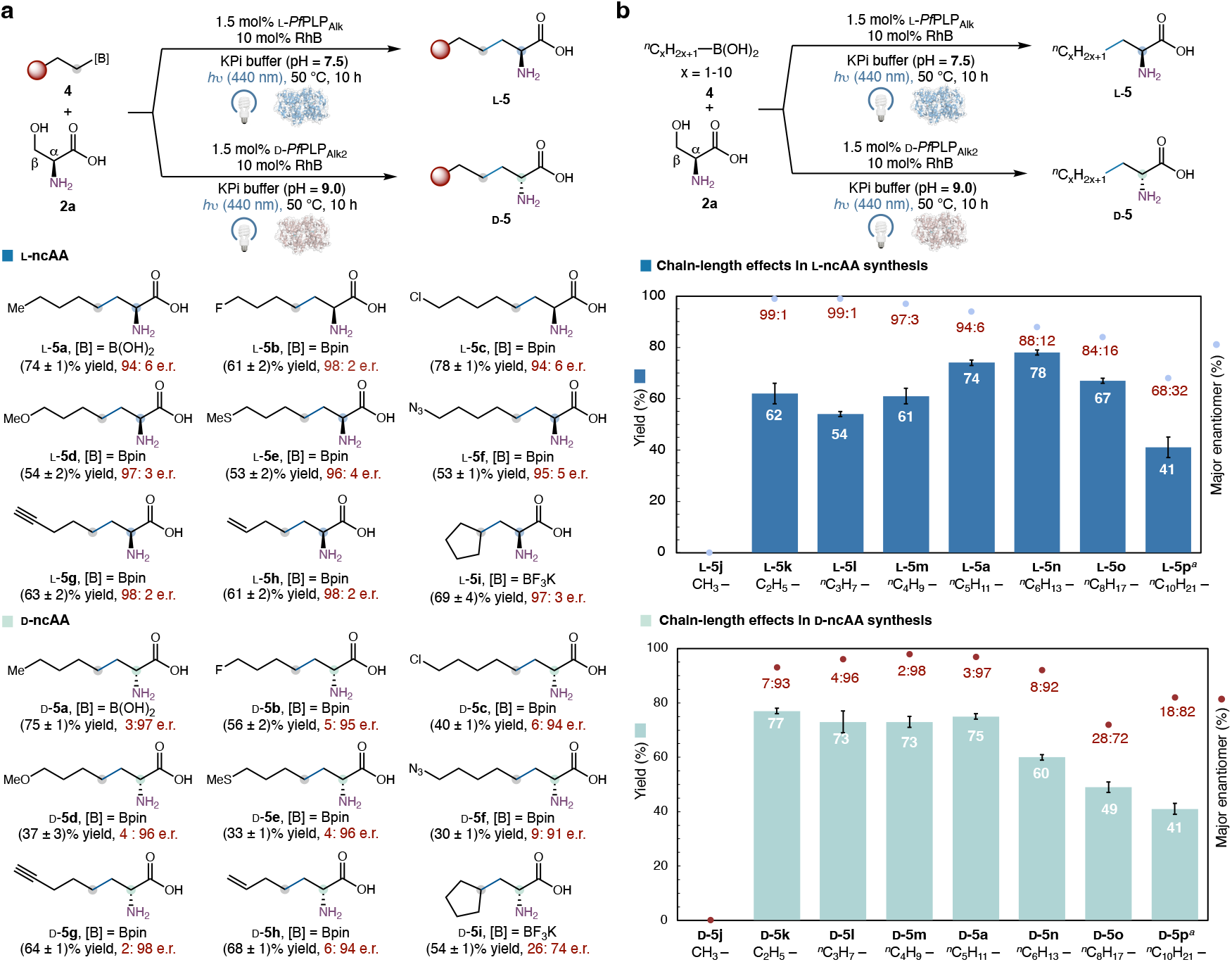
Enantiodivergent photobiocatalytic amino acid synthesis harnessing unstabilized, aliphatic radical intermediates. **a**, Substrate scope of functionalized alkylboron coupling partners. **b**, Chain-length effects in enantiodivergent photobiocatalytic coupling of alkyl radicals. Reaction conditions for L-amino acid synthesis: **4** (4.0 mM), **2a** (20.0 mM), 1.5 mol% L-*Pf*PLP_Alk_, 10 mol% RhB, *h*ν (440 nm, 4.0 W), 200 mM KPi buffer (pH = 7.5), DMSO (8% v/v), 50 °C, 10 h, under N_2_. Reaction conditions for D-amino acid synthesis: **4** (4.0 mM), **2a** (20.0 mM), 1.5 mol% D-*Pf*PLP_Alk2_ 10 mol% RhB, *h*ν (440 nm, 4.0 W), 200 mM KPi buffer (pH = 9.0), DMSO (8% v/v), 50 °C, 10 h, under N_2_. ^*a*^DMSO (15% v/v). All the reactions were performed as technical triplicates and averaged yields were reported; Error bars indicated standard deviation.

As summarized in Figure 3, a wide range of substituents on the pendant aromatic ring (L-**3b**–L-**3l**) were tolerated by the L-amino acid producing enzyme, affording the corresponding non-canonical amino acids with excellent enantiocontrol. Organoboron substrates bearing an electron-donating substituent at the para-position of the arene such as a methyl (L-**3b**), an ethyl (L-**3c**), a methoxy (L-**3d**), or a methylthio (L-**3e**) provided excellent yields and enantioselectivity. Halogenated substrates possessing a fluoro (L-**3f**), a chloro (L-**3g**), or a bromo (L-**3h**) substituent could also be transformed efficiently. Electron-withdrawing groups at the para-position of the aromatic ring including a cyano (L-**3i**) or a trifluoromethyl (L-**3j**) were also compatible, although the latter showed reduced yield. Additionally, meta-(L-**3k**) and ortho- (L-**3l**) substituted organoboron substrates were also accepted by the engineered biocatalyst, albeit with a slight decrease in enantioselectivity. Furthermore, heterocyclic substrates, including a furyl (L-**3m**), a thienyl (L-**3n**), or a pyridyl (L-**3o**) group were readily accommodated, highlighting the utility of the present photobiocatalytic method to access medicinally relevant amino acid products. Finally, a γ-aryl substrate (L-**3p**) was also converted with excellent yield and enantioselectivity. Notably, the substrate scope of the D-amino acid producing enzyme D-*Pf*PLP_Alk1_ closely mirrored that of the L-amino acid producing enzyme L-*Pf*PLP_Alk_ (D-**3b**–D-**3q**), allowing enantiodivergent access to a broad range of non-canonical amino acids via a convergent C–C coupling approach.

We next examined the substrate scope of aliphatic radical precursors lacking pendant aromatic groups (Figure 4). A broad spectrum of alkylboron substrates was efficiently transformed with excellent yields and enantioselectivity by either L-*Pf*PLP_Alk_ or D-*Pf*PLP_Alk2_. Functional groups including a fluoro- (L-**5b** and D-**5b**), a chloro (L-**5c** and D-**5c**), a methoxy (L-**5d** and D-**5d**), and a methylthio (L-**5e** and D-**5e**) substituent were well tolerated, providing the corresponding aliphatic non-canonical amino acid products with excellent enantioselectivity. Importantly, this system accommodated bioorthogonal functional group handles, including an azide (L-**5f** and D-**5f**), an alkyne (L-**5g** and D-**5g**), and an olefin (L-**5h** and D-**5h**), providing synthetically accessible ncAAs that are potentially useful for genetic code expansion^50-52^ and subsequent modification by click chemistry or thiol–ene coupling.^54^ Beyond primary alkyl radical precursors, secondary alkylboron reagents such as a cyclopentyl (L-**5i** and D-**5i**) an isopropyl and tertiary alkyl substrates such as a tert-butyl (Supplementary Figure S20) were also transformed. These results underscore the unusual capacity of this dual photobiocatalytic system to harness highly reactive, unstabilized radical intermediates across a broad range of substrates.

To date, there has been limited understanding of the key factors governing the efficiency of transformations involving unstabilized alkyl radicals in photobiocatalysis, hampering further optimization of biocatalytic reactions that harness these reactive intermediates. We reasoned that our newly engineered PLP enzymes, specifically developed to engage unstabilized alkyl radicals, could provide valuable mechanistic insights. To this end, we examined a series of linear alkylboronic acids (C1–C10, **5j**–**5p**) with the enantiocomplementary PLP enzymes L-*Pf*PLP_Alk_ and D-*Pf*PLP_Alk2_ (Figure 4b). Starting from the ethylboronic acid (C2, **4k**), all higher homologs were transformed into the corresponding aliphatic amino acid products with good to excellent enantioselectivity with both L-*Pf*PLP_Alk_ and D-*Pf*PLP_Alk2_.

With the L-amino acid producing enzyme L-*Pf*PLP_Alk_ (Figure 4b), the highest enantioselectivities were observed with short-chain alkyl substrates, with ethyl (L-**5k**) and n-propyl (L-**5l**) affording products in 99:1 e.r.. As the chain length increased from C2 to C10, enantioselectivity gradually decreased from 99:1 e.r. to 68:32 e.r.. In contrast to this monotonic decline, product yields followed a non-monotonic trend: yields improved from C3 to C6, peaking at n-pentyl (C5, L-**5a**) and n-hexyl (C6, L-**5n**), before declining with longer substrates.

Interestingly, the D-amino acid producing enzyme D-*Pf*PLP_Alk2_ exhibited distinct trends (Figure 4b). For C2– C5 substrates, enantioselectivity modestly increased (7:93 e.r. to 3:97 e.r.), but decreased slightly from C5 (D-**5a**) to C8 (D-**5o**). Yields were highest (73%–77%) for C2–C5 substrates (D-**5k**–D-**5m** and D-**5a**), while longer chains C6–C10 substrates (D-**5n**–D-**5p**) showed diminished activity. Increasing the amount of DMSO co-solvent (15%) improved yields for the poorly soluble *n*-decyl substrate (C10, **5p**) with both L-*Pf*PLP_Alk_ and D-*Pf*PLP_Alk2_. By contrast, methylboronic acid (**5j**) did not provide the desired coupling product, likely due to the highly reactive nature of the methyl radical generated. Collectively, this systematic investigation of chain-length effects revealed both general reactivity principles of unstabilized alkyl radicals and subtle enzyme-dependent influences on enantioselectivity and activity, underscoring fundamental distinctions between biocatalysis and small-molecule-catalysis. These insights may facilitate the development of cooperative photobiocatalytic reactions involving highly reactive alkyl radicals.

### Mechanistic studies

To elucidate the mechanism of this biocatalytic C(sp^3^)– C(sp^3^) coupling involving unstabilized alkyl radicals, a series of mechanistic studies was carried out. With either L-*Pf*PLP_Alk_ or D-*Pf*PLP_Alk2_, the reaction of cyclopropyl-containing radical clock substrate **4q** provided the ring-opened product **5h** exclusively in high yield and enantioselectivity (Figure 5a). Given the ring-opening rate constant of **4q** was previously determined to be 1.0 × 10^7^ s^−1^,^55^ these results establish a lower-bound radical lifetime of 10^−7^ s in this dual catalyst system with both L-*Pf*PLP_Alk_ and D-*Pf*PLP_Alk2_. When the 5-hexenyl-1-boronate substrate **4r** was applied, both the cyclized (**5r**) and the non-cyclized coupling product (**5s**) formed in a 95:5 ratio, suggesting that 5-exo-trig radical cyclization precedes the C–C bond formation with the enzymatic PLP covalent intermediate (Figure 5b).

**Figure 5.**
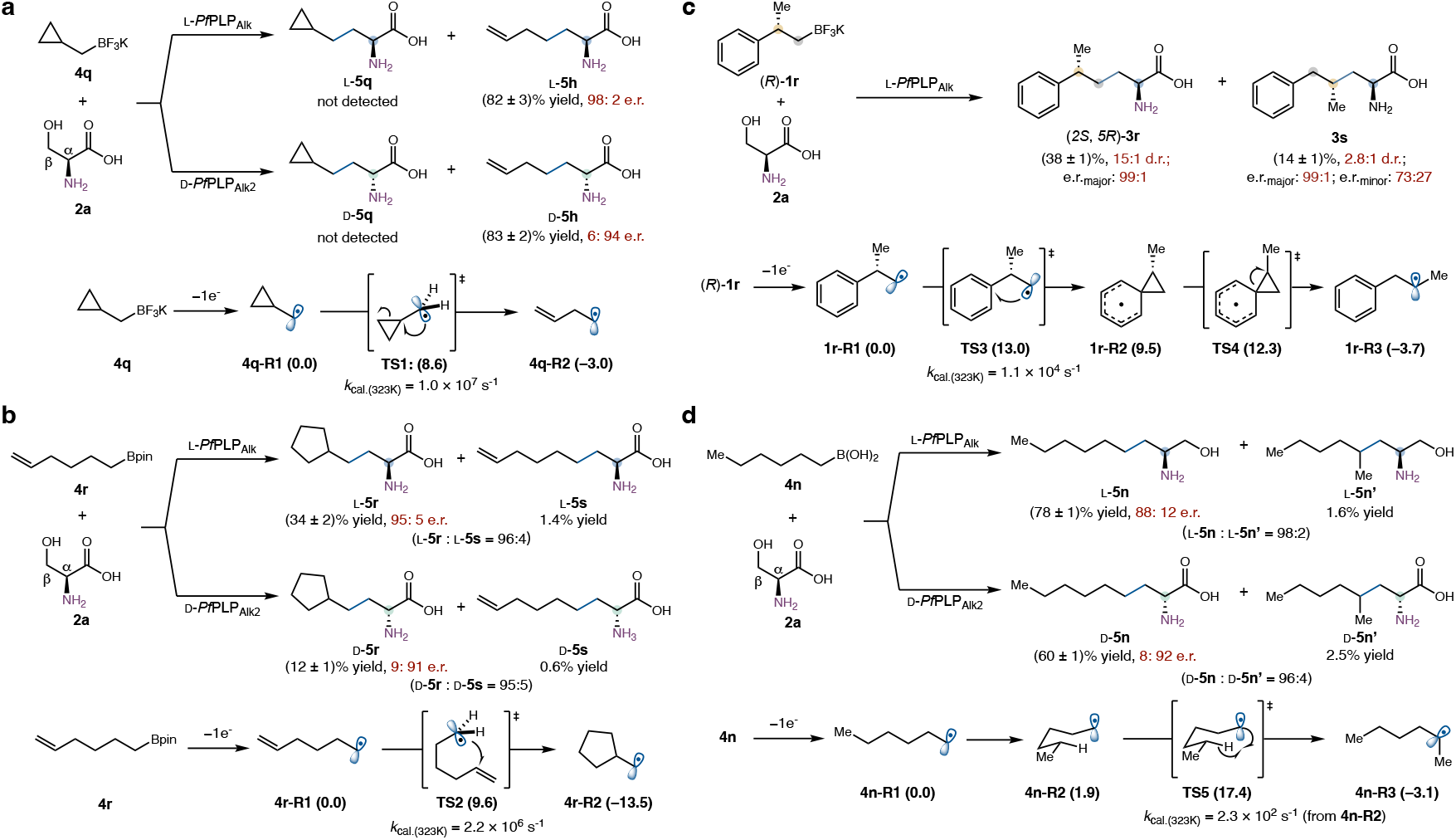
Probing the lifetime of unstabilized radical intermediates in photobiocatalytic C–C coupling. **a**, Radical-mediated ring opening of **4q. b**, Radical cyclization of **4r. c**, Radical-mediated 1,2-aryl migration of (*R*)-**1r. d**, Radical-mediated isomerization of **4n**. Density functional theory (DFT)-computed Gibbs free energies (in parentheses) are in kcal/mol with respect to corresponding primary radical intermediates.

Furthermore, photobiocatalytic C–C coupling of aryl substrate (*R*)-**1r** with L-*Pf*PLP_Alk_ provided amino acid product **3r** in 38% yield, 15:1 d.r. and >99:1 e.r. featuring a well-defined skipped α,δ-stereochemical dyad (Figure 5c). Interestingly, an isomerized product **3s** was also observed in 14% yield, 2.8:1 d.r. and 99:1 e.r. (major diastereomer) or 73:27 e.r. (minor diastereomer). This product likely arose from a radical-mediated 1,2-aryl migration, an uncommon rearrangement in biocatalytic radical processes. Density functional theory (DFT) calculations revealed a stepwise migration process with a rate constant of 1.1 × 10^4^ s^−1^, suggesting that the radical intermediate generated from **1r** has a lifetime exceeding or comparable to 10^−4^ s.

When *n*-hexylboronic acid (**4n**) was employed, in addition to the expected coupling product **5n**, a highly unusual isomerized product **5n’** was observed in 2–3% yield with either L-*Pf*PLP_Alk_ or D-*Pf*PLP_Alk2_ (Figure 5d). The identity of **5n’** was confirmed by comparison with an independently synthesized amino acid sample. We attribute its formation to a rarely reported 1,5-hydrogen atom transfer (1,5-HAT), converting a primary 1-hexyl radical into a slightly more stable secondary 2-hexyl radical. No analogous rearranged products were observed with shorter-chain alkylboronic acids such as ethyl (**4k**) and n-propylboronic acid (**4l**), consistent with our proposed 1,5-HAT mechanism to account for this isomerization. DFT computations further indicated a rate constant of 2.3 × 10^2^ s^−1^ for this 1,5-HAT process, proceeding from a bent (*g*^+^*g*^−^) conformer of the 1-hexyl radical (**4n**-**R2**), which likely represents the preferred conformation within the enzyme active site. For this unusual radical isomerization to occur, the lifetime of 1-hexyl radical is estimated to be on the order of 10^−2^ s^−1^, which is much longer than typical free radical lifetimes in solution-phase chemistry. Taken together, these radical rearrangement processes demonstrated that radical intermediates generated in this dual enzyme-photocatalyst system persist far longer than those in conventional homogenous systems. We hypothesized that active-site confinement played a critical role in extending the lifetimes of otherwise short-lived radical intermediates. Although a mechanism involving further single-electron oxidation of the radical intermediate to a carbocation, followed by carbocation-based rearrangement, cannot be ruled out, this pathway is less likely due to the limited stability of carbocationic intermediates under aqueous reaction conditions. Together, these findings provide insights into the conversion of unstabilized alkyl radicals in new biocatalytic processes.

Based on experimental results, a proposed mechanism for this PLP-enzyme catalyzed radical C–C coupling is described in Figure 6. The catalytic cycle begins with the internal aldimine **I** of the tryptophan synthase *β*-subunit,^56^ which undergoes transimination with the serine substrate to generate the corresponding external aldimine **II**. A lysine-mediated *α*-deprotonation of **II** affords the quinonoid intermediate **III**, which then undergoes *β*-elimination to afford the electrophilic aminoacrylate **IV**. An unstabilized alkyl radical **V**, generated from organoboron substrate **1** via photoinduced electron transfer, enters the enzyme active site, where it may be stabilized by active-site confinement.^48,57^ Stereoselective addition of **V** to the *β*-position of aminoacrylate **IV** affords the azaallyl radical intermediate **VI**, which undergoes single-electron reduction followed by enantioselective protonation to furnish the external aldimine **VII**. Protonation from the (*Re*)-face by the ε-ammonium group of the conserved lysine residue produces the (*S*)-configured aldimine, whereas protonation from the (*Si*)-face, likely mediated by water molecules, leads to inverted *α*-stereochemistry.^22^ Final transimination with the lysine residue regenerates internal aldimine **I** and releases the non-canonical amino acid product **3** or **5**, thus completing the catalytic cycle.

**Figure 6.**
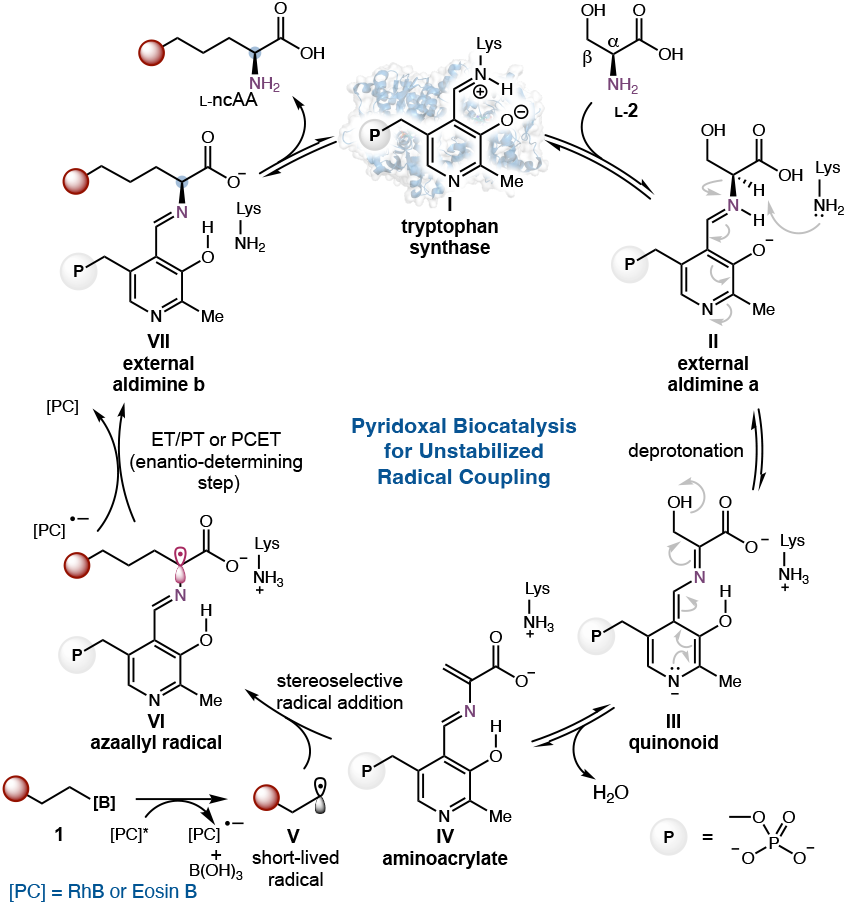
Proposed catalytic cycle.

## Conclusions

In summary, by exploiting enzyme-photocatalyst synergy, we reprogrammed and evolved PLP-dependent tryptophan synthases to harness unstabilized alkyl radical intermediates for the asymmetric synthesis of aliphatic non-canonical amino acids. For a long time, photobiocatalysis has been largely limited to the exploitation of stabilized radical intermediates such as benzyl and *α*-carbonyl radicals. Through careful engineering of PLP-dependent radical C–C coupling enzymes, we demonstrated that these traditionally challenging unstabilized alkyl radicals could be effectively transformed into valuable products. In particular, the development of enantiocomplementary PLP enzymes converting the same L-serine into either L- or D-non-canonical amino acid furnished a powerful biocatalytic tool for the convergent and stereocontrolled assembly of aliphatic amino acids that play an essential role in numerous biological and biochemical studies. Mechanistic studies further revealed the unexpectedly long-lived nature of unstabilized alkyl radicals within dual enzyme-photocatalyst systems, offering a blueprint for expanding photobiocatalysis to a much broader range of synthetically valuable radical species.

## ASSOCIATED CONTENT

### Supporting Information

The Supporting Information is available free of charge on the ACS Publications website.

Experimental procedures, DNA and protein sequences, characterization data, HPLC traces and NMR spectra (PDF).

## Notes

Y.Y., L.C. and Z.B. has filed a provisional patent on the compositions, systems and methods for biocatalytic amino acid synthesis based on the results described in this manuscript. Other authors have no competing interest to declare.

## ACKNOWLEDGMENT

This research is supported by the National Institutes of Health (R01EB036084 to Y.Y.), Army Research Office (W911NF-24-2-0246) and Howard Hughes Medical Institute (Y.Y.). Y.Y. is an Alfred P. Sloan Research Fellow (FG-2024-22244), a Camille Dreyfus Teacher-Scholar Awardee (TC-25-084), a David & Lucile Packard Fellow (2023-76169) and an HHMI Freeman Hrabowski Scholar. Computational study is supported by the National Science Foundation grant CHE-2400087 (P.L.). Computations were carried out at the University of Pittsburgh Center for Research Computing and Data and the Advanced Cyberinfrastructure Coordination Ecosystem: Services & Support (ACCESS) Program, supported by NSF award numbers OAC-2117681, OAC-1928147, and OAC-1928224.

---

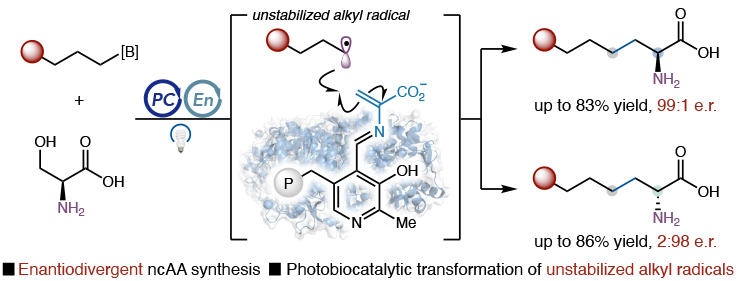

