## Supporting Information for "Engaging Unstabilized Alkyl Radicals with Pyridoxal Radical Biocatalysis: Enantiodivergent Synthesis of Aliphatic Non-Canonical Amino Acids"

#### I. General methods

**General.** Biocatalytic reactions were reperformed in a 1-dram vial (15 × 45 mm, 4 mL, VWR) with magnetic stirring (300 rpm) using Shanghai 3S Technology Co., Ltd parallel photoreactor (AF2 model) as the visible light source or in shell vials (8 × 30 mm, 0.8 mL, Thomas Scientific) using 96-well array (Lumidox® II 96-Position Discovery LED Array 527nm-Green). Unless otherwise noted, all chemicals and reagents were obtained from commercial suppliers (Sigma-Aldrich, VWR, Alfa Aesar, Combi-Blocks, AmBeed, Oakwood, and Enamine) and used without further purification. Silica gel chromatography was carried out using AMD Silica Gel 60, 230-400 mesh. $^{1}H$ and $^{13}C$ NMR spectra were recorded on a Bruker 400 or 500 MHz instrument in $CDCl_3$, $D_2O$ or methanol-$d_4$ ($CD_3OD$). $^{19}F$ NMR and $^{11}B$ NMR spectra (where applicable) were recorded on a Bruker 400 or 500 MHz ($^{1}H$ decoupled). Data for $^{1}H$ NMR are reported as follows: chemical shift (δ ppm), multiplicity (s = singlet, d = doublet, t = triplet, q = quartet, p = pentet, sext = sextet, m = multiplet, dd = doublet of doublets, dt = doublet of triplets, ddd = doublet of doublet of doublets, brs = broad singlet), coupling constant (Hz), integration. Sonication on a small scale was performed using a BioLogics ultrasonic homogenizer (model 150VT) equipped with a stepped microtip. Sonication on a large scale was performed using a Branson Digital Sonifier. 450 Ultrasonic Processor. All IR spectra were recorded on a Thermo Scientific Nicolet iS5 spectrometer (iD5 ATR, diamond). High-resolution mass spectrometry data was obtained at the Mass Spectrometry Facilities at the University of California Santa Barbara. High-resolution accurate mass (HRAM) ESI data was analyzed on a Waters LCT Premier mass spectrometer with a LEAP PAL autosampler with isocratic MeOH flow (no column; direct injection). Molecular formulas (MF) were validated by lock mass calibration to sodiated polyethylene glycol polymer or sodiated monoethyl ether polyethylene glycol polymer standards. High-resolution accurate mass (HRAM) EI data was acquired using a Waters LCT Premier time-of-flight (TOF) mass spectrometer. Masses of positively charged ions were calibrated using a methanol solution of polyethylene glycol or polyethylene glycol monomethyl ether as an internal standard. Synthetic reactions were monitored by thin layer chromatography (TLC, Silicycle TLG-R10014BK-323 gel plates) using a UV-lamp or an appropriate TLC stain for visualization. UV-vis spectra were collected on a UV1800 Shimadzu spectrophotometer.

*E. coli* cells were grown using Luria-Bertani medium (LB) with 0.1 mg/L ampicillin ($LB_{amp}$). Primer sequences for site-directed mutagenesis and site-saturation mutagenesis are provided below. T5 exonuclease, Phusion DNA polymerase, and *Taq* DNA ligase were purchased from New England Biolabs (NEB, Ipswich, MA). Potassium Phosphate Buffer (abbreviated as KPi buffer) was used as the buffering system for protein purification and storage unless otherwise specified.

**Chromatography.** Analytical reverse-phase high-performance liquid chromatography mass spectrometry (HPLC-MS) was performed using a Shimadzu 2040C 3D Plus system with a 2020 MS detector and a Poroshell 120 EC-C18 column (4.6 × 50 mm, 4 μm). Water (0.1% formic acid) and acetonitrile (0.1% formic acid) were used as the mobile phases. The yields of compounds **3a**–**3q** were determined by calibration curves using homophenylalanine as the internal standard. Compounds **5a**–**5p** were first derivatized using 2,4-dinitrofluorobenzene, and their yields were determined by calibration curves. The enantiomeric ratios (e.r.'s) of the non-canonical amino acid products were determined by Marfey's analysis as described below.

**Cloning, site-saturation mutagenesis.** pET-22b (+) was used as the cloning and expression vector for *Pyrococcus furiosus* tryptophan synthase β-subunit variants, including L-*Pf*$PLP_{Alk}$, D-*Pf*$PLP_{Alk}$. Genes of PLP-dependent enzymes were codon optimized and purchased from General Biol as plasmids using pET-22b(+) as the cloning vector. The gene of interest (GOI) was cloned into pET-22b(+) between restriction sites *Nde*I and *Xho*I and this construct has a C-terminal 6X His tag. Site-saturation mutagenesis was performed using the "22c-trick" method as described by Kille et al[1]. The PCR products were digested with DpnI, gel purified, and ligated using a Gibson mix prepared from 5X isothermal (ISO) reaction buffer (25% PEG-8000, 500 mM Tris-HCl pH 7.5, 50 mM $MgCl_2$, 50 mM DTT, 1 mM each of the dNTPs, and 5 mM NAD), T5 exonuclease, Phusion DNA polymerase, and *Taq* DNA ligase[2]. The ligation mixture was used directly to transform electrocompetent *E. coli* strain *E. cloni* BL21(DE3) cells (Lucigen).

**Expression of *Pf*$PLP_{Alk}$ variants in 24-well plates.** Single colonies from $LB_{amp}$ agar plates were picked using sterile toothpicks and cultured in deep-well 96-well plates containing $LB_{amp}$ (400 μL/well) at 37 °C with 250 rpm shaking overnight. An aliquot of these overnight cultures (200 μL per well) was then used to inoculate $TB_{amp}$ medium (3800 µL per well) in 24-well plates, followed by shaking at 37 °C and 230 rpm for 2.5 h. The plates were cooled on ice for 20 min before induction with isopropyl *β*-D-1-thiogalactopyranoside (IPTG) to a final concentration of 1.0 mM. Protein expression was carried out at 20 °C with 200 rpm shaking for 20 h.

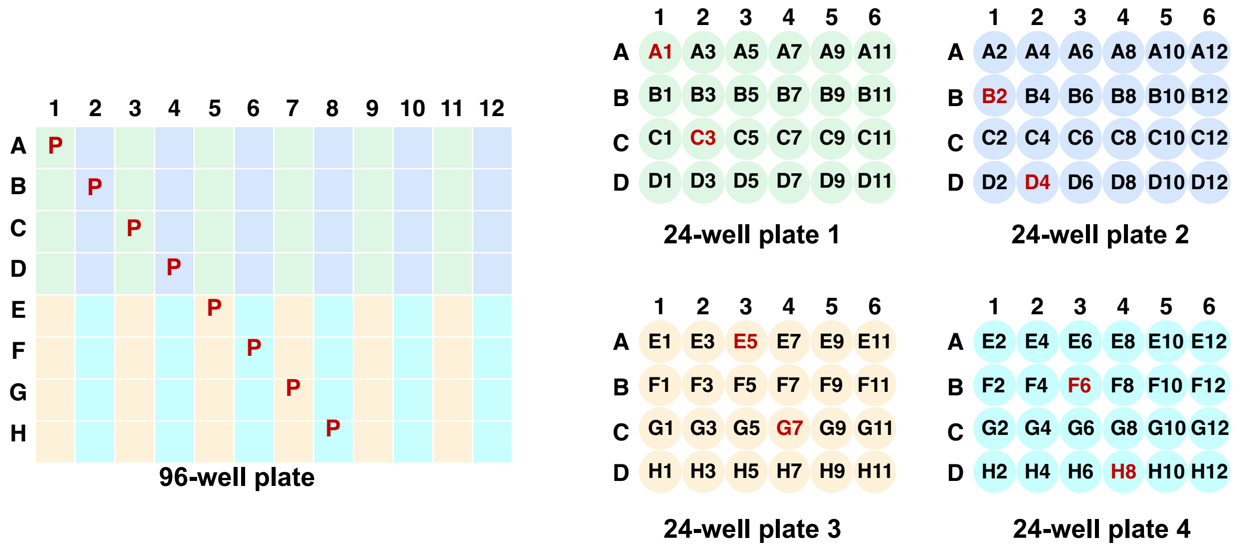

**Figure S1.** Inoculating four 24-well plates using overnight cultures from one 96-well plate. Parents (P, shown in red) were grown in the wells of A1, B2, C3, D4, E5, F6, G7, and H8.

**Reaction screening in 96-well plate format.** The cells grown in 24-well plates were then pelleted by centrifugation (3,434 *g*, 4 min, 4 °C) using an Eppendorf 5910R tabletop centrifuge. The cell pellets were resuspended in KPi buffer (200 mM, pH 8.0, 1200 µL/well) containing 0.2 mM PLP by gentle shaking (800 rpm, 3–5 min) on a Fisher Scientific microplate shaker. The cell suspension was lysed by sonication with a Qsonica Q500 sonicator equipped with a 24-tip horn. 24 wells per plate were lysed in each sonication round (e.g., wells A1, A3, A5, …, A11, C1, C3, C5, …, C11, E1, E3, E5, …, E11, G1, G3, G5, …, and G11). Complete lysis of all wells required four rounds of sonication under these conditions: 45% amplitude, 2 seconds on, 4 seconds off, totaling 9 min (3-minute effective sonication time). Following sonication, the lysates were heat-treated at 75 °C for 10 min and centrifuged (4,500 rpm, 25 min, 4 °C) to pellet cell debris by an Eppendorf tabletop centrifuge 5910R. An aliquot of supernatant (450 µL per well) was transferred to a reaction shell vial using an Eppendorf Xplorer plus, 12-channel, 1000 µL electronic pipette, after which the 96-

well plate was transferred into a Coy anaerobic chamber. Inside the chamber, 20 μL (2-phenylethyl)boronic acid or *n*-pentylboronic acid stock solution (100 mM in DMSO), 20 μL Eosin B stock solution (10.0 mM in DMSO) and 20 μL L-serine stock solution (500 mM in KPi buffer) were added into each well using an Eppendorf Xplorer 12-channel pipette (15–300 μL). The shell vials were sealed with aluminum foil and shaken at 400 rpm on a Corning microplate shaker in the Coy anaerobic chamber and illuminated by a 527 nm Lumidox 96-position LED array (80 mW) purchased from Analytical Sales and Services, Inc.. After 4 h, the vials were removed from the anaerobic chamber, the seals were removed, and the analytical-scale reactions were worked up as described below.

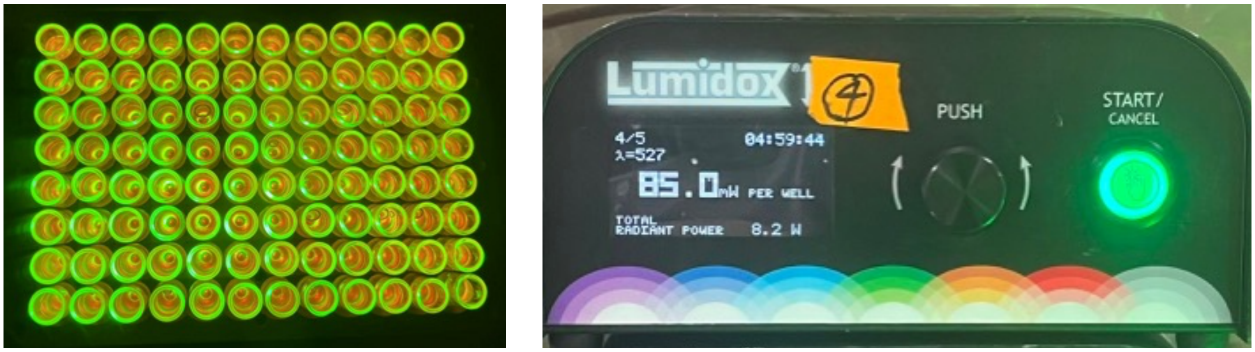

**Figure S2**. Lumidox® II 96-position LED array for directed evolution (wavelength: 527 nm).

**Product formation screening using reverse phase HPLC-MS.** The reaction mixture was transferred from the shell vial into a deep-well 96-well plate. To each well, 50 μL homophenylalanine stock solution (40.0 mM in $CH_3CN$/1 M aq. HCl) was added as the internal standard, followed by the addition of 1.5 mL of $CH_3CN$/1 M aq. HCl (v:v = 1:1). After vigorous mixing, the plate was centrifuged at 4,500 rpm for 25 min to remove the precipitate in the aqueous solution. An aliquot of the clear supernatant (200 μL per vial) was transferred to 500 μL vial inserts using an Eppendorf Xplorer 12-channel pipette (15–300 μL). The inserts were then placed in 2 mL vials and analyzed by reverse-phase LCMS. The yield increasing for **3a** was determined by LC-MS analysis using homophenylalanine as the internal standard with freshly developed calibration curves; the yield increasing for **5a** was determined by MS analysis using homophenylalanine as the internal standard. (Details of the LC-MS analysis method are provided in the following section on yield determination from analytical-scale reactions.)

**Enantiomeric ratio (e.r.) determination using Marfey's analysis.** Marfey's analysis is widely used to determine the stereochemical purity of amino acid products[3,4]. The protocol we used here is similar to that we described in our previous report[5]. To a 2.0 mL centrifuge tube were added 50 µL of the reaction mixture that was selected from the plate, 1.0 M aq. $NaHCO_3$ (100 µL), DMSO (50 µL) and a stock solution of Marfey's reagent, namely (*S*)-1-fluoro-2-4-dinitrophenyl-5-L-alanine amide ((*S*)-FDAA) or (*R*)-1-fluoro-2-4-dinitrophenyl-5-L-alanine amide ((*R*)-FDAA) (75 µL, 20 mM in acetone). The tubes were then placed in a microplate shaker and shaken at room temperature and 800 rpm for 8–12 h. The reaction mixture was diluted with 1:1 $CH_3CN$/1 M aq. HCl (1000 µL) and centrifuged at 15,000 rpm for 15 min to afford a clear solution which was then analyzed by LC-MS.

Short LC-MS analysis method (method A) is as follows (all analyses were performed using this method unless otherwise noted): Poroshell 120 EC-C18 column (4.6 × 50 mm, 4 µm); flow rate = 1.0 mL/min. The elution program was as follows: hold at 5% $CH_3CN$ (0.1% formic acid) in $H_2O$ (0.1% formic acid) for 0.5 min; ramp to 30% $CH_3CN$ over 2.5 min; further increase to 60% $CH_3CN$ over 7.0 min; ramp to 95% $CH_3CN$ over 0.5 min; hold at 95% $CH_3CN$ for 0.5 min. The total analysis time was 12.5 min.

Long LC-MS analysis method (method B) is as follows: Poroshell 120 EC-C18 column (4.6 × 150 mm, 2.7 µm); a flow rate = 1.0 mL/min. The elution program was as follows: hold at 5% $CH_3CN$ (0.1% formic acid) in $H_2O$ (0.1% formic acid) for 1 min; ramp to 30% $CH_3CN$ over 4 min; further increase to 55% $CH_3CN$ over 17 min; ramp to 95% $CH_3CN$ over 3 min; hold at 95% $CH_3CN$ for 1 min. The total runtime was 27 min.

**Expression and purification of *Pf*PLP$_{\mathbf{Alk}}$ variants.** A single colony from $LB_{amp}$ agar plate was picked using a sterile toothpick and cultured in $LB_{amp}$ media (25 mL) at 37 °C and 230 rpm overnight. A 15 mL aliquot of this preculture was used to inoculate a 1 L of $TB_{amp}$ media in a 4 L Erlenmeyer flask, and the expression culture was incubated at 37 °C with shaking at 230 rpm for ca. 3 h, until the $OD_{600}$ reached ca. 1.2. The culture was then cooled on ice for 20 min and induced with isopropyl-*β*-D-thiogalatopyranoside (IPTG) to a final concentration of 1.0 mM. Protein expression was conducted at 20 °C and 200 rpm for 20 h. *E. coli* cells were harvested by centrifugation at 4 °C and 4,500 rpm for 10 min using a Thermo Scientific Sorvall Lynx 6000 superspeed centrifuge. The cell pellet was then frozen with liquid nitrogen.

For protein purification, frozen cells were resuspended in HisTrap buffer A (25 mM KPi, 100 mM NaCl, 20 mM imidazole, pH 8.0, 2−3 mL/g of cell wet weight), following by the addition of the 10.0 mM PLP (stock solution in 50 mM KPi buffer, pH 8.0) to a final concentration of 200 μM. The mixture was suspended and lysed by sonication using a Qsonica Q500 sonicator. To pellet cell debris, lysates were centrifuged using a Lynx 6000 superspeed centrifuge (15,000 *g*, 60 min, 4 °C). L-*Pf*$PLP_{Alk}$ or D-*Pf*$PLP_{Alk}$ containing a C-terminal 6×His tag was purified with a Ni-NTA column (5.0 mL HisTrap HP column, GE Healthcare, Piscataway, NJ) using an AKTA Start protein purification system (GE healthcare). Proteins were eluted on a linear gradient from Histrap buffer A (25 mM KPi, 100 mM NaCl, 20 mM imidazole, pH 8.0) to Histrap buffer B (25 mM KPi, 100 mM NaCl, 500 mM imidazole, pH 8.0) over 10 column volumes (CVs). Proteins eluted at approximately 100 mM imidazole. Fractions containing L-*Pf*$PLP_{Alk}$ or D-*Pf*$PLP_{Alk}$ were combined, followed by the addition of a PLP stock solution (10.0 mM PLP in 50 mM KPi buffer, pH 8.0) to a final PLP concentration of 200 μM. The enzyme solution was concentrated and subjected to three rounds of buffer exchange with the storage buffer (buffer C, 50 mM KPi, pH 8.0) using ultracentrifugal filters (30 kDa molecular weight cut-off, Amicon Ultra, Sigma Millipore) to remove excess salt and imidazole. Protein concentration was measured with Nanodrop and normalized to ca. 50 mg/mL. Typically, 1 L of $TB_{amp}$ expression culture provided 250–300 mg protein. Protein concentration was also determined by NanoDrop and BCA assay prior to the setup of biocatalytic reactions. Concentrated proteins were aliquoted and flash-frozen in liquid $N_2$ and stored at –80 °C until further use. 10% glycerol was added to the final storage buffer as a cryoprotectant to enhance long-term protein stability upon storage.

**Protein concentration determination.** The concentration of holo PLP enzymes was also determined by PLP-specific UV-visible assays. 75 μL 0.2 M NaOH solution was added into a solution of purified PLP enzyme (5 μL) and carefully mixed. The solution was transformed into a quartz cuvette for UV-vis spectroscopic analysis. PLP concentration was determined using the equation: $\varepsilon^{PLP} = 6600\ M^{-1}cm^{-1}$ at 390 nm[6]. For PLP enzymes prepared using the procedure described above, the PLP cofactor loading (holo enzyme content) was (100 ± 10)%.

**General Procedure for analytical scale reactions using purified PLP radical enzymes**

**Photobiocatalytic synthesis of L-amino acids:** L-*Pf*PLP$_{Alk}$ (ca. 50 mg/mL) was allowed to thaw and kept on ice. In a Coy anaerobic chamber, the following stock solutions were prepared: organoboron substrate (RB(OH)$_2$, RBpin or RBF$_3$K) (100 mM in degassed DMSO), RhB (10.0 mM in degassed DMSO), and L-serine (500 mM in degassed 200 mM KPi buffer, pH 7.5). To a one-dram vial containing a stir bar were added 450 μL KPi buffer (200 mM, pH 7.5), 20 μL organoboron stock solution, 20 μL L-serine stock solution, 20 μL RhB stock solution, followed by L-*Pf*PLP$_{Alk}$ solution (1.0 mol%, ca. 17.0 μL, or 1.5 mol%, ca. 25.5 μL; the specific volume of the enzyme stock solution was determined by the concentration of the specific enzyme sample used). The reaction vial was then sealed and removed from the Coy anaerobic chamber and put in a 3S-Tech AF2 photoreactor at 50 °C. The reaction mixture was allowed to stir at 300 rpm for 3 min and then illuminated at 440 nm (power output: 4 W or 5 W) for 10 h.

**Photobiocatalytic synthesis of D-amino acids using D-*Pf*PLP$_{Alk1}$:** D-*Pf*PLP$_{Alk1}$ (ca. 50 mg/mL) was allowed to thaw and kept on ice. In a Coy anaerobic chamber, the following stock solutions were prepared: organoboron substrate (RB(OH)$_2$, RBpin or RBF$_3$K) (100 mM in degassed DMSO), Eosin B (5.0 mM in degassed DMSO), and L-serine (500 mM in degassed 200 mM KPi buffer, pH 9.0). To a one-dram vial containing a stir bar were added 450 μL KPi buffer (200 mM, pH 9.0), 20 μL organoboron substrate stock solution, 20 μL L-serine stock solution and 20 μL Eosin B stock solution, followed by the addition of D-*Pf*PLP$_{Alk1}$ solution (1.0 mol%, ca. 17 μL or 1.5 mol%, ca. 25.5 μL). The reaction mixture was left in a 3S-Tech AF2 photoreactor and allowed to stir at 300 rpm for 3 min and then illuminated at 525 nm (power output: 3 W) for 10 h.

**Photobiocatalytic synthesis of D-amino acids using D-*Pf*PLP$_{Alk2}$:** D-*Pf*PLP$_{Alk2}$ (ca. 50 mg/mL) was allowed to thaw and kept on ice. In a Coy anaerobic chamber, the following stock solutions were prepared: organoboron substrate (RB(OH)$_2$, RBpin or RBF$_3$K) (100 mM in degassed DMSO), RhB (10.0 mM in degassed DMSO), and L-serine (500 mM in degassed 200 mM KPi buffer, pH 9.0). To a one-dram vial containing a stir bar were added 450 μL KPi buffer (200 mM, pH 9.0), 20 μL organoboron substrate stock solution, 20 μL L-serine stock solution and 20 μL RhB stock solution, followed by the addition of D-*Pf*PLP$_{Alk2}$ solution (1.0 mol%, ca. 17 μL or 1.5 mol%, ca. 25.5 μL). The reaction mixture was left in a 3S-Tech AF2 photoreactor and allowed to stir at 300 rpm for 3 min and then illuminated at 440 nm (power output: 4 W) for 10 h.

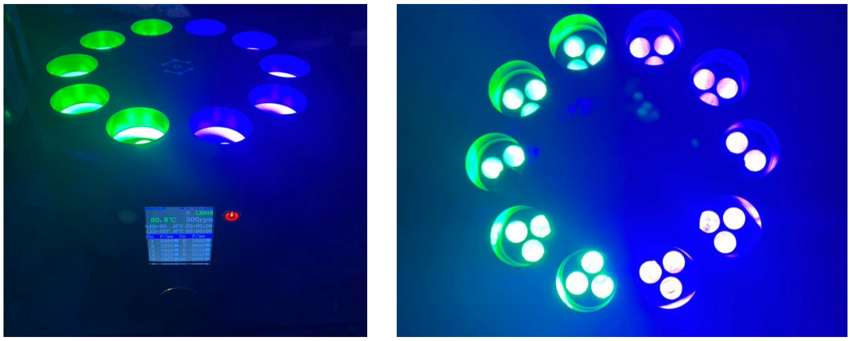

**Figure S3**. 3S-Tech AF2 photoreactor used in this study (wavelength: 525 nm, 440 nm).

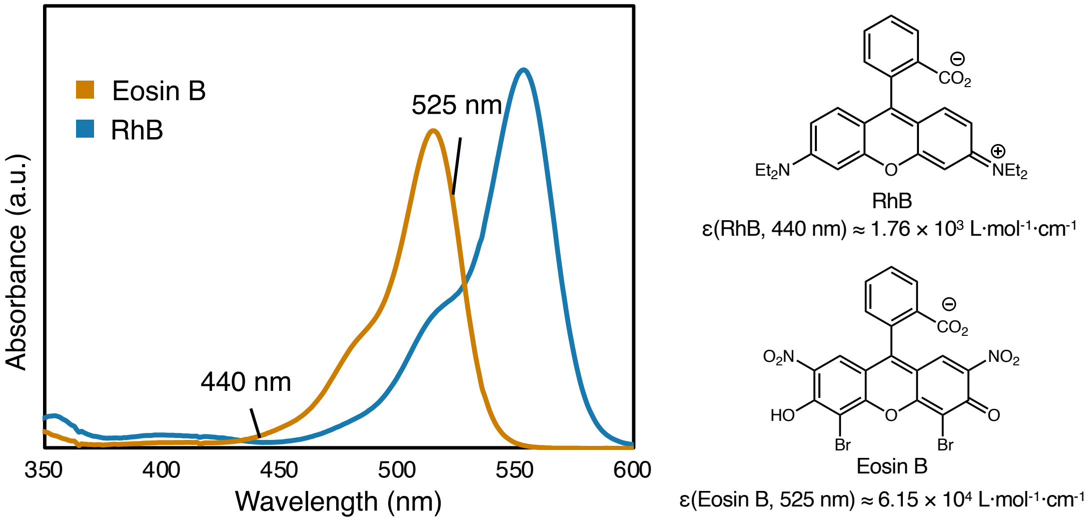

**Figure S4**. UV-visible spectra of rhodamine B (RhB) and eosin B (EB) in KPi buffer (pH = 8.0) used in this work

**Yield determination for UV-active amino acids using a PDA detector.**

**(a) Development of reverse-phase HPLC calibration curves using an internal standard**: Stock solutions of authentic products (40 mM in $CH_3CN$/1 M aq. HCl, 1:1, v/v) and internal standard (homophenylalanine, 40 mM in $CH_3CN$/1 M aq. HCl, 1:1, v/v) were freshly prepared. To a 1.5 mL microcentrifuge tube, 50 μL of the internal standard solution (4.0 μmol) was added, followed by 10, 20, 30, 40, or 50 μL of the authentic product solution. Subsequently, 940, 930, 920, 910, or 900 μL of $CH_3CN$/1 M aq. HCl (1:1, v/v) was added to bring the total volume to 1.0 mL. The mixtures were vortexed for 20 sec (three repetitions) to ensure thorough mixing and then transferred to 1.5 mL glass vials for HPLC analysis. Calibration curves were constructed by plotting the amount of product (y-axis) against the ratio of the product peak area to that of the internal standard (x-axis) obtained from HPLC analysis. Calibration samples were prepared to closely match the concentration range of analyte samples to ensure accurate interpolation.

Representative HPLC conditions were as follows: Poroshell 120 EC-C18 column (4.6 × 50 mm, 4 μm); flow rate = 1.0 mL/min; initial hold at 5% $CH_3CN$ (0.1% formic acid) in $H_2O$ (0.1% formic acid) for 0.30 min; linear gradient to 60% $CH_3CN$ over 3.45 min; ramp to 95% $CH_3CN$ over 0.25 min; hold at 95% $CH_3CN$ for 0.25 min; total runtime = 5.0 min.

(**b**) **Yield determination for analytical scale reactions**: Upon completion of the reaction, 50 μL of the homophenylalanine internal standard solution (40 mM in $CH_3CN$/1 M aq. HCl, 1:1, v/v) was added to each reaction well, followed by 1.0 mL of $CH_3CN$/1 M aq. HCl (1:1, v/v). The mixtures were vigorously mixed and centrifuged at 4,500 rpm for 15 min to remove insoluble material. A 1000 μL aliquot of the clear supernatant was transferred to a 1.5 mL glass vial for analysis. Product yields (e.g., for **3a**) were quantified using the calibration curves described above.

**Yield determination for UV-inactive amino acids.**

Product yields were determined using an external calibration method. Since the target amino acid products were UV-inactive in their native forms and could not be analyzed using a PDA detector, authentic standards were derivatized with 2,4-dinitrofluorobenzene (DNFB) to generate UV-active derivatives for yield determination.

**(a) Development of reverse-phase HPLC calibration curves amino acids derivatives.** Stock solutions of authentic products (2.0 mM in $CH_3CN$/1 M aq. HCl, 1:1, v/v) were freshly prepared. Aliquots of 5, 10, 15, 20, or 25 μL of each stock solution were transferred into individual 2.0 mL centrifuge tubes. To each tube, 100 μL of 1.0 M aqueous $NaHCO_3$, 75 μL of DNFB stock solution (20 mM in acetone), and 70, 65, 60, 55, or 50 μL of DMSO, respectively, were added in succession. The mixtures were incubated at room temperature in a microplate shaker at 800 rpm for 8–12 h to complete this DNFB derivatization. The reaction progress could be monitored by LC-MS analysis. Upon completion, each reaction was quenched by the addition of 1000 μL of $CH_3CN$/1 M aq. HCl (1:1, v/v), followed by centrifugation at 15,000 rpm for 15 min to remove precipitates. The resulting clear supernatants were subjected to LC-MS analysis. Calibration curves were constructed by plotting the amount of authentic standard (y-axis) against the corresponding peak area at 330 nm (x-axis). Calibration samples were prepared to closely match the concentration range of analyte samples to ensure accurate interpolation.

Representative LC-MS method: Poroshell 120 EC-C18 column (4.6 × 50 mm, 4 µm); flow rate = 1.0 mL/min. The gradient was programmed as follows: hold at 5% $CH_3CN$ (0.1% formic acid in $H_2O$) for 0.5 min; ramp to 60% $CH_3CN$ over 2.5 min; further ramp to 95% $CH_3CN$ over 3.0 min; hold at 95% $CH_3CN$ for 2.0 min. Total run time: 10.0 min.

(**b**) **Yield determination for analytical scale reactions**: Upon the completion of photobiocatalytic reaction, 1.0 mL of $CH_3CN$/1 M aq. HCl (1:1, v/v) was added to quench the reaction, and the total reaction mass was recorded using an analytical balance. A 35 µL aliquot of the crude mixture was transferred to a 2.0 mL centrifuge tube and weighed precisely. To this aliquot, 100 µL of 1.0 M aqueous $NaHCO_3$, 75 µL of DNFB stock solution (20 mM in acetone), and 50 µL of DMSO were added sequentially. The mixtures were incubated at room temperature with shaking at 800 rpm for 8–12 h. Reactions were quenched with 1000 µL of $CH_3CN$/1 M aq. HCl (1:1, v/v) and centrifuged at 15,000 rpm for 15 min to remove any precipitate. The clear supernatants were analyzed under the same LC-MS conditions described above. Product concentrations were interpolated from the calibration curve, and the overall yield was calculated based on the known initial substrate concentration and the total mass of the reaction mixture.

**Procedure for 100 mg-scale photobiocatalytic synthesis of non-canonical amino acids**

**Preparative photobiocatalytic L-amino acid synthesis:** L-*Pf*PLP$_{Alk1}$ (ca. 66 mg/mL, 1 mL/tube) was thawed on ice prior to use. All subsequent manipulations were carried out in a Coy anaerobic chamber. The following degassed stock solutions were prepared: (2-phenylethyl)boronic acid **1a** (100 mM in DMSO), RhB (10.0 mM in DMSO), and L-serine (500 mM in 200 mM KPi buffer, pH 9.0). To a 100 mL round-bottom flask equipped with a magnetic stir bar were added KPi buffer (58 mL, 200 mM, pH 7.5), (2-phenylethyl)boronic acid solution (3.3 mL, 0.33 mmol, 1.0 equiv), L-serine solution (3.3 mL, 1.7 mmol, 5.0 equiv), RhB solution (3.3 mL, 0.033 mmol, 10 mol%), and L-*Pf*PLP$_{Alk1}$ stock solution (ca. 3.2 mL, 1.5 mol%). The total reaction volume was around 71 mL, and the final concentrations of each reaction component were as follows: 4.7 mM (2-phenylethyl)boronic acid, 23.4 mM L-serine, 47.0 µM D-*Pf*PLP$_{Alk1}$, and 470 µM RhB. The flask was sealed, removed from the anaerobic chamber, and placed in a 50 °C water bath. After stirring at 450 rpm for 5 min, the mixture was irradiated with two Kessil LED lamps (440 nm, 45 W each, 10 cm distance between the LED lamp and the edge of the round-bottom flask) for 16 h. The reaction was quenched with a 1:1 mixture of $CH_3CN$/1 M aq. HCl (40 mL). Three batches of crude product were combined and concentrated *in vacuo* with the aid of a rotary evaporator to a final volume of <50 mL. MeOH (50 mL) was added, and the mixture was centrifuged at 4,500 rpm for 10 min using a Thermo Scientific Sorvall Lynx 6000 superspeed centrifuge. The supernatant was concentrated to near dryness, and the residue was re-dissolved in $H_2O$ (20 mL). The solution was filtered through a 2.2 μm membrane filter and loaded onto a C18 flash column (18 g) pre-equilibrated with $H_2O$. The column was washed with 10 CV of $H_2O$, followed by gradient elution (0–50% MeCN in $H_2O$ over 12 CV). Product-containing fractions were pooled and concentrated *in vacuo* with the aid of a rotary evaporator to afford the HCl adduct of L-**3a** as a white solid (167 mg, 0.73 mmol, 73% yield).

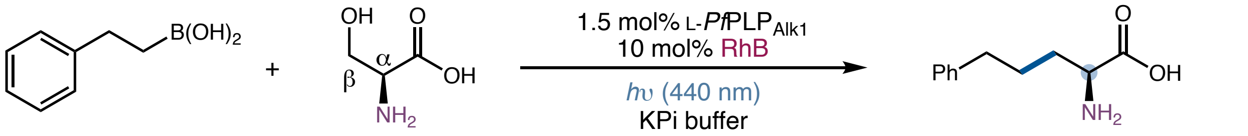

**1a** 0.33 mmol, 50 mg

**2a** 1.65 mmol, 174 mg

**L-3a** 73% yield, 95:5 e.r. 167 mg **L-3a**•HCl from 3 reactions (1 mmol in total)

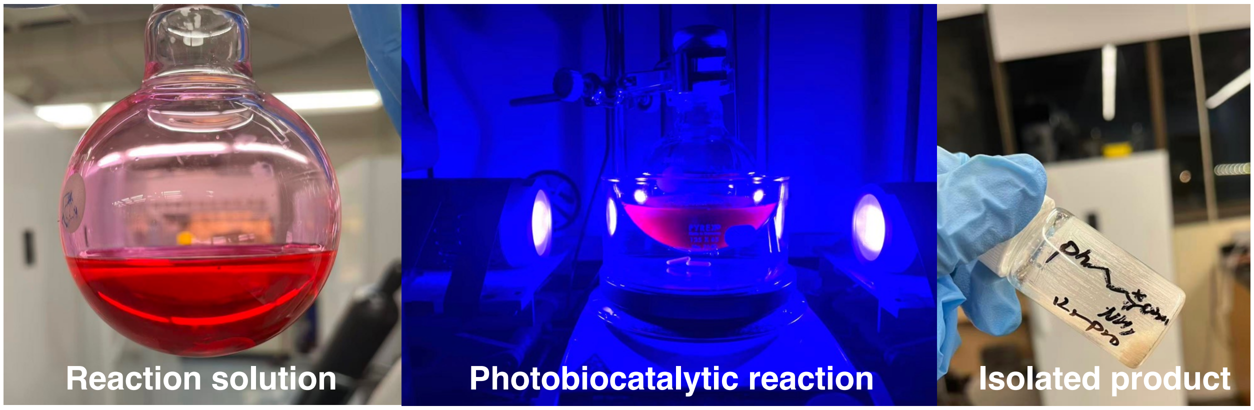

**Figure S5**. Preparative-scale photobiocatalytic reaction setup under blue light irradiation

**Preparative photobiocatalytic D-amino acid synthesis:** D-*Pf*PLP$_{Alk1}$ (ca. 55 mg/mL, 1 mL/tube) was thawed on ice prior to use. All subsequent manipulations were carried out in a Coy anaerobic chamber. The following degassed stock solutions were prepared: (2-phenylethyl)boronic acid **1a**

(100 mM in DMSO), Eosin B (5.0 mM in DMSO), and L-serine (500 mM in 200 mM KPi buffer, pH 9.0). To a 100 mL round-bottom flask equipped with a magnetic stir bar were added KPi buffer (32 mL, 200 mM, pH 9.0), (2-phenylethyl)boronic acid solution (2 mL, 0.2 mmol, 1.0 equiv), L-serine solution (2 mL, 1.0 mmol, 5.0 equiv), Eosin B solution (2 mL, 0.01 mmol, 5 mol%), and D-*Pf*PLP$_{Alk1}$ stock solution (ca. 1.54 mL, 1.0 mol%). The total reaction volume was around 40 mL, and the final concentrations of each reaction component were as follows: 5.0 mM (2-phenylethyl)boronic acid, 25.0 mM L-serine, 50.0 µM D-*Pf*PLP$_{Alk1}$, and 250 µM Eosin B. The flask was sealed, removed from the anaerobic chamber, and placed in a 50 °C water bath. After stirring at 450 rpm for 5 min, the mixture was irradiated with two Kessil LED lamps (525 nm, 22.5 W each, 10 cm distance between the LED lamp and the edge of the round-bottom flask) for 16 h. The reaction was quenched with a 1:1 mixture of $CH_3CN$/1 M aq. HCl (40 mL). Five batches of crude product were combined and concentrated *in vacuo* with the aid of a rotary evaporator to a final volume of <50 mL. MeOH (50 mL) was added, and the mixture was centrifuged at 4,500 rpm for 10 min using a Thermo Scientific Sorvall Lynx 6000. The supernatant was concentrated to near dryness, and the residue was re-dissolved in $H_2O$ (20 mL). The solution was filtered through a 2.2 µm membrane filter and loaded onto a C18 flash column (18 g) pre-equilibrated with $H_2O$. The column was washed with 10 CV of $H_2O$, followed by gradient elution (0–50% MeCN in $H_2O$ over 12 CV). Product-containing fractions were pooled and concentrated *in vacuo* with the aid of a rotary evaporator to afford the HCl adduct of D-**3a** as a white solid (154 mg, 0.67 mmol, 67% yield).

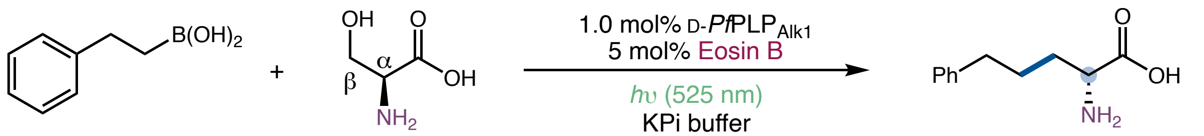

**1a**
0.2 mmol, 30 mg

**2a**
1.0 mmol, 105 mg

D-**3a**
67% yield, 5:95 e.r.
154 mg D-**3a**•HCl from 5 reactions
(1 mmol in total)

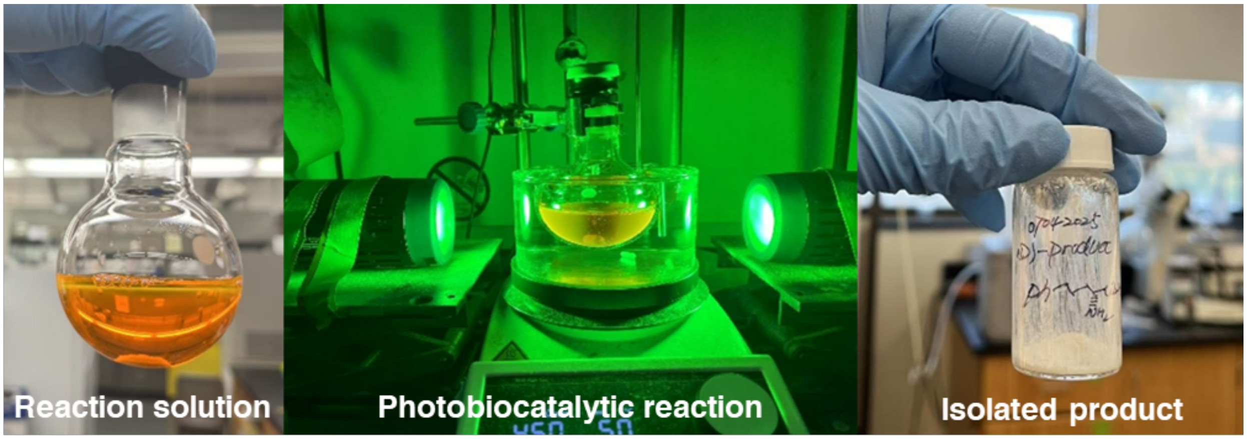

**Figure S6**. Preparative-scale photobiocatalytic reaction setup under green light irradiation

#### II. Directed evolution of *Pf*PLP$_{Alk}$ variants

##### Evolutionary trajectory of L-*Pf*PLP$_{Alk}$

Prior to mutagenesis, molecular docking simulations were conducted using AutoDock5 to identify active-site residues in proximity to the product-bound PLP covalent intermediate. Additionally, analysis of the crystal structure of L-*Pf*PLP$^{\beta}$ (PDB: 5VM5) using CAVER enabled visualization of the substrate access tunnel (teal mesh in Figure S7). Residues lining the tunnel and within the active site were selected for site-saturation mutagenesis (SSM) and screening to identify beneficial mutations.

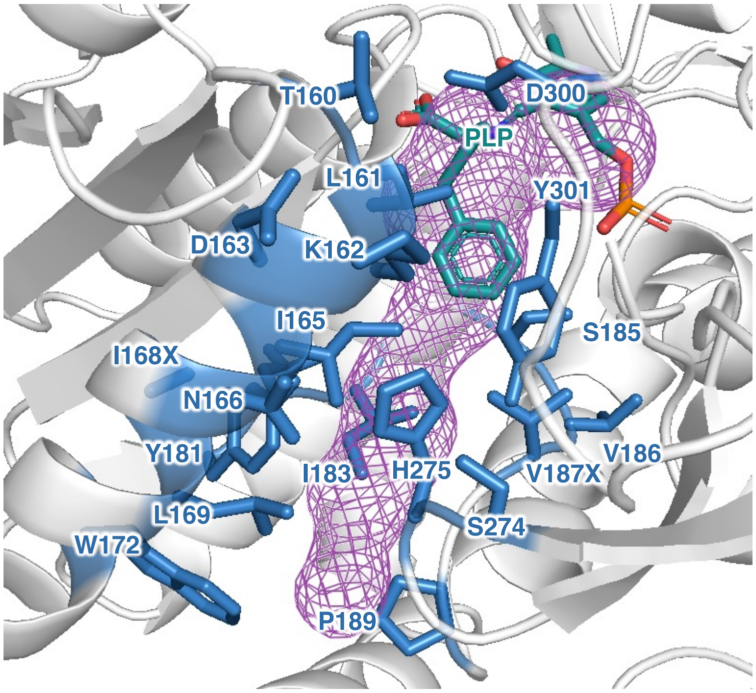

**Figure S7.** Docking model and substrate tunnel for L-*Pf*PLP$^{\beta}$ with (*S*)-**3a**.

**Table S1.** Summary of directed evolution of L-*Pf*PLP$_{Alk}$ (L-*Pf*PLP$^{\beta}$ Y301H I165T L161S Y181H H275D N166H T160K)

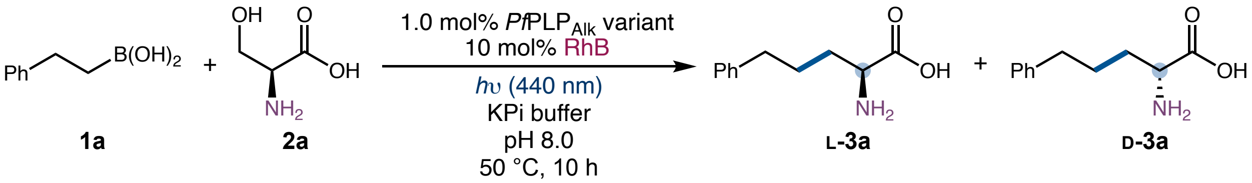

| parent | sites targeted by SSM | selection criterion | beneficial mutation |
|---|---|---|---|
| L-*Pf*PLP$^{\beta}$ Y301H (P) | H301X, I165X, L161X | activity | I165T |
| P I165T | I183X, L161X, W172X | activity, enantioselectivity | L161S |
| P I165T L161S | Y181X, S185X, K162X | activity, enantioselectivity | Y181H |
| P I165T L161S Y181H | H275X, D300X, D163X | enantioselectivity | H275D |
| P I165T L161S Y181H H275D | N166X, S274X, L182X | enantioselectivity | N166H |
| P I165T L161S Y181H H275D N166H | V186X, V187X, P189X, T160X | activity | T160K |
| P I165T L161S Y181H H275D N166H T160K | L169X, K162X, A168X | activity or enantioselectivity | No hits |

**Table S2.** Primers used for site-saturation mutagenesis using the 22-c trick method for directed evolution of L-*Pf*PLP$_{Alk}$ (L-*Pf*PLP$^{β}$ Y301H I165T L161S Y181H H275D N166H T160K)

| DNA template | primer name | primer sequence |
|---|---|---|
| L-*Pf*PLP$^{β}$ Y301H | H301X_fwd_NDT | CTCCATCGCACCAGGTCTGGAT**NDT**CCAGGTGTTG |
| | H301X_fwd_VHG | CTCCATCGCACCAGGTCTGGAT**VHG**CCAGGTGTTG |
| | H301X_fwd_TGG | CTCCATCGCACCAGGTCTGGAT**TGG**CCAGGTGTTG |
| | H301X_rev | CCAGACCTGGTGCGATGGAG |
| L-*Pf*PLP$^{β}$ Y301H | L161X_fwd_NDT | GTTAACTCCGGTTCTCGCACC**NDT**AAAGACGCAATC |
| | L161X_fwd_VHG | GTTAACTCCGGTTCTCGCACC**VHG**AAAGACGCAATC |
| | L161X_fwd_TGG | GTTAACTCCGGTTCTCGCACC**TGG**AAAGACGCAATC |
| | L161X_rev | GGTGCGAGAACCGGAGTTAAC |
| L-*Pf*PLP$^{β}$ Y301H | I165X_fwd_NDT | GGTTCTCGCACCCTGAAAGACGCA**NDT**AACGAGGCTC |
| | I165X_fwd_VHG | GGTTCTCGCACCCTGAAAGACGCA**VHG**AACGAGGCTC |
| | I165X_fwd_TGG | GGTTCTCGCACCCTGAAAGACGCA**TGG**AACGAGGCTC |
| | I165X_rev | GTCTTTCAGGGTGCGAGAACCGGAG |
| L-*Pf*PLP$^{β}$ Y301H I165T | L161X_fwd_NDT | GTTAACTCCGGTTCTCGCACC**NDT**AAAGACGCAAC |
| | L161X_fwd_VHG | GTTAACTCCGGTTCTCGCACC**VHG**AAAGACGCAAC |
| | L161X_fwd_TGG | GTTAACTCCGGTTCTCGCACC**TGG**AAAGACGCAAC |
| | L161X_rev | GGTGCGAGAACCGGAGTTAACTG |
| L-*Pf*PLP$^{β}$ Y301H I165T | W172X_fwd_NDT | CAACGAACGAGGCTCTGCGTGAT**NDT**GTGGCTACTTTTG |
| | W172X_fwd_VHG | CAACGAACGAGGCTCTGCGTGAT**VHG**GTGGCTACTTTTG |
| | W172X_fwd_TGG | CAACGAACGAGGCTCTGCGTGAT**TGG**GTGGCTACTTTTG |
| | W172X_rev | CACGCAGAGCCTCGTTCGTTGC |
| L-*Pf*PLP$^{β}$ Y301H I165T | I183X_fwd_NDT | CTACTTTTGAATACACCCACTACCTA**NDT**GGTTCCGTGG |
| | I183X_fwd_VHG | CTACTTTTGAATACACCCACTACCTA**VHG**GGTTCCGTGG |
| | I183X_fwd_TGG | CTACTTTTGAATACACCCACTACCTA**TGG**GGTTCCGTGG |
| | I183X_rev | GTGGGTGTATTCAAAAGTAGCCAC |
| L-*Pf*PLP$^{β}$ Y301H I165T | K162X_fwd_NDT | CTCCGGTTCTCGCACCAGT**NDT**GACGCAACGAAC |
| | K162X_fwd_VHG | CTCCGGTTCTCGCACCAGT**VHG**GACGCAACGAAC |

| | | |
|---|---|---|
| L161S | K162X_fwd_TGG | CTCCGGTTCTCGCACCAGT**TGG**GACGCAACGAAC |
| | K162X_rev | CACGGAACCGATTAGGTAGTGG |
| L-*Pf*PLP$^{\beta}$ Y301H I165T L161S | Y181X_fwd_NDT | CTACTTTTGAATACACCCAC**NDT**CTAATCGGTTC |
| | Y181X_fwd_VHG | CTACTTTTGAATACACCCAC**VHG**CTAATCGGTTC |
| | Y181X_fwd_TGG | CTACTTTTGAATACACCCAC**TGG**CTAATCGGTTC |
| | Y181X_rev | GTGGGTGTATTCAAAAGTAGCCAC |
| L-*Pf*PLP$^{\beta}$ Y301H I165T L161S | S185X_fwd_NDT | GAATACACCCACTACCTAATCGGT**NDT**GTGGTCGGTC |
| | S185X_fwd_VHG | GAATACACCCACTACCTAATCGGT**VHG**GTGGTCGGTC |
| | S185X_fwd_TGG | GAATACACCCACTACCTAATCGGT**TGG**GTGGTCGGTC |
| | S185X_rev | GATTAGGTAGTGGGTGTATTCAAAAGTAG |
| L-*Pf*PLP$^{\beta}$ Y301H I165T L161S Y181H | D163X_fwd_NDT | CTCCGGTTCTCGCACCAGTAAA**NDT**GCAACGAACG |
| | D163X_fwd_VHG | CTCCGGTTCTCGCACCAGTAAA**VHG**GCAACGAACG |
| | D163X_fwd_TGG | CTCCGGTTCTCGCACCAGTAAA**TGG**GCAACGAACG |
| | D163X_rev | CTGGTGCGAGAACCGGAGTTAAC |
| L-*Pf*PLP$^{\beta}$ Y301H I165T L161S Y181H | H275X_fwd_NDT | GGTCAGGTTGGTGTGTCC**NDT**GGCATGCTGTC |
| | H275X_fwd_VHG | GGTCAGGTTGGTGTGTCC**VHG**GGCATGCTGTC |
| | H275X_fwd_TGG | GGTCAGGTTGGTGTGTCC**TGG**GGCATGCTGTC |
| | H275X_rev | GGACACACCAACCTGACCTG |
| L-*Pf*PLP$^{\beta}$ Y301H I165T L161S Y181H | D300X_fwd_NDT | CTCCATCGCACCAGGTCTG**NDT**CATCCAGGTG |
| | D300X_fwd_VHG | CTCCATCGCACCAGGTCTG**VHG**CATCCAGGTG |
| | D300X_fwd_TGG | CTCCATCGCACCAGGTCTG**TGG**CATCCAGGTG |
| | D300X_rev | CAGACCTGGTGCGATGGAGTG |
| L-*Pf*PLP$^{\beta}$ Y301H I165T L161S Y181H H275D | N166X_fwd_NDT | CACCAGTAAAGACGCAACG**NDT**GAGGCTCTGC |
| | N166X_fwd_VHG | CACCAGTAAAGACGCAACG**VHG**GAGGCTCTGC |
| | N166X_fwd_TGG | CACCAGTAAAGACGCAACG**TGG**GAGGCTCTGC |
| | N166X_rev | CGTTGCGTCTTTACTGGTGCGAG |
| L-*Pf*PLP$^{\beta}$ Y301H I165T L161S Y181H | S274X_fwd_NDT | GCAGGTCAGGTTGGTGTG**NDT**GATGGCATGC |
| | S274X_fwd_VHG | GCAGGTCAGGTTGGTGTG**VHG**GATGGCATGC |
| | S274X_fwd_TGG | GCAGGTCAGGTTGGTGTG**TGG**GATGGCATGC |

| | | |
|---|---|---|
| H275D | S274X_rev | CACACCAACCTGACCTGCGTTC |
| L-*Pf*PLP$^{\beta}$ | L182X_fwd_NDT | CTTTTGAATACACCCACCAT**NDT**ATCGGTTCCGTG |
| Y301H I165T | L182X_fwd_VHG | CTTTTGAATACACCCACCAT**VHG**ATCGGTTCCGTG |
| L161S Y181H | L182X_fwd_TGG | CTTTTGAATACACCCACCAT**TGG**ATCGGTTCCGTG |
| H275D | L182X_rev | GGTGGGTGTATTCAAAAGTAG |
| L-*Pf*PLP$^{\beta}$ | V186X_fwd_NDT | CCACCATCTAATCGGTTCC**NDT**GTCGGTCCAC |
| Y301H I165T | V186X_fwd_VHG | CCACCATCTAATCGGTTCC**VHG**GTCGGTCCAC |
| L161S Y181H | V186X_fwd_TGG | CCACCATCTAATCGGTTCC**TGG**GTCGGTCCAC |
| H275D N166H | V186X_rev | GGAACCGATTAGATGGTGGGTG |
| L-*Pf*PLP$^{\beta}$ | V187X_fwd_NDT | CCATCTAATCGGTTCCGTG**NDT**GGTCCACATC |
| Y301H I165T | V187X_fwd_VHG | CCATCTAATCGGTTCCGTG**VHG**GGTCCACATC |
| L161S Y181H | V187X_fwd_TGG | CCATCTAATCGGTTCCGTG**TGG**GGTCCACATC |
| H275D N166H | V187X_rev | CACGGAACCGATTAGATGGTG |
| L-*Pf*PLP$^{\beta}$ | P189X_fwd_NDT | CTAATCGGTTCCGTGGTCGGT**NDT**CATCCGTATC |
| Y301H I165T | P189X_fwd_VHG | CTAATCGGTTCCGTGGTCGGT**VHG**CATCCGTATC |
| L161S Y181H | P189X_fwd_TGG | CTAATCGGTTCCGTGGTCGGT**TGG**CATCCGTATC |
| H275D N166H | P189X_rev | CGACCACGGAACCGATTAGATG |
| L-*Pf*PLP$^{\beta}$ | T160X_fwd_NDT | CCAGTTAACTCCGGTTCTCGC**NDT**AGTAAAGACG |
| Y301H I165T | T160X_fwd_VHG | CCAGTTAACTCCGGTTCTCGC**VHG**AGTAAAGACG |
| L161S Y181H | T160X_fwd_TGG | CCAGTTAACTCCGGTTCTCGC**TGG**AGTAAAGACG |
| H275D N166H | T160X_rev | CGAGAACCGGAGTTAACTGGAATTAC |
| L-*Pf*PLP$^{\beta}$ | K162X_fwd_NDT | CTCCGGTTCTCGCACCAGTNDTGACGCAACGCAT |
| Y301H I165T | K162X_fwd_VHG | CTCCGGTTCTCGCACCAGTVHGGACGCAACGCAT |
| L161S Y181H | K162X_fwd_TGG | CTCCGGTTCTCGCACCAGTTGGGACGCAACGCAT |
| H275D N166H T160K | K162X_rev | CTGGTGCGAGAACCGGAGTTAACTG |
| L-*Pf*PLP$^{\beta}$ | A168X_fwd_NDT | GTAAAGACGCAACGCATGAG**NDT**CTGCGTGATTG |
| Y301H I165T | A168X_fwd_VHG | GTAAAGACGCAACGCATGAG**VHG**CTGCGTGATTG |
| L161S Y181H | A168X_fwd_TGG | GTAAAGACGCAACGCATGAG**TGG**CTGCGTGATTG |
| H275D N166H | A168X_rev | CTCATGCGTTGCGTCTTTACTGG |

| | | |
|---|---|---|
| T160K | | |
| L-*Pf*PLP$^{\beta}$ L-*Pf*PLP$^{\beta}$ Y301H I165T L161S Y181H H275D N166H T160K | L169X_fwd_NDT | TAAAGACGCAACGCATGAGGCT**NDT**CGTGATTGGGTG |
| | L169X_fwd_VHG | TAAAGACGCAACGCATGAGGCT**VHG**CGTGATTGGGTG |
| | L169X_fwd_TGG | TAAAGACGCAACGCATGAGGCT**TGG**CGTGATTGGGTG |
| | L169X_rev | CCTCATGCGTTGCGTCTTTACTG |

**Table S3.** Enzyme activity and selectivity summary for photobiocatalytic synthesis of L-**3a**

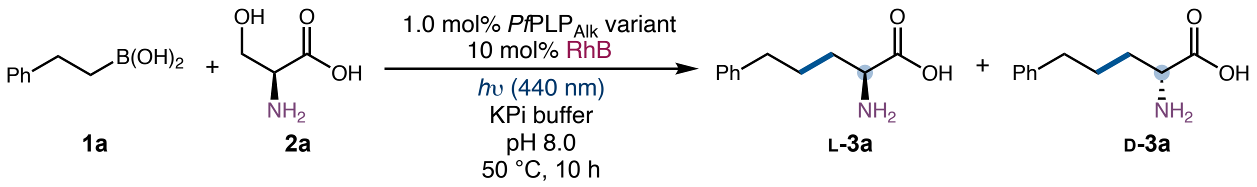

| *Pf*PLP$^{\beta}$ variant | yield of **3a** | e.r. (L-**3a**:D-**3a**) |
|---|---|---|
| L-*Pf*PLP$^{\beta}$ | (1.9 ± 0.2)% | 29:71 |
| L-*Pf*PLP$^{\beta}$ Y301H (P) | (15 ± 1)% | 52:48 |
| P I165T | (23 ± 4)% | 40:60 |
| P I165T L161S | (30 ± 3)% | 68:32 |
| P I165T L161S Y181H | (48 ± 3)% | 82:18 |
| P I165T L161S Y181H H275D | (50 ± 3)% | 88:12 |
| P I165T L161S Y181H H275D N166H | (53 ± 2)% | 94:6 |
| **P I165T L161S Y181H H275D N166H T160K (L-*Pf*PLP$_{Alk}$)** | **(63 ± 1)%** | **93:7** |

Reaction conditions: **1a** (4.0 mM), **2a** (20.0 mM), 1.0 mol% *Pf*PLP$_{Alk}$ variant, 10 mol% RhB, *hν* (440 nm, 5 W), 200 mM KPi buffer (pH = 8.0), DMSO (8% v/v), 50 °C, 10 h. All the reactions were performed in triplicates and averaged yields were reported.

**Table S4.** pH effect on photobiocatalytic synthesis of L-**3a**

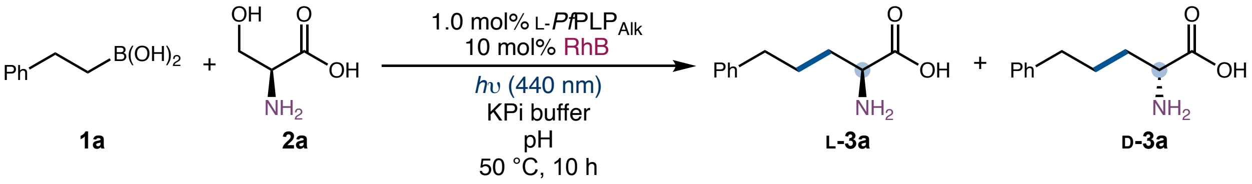

| entry | pH | yield of **3a** | e.r. (L-**3a**:D-**3a**) |
|---|---|---|---|
| 1 | 8.0 | 64% | 93:7 |
| **2** | **7.5** | **63%** | **95:5** |
| 3 | 7.0 | 57% | 96:4 |
| 4 | 6.5 | 53% | 97:3 |

Reaction conditions: **1a** (4.0 mM), **2a** (20.0 mM), 1.0 mol% L-*Pf*PLP$_{Alk}$, 10 mol% RhB, *h*ν (440 nm, 5 W), 200 mM KPi buffer, DMSO (8% v/v), 50 °C, 10 h.

**Table S5.** Photoredox catalyst effect on photobiocatalytic synthesis of L-**3a**

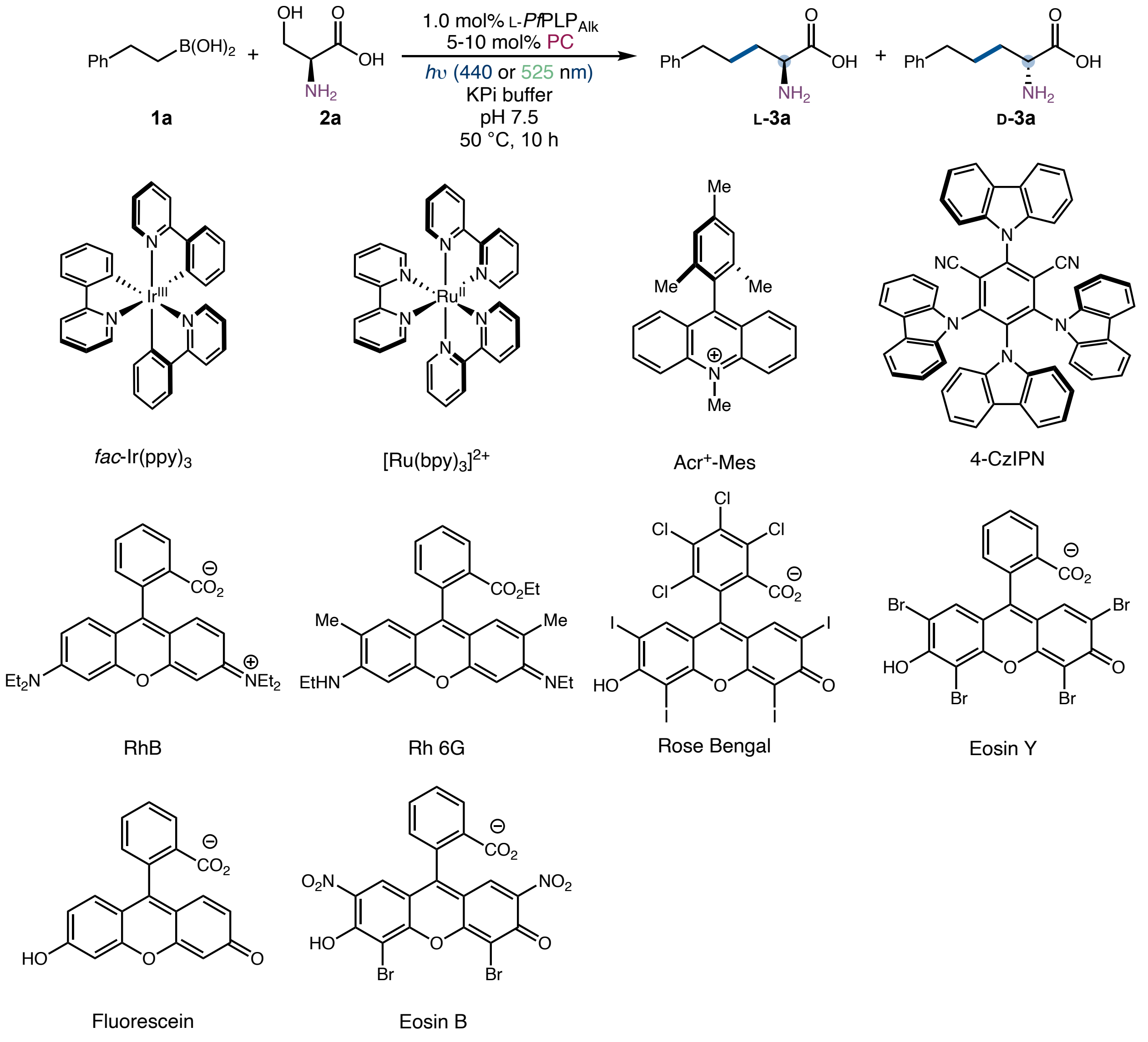

| entry | photocatalyst | irradiation wavelength | yield of **3a** | e.r. (L-**3a**:D-**3a**) |
|---|---|---|---|---|
| 1 | *fac*-Ir(ppy)$_3$ (5 mol%) | 440 nm | 0% | - |
| 2 | Ru(bpy)$_3$Cl$_2$ (5 mol%) | 440 nm | 0% | - |
| 3 | MesAcr (10 mol%) | 440 nm | <1% | 91:9 |
| 4 | 4CzIPN (10 mol%) | 440 nm | trace | - |

| | | | | |
|---|---|---|---|---|
| **5** | **RhB (10 mol%)** | **440 nm** | **63%** | **95:5** |
| 6 | Rh6G (10 mol%) | 440 nm | 41% | 95:5 |
| 7 | Rose Bengal (10 mol%) | 440 nm | 29% | 95:5 |
| 8 | Eosin Y (10 mol%) | 440 nm | 30% | 94:6 |
| 9 | Fluorescein (10 mol%) | 440 nm | 26% | 94:6 |
| 10 | Eosin B (10 mol%) | 440 nm | 23% | 95:5 |
| 11 | RhB (5 mol%) | 525 nm | 5% | 95:5 |
| 12 | Eosin B (5 mol%) | 525 nm | 36% | 95:5 |
| **13** | **RhB (10 mol%) (1.5 mol% L-*Pf*PLP$_{Alk}$)** | **440 nm** | **83%** | **95:5** |

Reaction conditions: **1a** (4.0 mM), **2a** (20.0 mM), 1.0 mol% L-*Pf*PLP$_{Alk}$, 5-10 mol% photocatalyst, *h*ν (440 nm and 5.0 W; or 525 nm and 3.0 W), 200 mM KPi buffer (pH = 7.5), DMSO (8% v/v), 50 °C, 10 h.

**Table S6.** Effect of different radical precursors on photobiocatalytic synthesis of L-**3a**

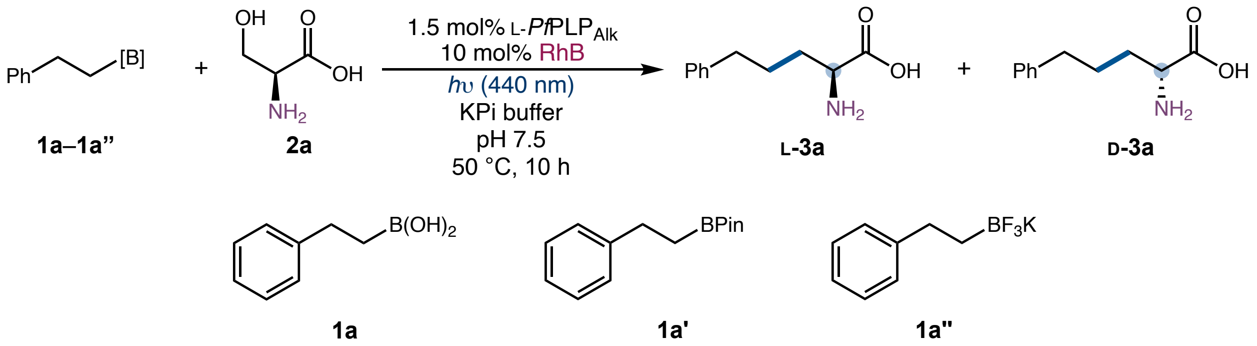

| entry | [B] | yield of **3a** | e.r. (L-**3a**:D-**3a**) |
|---|---|---|---|
| 1 | **1a** | 83% | 95:5 |
| 2 | **1a’** | 83% | 95:5 |
| 3 | **1a’’** | 81% | 95:5 |

Reaction conditions: **1a** (4.0 mM), **2a** (20.0 mM), 1.5 mol% L-*Pf*PLP$_{Alk}$, 10 mol% RhB, *h*ν (440 nm, 5.0 W), 200 mM KPi buffer (pH = 7.5), DMSO (8% v/v), 50 °C, 10 h.

As can be seen from results in Table S6, (2-phenylethyl)boronic acid (**1a**), (2-phenylethyl)boronic acid pinacol ester (**1a'**), and potassium 2-phenylethyl trifluoroborate (**1a''**) exhibited comparable activity under the standard conditions with the evolved enzyme L-*Pf*PLP$_{Alk}$.

**Table S7.** Enzyme activity and selectivity summary for photobiocatalytic synthesis of L-**5a**

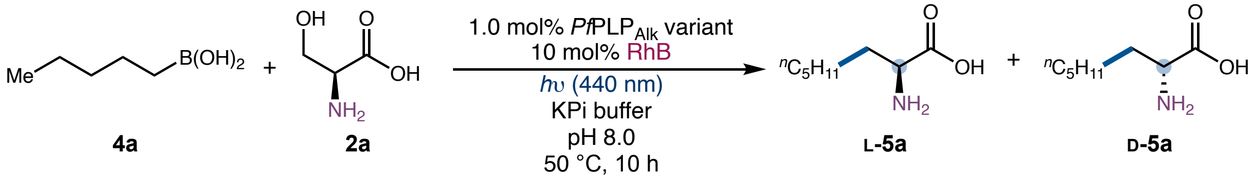

| *Pf*PLP$^{β}$ variant | yield of **5a** | e.r. (L-**5a**:D-**5a**) |
|---|---|---|
| L-*Pf*PLP$^{β}$ Y301H (P) | (12 ± 1)% | 83:17 |
| P I165T | (19 ± 4)% | 83:17 |
| P I165T L161S | (26 ± 3)% | 77:23 |
| P I165T L161S Y181H | (34 ± 3)% | 83:17 |
| P I165T L161S Y181H H275D | (27 ± 1)% | 94:6 |
| P I165T L161S Y181H H275D N166H | (39 ± 3)% | 95:5 |
| **P I165T L161S Y181H H275D N166H T160K (L-*Pf*PLP$_{Alk}$)** | **(59 ± 1)%** | **91:9** |

Reaction conditions: **4a** (4.0 mM), **2a** (20.0 mM), 1.0 mol% *Pf*PLP$_{Alk}$ variant, 10 mol% RhB, *h*ν (440 nm, 5.0 W), 200 mM KPi buffer (pH = 8.0), DMSO (8% v/v), 50 °C, 10 h. All the reactions were performed in triplicates and averaged yields were reported.

**Table S8.** Selective conditions optimization results for photobiocatalytic synthesis of L-**5a**

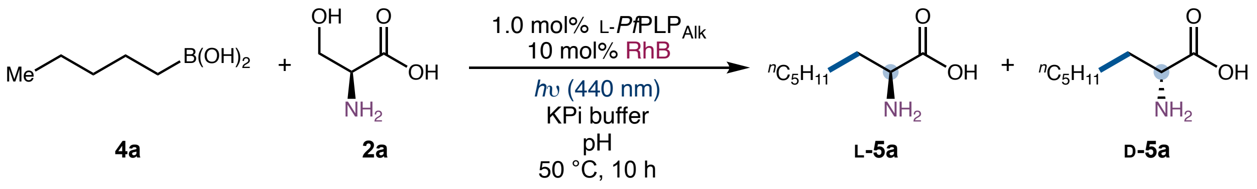

| entry | pH | light intensity | yield of **5a** | e.r. (L-**5a**:D-**5a**) |
|---|---|---|---|---|
| 1 | 8.5 | 5 W | 62% | 88:22 |
| 2 | 8.0 | 5 W | 59% | 91:9 |
| **3** | **7.5** | **5 W** | **59%** | **94:6** |
| 4 | 7.0 | 5 W | 55% | 95:5 |
| 5 | 6.5 | 5 W | 49% | 96:4 |
| 8 | 7.5 | 6 W | 52% | 94:6 |
| **8** | **7.5** | **4 W** | **62%** | **94:6** |
| 9 | 7.5 | 3 W | 56% | 96:4 |
| **10** | **7.5 (1.5 mol% L-*Pf*PLP$_{Alk}$)** | **4 W** | **74%** | **94:6** |

Reaction conditions: **4a** (4.0 mM), **2a** (20.0 mM), 1.0 mol% L-*Pf*PLP$_{Alk}$, 10 mol% RhB, *h*ν (440 nm), 200 mM KPi buffer, DMSO (8% v/v), 50 °C, 10 h.

**Evolutionary trajectory of D-*Pf*PLP$_{Alk}$**

Similarly, (2-phenylethyl)boronic acid and *n*-pentylboronic acid were selected as model substrates for the directed evolution of D-*Pf*PLP$_{Alk}$. Results of molecular docking with AutoDock5 and tunnel analysis with CAVER as well as residues evaluated in site-saturation mutagenesis and screening were described in Figure S8.

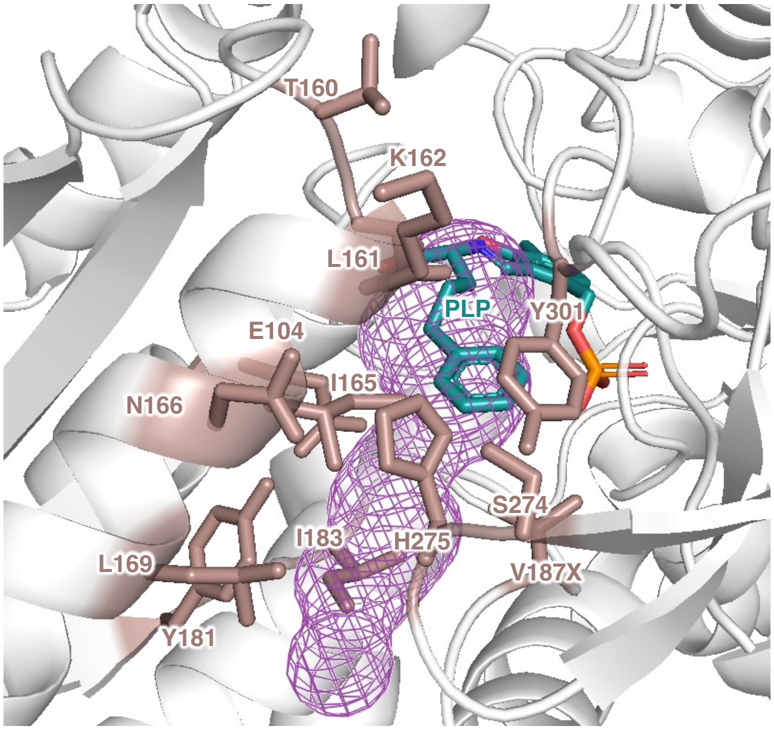

**Figure S8.** Docking model and substrate tunnel for L-*Pf*PLP$^{\beta}$ with D-**3a**.

**Table S9.** Summary of directed evolution of D-*Pf*PLP$_{Alk1}$ (L-*Pf*PLP$^{\beta}$ I165G Y181W H275N)

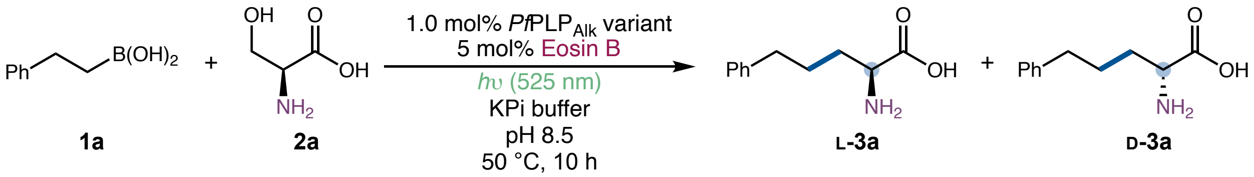

| parent | sites targeted by SSM | selection criterion | beneficial mutation |
|---|---|---|---|
| L-*Pf*PLP$^{\beta}$ I165G (P) | E104X, L161X, N166X Y181X, I183X | activity | Y181W |
| P Y181W | L169X, V187X, H275X | activity | H275N |
| P Y181W H275N | K162X, A164X, N166X, A168X, S274X, Y301X | activity, enantioselectivity | No hits |

**Table S10.** Primers used for site-saturation mutagenesis (22c trick) for directed evolution of D-*Pf*PLP$_{Alk1}$

| DNA template | primer name | primer sequence |
|---|---|---|
| L-*Pf*PLP$^{\beta}$ I165G | E104X_fwd_NDT | GGTAAAACTCGTCTGATCGCT**NDT**ACCGGTGCTGG |
| | E104X_fwd_VHG | GGTAAAACTCGTCTGATCGCT**VHG**ACCGGTGCTGG |
| | E104X_fwd_TGG | GGTAAAACTCGTCTGATCGCT**TGG**ACCGGTGCTGG |
| | E104X_rev | CGATCAGACGAGTTTTACCCATG |
| L-*Pf*PLP$^{\beta}$ I165G | L161X_fwd_NDT | GTTAACTCCGGTTCTCGCACC**NDT**AAAGACGCAG |
| | L161X_fwd_VHG | GTTAACTCCGGTTCTCGCACC**VHG**AAAGACGCAG |
| | L161X_fwd_TGG | GTTAACTCCGGTTCTCGCACC**TGG**AAAGACGCAG |
| | L161X_rev | GGTGCGAGAACCGGAGTTAACTG |
| L-*Pf*PLP$^{\beta}$ I165G | N166X_fwd_NDT | GCACCCTGAAAGACGCAGGT**NDT**GAGGCTCTGC |
| | N166X_fwd_VHG | GCACCCTGAAAGACGCAGGT**VHG**GAGGCTCTGC |
| | N166X_fwd_TGG | GCACCCTGAAAGACGCAGGT**TGG**GAGGCTCTGC |

| | | |
|---|---|---|
| | N166X_rev | CCTGCGTCTTTCAGGGTGCGAG |
| L-*Pf*PLP$^{\beta}$ I165G | Y181X_fwd_NDT | CTACTTTTGAATACACCCAC**NDT**CTAATCGGTTC |
| | Y181X_fwd_VHG | CTACTTTTGAATACACCCAC**VHG**CTAATCGGTTC |
| | Y181X_fwd_TGG | CTACTTTTGAATACACCCAC**TGG**CTAATCGGTTC |
| | Y181X_rev | GTGGGTGTATTCAAAAGTAGCCAC |
| L-*Pf*PLP$^{\beta}$ I165G | I183X_fwd_NDT | CTACTTTTGAATACACCCACTACCTA**NDT**GGTTCCGTGG |
| | I183X_fwd_VHG | CTACTTTTGAATACACCCACTACCTA**VHG**GGTTCCGTGG |
| | I183X_fwd_TGG | CTACTTTTGAATACACCCACTACCTA**TGG**GGTTCCGTGG |
| | I183X_rev | GTGGGTGTATTCAAAAGTAGCCAC |
| L-*Pf*PLP$^{\beta}$ I165G Y181W | L169X_fwd_NDT | GAAAGACGCAGGTAACGAGGCT**NDT**CGTGATTGGGTG |
| | L169X_fwd_VHG | GAAAGACGCAGGTAACGAGGCT**VHG**CGTGATTGGGTG |
| | L169X_fwd_TGG | GAAAGACGCAGGTAACGAGGCT**TGG**CGTGATTGGGTG |
| | L169X_rev | CCTCGTTACCTGCGTCTTTCAGG |
| L-*Pf*PLP$^{\beta}$ I165G Y181W | V187X_fwd_NDT | CTGGCTAATCGGTTCCGTG**NDT**GGTCCACATC |
| | V187X_fwd_VHG | CTGGCTAATCGGTTCCGTG**VHG**GGTCCACATC |
| | V187X_fwd_TGG | CTGGCTAATCGGTTCCGTG**TGG**GGTCCACATC |
| | V187X_rev | CACGGAACCGATTAGCCAGTG |
| L-*Pf*PLP$^{\beta}$ I165G Y181W | H275X_fwd_NDT | GGTCAGGTTGGTGTGTCC**NDT**GGCATGCTGTC |
| | H275X_fwd_VHG | GGTCAGGTTGGTGTGTCC**VHG**GGCATGCTGTC |
| | H275X_fwd_TGG | GGTCAGGTTGGTGTGTCC**TGG**GGCATGCTGTC |
| | H275X_rev | GGACACACCAACCTGACCTG |
| L-*Pf*PLP$^{\beta}$ I165G Y181W H275N | N166X_fwd_NDT | GCACCCTGAAAGACGCAGGT**NDT**GAGGCTCTGC |
| | N166X_fwd_VHG | GCACCCTGAAAGACGCAGGT**VHG**GAGGCTCTGC |
| | N166X_fwd_TGG | GCACCCTGAAAGACGCAGGT**TGG**GAGGCTCTGC |
| | N166X_rev | CCTGCGTCTTTCAGGGTGCGAG |
| L-*Pf*PLP$^{\beta}$ I165G Y181W H275N | K162X_fwd_NDT | GTTAACTCCGGTTCTCGCACCCTG**NDT**GACGCAGGTAAC |
| | K162X_fwd_VHG | GTTAACTCCGGTTCTCGCACCCTG**VHG**GACGCAGGTAAC |
| | K162X_fwd_TGG | GTTAACTCCGGTTCTCGCACCCTG**TGG**GACGCAGGTAAC |
| | K162X_rev | GGTGCGAGAACCGGAGTTAACTG |

| | | |
|---|---|---|
| L-*Pf*PLP$^{\beta}$ I165G Y181W H275N | A164X_fwd_NDT | GTTCTCGCACCCTGAAAGAC**NDT**GGTAACGAGG |
| | A164X_fwd_VHG | GTTCTCGCACCCTGAAAGAC**VHG**GGTAACGAGG |
| | A164X_fwd_TGG | GTTCTCGCACCCTGAAAGAC**TGG**GGTAACGAGG |
| | A164X_rev | GTCTTTCAGGGTGCGAGAACC |
| L-*Pf*PLP$^{\beta}$ I165G Y181W H275N | A168X_fwd_NDT | GAAAGACGCAGGTAACGAG**NDT**CTGCGTGATTG |
| | A168X_fwd_VHG | GAAAGACGCAGGTAACGAG**VHG**CTGCGTGATTG |
| | A168X_fwd_TGG | GAAAGACGCAGGTAACGAG**TGG**CTGCGTGATTG |
| | A168X_rev | CTCGTTACCTGCGTCTTTCAGG |
| L-*Pf*PLP$^{\beta}$ I165G Y181W H275N | S274X_fwd_NDT | GCAGGTCAGGTTGGTGTG**NDT**AATGGCATGC |
| | S274X_fwd_VHG | GCAGGTCAGGTTGGTGTG**VHG**AATGGCATGC |
| | S274X_fwd_TGG | GCAGGTCAGGTTGGTGTG**TGG**AATGGCATGC |
| | S274X_rev | CACACCAACCTGACCTGCGTTC |
| L-*Pf*PLP$^{\beta}$ I165G Y181W H275N | Y301X_fwd_NDT | CTCCATCGCACCAGGTCTGGAT**NDT**CCAGGTGTTG |
| | Y301X_fwd_VHG | CTCCATCGCACCAGGTCTGGAT**VHG**CCAGGTGTTG |
| | Y301X_fwd_TGG | CTCCATCGCACCAGGTCTGGAT**TGG**CCAGGTGTTG |
| | Y301X_rev | CCAGACCTGGTGCGATGGAG |

**Table S11.** Enzyme activity and selectivity summary for photobiocatalytic synthesis of D-**3a**

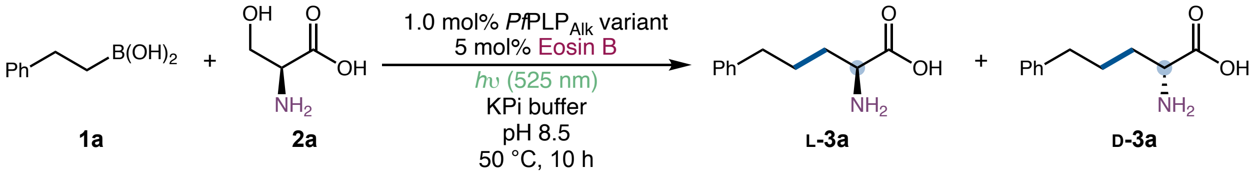

| *Pf*PLP$^{\beta}$ variant | yield of **3a** | e.r. (L-**3a**:D-**3a**) |
|---|---|---|
| L-*Pf*PLP$^{\beta}$ E104G (D-*Pf*PLP$^{\beta}$) | (1.1 ± 0.3)% | 10:90 |
| L-*Pf*PLP$^{\beta}$ E104P (D-*Pf*PLP$^{\beta\ 2.0}$) | (5 ± 1)% | 9:91 |
| L-*Pf*PLP$^{\beta}$ I165G | (27 ± 1)% | 4:96 |
| L-*Pf*PLP$^{\beta}$ I165G Y181W | (62 ± 3)% | 5:95 |

| | | |
|---|---|---|
| **L-*Pf*PLP$^{\beta}$ I165G Y181W H275N (D-*Pf*PLP$_{Alk1}$)** | **(76 ± 1)%** | **6:94** |

Reaction conditions: **1a** (4.0 mM), **2a** (20.0 mM), 1.0 mol% *Pf*PLP$_{Alk}$ variant, 5 mol% Eosin B, *h*ν (525 nm, 3.0 W), 200 mM KPi buffer (pH = 8.5), DMSO (8% v/v), 50 °C, 10 h. All the reactions were performed in triplicates and averaged yields were reported.

**Table S12**. pH effect on photobiocatalytic synthesis of D-**3a**

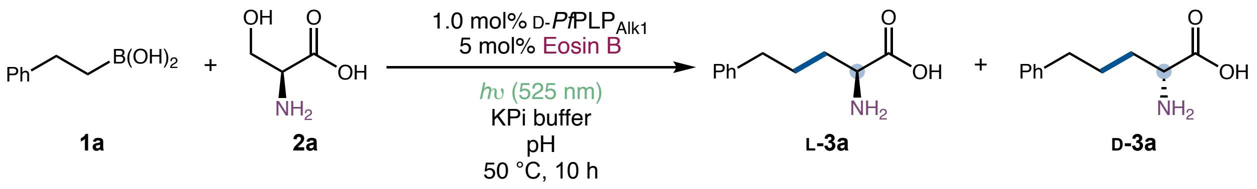

| entry | pH | yield of **3a** | e.r. (L-**3a**:D-**3a**) |
|---|---|---|---|
| 1 | 8.0 | 71% | 7:93 |
| 2 | 8.5 | 75% | 5:95 |
| **3** | **9.0** | **75%** | **4:96** |
| 4 | 9.5 | 66% | 4:96 |

Reaction conditions: **1a** (4.0 mM), **2a** (20.0 mM), 1.0 mol% D-*Pf*PLP$_{Alk1}$, 5 mol% Eosin B, *h*ν (525 nm, 3.0 W), 200 mM KPi buffer (pH), DMSO (8% v/v), 50 °C, 10 h.

**Table S13**. Photoredox catalyst effect on photobiocatalytic synthesis of D-**3a**

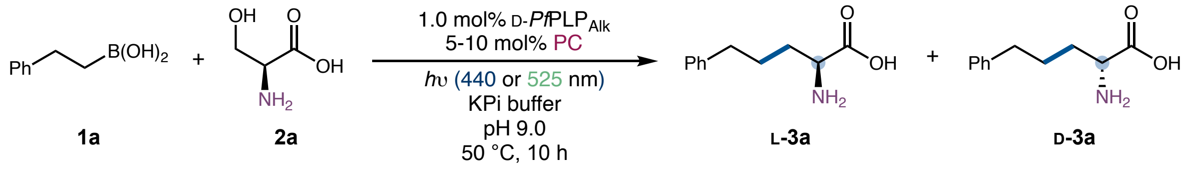

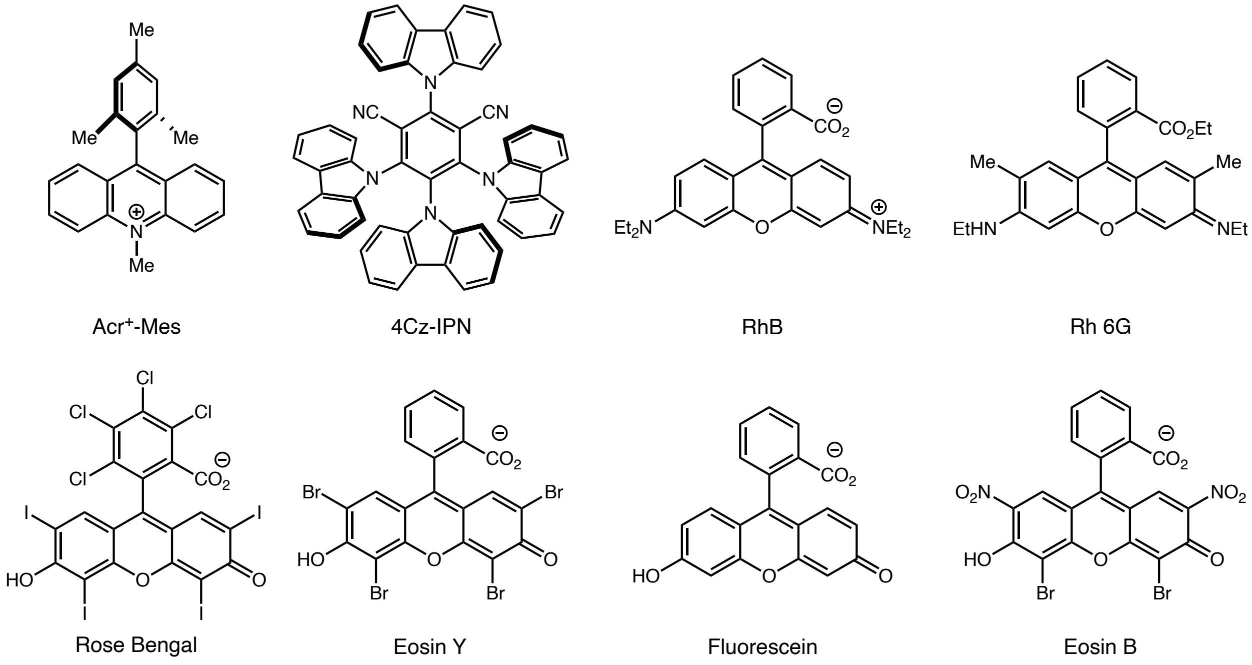

| entry | photocatalyst | light source | yield of **3a** | e.r. (L-**3a**:D-**3a**) |
|---|---|---|---|---|
| 1 | MesAcr (10 mol%) | 440 nm | <1% | 10:90 |
| 2 | 4CzIPN (10 mol%) | 440 nm | trace | - |
| 3 | RhB (10 mol%) | 440 nm | 72% | 4:96 |
| 4 | Rh6G (10 mol%) | 440 nm | 33% | 4:96 |
| 5 | Rose Bengal (10 mol%) | 440 nm | 54% | 8:92 |
| 6 | Eosin Y (10 mol%) | 440 nm | 63% | 5:95 |
| 7 | Eosin B (10 mol%) | 440 nm | 50% | 4:96 |
| 8 | Rose Bengal (5 mol%) | 525 nm | 45% | 6:94 |
| 9 | Eosin Y (5 mol%) | 525 nm | 72% | 5:95 |
| 10 | Fluorescein (5 mol%) | 525 nm | 60% | 4:96 |

| | | | | |
|---|---|---|---|---|
| 11 | RhB (5 mol%) | 525 nm | 8% | 4:96 |
| **12** | **Eosin B (5 mol%)** | **525 nm** | **75%** | **4:96** |
| **13** | **Eosin B (5 mol%) (1.5 mol% D-*Pf*PLP$_{Alk1}$)** | **525 nm** | **85%** | **4:96** |

Reaction conditions: **1a** (4.0 mM), **2a** (20.0 mM), 1.0 mol% D-*Pf*PLP$_{Alk1}$, 5–10 mol% photocatalyst, *h*ν (440 nm, 5.0 W or 525 nm, 3.0 W), 200 mM KPi buffer (pH = 9.0), DMSO (8% v/v), 50 °C, 10 h.

**Table S14.** Summary of directed evolution of D-*Pf*PLP$_{Alk2}$ (L-*Pf*PLP$^{\beta}$ E104P Y181K)

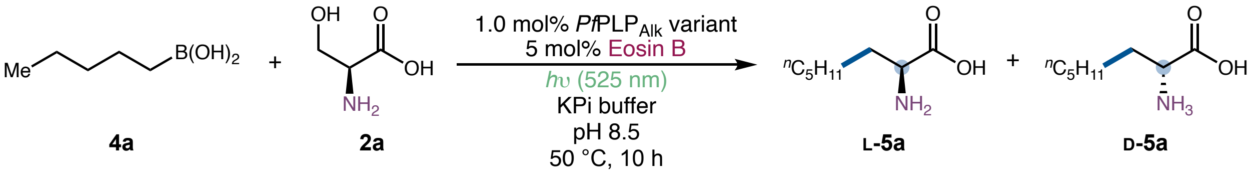

**4a** **2a** **L-5a** **D-5a**

| parent | sites targeted by SSM | selection criterion | beneficial mutation |
|---|---|---|---|
| L-*Pf*PLP$^{\beta}$ E104P (P) (D-*Pf*PLP$^{\beta\,2.0}$) | Y181X, I165X, H275X | activity | Y181K |
| P Y181K | A103X, L161X, T105X, A168X, L169X, I183X, Y301X | activity, enantioselectivity | No hits |

**Table S15.** Primers used for site-saturation mutagenesis (22c trick) for directed evolution of D-*Pf*PLP$_{Alk2}$

| DNA template | primer name | primer sequence |
|---|---|---|
| L-*Pf*PLP$^{\beta}$ E104P | I165X_fwd_NDT | GGTTCTCGCACCCTGAAAGACGCA**NDT**AACGAGGCTC |
| | I165X_fwd_VHG | GGTTCTCGCACCCTGAAAGACGCA**VHG**AACGAGGCTC |
| | I165X_fwd_TGG | GGTTCTCGCACCCTGAAAGACGCA**TGG**AACGAGGCTC |
| | I165X_rev | CGTCTTTCAGGGTGCGAGAACCGGAG |
| L-*Pf*PLP$^{\beta}$ E104P | Y181X_fwd_NDT | CTACTTTTGAATACACCCAC**NDT**CTAATCGGTTC |
| | Y181X_fwd_VHG | CTACTTTTGAATACACCCAC**VHG**CTAATCGGTTC |
| | Y181X_fwd_TGG | CTACTTTTGAATACACCCAC**TGG**CTAATCGGTTC |
| | Y181X_rev | GTGGGTGTATTCAAAAGTAGCCAC |
| L-*Pf*PLP$^{\beta}$ E104P | H275X_fwd_NDT | GGTCAGGTTGGTGTGTCC**NDT**GGCATGCTGTC |
| | H275X_fwd_VHG | GGTCAGGTTGGTGTGTCC**VHG**GGCATGCTGTC |
| | H275X_fwd_TGG | GGTCAGGTTGGTGTGTCC**TGG**GGCATGCTGTC |

| | | |
|---|---|---|
| | H275X_rev | GGACACACCAACCTGACCTG |
| L-*Pf*PLP$^{\beta}$ E104P Y181K | A103X_fwd_NDT | CATGGGTAAAACTCGTCTGATC**NDT**CCGACCGGTG |
| | A103X_fwd_VHG | CATGGGTAAAACTCGTCTGATC**VHG**CCGACCGGTG |
| | A103X_fwd_TGG | CATGGGTAAAACTCGTCTGATC**TGG**CCGACCGGTG |
| | A103X_rev | GATCAGACGAGTTTTACCCATGAG |
| L-*Pf*PLP$^{\beta}$ E104P Y181K | T105X_fwd_NDT | GTAAAACTCGTCTGATCGCTCCG**NDT**GGTGCTGGTC |
| | T105X_fwd_VHG | GTAAAACTCGTCTGATCGCTCCG**VHG**GGTGCTGGTC |
| | T105X_fwd_TGG | GTAAAACTCGTCTGATCGCTCCG**TGG**GGTGCTGGTC |
| | T105X_rev | GGAGCGATCAGACGAGTTTTAC |
| L-*Pf*PLP$^{\beta}$ E104P Y181K | L161X_fwd_NDT | GTTAACTCCGGTTCTCGCACC**NDT**AAAGACGCAATC |
| | L161X_fwd_VHG | GTTAACTCCGGTTCTCGCACC**VHG**AAAGACGCAATC |
| | L161X_fwd_TGG | GTTAACTCCGGTTCTCGCACC**TGG**AAAGACGCAATC |
| | L161X_rev | GGTGCGAGAACCGGAGTTAACTG |
| L-*Pf*PLP$^{\beta}$ E104P Y181K | A168X_fwd_NDT | GAAAGACGCAATCAACGAG**NDT**CTGCGTGATTG |
| | A168X_fwd_VHG | GAAAGACGCAATCAACGAG**VHG**CTGCGTGATTG |
| | A168X_fwd_TGG | GAAAGACGCAATCAACGAG**TGG**CTGCGTGATTG |
| | A168X_rev | CTCGTTGATTGCGTCTTTCAGG |
| L-*Pf*PLP$^{\beta}$ E104P Y181K | L169X_fwd_NDT | GAAAGACGCAATCAACGAGGCT**NDT**CGTGATTGGGTG |
| | L169X_fwd_VHG | GAAAGACGCAATCAACGAGGCT**VHG**CGTGATTGGGTG |
| | L169X_fwd_TGG | GAAAGACGCAATCAACGAGGCT**TGG**CGTGATTGGGTG |
| | L169X_rev | CCTCGTTGATTGCGTCTTTCAGG |
| L-*Pf*PLP$^{\beta}$ E104P Y181K | N166X_fwd_NDT | CACCCTGAAAGACGCAATC**NDT**GAGGCTCTGCGTG |
| | N166X_fwd_VHG | CACCCTGAAAGACGCAATC**VHG**GAGGCTCTGCGTG |
| | N166X_fwd_TGG | CACCCTGAAAGACGCAATC**TGG**GAGGCTCTGCGTG |
| | N166X_rev | GATTGCGTCTTTCAGGGTGCGAG |
| L-*Pf*PLP$^{\beta}$ E104P Y181K | I183X_fwd_NDT | CTTTTGAATACACCCACAAGCTA**NDT**GGTTCCGTGG |
| | I183X_fwd_VHG | CTTTTGAATACACCCACAAGCTA**VHG**GGTTCCGTGG |
| | I183X_fwd_TGG | CTTTTGAATACACCCACAAGCTA**TGG**GGTTCCGTGG |
| | K162X_rev | GCTTGTGGGTGTATTCAAAAGTAGC |

| | | |
|---|---|---|
| L-*Pf*PLP$^{\beta}$ E104P Y181K | Y301X_fwd_NDT | CTCCATCGCACCAGGTCTGGAT**NDT**CCAGGTGTTG |
| | Y301X_fwd_VHG | CTCCATCGCACCAGGTCTGGAT**VHG**CCAGGTGTTG |
| | Y301X_fwd_TGG | CTCCATCGCACCAGGTCTGGAT**TGG**CCAGGTGTTG |
| | Y301X_rev | CCAGACCTGGTGCGATGGAG |

**Table S16.** Enzyme activity and selectivity summary for photobiocatalytic synthesis of D-**5a**

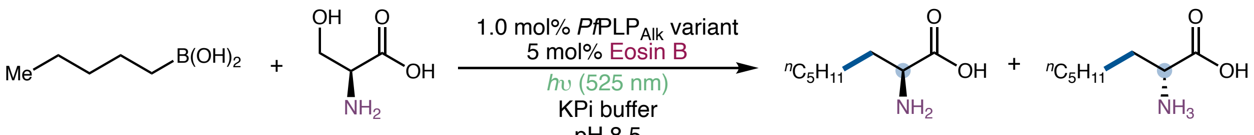

**4a** **2a** **L-5a** **D-5a**

| *Pf*PLP$^{\beta}$ variant | Yield of **5a** | e.r. (L-**5a**:D-**5a**) |
|---|---|---|
| L-*Pf*PLP$^{\beta}$ I165G | (15 ± 0.4)% | 22:78 |
| L-*Pf*PLP$^{\beta}$ I165G Y181W H275N (D-*Pf*PLP$_{\text{Alk1}}$) | (55 ± 1)% | 34:66 |
| L-*Pf*PLP$^{\beta}$ E104P (D-*Pf*PLP$^{\beta\ 2.0}$) | (16 ± 1)% | 5:95 |
| **L-*Pf*PLP$^{\beta}$ E104P Y181K (D-*Pf*PLP$_{\text{Alk2}}$)** | **(47 ± 4)%** | **4:96** |

Reaction conditions: **4a** (4.0 mM), **2a** (20.0 mM), 1.0 mol% *Pf*PLP$_{\text{Alk}}$ variant, 5 mol% Eosin B, *h*ν (525 nm, 3.0 W), 200 mM KPi buffer (pH = 8.5), DMSO (8% v/v), 50 °C, 10 h. All the reactions were performed in triplicates and averaged yields were reported.

**Table S17.** Selected reaction conditions optimization for photobiocatalytic synthesis of D-**5a**

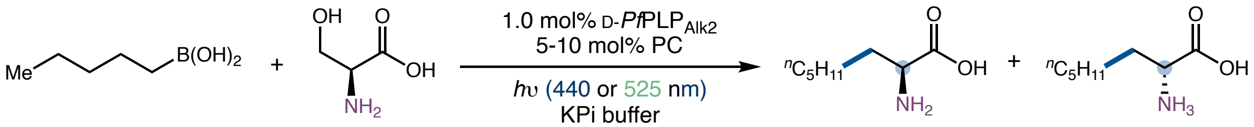

**4a** **2a** **L-5a** **D-5a**

| entry | pH | photocatalytic conditions | yield of **5a** | e.r. (L-**5a**:D-**5a**) |
|---|---|---|---|---|
| 1 | 8.5 | 5% Eosin B, 525 nm, 3 W | 47% | 4:96 |
| 2 | 8.5 | 10% RhB, 440 nm, 5 W | 58% | 4:96 |
| **3** | **8.5** | **10% RhB, 440 nm, 4 W** | **64%** | **4:96** |
| 4 | 8.5 | 10% RhB, 440 nm, 3 W | 61% | 4:96 |
| 5 | 8.0 | 10% RhB, 440 nm, 4 W | 62% | 5:95 |
| **6** | **9.0** | **10% RhB, 440 nm, 4 W** | **64%** | **3:97** |
| **7** | **9.0 (1.5 mol% D-*Pf*PLP$_{\mathbf{Alk2}}$)** | **10% RhB, 440 nm, 4 W** | **75%** | **3:97** |

Reaction conditions using Eosin B: **4a** (4.0 mM), **2a** (20.0 mM), 1.0 mol% D-*Pf*PLP$_{Alk2}$, 5 mol% Eosin B, *h*ν (525 nm, 3.0 W), 200 mM KPi buffer (pH), DMSO (8% v/v), 50 °C, 10 h. Reaction conditions using RhB: **4a** (4.0 mM), **2a** (20.0 mM), 1.0 mol% D-*Pf*PLP $_{Alk2}$, 10 mol% RhB, *h*ν (440 nm, 3–5 W), 200 mM KPi buffer (pH), DMSO (8% v/v), 50 °C, 10 h.

#### III. DNA and protein sequences

DNA and amino acids sequence of L-***Pf*PLP**$^{\beta}$:

##### DNA sequence

ATGTGGTTCGGTGAATTTGGTGGTCAGTACGTGCCAGAAACGCTGGTTGGACCCCTGAAAGAGC
TGGAAAAAGCTTACAAACGTTTCAAAGATGACGAAGAATTCAATCGTCAGCTGAATTACTACCT
GAAAACCTGGGCAGGTCGTCCAACCCCACTGTACTACGCAAAACGCCTGACTGAAAAAATCGGT
GGTGCTAAAGTCTACCTGAAACGTGAAGACCTGGTTCACGGTGGTGCACACAAGACCAACAACG
CCATCGGTCAGGCACTGCTGGCAAAGCTCATGGGTAAAACTCGTCTGATCGCTGAGACCGGTGC
TGGTCAGCACGGCGTAGCGACTGCAATGGCTGGTGCACTGCTGGGCATGAAAGTGGACATTTAC
ATGGGTGCTGAGGACGTAGAACGTCAGAAAATGAACGTATTCCGTATGAAGCTGCTGGGTGCAA
ACGTAATTCCAGTTAACTCCGGTTCTCGCACCCTGAAAGACGCAATCAACGAGGCTCTGCGTGA
TTGGGTGGCTACTTTTGAATACACCCACTACCTAATCGGTTCCGTGGTCGGTCCACATCCGTAT
CCGACCATCGTTCGTGATTTTCAGTCTGTTATCGGTCGTGAGGCTAAAGCGCAGATCCTGGAGG
CTGAAGGTCAGCTGCCAGATGTAATCGTTGCTTGTGTTGGTGGTGGCTCTAACGCGATGGGTAT
CTTTTACCCGTTCGTGAACGACAAAAAAGTTAAGCTGGTTGGCGTTGAGGCTGGTGGTAAAGGC
CTGGAATCTGGTAAGCATTCCGCTAGCCTGAACGCAGGTCAGGTTGGTGTGTCCCATGGCATGC
TGTCCTACTTTCTGCAGGACGAAGAAGGTCAGATCAAACCAAGCCACTCCATCGCACCAGGTCT
GGATTATCCAGGTGTTGGTCCAGAACACGCTTACCTGAAAAAAATTCAGCGTGCTGAATACGTG
GCTGTAACCGATGAAGAAGCACTGAAAGCGTTCCATGAACTGAGCCGTACCGAAGGTATCATCC
CAGCTCTGGAATCTGCGCATGCTGTGGCTTACGCTATGAAACTGGCTAAGGAAATGTCTCGTGA
TGAGATCATCATCGTAAACCTGTCTGGTCGTGGTGACAAAGACCTGGATATTGTCCTGAAAGCG
TCTGGCAACGTGCTCGAGCACCACCACCACCACCACTGA

##### Protein sequence

MWFGEFGGQYVPETLVGPLKELEKAYKRFKDDEEFNRQLNYYLKTWAGRPTPLYYAKRLTEKIG
GAKVYLKREDLVHGGAHKTNNAIGQALLAKLMGKTRLIAETGAGQHGVATAMAGALLGMKVDIY
MGAEDVERQKMNVFRMKLLGANVIPVNSGSRTLKDAINEALRDWVATFEYTHYLIGSVVGPHPY
PTIVRDFQSVIGREAKAQILEAEGQLPDVIVACVGGGSNAMGIFYPFVNDKKVKLVGVEAGGKG

LESGKHSASLNAGQVGVSHGMLSYFLQDEEGQIKPSHSIAPGLDYPGVGPEHAYLKKIQRAEYV
AVTDEEALKAFHELSRTEGIIPALESAHAVAYAMKLAKEMSRDEIIIVNLSGRGDKDLDIVLKA
SGNVLEHHHHHH*

DNA and amino acids sequence of **L-*Pf*PLP$_{\mathbf{Alk}}$**:

L-*Pf*PLP$^{\beta}$ Y301H I165T L161S Y181H H275D N166H T160K

**DNA sequence**

ATGTGGTTCGGTGAATTTGGTGGTCAGTACGTGCCAGAAACGCTGGTTGGACCCCTGAAAGAGC
TGGAAAAAGCTTACAAACGTTTCAAAGATGACGAAGAATTCAATCGTCAGCTGAATTACTACCT
GAAAACCTGGGCAGGTCGTCCAACCCCACTGTACTACGCAAAACGCCTGACTGAAAAAATCGGT
GGTGCTAAAGTCTACCTGAAACGTGAAGACCTGGTTCACGGTGGTGCACACAAGACCAACAACG
CCATCGGTCAGGCACTGCTGGCAAAGCTCATGGGTAAAACTCGTCTGATCGCTGAGACCGGTGC
TGGTCAGCACGGCGTAGCGACTGCAATGGCTGGTGCACTGCTGGGCATGAAAGTGGACATTTAC
ATGGGTGCTGAGGACGTAGAACGTCAGAAAATGAACGTATTCCGTATGAAGCTGCTGGGTGCAA
ACGTAATTCCAGTTAACTCCGGTTCTCGC**AAGAGT**AAAGACGCA**ACGCAT**GAGGCTCTGCGTGA
TTGGGTGGCTACTTTTGAATACACCCAC**CAT**CTAATCGGTTCCGTGGTCGGTCCACATCCGTAT
CCGACCATCGTTCGTGATTTTCAGTCTGTTATCGGTCGTGAGGCTAAAGCGCAGATCCTGGAGG
CTGAAGGTCAGCTGCCAGATGTAATCGTTGCTTGTGTTGGTGGTGGCTCTAACGCGATGGGTAT
CTTTTACCCGTTCGTGAACGACAAAAAAGTTAAGCTGGTTGGCGTTGAGGCTGGTGGTAAAGGC
CTGGAATCTGGTAAGCATTCCGCTAGCCTGAACGCAGGTCAGGTTGGTGTGTCC**GAT**GGCATGC
TGTCCTACTTTCTGCAGGACGAAGAAGGTCAGATCAAACCAAGCCACTCCATCGCACCAGGTCT
GGAT**CAT**CCAGGTGTTGGTCCAGAACACGCTTACCTGAAAAAAATTCAGCGTGCTGAATACGTG
GCTGTAACCGATGAAGAAGCACTGAAAGCGTTCCATGAACTGAGCCGTACCGAAGGTATCATCC
CAGCTCTGGAATCTGCGCATGCTGTGGCTTACGCTATGAAACTGGCTAAGGAAATGTCTCGTGA
TGAGATCATCATCGTAAACCTGTCTGGTCGTGGTGACAAAGACCTGGATATTGTCCTGAAAGCG
TCTGGCAACGTGCTCGAGCACCACCACCACCACCACTGA

**Protein sequence**

MWFGEFGGQYVPETLVGPLKELEKAYKRFKDDEEFNRQLNYYLKTWAGRPTPLYYAKRLTEKIG

GAKVYLKREDLVHGGAHKTNNAIGQALLAKLMGKTRLIAETGAGQHGVATAMAGALLGMKVDIY

MGAEDVERQKMNVFRMKLLGANVIPVNSGSR**KS**KDA**TH**EALRDWVATFEYTH**H**LIGSVVGPHPY

PTIVRDFQSVIGREAKAQILEAEGQLPDVIVACVGGGSNAMGIFYPFVNDKKVKLVGVEAGGKG

LESGKHSASLNAGQVGVS**D**GMLSYFLQDEEGQIKPSHSIAPGLD**H**PGVGPEHAYLKKIQRAEYV

AVTDEEALKAFHELSRTEGIIPALESAHAVAYAMKLAKEMSRDEIIIVNLSGRGDKDLDIVLKA

SGNVLEHHHHHH*

DNA and amino acids sequence of **D-*Pf*PLP$_{Alk1}$**:

L-*Pf*PLP$^{\beta}$ I165G Y181W H275N

**DNA sequence**

ATGTGGTTCGGTGAATTTGGTGGTCAGTACGTGCCAGAAACGCTGGTTGGACCCCTGAAAGAGC

TGGAAAAAGCTTACAAACGTTTCAAAGATGACGAAGAATTCAATCGTCAGCTGAATTACTACCT

GAAAACCTGGGCAGGTCGTCCAACCCCACTGTACTACGCAAAACGCCTGACTGAAAAAATCGGT

GGTGCTAAAGTCTACCTGAAACGTGAAGACCTGGTTCACGGTGGTGCACACAAGACCAACAACG

CCATCGGTCAGGCACTGCTGGCAAAGCTCATGGGTAAAACTCGTCTGATCGCTGAGACCGGTGC

TGGTCAGCACGGCGTAGCGACTGCAATGGCTGGTGCACTGCTGGGCATGAAAGTGGACATTTAC

ATGGGTGCTGAGGACGTAGAACGTCAGAAAATGAACGTATTCCGTATGAAGCTGCTGGGTGCAA

ACGTAATTCCAGTTAACTCCGGTTCTCGCACCCTGAAAGACGCA**GGT**AACGAGGCTCTGCGTGA

TTGGGTGGCTACTTTTGAATACACCCAC**TGG**CTAATCGGTTCCGTGGTCGGTCCACATCCGTAT

CCGACCATCGTTCGTGATTTTCAGTCTGTTATCGGTCGTGAGGCTAAAGCGCAGATCCTGGAGG

CTGAAGGTCAGCTGCCAGATGTAATCGTTGCTTGTGTTGGTGGTGGCTCTAACGCGATGGGTAT

CTTTTACCCGTTCGTGAACGACAAAAAAGTTAAGCTGGTTGGCGTTGAGGCTGGTGGTAAAGGC

CTGGAATCTGGTAAGCATTCCGCTAGCCTGAACGCAGGTCAGGTTGGTGTGTCC**AAT**GGCATGC

TGTCCTACTTTCTGCAGGACGAAGAAGGTCAGATCAAACCAAGCCACTCCATCGCACCAGGTCT

GGATTATCCAGGTGTTGGTCCAGAACACGCTTACCTGAAAAAAATTCAGCGTGCTGAATACGTG

GCTGTAACCGATGAAGAAGCACTGAAAGCGTTCCATGAACTGAGCCGTACCGAAGGTATCATCC

CAGCTCTGGAATCTGCGCATGCTGTGGCTTACGCTATGAAACTGGCTAAGGAAATGTCTCGTGA

TGAGATCATCATCGTAAACCTGTCTGGTCGTGGTGACAAAGACCTGGATATTGTCCTGAAAGCG
TCTGGCAACGTGCTCGAGCACCACCACCACCACCACTGA

**Protein sequence**

MWFGEFGGQYVPETLVGPLKELEKAYKRFKDDEEFNRQLNYYLKTWAGRPTPLYYAKRLTEKIG
GAKVYLKREDLVHGGAHKTNNAIGQALLAKLMGKTRLIAETGAGQHGVATAMAGALLGMKVDIY
MGAEDVERQKMNVFRMKLLGANVIPVNSGSRTLKDA**G**NEALRDWVATFEYTH**W**LIGSVVGPHPY
PTIVRDFQSVIGREAKAQILEAEGQLPDVIVACVGGGSNAMGIFYPFVNDKKVKLVGVEAGGKG
LESGKHSASLNAGQVGVS**N**GMLSYFLQDEEGQIKPSHSIAPGLDYPGVGPEHAYLKKIQRAEYV
AVTDEEALKAFHELSRTEGIIPALESAHAVAYAMKLAKEMSRDEIIIVNLSGRGDKDLDIVLKA
SGNVLEHHHHHH*

DNA and amino acids sequence of **D-*Pf*PLP$_{Alk2}$**:

L-*Pf*PLP$^{\beta}$ E104P Y181K

**DNA sequence**

ATGTGGTTCGGTGAATTTGGTGGTCAGTACGTGCCAGAAACGCTGGTTGGACCCCTGAAAGAGC
TGGAAAAAGCTTACAAACGTTTCAAAGATGACGAAGAATTCAATCGTCAGCTGAATTACTACCT
GAAAACCTGGGCAGGTCGTCCAACCCCACTGTACTACGCAAAACGCCTGACTGAAAAAATCGGT
GGTGCTAAAGTCTACCTGAAACGTGAAGACCTGGTTCACGGTGGTGCACACAAGACCAACAACG
CCATCGGTCAGGCACTGCTGGCAAAGCTCATGGGTAAAACTCGTCTGATCGCT**CCG**ACCGGTGC
TGGTCAGCACGGCGTAGCGACTGCAATGGCTGGTGCACTGCTGGGCATGAAAGTGGACATTTAC
ATGGGTGCTGAGGACGTAGAACGTCAGAAAATGAACGTATTCCGTATGAAGCTGCTGGGTGCAA
ACGTAATTCCAGTTAACTCCGGTTCTCGCACCCTGAAAGACGCAATCAACGAGGCTCTGCGTGA
TTGGGTGGCTACTTTTGAATACACCCAC**AAG**CTAATCGGTTCCGTGGTCGGTCCACATCCGTAT
CCGACCATCGTTCGTGATTTTCAGTCTGTTATCGGTCGTGAGGCTAAAGCGCAGATCCTGGAGG
CTGAAGGTCAGCTGCCAGATGTAATCGTTGCTTGTGTTGGTGGTGGCTCTAACGCGATGGGTAT
CTTTTACCCGTTCGTGAACGACAAAAAAGTTAAGCTGGTTGGCGTTGAGGCTGGTGGTAAAGGC
CTGGAATCTGGTAAGCATTCCGCTAGCCTGAACGCAGGTCAGGTTGGTGTGTCCCATGGCATGC

TGTCCTACTTTCTGCAGGACGAAGAAGGTCAGATCAAACCAAGCCACTCCATCGCACCAGGTCT
GGATTATCCAGGTGTTGGTCCAGAACACGCTTACCTGAAAAAAATTCAGCGTGCTGAATACGTG
GCTGTAACCGATGAAGAAGCACTGAAAGCGTTCCATGAACTGAGCCGTACCGAAGGTATCATCC
CAGCTCTGGAATCTGCGCATGCTGTGGCTTACGCTATGAAACTGGCTAAGGAAATGTCTCGTGA
TGAGATCATCATCGTAAACCTGTCTGGTCGTGGTGACAAAGACCTGGATATTGTCCTGAAAGCG
TCTGGCAACGTGCTCGAGCACCACCACCACCACCACTGA

**Protein sequence**

MWFGEFGGQYVPETLVGPLKELEKAYKRFKDDEEFNRQLNYYLKTWAGRPTPLYYAKRLTEKIG
GAKVYLKREDLVHGGAHKTNNAIGQALLAKLMGKTRLIA**P**TGAGQHGVATAMAGALLGMKVDIY
MGAEDVERQKMNVFRMKLLGANVIPVNSGSRTLKDAINEALRDWVATFEYTH**K**LIGSVVGPHPY
PTIVRDFQSVIGREAKAQILEAEGQLPDVIVACVGGGSNAMGIFYPFVNDKKVKLVGVEAGGKG
LESGKHSASLNAGQVGVSHGMLSYFLQDEEGQIKPSHSIAPGLDYPGVGPEHAYLKKIQRAEYV
AVTDEEALKAFHELSRTEGIIPALESAHAVAYAMKLAKEMSRDEIIIVNLSGRGDKDLDIVLKA
SGNVLEHHHHHH*

### IV. Mechanistic studies

#### 1. Radical clock experiments

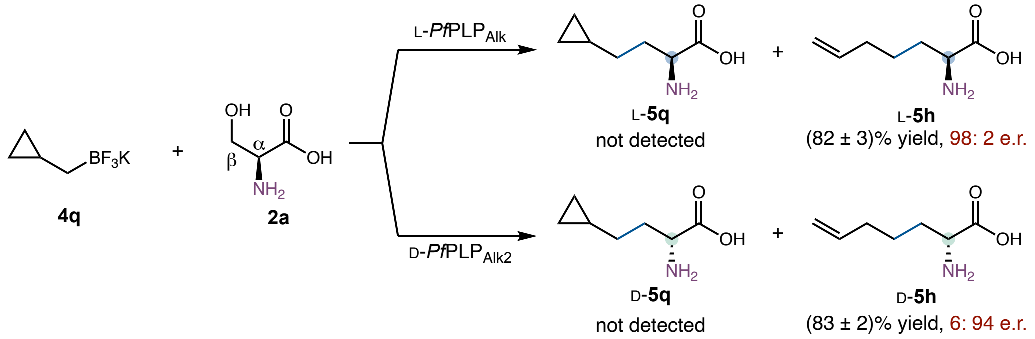

Radical clock substrate **4q** was subjected to the standard conditions. Following derivatization and comparison with authentic standards, only the ring-opened product **5h** was detected; no cyclized product **5q** was observed. The LC-MS trace is shown in Figure S9.

a) LC trace for Marfey's derivatization of authentic products

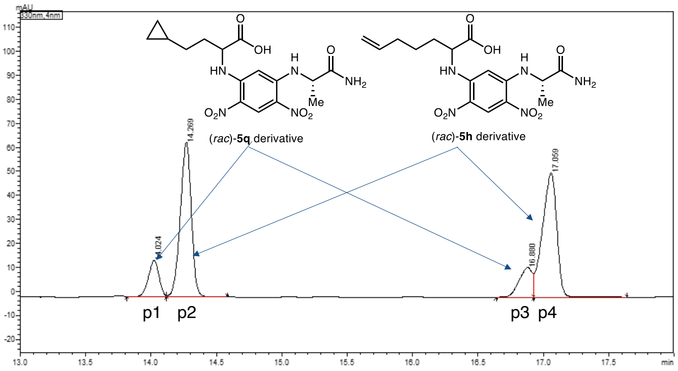

Peak Table | Compound | Group | Calibration Curve

| Peak# | Peak Start | Peak End | Ret. Time | Area | Area% |
|---|---|---|---|---|---|
| 1 | 13.816 | 14.120 | 14.024 | 84905 | 8.976 |
| 2 | 14.120 | 14.584 | 14.269 | 376843 | 39.841 |
| 3 | 16.640 | 16.928 | 16.880 | 88218 | 9.327 |
| 4 | 16.928 | 17.640 | 17.059 | 395912 | 41.857 |
| Total | | | | 945878 | 100.000 |

P1 and P3 are the enantiomers of the (*rac*)-**5r** derivative;
P2 and P4 are the enantiomers of the (*rac*)-**5m** derivative;

b) LC trace of Marfey's derivatization for reaction using **L-*Pf*PLP$_{alk}$**

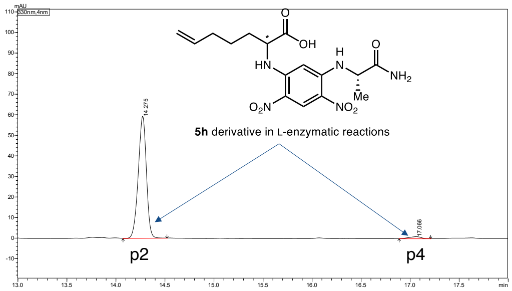

Peak Table | Compound | Group | Calibration Curve

| Peak# | Peak Start | Peak End | Ret. Time | Area | Area% |
|---|---|---|---|---|---|
| 1 | 14.072 | 14.520 | 14.275 | 341051 | 98.002 |
| 2 | 16.888 | 17.208 | 17.066 | 6954 | 1.998 |
| Total | | | | 348005 | 100.000 |

c) LC trace of Marfey's derivatization for reaction using **D-*Pf*PLP$^{\beta\ 3.0b}$**

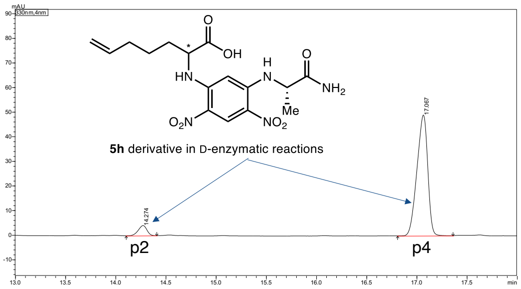

Peak Table | Compound | Group | Calibration Curve

| Peak# | Peak Start | Peak End | Ret. Time | Area | Area% |
|---|---|---|---|---|---|
| 1 | 14.104 | 14.408 | 14.274 | 24659 | 6.321 |
| 2 | 16.808 | 17.360 | 17.067 | 365442 | 93.679 |
| Total | | | | 390100 | 100.000 |

**Figure S9.** LC-MS analysis of Marfey derivatization of amino acid product derived from **4q**

HPLC conditions for Marfey's analysis: Poroshell 120 EC-C18 column (4.6 × 150 mm, 2.7 µm) at a flow rate of 1.0 mL/min. The elution program was as follows: hold at 5% $CH_3CN$ (0.1% formic acid) in $H_2O$ (0.1% formic acid) for 1 min; linear gradient to 30% $CH_3CN$ over 4 min; further gradient to 55% $CH_3CN$ over 17 min; ramp to 95% $CH_3CN$ over 3 min; hold at 95% $CH_3CN$ for 1 min. The total runtime was 27 min.

**2-Aminohept-6-enoic acid (5h)**

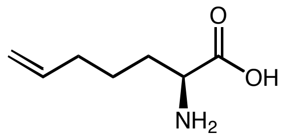

**L-5h** was prepared according to General Procedure using L-*Pf*PLP$_{Alk}$ as the enzyme and **4q** as the substrate. The reactions were carried out in triplicatess. **L-5h** was derivatized using (*S*)-Marfey's reagent and analyzed by LC-MS.

**Yield:** run 1: 83.2%; run 2: 78.8%; run 3: 84.9%; average yield: (82 ± 3)%.

**Enantioselectivity:** run 1: 98:2 e.r.; run 2: 98:2 e.r.; run 3: 98:2 e.r.; average e.r.: 98:2.

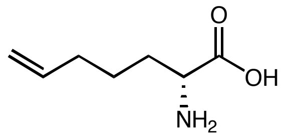

**D-5h** was prepared according to General Procedure using D-*Pf*PLP$_{Alk2}$ as the enzyme and **4q** as the substrate. The reactions were carried out in triplicatess. **D-5h** was derivatized using (*S*)-Marfey's reagent and analyzed by LC-MS.

**Yield:** run 1: 85.0%; run 2: 81.9%; run 3: 80.8%; average yield: (83 ± 2)%.

**Enantioselectivity:** run 1: 6:94 e.r.; run 2: 6:94 e.r.; run 3: 6:94 e.r.; average e.r.: 6:94.

#### 2. Radical cyclization

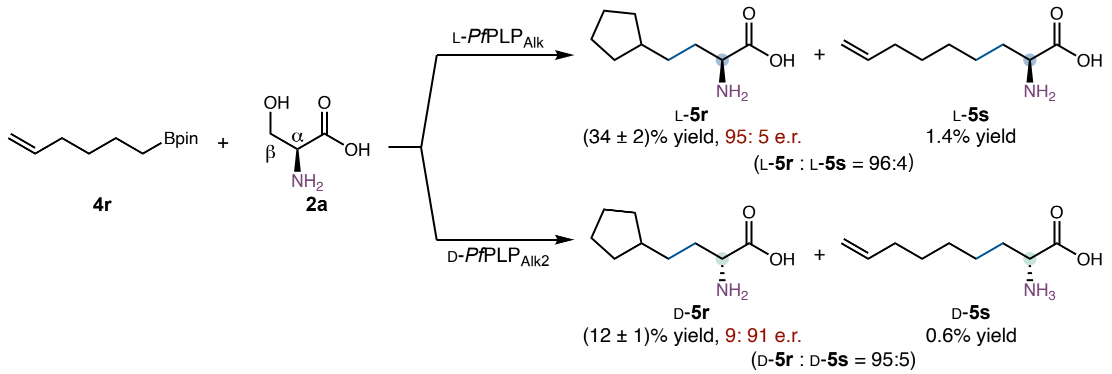

Following the General Procedure, the reaction was carried out using **4r** as the substrate. After dinitrofluorobenzene (DNFB) derivatization and comparison with authentic standards, both the cyclized product **5r** and the non-cyclized product **5s** were identified (Figure S10B).

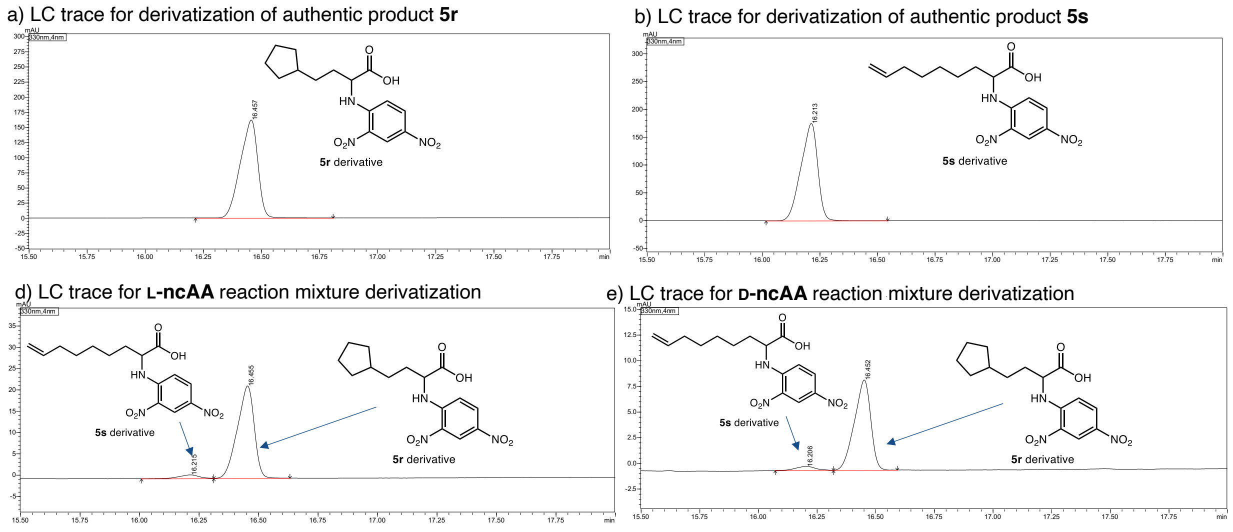

| Peak# | Peak Start | Peak End | Ret. Time | Area | Area% |
|---|---|---|---|---|---|
| 1 | 16.008 | 16.312 | 16.215 | 4857 | 4.346 |
| 2 | 16.312 | 16.632 | 16.455 | 106897 | 95.654 |
| Total | | | | 111755 | 100.000 |

| Peak# | Peak Start | Peak End | Ret. Time | Area | Area% |
|---|---|---|---|---|---|
| 1 | 16.072 | 16.320 | 16.206 | 2182 | 4.876 |
| 2 | 16.320 | 16.592 | 16.452 | 42555 | 95.124 |
| Total | | | | 44737 | 100.000 |

**Figure S10.** HPLC analysis of Marfey derivatization products of amino acid derived from **4r**.

HPLC conditions for DNFB derivation analysis: Poroshell 120 EC-C18 column (4.6 × 150 mm, 2.7 μm) at a flow rate of 1.0 mL/min. The elution program was as follows: hold at 5% $CH_3CN$ (0.1% formic acid) in $H_2O$ (0.1% formic acid) for 1 min; linear gradient to 30% $CH_3CN$ over 4 min; further gradient to 70% $CH_3CN$ over 15 min; ramp to 95% $CH_3CN$ over 1 min; hold at 95% $CH_3CN$ for 1 min. The total runtime was 23 min.

**2-Amino-4-cyclopentylbutanoic acid (5r)**

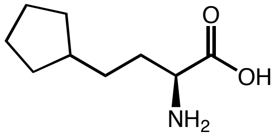

**L-5r** was prepared according to General Procedure using L-*Pf*PLP$_{Alk}$ as the enzyme and **4r** as the substrate. The reactions were carried out in triplicates. **L-5r** was derivatized using (*S*)-Marfey's reagent and analyzed by LC-MS.

**Yield:** run 1: 36.1%; run 2: 33.7%; run 3: 31.6%; average yield: (34 ± 2)%.

**Enantioselectivity:** run 1: 95:5 e.r.; run 2: 95:5 e.r.; run 3: 95:5 e.r.; average e.r.: 95:5.

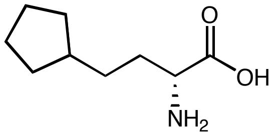

**D-5r** was prepared according to General Procedure using D-*Pf*PLP$_{Alk2}$ as the enzyme and **4r** as the substrate. The reactions were carried out in triplicates. **D-5r** was derivatized using (*S*)-Marfey's reagent and analyzed by LC-MS.

**Yield:** run 1: 12.4%; run 2: 10.6%; run 3: 13.9%; average yield: (12 ± 2)%.

**Enantioselectivity:** run 1: 9:91 e.r.; run 2: 9:91 e.r.; run 3: 9:91 e.r.; average e.r.: 9:91.

#### 3. Radical aryl migration

Following the general procedure, the reaction was conducted using (*R*)-**1r** as the substrate. After Marfey's derivatization and analysis using authentic standards, both the desired product **3r** and the aryl migration product **3s** were observed (Figure S11).

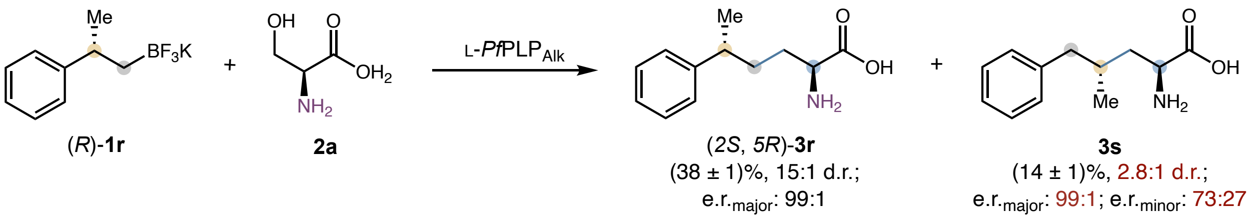

**HPLC Analysis of Marfey's derivatization products and stereochemical assignment**: HPLC analysis of Marfey derivatization products of the authentic standards (*rac*)-**3r** and (*rac*)-**3s** is shown in Figures S11. Peaks **p1** and **p3** represent the enantiomers of the major diastereomer of **3r**, while p2 and p4 corresponds to the enantiomers of the minor diastereomer of **3r** (Figure S11a). Similarly, p5 and p7 correspond to the enantiomers of the major diastereomer of **3s**, while p6 and p8 are the enantiomers of the minor diastereomer of **3s** (Figure S11b). The HPLC trace of Marfey derivatization mixture of (*rac*)-**3r** and (*rac*)-**3s** is shown in Figure S11c. As can be seen from Figure S11c, stereoisomer p1 of **3r** co-eluded with stereoisomer p5 of **3s**, while stereoisomer p3 of **3r** co-eluded with stereoisomer p7 of **3s**, complicating the analysis of stereoisomeric ratios. Finally, the HPLC trace of Marfey derivatization products of amino acids derived from (*R*)-**1r** is provided in Figure S11d.

a) LC trace for derivatization of (*rac*)-**3r**

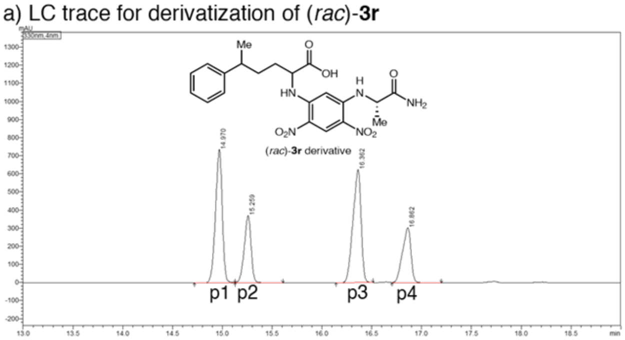

| Peak# | Peak Start | Peak End | Ret. Time | Area | Area% |
|---|---|---|---|---|---|
| 1 | 14.720 | 15.128 | 14.970 | 3470456 | 32.881 |
| 2 | 15.128 | 15.608 | 15.259 | 1761411 | 16.689 |
| 3 | 16.144 | 16.512 | 16.362 | 3544672 | 33.584 |
| 4 | 16.704 | 17.200 | 16.862 | 1777975 | 16.846 |
| Total | | | | 10554513 | 100.000 |

b) LC trace for derivatization of (*rac*)-**3s**

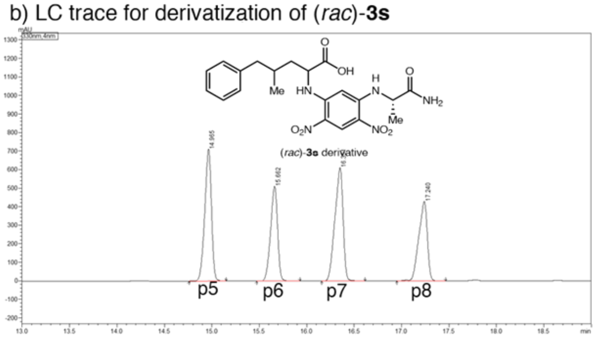

| Peak# | Peak Start | Peak End | Ret. Time | Area | Area% |
|---|---|---|---|---|---|
| 1 | 14.760 | 15.152 | 14.965 | 3512302 | 28.095 |
| 2 | 15.472 | 15.928 | 15.662 | 2627311 | 21.016 |
| 3 | 16.160 | 16.616 | 16.351 | 3662773 | 29.299 |
| 4 | 16.952 | 17.464 | 17.240 | 2699120 | 21.590 |
| Total | | | | 12501507 | 100.000 |

c) LC trace for derivatization of mixture of (*rac*)-**3r and** (*rac*)-**3s**

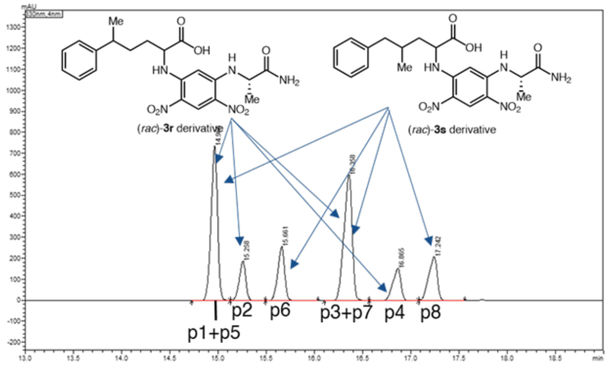

| Peak# | Peak Start | Peak End | Ret. Time | Area | Area% |
|---|---|---|---|---|---|
| 1 | 14.728 | 15.128 | 14.965 | 3449000 | 30.333 |
| 2 | 15.128 | 15.496 | 15.258 | 889443 | 7.822 |
| 3 | 15.496 | 16.040 | 15.661 | 1272280 | 11.189 |
| 4 | 16.104 | 16.568 | 16.358 | 3548612 | 31.209 |
| 5 | 16.568 | 17.080 | 16.865 | 915748 | 8.054 |
| 6 | 17.080 | 17.560 | 17.242 | 1295328 | 11.392 |
| Total | | | | 11370411 | 100.000 |

d) LC trace for derivatization of **L-amino acid synthase** reaction

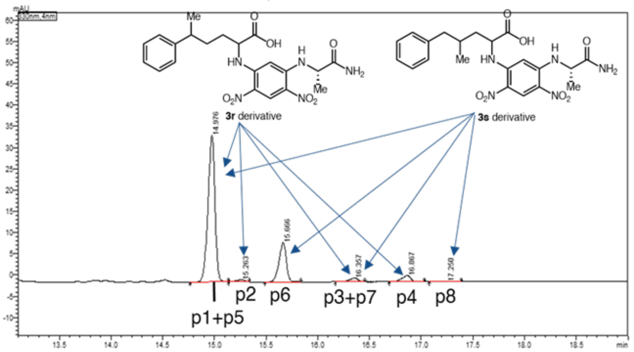

| Peak# | Peak Start | Peak End | Ret. Time | Area | Area% |
|---|---|---|---|---|---|
| 1 | 14.768 | 15.136 | 14.976 | 159740 | 72.456 |
| 2 | 15.136 | 15.336 | 15.263 | 2467 | 1.119 |
| 3 | 15.488 | 15.840 | 15.666 | 45073 | 20.445 |
| 4 | 16.168 | 16.456 | 16.357 | 4756 | 2.157 |
| 5 | 16.688 | 17.032 | 16.867 | 8060 | 3.656 |
| 6 | 17.080 | 17.392 | 17.250 | 367 | 0.167 |
| Total | | | | 220464 | 100.000 |

**Figure S11.** HPLC trace for the product analysis using (*R*)-**1r** as substrate

To unambiguously determine product stereoisomer distribution, a parallel reaction using **1s** as the substrate was conducted under the same photobiocatalytic conditions (Figure S12). As the same homobenzyl radical intermediate is generated either directly from **1s** or from **1r** rearrangement, the diastereo- and enantiomeric ratios of 3s derived from **1s** are likely the same as that derived from (*R*)-**1r**. The reaction of **1s** provided product **3s** in 83% yield, with a diastereomeric ratio (d.r.) of 2.8:1 as determined by the peak area ratio [*A*(p6) + *A*(p8)] / [*A*(p5) + *A*(p7)]. The major diastereomer of **3s** showed an enantiomeric ratio (e.r.) of 99:1 as determined by the peak area ratio *A*(p6)/*A*(p8), while the minor diastereomer of **3s** showed an e.r. of 73:27 as determined by the peak area ratio *A*(p5)/*A*(p7) (Figure S13).

Based on these results, the diastereomeric ratio of **3r** derived from **1r** was determined to be 15:1 as calculated by the peak area ratio [*A*(p1) + *A*(p3)] / [*A*(p2) + *A*(p4)]. The enantiomeric ratio of the major enantiomer of **3r** derived from **1r** was determined to be 99:1 as calculated by the peak area ratio *A*(p1)/*A*(p3). The yields of **3r** and **3s** were determined to be (38 ± 1)% and (14 ± 1)%, respectively, based HPLC analysis with the aid of calibration curves using an internal standard.

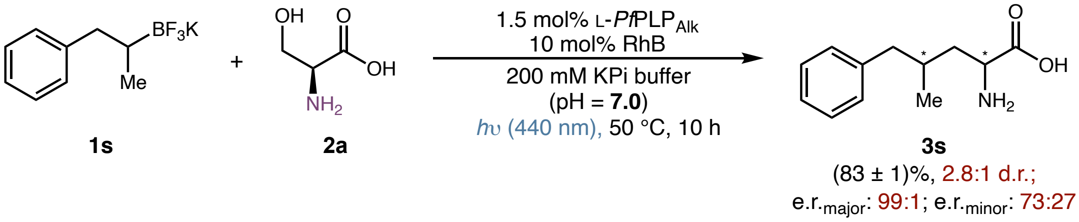

**Figure S12.** Photobiocatalytic reaction of **1s** using L-*Pf*PLP$_{Alk}$

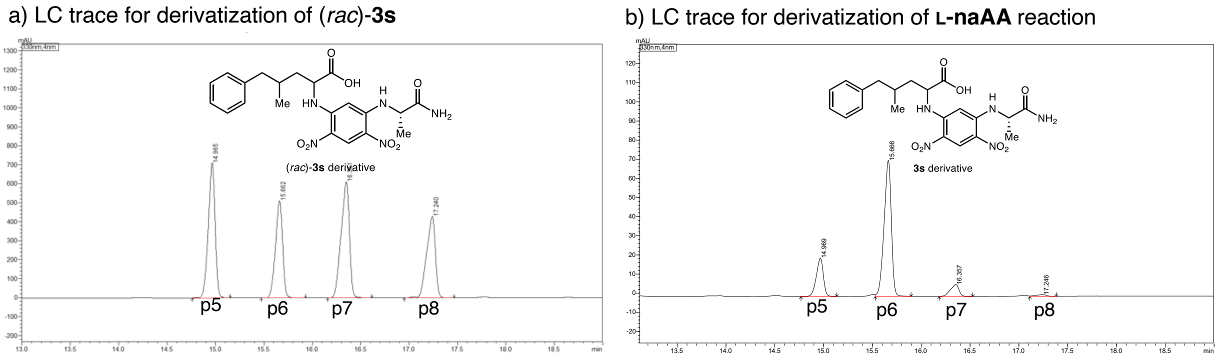

a)

| Peak# | Peak Start | Peak End | Ret. Time | Area | Area% |
|---|---|---|---|---|---|
| 1 | 14.760 | 15.152 | 14.965 | 3512302 | 28.095 |
| 2 | 15.472 | 15.928 | 15.662 | 2627311 | 21.016 |
| 3 | 16.160 | 16.616 | 16.351 | 3662773 | 29.299 |
| 4 | 16.952 | 17.464 | 17.240 | 2699120 | 21.590 |
| Total | | | | 12501507 | 100.000 |

b)

| Peak# | Peak Start | Peak End | Ret. Time | Area | Area% |
|---|---|---|---|---|---|
| 1 | 14.768 | 15.136 | 14.969 | 99115 | 19.519 |
| 2 | 15.528 | 15.896 | 15.666 | 366259 | 72.130 |
| 3 | 16.184 | 16.528 | 16.357 | 36827 | 7.253 |
| 4 | 17.112 | 17.384 | 17.246 | 5577 | 1.098 |
| Total | | | | 507777 | 100.000 |

**Figure S13.** HPLC trace for the product analysis using **1s** as substrate

#### 2-Amino-5-phenylhexanoic acid (3r)

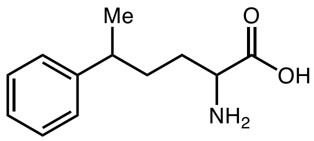

$^{1}$H NMR (400 MHz, $D_2O$) δ: 7.64 – 6.98 (m, 5H), 3.96 (dt, *J* = 9.1, 6.1 Hz, 1H), 2.71 (q, *J* = 7.0 Hz, 1H), 2.23 – 1.47 (m, 4H), 1.17 (dd, *J* = 7.0, 1.4 Hz, 3H) ppm. Major diastereomer $^{13}$C NMR (101 MHz, $D_2O$) δ: 183.1, 147.8, 128.6, 127.1, 56.0 (d, *J* = 2.8 Hz), 39.2 (d, *J* = 4.8 Hz),

33.5, 33.2, 21.7 ppm. Minor diastereomer: $^{13}$C NMR (101 MHz, $D_2O$) δ: 183.2, 147.8, 128.6, 126.0, 56.1 (d, $J$ = 2.9 Hz), 39.15 (d, $J$ = 4.8 Hz), 33.7, 33.3, 21.8 ppm. IR (neat) $\nu_{max}$: 3394, 2847, 1722, 1498, 1208, 1094, 831, 695, 634 cm$^{-1}$. HRMS (ESI) (m/z) calculated for $C_{12}H_{18}NO_2$ [M+H]$^+$: 208.1338, found 208.1342.

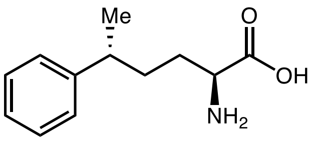

**3r** was prepared according to General Procedure using L-*Pf*PLP$_{Alk}$ as the enzyme and **1r** as the substrate at pH 7.0 KPi buffer. The reactions were carried out in triplicates. **3r** was derived using (*S*)-Marfey's reagent and analyzed by LC-MS.

**Yield:** run 1: 38.9%; run 2: 37.0%; run 3: 37.7%; average yield: (38 ± 1)%.

**Diastereomeric ratio by LC-MS**: run 1: 93.2:1.6:0.1:5.1; run 2: 93.6:1.3:0.1:5.0; run 3: 93.4:1.4:0.1:5.0.

**Diastereoselectivity by LC-MS** ($A_1$+$A_3$)/($A_2$+$A_4$)**:** run 1: 14:1 d.r.; run 2: 15:1 d.r.; run 3: 15:1 d.r.; average d.r.: 15:1.

**Enantioselectivity of major diastereomer by LC-MS** $A_1$/$A_3$: run 1: >99:1 e.r.; run 2: >99:1 e.r.; run 3: >99:1 e.r.; average e.r.: >99:1.

**2-Amino-4-methyl-5-phenylpentanoic acid (3s)**

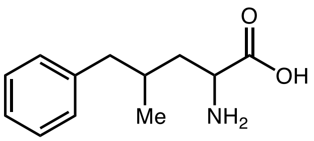

$^1$H NMR (400 MHz, $D_2O$) δ: 8.01 – 6.40 (m, 5H), 3.87 (dt, $J$ = 9.1, 6.1 Hz, 1H), 2.63 (q, $J$ = 7.0 Hz, 1H), 2.12 – 1.30 (m, 4H), 1.09 (dd, $J$ = 7.0, 1.4 Hz, 3H) ppm. Major diastereomer $^{13}$C NMR (101 MHz, $D_2O$) δ: 183.5, 141.4, 129.4, 128.3, 125.8, 54.5 (d, $J$ = 2.7 Hz), 43.2, 42.4, 31.4, 19.2. ppm; Minor diastereomer $^{13}$C NMR (126 MHz, $D_2O$) δ: 183.8, 141.4, 129.4, 125.8, 54.10 (d, $J$ = 2.8 Hz), 42.7, 42.3, 31.4, 18.0. IR (neat) $\nu_{max}$: 3421, 3078, 2924, 1618, 1585, 1411, 1307, 837, 679 cm$^{-1}$. HRMS (ESI) (m/z) calculated for $C_{12}H_{18}NO_2$ [M+H]$^+$: 208.1338, found 208.1342.

**Yield:** run 1: 14.0%; run 2: 14.2%; run 3: 14.3%; average yield: (14 ± 1)%.

**Diastereomeric ratio by LC-MS**: run 1: 19.5:72.1:7.2:1.1; run 2: 19.3:72.6:7.1:1.0; run 3: 19.3:72.4:7.1:1.1.

**Diastereoselectivity by LC-MS** ($A_1$+$A_3$)/($A_2$+$A_4$)**:** run 1: 2.7:1 d.r.; run 2: 2.7:1 d.r.; run 3: 2.8:1 d.r.; average d.r.: 2.7:1.

**Enantioselectivity of major diastereomer diastereomer by LC-MS** $A_2/A_4$: run 1: 98:2 e.r.; run 2: 99:1 e.r.; run 3: 98:2 e.r.; average e.r.: 98:2. **Minor diastereomer by LC-MS** $A_1/A_3$: run 1: 73:27 e.r.; run 2: 73:27 e.r.; run 3: 73:27 e.r.; average e.r.: 73:27.

#### 4. Isomerization via radical-mediated 1,5-H atom transfer (1,5-HAT)

When substrates **4n** (C6), **4o** (C8), and **4p** (C10) were used for photobiocatalytic coupling, isomerization products **5n'**, **5o'** and **5p'** were observed, respectively (Figure S14). Using **4n** (C6), **4o** (C8), and **4p** (C10) as the substrate and L-*Pf*$PLP_{Alk}$ as the enzyme, the unisomerized : isomerized product ratios were 42:1, 40:1 and 65:1, respectively. Using **4n** (C6), **4o** (C8), and **4p** (C10) as the substrate and D-*Pf*$PLP_{Alk2}$ as the enzyme, the unisomerized : isomerized product ratios were 23:1, 27:1 and 80:1, respectively. The formation of isomerized products is proposed to involve an intramolecular **1**,5-hydrogen atom transfer (1,5-HAT) mechanism to generate a more stable secondary radical.

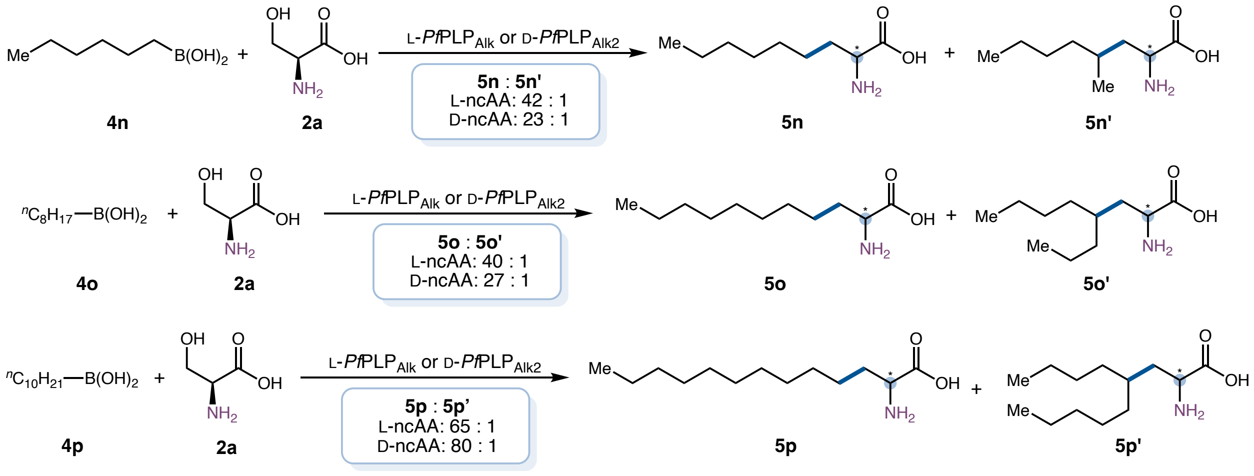

**Figure S14.** Analysis of byproduct formation using **4n**, **4o** and **4p** as substrates

#### 5. Radical trapping experiments

##### 5.1 TEMPO trapping experiments

To probe the involvement of radical intermediates, the model reaction using **1a** as the substrate and L-*Pf*PLP$_{Alk}$ as the enzyme was carried out in the presence of 3 equiv TEMPO. Under these conditions, the formation of desired product **3a** was not observed. Instead, TEMPO trapping adduct **S1** (*m/z* = 261) was detected by mass spectrometry, consistent with the proposed radical mechanism (Figure S15). Similar results were obtained when **4a** was used as the substrate (Figure S16).

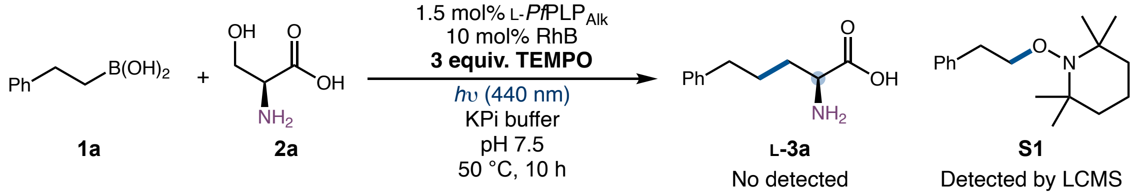

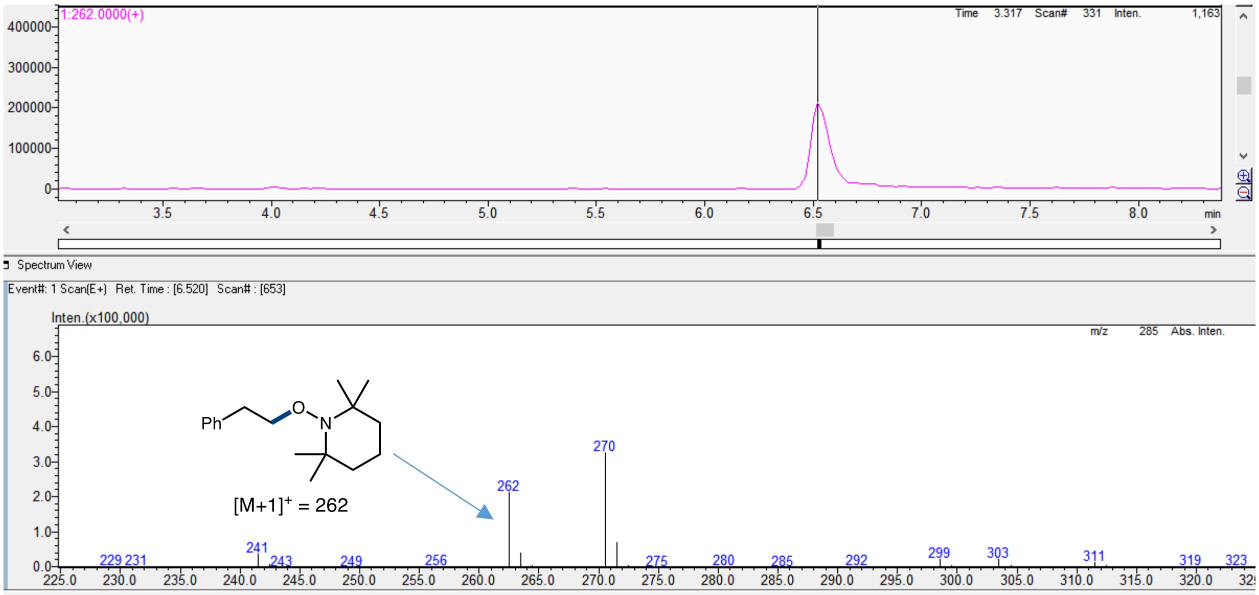

**Figure S15.** Radical trapping experiment with TEMPO using **1a** as the substrate

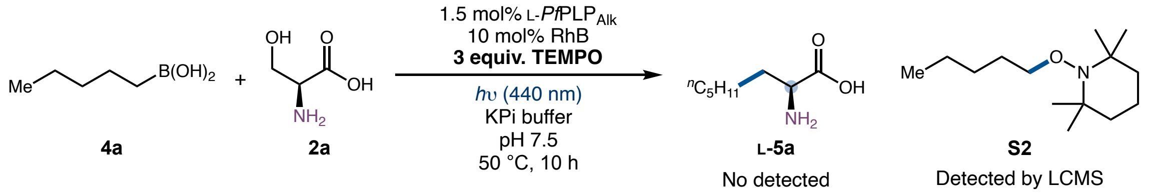

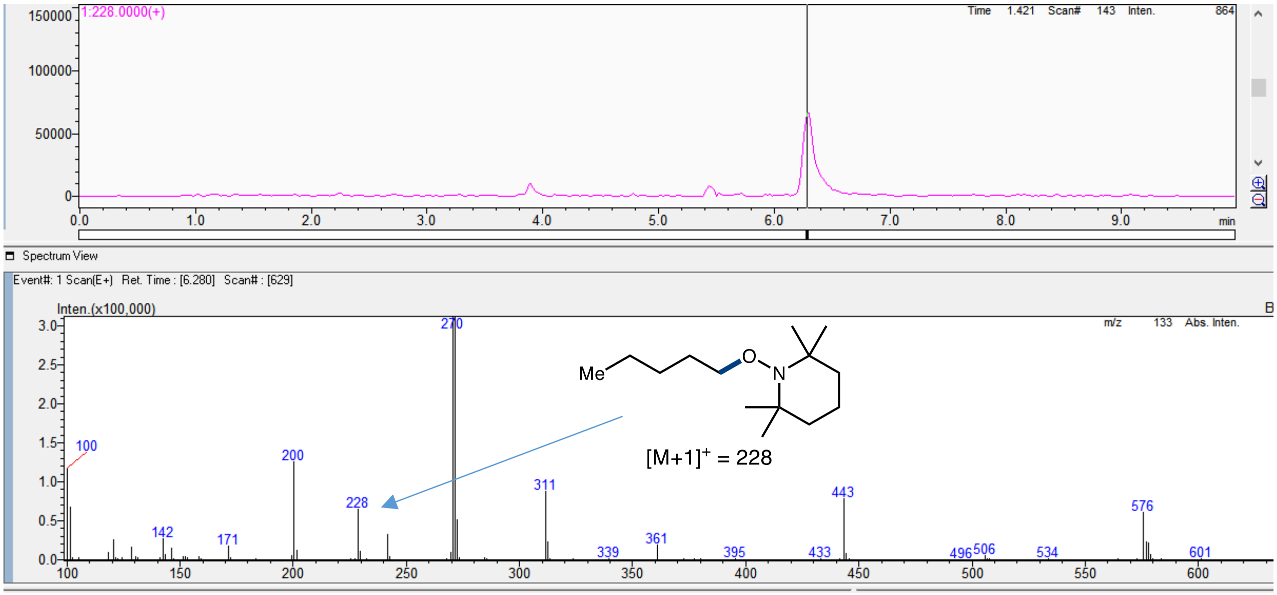

**Figure S16.** Radical trapping experiment with TEMPO using **4a** as the substrate

#### 5.2 Radical Addition to Michael Acceptors

Radical-trapping experiments using **1a** and methyl methacrylate were carried out under the standard reaction conditions in the absence of the PLP enzyme. The corresponding radical addition products were detected in GC-MS. $^1$H NMR spectroscopic analysis showed the yield of radical addition adducts **1** and **2** to be 35% and 27%, respectively. These results indicated that radical intermediates could form in the absence of the PLP enzyme (Figure S17).

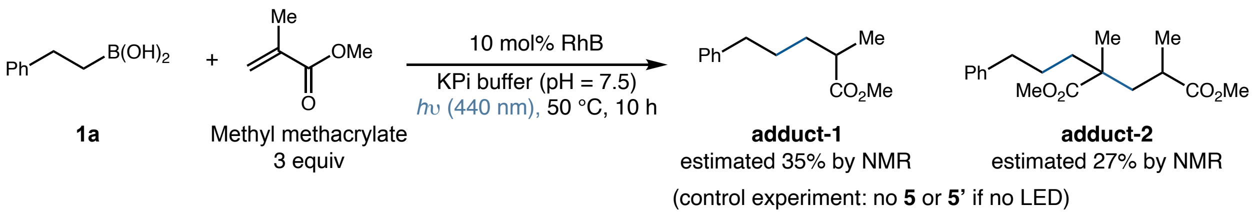

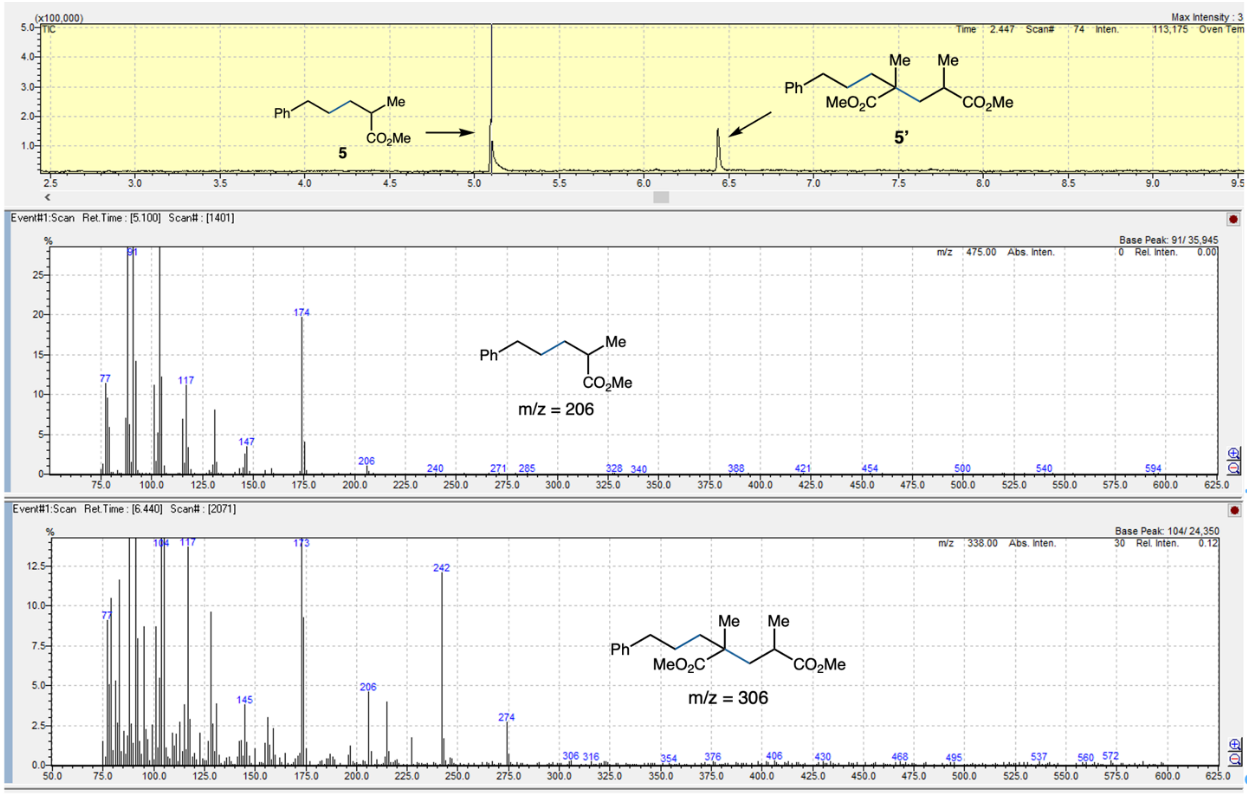

**Figure S17.** Radical trapping experiment with methyl methacrylate using **1a** as the substrate

### 6. Kinetic profiles of the model reactions

#### 6.1 Kinetic profile of L-*Pf*PLP$_{Alk}$/RhB-catalyzed reaction

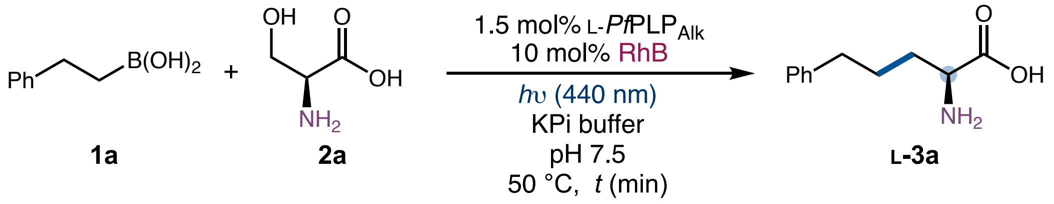

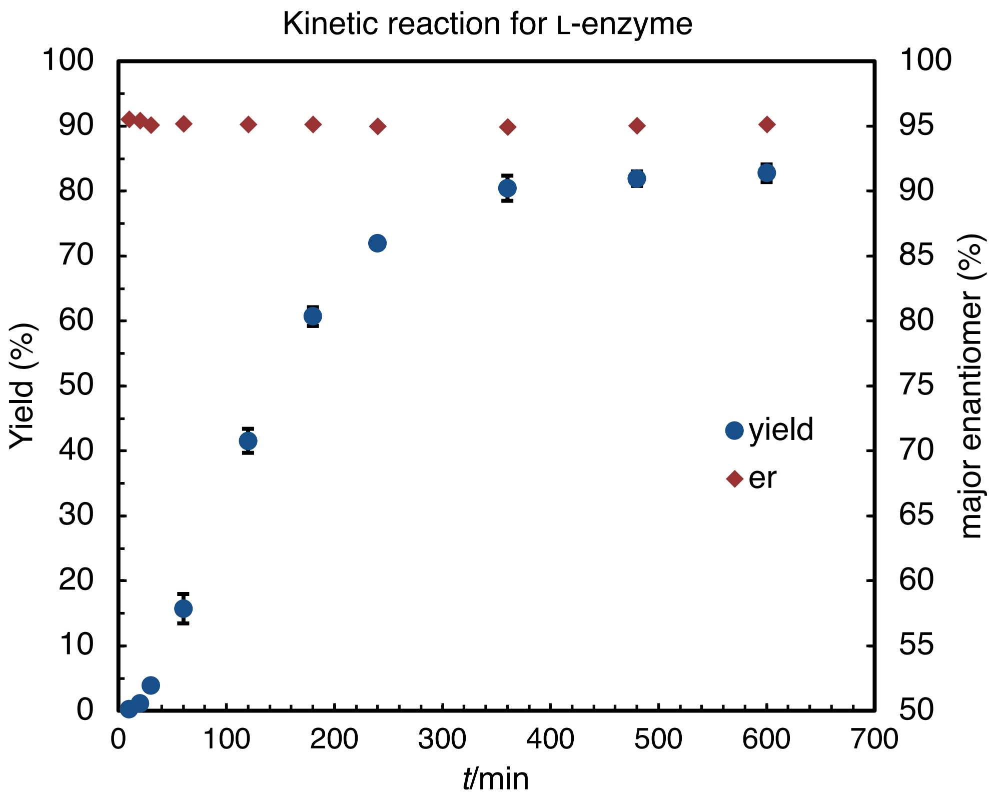

**Figure S18.** Kinetic profile of L-*Pf*PLP$_{Alk}$/RhB-catalyzed reaction

**Experimental procedure.** Following the General Procedure, L-*Pf*PLP$_{Alk}$/RhB-catalyzed reaction was quenched and analyzed at 10, 20, 30, 60, 120, 240, 360, 480, and 720 min, respectively. Product yields and enantioselectivities were plotted as a function of time (Figure S18). This dual catalytic reaction was found to reach full conversion between 360 and 480 min. An initial induction period was observed within the first 30 min, during which less than 5% product formation was detected. The enantioselectivity remained constant throughout the course of the transformation.

##### 6.2 Kinetic profile of D-*Pf*PLP$_{Alk1}$/Eosin B-catalyzed reaction

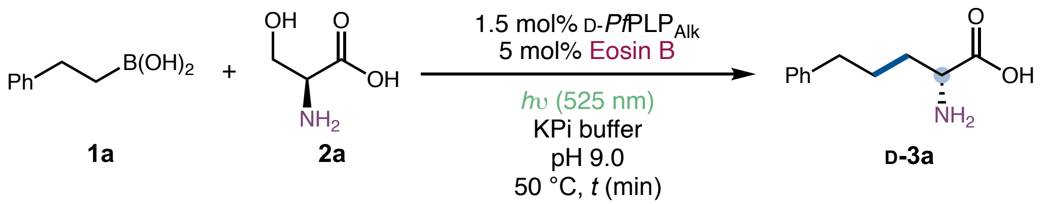

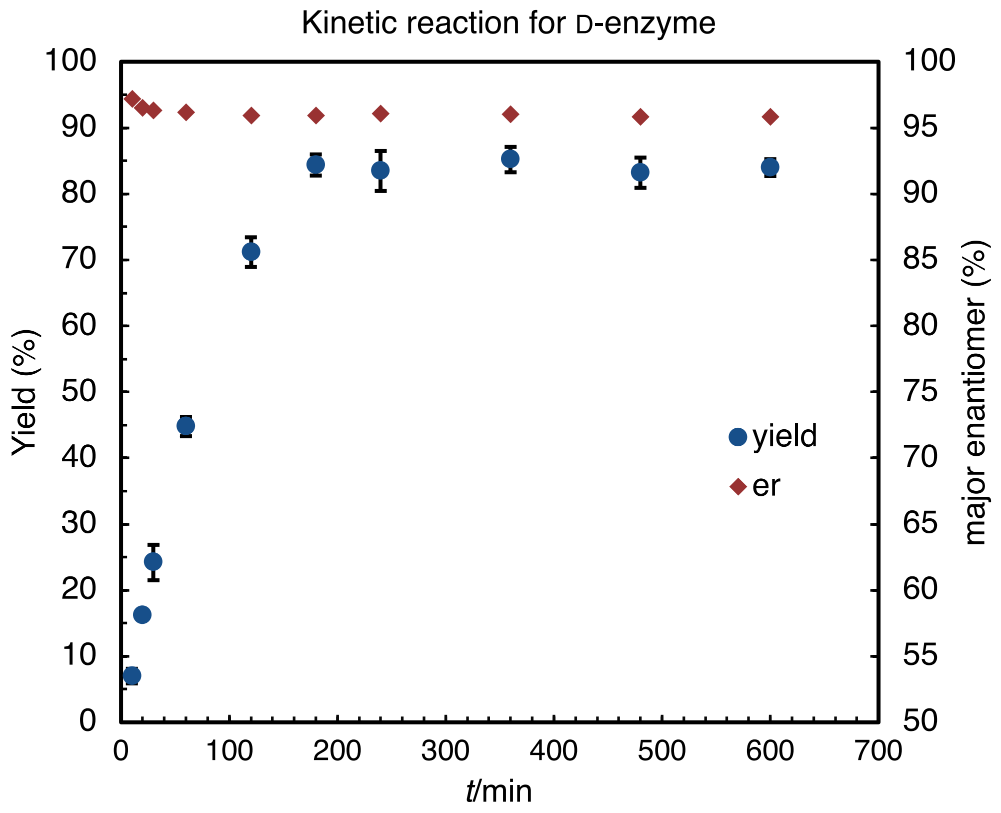

**Figure S19.** Kinetic profile of D-*Pf*PLP$_{Alk1}$/Eosin B-catalyzed reaction

Following the General Procedure, D-*Pf*PLP$_{Alk1}$/Eosin B-catalyzed reaction was quenched and analyzed at 10, 20, 30, 60, 120, 240, 360, 480, and 720 min, respectively. Product yields and enantioselectivities were plotted as a function of time (Figure S19). In contrast to L-*Pf*PLP$_{Alk}$/RhB-catalyzed reaction, D-*Pf*PLP$_{Alk1}$/Eosin B-catalyzed reaction reached full conversion within 180 min. No induction period was observed.

#### 7. Photobiocatalytic reaction in $D_2O$ buffer

**Table S18.** Buffer effect on L-*Pf*PLP$_{Alk}$/RhB-catalyzed synthesis of L-**3a**

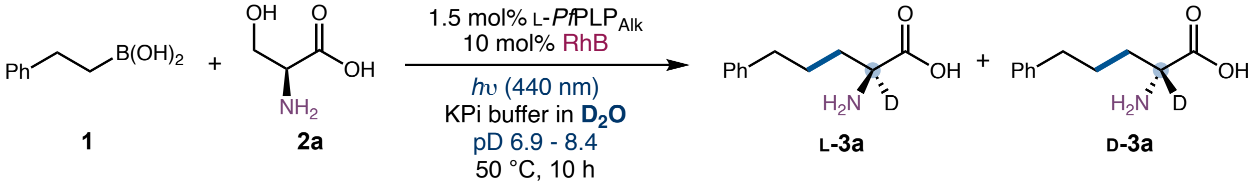

| entry | pD | yield of **3a** | e.r. (L-**3a**: D-**3a**) |
|---|---|---|---|
| 1 | 8.4 | 80% (82%)[a] | 97:3 (93:7)[a] |
| 2 | 7.9 | 79% (83%)[a] | 98:2 (95:5)[a] |
| 3 | 7.4 | 76% (79%)[a] | 99:1 (96:4)[a] |
| 4 | 6.9 | 75% (76%)[a] | 99:1 (97:3)[a] |

Reaction conditions: **1a** (4.0 mM), **2a** (20.0 mM), 1.5 mol% L-*Pf*PLP$_{Alk}$, 10 mol% RhB, *h*ν (440 nm, 5.0 W), 200 mM KPi buffer ($D_2O$), DMSO (8% v/v), 50 °C, 10 h. [a]Yield and e.r. of **3a** obtained with $H_2O$-based KPi buffer were reported in parentheses. pH = pD – 0.4.

**Table S19.** Buffer effect on L-*Pf*PLP$_{Alk}$/RhB-catalyzed synthesis of L-**5a**

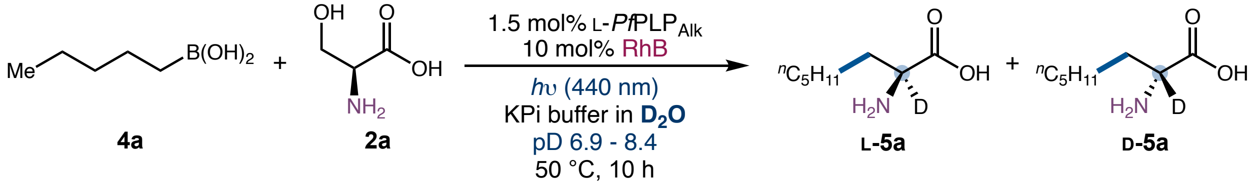

| entry | pD | yield of **5a** | e.r. (L-**5a**: D-**5a**) |
|---|---|---|---|
| 1 | 8.4 | 58% | 96:4 |

| | | | |
|---|---|---|---|
| | | (71%) [a] | (90:10) [a] |
| 2 | 7.9 | 63%<br>(74%) [a] | 97:3<br>(94:6) [a] |
| 3 | 7.4 | 62%<br>(69%) [a] | 99:1<br>(96:4) [a] |
| 4 | 6.9 | 62%<br>(66%) [a] | 99:1<br>(97:3) [a] |

Reaction conditions: **4a** (4.0 mM), **2a** (20.0 mM), 1.5 mol% L-*Pf*PLP$_{Alk}$, 10 mol% RhB, *h*ν (440 nm, 4.0 W), 200 mM KPi buffer ($D_2O$), DMSO (8% v/v), 50 °C, 10 h. [a] Yields and e.r. of **5a** obtained with $H_2O$-based KPi buffer were reported in parentheses. pH = pD – 0.4.

For L-*Pf*PLP$_{Alk}$/RhB-catalyzed reaction of **1a** and **4a** in $D_2O$-based buffer, a slight decrease in yield was observed, while the overall trends in yield and selectivity remained consistent with those observed under varying pH conditions in $H_2O$-based buffers (Table S18 and S19). Switching the reaction medium from $H_2O$ to $D_2O$ led to enhancement in the enantiomeric purity of L-**3a** and L-**5a**, suggesting that the formation of D-**3a** and D-**5a** is slower in $D_2O$.

**Table S20.** Buffer effect on D-*Pf*PLP$_{Alk1}$/Eosin B-catalyzed synthesis of D-**3a**

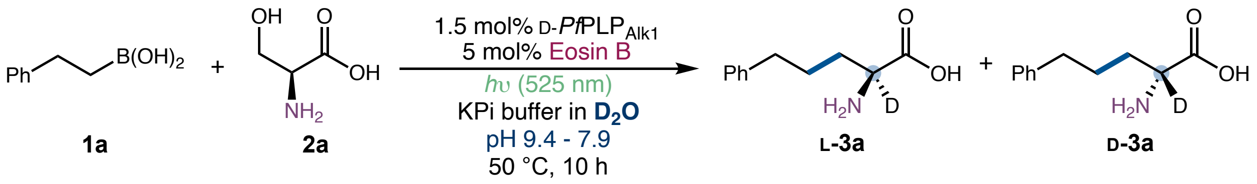

| entry | pD | yield of **3a** | e.r. (L-**3a**: D-**3a**) |
|---|---|---|---|
| 1 | 9.4 | 70%<br>(84%) [a] | 23:77<br>(4:96) [a] |
| 2 | 8.9 | 72%<br>(84%) [a] | 24:76<br>(5:95) [a] |

| | | | |
|---|---|---|---|
| 3 | 8.4 | 73% (86%) [a] | 27:73 (7:93) [a] |
| 4 | 7.9 | 70% (82%) [a] | 33:67 (11:89) [a] |

Reaction conditions: **4a** (4.0 mM), **2a** (20.0 mM), 1.5 mol% D-*Pf*PLP$_{Alk1}$, 5 mol% Eosin B, *h*ν (525 nm, 3.0 W), 200 mM KPi buffer ($D_2O$), DMSO (8% v/v), 50 °C, 10 h. [a] Yields and e.r. of **3a** obtained with $H_2O$-based KPi buffer were reported in parentheses. pH = pD – 0.4.

For D-*Pf*PLP$_{Alk1}$/Eosin B-catalyzed reaction of **1a** in $D_2O$-based buffers, a slight decrease in yield was observed, while the overall trends in yield and enantioselectivity remained consistent with those observed under varying pH conditions in $H_2O$-based buffers (Table S20). When the reaction medium was switched from $H_2O$ to $D_2O$, a significant decrease in the enantiomeric purity of D-**3a** was observed, suggesting that the formation of D-**3a** proceeds more slowly in $D_2O$.

**Table S21.** Buffer effect on D-*Pf*PLP$_{Alk2}$/RhB-catalyzed synthesis of D-**5a**

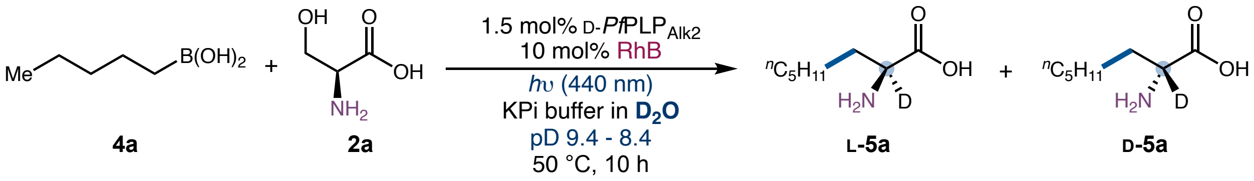

| entry | pD | yield of **3a** | e.r. (L-**3a**: D-**3a**) |
|---|---|---|---|
| 1 | 9.4 | 31% (77%) [a] | 14:86 (3:97) [a] |
| 2 | 8.9 | 36% (76%) [a] | 18:82 (4:96) [a] |
| 3 | 8.4 | 34% (76%) [a] | 19:81 (5:95) [a] |

Reaction conditions: **4a** (4.0 mM), **2a** (20.0 mM), 1.5 mol% D-*Pf*PLP$_{Alk2}$, 10 mol% RhB, *h*ν (440 nm, 4.0 W), 200 mM KPi buffer ($D_2O$), DMSO (8% v/v), 50 °C, 10 h. [a] Yields and e.r. of **5a** obtained with $H_2O$-based KPi buffer were reported in parentheses. pH = pD − 0.4.

For D-*Pf*PLP$_{Alk2}$/RhB-catalyzed reaction of **4a**, switching the reaction medium from $H_2O$ to $D_2O$ resulted in a more significant decrease in both the yield and enantiomeric purity of D-**5a** (Table S21). Similar to D-*Pf*PLP$_{Alk1}$/Eosin B-catalyzed reaction (Table S20), this observation suggests that the formation of D-configured products proceeds more slowly in $D_2O$. The E104P mutation may contribute to the observed decrease in yield[7].

### V. Extended scope

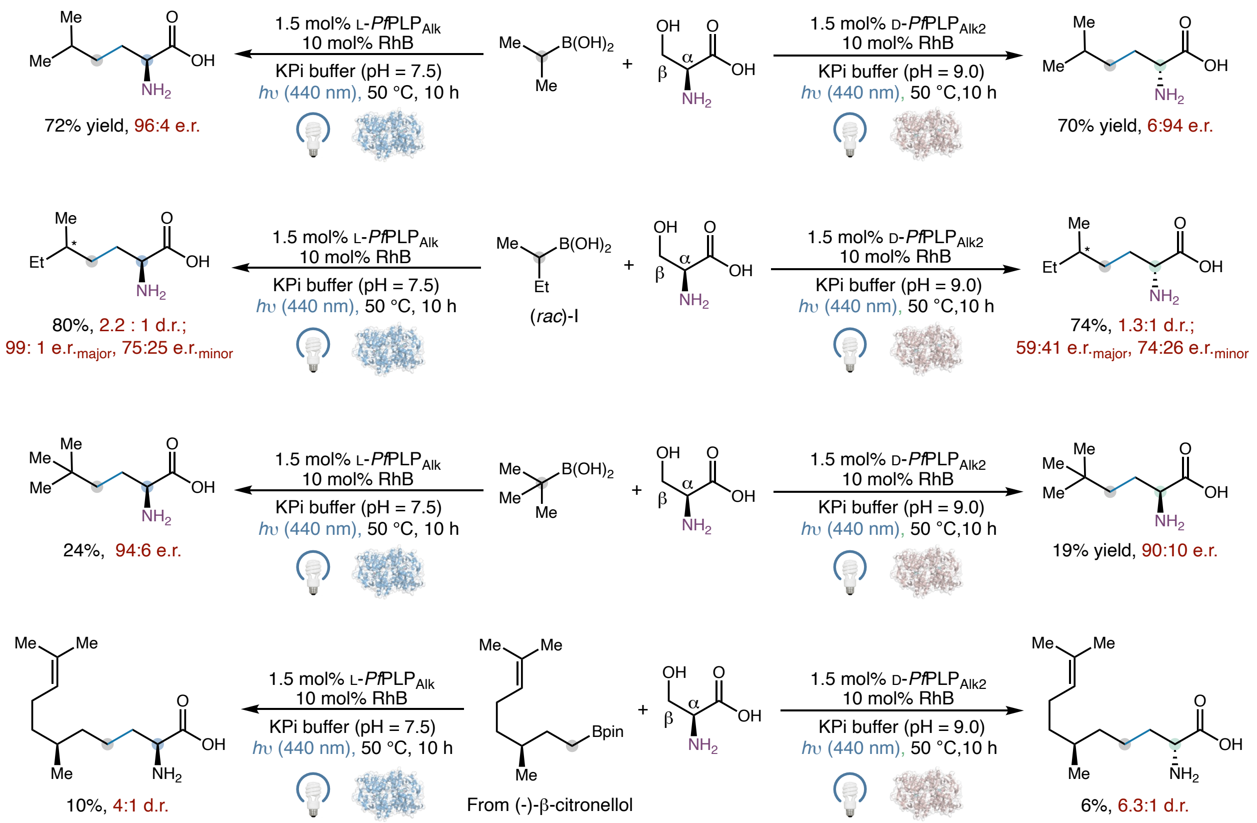

**Figure S20.** Extended substrate scope results.

#### VI. Synthesis and characterization of substrates

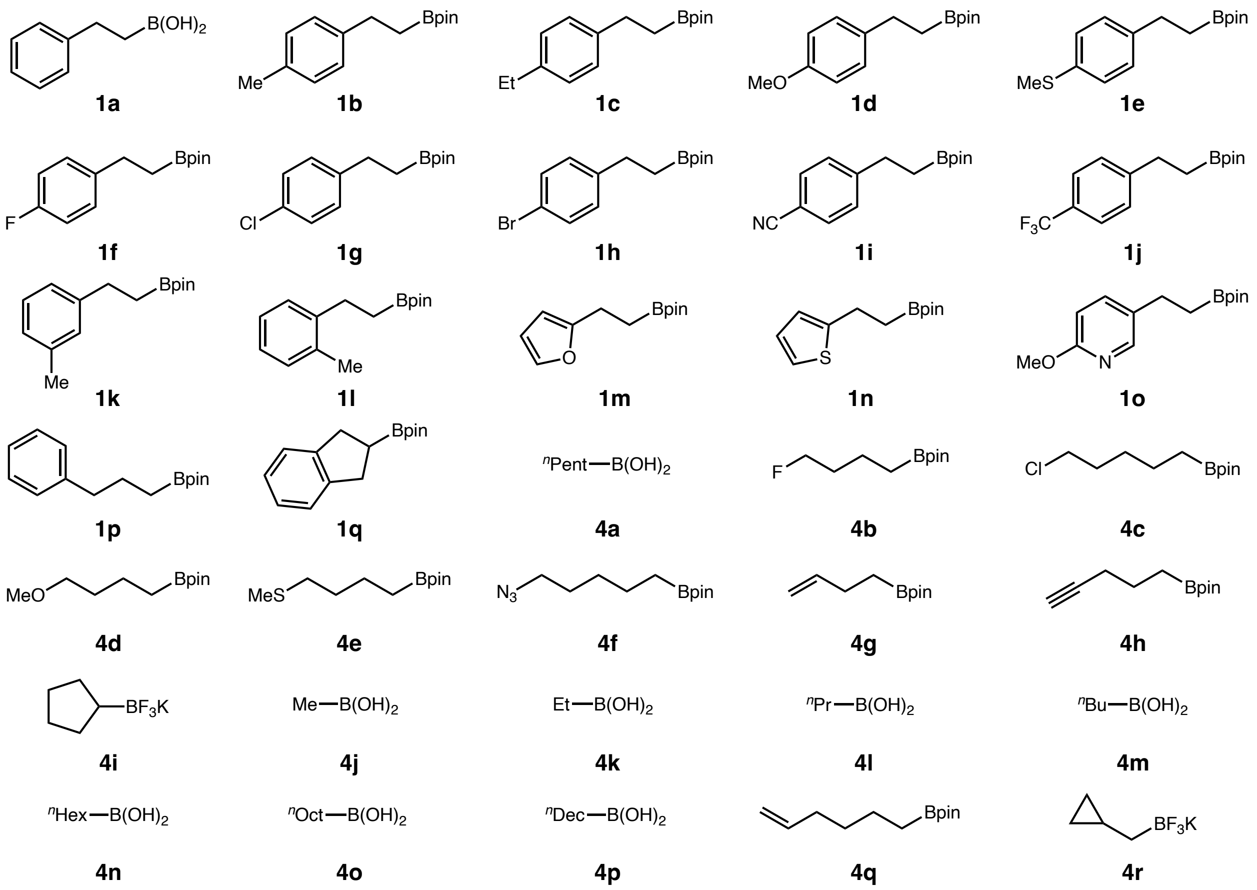

Compounds **1a, 1q, 4a, 4g**, **4i**–**4r** were commercially available from Combi-Blocks and they were used as received without further purification. All other substrates were synthesized according to General Procedure A, B, and C as detailed below.

##### General Procedure A

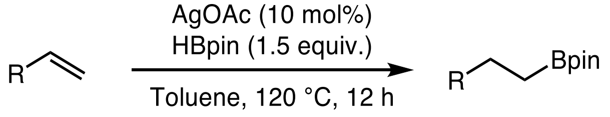

This procedure was modified from a previous method[8]. To a 100 mL round-bottom flask equipped with a stir bar were added AgOAc (1.0 mmol, 0.1 equiv), alkene (10.0 mmol, 1.0 equiv) and toluene (0.50 M). The flask was evacuated and backfilled with nitrogen (this process was repeated three times). HBpin (15.0 mol, 1.5 equiv) was added dropwise by syringe over 2 min under nitrogen. The resulting mixture was stirred at 120 °C for 12 h. After cooling to room temperature, the reaction mixture was diluted with 30 mL EtOAc and filtered through a celite plug, eluting with 10–20 mL of EtOAc. The combined organic layers were dried over $MgSO_4$ and concentrated *in vacuo* with the aid of a rotary evaporator to provide the crude product, which was further purified by column chromatography with the aid of a Biotage Isolera to afford the desired product.

Compounds **1b**[8], **1c**[8], **1d**[9], **1e**[8], **1f**[8], **1g**[10], **1h**[8], **1i**[9], **1j**[9], **1k**[10], **1l**[11], **1o, 1p**[8], and **4e**[12] were prepared according to General Procedure A.

**General Procedure B**[13]

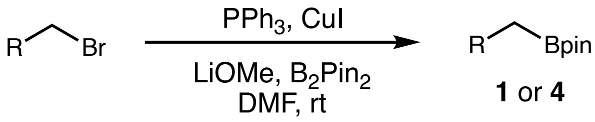

CuI (1.5 mmol, 0.1 equiv), $PPh_3$ (2.0 mmol, 0.13 equiv), LiOMe (30.0 mmol, 2.0 equiv) and bis(pinacolato)diboron (22.5 mmol, 1.5 equiv) were added to a 100 mL round-bottom flask. The flask was evacuated and backfilled with nitrogen (this process was repeated three times). DMF (30 mL) and benzyl bromide (15.0 mmol, 1.0 equiv) were then added under nitrogen. The resulting mixture was allowed to stir at 25 °C for 18 h. The reaction mixture was then diluted with EtOAc (100 mL) and filtered through a silica plug. The filtrate was washed with brine (3 × 100 mL). The organic layer was dried over $MgSO_4$ and concentrated *in vacuo* with the aid of a rotary evaporator to provide the crude product, which was further purified by column chromatography with the aid of a Biotage Isolera to afford the desired product.

Compound **1n**[9], **4b**, **4c**[14] and **4d**[15] was prepared following General Procedure B.

**General Procedure C**

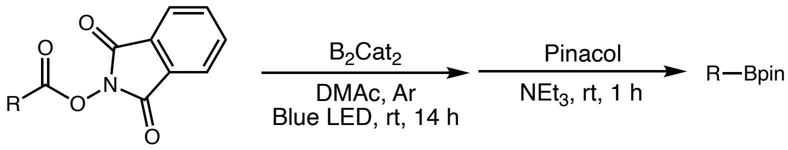

This procedure was modified from a previous method[16]. NHPI ester (0.4 mmol, 1.0 equiv) and $B_2cat_2$ (0.5 mmol, 1.25 equiv) were carefully weighed into a flame-dried 20 mL vial containing a small magnetic stirrer bar. DMAc (4 mL, 0.1 M) was added, then the vial was evacuated and backfilled with nitrogen (this process was repeated three times). The vial was tightly sealed and stirred under blue LED irradiation at room temperature for 14 h. Pinacol (1.6 mmol, 4 equiv) was dissolved in $Et_3N$ (1.5 mL), added to the reaction mixture and stirred for 1 h. The reaction mixture was quenched by 30 mL brine. 30 mL $H_2O$ was added, and the mixture was extracted with EtOAc for 3 times. The organic layers were combined, dried over $MgSO_4$ and concentrated *in vacuo* to provide the crude product, which was further purified by column chromatography with the aid of a Biotage Isolera to afford the desired product.

Compounds **1m** and **4g**[16] were prepared following General Procedure C.

#### General Procedure D

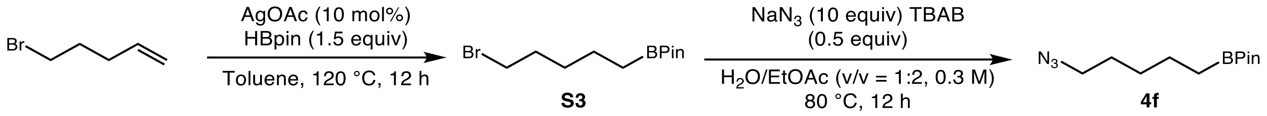

The bromo-substituted alkylboronate **S3** was synthesized according to General Procedure A.

To a round bottom flask were added the alkylboronate (5 mmol, 1 equiv), tetrabutylammonium bromide (TBAB, 2.5 mmol, 0.5 equiv) and sodium azide (50.0 mmol, 10 equiv), followed by 5 mL water and 10 ml EtOAc. The reaction mixture was heated at 80 °C overnight. The mixture was then quenched with water and extracted with EtOAc for 3 times. The organic layers were dried over $MgSO_4$, and concentrated *in vacuo*. The residue was loaded onto a pad of silica eluting with ether. The collected ether fractions were concentrated *in vacuo* to provide the azidoalkylboronate **4f** as colorless oil.

#### Characterization data for previously unknown substrates

**2-(2-(Furan-2-yl)ethyl)-4,4,5,5-tetramethyl-1,3,2-dioxaborolane (1m)**

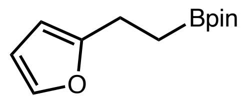

Purified by Biotage (50 g iLOK cartridge, 0–10% EtOAc/hexanes for 10 CVs, then 10% EtOAc/hexanes for 5 CVs) to afford the product as a colorless oil. $^{1}$H NMR (400 MHz, $CDCl_3$) δ: 7.27 (d, *J* = 1.9 Hz, 1H), 6.25 (dd, *J* = 3.2, 1.9 Hz, 1H), 5.96 (d, *J* = 3.2 Hz, 1H), 2.74 (t, *J* = 7.9 Hz, 2H), 1.23 (s, 12H), 1.13 (t, *J* = 7.9 Hz, 2H) ppm. $^{13}$C NMR (101 MHz, $CDCl_3$) δ: 157.9, 140.6, 110.0, 103.9, 83.2, 24.8, 22.4 ppm. $^{11}$B NMR (160 MHz, $CDCl_3$) δ: 34.13 ppm. IR (neat) $\nu_{max}$: 2978, 2934, 1372, 1318, 1143, 1006, 963, 848 $cm^{-1}$. HRMS (EI) (m/z) calculated for $C_{12}H_{19}BO_3$ $[M]^+$: 222.1427, found 222.1431.

**2-Methoxy-5-(2-(4,4,5,5-tetramethyl-1,3,2-dioxaborolan-2-yl)ethyl)pyridine (1o)**

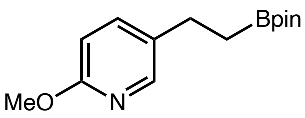

Purified by Biotage (50 g iLOK cartridge, 0–15% EtOAc/hexanes for 10 CVs, then 15% EtOAc/hexanes for 5 CVs) to afford the product as a colorless oil. $^{1}$H NMR (400 MHz, $CDCl_3$) δ: 7.98 (d, *J* = 2.4 Hz, 1H), 7.42 (dd, *J* = 8.5, 2.4 Hz, 1H), 6.64 (d, *J* = 8.5 Hz, 1H), 3.88 (s, 3H), 2.64 (t, *J* = 8.0 Hz, 2H), 1.20 (s, 13H), 1.08 (t, *J* = 8.0 Hz, 2H) ppm. $^{13}$C NMR (101 MHz, $CDCl_3$) δ: 162.5, 145.7, 138.7, 132.2, 110.2, 83.2, 53.3, 26.2, 24.8 ppm. $^{11}$B NMR (160 MHz, $CDCl_3$) δ: 34.11 ppm. IR (neat) $\nu_{max}$: 2978, 2940, 1607, 1487, 1372, 1285, 1143, 1034, 968, 832 $cm^{-1}$. HRMS (EI) (m/z) calculated for $C_{14}H_{22}BNO_3$ $[M]^+$: 263.1693, found 263.1696.

**2-(4-Fluorobutyl)-4,4,5,5-tetramethyl-1,3,2-dioxaborolane (4b)**

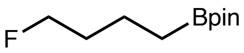

Purified by Biotage (50 g iLOK cartridge, 0–8% EtOAc/hexanes for 10 CVs, then 8–15% EtOAc/hexanes for 5 CVs) to afford the product as a colorless oil. $^{1}$H NMR (500 MHz, $CDCl_3$) δ: 4.45 (t, *J* = 6.2 Hz, 1H), 4.36 (t, *J* = 6.2 Hz, 1H), 1.75 – 1.61 (m, 2H), 1.54 – 1.46 (m, 2H), 1.22 (s, 12H), 0.79 (t, *J* = 7.9 Hz, 2H) ppm. $^{13}$C NMR (126 MHz, $CDCl_3$) δ: 83.9 (d, *J* = 164.0 Hz), 83.0, 32.8 (d, *J* = 19.2 Hz), 24.8, 19.64 (d, *J* = 6.1 Hz) ppm. $^{11}$B NMR (160 MHz, $CDCl_3$) δ: 34.09 ppm. $^{19}$F NMR (376 MHz, $CDCl_3$) δ: -218.23 ppm. IR (neat) $\nu_{max}$: 2979, 2936, 1377, 1323, 1138, 970, 840 $cm^{-1}$. HRMS (EI) (m/z) calculated for $C_{10}H_{20}BFO_2$ $[M]^+$: 202.1540, found 202.1545.

**2-(5-Azidopentyl)-4,4,5,5-tetramethyl-1,3,2-dioxaborolane (4f)**

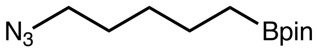

Purified by Biotage (50 g iLOK cartridge, 0–8% EtOAc/hexanes for 10 CVs, then 8–15% EtOAc/hexanes for 5 CVs) to afford the product as a colorless oil. $^{1}$H NMR (400 MHz, $CDCl_3$) δ: 3.23 (t, $J$ = 7.0 Hz, 2H), 1.59 (p, $J$ = 7.1 Hz, 2H), 1.49 – 1.31 (m, 4H), 1.23 (s, 12H), 0.77 (t, $J$ = 7.5 Hz, 2H) ppm. $^{13}$C NMR (126 MHz, $CDCl_3$) δ: 82.9, 51.4, 29.3, 28.6, 24.8, 23.5 ppm. $^{11}$B NMR (128 MHz, $CDCl_3$) δ: 34.03 ppm. IR (neat) $\nu_{max}$: 2978, 2929, 2864, 2093, 1367, 1318, 1143, 963, 842 cm$^{-1}$. HRMS (EI) (m/z) calculated for $C_{11}H_{22}BN_3O_2$ $[M]^+$: 239.1805, found 239.1809.

#### VII. Synthesis and characterization of products

##### General procedure for the synthesis of racemic products

Compounds **5a, 5j–5n** were commercially available from Combi-Blocks and they were used as received. All other products were synthesized according to General Procedure E and F as detailed below.

##### General Procedure E

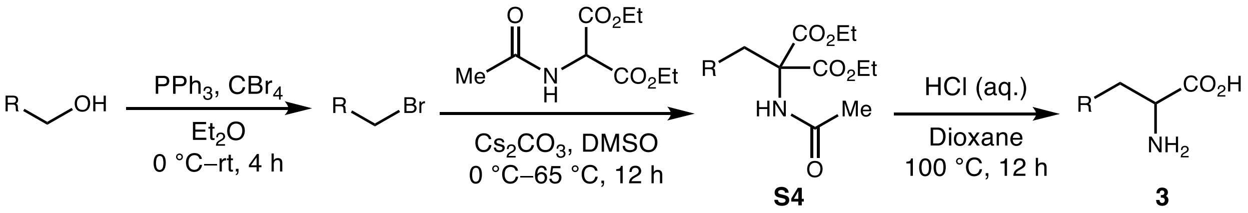

This procedure was modified from our previously reported procedure.[5] A round bottom flask was evacuated and backfilled with nitrogen and this procedure was repeated for a total of three time. The alcohol (10 mmol, 1.0 equiv) and triphenylphosphine (12 mmol, 1.2 equiv) were added, followed by $Et_2O$ (50 mL). The resulting solution was cooled to 0 °C in an ice bath. In a separate flask, $CBr_4$ (12 mmol, 1.2 equiv) was dissolved in $Et_2O$ (20 mL) and this solution was added dropwise to the reaction mixture over 5 min. The resulting mixture was stirred at 25 °C for 4 h. Upon completion of the reaction as indicated by TLC analysis, the mixture was concentrated under reduced pressure and purified by column chromatography using a Biotage Isolera system to afford the bromide product.

To a solution of diethyl acetamidomalonate (14.4 mol, 2.0 equiv) in DMSO (30 mL) was added $Cs_2CO_3$ (18.0 mol, 2.5 equiv) at 0 °C. The reaction solution was stirred at rt for 1 h. The alkyl bromide (7.2 mol, 1.0 equiv) was added to the reaction mixture at room temperature and the reaction mixture was stirred at 65 °C for 8 h. Upon the completion of this reaction, the mixture was added into ice water (80 mL) and stirred for 1 h. The precipitate was filtered and dried under vacuum to give the alkylation product **S4** as an off-white solid, which was used directly for the next step without further purification.

A suspension of this crude product (5.00 mmol, 0.2 M) in 1:1 of aq. HCl (6.0 M solution for **3a**, **3b**–**3c**, **3f**–**3l**, **3o** and **3q**; 3.0 M solution for **3d**, **3e**) and 1,4-dioxane was heated to reflux for

12 h. Upon the completion of the reaction as indicated by LC-MS analysis, the reaction mixture was concentrated *in vacuo* with the aid of a rotary evaporator and the residue was recrystallized from methanol and diethyl ether. The solid amino acid product was collected by filtration. After several times of recrystallization, this collected solid was found to be analytically pure. It can be further purified by C18 silica gel chromatography with the aid of a Biotage Isolera if necessary. Compounds **3a**−**3l**, **3o**, **3q** were prepared following General Procedure E.

**General Procedure F**

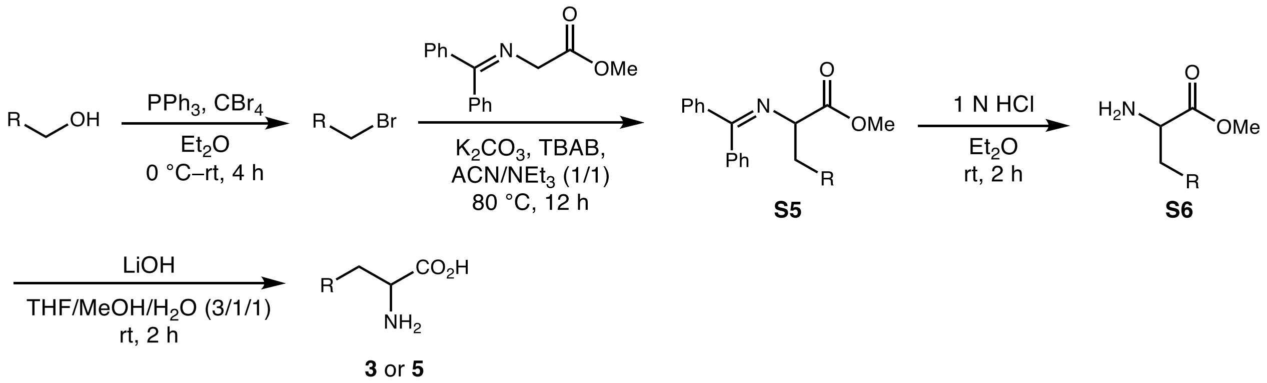

The alkyl bromides were prepared using the same procedure detailed in General Procedure E. *For alkyl bromides with a low boiling point, concentration under reduced pressure at low temperature was necessary to minimize product loss.*

The alkylation reaction using this alkyl bromide was modified from a previously reported procedure.[17] An oven-dried round bottom flask was evacuated and backfilled with nitrogen and this procedure was repeated for a total of three time.. To this flask were added methyl 2-((diphenylmethylene)amino)acetate (15 mmol 1 equiv), $K_2CO_3$ (45 mmol, 3 equiv), tetrabutylammonium bromide (TBAB, 1.5 mmol, 0.1 equiv), acetonitrile (40 mL) and alkyl bromide (22.5 mmol, 1.5 equiv). The reaction mixture was stirred at 80 °C for 12 h. After cooling to room temperature, the mixture was filtered through a celite plug, and the filtrate was concentrated under reduced pressure. The residue was purified by column chromatography with the aid of a Biotage (50 g iLOK cartridge; 0–10% EtOAc/hexanes for 10 column volumes, followed by 10–20% EtOAc/hexanes for 5 column volumes) to afford the alkylation product **S5** as a pale yellow oil.

To a solution of **S5** (10 mmol) in $Et_2O$ (50 mL) was added 1 M HCl (30 mL, 3.0 equiv). The suspension was stirred at 25 °C for 4 h. The organic phase was separated from the aqueous phase, and the aqueous layer was washed with $Et_2O$ (2 × 30 mL) and concentrated *in vacuo* to afford the hydrolysis product **S6** as a viscous residue, which was used directly in the next step without further purification.

To this crude **S6** (10 mmol) were added MeOH (10 mL), $H_2O$ (10 mL), and THF (30 mL). LiOH (100 mmol, 10 equiv) was then added, and the mixture was stirred at 25 °C for 4 h. Upon completion of the reaction as indicated by LC-MS analysis, the mixture was filtered through a celite plug, and the pH was adjusted to 7.0 with 1 M HCl. The solution was concentrated *in vacuo* with the aid of a rotary evaporator. The residue was recrystallized from methanol and diethyl ether, and the resulting amino acid product was collected by filtration. After several cycles of recrystallization, the product was determined to be analytically pure. Further purification, if necessary, was performed by C18 silica gel chromatography using a Biotage Isolera system.

Compounds **3m, 3n**, **3p**, **5b–5i, 5o–5p** were prepared following General Procedure F.

#### Characterization data for non-canonical amino acids

##### 2-Amino-5-phenylpentanoic acid (3a)[18]

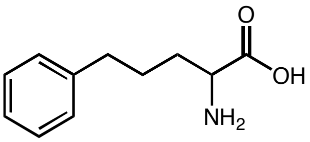

$^1$H NMR (400 MHz, $D_2O$) δ: 7.35 – 7.10 (m, 5H), 3.93 (t, $J$ = 6.3 Hz, 1H), 2.60 (t, $J$ = 7.3 Hz, 2H), 1. 95 – 1.51 (m, 4H) ppm. $^{13}$C NMR (101 MHz, $D_2O$) δ: 172.4, 141.8, 128.7, 128.6, 126.2, 53.0, 34.2, 29.21, 25.9 ppm.

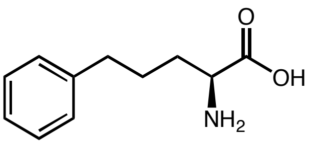

**L-3a** was prepared according to General Procedure using L-*Pf*PLP$_{Alk}$. The reactions were carried out in triplicates. **L-3a** was derivatized using (*S*)-FDAA and analyzed by LC-MS.

**Yield:** run 1: 83.4%; run 2: 81.3%; run 3: 83.6%; average yield: (83 ± 1)%.

**Enantioselectivity:** run 1: 95:5 e.r.; run 2: 95:5 e.r.; run 3: 95:5 e.r.; average e.r.: 95:5.

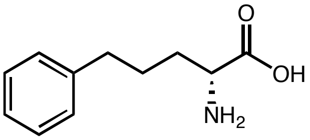

D-**3a** was prepared according to General Procedure using D-*Pf*PLP$_{Alk1}$. The reactions were carried out in triplicates. D-**3a** was derivatized using (*S*)-FDAA and analyzed by LC-MS.

**Yield:** run 1: 87.0%; run 2: 86.0%; run 3: 83.5%; average yield: (86 ± 2)%.

**Enantioselectivity:** run 1: 4:96 e.r.; run 2: 4:96 e.r.; run 3: 4:96 e.r.; average e.r.: 4:96.

##### 2-Amino-5-(p-tolyl)pentanoic acid (3b)

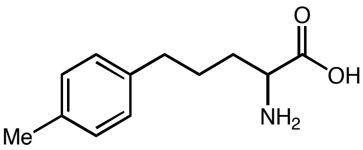

$^1$H NMR (500 MHz, $D_2O$) δ: 7.29 – 6.93 (m, 4H), 3.98 (t, $J$ = 6.2 Hz, 1H), 2.56 (t, $J$ = 7.0 Hz, 2H), 2.19 (s, 3H), 1.93 – 1.73 (m, 2H), 1.73 – 1.50 (m, 2H) ppm. $^{13}$C NMR (126 MHz, $D_2O$) δ: 172.0, 138.6, 136.1, 129.2, 128.5, 52.7, 33.7, 29.0, 25.9, 19.9 ppm. IR (neat) $\nu_{max}$: 2920, 2605, 1991, 1733, 1601, 1496, 1421, 1223, 1181, 802, 541 cm$^{-1}$. HRMS (ESI) (m/z) calculated for $C_{12}H_{18}NO_2$ $[M+H]^+$: 208.1338, found 208.1334.

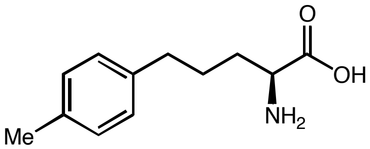

**L-3b** was prepared according to General Procedure using L-*Pf*PLP$_{Alk}$. The reactions were carried out in triplicates. **L-3b** was derivatized using (*S*)-FDAA and analyzed by LC-MS.

**Yield:** run 1: 73.6%; run 2: 81.4%; run 3: 80.0%; average yield: (78 ± 4)%.

**Enantioselectivity:** run 1: 96:4 e.r.; run 2: 96:4 e.r.; run 3: 96:4 e.r.; average e.r. 96:4.

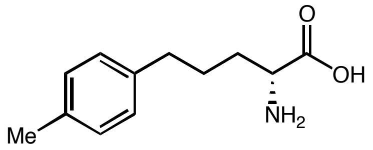

**D-3b** was prepared according to General Procedure using D-*Pf*PLP$_{Alk1}$. The reactions were carried out in triplicates. D-**3b** was derivatized using (*S*)-Marfey's reagent and analyzed by LC-MS.

**Yield:** run 1: 70.5%; run 2: 70.0%; run 3: 70.1%; average yield: (70 ± 1)%.

**Enantioselectivity:** run 1: 7:93 e.r.; run 2: 7:93 e.r.; run 3: 7:93 e.r.; average e.r.: 7:93.

**2-Amino-5-(4-ethylphenyl)pentanoic acid (3c)**

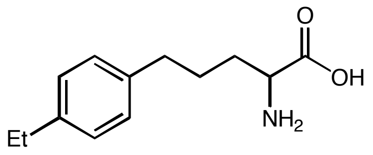

$^1$H NMR (500 MHz, $D_2O$) δ: 7.66 – 6.54 (m, 4H), 3.86 (t, *J* = 6.3 Hz, 1H), 2.97 – 2.23 (m, 4H), 1.90 – 1.58 (m, 2H), 1.58 – 1.31 (m, 2H), 0.93 (t, *J* = 7.6 Hz, 3H) ppm. $^{13}$C NMR (126 MHz, $D_2O$) δ: 183.7, 142.3, 140.1, 128.7, 127.9, 55.9, 34.5, 34.3, 27.8, 27.1, 15.2 ppm. IR (neat) $\nu_{max}$: 3017, 2929, 1749, 1509, 1214, 804, 564 cm$^{-1}$. HRMS (ESI) (m/z) for $[M+H]^+$ $C_{13}H_{20}NO_2$ requires 222.1489, observed 222.1493.

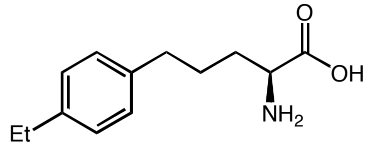

**L-3c** was prepared according to General Procedure using L-*Pf*PLP$_{Alk}$. The reactions were carried out in triplicates. **L-3c** was derived with (*S*)-Marfey's reagent and analyzed by LC-MS.

**Yield:** run 1: 59.8%; run 2: 63.2%; run 3: 61.9%; average yield: (62 ± 2)%.

**Enantioselectivity:** run 1: 96:4 e.r.; run 2: 96:4 e.r.; run 3: 96:4 e.r.; average e.r.: 96:4.

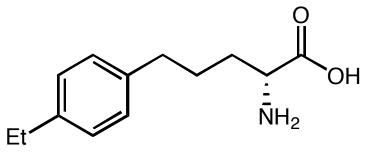

**D-3c** was prepared according to General Procedure using D-*Pf*PLP$_{Alk1}$. The reactions were carried out in triplicates. **D-3c** was derived as diastereomers using (*S*)-Marfey's reagent and analyzed by LC-MS.

**Yield:** run 1: 56.0%; run 2: 55.8%; run 3: 57.8%; average yield: (56 ± 1)%.

**Enantioselectivity:** run 1: 8:92 e.r.; run 2: 8:92 e.r.; run 3: 8:92 e.r.; average e.r.: 8:92.

**2-Amino-5-(4-methoxyphenyl)pentanoic acid (3d)**[19]

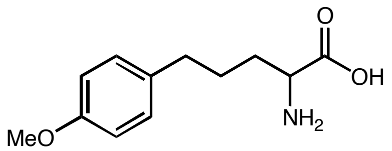

$^1$H NMR (500 MHz, $D_2O$) δ: 7.13 (d, $J$ = 8.2 Hz, 2H), 6.86 (d, $J$ = 8.2 Hz, 2H), 3.98 (t, $J$ = 6.3 Hz, 1H), 3.71 (s, 3H), 2.55 (t, $J$ = 7.4 Hz, 2H), 2.15 – 1.72 (m, 2H), 1.70 – 1.51 (m, 2H) ppm. $^{13}$C NMR (126 MHz, $D_2O$) δ: 172.0, 157.1, 134.3, 129.7, 114.1, 55.4, 52.7, 33.2, 29.0, 26.0.

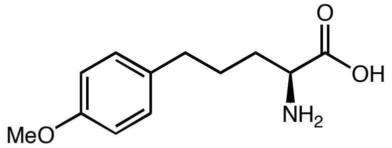

L-**3d** was prepared according to General Procedure L-*Pf*PLP$_{Alk}$. The reactions were carried out in triplicates. L-**3d** was derived with (*S*)-Marfey's reagent and analyzed by LC-MS.

**Yield:** run 1: 73.4%; run 2: 68.4%; run 3: 69.8%; average yield: (71 ± 3)%.

**Enantioselectivity:** run 1: 99:1 e.r.; run 2: 99:1e.r.; run 3: 99:1 e.r.; average e.r. 99:1.

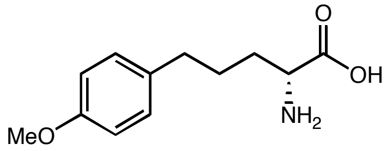

D-**3d** was prepared according to General Procedure using D-*Pf*PLP$_{Alk1}$, 10% Eosin B and 2 W power output was used. The reactions were carried out in triplicates. D-**3d** was derivatized using (*S*)-Marfey's reagent and analyzed by LC-MS.

**Yield:** run 1: 58.1%; run 2: 59.6%; run 3: 58.2%; average yield: (59 ± 1)%.

**Enantioselectivity:** run 1: 10:90 e.r.; run 2: 10:90 e.r.; run 3: 10:90 e.r.; average e.r.: 10:90.

**2-Amino-5-(4-(methylthio)phenyl)pentanoic acid (3e)**

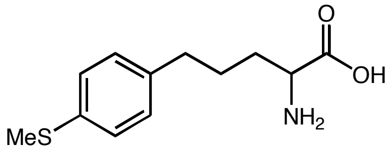

$^1$H NMR (500 MHz, $D_2O$) δ: 7.07 (d, $J$ = 8.1 Hz, 2H), 7.01 (d, $J$ = 8.1 Hz, 2H), 3.87 (t, $J$ = 6.3 Hz, 1H), 2.44 (t, $J$ = 7.7 Hz, 2H), 2.25 (s, 3H), 1.87 – 1.62 (m, 2H), 1.60 – 1.41 (m, 2H) ppm. $^{13}$C NMR (126 MHz, $D_2O$) δ: 171.8, 138.9, 134.6, 129.2, 126.8, 52.5, 33.5, 28.9, 25.7, 14.9 ppm. IR (neat) $\nu_{max}$: 2940, 2749, 1607, 1569, 1421, 1334, 804, 558, 482 cm$^{-1}$. HRMS (ESI) (m/z) for $[M+H]^+$ $C_{12}H_{18}NO_2S$ requires 240.1053, observed 240.1057.

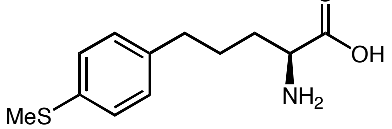

L-**3e** was prepared according to General Procedure using L-*Pf*PLP$_{Alk}$. The reactions were carried out in triplicates. L-**3e** was derived with (*S*)-Marfey's reagent and analyzed by LC-MS.

**Yield:** run 1: 33.5%; run 2: 36.8%; run 3: 35.7%; average yield: (35 ± 2)%.

**Enantioselectivity:** run 1: 97:3 e.r.; run 2: 97:3 e.r.; run 3: 97:3 e.r.; average e.r.: 97:3.

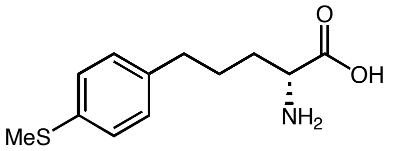

**D-3e** was prepared according to General Procedure using D-*Pf*PLP$_{Alk1}$, 10% Eosin B and 2 W power output was used. The reactions were carried out in triplicates. **D-3e** was derived as diastereomers using (*S*)-Marfey's reagent and analyzed by LC-MS.

**Yield:** run 1: 42.4%; run 2: 43.3%; run 3: 40.3%; average yield: (42 ± 2)%.

**Enantioselectivity:** run 1: 12:88 e.r.; run 2: 12:88 e.r.; run 3: 11:89 e.r.; average e.r.: 12:88.

**2-Amino-5-(4-fluorophenyl)pentanoic acid (3f)**

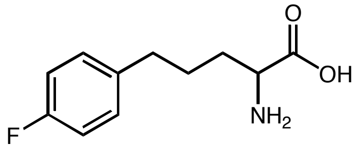

$^{1}$H NMR (500 MHz, $D_2O$) δ: 7.13 (dd, *J* = 8.3, 5.5 Hz, 2H), 6.94 (t, *J* = 8.3 Hz, 2H), 3.96 (t, *J* = 6.2 Hz, 1H), 2.67 – 2.15 (m, 2H), 1.93 – 1.72 (m, 2H), 1.71 – 1.47 (m, 2H) ppm. $^{13}$C NMR (126 MHz, $D_2O$) δ: 171.9, 161.1 (d, *J* = 240.9 Hz), 137.3 (d, *J* = 2.9 Hz), 130.0 (d, *J* = 7.9 Hz), 115.0 (d, *J* = 21.1 Hz), 52.6, 33.3, 28.9, 25.9 ppm. $^{19}$F NMR (471 MHz, $D_2O$) δ: -118.12 (tt, *J* = 9.4, 5.4 Hz). IR (neat) $\nu_{max}$: 3137, 2984, 2803, 1728, 1591, 1504, 1214, 804, 504 cm$^{-1}$. HRMS (ESI) (m/z) calculated for $C_{11}H_{15}FNO_2$ $[M+H]^+$: 212.1081, found 212.1085.

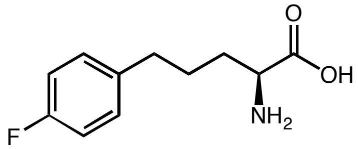

**L-3f** was prepared according to General Procedure using procedure L-*Pf*PLP$_{Alk}$. The reactions were carried out in triplicates. **L-3f** was derived with (*S*)-Marfey's reagent and analyzed by LC-MS.

**Yield:** run 1: 74.4%; run 2: 75.4%; run 3: 73.7%; average yield: (75 ± 1)%.

**Enantioselectivity:** run 1: 97:3 e.r.; run 2: 97:3 e.r.; run 3: 97:3 e.r.; average e.r.: 97:3.

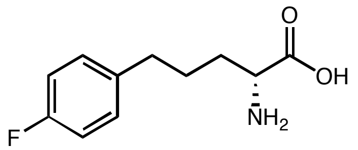

**D-3f** was prepared according to General Procedure using D-*Pf*PLP$_{Alk1}$. The reactions were carried out in triplicates. **D-3f** was derived with (*S*)-Marfey's reagent and analyzed by LC-MS.

**Yield:** run 1: 70.2%; run 2: 71.7%; run 3: 69.5%; average yield: (71 ± 1)%.

**Enantioselectivity:** run 1: 5:95 e.r.; run 2: 5:95 e.r.; run 3: 5:95 e.r.; average e.r.: 5:95.

**2-Amino-5-(4-chlorophenyl)pentanoic acid (3g)**

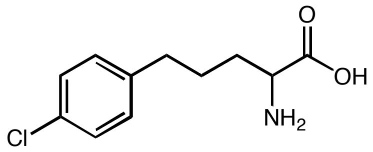

$^{1}$H NMR (500 MHz, $D_2O$) δ: 7.32 – 7.16 (m, 2H), 7.16 – 7.04 (m, 2H), 3.96 (t, $J$ = 6.2 Hz, 1H), 2.55 (td, $J$ = 7.3, 2.1 Hz, 2H), 1.90 – 1.71 (m, 2H), 1.70 – 1.45 (m, 2H) ppm. $^{13}$C NMR (126 MHz, $D_2O$) δ: 171.9, 140.2, 131.1, 130.0, 128.3, 52.6, 33.5, 28.9, 25.7 ppm. IR (neat) $\nu_{max}$: 3131, 2984, 2787, 1728, 1607, 1493, 1422, 1241, 1088, 837, 782, 657 cm$^{-1}$. HRMS (ESI) (m/z) calculated for $C_{11}H_{15}ClNO_2$ $[M+H]^+$: 228.0786, found 228.0790.

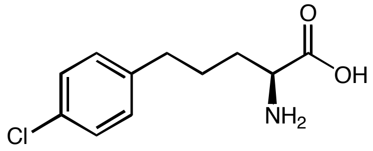

**L-3g** was prepared according to General Procedure using L-*Pf*PLP$_{Alk}$. The reactions were carried out in triplicates. **L-3g** was derived with (*S*)-Marfey's reagent and analyzed by LC-MS.

**Yield:** run 1: 75.8%; run 2: 69.5%; run 3: 71.1%; average yield: (72 ± 3)%.

**Enantioselectivity:** run 1: 97:3 e.r.; run 2: 97:3 e.r.; run 3: 97:3 e.r.; average e.r.: 97:3.

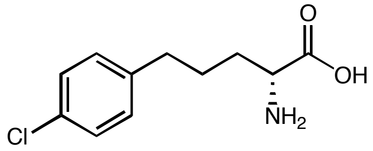

**D-3g** was prepared according to General Procedure using D-*Pf*PLP$_{Alk1}$. The reactions were carried out in triplicates. **D-3g** was derived as diastereomers using (*S*)-Marfey's reagent and analyzed by LC-MS.

**Yield:** run 1: 55.5%; run 2: 52.2%; run 3: 57.9%; average yield: (55 ± 3)%.

**Enantioselectivity:** run 1: 10:90 e.r.; run 2: 10:90 e.r.; run 3: 10:90 e.r.; average e.r.: 10:90.

**2-Amino-5-(4-bromophenyl)pentanoic acid (3h)**

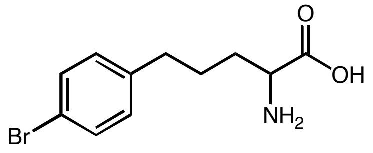

$^{1}$H NMR (400 MHz, $D_2O$) δ: 7.37 (d, $J$ = 8.0 Hz, 2H), 7.07 (d, $J$ = 8.0 Hz, 2H), 3.12 (t, $J$ = 5.4 Hz, 1H), 2.49 (t, $J$ = 7.0 Hz, 2H), 2.08 – 1.24 (m, 4H) ppm. $^{13}$C NMR (101 MHz, $D_2O$) δ: 183.6, 141.9, 131.3, 130.5, 118.8, 55.9, 34.3, 34.3, 26.8 ppm. IR (neat) $\nu_{max}$: 2989, 2913, 1733, 1498, 1225, 1006, 793, 531 cm$^{-1}$. HRMS (ESI) (m/z) for $[M+H]^+$ $C_{11}H_{15}BrNO_2$ requires 272.0281, observed 272.0286.

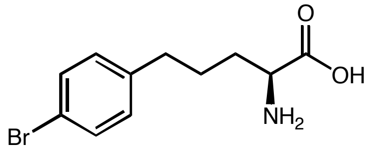

**L-3h** was prepared according to General Procedure using L-*Pf*PLP$_{Alk}$. The reactions were carried out in triplicates. **L-3h** was derived with (*S*)-Marfey's reagent and analyzed by LC-MS.

**Yield:** run 1: 46.3%; run 2: 44.9%; run 3: 43.6%; average yield: (45 ± 1)%.

**Enantioselectivity:** run 1: 97:3 e.r.; run 2: 97:3 e.r.; run 3: 97:3 e.r.; average e.r.: 97:3.

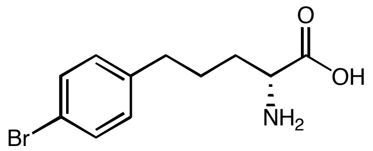

**D-3h** was prepared according to General Procedure using D-*Pf*PLP$_{Alk1}$. The reactions were carried out in triplicates. **D-3h** was derived as diastereomers using (*S*)-Marfey's reagent and analyzed by LC-MS.

**Yield:** run 1: 33.7%; run 2: 34.4%; run 3: 32.4%; average yield: (34 ± 1)%.

**Enantioselectivity:** run 1: 10: 90 e.r.; run 2: 10: 90 e.r.; run 3: 10: 90 e.r.; average e.r.: 10: 90.

**2-Amino-5-(4-cyanophenyl)pentanoic acid (3i)**

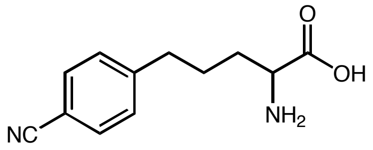

$^1$H NMR (400 MHz, $D_2O$) δ: 7.46 (d, *J* = 8.0 Hz, 2H), 7.19 (d, *J* = 8.0 Hz, 2H), 3.89 (t, *J* = 6.1 Hz, 1H), 2.55 (t, *J* = 7.3 Hz, 2H), 1.86 – 1.40 (m, 4H) ppm. $^{13}$C NMR (101 MHz, $D_2O$) δ: 171.7, 147.6, 132.4, 129.2, 119.8, 108.2, 52.5, 34.3, 28.9, 25.3 ppm. IR (neat) $\nu_{max}$: 3022, 2929, 2230, 1689, 1471, 1170, 809, 542 cm$^{-1}$. HRMS (ESI) (m/z) for $[M+H]^+$ $C_{12}H_{15}N_2O_2$ requires 219.1128, observed 219.1133.

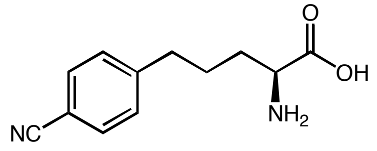

**L-3i** was prepared according to General Procedure using L-*Pf*PLP$_{Alk}$. The reactions were carried out in triplicates. **L-3i** was derived with (*S*)-Marfey's reagent and analyzed by LC-MS.

**Yield:** run 1: 57.5%; run 2: 59.5%; run 3: 61.7%; average yield: (60 ± 2)%.

**Enantioselectivity:** run 1: 99:1e.r.; run 2: 99:1 e.r.; run 3: 99:1 e.r.; average e.r.: 99:1.

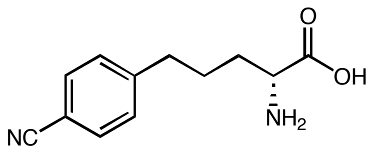

**D-3i** was prepared according to General Procedure using D-*Pf*PLP$_{Alk1}$, pH 8.5 KPi buffer, 10% Eosin B and 2 W power output was used. The reactions were carried out in triplicates. **D-3i** was derived as diastereomers using (*S*)-Marfey's reagent and analyzed by LC-MS.

**Yield:** run 1: 45.7%; run 2: 44.8%; run 3: 44.3%; average yield: (45 ± 1)%.

**Enantioselectivity:** run 1: 9:91 e.r.; run 2: 9:91 e.r.; run 3: 9:91 e.r.; average e.r.: 9:91.

**2-Amino-5-(4-(trifluoromethyl)phenyl)pentanoic acid (3j)**

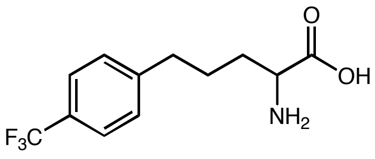

$^1H$ NMR (500 MHz, $D_2O$) δ: 7.40 (d, *J* = 8.0 Hz, 2H), 7.18 (d, *J* = 8.0 Hz, 2H), 3.86 (t, *J* = 6.2 Hz, 1H), 2.53 (t, *J* = 7.4 Hz, 2H), 2.04 – 1.37 (m, 4H) ppm. $^{13}C$ NMR (126 MHz, $D_2O$) δ: 171.7, 145.8, 128.8, 127.4 (d, *J* = 32.1 Hz), 125.2 (q, *J* = 3.7 Hz), 124.3 (d, *J* = 271.0 Hz), 52.5, 33.9, 28.8, 25.4 ppm. $^{19}F$ NMR (471 MHz, $D_2O$) δ: -62.16 ppm. IR (neat) $\nu_{max}$: 2973, 2929, 1738, 1323, 1198, 1110, 1067, 815, 531 $cm^{-1}$. HRMS (ESI) (m/z) for $[M+H]^+$ $C_{12}H_{15}F_3NO_2$ requires 262.1049, observed 262.1058.

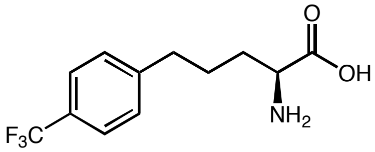

**L-3j** was prepared according to General Procedure using L-*Pf*PLP$_{Alk}$. The reactions were carried out in triplicates. **L-3j** was derived with (*S*)-Marfey's reagent and analyzed by LC-MS.

**Yield:** run 1: 13.3%; run 2: 14.3%; run 3: 16.6%; average yield: (15 ± 2)%.

**Enantioselectivity:** run 1: 94:6 e.r.; run 2: 94:6 e.r.; run 3: 93:7 e.r.; average e.r.: 94:6.

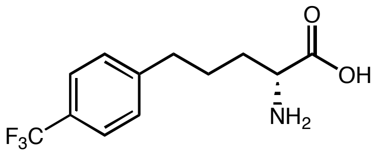

**D-3j** was prepared according to General Procedure using D-*Pf*PLP$_{Alk1}$ The reactions were carried out in triplicates. **D-3j** was derived as diastereomers using (*S*)-Marfey's reagent and analyzed by LC-MS.

**Yield:** run 1: 25.6%; run 2: 21.8%; run 3: 22.6%; average yield: (23 ± 2)%.

**Enantioselectivity:** run 1: 11:89 e.r.; run 2: 11:89 e.r.; run 3: 11:89 e.r.; average e.r.: 11:89.**2-**

**Amino-5-(m-tolyl)pentanoic acid (3k)**

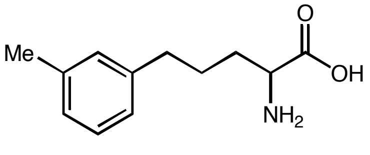

[1]H NMR (500 MHz, $D_2O$) δ: 7.18 (t, *J* = 7.5 Hz, 1H), 7.11 – 6.84 (m, 3H), 3.99 (t, *J* = 6.3 Hz, 1H), 2.58 (t, *J* = 7.4 Hz, 2H), 2.22 (s, 3H), 2.05 – 1.76 (m, 2H), 1.76 – 1.51 (m, 2H) ppm. [13]C NMR (126 MHz, $D_2O$) δ: 172.0, 141.8, 138.8, 129.2, 128.7, 126.8, 125.5, 52.7, 34.1, 29.1, 25.9, 20.3 ppm. IR (neat) $\nu_{max}$: 2990, 2914, 1730, 1491, 1220, 1182, 775, 699, 519 $cm^{-1}$. HRMS (ESI) (m/z) for $[M+H]^+$ $C_{12}H_{18}NO_2$ requires 208.1338, observed 208.1338.

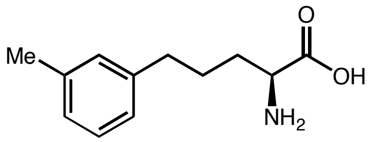

**L-3k** was prepared according to General Procedure using L-*Pf*$PLP_{Alk}$. The reactions were carried out in triplicates. **L-3k** was derived with (*S*)-Marfey's reagent and analyzed by LC-MS.

**Yield:** run 1: 59.3%; run 2: 62.3%; run 3: 61.9%; average yield: (61 ± 2)%.

**Enantioselectivity:** run 1: 88:12 e.r.; run 2: 88:12 e.r.; run 3: 88:12 e.r.; average e.r.: 88:12.

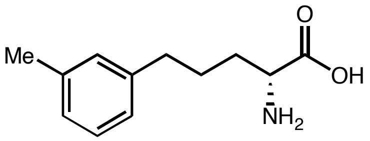

**D-3k** was prepared according to General Procedure using D-*Pf*$PLP_{Alk1}$. The reactions were carried out in triplicates. **D-3k** was derived as diastereomers using (*S*)-Marfey's reagent and analyzed by LC-MS.

**Yield:** run 1: 57.7%; run 2: 59.4%; run 3: 59.5%; average yield: (59 ± 1)%.

**Enantioselectivity:** run 1: 3:97 e.r.; run 2: 3:97 e.r.; run 3: 3:97 e.r.; average e.r.: 3:97.

**2-Amino-5-(o-tolyl)pentanoic acid (3l)**

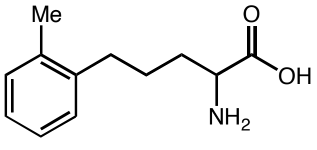

[1]H NMR (500 MHz, $D_2O$) δ: 7.36 – 6.71 (m, 4H), 4.00 (t, *J* = 6.3 Hz, 1H), 2.61 (t, *J* = 7.6 Hz, 2H), 2.21 (s, 3H), 2.06 – 1.78 (m, 2H), 1.71 – 1.36 (m, 2H) ppm. [13]C NMR (126 MHz, $D_2O$) δ: 172.0, 139.9, 136.6, 130.3, 129.1, 126.5, 126.1, 52.7, 31.7, 29.4, 24.8, 18.2. IR (neat) $\nu_{max}$: 2989, 2918, 1733, 1493 1225, 793, 739 $cm^{-1}$. HRMS (ESI) (m/z) for $[M+H]^+$ $C_{12}H_{18}NO_2$ requires 208.1338, observed 208.1338.

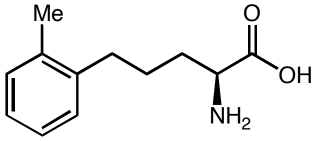

**L-3l** was prepared according to General Procedure using L-*Pf*PLP$_{Alk}$. The reactions were carried out in triplicates. **L-3l** was derived with (*S*)-Marfey's reagent and analyzed by LC-MS.

**Yield:** run 1: 37.0%; run 2: 35.3%; run 3: 34.6%; average yield: (36 ± 1)%.

**Enantioselectivity:** run 1: 62:38 e.r.; run 2: 62:38 e.r.; run 3: 62:38 e.r.; average e.r.: 62:38.

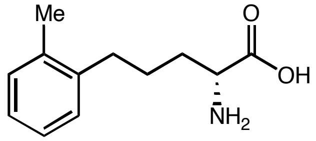

**D-3l** was prepared according to General Procedure using D-*Pf*PLP$_{Alk1}$. The reactions were carried out in triplicates. **D-3l** was derived with (*S*)-Marfey's reagent and analyzed by LC-MS.

**Yield:** run 1: 51.0%; run 2: 46.7%; run 3: 52.1%; average yield: (50 ± 3)%.

**Enantioselectivity:** run 1: 4:96 e.r.; run 2: 4:96 e.r.; run 3: 4:96 e.r.; average e.r.: 4:96.

**2-Amino-5-(furan-2-yl)pentanoic acid (3m)**

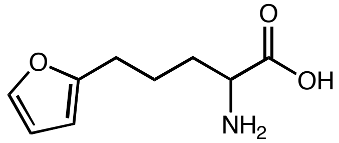

$^{1}$H NMR (500 MHz, $D_2O$) δ: 7.26 (d, $J$ = 2.0 Hz, 1H), 6.24 (dt, $J$ = 3.2, 1.5 Hz, 1H), 5.99 (d, $J$ = 3.2 Hz, 1H), 3.94 (t, $J$ = 6.3 Hz, 1H), 2.56 (t, $J$ = 7.2 Hz, 2H), 2.09 – 1.72 (m, 2H), 1.70 – 1.52 (m, 2H) ppm. $^{13}$C NMR (101 MHz, $D_2O$) δ: 171.8, 155.1, 141.5, 110.3, 105.4, 52.5, 28.9, 26.3, 22.7 ppm. IR (neat) $v_{max}$: 2989, 2924, 1607, 1574, 1493, 1389, 1279, 1077, 1017, 826, 733, 542 cm$^{-1}$. HRMS (ESI) (m/z) for $[M+H]^+$ $C_9H_{14}NO_3$ requires 184.0968, observed 184.0975.

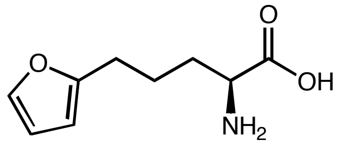

**L-3m** was prepared according to General Procedure using L-*Pf*PLP$_{Alk}$. The reactions were carried out in triplicates. **L-3m** was derived with (*S*)-Marfey's reagent and analyzed by LC-MS.

**Yield:** run 1: 55.9%; run 2: 60.4%; run 3: 53.6%; average yield: (57 ± 4)%.

**Enantioselectivity:** run 1: 98:2 e.r.; run 2: 98:2 e.r.; run 3: 98:2 e.r.; average e.r.: 98:2.

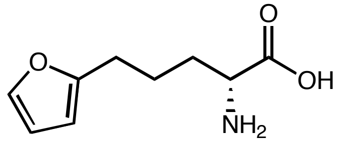

**D-3m** was prepared according to General Procedure using D-*Pf*PLP$_{Alk1}$. pH 8.5 KPi buffer, 10% Eosin B and 2 W power output was used. The reactions were carried out in triplicates. **D-3m** was derived with (*S*)-Marfey's reagent and analyzed by LC-MS.

**Yield:** run 1: 48.6%; run 2: 41.6%; run 3: 44.2%; average yield: (45 ± 4)%.

**Enantioselectivity:** run 1: 2: 98 e.r.; run 2: 2: 98 e.r.; run 3: 2: 98 e.r.; average e.r. 2: 98.

**2-Amino-5-(thiophen-2-yl)pentanoic acid (3n)**

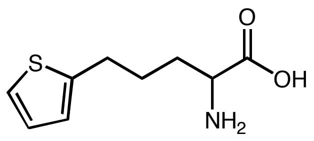

$^1$H NMR (500 MHz, $D_2O$) δ: 7.15 (dd, *J* = 5.1, 1.2 Hz, 1H), 6.87 (dd, *J* = 5.1, 3.4 Hz, 1H), 6.83 – 6.72 (m, 1H), 3.97 (t, *J* = 6.2 Hz, 1H), 2.80 (t, *J* = 7.2 Hz, 2H), 2.21 – 1.78 (m, 2H), 1.76 – 1.53 (m, 2H) ppm. $^{13}$C NMR (126 MHz, $D_2O$) δ: 171.8, 144.5, 127.1, 124.9, 123.8, 52.5, 28.8, 28.2, 26.3 ppm. IR (neat) $\nu_{max}$: 3050, 2941, 1660, 1600, 1415, 1356, 1144, 693, 552, 438 cm$^{-1}$. HRMS (ESI) (m/z) for $[M+H]^+$ $C_9H_{14}NO_2S$ requires 200.0740, observed 200.0747.

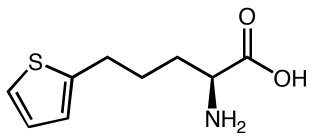

**L-3n** was prepared according to General Procedure using L-*Pf*PLP$_{Alk}$. The reactions were carried out in triplicates. **L-3n** was derived with (*S*)-Marfey's reagent and analyzed by LC-MS.

**Yield:** run 1: 72.4%; run 2: 68.5%; run 3: 78.6%; average yield: (73 ± 5)%.

**Enantioselectivity:** run 1: 97:3 e.r.; run 2: 96:4 e.r.; run 3: 96:4 e.r.; average e.r.: 96:4.

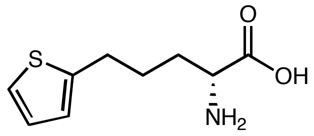

**D-3n** was prepared according to General Procedure using D-*Pf*PLP$_{Alk1}$, pH 8.5 KPi buffer, 10% Eosin B and 2 W power output was used. The reactions were carried out in triplicates. **D-3n** was derived with (*S*)-Marfey's reagent and analyzed by LC-MS.

**Yield:** run 1: 72.4%; run 2: 73.4%; run 3: 71.7%; average yield: (73 ± 1)%.

**Enantioselectivity:** run 1: 13:87 e.r.; run 2: 13:87 e.r.; run 3: 13:87 e.r.; average e.r.: 13:87.

**2-Amino-5-(6-methoxypyridin-3-yl)pentanoic acid (3o)**

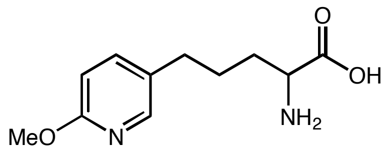

$^1$H NMR (400 MHz, $D_2O$) δ: 8.00 – 7.81 (m, 2H), 7.12 – 6.94 (m, 1H), 3.90 (s, 3H), 3.65 (t, *J* = 6.0 Hz, 1H), 2.57 (t, *J* = 7.4 Hz, 2H), 2.00 – 1.43 (m, 4H) ppm. $^{13}$C NMR (101 MHz, $D_2O$) δ: 174.4, 160.7, 145.4, 140.7, 131.5, 110.1, 56.1, 54.4, 30.3, 29.6, 25.5 ppm.

IR (neat) $\nu_{max}$: 2984, 2924, 1678, 1607, 1487, 1394, 1285, 1258, 1012, 826, 586 cm$^{-1}$. HRMS (ESI) (m/z) for $[M+H]^+$ $C_{11}H_{17}N_2O_3$ requires 225.1239, observed 225.1244.

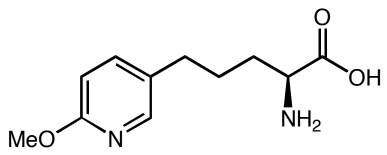

**L-3o** was prepared according to General Procedure using L-*Pf*PLP$_{Alk}$. The reactions were carried out in triplicates. **L-3o** was derived with (*S*)-Marfey's reagent and analyzed by LC-MS.

**Yield:** run 1: 48.4%; run 2: 47.9%; run 3: 48.6%; average yield: (48 ± 1)%.

**Enantioselectivity:** run 1: 99:1 e.r.; run 2: 99:1 e.r.; run 3: 99:1 e.r.; average e.r.: 99:1.

**2-Amino-6-phenylhexanoic acid (3p)**[5]

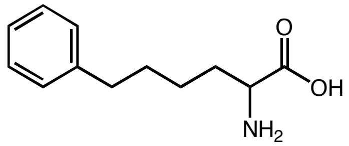

$^1$H NMR (400 MHz, $D_2O$) δ: 7.29 – 6.89 (m, 5H), 3.88 (t, $J$ = 6.3 Hz, 1H), 2.46 (t, $J$ = 7.5 Hz, 2H), 1.96 – 1.62 (m, 2H), 1.48 (p, $J$ = 7.5 Hz, 2H), 1.35 – 1.06 (m, 2H) ppm. $^{13}$C NMR (126 MHz, $D_2O$) δ: 171.9, 142.6, 128.5, 125.8, 52.7, 34.3, 29.9, 29.3, 23.4 ppm.

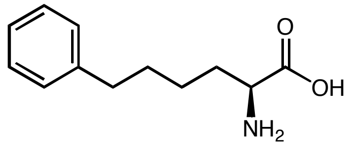

**L-3p** was prepared according to General Procedure using L-*Pf*PLP$^β$ Y301H I165T L161S Y181H H275D N166H. The reactions were carried out in triplicates. **L-3p** was derived with (*S*)-Marfey's reagent and analyzed by LC-MS.

**Yield:** run 1: 64.6%; run 2: 64.1%; run 3: 65.7%; average yield: (65 ± 1)%.

**Enantioselectivity:** run 1: 96:4 e.r.; run 2: 96:4 e.r.; run 3: 96:4 e.r.; average e.r.: 96:4.

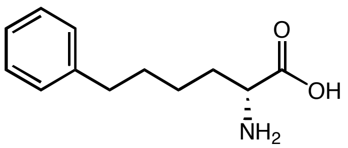

**D-3p** was prepared according to General Procedure using L-*Pf*PLP$^β$ I165G Y181W. The reactions were carried out in triplicates. **D-3p** was derived as diastereomers using (*S*)-Marfey's reagent and analyzed by LC-MS.

**Yield:** run 1: 61.2%; run 2: 63.3%; run 3: 62.1%; average yield: (62 ± 1)%.

**Enantioselectivity:** run 1: 10:90 e.r.; run 2: 9:91 e.r.; run 3: 9:91 e.r.; average e.r.: 9:91.

**2-Amino-3-(2,3-dihydro-1H-inden-2-yl)propanoic acid (3q)**

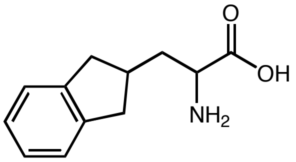

$^{1}$H NMR (500 MHz, $D_2O$) δ: 7.21 – 7.07 (m, 2H), 7.05 – 6.93 (m, 2H), 3.98 (td, $J$ = 6.9, 1.3 Hz, 1H), 3.15 (d, $J$ = 1.0 Hz, 3H), 2.95 (dq, $J$ = 14.8, 6.5 Hz, 2H), 2.70 – 2.34 (m, 3H), 2.14 – 1.96 (m, 1H), 1.94 – 1.71 (m, 1H) ppm. $^{13}$C NMR (126 MHz, $D_2O$) δ: 172.1, 142.8, 142.6, 126.4, 124.5, 124.5, 51.9, 48.8, 38.0, 37.9, 35.6, 35.4 ppm. IR (neat) $\nu_{max}$: 3290, 3181 2875, 1706, 1476, 1274, 1006, 952, 810, 744 cm$^{-1}$. HRMS (ESI) (m/z) for $[M+H]^+$ $C_{12}H_{16}NO_2$ requires 206.1176, observed 206.1183.

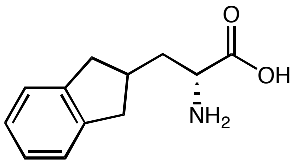

**D-3q** was prepared according to General Procedure using D-*Pf*PLP$_{Alk2}$. The reactions were carried out in triplicates. **D-3q** was derived as diastereomers using (*S*)-Marfey's reagent and analyzed by LC-MS.

**Yield:** run 1: 66.1%; run 2: 61.8%; run 3: 62.5%; average yield: (64 ± 2)%.

**Enantioselectivity:** run 1: 30:70 e.r.; run 2: 30:70 e.r.; run 3: 30:70 e.r.; average e.r.: 30:70.

**2-Aminooctanoic acid (5a)**[18]

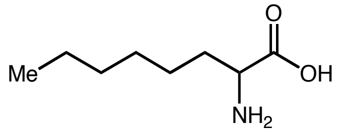

$^{1}$H NMR (400 MHz, $D_2O$) δ: 4.00 – 3.81 (m, 1H), 2.12 – 1.67 (m, 2H), 1.43 – 1.03 (m, 8H), 0.71 (t, $J$ = 6.8 Hz, 3H) ppm. $^{13}$C NMR (101 MHz, $D_2O$) δ: 174.6, 55.3, 33.0, 32.1, 30.3, 26.4, 24.2, 15.8 ppm.

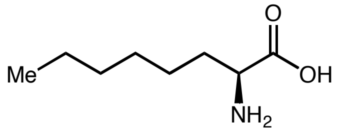

**L-5a** was prepared according to General Procedure using L-*Pf*PLP$_{Alk}$. The reactions were carried out in triplicates. **L-5a** was derived as diastereomers using (*S*)-Marfey's reagent and analyzed by LC-MS.

**Yield:** run 1: 74.3%; run 2: 74.0%; run 3: 73.5%; average yield: (74 ± 1)%.

**Enantioselectivity:** run 1: 94:6 e.r.; run 2: 94:6 e.r.; run 3: 94:6 e.r.; average e.r.: 94:6.

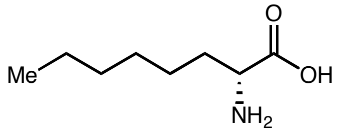

**D-5a** was prepared according to General Procedure using D-*Pf*PLP$_{Alk2}$. The reactions were carried out in triplicates. **D-5a** was derived as diastereomers using (*S*)-Marfey's reagent and analyzed by LC-MS.

**Yield:** run 1: 74.9%; run 2: 75.4%; run 3: 74.8%; average yield: (75 ± 1)%.

**Enantioselectivity:** run 1: 3:97 e.r.; run 2: 3:97 e.r.; run 3: 3:97 e.r.; average e.r.: 3:97.

**2-Amino-7-fluoroheptanoic acid (5b)**[20]

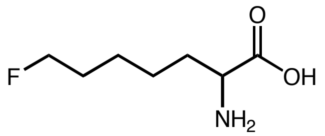

$^1$H NMR (400 MHz, $D_2O$) δ: 4.49 (td, *J* = 6.1, 1.1 Hz, 1H), 4.37 (td, *J* = 6.1, 1.1 Hz, 1H), 3.94 (t, *J* = 6.3 Hz, 1H), 1.94 – 1.77 (m, 2H), 1.76 – 1.53 (m, 2H), 1.48 – 1.28 (m, 4H) ppm. $^{13}$C NMR (101 MHz, $D_2O$) δ: 172.6, 85.3 (d, *J* = 157.2 Hz), 53.1, 29.7, 29.1 (d, *J* = 19.0 Hz), 24.0 (d, *J* = 5.6 Hz), 23.8. $^{19}$F NMR (376 MHz, $D_2O$) δ: -216.62.

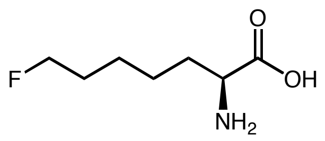

**L-5b** was prepared according to General Procedure using L-*Pf*PLP$_{Alk}$. The reactions were carried out in triplicates. **L-5b** was derived as diastereomers using (*S*)-Marfey's reagent and analyzed by LC-MS.

**Yield:** run 1: 59.8%; run 2: 62.9%; run 3: 61.0%; average yield: (61 ± 2)%.

**Enantioselectivity:** run 1: 98:2 e.r.; run 2: 98:2 e.r.; run 3: 98:2 e.r.; average e.r.: 98:2.

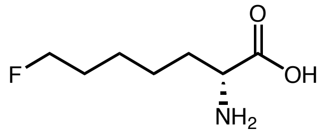

**D-5b** was prepared according to General Procedure using D-*Pf*PLP$_{Alk2}$. The reactions were carried out in triplicates. **D-5b** was derived as diastereomers using (*S*)-Marfey's reagent and analyzed by LC-MS.

**Yield:** run 1: 58.9%; run 2: 54.8%; run 3: 54.7%; average yield: (56 ± 2)%.

**Enantioselectivity:** run 1: 5:95 e.r.; run 2: 5:95 e.r.; run 3: 5:95 e.r.; average e.r.: 5:95.

**2-Amino-8-chlorooctanoic acid (5c)**

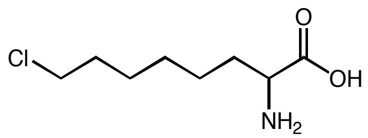

$^1$H NMR (400 MHz, $D_2O$) δ: 3.94 (t, *J* = 6.3 Hz, 1H), 3.46 (t, *J* = 6.7 Hz, 2H), 2.06 – 1.71 (m, 2H), 1.68 – 1.53 (m, 2H), 1.47 – 1.06 (m, 6H) ppm. $^{13}$C NMR (101 MHz, $D_2O$) δ: 172.1, 52.7, 45.6, 31.6, 29.5, 27.3, 25.5, 23.8 ppm. IR (neat) $\nu_{max}$: 3022, 2924, 2853, 1656, 1580, 1509, 1411, 1345, 722, 651, 564 cm$^{-1}$. HRMS (ESI) (m/z) for $[M+H]^+$ $C_8H_{17}ClNO_2$ requires 194.0948, observed 194.0950.

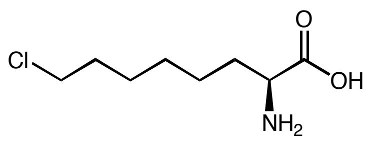

**L-5c** was prepared according to General Procedure using L-*Pf*PLP$_{Alk}$. The reactions were carried out in triplicates. **L-5c** was derived as diastereomers using (*S*)-Marfey's reagent and analyzed by LC-MS.

**Yield:** run 1: 77.5%; run 2: 78.9%; run 3: 79.1%; average yield: (78 ± 1)%.

**Enantioselectivity:** run 1: 94:6 e.r.; run 2: 94:6 e.r.; run 3: 94:6 e.r.; average e.r.: 94:6.

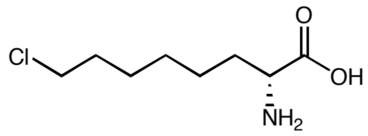

D-**5c** was prepared according to General Procedure using D-*Pf*PLP$_{Alk2}$. The reactions were carried out in triplicates. D-**5c** was derived as diastereomers using (*S*)-Marfey's reagent and analyzed by LC-MS.

**Yield:** run 1: 39.6%; run 2: 38.7%; run 3: 41.1%; average yield: (40 ± 1)%.

**Enantioselectivity:** run 1: 6:94 e.r.; run 2: 6:94 e.r.; run 3: 6:94 e.r.; average e.r.: 6:94.

**2-Amino-7-methoxyheptanoic acid (5d)**

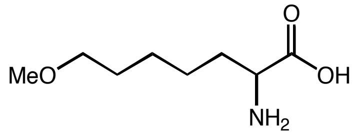

$^1$H NMR (400 MHz, $D_2O$) δ: 3.86 (t, *J* = 6.2 Hz, 1H), 3.38 (t, *J* = 6.6 Hz, 2H), 3.23 (s, 3H), 1.89 – 1.71 (m, 2H), 1.60 – 1.45 (m, 2H), 1.43 – 1.15 (m, 4H) ppm. $^{13}$C NMR (101 MHz, $D_2O$) δ: 173.1, 72.4, 57.6, 53.5, 29.8, 28.0, 24.8, 23.9 ppm. IR (neat) $\nu_{max}$: 3164, 2891, 1733, 1602, 1487, 1421, 1214, 1121, 903, 848, 640 cm$^{-1}$. HRMS (ESI) (m/z) for $[M+H]^+$ $C_8H_{18}NO_3$ requires 176.1287, observed 176.1287.

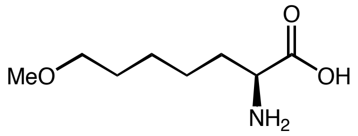

L-**5d** was prepared according to General Procedure using L-*Pf*PLP$_{Alk}$. The reactions were carried out in triplicates. L-**5d** was derived as diastereomers using (*S*)-Marfey's reagent and analyzed by LC-MS.

**Yield:** run 1: 55.4%; run 2: 55.5%; run 3: 51.3%; average yield: (54 ± 2)%.

**Enantioselectivity:** run 1: 97:3 e.r.; run 2: 97:3 e.r.; run 3: 97:3 e.r.; average e.r.: 97:3.

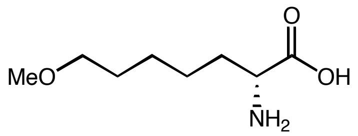

D-**5d** was prepared according to General Procedure using D-*Pf*PLP$_{Alk2}$. The reactions were carried out in triplicates. D-**5d** was derived as diastereomers using (*S*)-Marfey's reagent and analyzed by LC-MS.

**Yield:** run 1: 39.6%; run 2: 38.7%; run 3: 41.1%; average yield: (40 ± 1)%.

**Enantioselectivity:** run 1: 3:97 e.r.; run 2: 4:96 e.r.; run 3: 4:96 e.r.; average e.r.: 4:96.

**2-Amino-7-(methylthio)heptanoic acid (5e)**

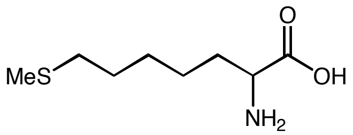

$^{1}$H NMR (400 MHz, $D_2O$) δ: 3.95 (t, *J* = 6.3 Hz, 1H), 2.39 (t, *J* = 7.3 Hz, 2H), 1.94 (s, 3H), 1.88 – 1.60 (m, 2H), 1.68 – 1.40 (m, 2H), 1.42 – 1.16 (m, 4H) ppm. $^{13}$C NMR (101 MHz, $D_2O$) δ: 172.1, 52.7, 33.0, 29.5, 27.7, 27.2, 23.6, 14.1 ppm. IR (neat) $\nu_{max}$: 3000, 2913, 2853, 1728, 1487, 1422, 1219, 1138, 837, 799, 722, 537 cm$^{-1}$. HRMS (ESI) (m/z) for $[M+H]^+$ $C_8H_{18}NO_2S$ requires 192.1058, observed 192.1060.

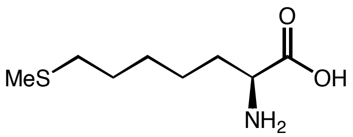

**L-5e** was prepared according to General Procedure using L-*Pf*PLP$_{Alk}$. The reactions were carried out in triplicates. **L-5e** was derived as diastereomers using (*S*)-Marfey's reagent and analyzed by LC-MS.

**Yield:** run 1: 55.1%; run 2: 53.1%; run 3: 50.2%; average yield: (53 ± 3)%.

**Enantioselectivity:** run 1: 96:4 e.r.; run 2: 96:4 e.r.; run 3: 96:4 e.r.; average e.r.: 96:4.

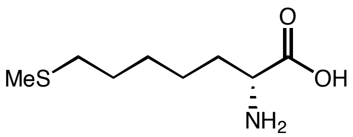

**D-5e** was prepared according to General Procedure using D-*Pf*PLP$_{Alk2}$. The reactions were carried out in triplicates. **D-5e** was derived as diastereomers using (*S*)-Marfey's reagent and analyzed by LC-MS.

**Yield:** run 1: 34.1%; run 2: 33.0%; run 3: 32.0%; average yield: (33 ± 1)%.

**Enantioselectivity:** run 1: 7:93 e.r.; run 2: 7:93 e.r.; run 3: 7:93 e.r.; average e.r.: 7:93.

**2-Amino-8-azidooctanoic acid (5f)**

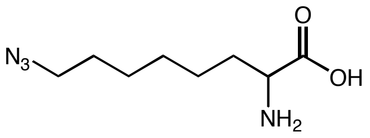

$^{1}$H NMR (400 MHz, $D_2O$) δ: 3.97 (t, *J* = 6.3 Hz, 1H), 3.19 (t, *J* = 6.8 Hz, 2H), 1.94 – 1.69 (m, 2H), 1.63 – 1.42 (m, 2H), 1.41 – 1.16 (m, 6H) ppm. $^{13}$C NMR (101 MHz, $D_2O$) δ: 172.1, 52.8, 51.1, 29.5, 27.8, 27.6, 25.5, 23.9 ppm. IR (neat) $\nu_{max}$: 3049, 2929, 2864, 2099, 1678, 1509, 1482, 1263, 1127, 728, 580 cm$^{-1}$. HRMS (ESI) (m/z) for $[M+H]^+$ $C_8H_{17}N_4O_2$ requires 201.1352, observed 201.1355.

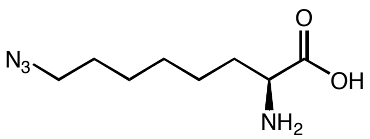

**L-5f** was prepared according to General Procedure using L-*Pf*PLP$_{Alk}$. The reactions were carried out in triplicates. **L-5f** was derived as diastereomers using (*S*)-Marfey's reagent and analyzed by LC-MS.

**Yield:** run 1: 53.4%; run 2: 52.1%; run 3: 53.9%; average yield: (53 ± 1)%.

**Enantioselectivity:** run 1: 95:5 e.r.; run 2: 95:5 e.r.; run 3: 95:5 e.r.; average e.r.: 95:5.

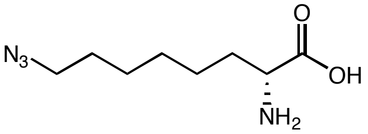

**D-5f** was prepared according to General Procedure using D-*Pf*PLP$_{\text{Alk2}}$. The reactions were carried out in triplicates. **D-5f** was derived as diastereomers using (*S*)-Marfey's reagent and analyzed by LC-MS.

**Yield:** run 1: 30.4%; run 2: 30.8%; run 3: 29.9%; average yield: (30 ± 1)%.

**Enantioselectivity:** run 1: 9:91 e.r.; run 2: 9:91 e.r.; run 3: 9:91 e.r.; average e.r.: 9:91.

**2-Aminooct-7-ynoic acid (5g)**[21]

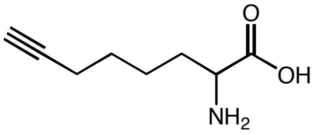

$^{1}$H NMR (400 MHz, $D_2O$) δ: 3.98 (t, *J* = 6.4 Hz, 1H), 2.34 – 2.20 (m, 1H), 2.19 – 2.06 (m, 2H), 1.98 – 1.69 (m, 2H), 1.54 – 1.29 (m, 4H) ppm. $^{13}$C NMR (101 MHz, $D_2O$) δ: 172.0, 85.5, 69.4, 52.7, 29.1, 26.9, 23.2, 17.1 ppm.

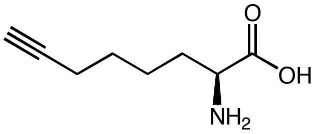

**L-5g** was prepared according to General Procedure using L-*Pf*PLP$_{\text{Alk}}$. The reactions were carried out in triplicates. **L-5g** was derived as diastereomers using (*S*)-Marfey's reagent and analyzed by LC-MS.

**Yield:** run 1: 64.8%; run 2: 61.5%; run 3: 63.3%; average yield: (63 ± 2)%.

**Enantioselectivity:** run 1: 98:2 e.r.; run 2: 98:2 e.r.; run 3: 98:2 e.r.; average e.r.: 98:2.

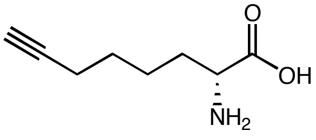

**D-5g** was prepared according to General Procedure using D-*Pf*PLP$_{\text{Alk2}}$. The reactions were carried out in triplicates. **D-5g** was derived as diastereomers using (*S*)-Marfey's reagent and analyzed by LC-MS.

**Yield:** run 1: 65.3%; run 2: 63.9%; run 3: 64.4%; average yield: (65 ± 1)%.

**Enantioselectivity:** run 1: 2:98 e.r.; run 2: 2:98 e.r.; run 3: 2:98 e.r.; average e.r.: 2:98.

**2-Aminohept-6-enoic acid (5h)**[22]

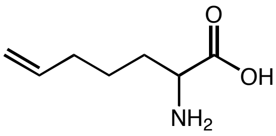

$^{1}$H NMR (400 MHz, $D_2O$) δ: 5.97 – 5.64 (m, 1H), 5.32 – 4.87 (m, 2H), 3.15 (t, *J* = 6.2 Hz, 1H), 2.00 (q, *J* = 6.6 Hz, 2H), 1.74 – 1.41 (m, 2H), 1.33

(p, $J$ = 7.7 Hz, 2H) ppm. $^{13}$C NMR (101 MHz, $D_2O$) δ: 183.9, 139.7, 114.5, 55.9, 34.2, 33.0, 24.4 ppm.

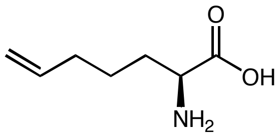

**L-5h** was prepared according to General Procedure using L-*Pf*PLP$_{Alk}$. The reactions were carried out in triplicates. **L-5h** was derived as diastereomers using (*S*)-Marfey's reagent and analyzed by LC-MS.

**Yield:** run 1: 62.9%; run 2: 62.6%; run 3: 59.1%; average yield: (62 ± 2)%.

**Enantioselectivity:** run 1: 98:2 e.r.; run 2: 98:2 e.r.; run 3: 98:2 e.r.; average e.r.: 98:2.

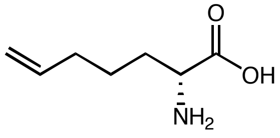

**D-5m** was prepared according to General Procedure using D-*Pf*PLP$_{Alk2}$. The reactions were carried out in triplicates. **D-5h** was derived as diastereomers using (*S*)-Marfey's reagent and analyzed by LC-MS.

**Yield:** run 1: 67.0%; run 2: 68.5%; run 3: 67.9%; average yield: (68 ± 1)%.

**Enantioselectivity:** run 1: 5:95 e.r.; run 2: 6:94 e.r.; run 3: 6:94 e.r.; average e.r.: 6:94.

**2-Amino-3-cyclopentylpropanoic acid (5i)**[5]

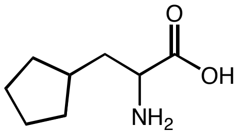

$^{1}$H NMR (500 MHz, MeOD) δ: 3.91 (dd, $J$ = 7.4, 5.9 Hz, 1H), 2.09 – 1.94 (m, 2H), 1.93 – 1.81 (m, 3H), 1.76 – 1.65 (m, 2H), 1.65 – 1.55 (m, 2H), 1.28 – 1.11 (m, 2H) ppm. $^{13}$C NMR (126 MHz, MeOD) δ: 172.2, 53.5, 37.9, 37.3, 33.4, 33.3, 26.0, 25.8 ppm.

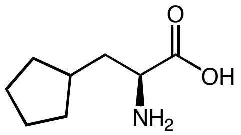

**L-5i** was prepared according to General Procedure using L-*Pf*PLP$_{Alk}$. The reactions were carried out in triplicates. **L-5i** was derived as diastereomers using (*S*)-Marfey's reagent and analyzed by LC-MS.

**Yield:** run 1: 67.9%; run 2: 65.6%; run 3: 73.1%; average yield: (69 ± 4)%.

**Enantioselectivity:** run 1: 97:3 e.r.; run 2: 97:3 e.r.; run 3: 97:3 e.r.; average e.r.: 97:3.

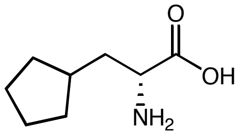

**D-5i** was prepared according to General Procedure using D-*Pf*PLP$_{Alk2}$. The reactions were carried out in triplicates. **D-5i** was derived as diastereomers using (*S*)-Marfey's reagent and analyzed by LC-MS.

**Yield:** run 1: 53.0%; run 2: 54.7%; run 3: 54.1%; average yield: (54 ± 1)%.

**Enantioselectivity:** run 1: 26:74 e.r.; run 2: 26:74 e.r.; run 3: 26:74 e.r.; average e.r.: 26:74.

**2-Aminopentanoic acid (5k)**[23]

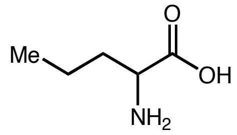

$^1$H NMR (400 MHz, $D_2O$) δ: 3.93 (t, *J* = 6.3 Hz, 1H), 2.26 – 1.69 (m, 2H), 1.59 – 1.27 (m, 2H), 0.93 (t, *J* = 7.3 Hz, 3H) ppm. $^{13}$C NMR (101 MHz, $D_2O$) δ: 173.4, 53.5, 32.0, 17.7, 12.8 ppm.

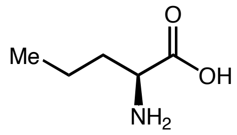

**L-5k** was prepared according to General Procedure using L-*Pf*PLP$_{Alk}$. The reactions were carried out in triplicates. **L-5k** was derived as diastereomers using (*S*)-Marfey's reagent and analyzed by LC-MS.

**Yield:** run 1: 66.0%; run 2: 60.2%; run 3: 59.0%; average yield: (62 ± 4)%.

**Enantioselectivity:** run 1: 99:1 e.r.; run 2: 99:1 e.r.; run 3: 99:1 e.r.; average e.r.: 99:1.

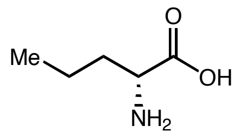

**D-5k** was prepared according to General Procedure using D-*Pf*PLP$_{Alk2}$. The reactions were carried out in triplicates. **D-5k** was derived as diastereomers using (*S*)-Marfey's reagent and analyzed by LC-MS.

**Yield:** run 1: 77.1%; run 2: 76.3%; run 3: 76.6%; average yield: (77 ± 1)%.

**Enantioselectivity:** run 1: 7:93 e.r.; run 2: 7:93 e.r.; run 3: 7:93 e.r.; average e.r.: 7:93.

**2-Aminohexanoic acid (5l)**[24]

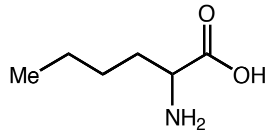

$^1$H NMR (400 MHz, $D_2O$) δ: 4.05 (t, *J* = 6.3 Hz, 1H), 2.32 – 1.79 (m, 2H), 1.60 – 1.27 (m, 4H), 0.89 (t, *J* = 7.0 Hz, 3H) ppm. $^{13}$C NMR (101 MHz, $D_2O$) δ: 172.5, 53.0, 29.4, 26.2, 21.5,12.9 ppm.

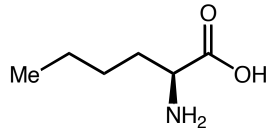

**L-5l** was prepared according to General Procedure using L-*Pf*PLP$_{Alk}$. The reactions were carried out in triplicates. **L-5l** was derived as diastereomers using (*S*)-Marfey's reagent and analyzed by LC-MS.

**Yield:** run 1: 55.3%; run 2: 53.4%; run 3: 54.8%; average yield: (55 ± 1)%.

**Enantioselectivity:** run 1: 99:1 e.r.; run 2: 99:1 e.r.; run 3: 99:1 e.r.; average e.r.: 99:1.

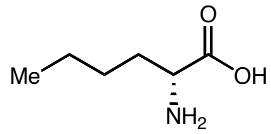

**D-5l** was prepared according to General Procedure using D-*Pf*PLP$_{Alk2}$. The reactions were carried out in triplicates. **D-5l** was derived as diastereomers using (*S*)-Marfey's reagent and analyzed by LC-MS.

**Yield:** run 1: 74.2%; run 2: 69.1%; run 3: 76.6%; average yield: (73 ± 4)%.

**Enantioselectivity:** run 1: 4:96 e.r.; run 2: 4:96 e.r.; run 3: 4:96 e.r.; average e.r.: 4:96.

**2-Aminoheptanoic acid (5m)**[18]

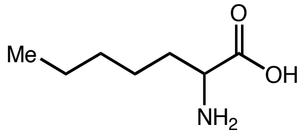

$^{1}$H NMR (400 MHz, $D_2O$) δ: 4.06 (t, $J$ = 6.3 Hz, 1H), 2.19 – 1.81 (m, 2H), 1.63 – 1.11 (m, 7H), 0.87 (t, $J$ = 7.0 Hz, 3H) ppm. $^{13}$C NMR (101 MHz, $D_2O$) δ: 3.24 (t, $J$ = 6.4 Hz, 1H), 1.77 – 1.46 (m, 2H), 1.31 (d, $J$ = 105.3 Hz, 11H), 0.89 (t, $J$ = 6.6 Hz, 3H) ppm. $^{13}$C NMR (101 MHz, $CDCl_3$) δ: 175.0, 55.6, 32.9, 32.2, 26.2, 24.1, 15.7.

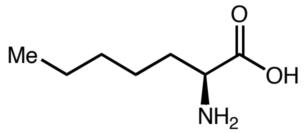

**L-5m** was prepared according to General Procedure using L-*Pf*PLP$_{Alk}$. The reactions were carried out in triplicates. **L-5m** was derived as diastereomers using (*S*)-Marfey's reagent and analyzed by LC-MS.

**Yield:** run 1: 63.2%; run 2: 63.6%; run 3: 57.6%; average yield: (62 ± 3)%.

**Enantioselectivity:** run 1: 98:2 e.r.; run 2: 98:2 e.r.; run 3: 97.5:2.5 e.r.; average e.r.: 98:2.

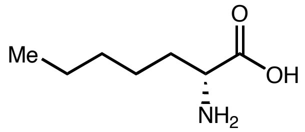

**D-5m** was prepared according to General Procedure using D-*Pf*PLP$_{Alk2}$. The reactions were carried out in triplicates. **D-5m** was derived as diastereomers using (*S*)-Marfey's reagent and analyzed by LC-MS.

**Yield:** run 1: 73.0%; run 2: 74.7%; run 3: 71.6%; average yield: (73 ± 2)%.

**Enantioselectivity:** run 1: 2:98 e.r.; run 2: 2.5:97.5 e.r.; run 3: 2.5:97.5 e.r.; average e.r.: 2:98.

**2-Aminononanoic acid (5n)**[18]

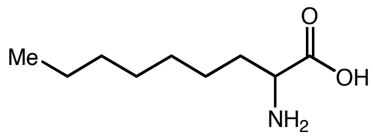

$^{1}$H NMR (400 MHz, $D_2O$) δ: 3.24 (t, $J$ = 6.4 Hz, 1H), 1.77 – 1.46 (m, 2H), 1.41 – 1.21 (m, 10H), 0.89 (t, $J$ = 6.6 Hz, 3H) ppm. $^{13}$C NMR (101 MHz, $D_2O$) δ: 186.6, 58.6, 37.3, 33.6, 31.3, 30.9, 27.5, 24.6, 16.0 ppm.

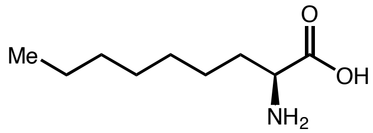

**L-5n** was prepared according to General Procedure using L-*Pf*PLP$_{Alk}$. The reactions were carried out in triplicates. **L-5n** was derived as diastereomers using (*S*)-Marfey's reagent and analyzed by LC-MS.

**Yield:** run 1: 77.7%; run 2: 79.4%; run 3: 77.7%; average yield: (78 ± 1)%.

**Enantioselectivity:** run 1: 88:12 e.r.; run 2: 88:12 e.r.; run 3: 88:12 e.r.; average e.r.: 88:12.

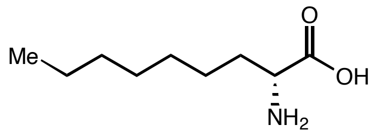

**D-5n** was prepared according to General Procedure using D-*Pf*PLP$_{Alk2}$. The reactions were carried out in triplicates. **D-5n** was derived as diastereomers using (*S*)-Marfey's reagent and analyzed by LC-MS.

**Yield:** run 1: 59.7%; run 2: 60.7%; run 3: 60.3%; average yield: (60 ± 1)%.

**Enantioselectivity:** run 1: 8:92 e.r.; run 2: 8:92 e.r.; run 3: 8:92 e.r.; average e.r.: 8:92.

**2-amino-4-methyloctanoic acid (5n')**

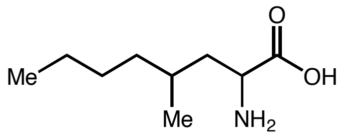

**5s** was isolated as a mixture of two isomers. $^{1}$H NMR (400 MHz, $D_2O$) δ: 3.33 – 2.97 (m, 1H), 1.75 – 0.95 (m, 9H), 0.90 – 0.67 (m, 6H) ppm. $^{13}$C NMR (101 MHz, $D_2O$) for major diastereomer δ: 184.4, 54.6, 42.7, 35.6, 29.1, 28.3, 22.3, 19.4, 13.4 ppm; for major diastereomer δ: 184.6, 54.3, 42.5, 36.5, 28.9, 28.4, 22.3, 18.5, 13.4 ppm. IR (neat) $\nu_{max}$: 3421, 3077, 2957, 2924, 1585, 1514, 1411, 1307, 1127, 837, 679, 564 cm$^{-1}$. HRMS (ESI) (m/z) for $[M+H]^+$ $C_9H_{20}NO_2$ requires 174.1494, observed 174.1496.

**2-Aminoundecanoic acid (5o)**[18]

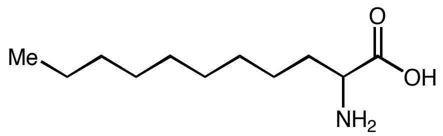

$^{1}$H NMR (400 MHz, $D_2O$) δ: 3.17 (dd, *J* = 7.6, 5.3 Hz, 1H), 2.03 – 1.52 (m, 1H), 1.54 – 1.38 (m, 1H), 1.38 – 1.16 (m, 14H), 0.86 (t, *J* = 6.4 Hz, 3H) ppm. $^{13}$C NMR (101 MHz, $D_2O$) δ: 183.0, 56.3, 35.7, 31.8, 29.6, 29.5, 29.2, 25.8, 22.5, 13.7 ppm.

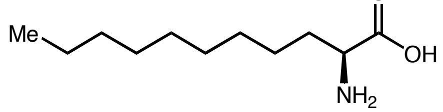

**L-5o** was prepared according to General Procedure using L-*Pf*PLP$_{Alk}$. The reactions were carried out in triplicates. **L-5o** was derived as diastereomers using (*S*)-Marfey's reagent and analyzed by LC-MS.

**Yield:** run 1: 67.6%; run 2: 65.7%; run 3: 66.7%; average yield: (67 ± 1)%.

**Enantioselectivity:** run 1: 84.5:15.5 e.r.; run 2: 84.5:15.5 e.r.; run 3: 84.5:15.5 e.r.; average e.r.: 84.5:14.5.

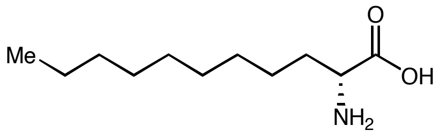

**D-5o** was prepared according to General Procedure using D-*Pf*PLP$_{\text{Alk2}}$. The reactions were carried out in triplicates. **D-5f** was derived as diastereomers using (*S*)-Marfey's reagent and analyzed by LC-MS.

**Yield:** run 1: 47.9%; run 2: 48.8%; run 3: 50.9%; average yield: (49 ± 2)%.

**Enantioselectivity:** run 1: 28:72 e.r.; run 2: 28:72 e.r.; run 3: 28:72 e.r.; average e.r.: 28:72.

**2-Aminotridecanoic acid (5p)**[25]

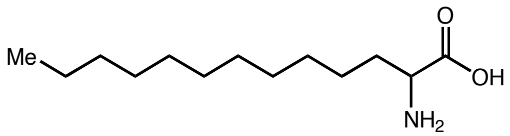

$^{1}$H NMR (400 MHz, $D_2O$) δ: 3.05 (dd, *J* = 7.8, 5.1 Hz, 1H), 1.76 – 1.45 (m, 1H), 1.40 – 1.26 (m, 1H), 1.23 – 1.09 (m, 18H), 0.74 (t, *J* = 6.2 Hz, 3H) ppm. $^{13}$C NMR (101 MHz, $D_2O$) δ: 182.7, 56.4, 36.0, 32.0, 30.0, 29.9, 29.9, 29.8, 29.8, 29.5, 26.0, 22.6, 13.8 ppm.

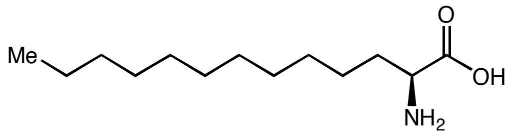

**L-5p** was prepared according to General Procedure using L-*Pf*PLP$_{\text{Alk}}$ and additional DMSO was added to (15% v/v). The reactions were carried out in triplicates. **L-5p** was derived as diastereomers using (*S*)-Marfey's reagent and analyzed by LC-MS.

**Yield:** run 1: 43.4%; run 2: 43.4%; run 3: 37.0%; average yield: (41 ± 4)%.

**Enantioselectivity:** run 1: 68:32 e.r.; run 2: 68:32 e.r.; run 3: 68:32 e.r.; average e.r.: 68:32.

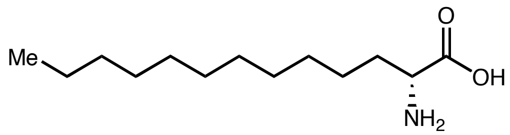

**D-5p** was prepared according to General Procedure using D-*Pf*PLP$_{\text{Alk2}}$ and additional DMSO was added to (15% v/v). The reactions were carried out in triplicates. **D-5p** was derived as diastereomers using (*S*)-Marfey's reagent and analyzed by LC-MS.

**Yield:** run 1: 42.8%; run 2: 41.0%; run 3: 38.6%; average yield: (41 ± 2)%.

**Enantioselectivity:** run 1: 18:82 e.r.; run 2: 18:82 e.r.; run 3: 18:82 e.r.; average e.r.: 18:82.

**2-Amino-4-cyclopropylbutanoic acid (5q)**

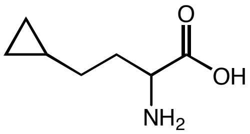

$^{1}$H NMR (400 MHz, $D_2O$) δ: 3.97 (t, $J$ = 6.2 Hz, 1H), 2.80 – 1.80 (m, 2H), 1.52 – 1.23 (m, 2H), 0.88 – 0.62 (m, 1H), 0.51 – 0.42 (m, 2H), 0.09 (dd, $J$ = 5.0, 1.5 Hz, 2H) ppm. $^{13}$C NMR (101 MHz, $D_2O$) δ: 172.1, 52.6, 29.8, 29.1, 9.5, 3.7, 3.6 ppm. IR (neat) $\nu_{max}$: 3383, 2910, 1734, 1628, 1584, 1500, 1212, 1140, 1049, 899, 593 cm$^{-1}$. HRMS (ESI) (m/z) for $[M+H]^+$ $C_7H_{14}NO_2$ requires 144.1019, observed 144.1025.

**2-Amino-4-cyclopentylbutanoic acid (5r)**

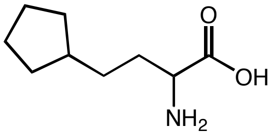

$^{1}$H NMR (400 MHz, $D_2O$) δ: 3.18 (t, $J$ = 6.3 Hz, 1H), 1.82 – 1.65 (m, 3H), 1.63 – 1.39 (m, 6H), 1.34 – 1.20 (m, 2H), 1.15 – 0.98 (m, 2H) ppm. $^{13}$C NMR (101 MHz, $D_2O$) δ: 184.0, 56.2, 39.4, 33.9, 32.1, 32.1, 31.5, 24.7 ppm. IR (neat) $\nu_{max}$: 2951, 2864, 1706, 1602, 1520, 1482, 1263, 1138, 558 cm$^{-1}$. HRMS (ESI) (m/z) for $[M+H]^+$ $C_9H_{18}NO_2$ requires 172.1338, observed 172.1341.

**2-Aminonon-8-enoic acid (5s)**[26]

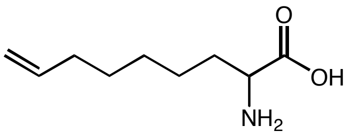

$^{1}$H NMR (400 MHz, $D_2O$) δ: 6.01 – 5.75 (m, 1H), 4.98 (dd, $J$ = 17.3, 10.1 Hz, 2H), 3.19 (t, $J$ = 6.7 Hz, 1H), 2.03 (q, $J$ = 6.5 Hz, 2H), 1.57 – 1.52 (m, 2H), 1.42 – 1.15 (m, 6H) ppm. $^{13}$C NMR (101 MHz, $D_2O$) δ: 184.0, 140.3, 114.0, 56.0, 34.6, 33.0, 28.2, 27.9, 24.7 ppm.

#### VIII. HPLC calibration curves of amino acid products

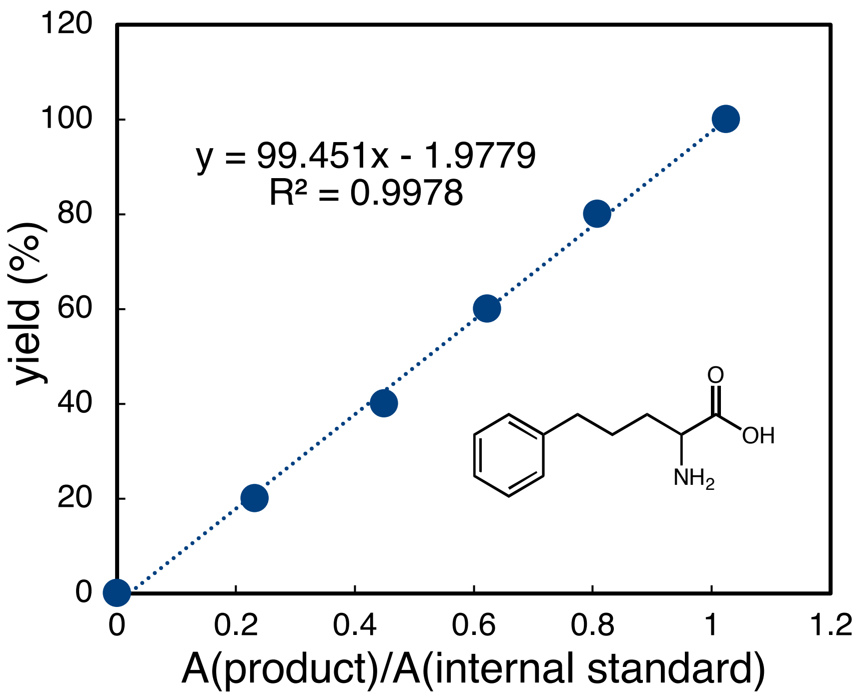

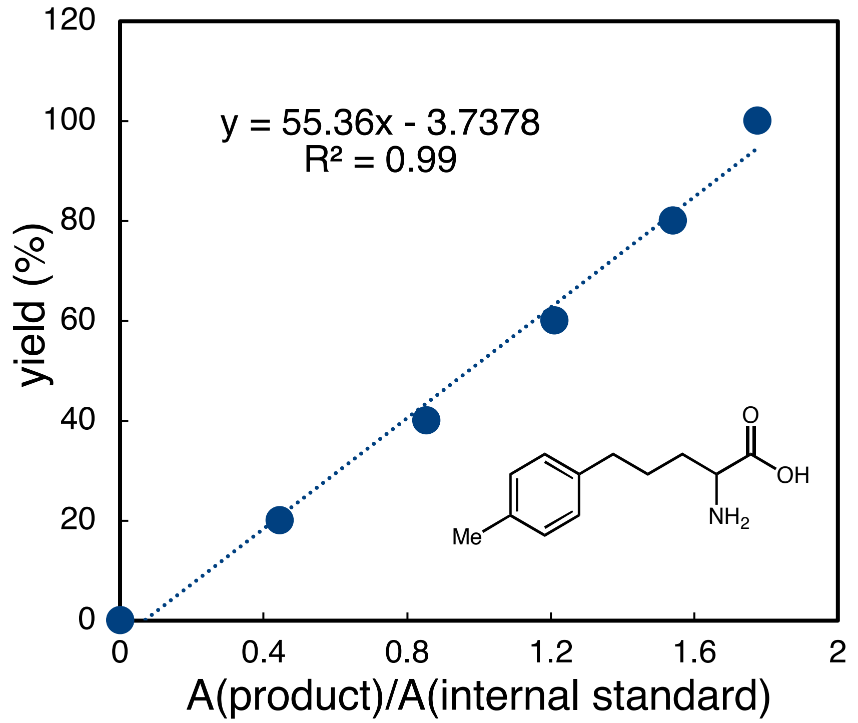

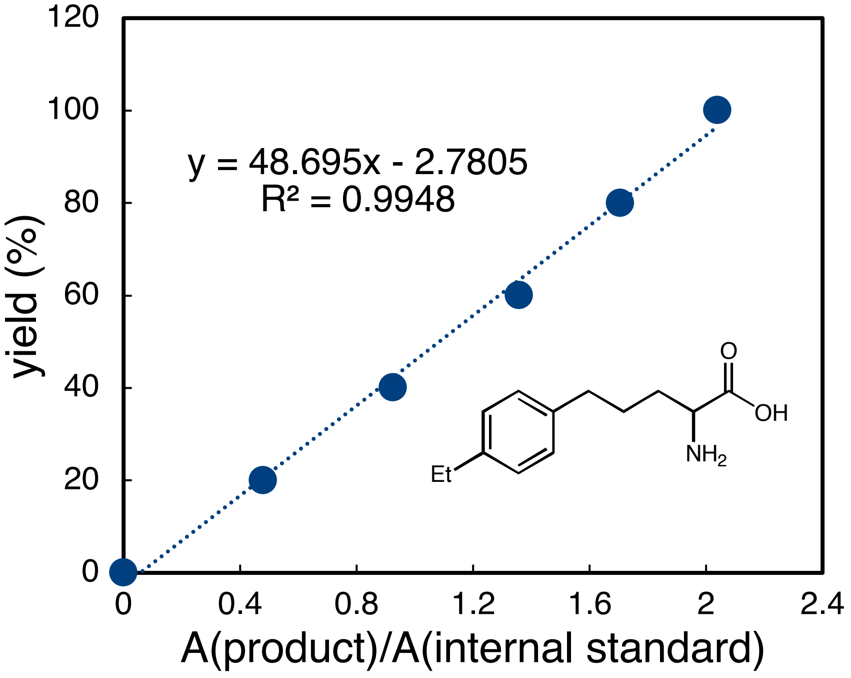

y = 48.695x - 2.7805
R² = 0.9948
yield (%)
A(product)/A(internal standard)
Et
NH2
O
OH

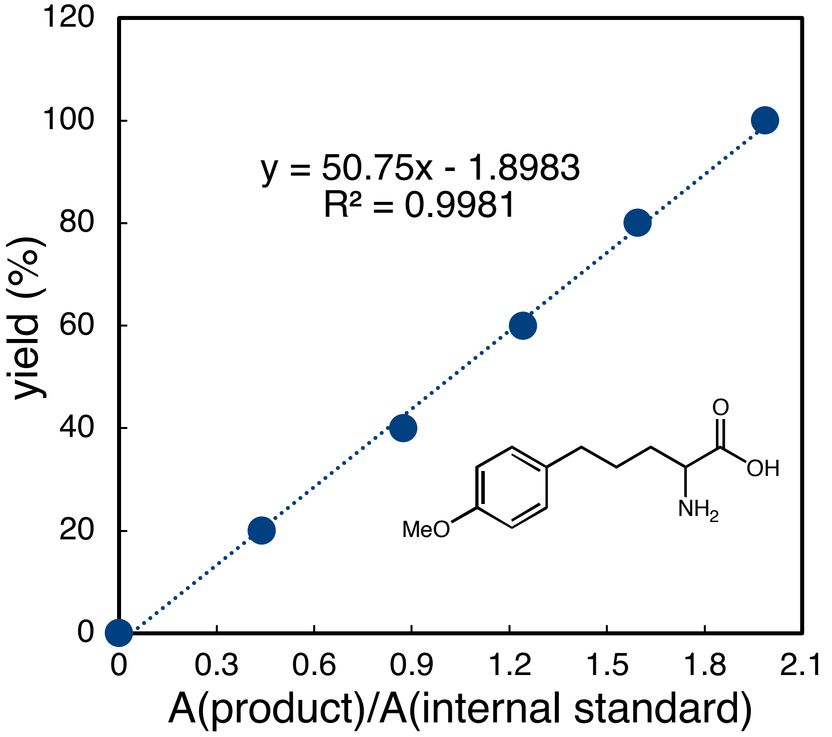

y = 50.75x - 1.8983
R² = 0.9981
yield (%)
A(product)/A(internal standard)
MeO
NH2
O
OH

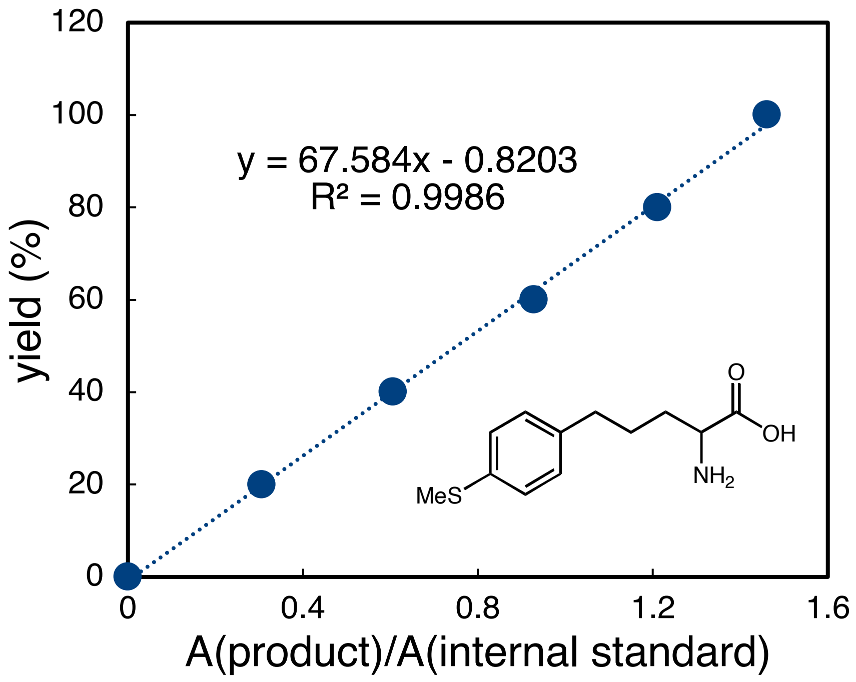

120
100
80
60
40
20
0
yield (%)
y = 67.584x - 0.8203
R² = 0.9986
O
OH
MeS
NH2
0
0.4
0.8
1.2
1.6
A(product)/A(internal standard)

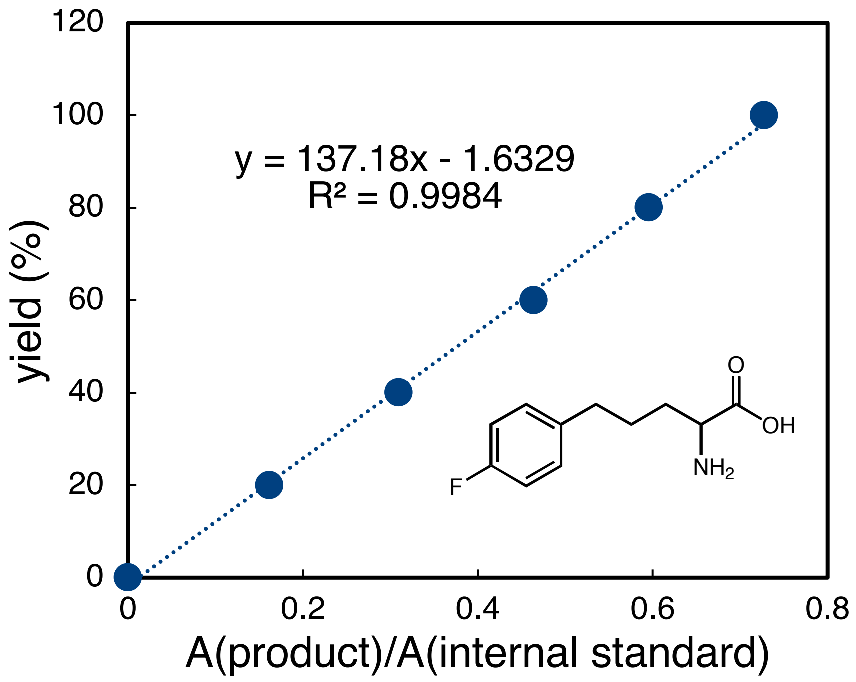

120
100
80
60
40
20
0
yield (%)
y = 137.18x - 1.6329
R² = 0.9984
O
OH
F
NH2
0
0.2
0.4
0.6
0.8
A(product)/A(internal standard)

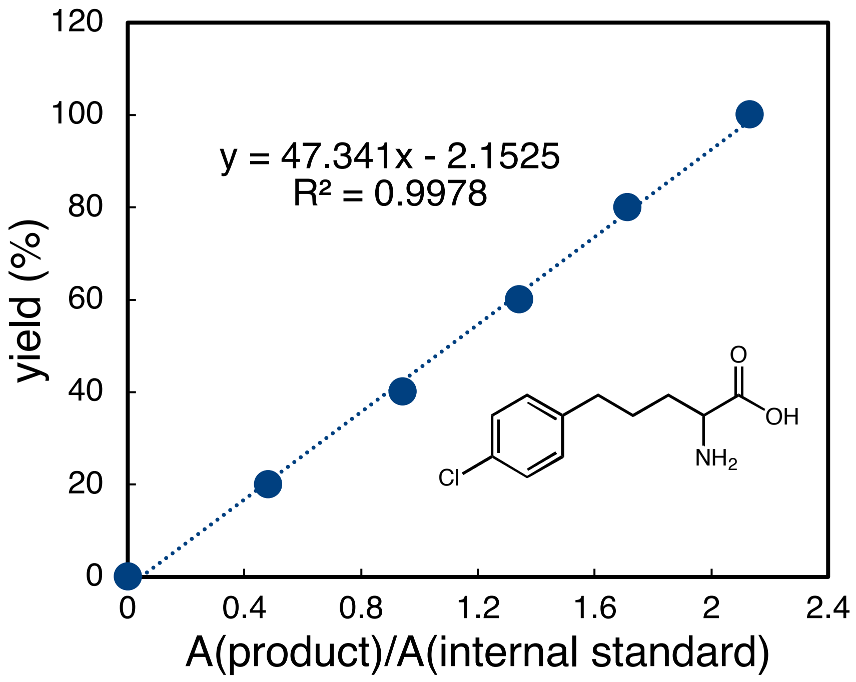
120
100
80
60
40
20
0
yield (%)
y = 47.341x - 2.1525
R² = 0.9978
O
OH
NH2
Cl
0
0.4
0.8
1.2
1.6
2
2.4
A(product)/A(internal standard)

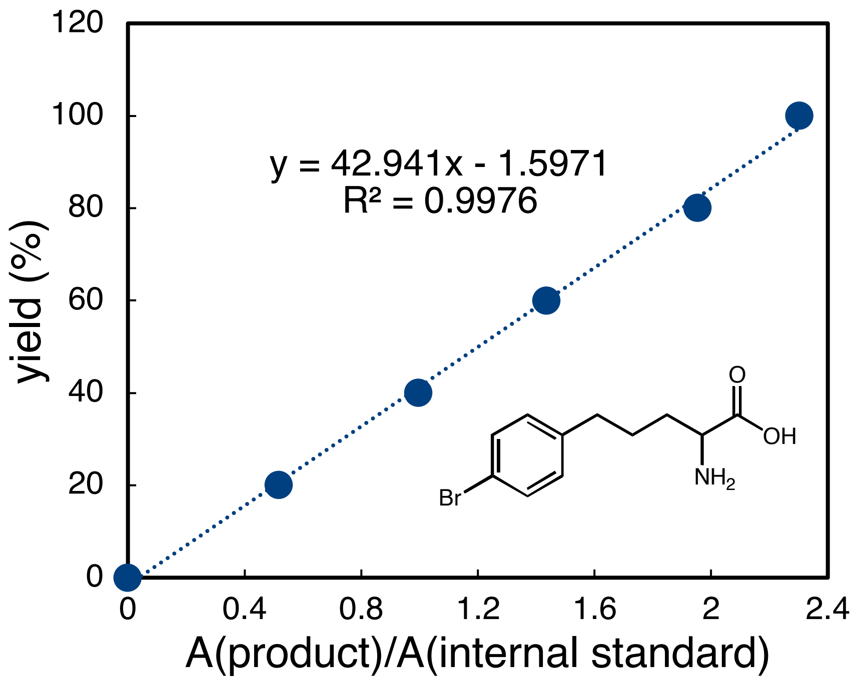
120
100
80
60
40
20
0
yield (%)
y = 42.941x - 1.5971
R² = 0.9976
O
OH
NH2
Br
0
0.4
0.8
1.2
1.6
2
2.4
A(product)/A(internal standard)

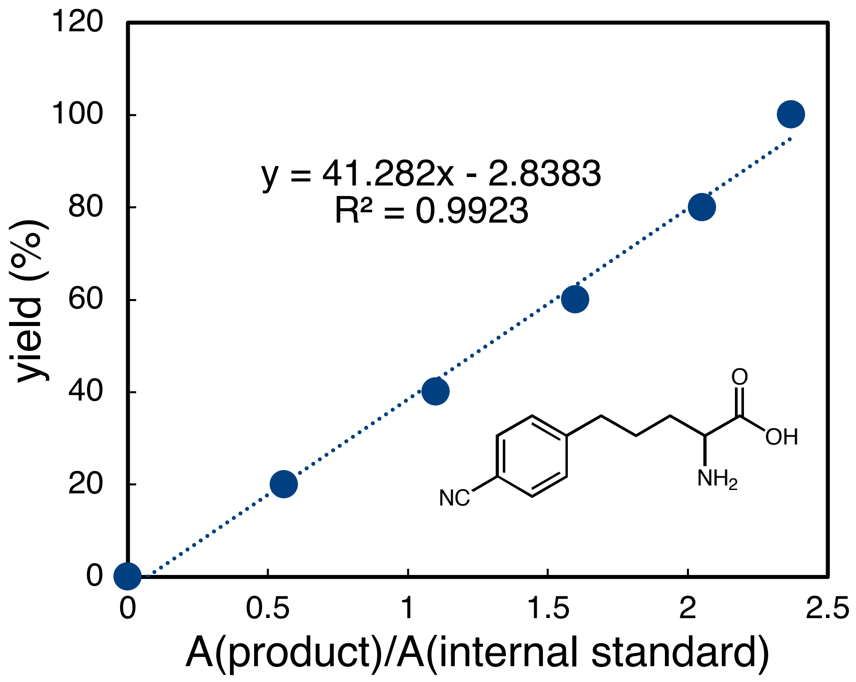
120
100
80
60
40
20
0
yield (%)
y = 41.282x - 2.8383
R² = 0.9923
O
OH
NH2
NC
0
0.5
1
1.5
2
2.5
A(product)/A(internal standard)

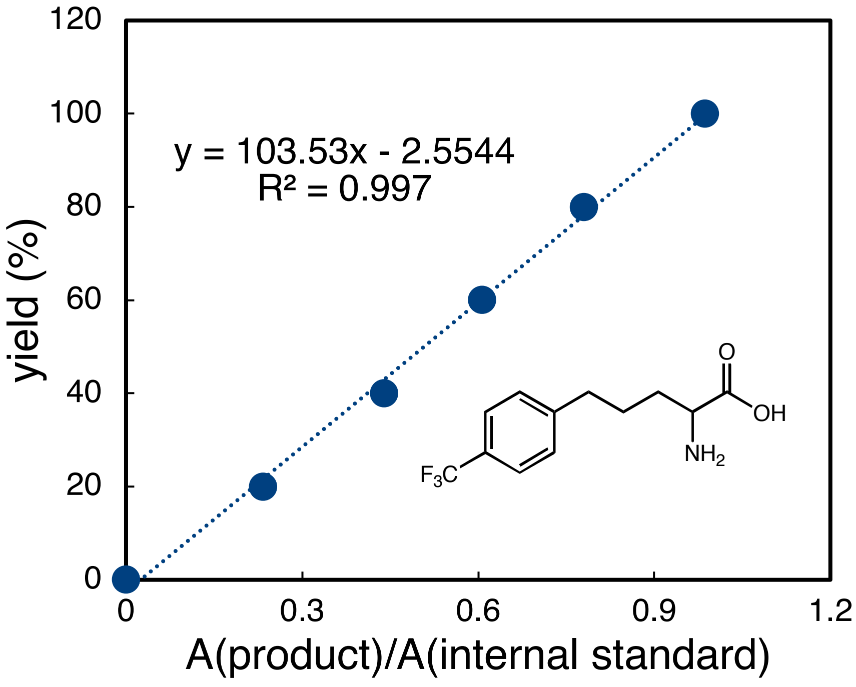
120
100
80
60
40
20
0
yield (%)
y = 103.53x - 2.5544
R² = 0.997
O
OH
NH2
F3C
0
0.3
0.6
0.9
1.2
A(product)/A(internal standard)

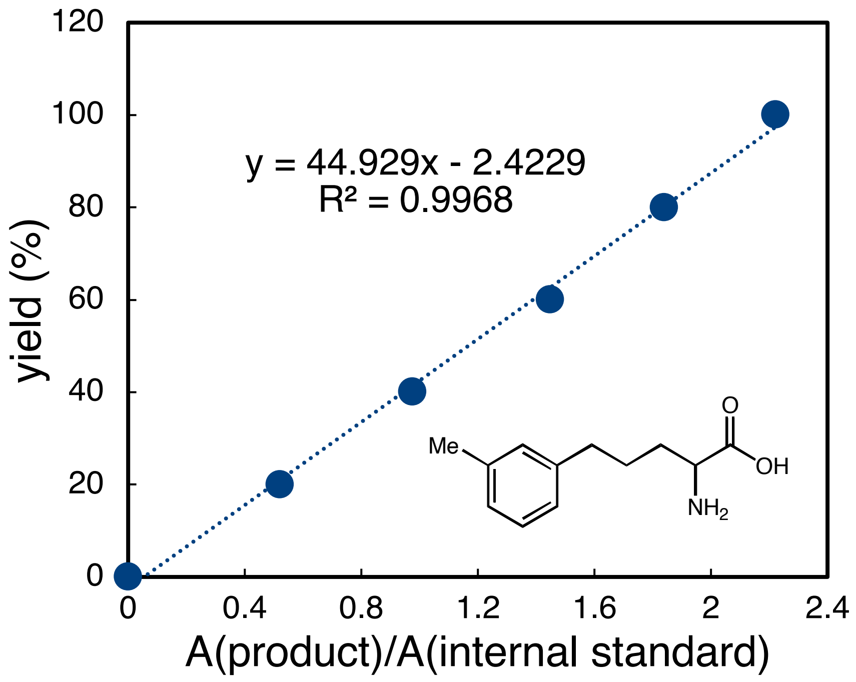

120
100
80
60
40
20
0
yield (%)
0
0.4
0.8
1.2
1.6
2
2.4
A(product)/A(internal standard)
y = 44.929x - 2.4229
R² = 0.9968
Me
O
OH
NH2

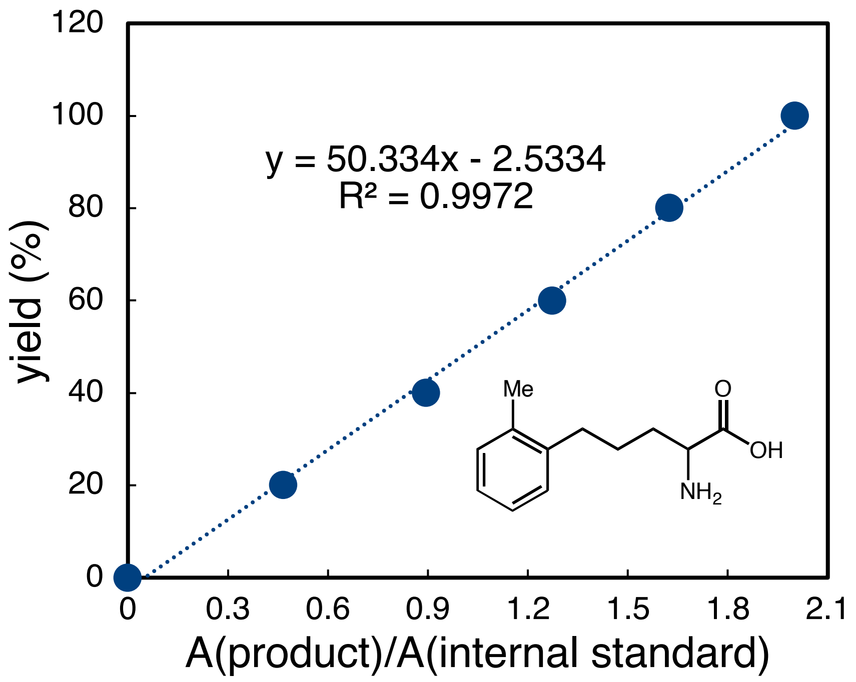

120
100
80
60
40
20
0
yield (%)
0
0.3
0.6
0.9
1.2
1.5
1.8
2.1
A(product)/A(internal standard)
y = 50.334x - 2.5334
R² = 0.9972
Me
O
OH
NH2

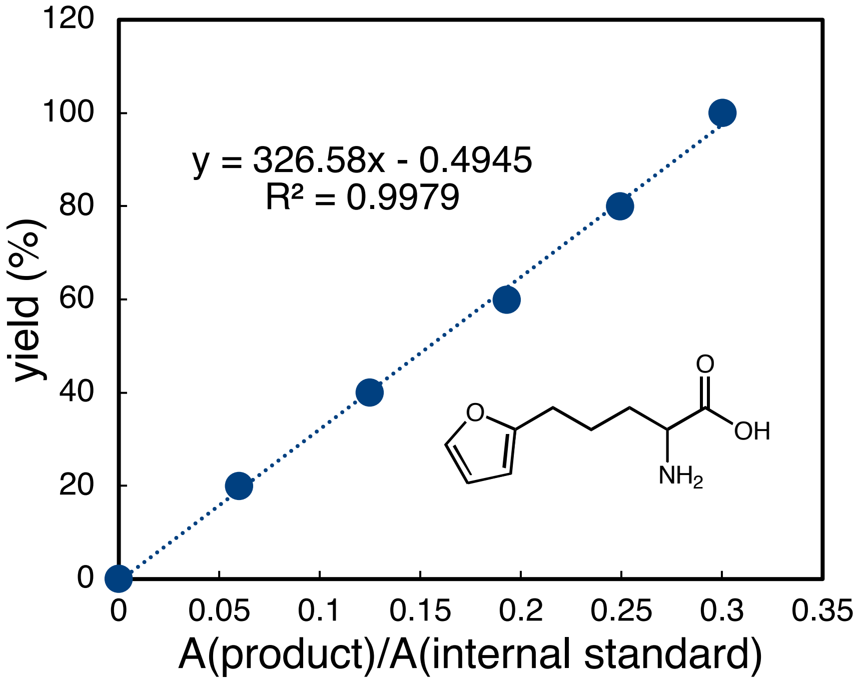
y = 326.58x - 0.4945
R² = 0.9979
yield (%)
A(product)/A(internal standard)
O
OH
NH2
0
20
40
60
80
100
120
0
0.05
0.1
0.15
0.2
0.25
0.3
0.35

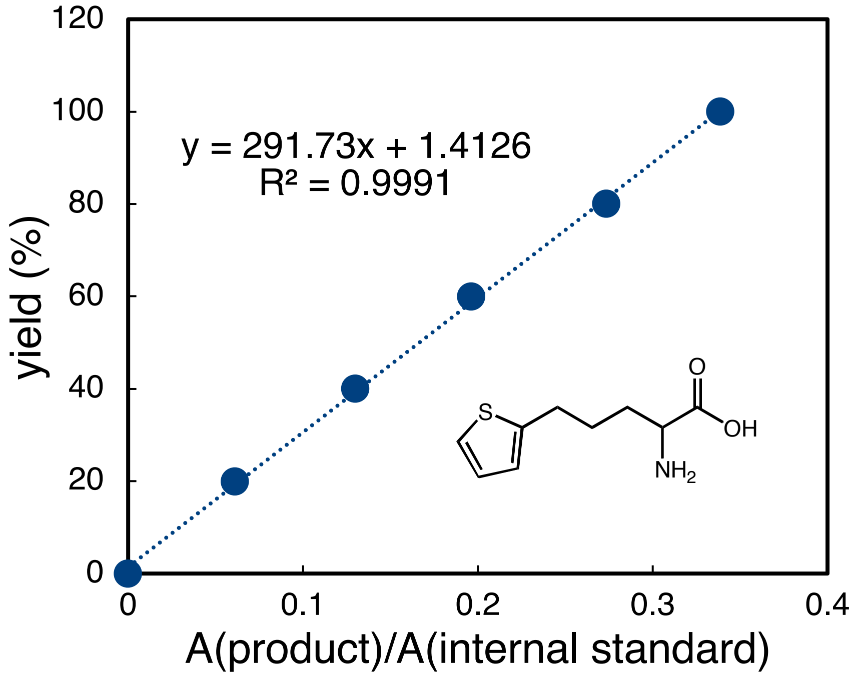
y = 291.73x + 1.4126
R² = 0.9991
yield (%)
A(product)/A(internal standard)
S
O
OH
NH2
0
20
40
60
80
100
120
0
0.1
0.2
0.3
0.4

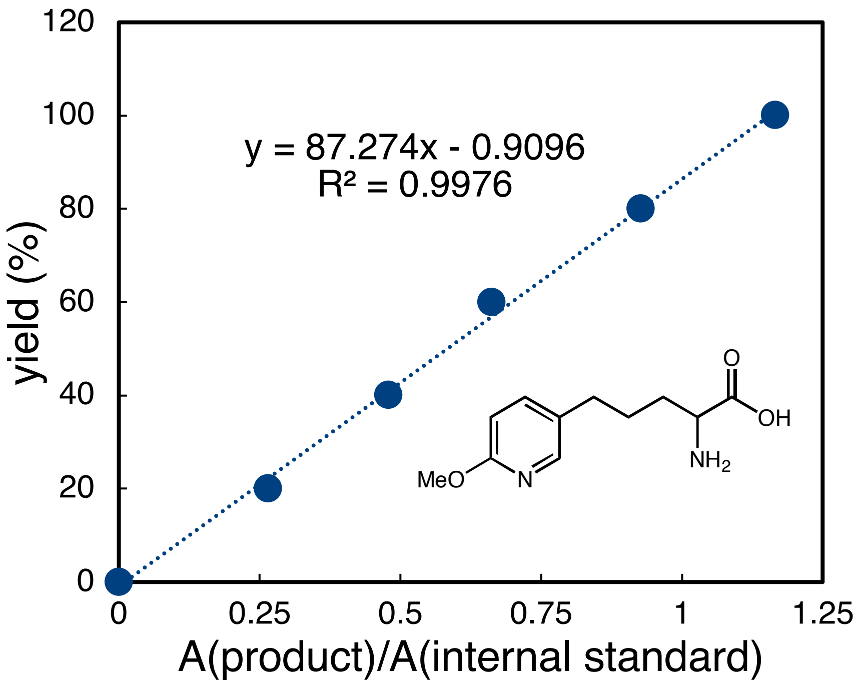

y = 87.274x - 0.9096
R² = 0.9976
yield (%)
A(product)/A(internal standard)
MeO
N
O
OH
NH2

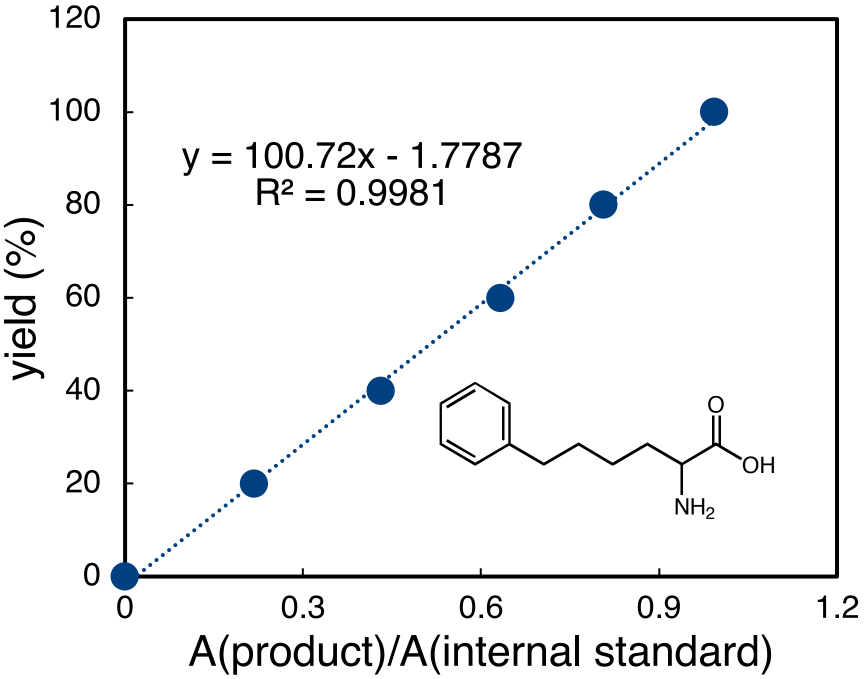

y = 100.72x - 1.7787
R² = 0.9981
yield (%)
A(product)/A(internal standard)
O
OH
NH2

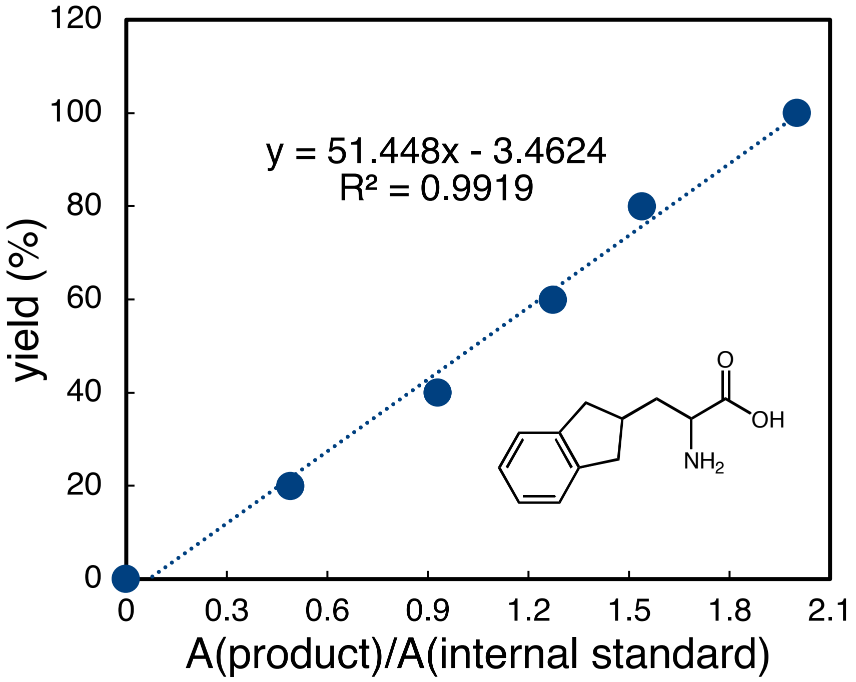
y = 51.448x - 3.4624
R² = 0.9919
yield (%)
120
100
80
60
40
20
0
0
0.3
0.6
0.9
1.2
1.5
1.8
2.1
A(product)/A(internal standard)
O
OH
NH2

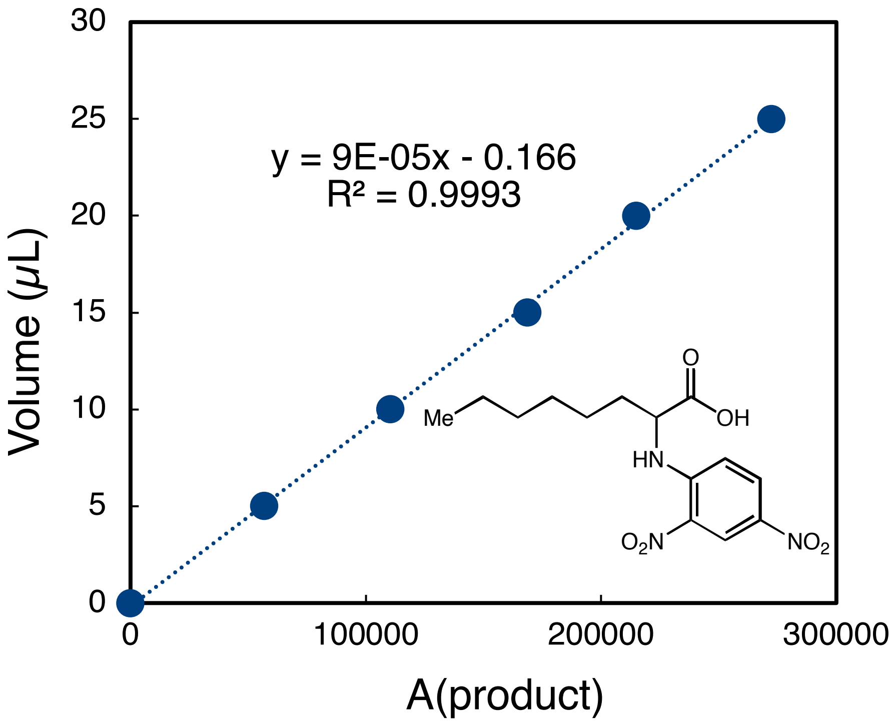
y = 9E-05x - 0.166
R² = 0.9993
Volume (μL)
A(product)
0
5
10
15
20
25
30
0
100000
200000
300000
Me
O
OH
HN
O2N
NO2

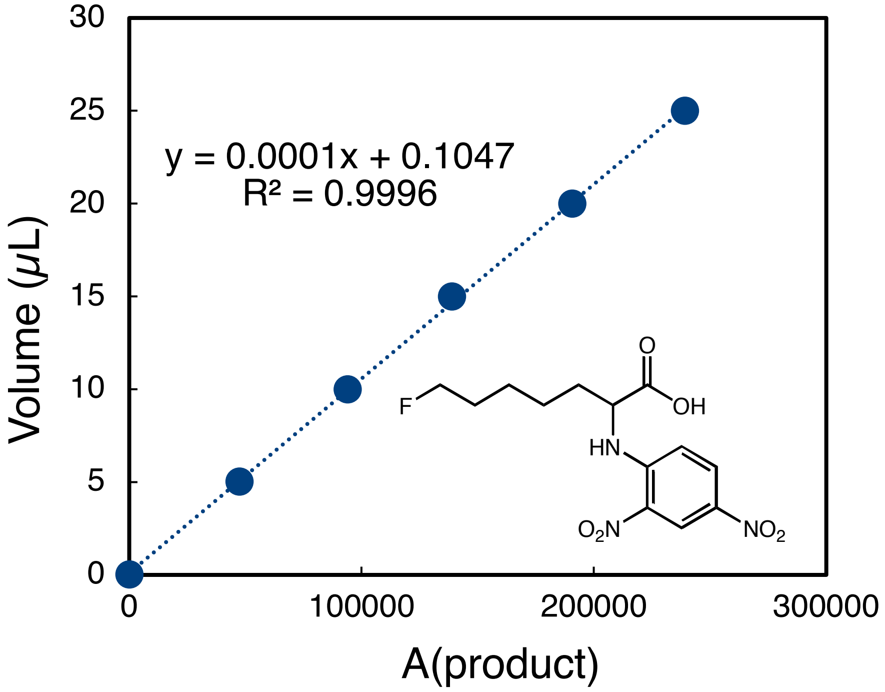
y = 0.0001x + 0.1047
R² = 0.9996
Volume (μL)
A(product)
0
5
10
15
20
25
30
0
100000
200000
300000
F
O
OH
HN
O2N
NO2

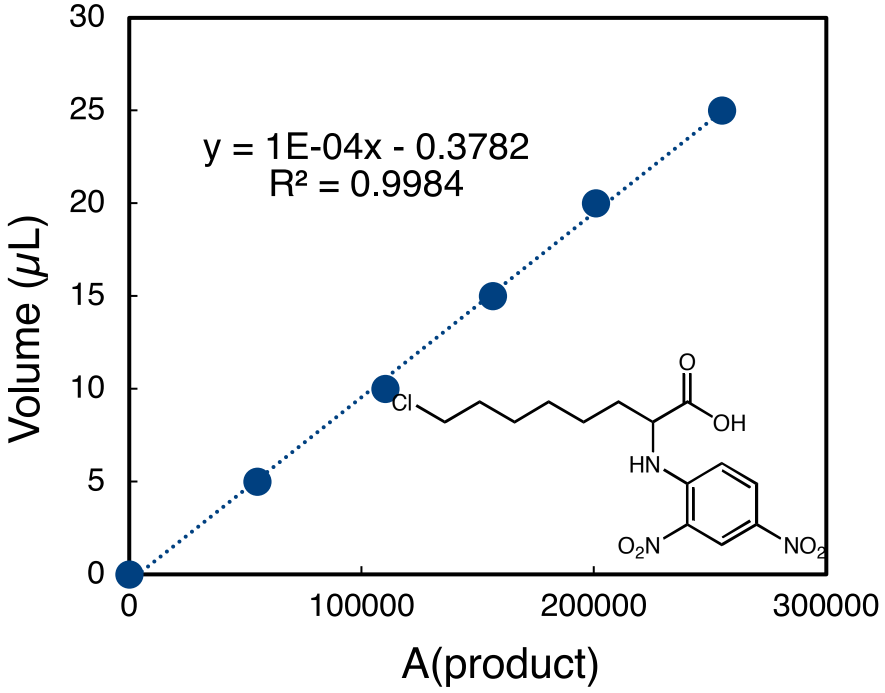

y = 1E-04x - 0.3782
R² = 0.9984
Volume (μL)
A(product)
0
5
10
15
20
25
30
0
100000
200000
300000
Cl
O
OH
HN
O2N
NO2

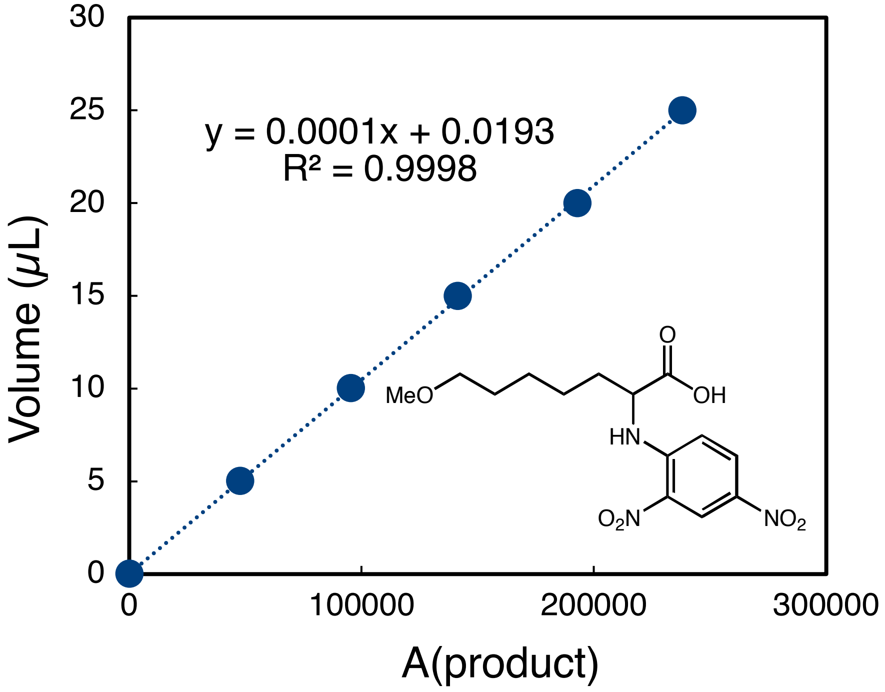

y = 0.0001x + 0.0193
R² = 0.9998
Volume (μL)
A(product)
0
5
10
15
20
25
30
0
100000
200000
300000
MeO
O
OH
HN
O2N
NO2

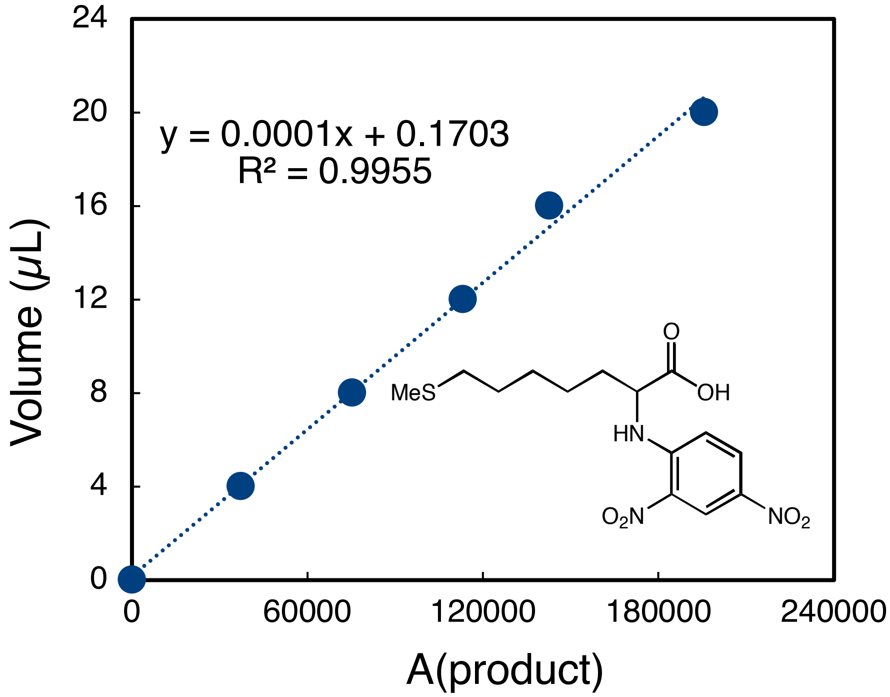

y = 0.0001x + 0.1703
R² = 0.9955
Volume (μL)
A(product)
0
4
8
12
16
20
24
0
60000
120000
180000
240000
MeS
O
OH
HN
O2N
NO2

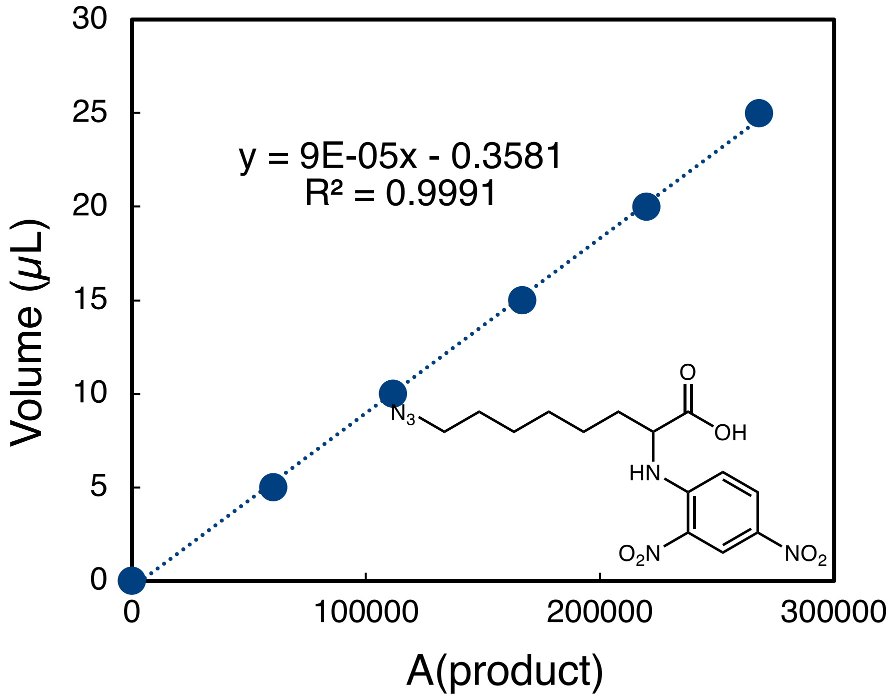

y = 9E-05x - 0.3581
R² = 0.9991
Volume (μL)
A(product)
0
5
10
15
20
25
30
0
100000
200000
300000
N3
O
OH
HN
O2N
NO2

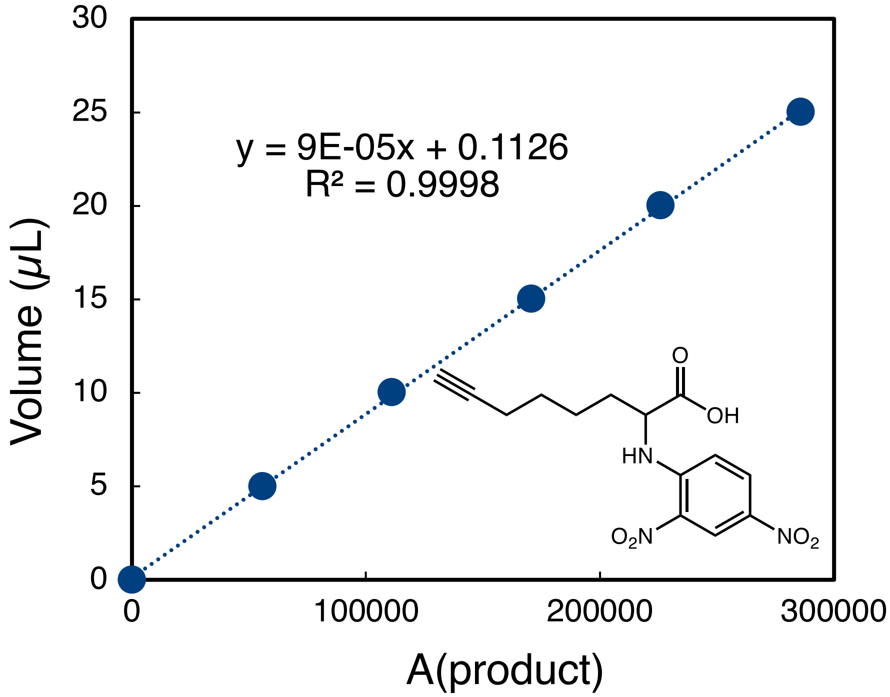
y = 9E-05x + 0.1126
R² = 0.9998
O
OH
HN
O2N
NO2
Volume (μL)
A(product)
0
5
10
15
20
25
30
0
100000
200000
300000

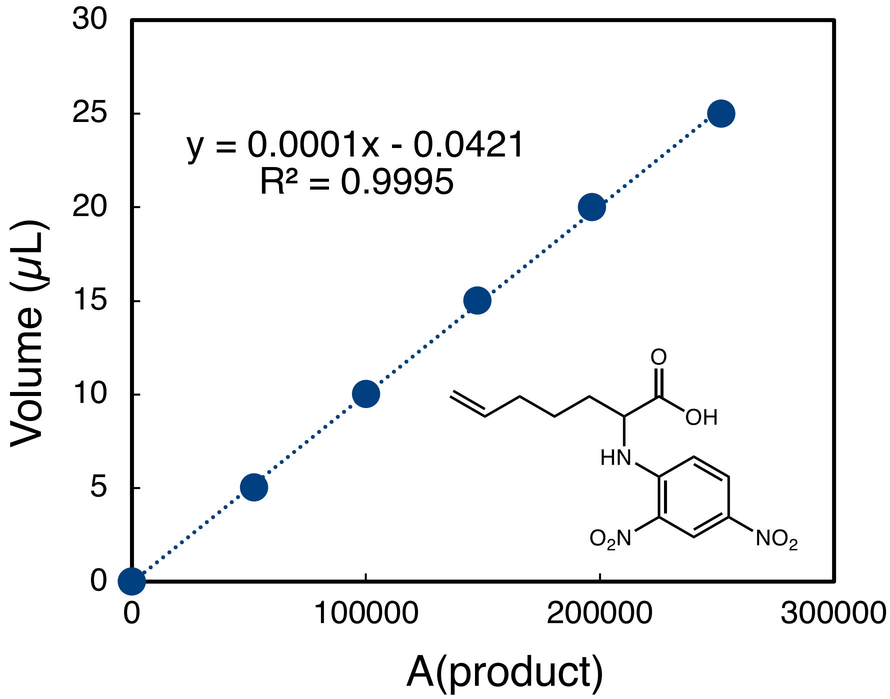
y = 0.0001x - 0.0421
R² = 0.9995
O
OH
HN
O2N
NO2
Volume (μL)
A(product)
0
5
10
15
20
25
30
0
100000
200000
300000

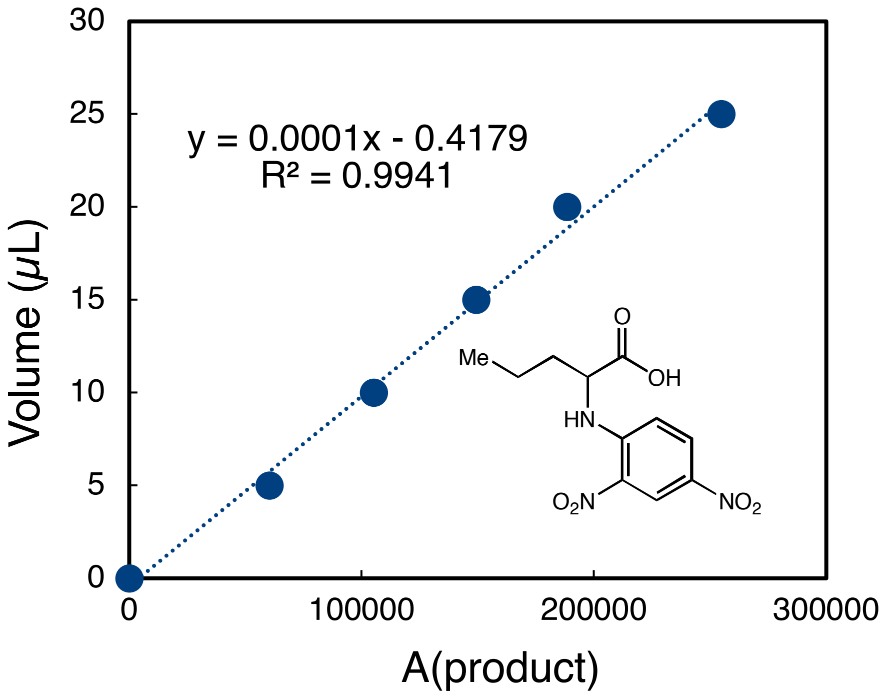
y = 0.0001x - 0.4179
R² = 0.9941
Volume (μL)
A(product)
Me
O
OH
HN
O2N
NO2

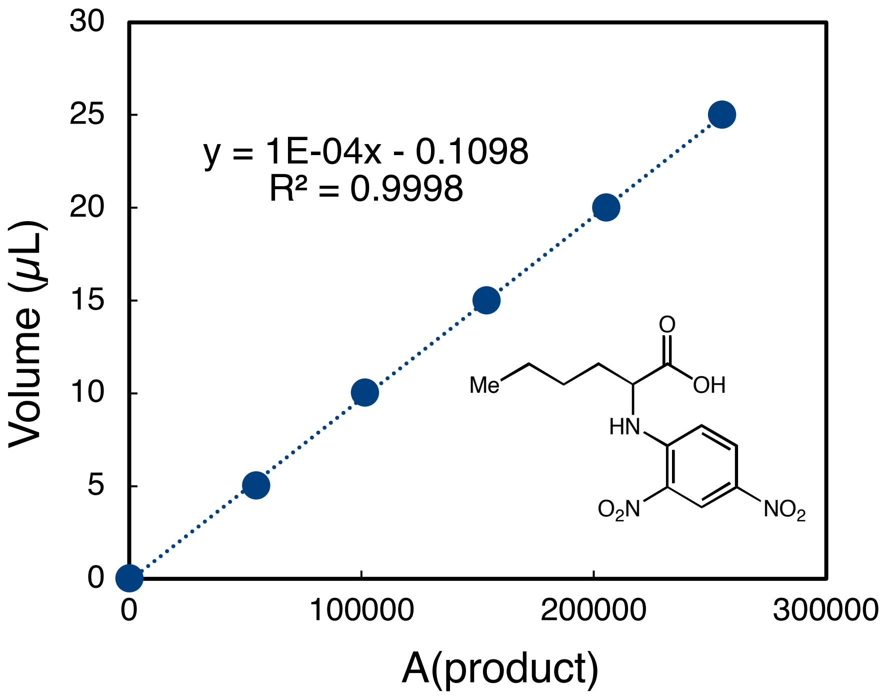
y = 1E-04x - 0.1098
R² = 0.9998
Volume (μL)
A(product)
Me
O
OH
HN
O2N
NO2

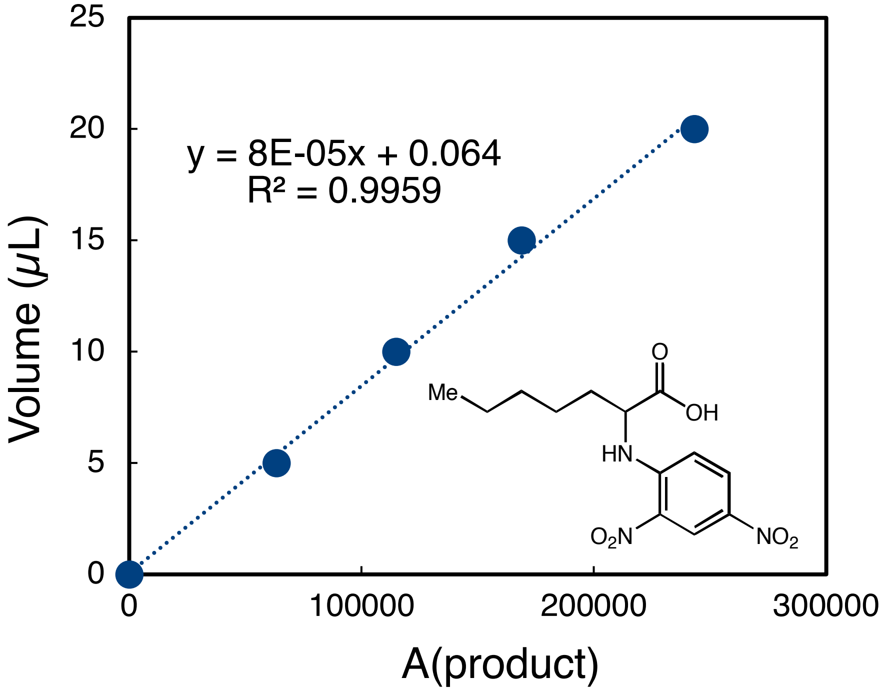

Volume (μL)
A(product)
y = 8E-05x + 0.064
R² = 0.9959
0
5
10
15
20
25
0
100000
200000
300000
Me
O
OH
HN
O2N
NO2

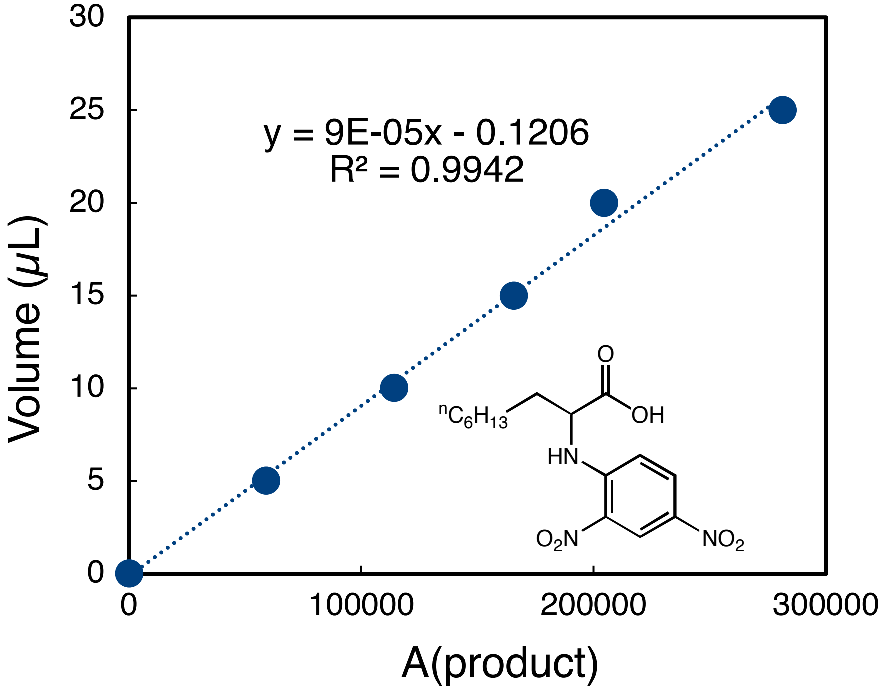

Volume (μL)
A(product)
y = 9E-05x - 0.1206
R² = 0.9942
0
5
10
15
20
25
30
0
100000
200000
300000
nC6H13
O
OH
HN
O2N
NO2

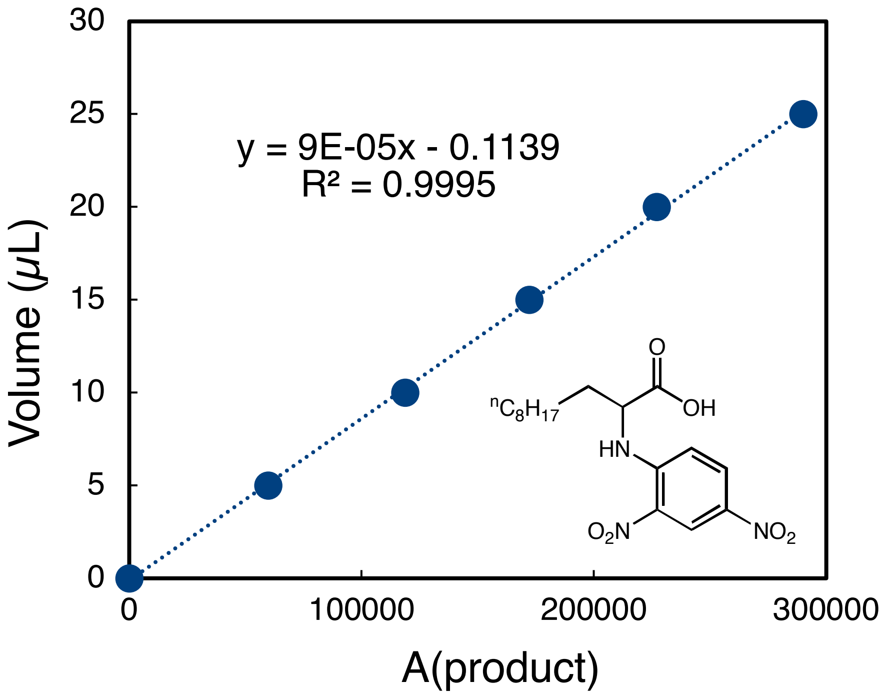

y = 9E-05x - 0.1139
R² = 0.9995
Volume (μL)
A(product)
0
5
10
15
20
25
30
0
100000
200000
300000
O
OH
nC8H17
HN
O2N
NO2

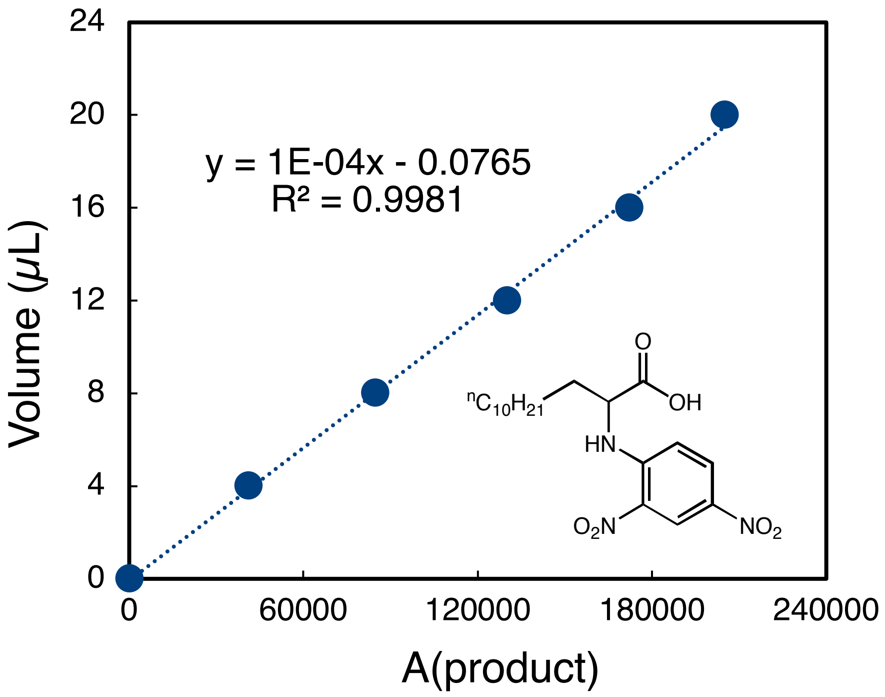

y = 1E-04x - 0.0765
R² = 0.9981
Volume (μL)
A(product)
0
4
8
12
16
20
24
0
60000
120000
180000
240000
O
OH
nC10H21
HN
O2N
NO2

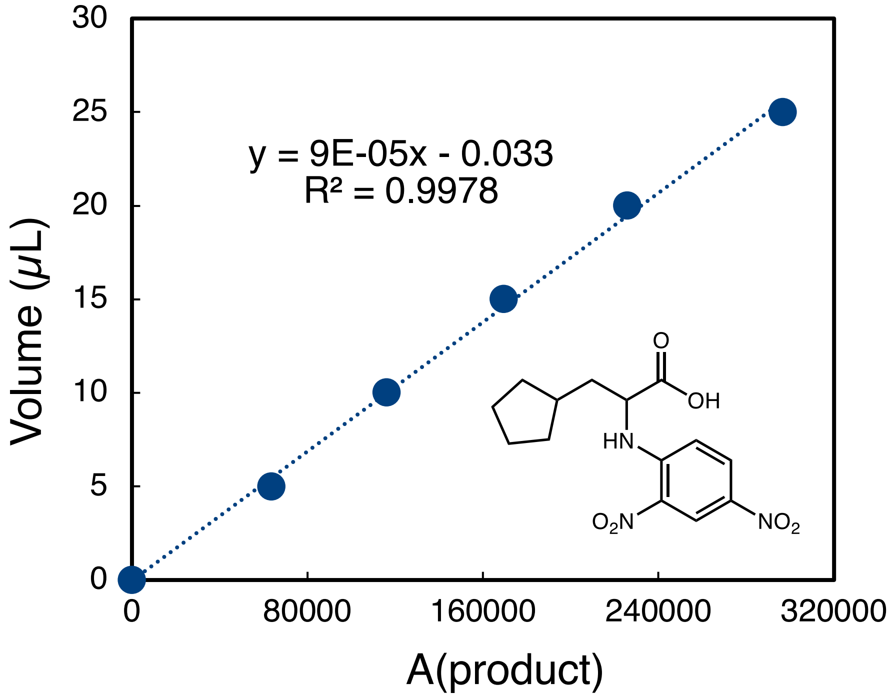
y = 9E-05x - 0.033
R² = 0.9978
Volume (μL)
A(product)
O
OH
HN
O2N
NO2

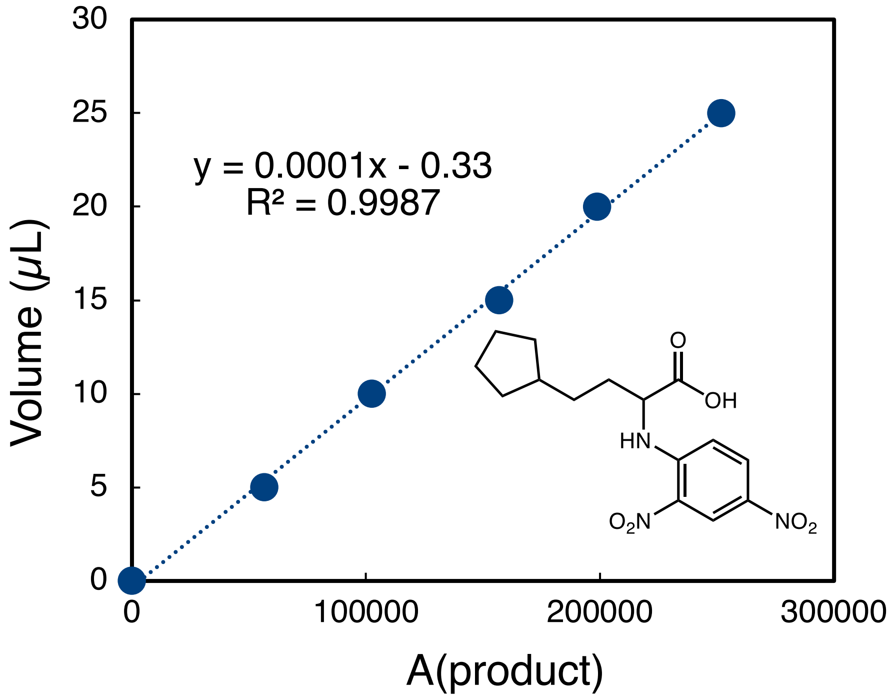
y = 0.0001x - 0.33
R² = 0.9987
Volume (μL)
A(product)
O
OH
HN
O2N
NO2

### IX. HPLC traces for Marfey's analysis

#### Determination of absolute configuration with HPLC analysis

The absolute configuration of (*S*)-**3a** and (*S*)-**5a** was determined by comparing their HPLC retention times with those of standard chiral compounds using Marfey's analysis.

#### 2-Amino-5-phenylpentanoic acid (3a)

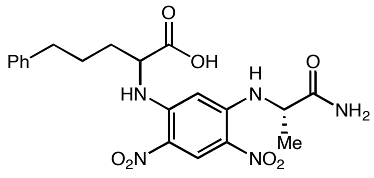

(*rac*)-**3a** derivative

Marfey's analysis of (*rac*)-**3a**

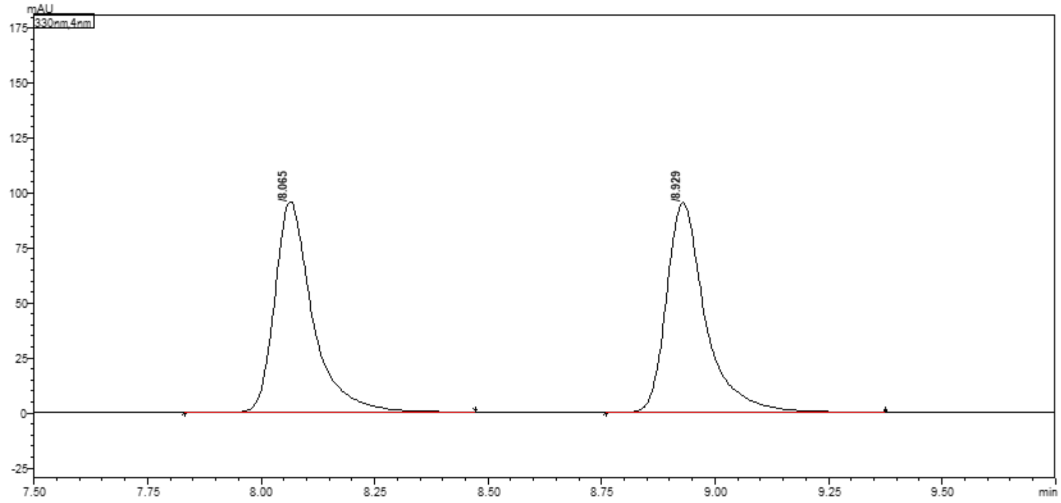

| Peak# | Ret. Time | Height | Area | Area% |
|---|---|---|---|---|
| 1 | 8.065 | 95254 | 548438 | 49.310 |
| 2 | 8.929 | 94789 | 563777 | 50.690 |
| Total | | 190043 | 1112215 | 100.000 |

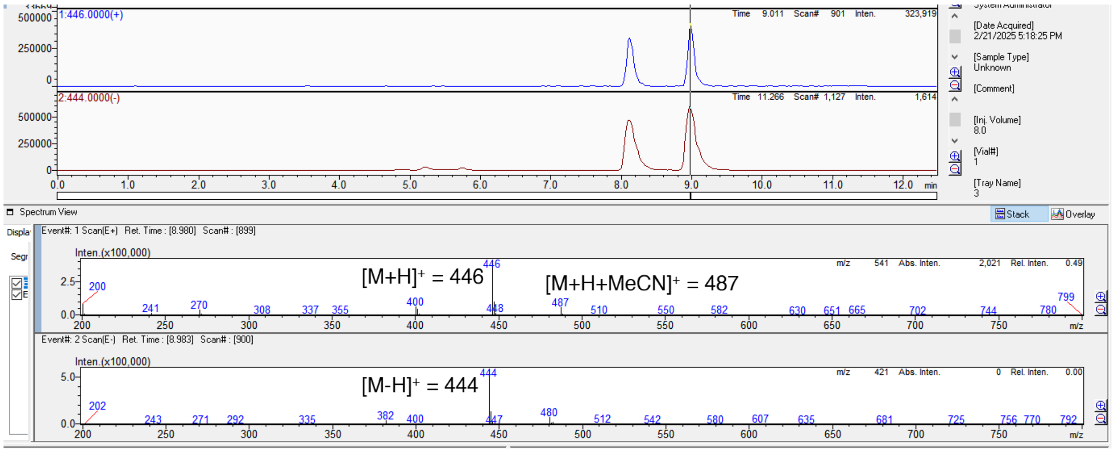

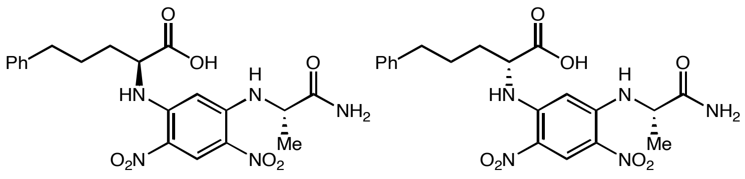

L-**3a** derivative

D-**3a** derivative

Marfey's analysis of L-**3a**

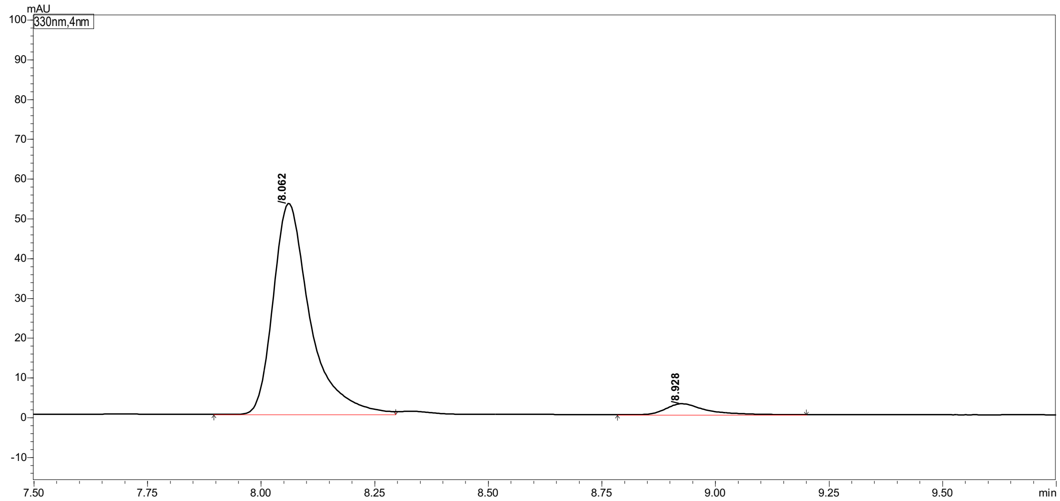

Peak Table | Compound | Group | Calibration Curve

| Peak# | Ret. Time | Height | Area | Area% |
|---|---|---|---|---|
| 1 | 8.062 | 52969 | 299512 | 94.791 |
| 2 | 8.928 | 2739 | 16458 | 5.209 |
| Total | | 55708 | 315970 | 100.000 |

Marfey's analysis of D-**3a**

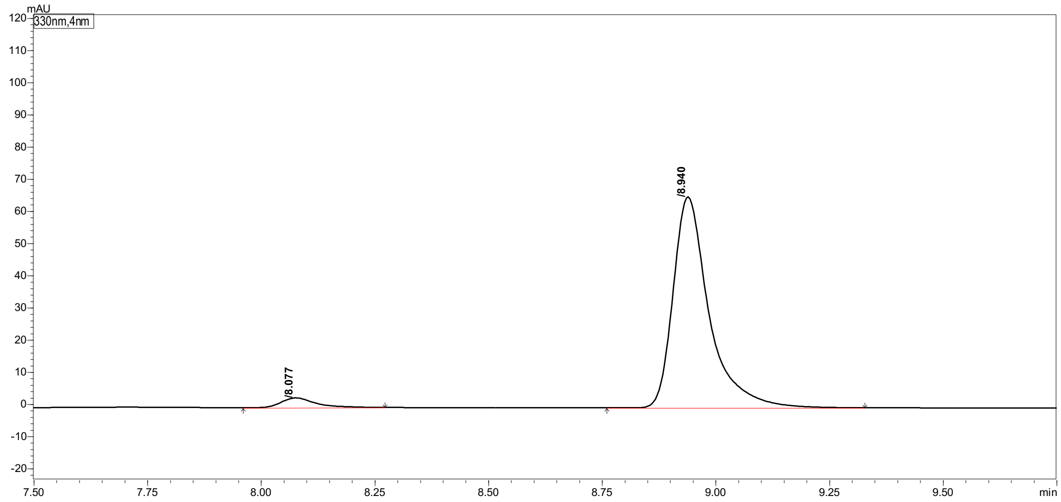

Peak Table | Compound | Group | Calibration Curve

| Peak# | Ret. Time | Height | Area | Area% |
|---|---|---|---|---|
| 1 | 8.077 | 2995 | 15430 | 4.107 |
| 2 | 8.940 | 65237 | 360271 | 95.893 |
| Total | | 68231 | 375700 | 100.000 |

#### 2-Amino-5-(p-tolyl)pentanoic acid (3b)

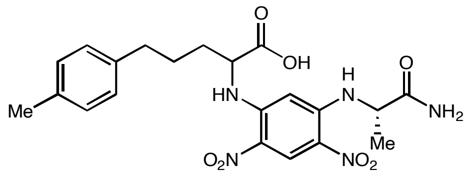

(*rac*)-**3b** derivative

#### Marfey's analysis of (*rac*)-3b

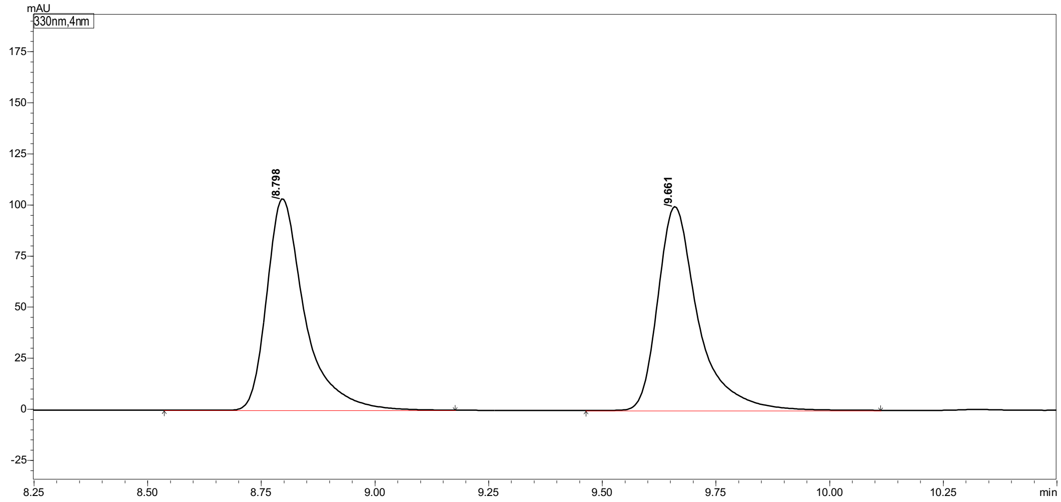

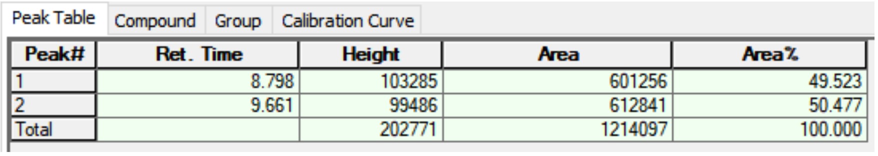

Peak Table | Compound | Group | Calibration Curve

| Peak# | Ret. Time | Height | Area | Area% |
|---|---|---|---|---|
| 1 | 8.798 | 103285 | 601256 | 49.523 |
| 2 | 9.661 | 99486 | 612841 | 50.477 |
| Total | | 202771 | 1214097 | 100.000 |

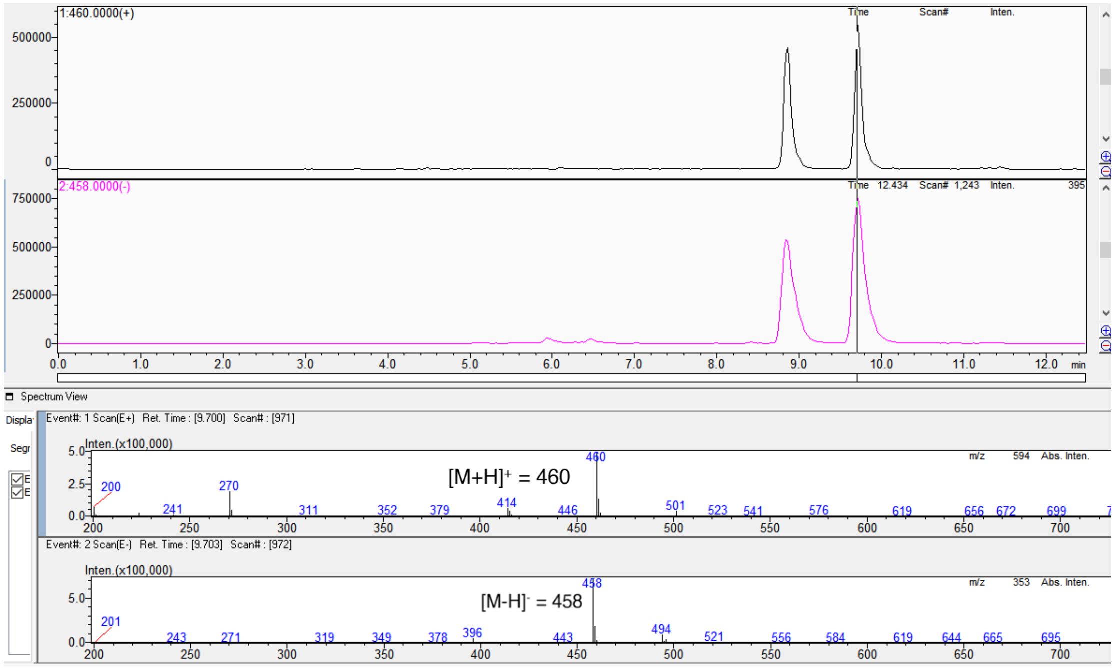

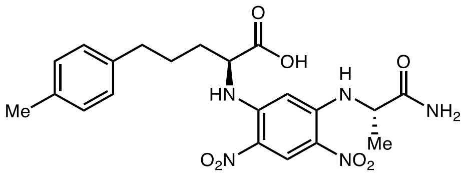

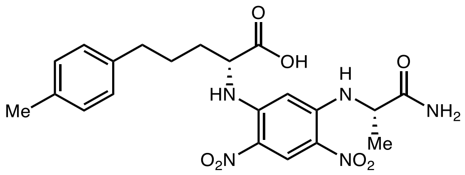

L-**3b** derivative

D-**3b** derivative

Marfey's analysis of L-**3b**

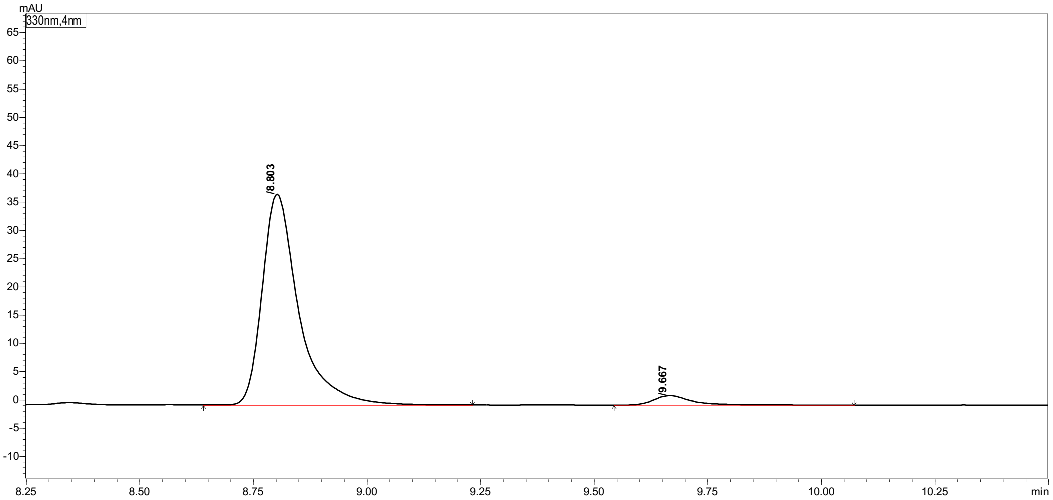

| Peak# | Ret. Time | Height | Area | Area% |
|---|---|---|---|---|
| 1 | 8.803 | 37095 | 207998 | 95.459 |
| 2 | 9.667 | 1682 | 9895 | 4.541 |
| Total | | 38777 | 217894 | 100.000 |

Marfey's analysis of D-**3b**

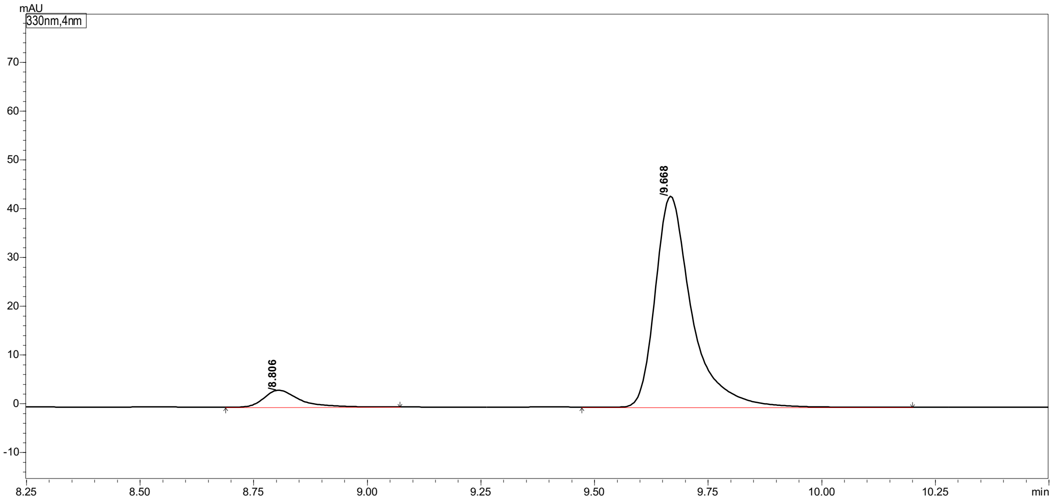

| Peak# | Ret. Time | Height | Area | Area% |
|---|---|---|---|---|
| 1 | 8.806 | 3467 | 18398 | 7.081 |
| 2 | 9.668 | 43093 | 241441 | 92.919 |
| Total | | 46560 | 259839 | 100.000 |

#### 2-Amino-5-(4-ethylphenyl)pentanoic acid (3c)

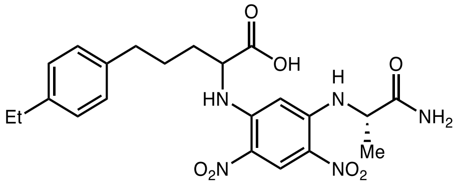

(*rac*)-**3c** derivative

Marfey's analysis of (*rac*)-**3g**

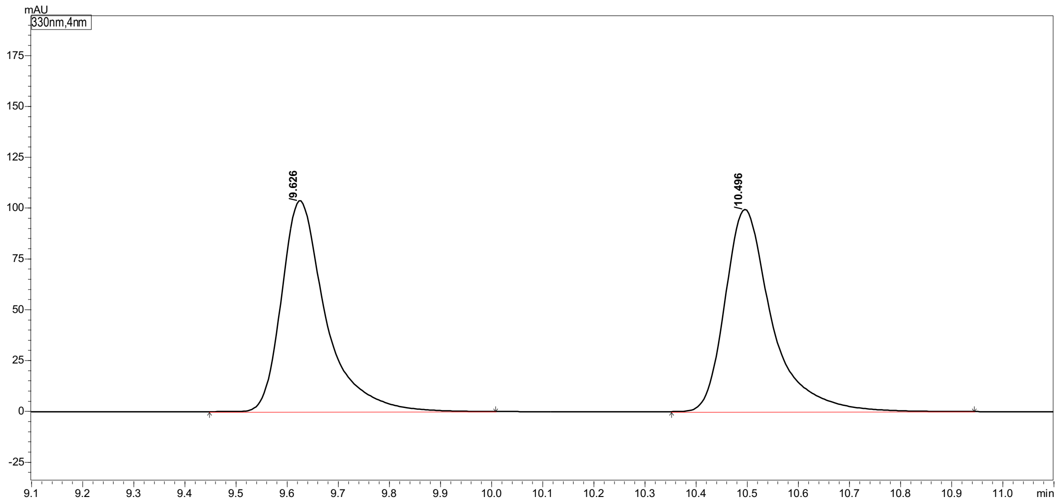

Peak Table | Compound | Group | Calibration Curve

| Peak# | Ret. Time | Height | Area | Area% |
|---|---|---|---|---|
| 1 | 9.626 | 102905 | 620001 | 49.487 |
| 2 | 10.496 | 99141 | 632852 | 50.513 |
| Total | | 202046 | 1252853 | 100.000 |

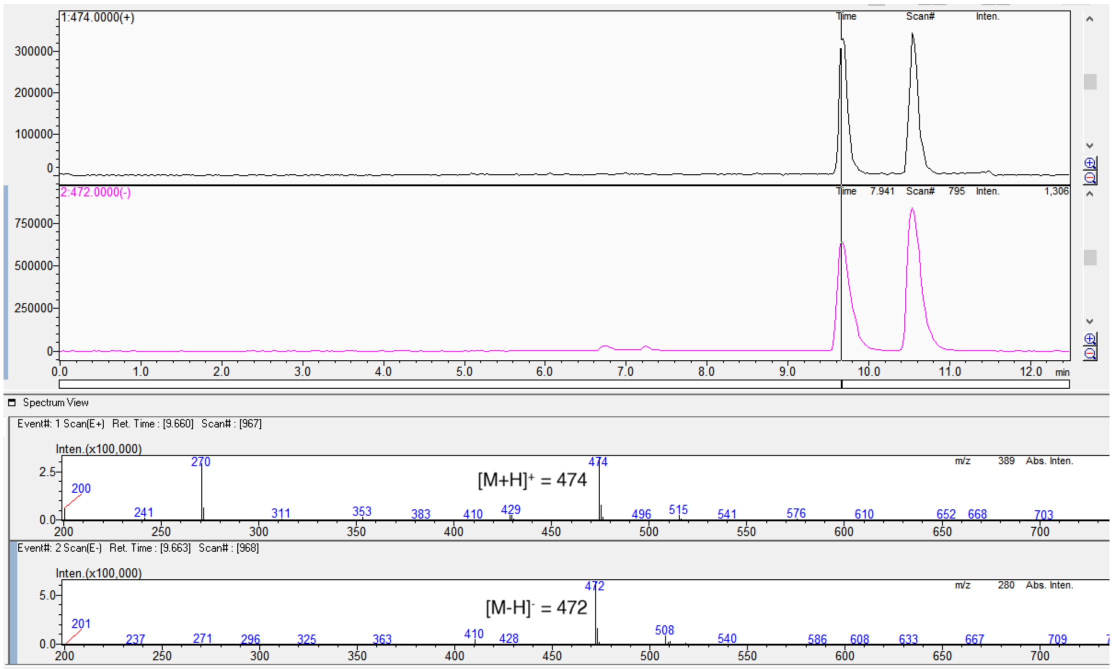

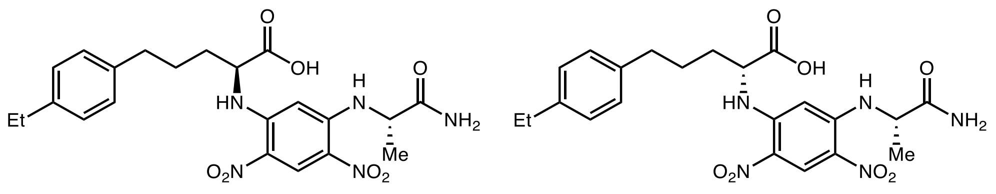

L-**3c** derivative

D-**3c** derivative

Marfey’s analysis of L-**3c**

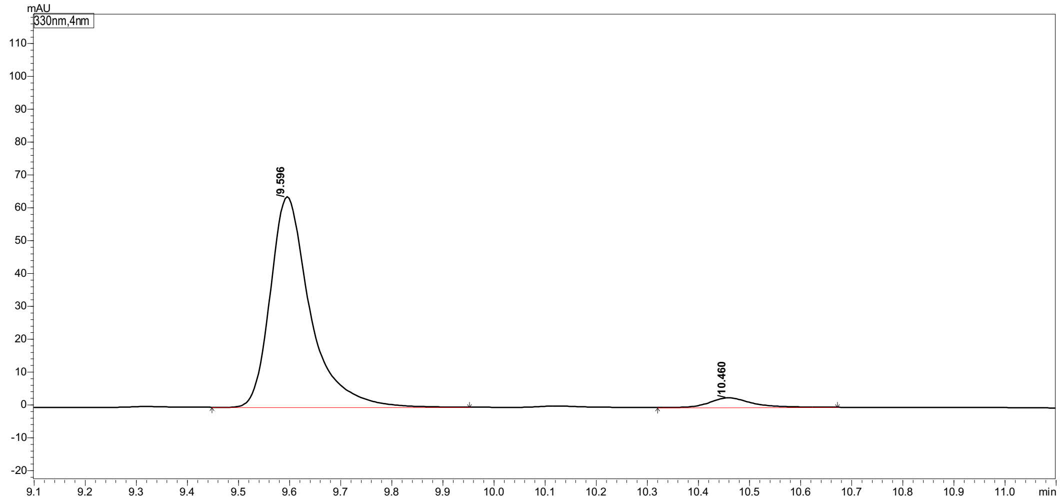

| Peak# | Ret. Time | Height | Area | Area% |
|---|---|---|---|---|
| 1 | 9.596 | 63828 | 347084 | 95.567 |
| 2 | 10.460 | 2901 | 16101 | 4.433 |
| Total | | 66729 | 363185 | 100.000 |

Marfey’s analysis of D-**3c**

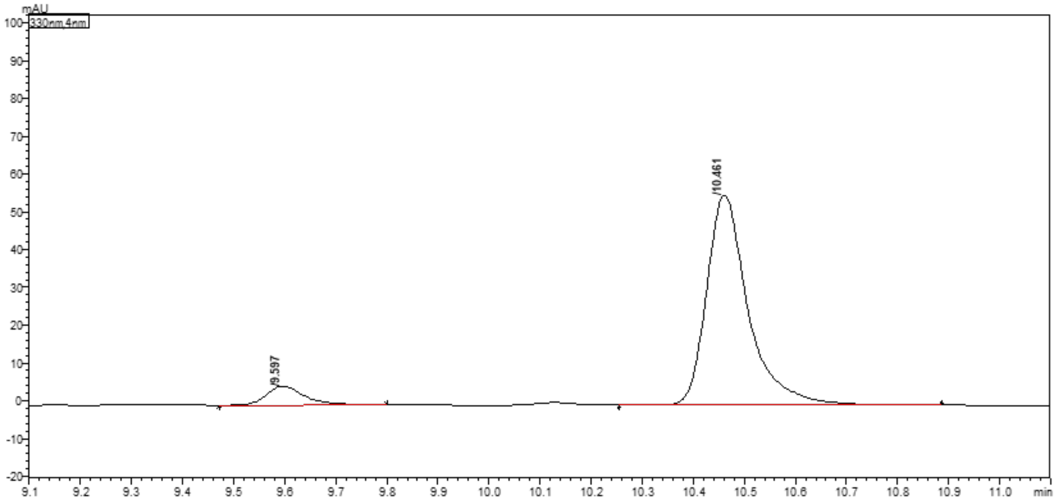

| Peak# | Ret. Time | Height | Area | Area% |
|---|---|---|---|---|
| 1 | 9.597 | 4995 | 26375 | 7.793 |
| 2 | 10.461 | 55459 | 312058 | 92.207 |
| Total | | 60454 | 338434 | 100.000 |

#### 2-Amino-5-(4-methoxyphenyl)pentanoic acid (3d)

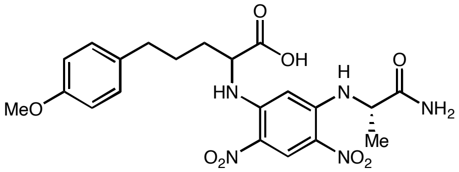

(*rac*)-**3d** derivative

#### Marfey's analysis of (*rac*)-**3d**

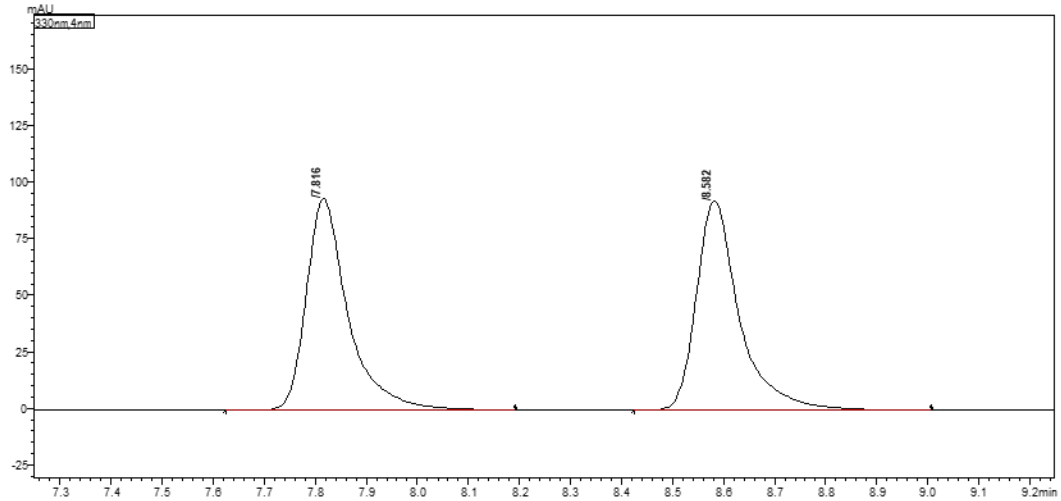

| Peak# | Ret. Time | Height | Area | Area% |
|---|---|---|---|---|
| 1 | 7.816 | 92597 | 526410 | 49.645 |
| 2 | 8.582 | 92260 | 533947 | 50.355 |
| Total | | 184858 | 1060358 | 100.000 |

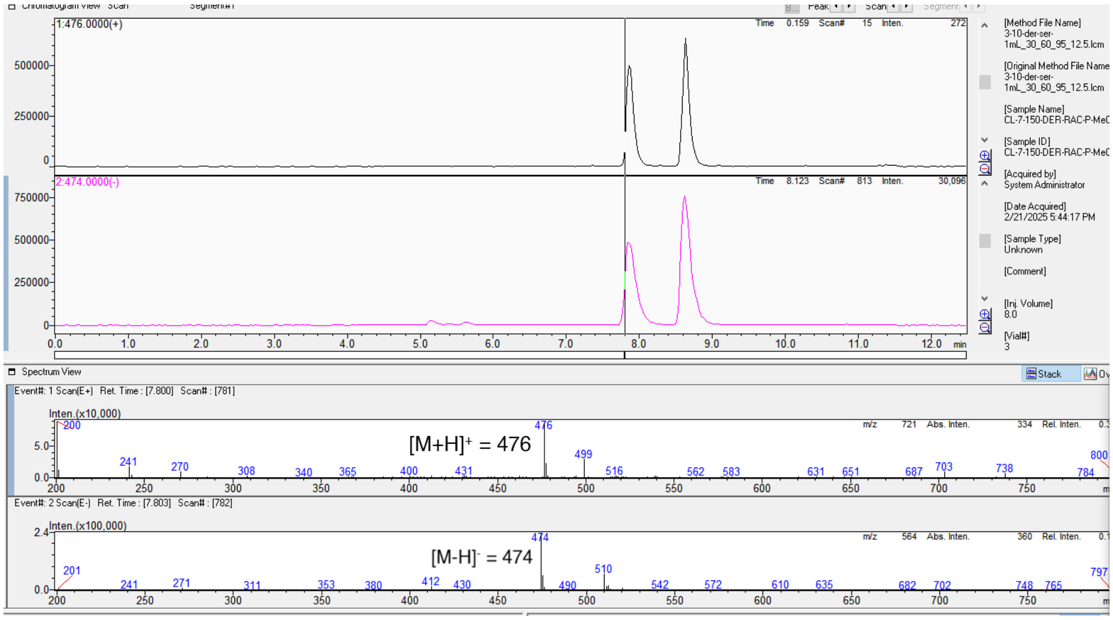

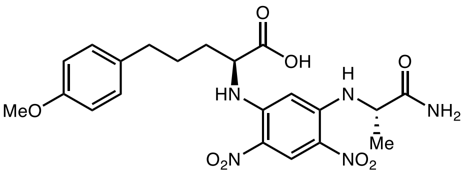

L-**3d** derivative

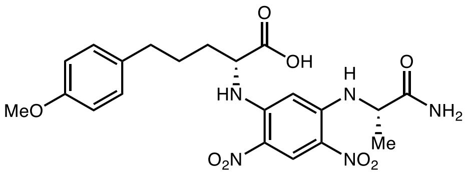

D-**3d** derivative

Marfey's analysis of L-**3d**

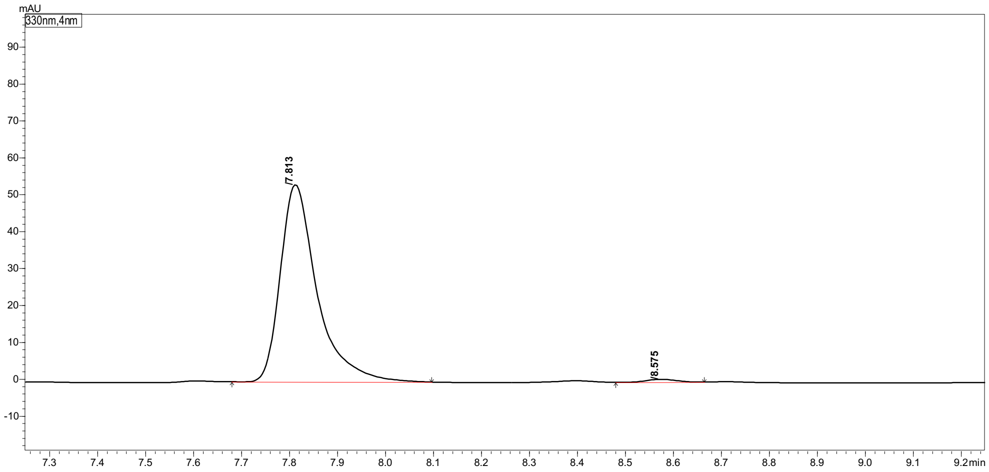

Peak Table | Compound | Group | Calibration Curve

| Peak# | Ret. Time | Height | Area | Area% |
|---|---|---|---|---|
| 1 | 7.813 | 53290 | 284363 | 98.872 |
| 2 | 8.575 | 777 | 3243 | 1.128 |
| Total | | 54067 | 287606 | 100.000 |

Marfey's analysis of D-**3d**

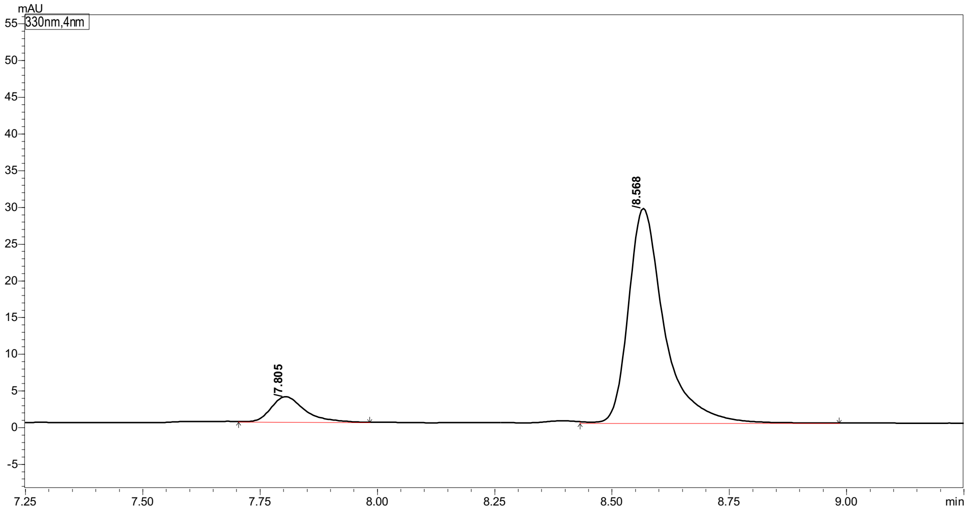

Peak Table | Compound | Group | Calibration Curve

| Peak# | Ret. Time | Height | Area | Area% |
|---|---|---|---|---|
| 1 | 7.805 | 3409 | 16657 | 9.756 |
| 2 | 8.568 | 29106 | 154073 | 90.244 |
| Total | | 32515 | 170731 | 100.000 |

#### 2-Amino-5-(4-(methylthio)phenyl)pentanoic acid (3e)

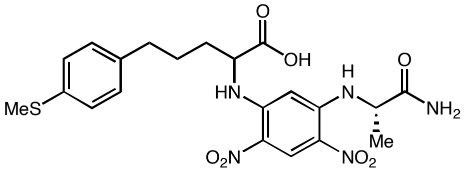

(*rac*)-**3e** derivative

Marfey's analysis of (*rac*)-**3e**

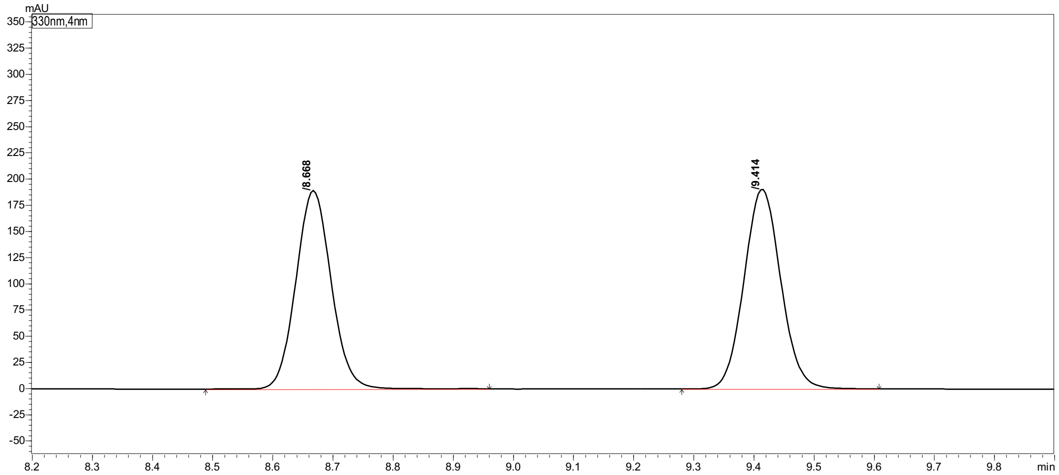

| Peak# | Ret. Time | Height | Area | Area% |
|---|---|---|---|---|
| 1 | 8.668 | 188271 | 770960 | 48.712 |
| 2 | 9.414 | 190317 | 811722 | 51.288 |
| Total | | 378588 | 1582682 | 100.000 |

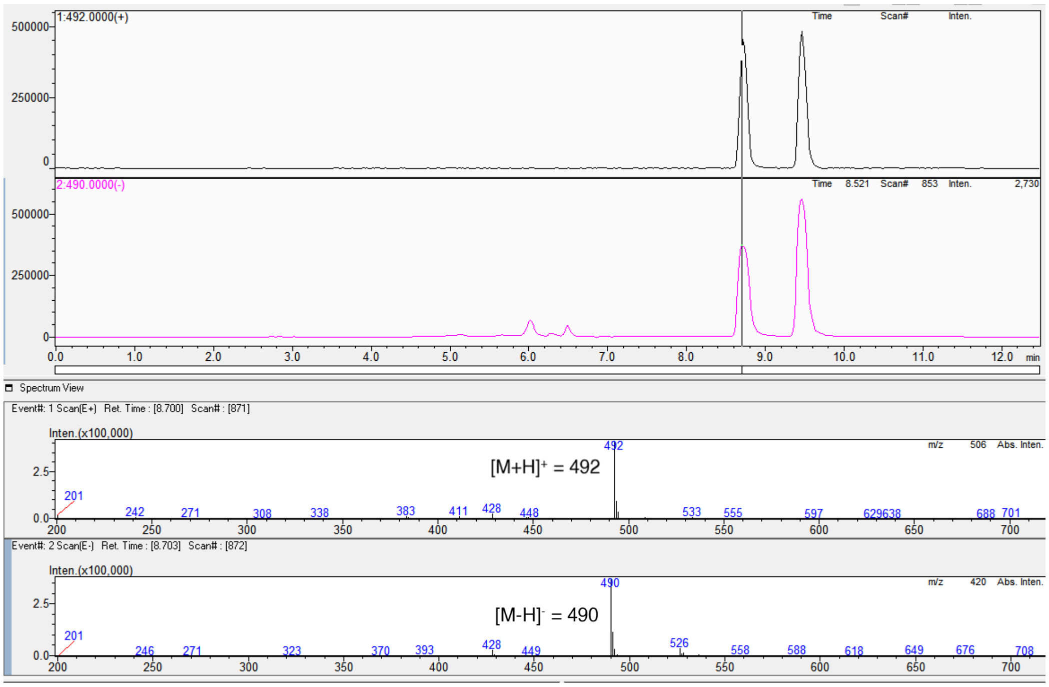

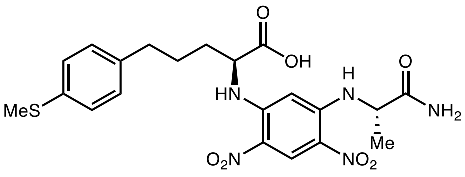

L-**3e** derivative

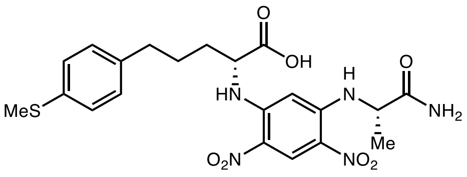

D-**3e** derivative

Marfey's analysis of L-**3e**

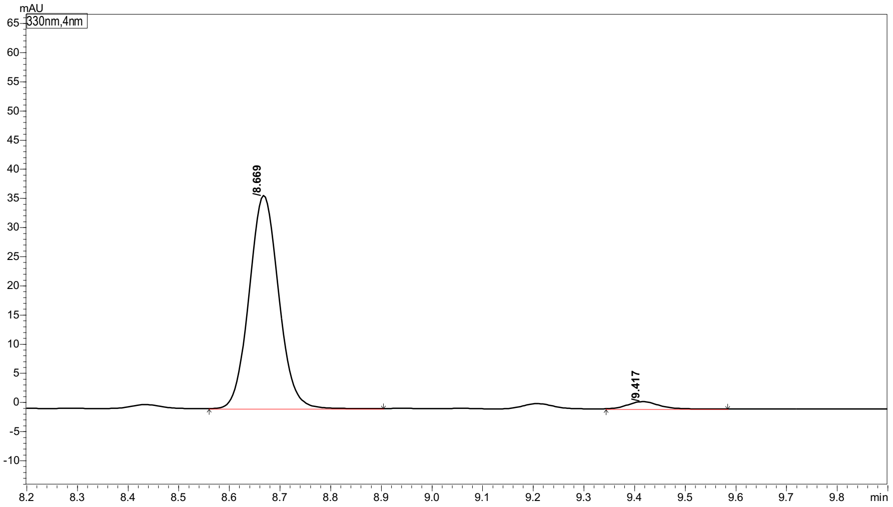

Peak Table | Compound | Group | Calibration Curve

| Peak# | Ret. Time | Height | Area | Area% |
|---|---|---|---|---|
| 1 | 8.673 | 31317 | 127772 | 96.595 |
| 2 | 9.422 | 1081 | 4504 | 3.405 |
| Total | | 32398 | 132276 | 100.000 |

Marfey's analysis of D-**3e**

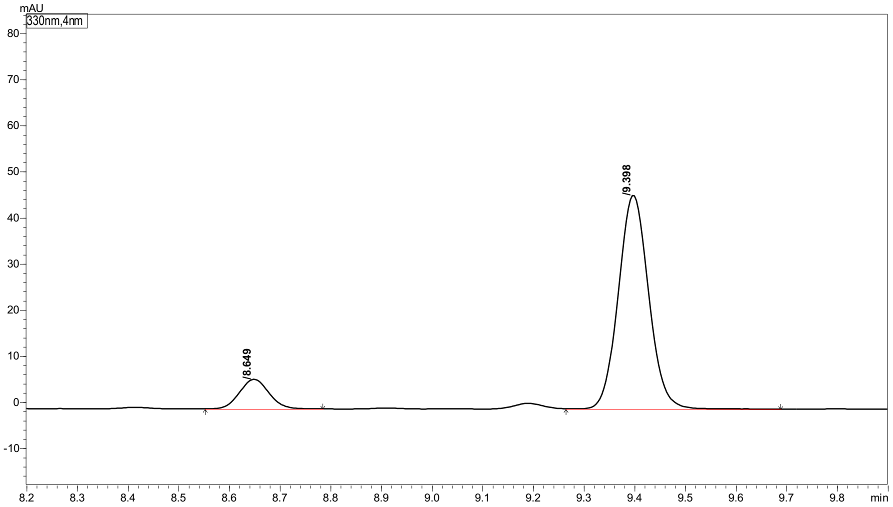

Peak Table | Compound | Group | Calibration Curve

| Peak# | Ret. Time | Height | Area | Area% |
|---|---|---|---|---|
| 1 | 8.649 | 6364 | 25837 | 11.820 |
| 2 | 9.398 | 46262 | 192757 | 88.180 |
| Total | | 52626 | 218594 | 100.000 |

#### 2-Amino-5-(4-fluorophenyl)pentanoic acid (3f)

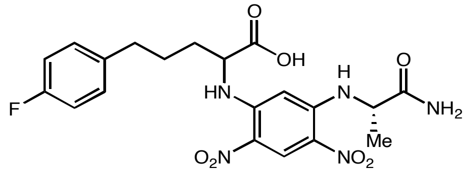

(*rac*)-**3f** derivative

Marfey's analysis of (*rac*)-**3f**

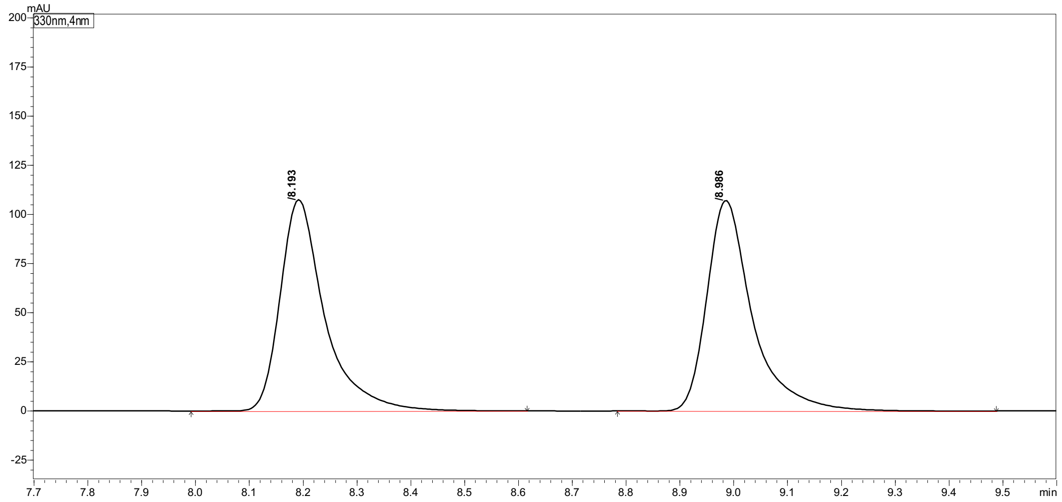

Peak Table | Compound | Group | Calibration Curve

| Peak# | Ret. Time | Height | Area | Area% |
|---|---|---|---|---|
| 1 | 8.193 | 106915 | 617635 | 49.614 |
| 2 | 8.986 | 106069 | 627241 | 50.386 |
| Total | | 212984 | 1244876 | 100.000 |

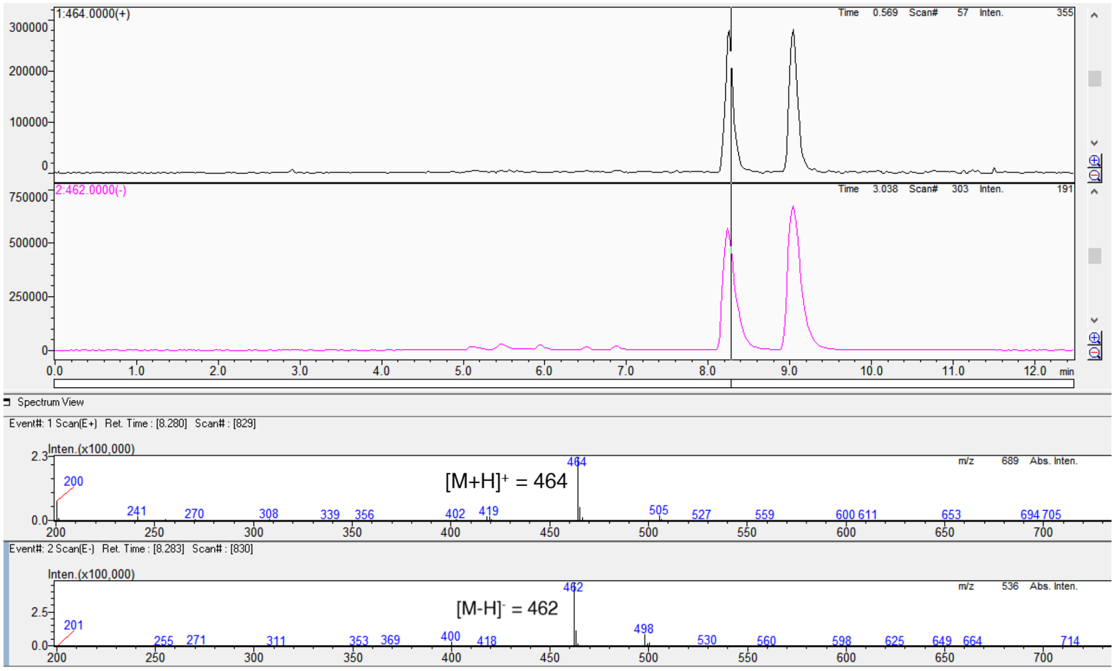

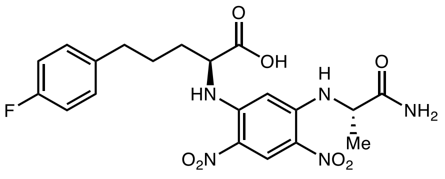

L-**3f** derivative

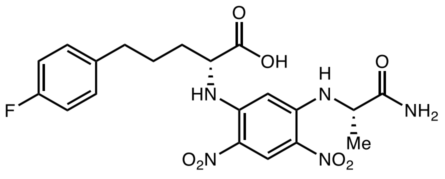

D-**3f** derivative

Marfey's analysis of L-**3f**

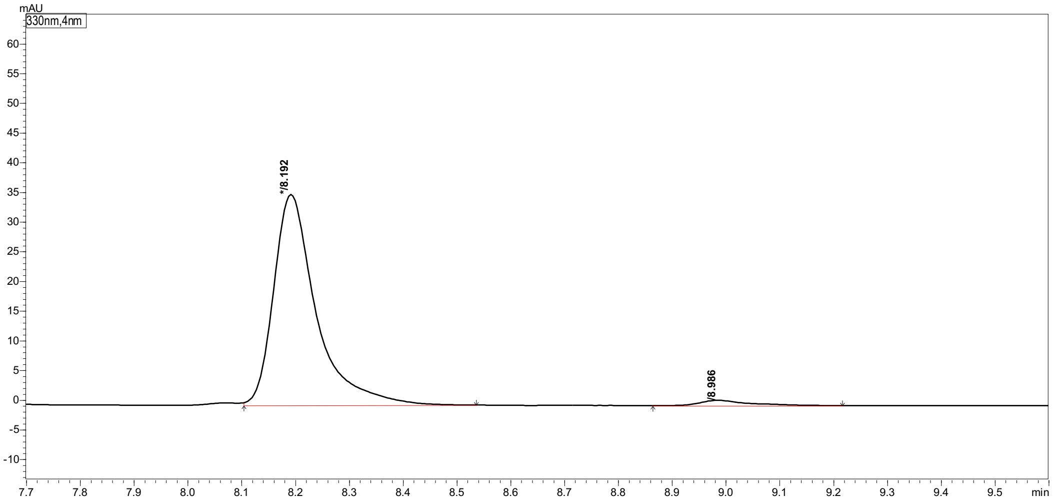

| Peak# | Ret. Time | Height | Area | Area% |
|---|---|---|---|---|
| 1 | 8.192 | 35322 | 194339 | 97.259 |
| 2 | 8.986 | 870 | 5477 | 2.741 |
| Total | | 36192 | 199816 | 100.000 |

Marfey's analysis of D-**3f**

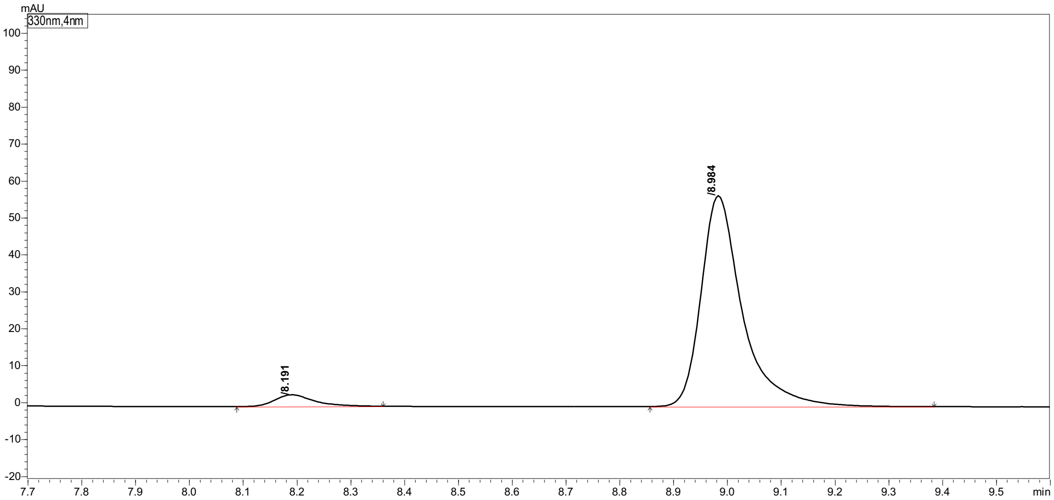

| Peak# | Ret. Time | Height | Area | Area% |
|---|---|---|---|---|
| 1 | 8.191 | 3136 | 15827 | 4.934 |
| 2 | 8.984 | 56859 | 304948 | 95.066 |
| Total | | 59996 | 320774 | 100.000 |

#### 2-Amino-5-(4-chlorophenyl)pentanoic acid (3g)

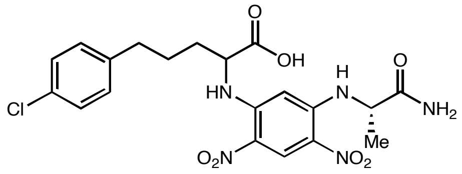

(*rac*)-**3g** derivative

#### Marfey's analysis of (*rac*)-**3g**

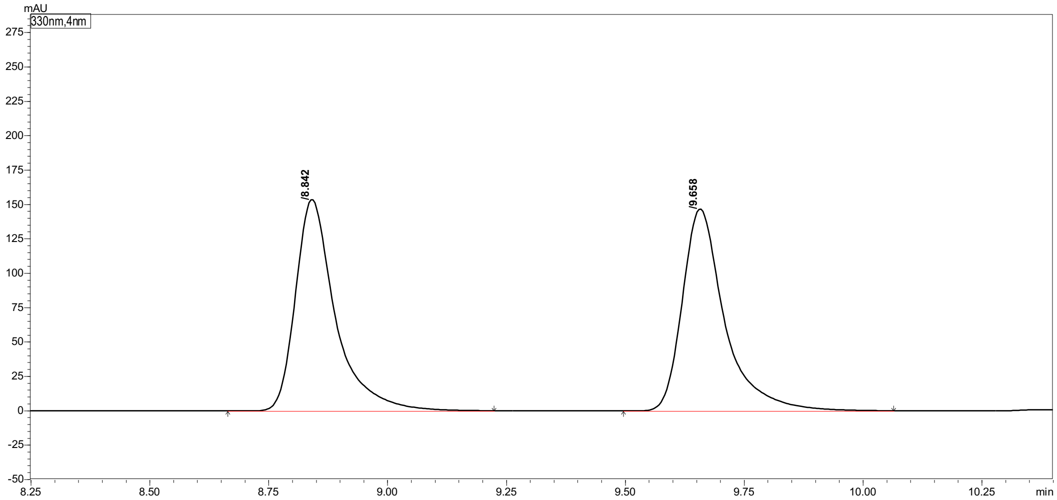

| Peak# | Ret. Time | Height | Area | Area% |
|---|---|---|---|---|
| 1 | 8.842 | 152249 | 895650 | 49.615 |
| 2 | 9.658 | 145559 | 909542 | 50.385 |
| Total | | 297808 | 1805192 | 100.000 |

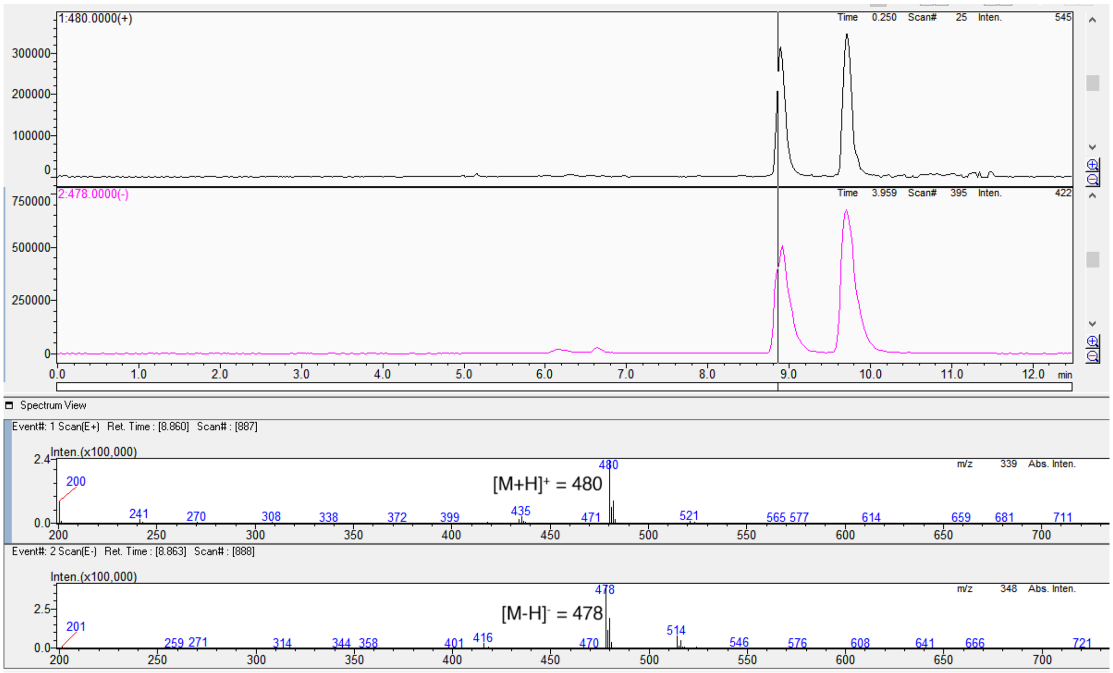

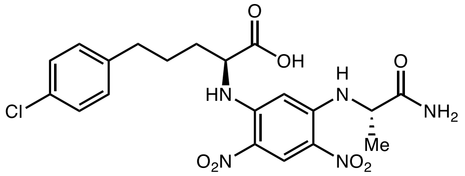

L-**3g** derivative

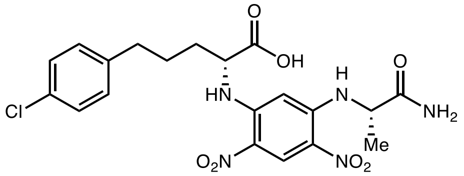

D-**3g** derivative

Marfey's analysis of L-**3g**

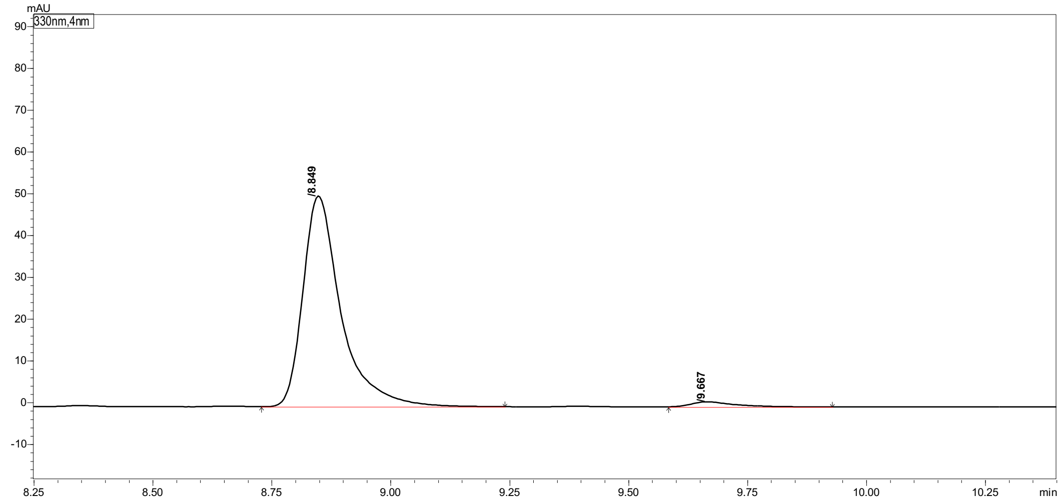

Peak Table | Compound | Group | Calibration Curve

| Peak# | Ret. Time | Height | Area | Area% |
|---|---|---|---|---|
| 1 | 8.849 | 50145 | 280333 | 97.436 |
| 2 | 9.667 | 1186 | 7377 | 2.564 |
| Total | | 51331 | 287709 | 100.000 |

Marfey's analysis of D-**3g**

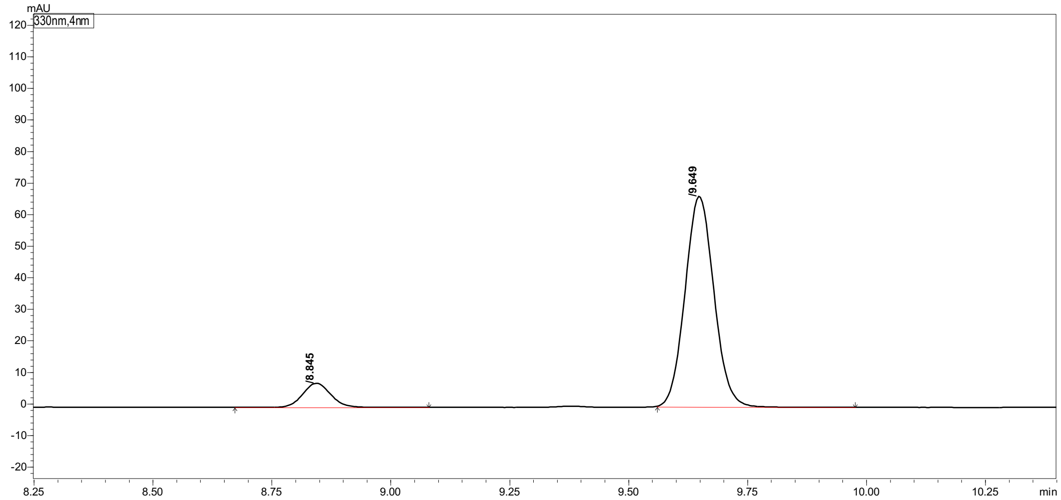

Peak Table | Compound | Group | Calibration Curve

| Peak# | Ret. Time | Height | Area | Area% |
|---|---|---|---|---|
| 1 | 8.814 | 6836 | 27481 | 9.813 |
| 2 | 9.617 | 58891 | 252562 | 90.187 |
| Total | | 65727 | 280043 | 100.000 |

#### 2-Amino-5-(4-bromophenyl)pentanoic acid (3h)

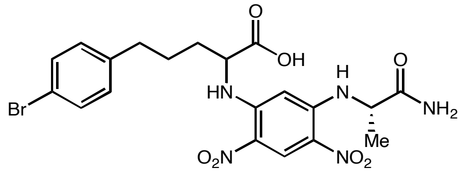

(*rac*)-**3h** derivative

Marfey's analysis of (*rac*)-**3h** (Long HPLC **method b** was used for separation)

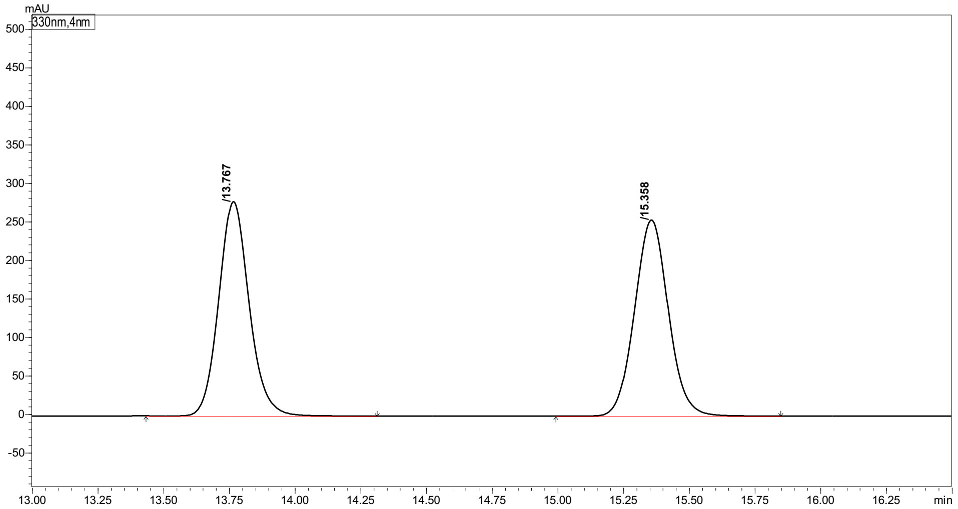

| Peak# | Ret. Time | Height | Area | Area% |
|---|---|---|---|---|
| 1 | 13.767 | 277768 | 2239218 | 49.644 |
| 2 | 15.358 | 254333 | 2271348 | 50.356 |
| Total | | 532101 | 4510566 | 100.000 |

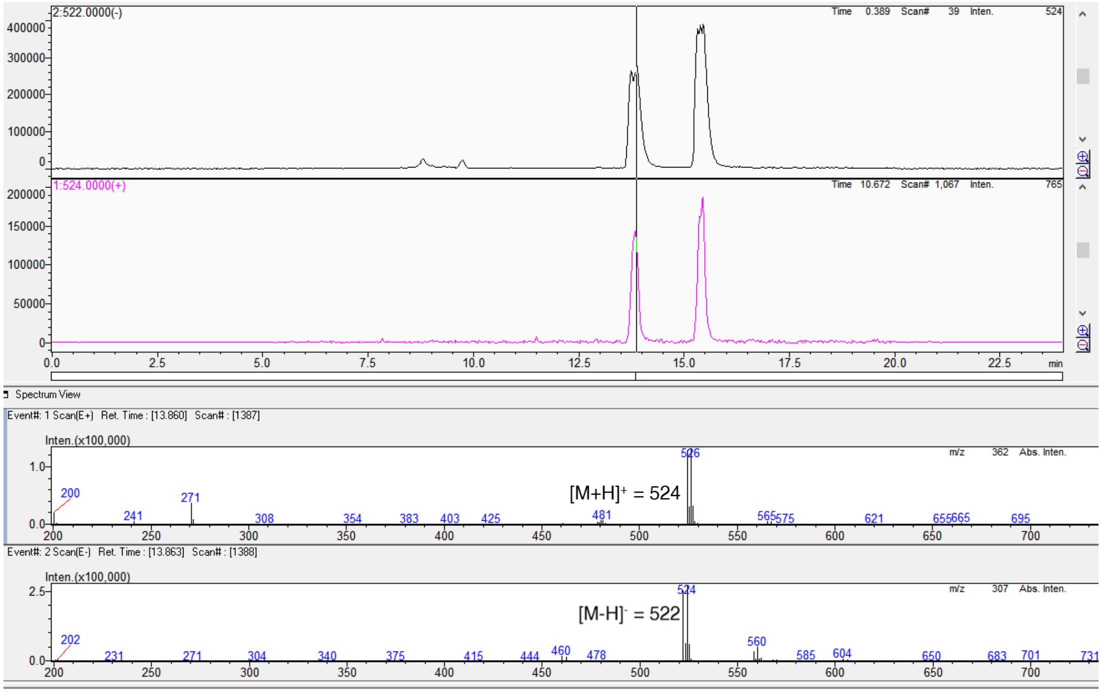

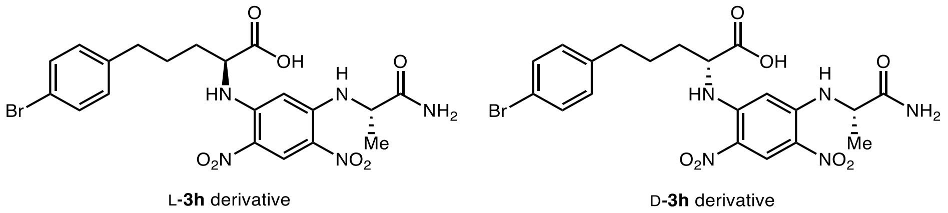

Marfey's analysis of L-**3h**

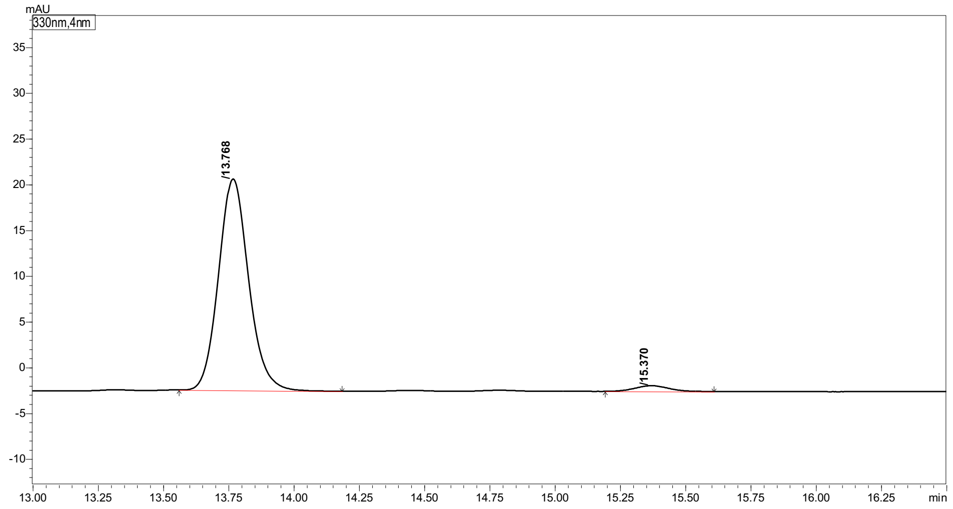

| Peak# | Ret. Time | Height | Area | Area% |
|---|---|---|---|---|
| 1 | 13.768 | 23087 | 185450 | 97.080 |
| 2 | 15.370 | 622 | 5577 | 2.920 |
| Total | | 23709 | 191027 | 100.000 |

Marfey's analysis of D-**3h**

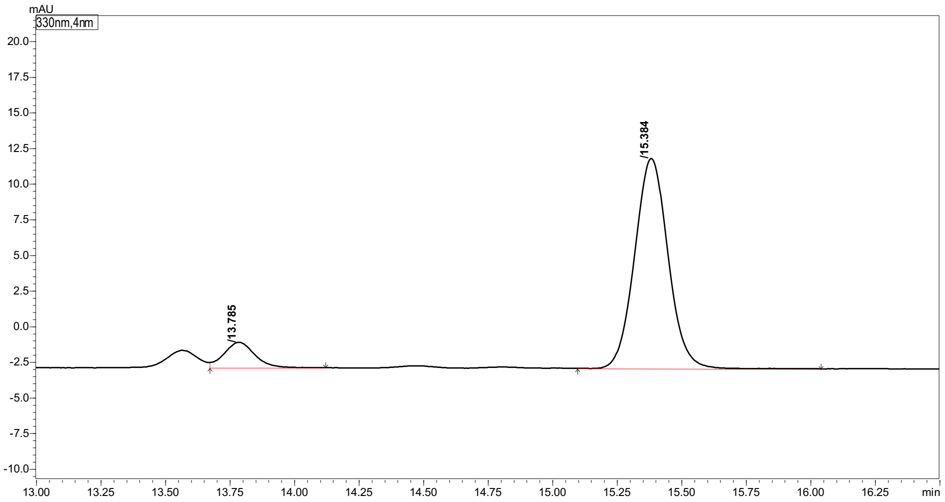

| Peak# | Ret. Time | Height | Area | Area% |
|---|---|---|---|---|
| 1 | 13.785 | 1779 | 14735 | 10.008 |
| 2 | 15.384 | 14720 | 132488 | 89.992 |
| Total | | 16499 | 147223 | 100.000 |

#### 2-Amino-5-(4-cyanophenyl)pentanoic acid (3i)

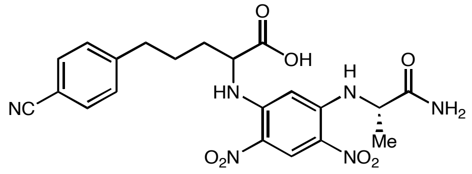

(*rac*)-**3i** derivative

#### Marfey's analysis of (*rac*)-**3i**

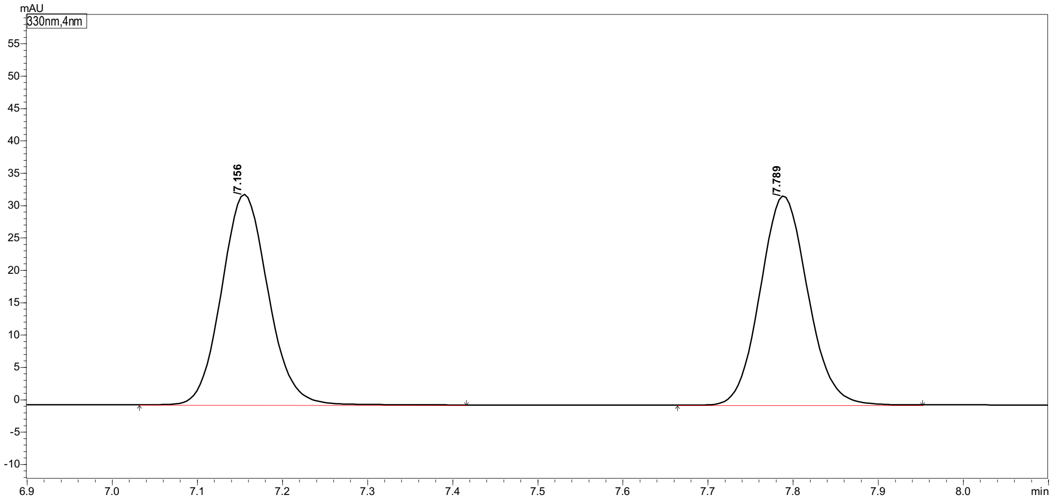

| Peak# | Ret. Time | Height | Area | Area% |
|---|---|---|---|---|
| 1 | 7.156 | 32255 | 123454 | 49.559 |
| 2 | 7.789 | 32215 | 125653 | 50.441 |
| Total | | 64471 | 249108 | 100.000 |

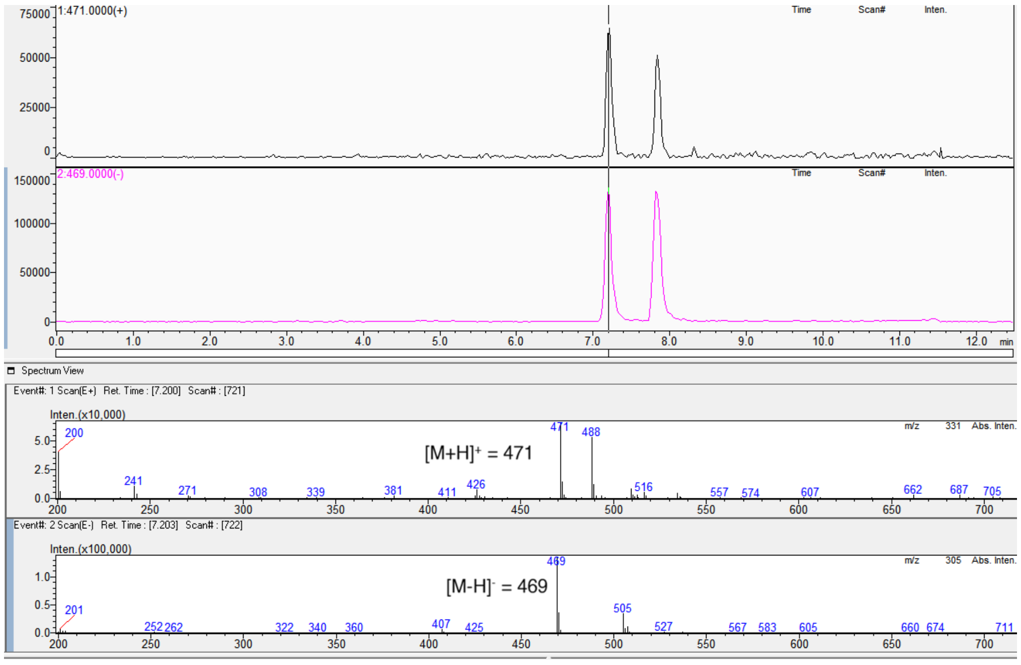

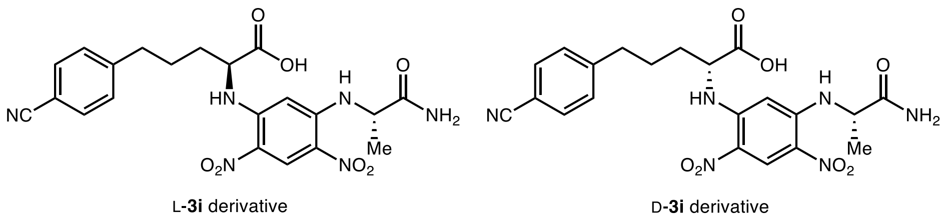

Marfey's analysis of L-**3i**

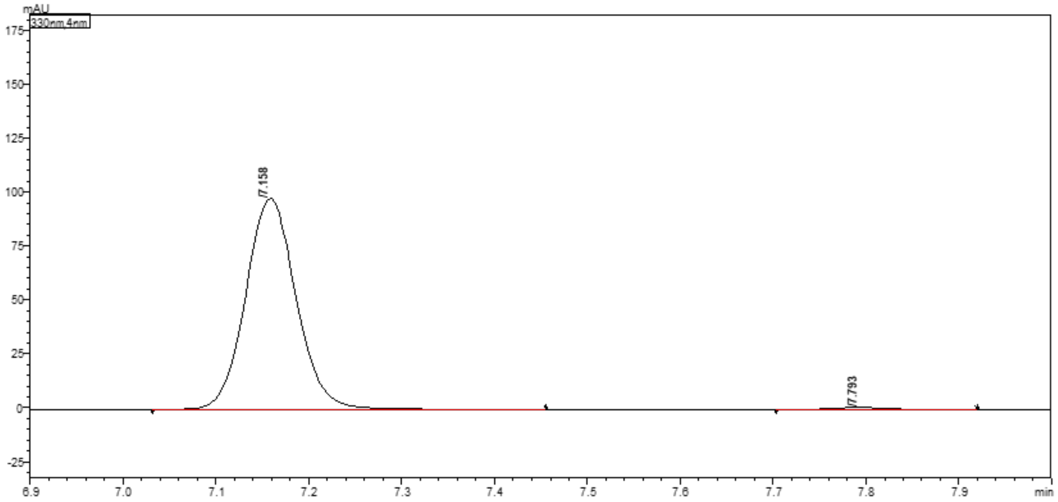

| Peak# | Ret. Time | Height | Area | Area% |
|---|---|---|---|---|
| 1 | 7.158 | 97488 | 368194 | 99.295 |
| 2 | 7.793 | 707 | 2616 | 0.705 |
| Total | | 98194 | 370810 | 100.000 |

Marfey's analysis of D-**3i**

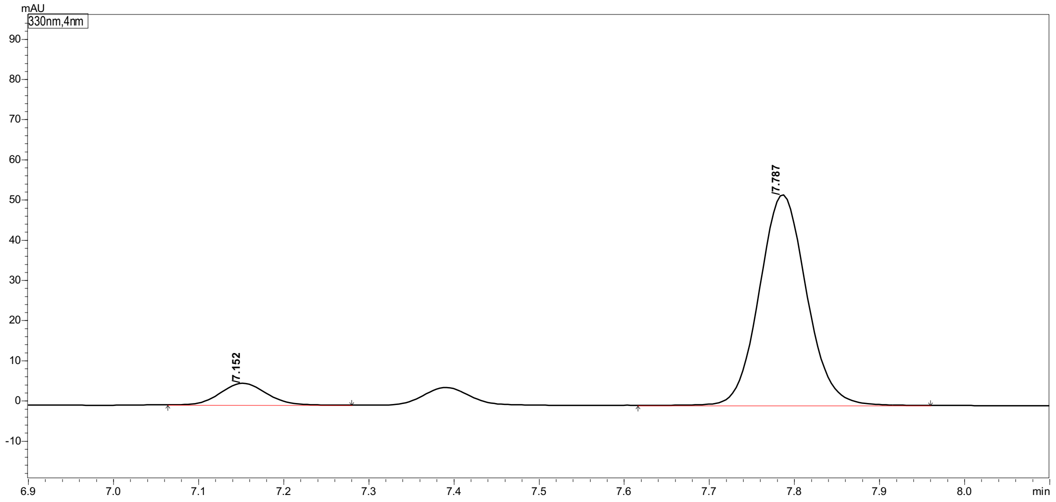

| Peak# | Ret. Time | Height | Area | Area% |
|---|---|---|---|---|
| 1 | 7.152 | 5369 | 20063 | 9.050 |
| 2 | 7.787 | 51766 | 201637 | 90.950 |
| Total | | 57135 | 221700 | 100.000 |

#### 2-Amino-5-(4-(trifluoromethyl)phenyl)pentanoic acid (3j)

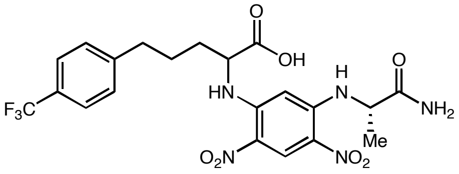

(*rac*)-**3j** derivative

#### Marfey's analysis of (*rac*)-**3j**

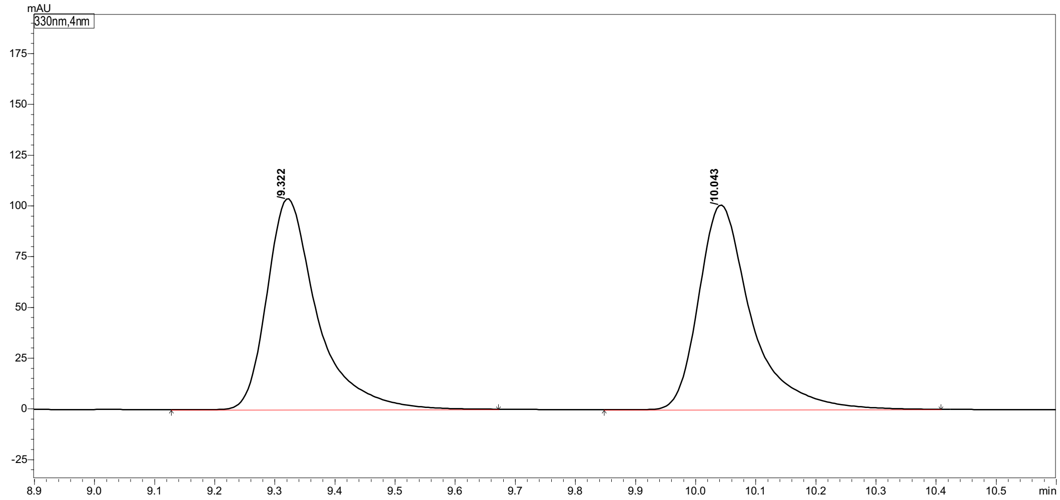

| Peak# | Ret. Time | Height | Area | Area% |
|---|---|---|---|---|
| 1 | 9.322 | 103055 | 601670 | 49.543 |
| 2 | 10.043 | 100235 | 612779 | 50.457 |
| Total | | 203290 | 1214449 | 100.000 |

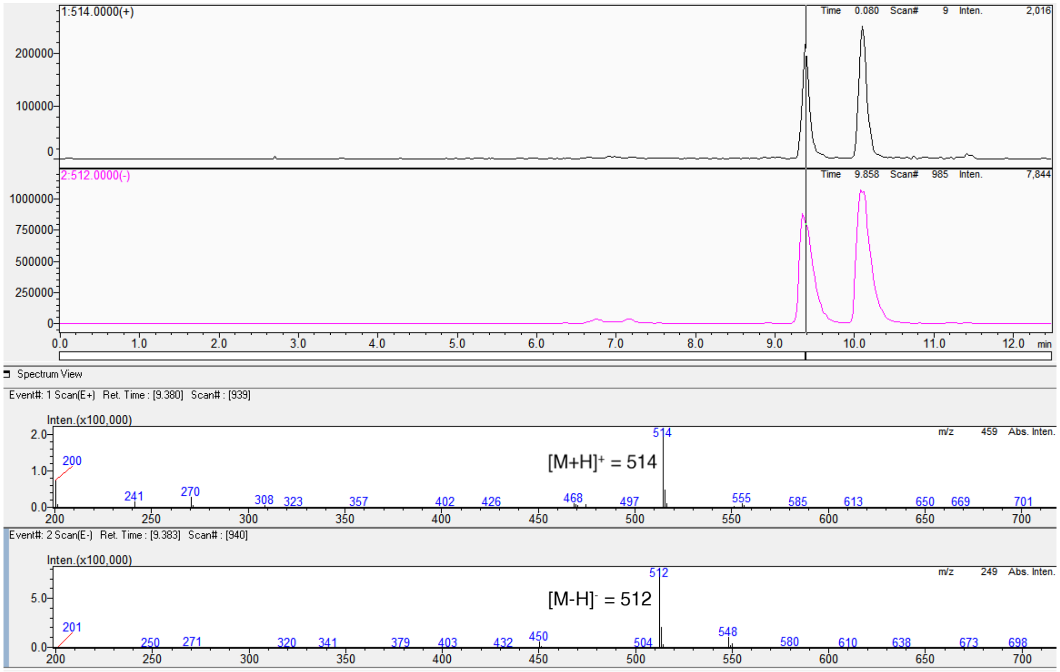

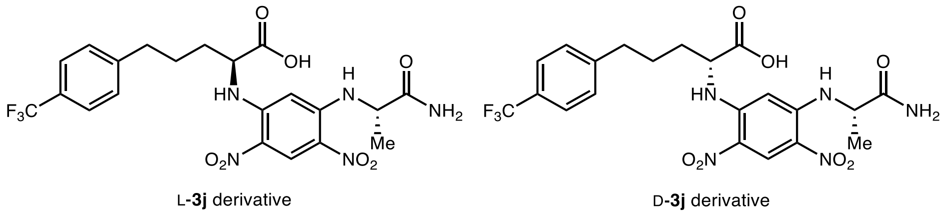

Marfey's analysis of L-**3j**

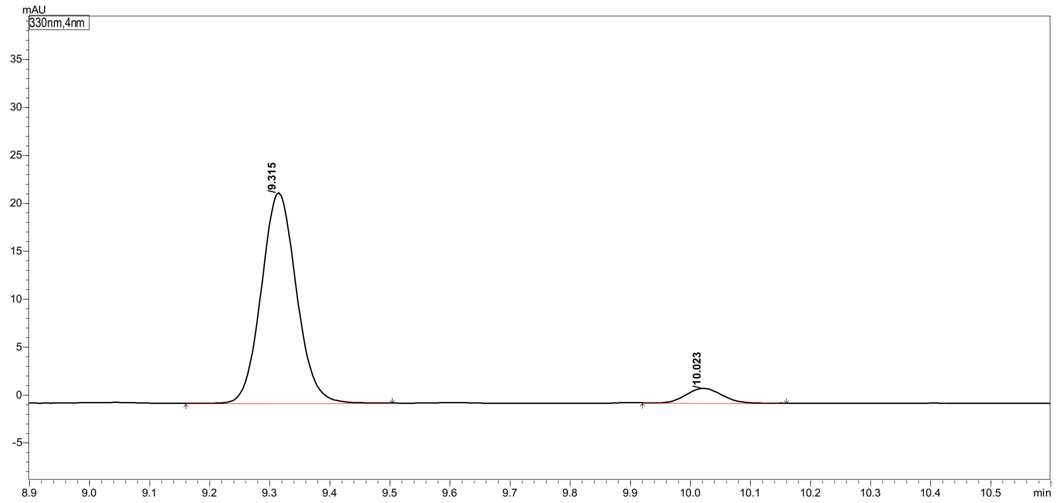

| Peak# | Ret. Time | Height | Area | Area% |
|---|---|---|---|---|
| 1 | 9.315 | 21774 | 88858 | 93.621 |
| 2 | 10.023 | 1514 | 6054 | 6.379 |
| Total | | 23288 | 94912 | 100.000 |

Marfey's analysis of D-**3j**

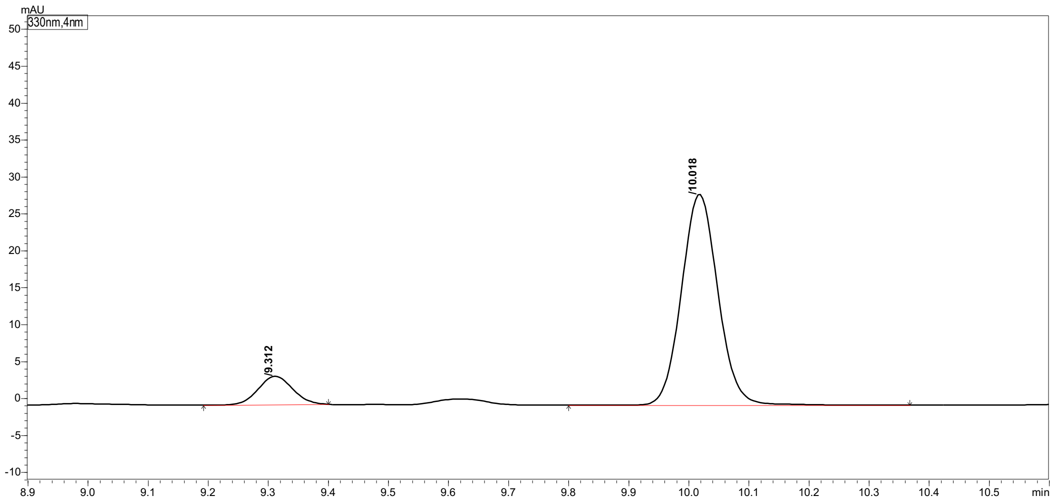

| Peak# | Ret. Time | Height | Area | Area% |
|---|---|---|---|---|
| 1 | 9.312 | 3809 | 15166 | 11.212 |
| 2 | 10.018 | 28232 | 120098 | 88.788 |
| Total | | 32042 | 135265 | 100.000 |

#### 2-Amino-5-(m-tolyl)pentanoic acid (3k)

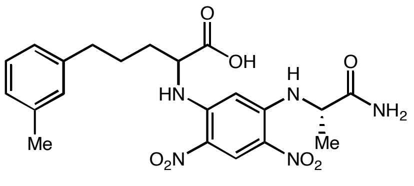

(*rac*)-**3k** derivative

Marfey's analysis of (*rac*)-**3k**

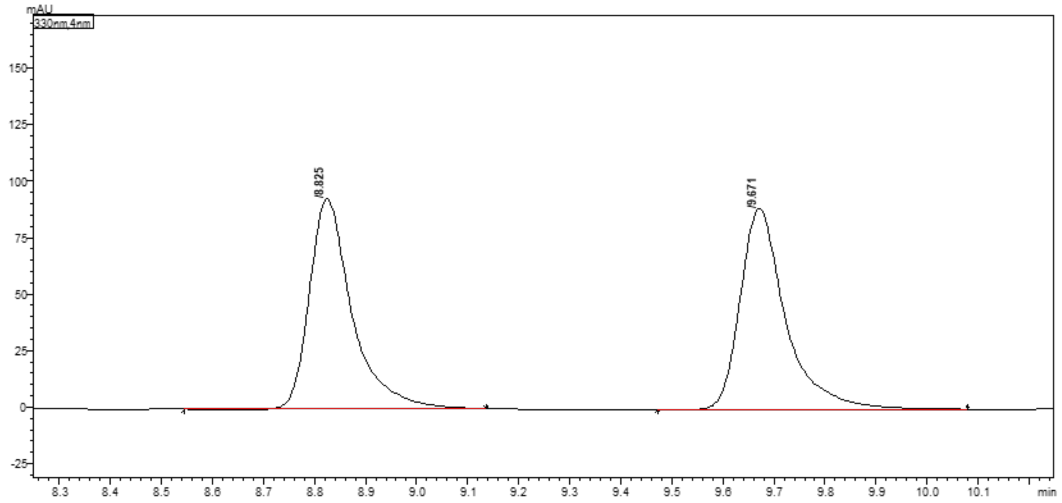

| Peak# | Ret. Time | Height | Area | Area% |
|---|---|---|---|---|
| 1 | 8.825 | 92540 | 537214 | 49.580 |
| 2 | 9.671 | 88578 | 546322 | 50.420 |
| Total | | 181118 | 1083536 | 100.000 |

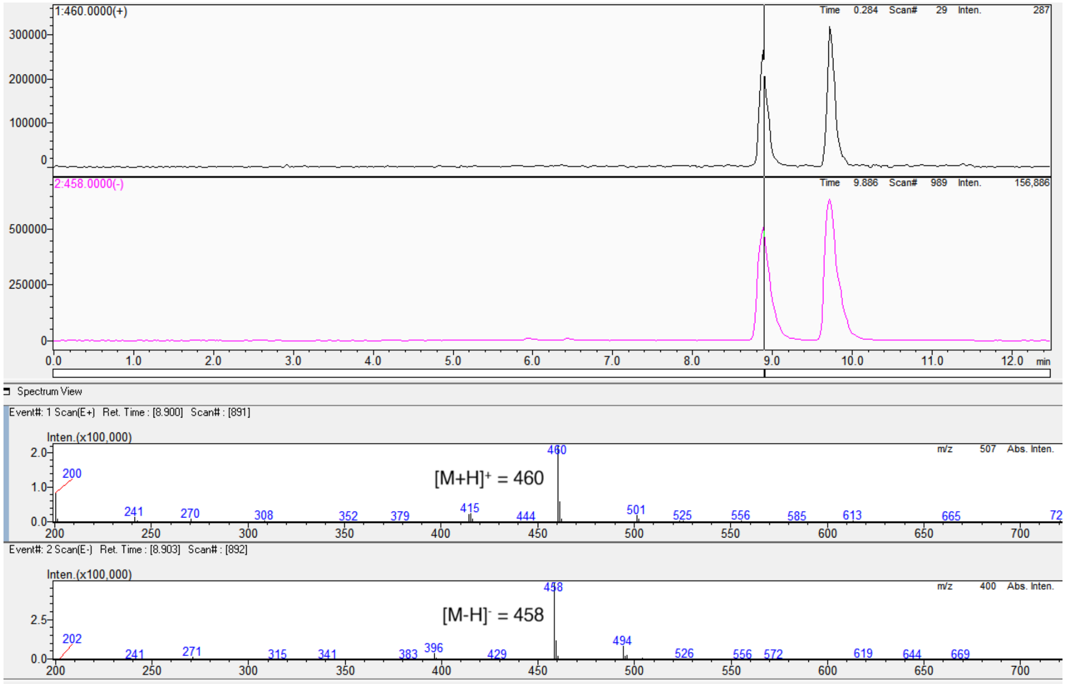

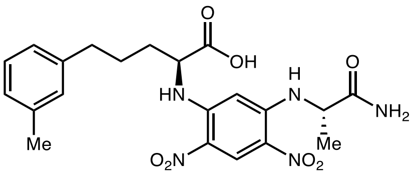

L-**3k** derivative

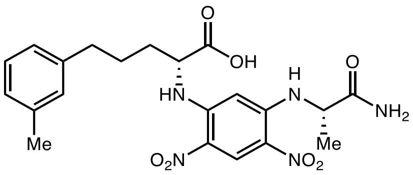

D-**3k** derivative

#### Marfey's analysis of L-**3k**

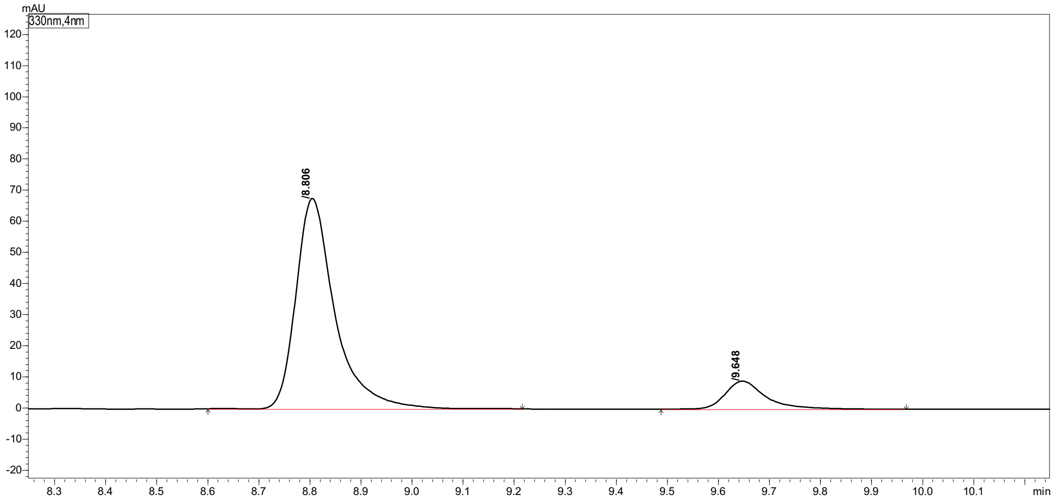

Peak Table | Compound | Group | Calibration Curve

| Peak# | Ret. Time | Height | Area | Area% |
|---|---|---|---|---|
| 1 | 8.806 | 67534 | 358698 | 87.824 |
| 2 | 9.648 | 8970 | 49732 | 12.176 |
| Total | | 76504 | 408430 | 100.000 |

#### Marfey's analysis of D-**3k**

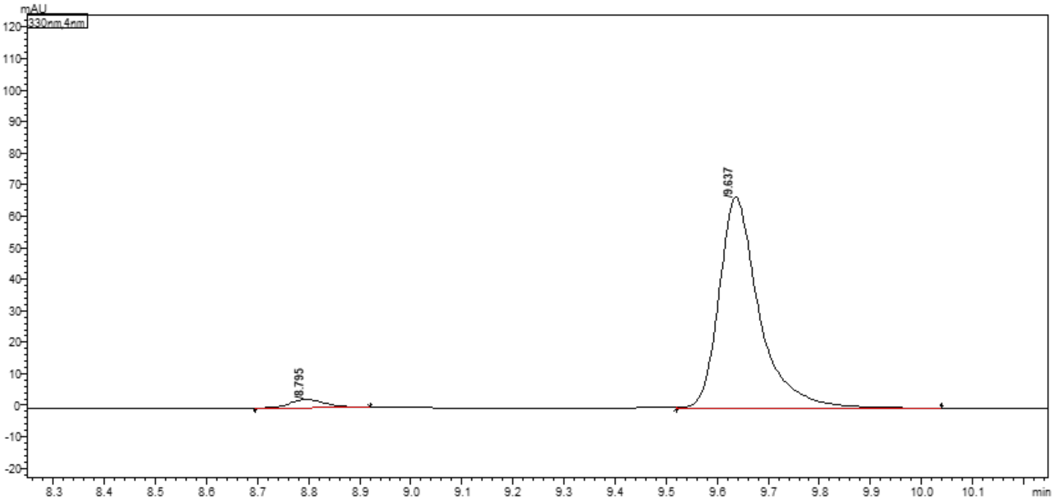

Peak Table | Compound | Group | Calibration Curve

| Peak# | Ret. Time | Height | Area | Area% |
|---|---|---|---|---|
| 1 | 8.795 | 2556 | 11899 | 3.153 |
| 2 | 9.637 | 66658 | 365482 | 96.847 |
| Total | | 69215 | 377380 | 100.000 |

#### 2-Amino-5-(o-tolyl)pentanoic acid (3l)

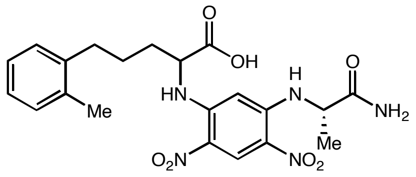

(*rac*)-**3l** derivative

#### Marfey's analysis of (*rac*)-**3l**

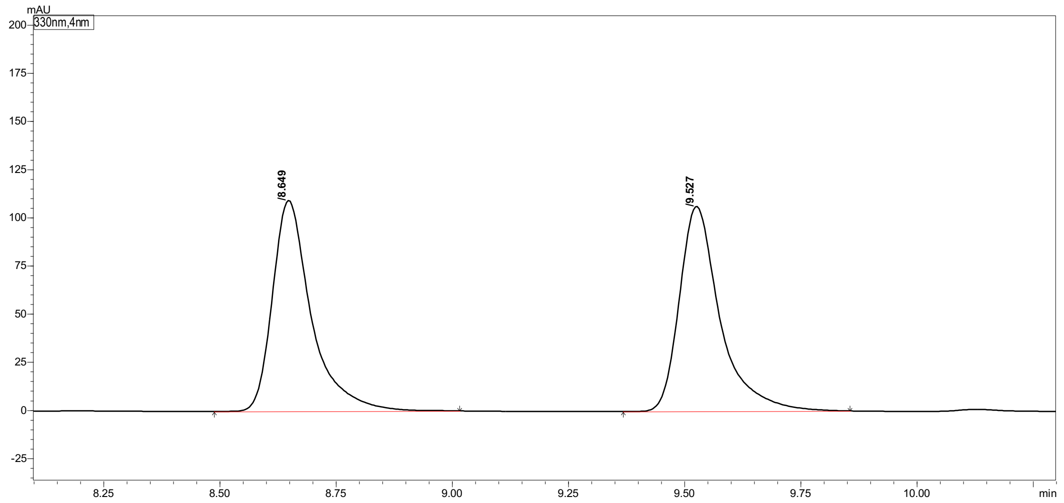

| Peak# | Ret. Time | Height | Area | Area% |
|---|---|---|---|---|
| 1 | 8.649 | 108899 | 634044 | 49.385 |
| 2 | 9.527 | 106273 | 649839 | 50.615 |
| Total | | 215172 | 1283883 | 100.000 |

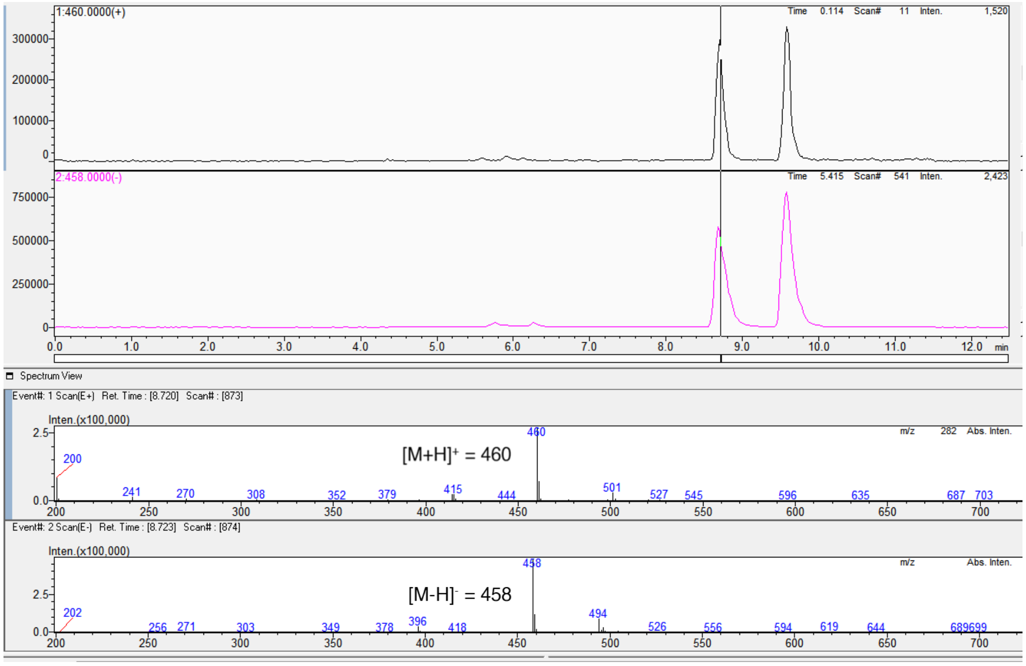

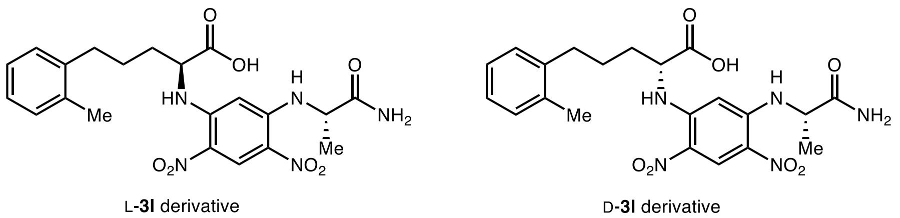

Marfey's analysis of L-**3l**

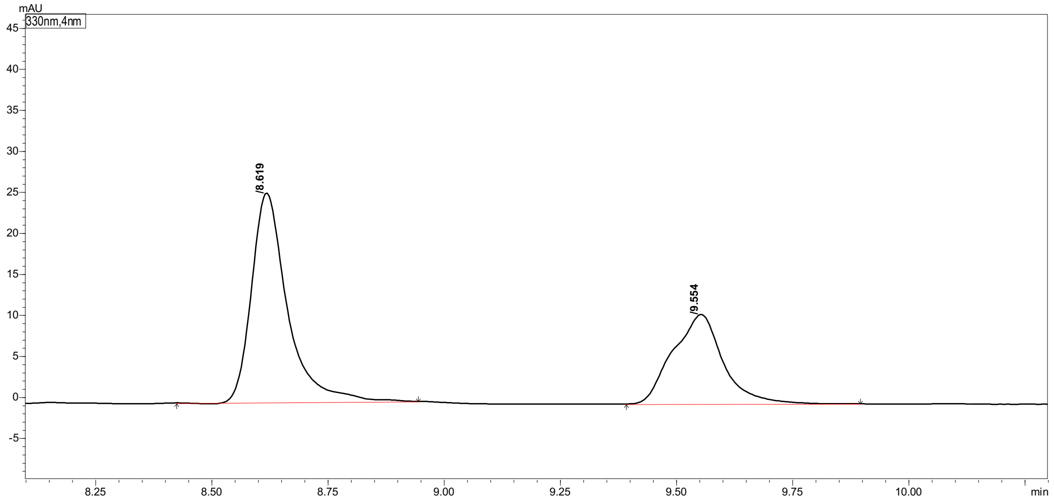

| Peak# | Ret. Time | Height | Area | Area% |
|---|---|---|---|---|
| 1 | 8.619 | 25324 | 136243 | 62.497 |
| 2 | 9.554 | 10838 | 81757 | 37.503 |
| Total | | 36162 | 218000 | 100.000 |

Marfey's analysis of D-**3l**

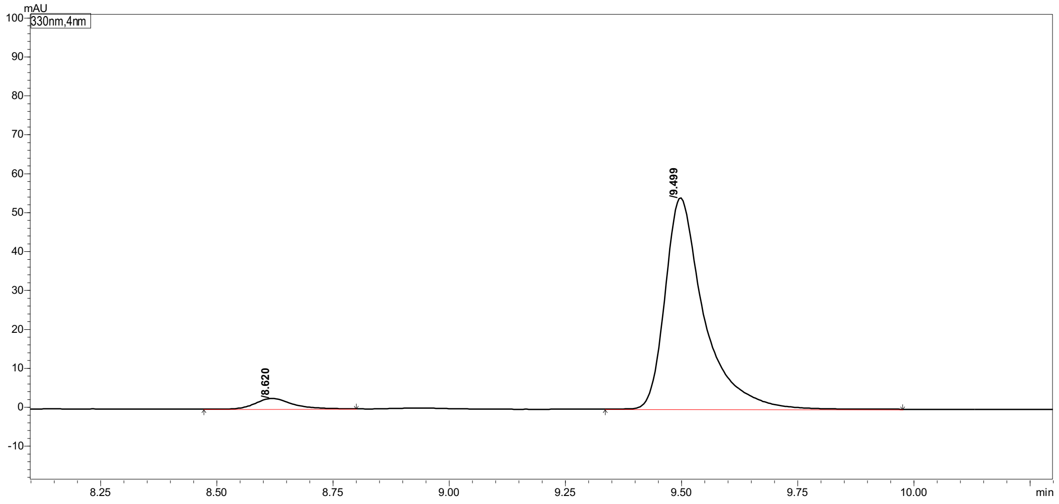

| Peak# | Ret. Time | Height | Area | Area% |
|---|---|---|---|---|
| 1 | 8.620 | 2712 | 14511 | 4.390 |
| 2 | 9.499 | 53898 | 316067 | 95.610 |
| Total | | 56609 | 330578 | 100.000 |

#### 2-Amino-5-(furan-2-yl)pentanoic acid (3m)

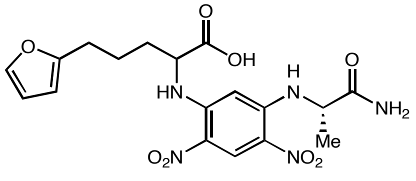

(*rac*)-**3m** derivative

##### Marfey's analysis of (*rac*)-**3m**

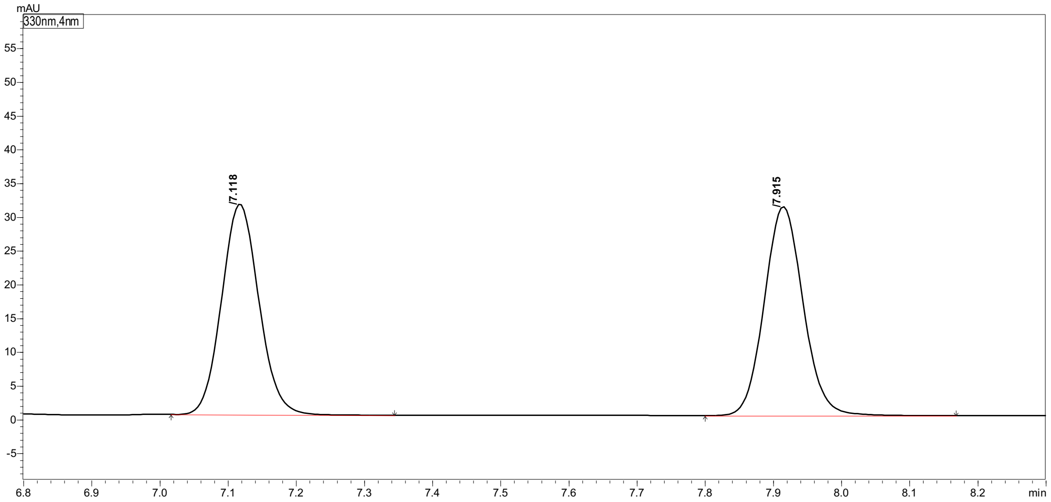

Peak Table | Compound | Group | Calibration Curve

| Peak# | Ret. Time | Height | Area | Area% |
|---|---|---|---|---|
| 1 | 7.118 | 31100 | 118546 | 49.183 |
| 2 | 7.915 | 30673 | 122487 | 50.817 |
| Total | | 61773 | 241033 | 100.000 |

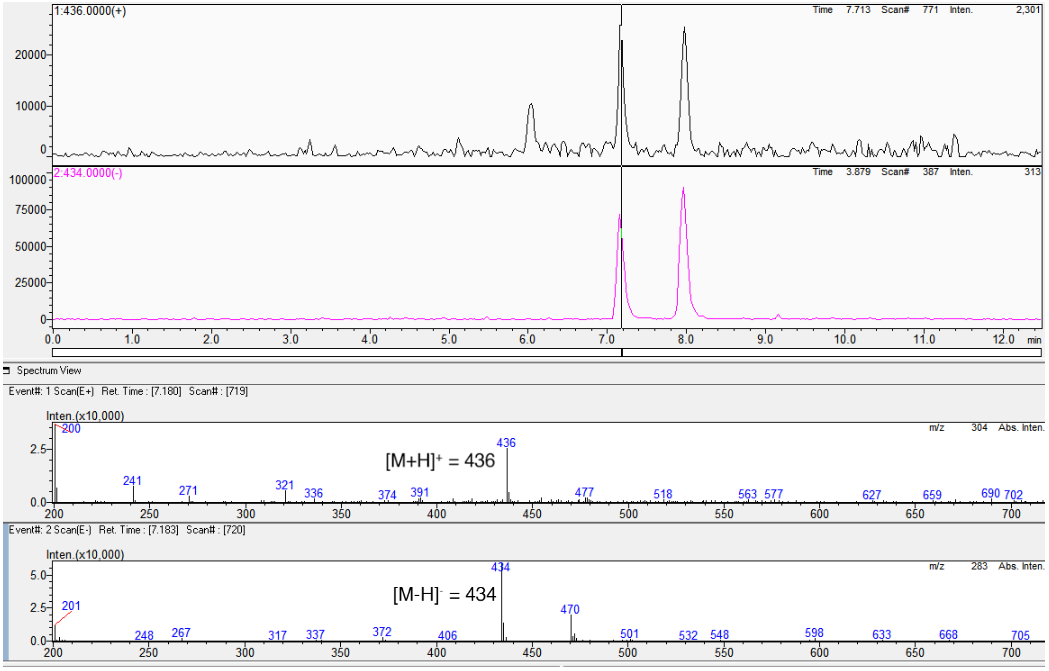

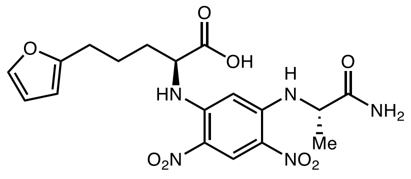

L-**3m** derivative

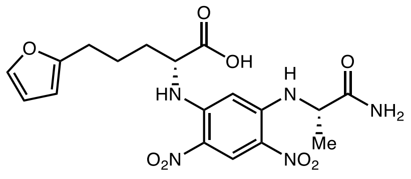

D-**3m** derivative

Marfey's analysis of L-**3m**

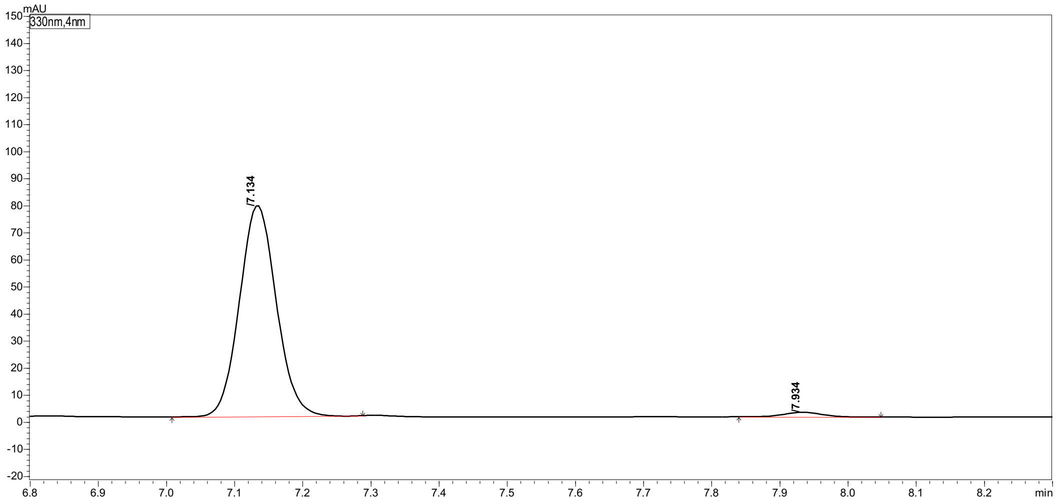

| Peak# | Ret. Time | Height | Area | Area% |
|---|---|---|---|---|
| 1 | 7.134 | 77696 | 292436 | 97.847 |
| 2 | 7.934 | 1698 | 6435 | 2.153 |
| Total | | 79394 | 298871 | 100.000 |

Marfey's analysis of D-**3m**

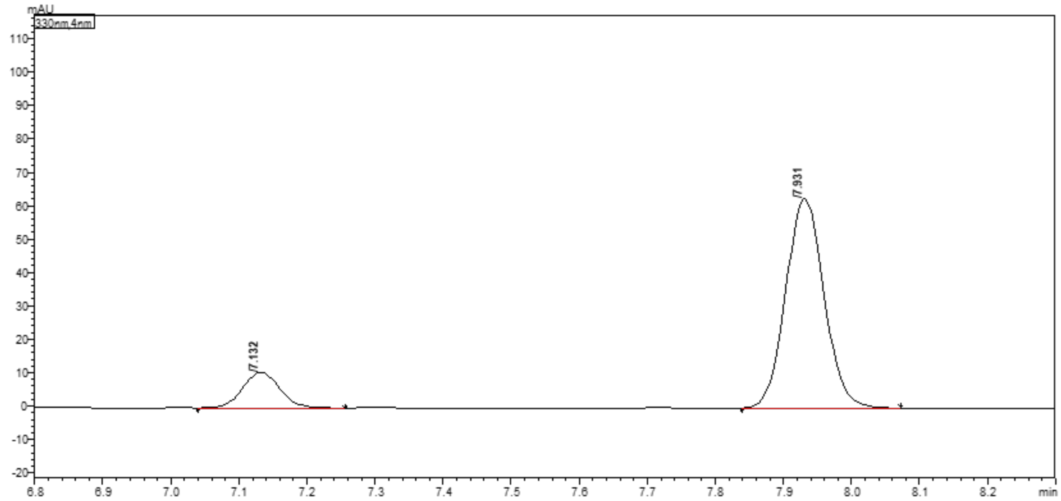

| Peak# | Ret. Time | Height | Area | Area% |
|---|---|---|---|---|
| 1 | 7.132 | 10800 | 41707 | 14.536 |
| 2 | 7.931 | 62306 | 245208 | 85.464 |
| Total | | 73106 | 286915 | 100.000 |

#### 2-Amino-5-(thiophen-2-yl)pentanoic acid (3n)

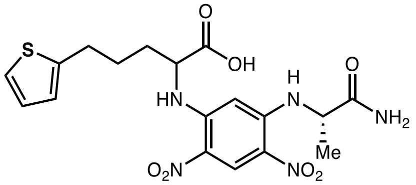

(*rac*)-**3n** derivative

#### Marfey's analysis of (*rac*)-**3n**

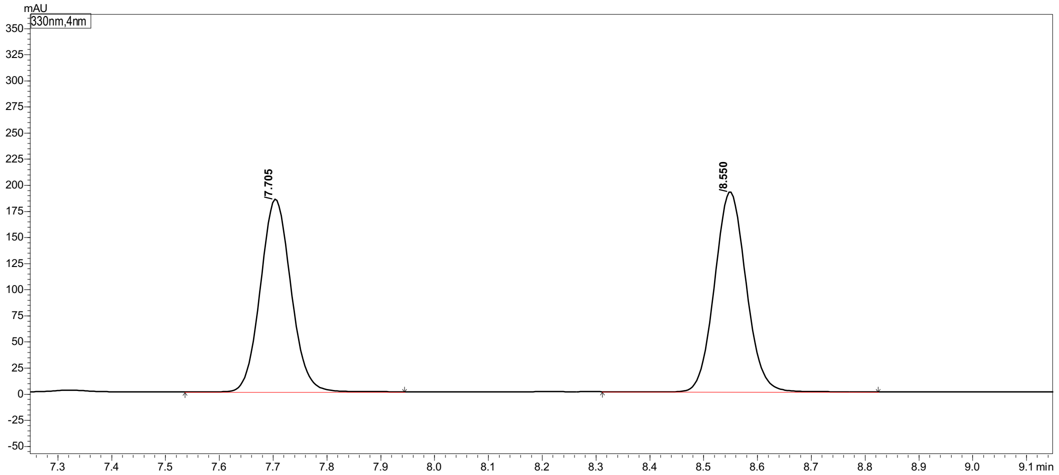

Peak Table | Compound | Group | Calibration Curve

| Peak# | Ret. Time | Height | Area | Area% |
|---|---|---|---|---|
| 1 | 7.705 | 182626 | 726447 | 48.329 |
| 2 | 8.550 | 191055 | 776679 | 51.671 |
| Total | | 373681 | 1503126 | 100.000 |

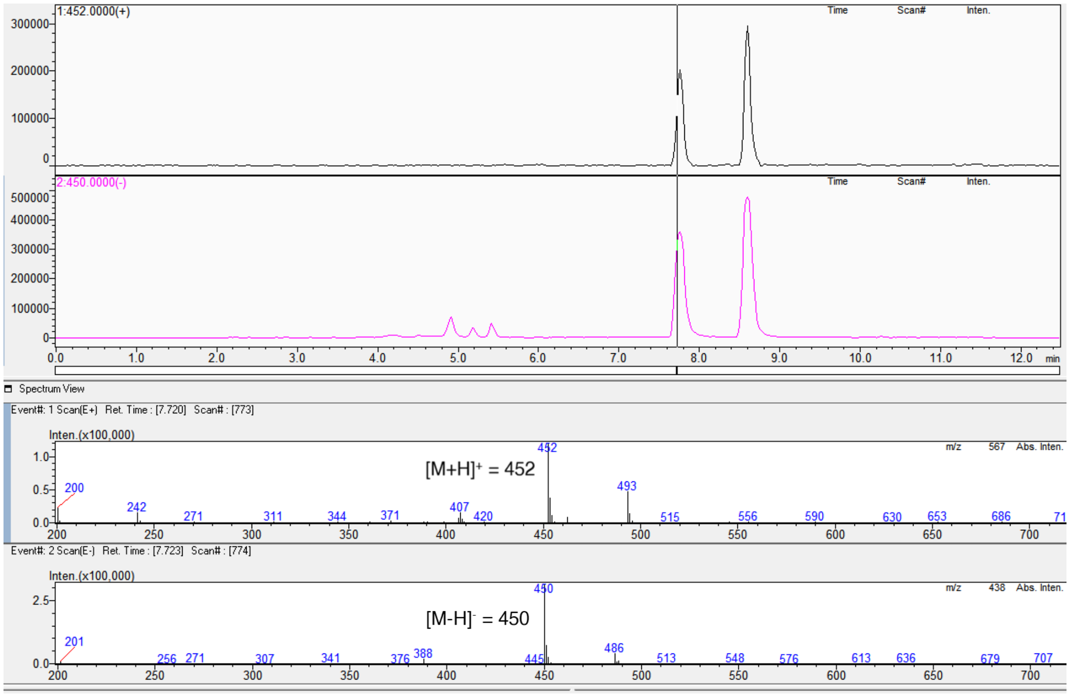

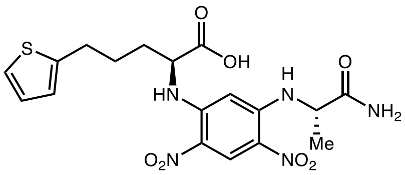

L-**3n** derivative

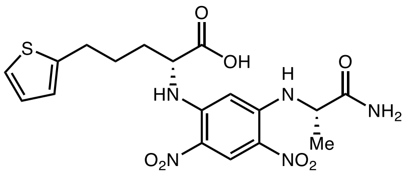

D-**3n** derivative

Marfey's analysis of L-**3n**

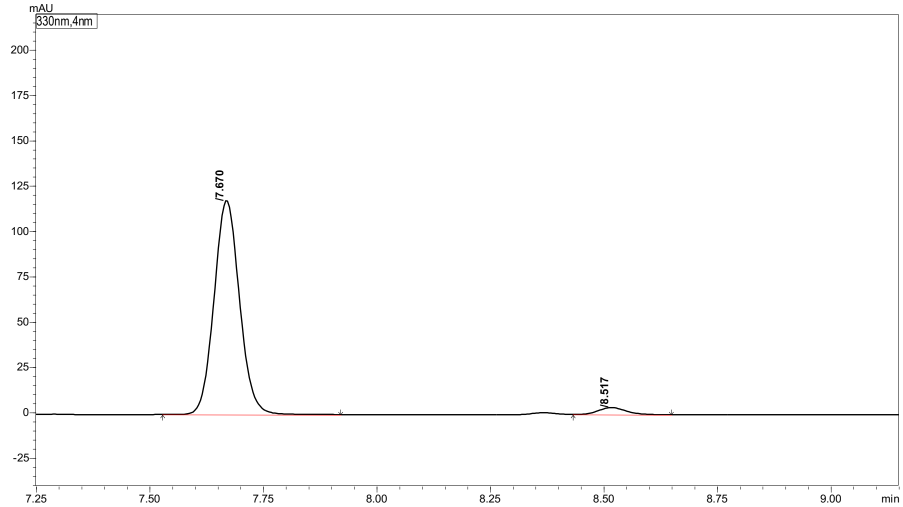

| Peak# | Ret. Time | Height | Area | Area% |
|---|---|---|---|---|
| 1 | 7.670 | 117716 | 459066 | 96.618 |
| 2 | 8.517 | 3839 | 16069 | 3.382 |
| Total | | 121556 | 475136 | 100.000 |

Marfey's analysis of D-**3n**

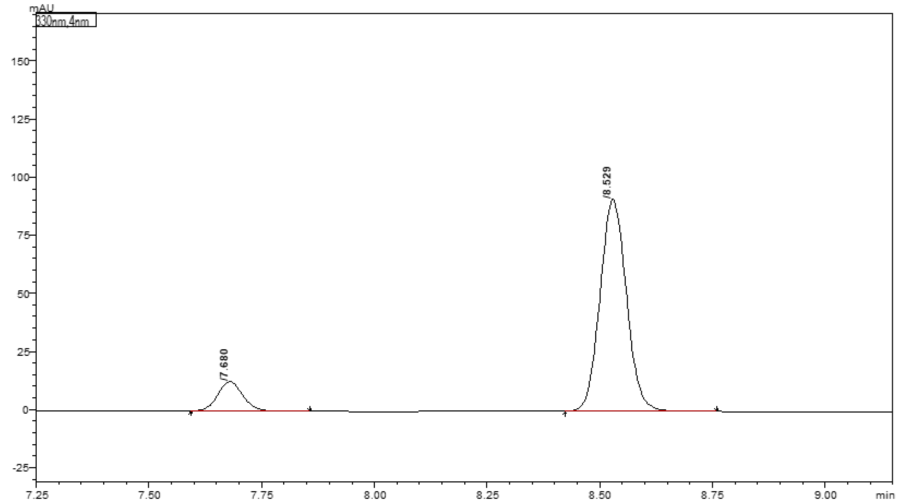

| Peak# | Ret. Time | Height | Area | Area% |
|---|---|---|---|---|
| 1 | 7.680 | 12470 | 48344 | 11.492 |
| 2 | 8.529 | 90561 | 372319 | 88.508 |
| Total | | 103030 | 420663 | 100.000 |

#### 2-Amino-5-(6-methoxypyridin-3-yl)pentanoic acid (3o)

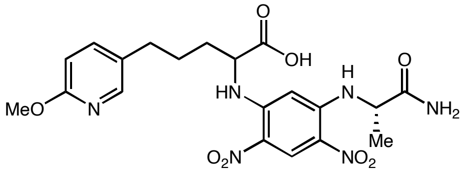

(*rac*)-**3p** derivative

#### Marfey's analysis of (*rac*)-**3o**

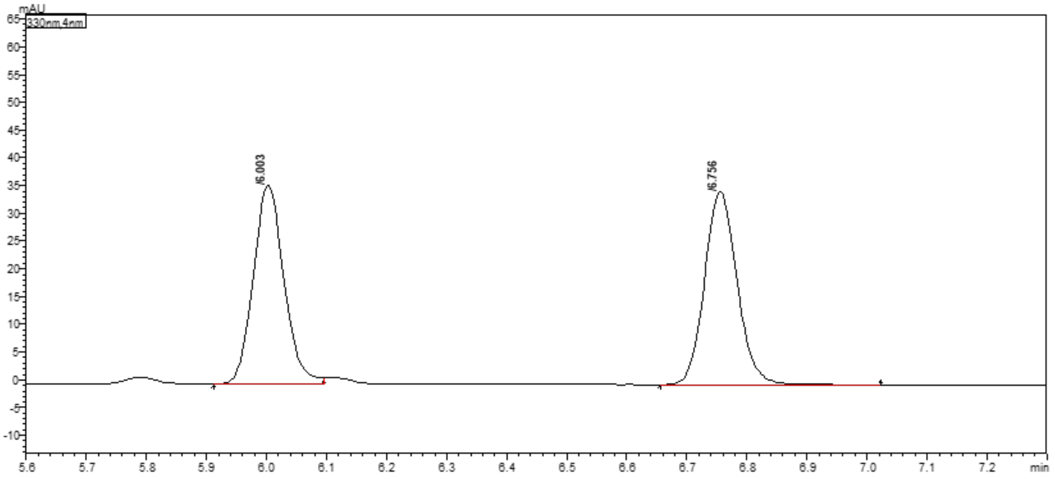

Peak Table | Compound | Group | Calibration Curve

| Peak# | Ret. Time | Height | Area | Area% |
|---|---|---|---|---|
| 1 | 6.003 | 35330 | 128219 | 49.478 |
| 2 | 6.756 | 34611 | 130925 | 50.522 |
| Total | | 69940 | 259144 | 100.000 |

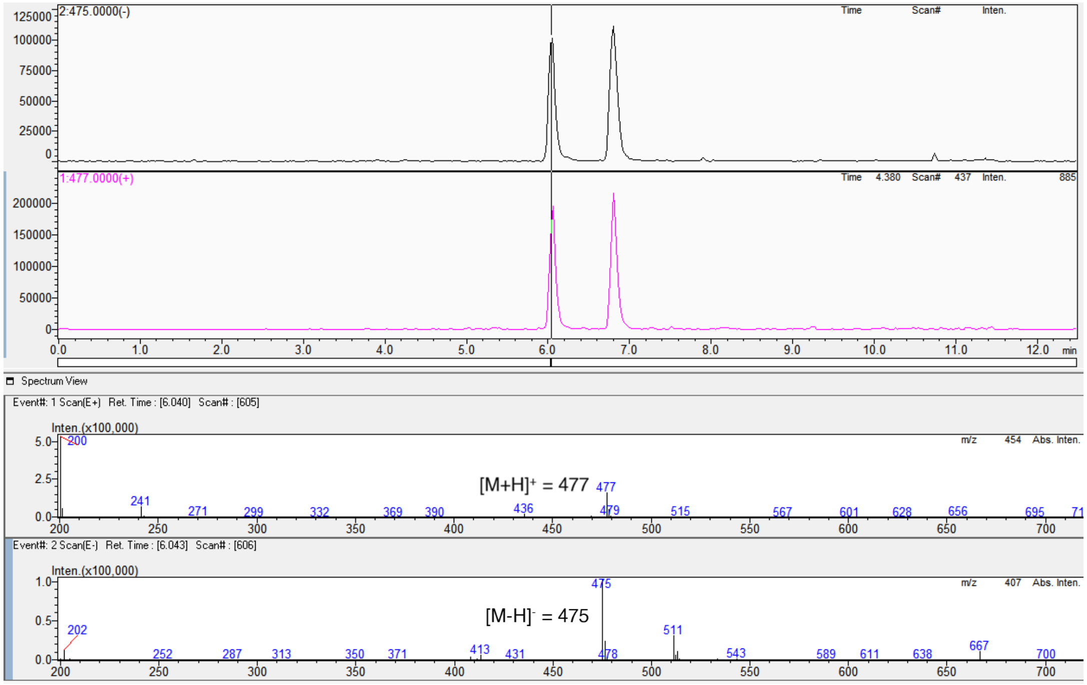

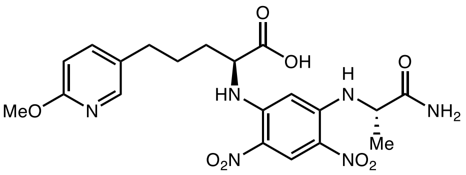

L-**3p** derivative

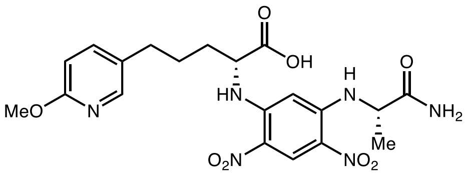

D-**3p** derivative

#### Marfey's analysis of L-**3o**

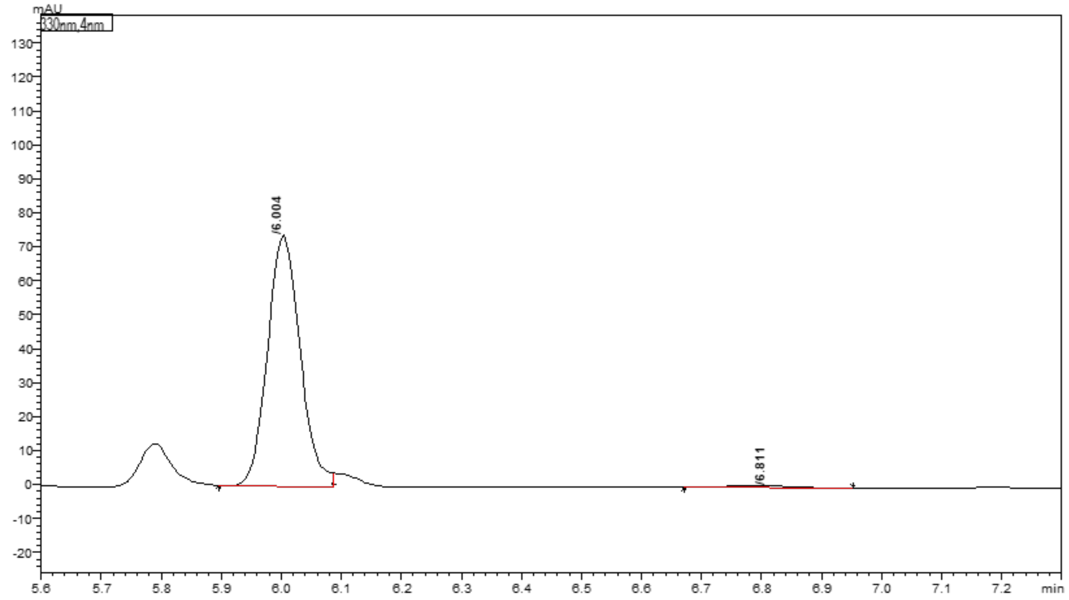

Peak Table | Compound | Group | Calibration Curve

| Peak# | Ret. Time | Height | Area | Area% |
|---|---|---|---|---|
| 1 | 6.004 | 73351 | 278818 | 98.864 |
| 2 | 6.811 | 574 | 3204 | 1.136 |
| Total | | 73926 | 282022 | 100.000 |

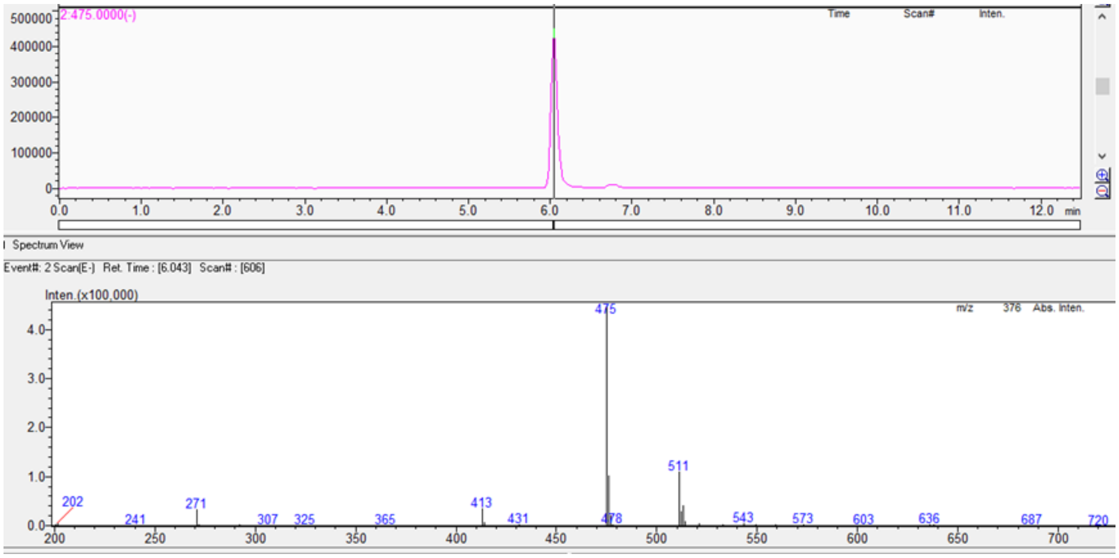

#### 2-Amino-6-phenylhexanoic acid (3p)

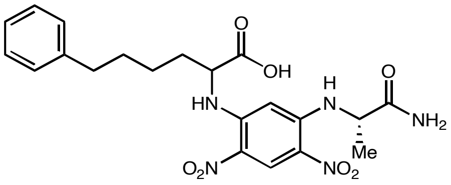

(*rac*)-**3o** derivative

Marfey's analysis of (*rac*)-**3p**

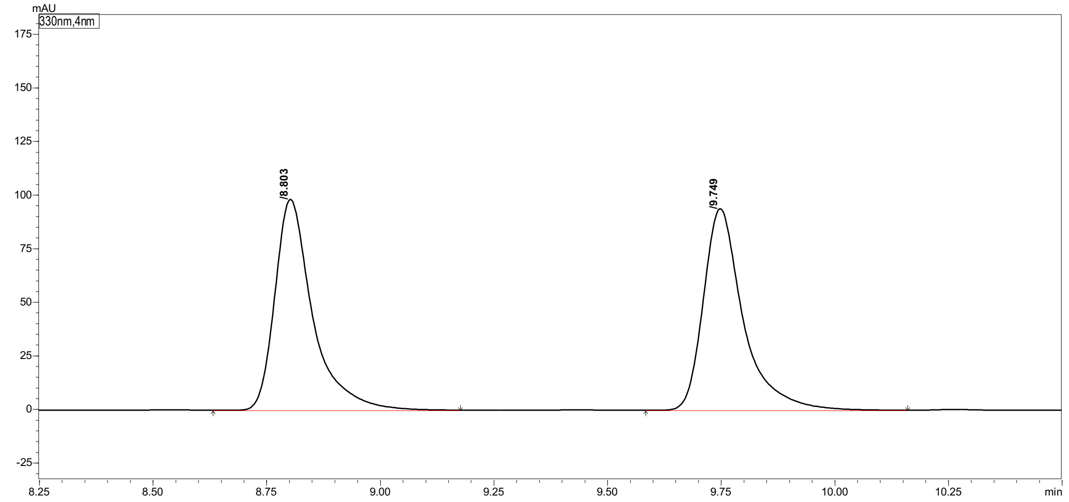

Peak Table | Compound | Group | Calibration Curve

| Peak# | Ret. Time | Height | Area | Area% |
|---|---|---|---|---|
| 1 | 8.803 | 97922 | 569218 | 49.812 |
| 2 | 9.749 | 93879 | 573507 | 50.188 |
| Total | | 191801 | 1142725 | 100.000 |

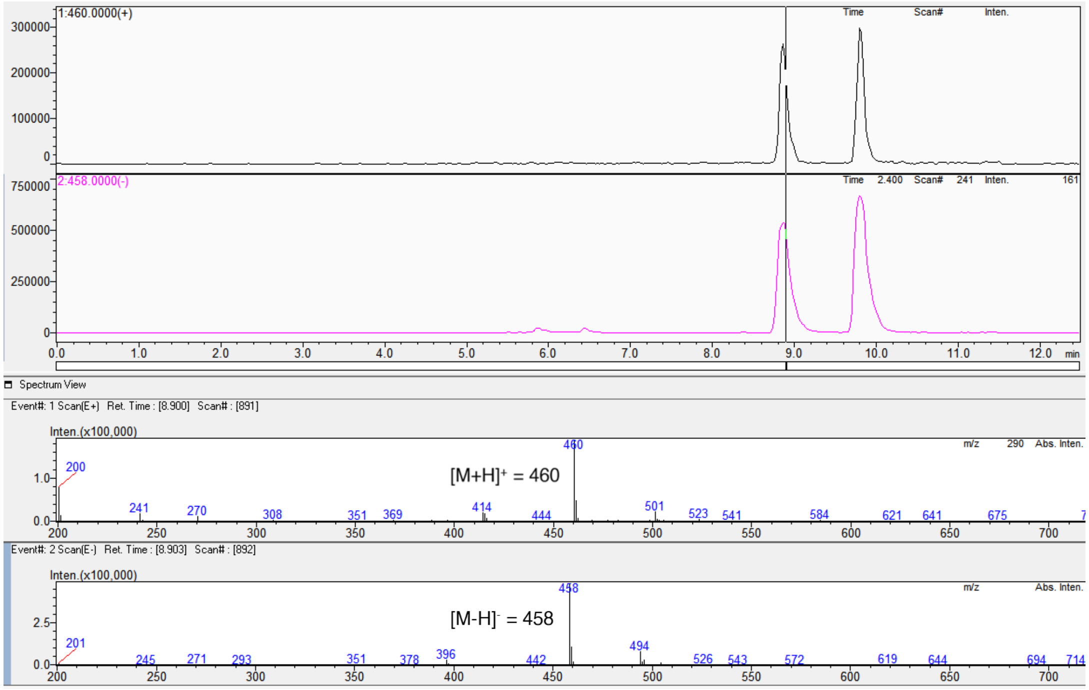

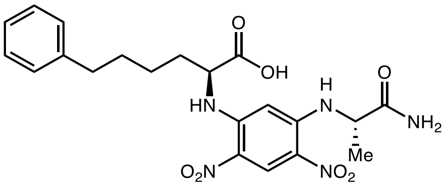

L-**3o** derivative

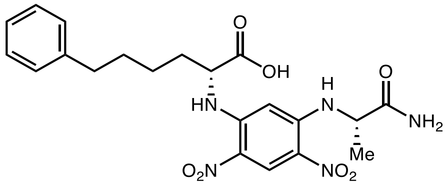

D-**3o** derivative

#### Marfey's analysis of L-**3p**

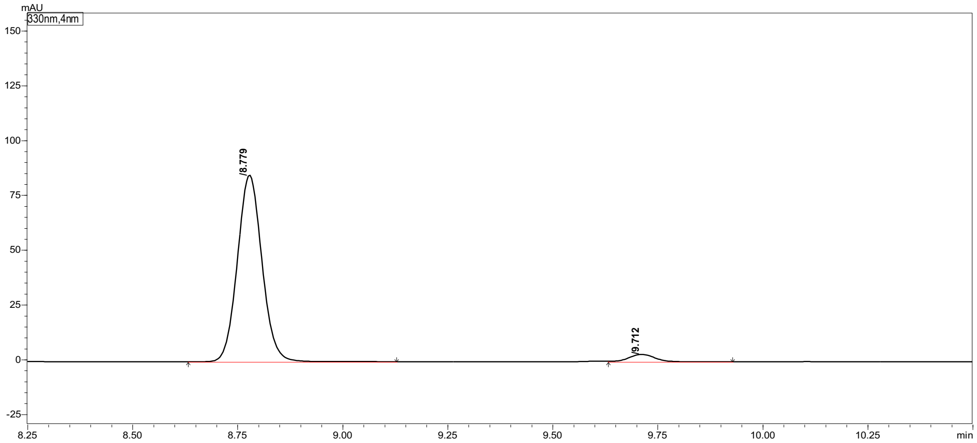

Peak Table | Compound | Group | Calibration Curve

| Peak# | Ret. Time | Height | Area | Area% |
|---|---|---|---|---|
| 1 | 8.779 | 84340 | 346601 | 96.004 |
| 2 | 9.712 | 3378 | 14428 | 3.996 |
| Total | | 87717 | 361028 | 100.000 |

#### Marfey's analysis of D-**3p**

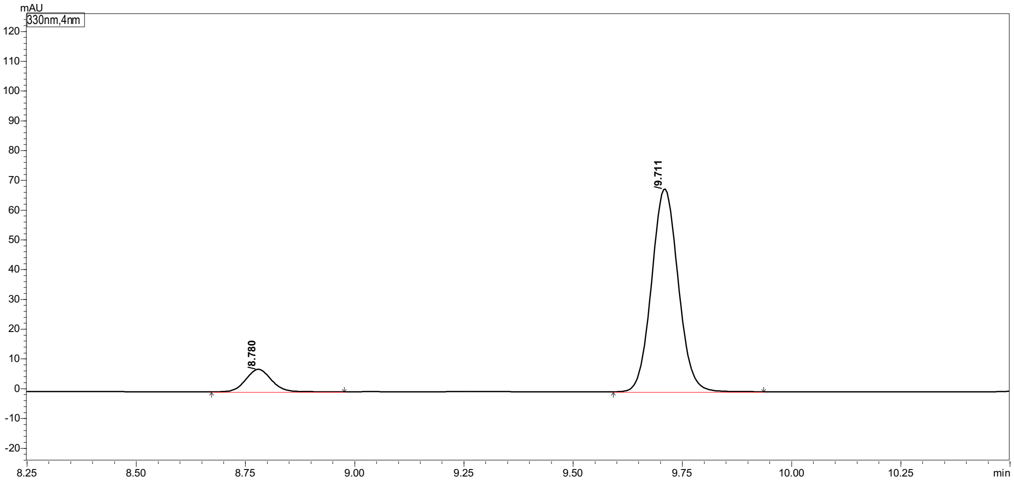

Peak Table | Compound | Group | Calibration Curve

| Peak# | Ret. Time | Height | Area | Area% |
|---|---|---|---|---|
| 1 | 8.780 | 7506 | 30612 | 9.613 |
| 2 | 9.711 | 68008 | 287838 | 90.387 |
| Total | | 75514 | 318449 | 100.000 |

#### 2-Amino-3-(2,3-dihydro-1H-inden-2-yl)propanoic acid (3q)

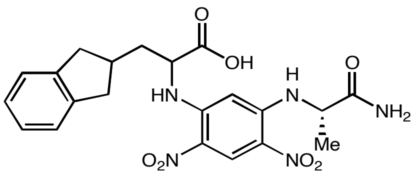

(*rac*)-**3q** derivative

#### Marfey's analysis of (*rac*)-**3q**

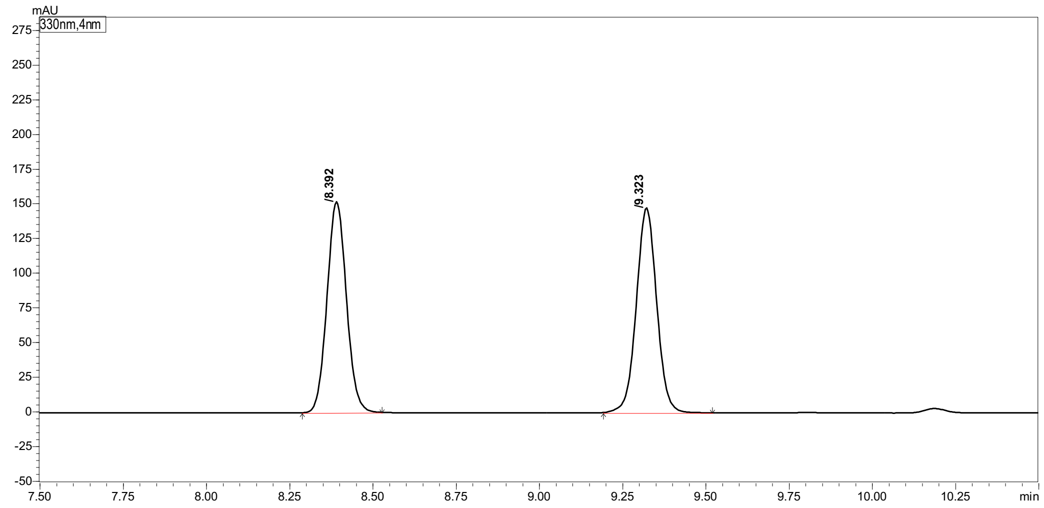

| Peak# | Ret. Time | Height | Area | Area% |
|---|---|---|---|---|
| 1 | 8.392 | 150884 | 615786 | 49.507 |
| 2 | 9.323 | 146462 | 628039 | 50.493 |
| Total | | 297346 | 1243825 | 100.000 |

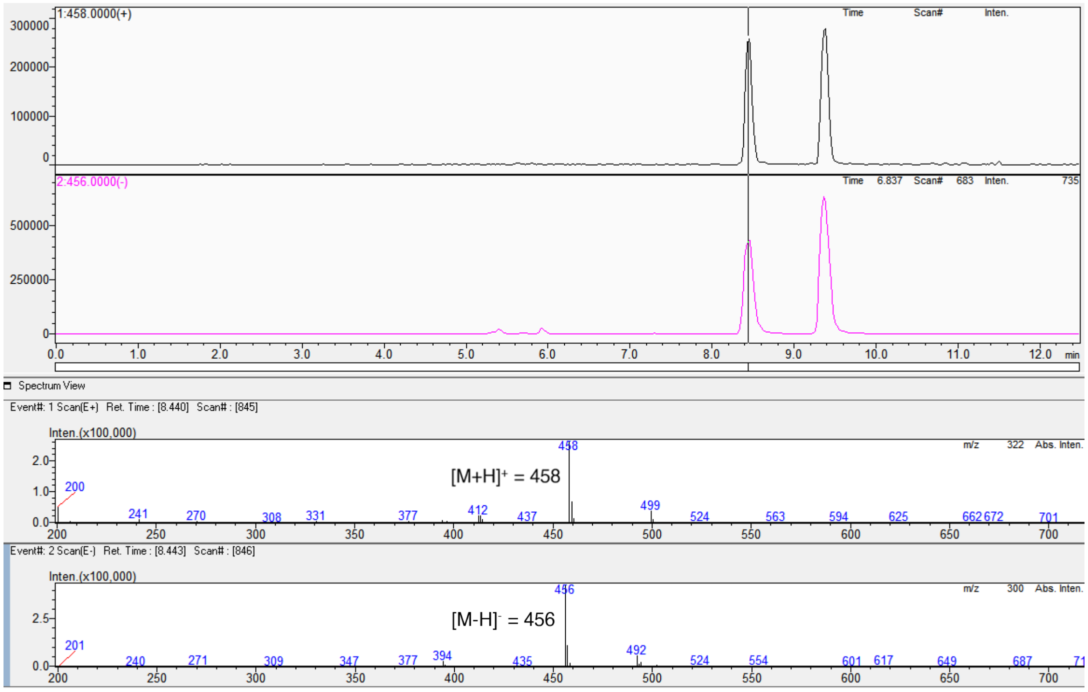

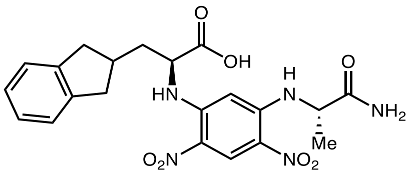

L-**3q** derivative

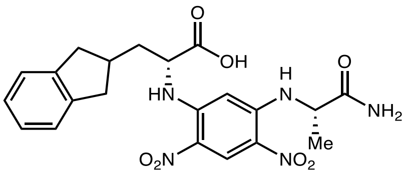

D-**3q** derivative

Marfey's analysis of D-**3q**

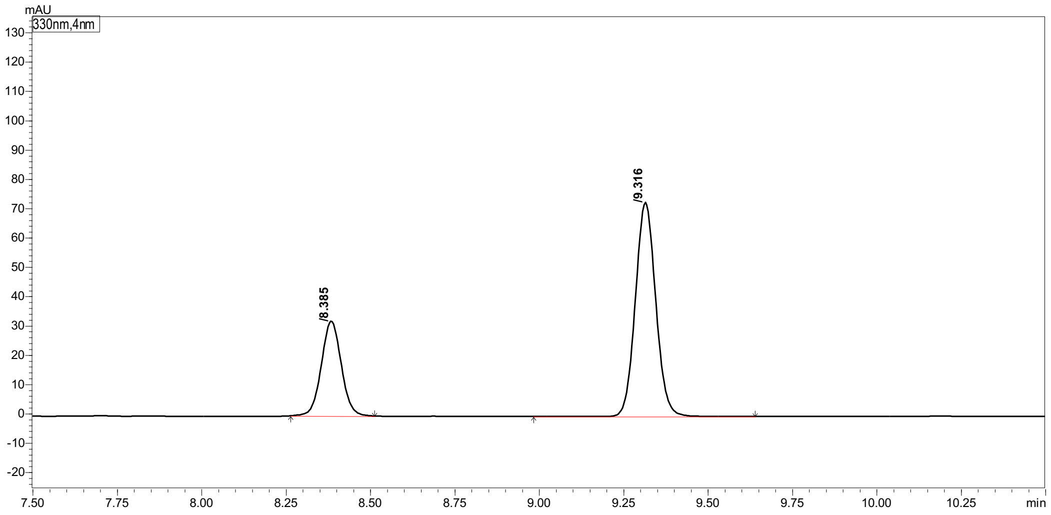

| Peak# | Ret. Time | Height | Area | Area% |
|---|---|---|---|---|
| 1 | 8.385 | 31967 | 132421 | 30.154 |
| 2 | 9.316 | 72649 | 306733 | 69.846 |
| Total | | 104617 | 439154 | 100.000 |

#### 2-Aminooctanoic acid (5a)

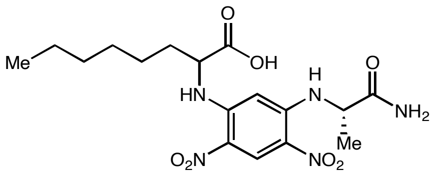

(*rac*)-**5a** derivative

#### Marfey's analysis of (*rac*)-**5a**

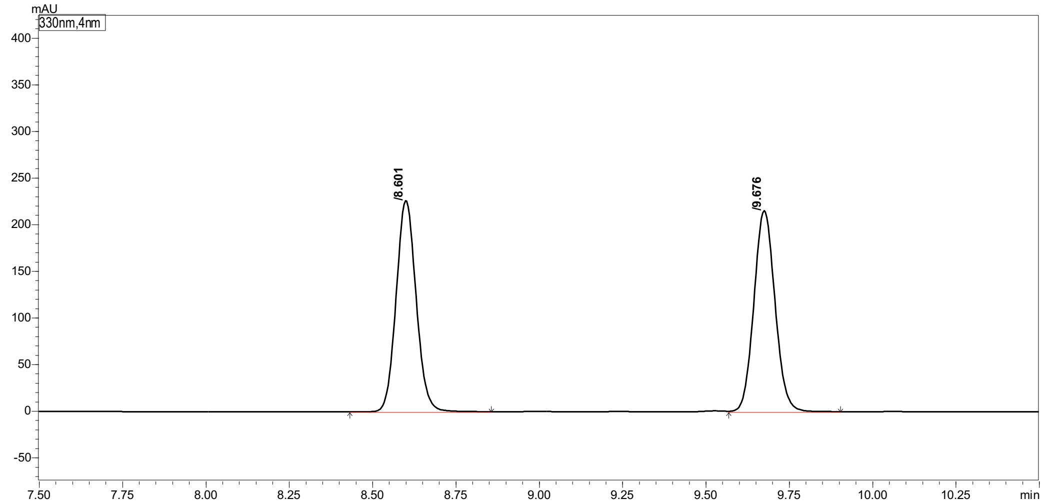

Peak Table | Compound | Group | Calibration Curve

| Peak# | Ret. Time | Height | Area | Area% |
|---|---|---|---|---|
| 1 | 8.601 | 224282 | 942566 | 49.869 |
| 2 | 9.676 | 214844 | 947501 | 50.131 |
| Total | | 439126 | 1890067 | 100.000 |

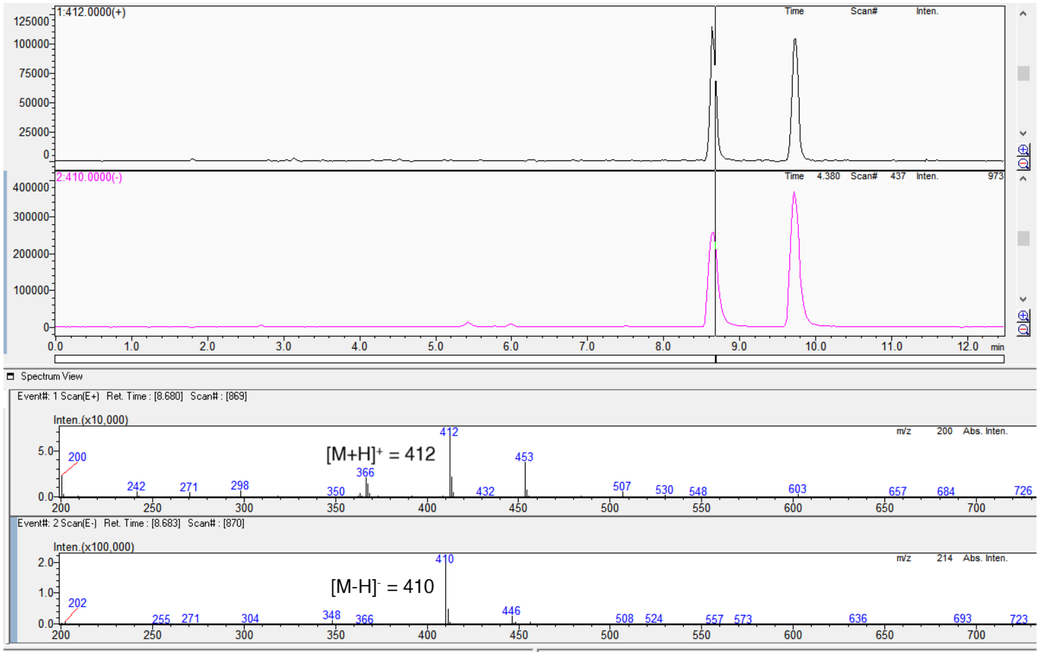

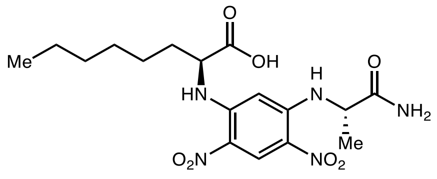

L-**5a** derivative

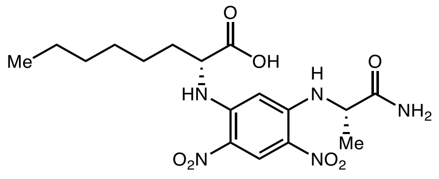

D-**5a** derivative

Marfey's analysis of L-**5a**

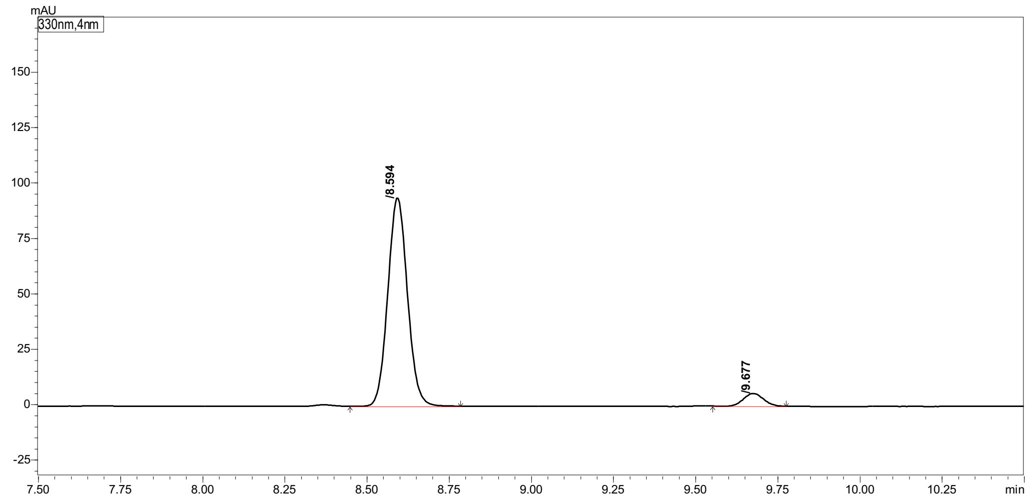

Peak Table | Compound | Group | Calibration Curve

| Peak# | Ret. Time | Height | Area | Area% |
|---|---|---|---|---|
| 1 | 8.594 | 92683 | 390269 | 94.334 |
| 2 | 9.677 | 5673 | 23443 | 5.666 |
| Total | | 98355 | 413712 | 100.000 |

Marfey's analysis of D-**5a**

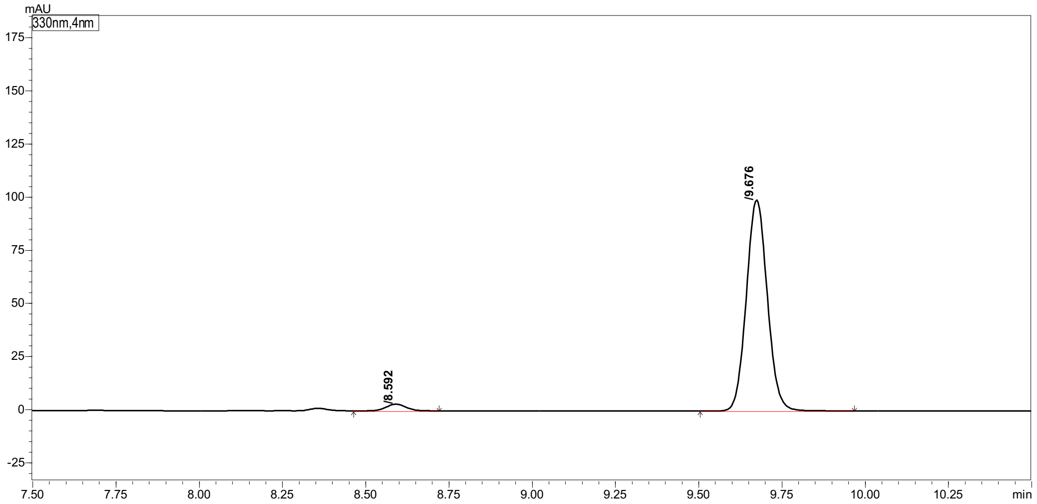

Peak Table | Compound | Group | Calibration Curve

| Peak# | Ret. Time | Height | Area | Area% |
|---|---|---|---|---|
| 1 | 8.592 | 3107 | 12757 | 2.858 |
| 2 | 9.676 | 98701 | 433678 | 97.142 |
| Total | | 101807 | 446435 | 100.000 |

#### 2-Amino-7-fluoroheptanoic acid (5b)

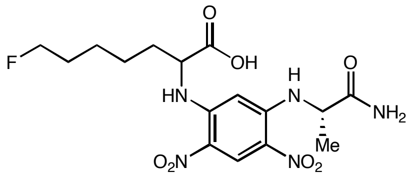

(*rac*)-**5h** derivative

Marfey's analysis of (*rac*)-**5b**

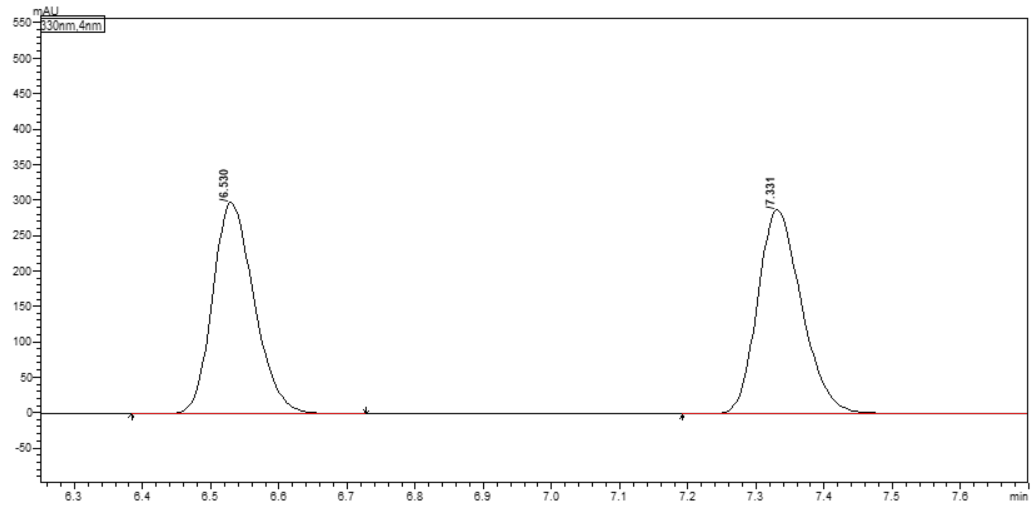

Peak Table | Compound | Group | Calibration Curve

| Peak# | Ret. Time | Height | Area | Area% |
|---|---|---|---|---|
| 1 | 6.530 | 295020 | 1265637 | 49.358 |
| 2 | 7.331 | 285796 | 1298555 | 50.642 |
| Total | | 580816 | 2564192 | 100.000 |

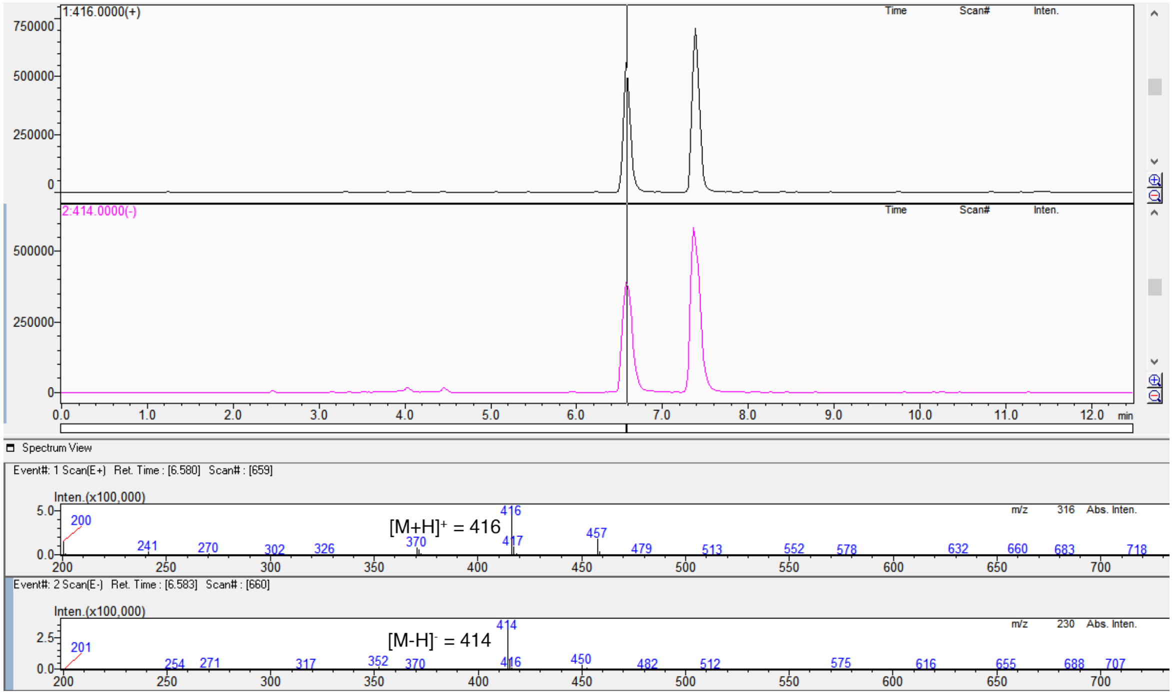

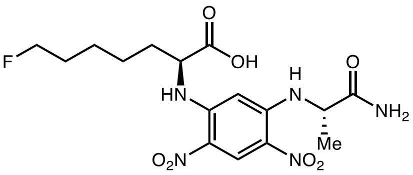

L-**5b** derivative

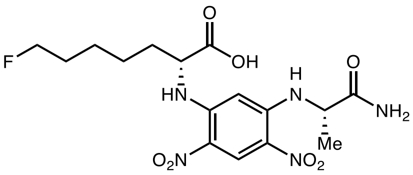

D-**5b** derivative

Marfey's analysis of L-**5b**

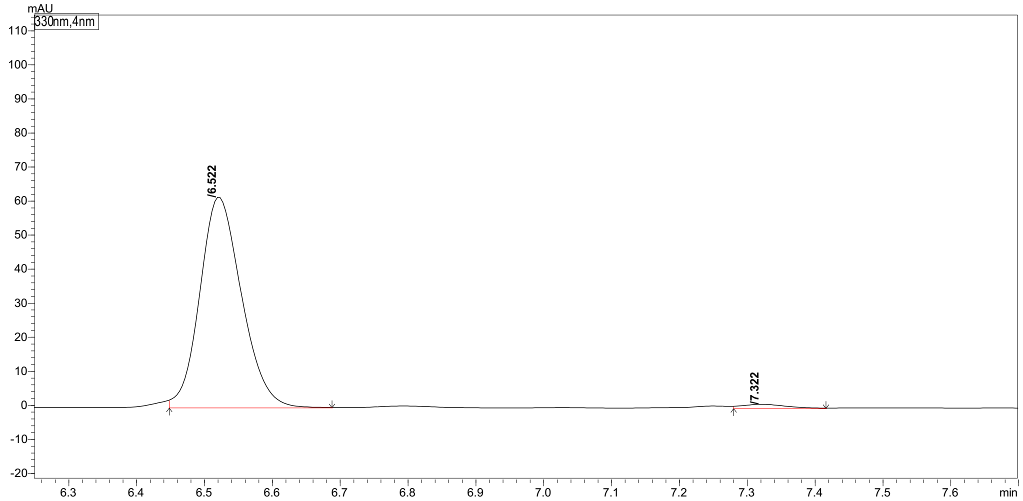

| Peak# | Ret. Time | Height | Area | Area% |
|---|---|---|---|---|
| 1 | 6.522 | 61060 | 263031 | 98.325 |
| 2 | 7.322 | 1070 | 4480 | 1.675 |
| Total | | 62130 | 267510 | 100.000 |

Marfey's analysis of D-**5b**

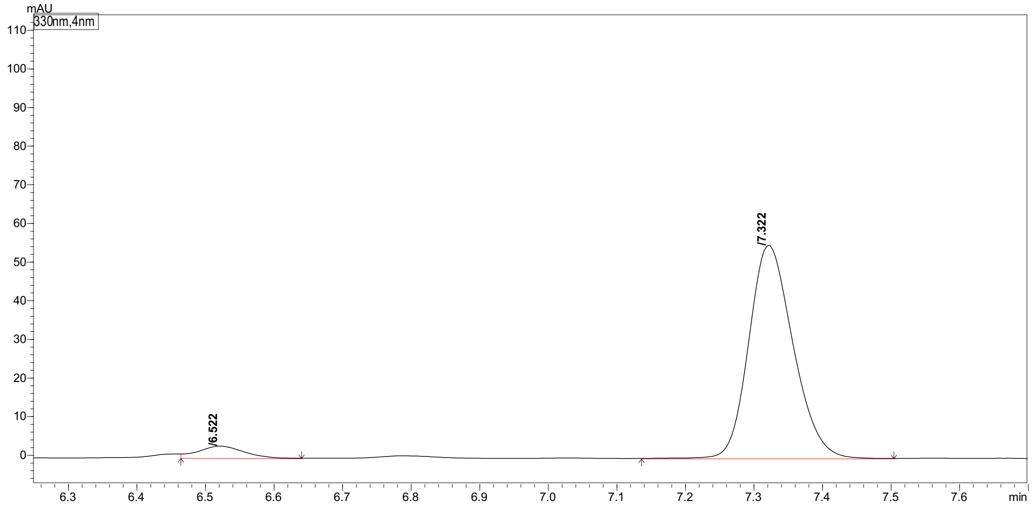

| Peak# | Ret. Time | Height | Area | Area% |
|---|---|---|---|---|
| 1 | 6.522 | 3050 | 13834 | 5.310 |
| 2 | 7.322 | 54500 | 246681 | 94.690 |
| Total | | 57549 | 260516 | 100.000 |

#### 2-Amino-8-chlorooctanoic acid (5c)

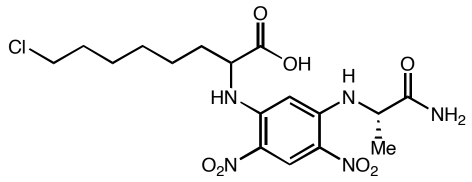

(*rac*)-**5c** derivative

#### Marfey's analysis of (*rac*)-**5c**

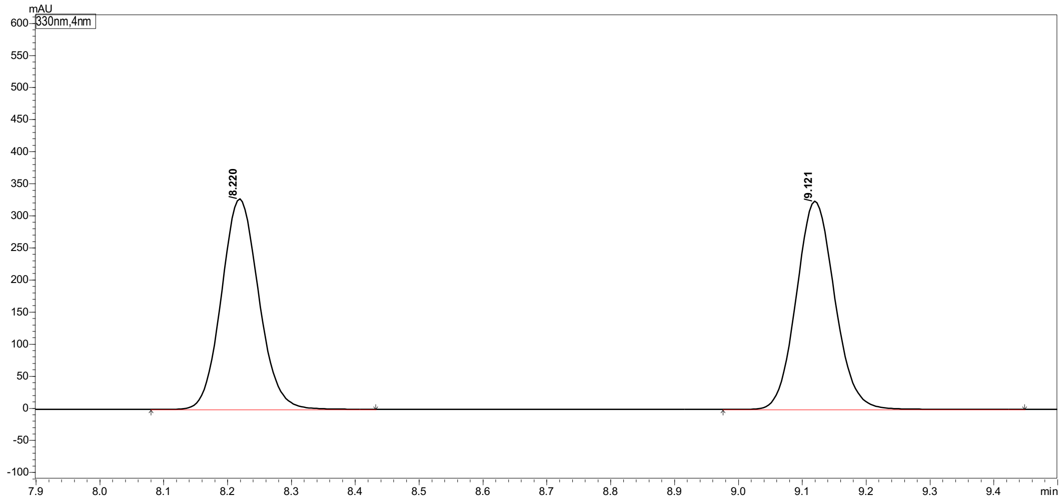

| Peak# | Ret. Time | Height | Area | Area% |
|---|---|---|---|---|
| 1 | 8.220 | 325872 | 1330887 | 49.543 |
| 2 | 9.121 | 322530 | 1355451 | 50.457 |
| Total | | 648402 | 2686338 | 100.000 |

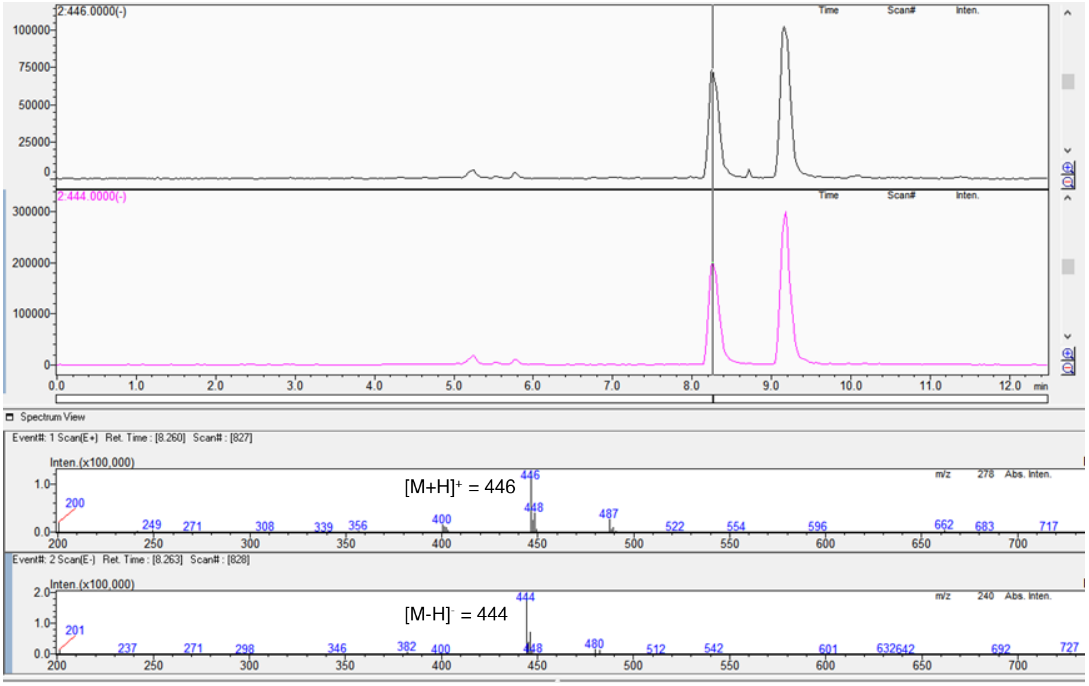

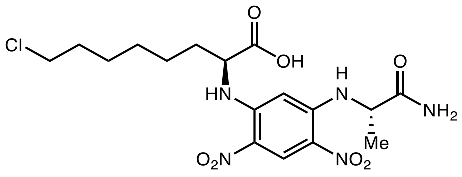

L-**5c** derivative

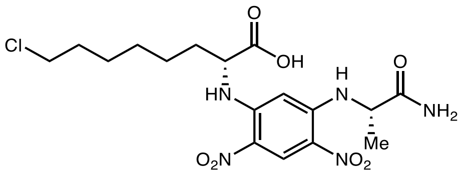

D-**5c** derivative

Marfey's analysis of L-**5c**

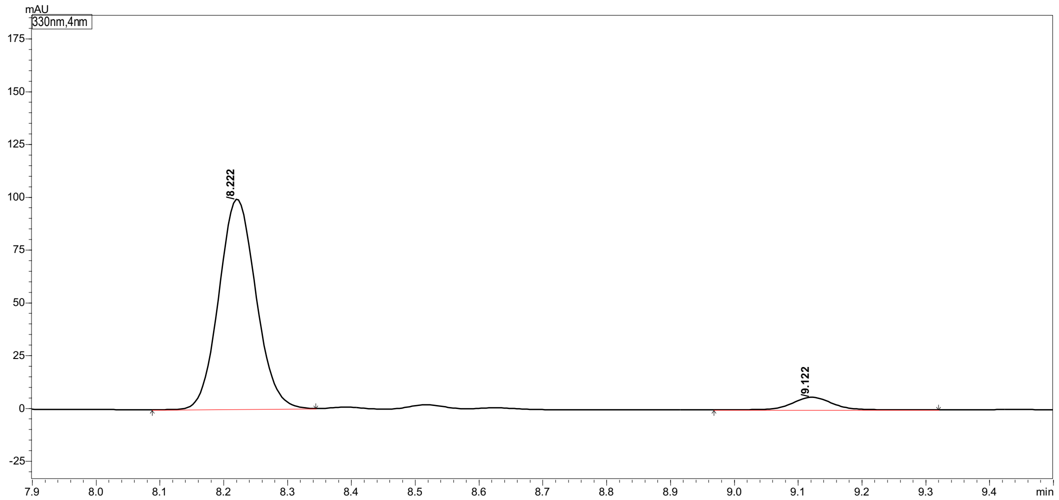

Peak Table | Compound | Group | Calibration Curve

| Peak# | Ret. Time | Height | Area | Area% |
|---|---|---|---|---|
| 1 | 8.222 | 99257 | 400788 | 94.104 |
| 2 | 9.122 | 5830 | 25112 | 5.896 |
| Total | | 105087 | 425900 | 100.000 |

Marfey's analysis of D-**5c**

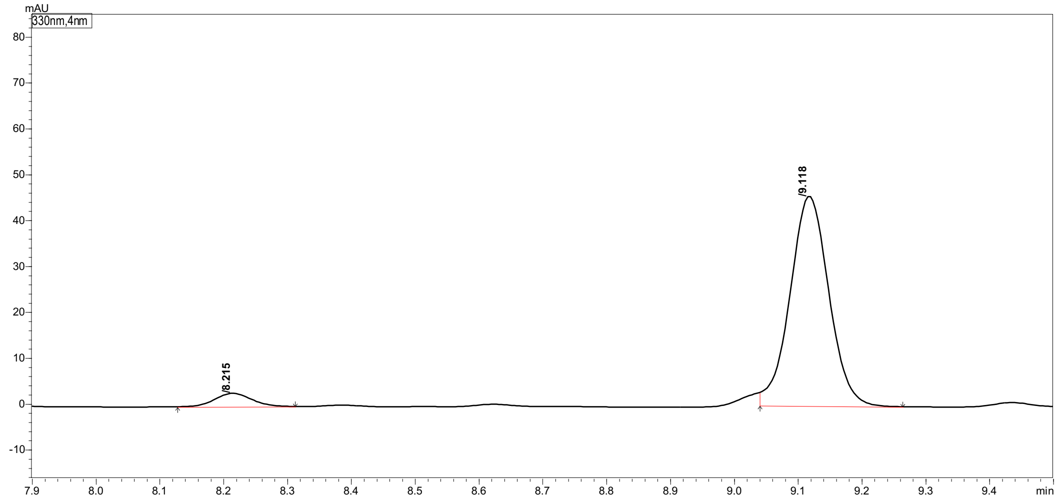

Peak Table | Compound | Group | Calibration Curve

| Peak# | Ret. Time | Height | Area | Area% |
|---|---|---|---|---|
| 1 | 8.215 | 2885 | 11566 | 5.652 |
| 2 | 9.118 | 45658 | 193071 | 94.348 |
| Total | | 48543 | 204637 | 100.000 |

**2-Amino-7-methoxyheptanoic acid (5d)**

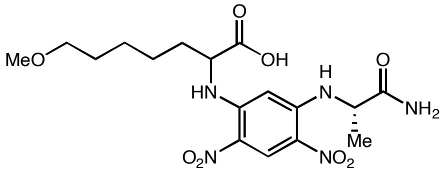

(*rac*)-**5d** derivative

Marfey's analysis of (*rac*)-**5d** (Long HPLC **method b** was used for separation)

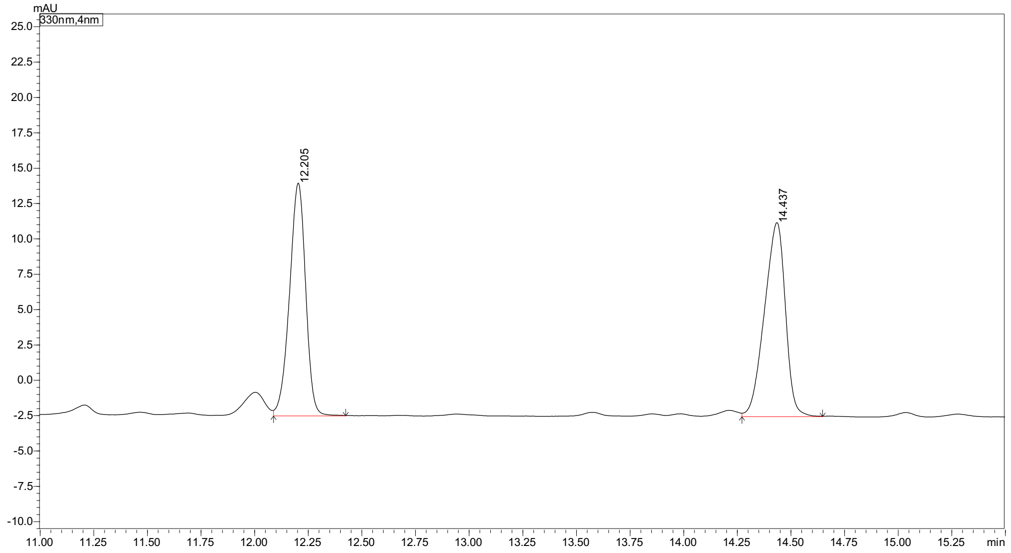

Peak Table | Compound | Group | Calibration Curve

| Peak# | Peak Start | Peak End | Ret. Time | Area | Area% |
|---|---|---|---|---|---|
| 1 | 12.088 | 12.424 | 12.205 | 85674 | 48.051 |
| 2 | 14.272 | 14.648 | 14.437 | 92623 | 51.949 |
| Total | | | | 178297 | 100.000 |

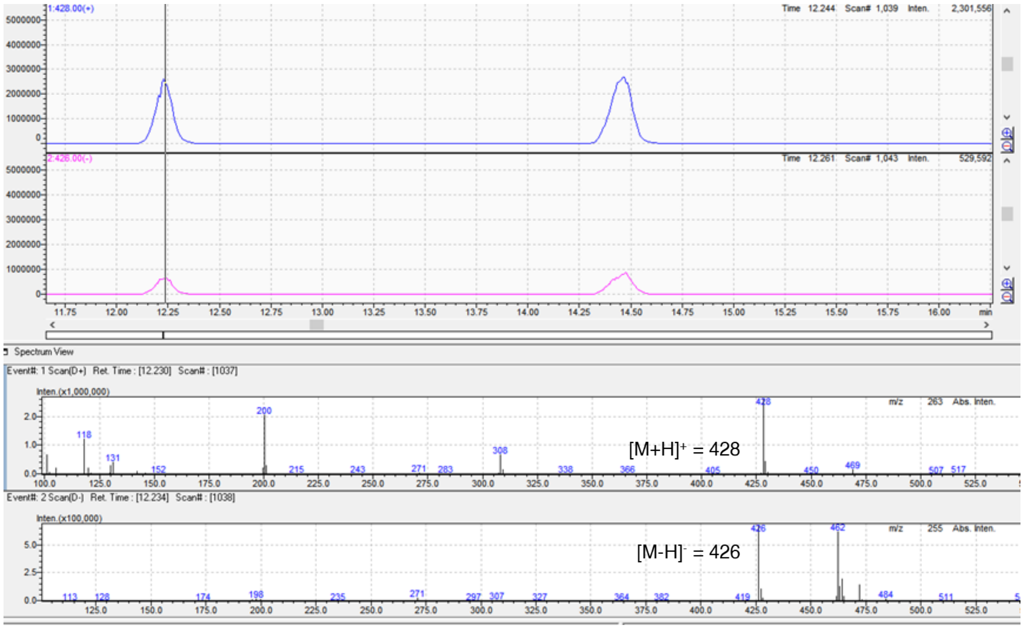

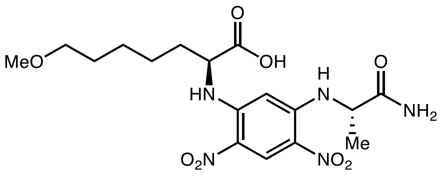

L-**5d** derivative

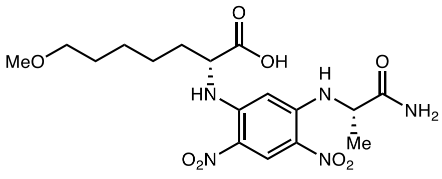

D-**5d** derivative

Marfey's analysis of L-**5d**

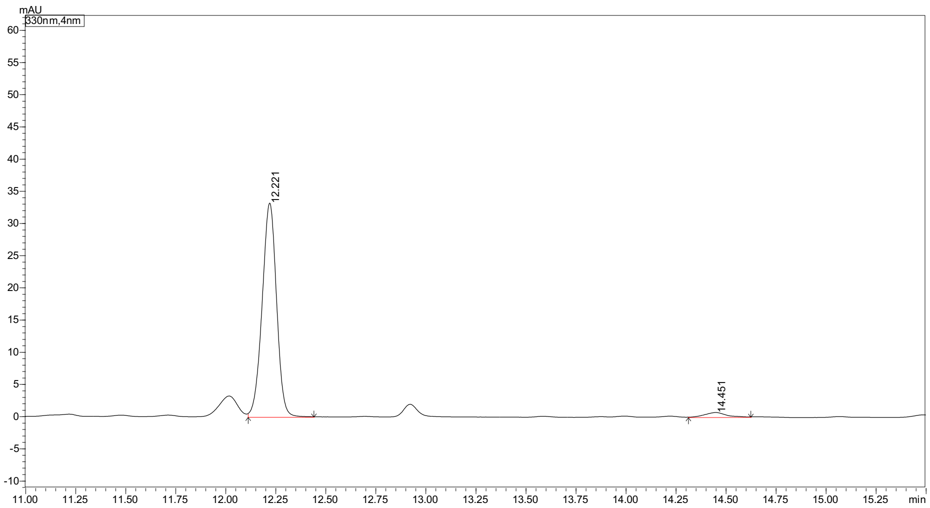

Peak Table | Compound | Group | Calibration Curve

| Peak# | Peak Start | Peak End | Ret. Time | Area | Area% |
|---|---|---|---|---|---|
| 1 | 12.112 | 12.440 | 12.221 | 165302 | 97.135 |
| 2 | 14.312 | 14.624 | 14.451 | 4875 | 2.865 |
| Total | | | | 170177 | 100.000 |

Marfey's analysis of D-**5d**

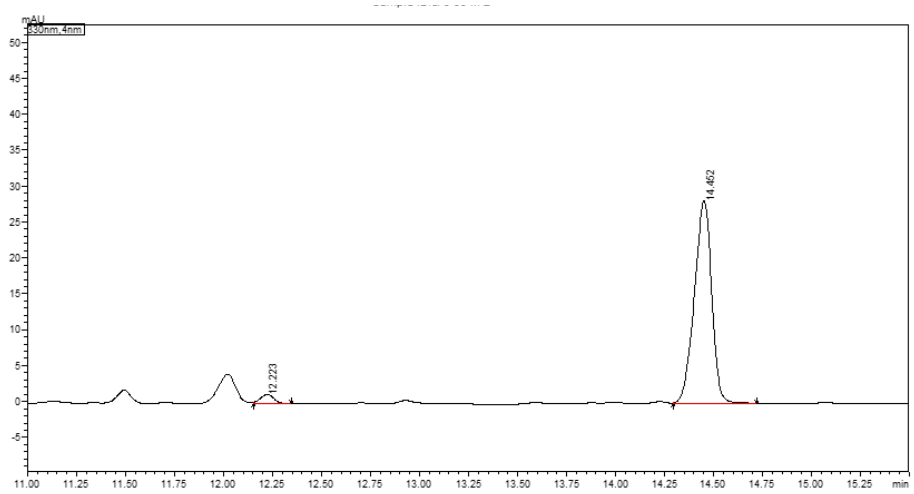

Peak Table | Compound | Group | Calibration Curve

| Peak# | Peak Start | Peak End | Ret. Time | Area | Area% |
|---|---|---|---|---|---|
| 1 | 12.152 | 12.344 | 12.223 | 6117 | 3.411 |
| 2 | 14.296 | 14.720 | 14.452 | 173226 | 96.589 |
| Total | | | | 179343 | 100.000 |

#### 2-Amino-7-(methylthio)heptanoic acid (5e)

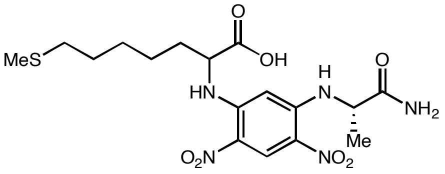

(*rac*)-**5e** derivative

Marfey's analysis of (*rac*)-**5e** (Long HPLC **method b** was used for separation)

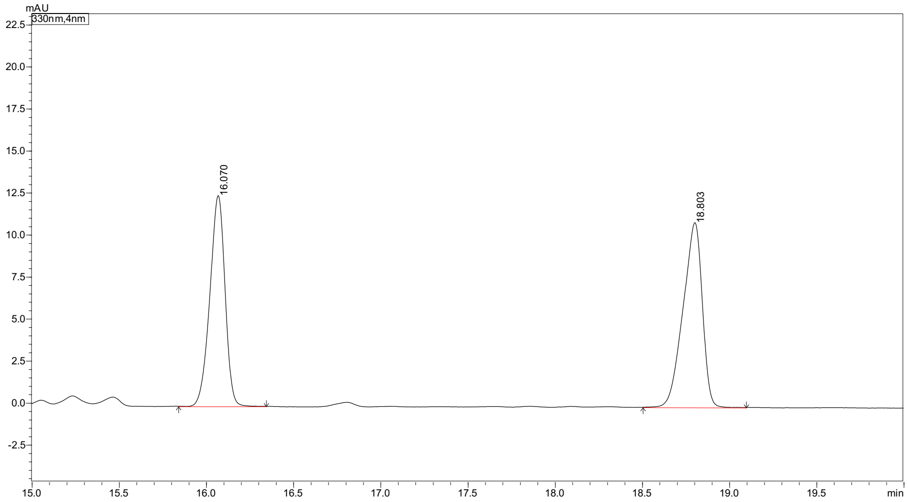

| Peak# | Peak Start | Peak End | Ret. Time | Area | Area% |
|---|---|---|---|---|---|
| 1 | 15.840 | 16.344 | 16.070 | 75718 | 47.322 |
| 2 | 18.504 | 19.096 | 18.803 | 84289 | 52.678 |
| Total | | | | 160007 | 100.000 |

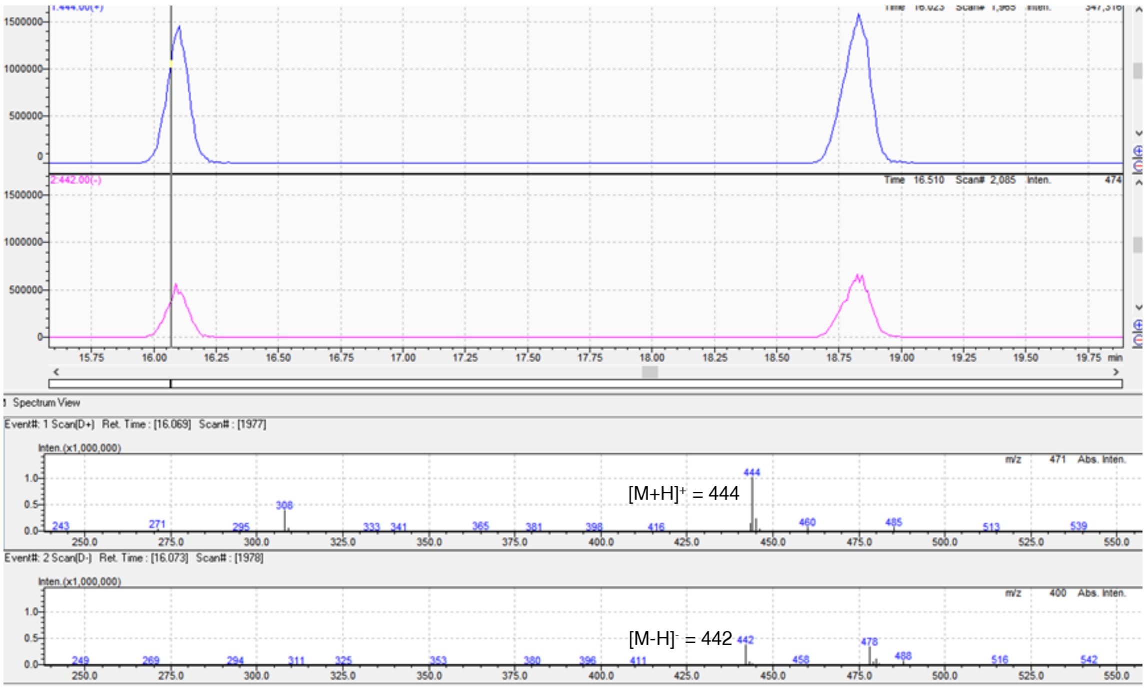

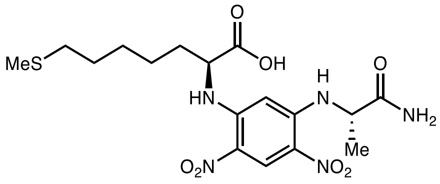

L-**5e** derivative

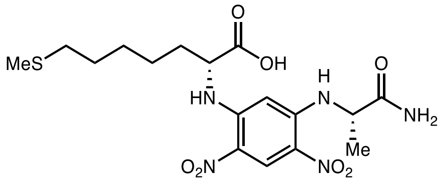

D-**5e** derivative

Marfey's analysis of L-**5e**

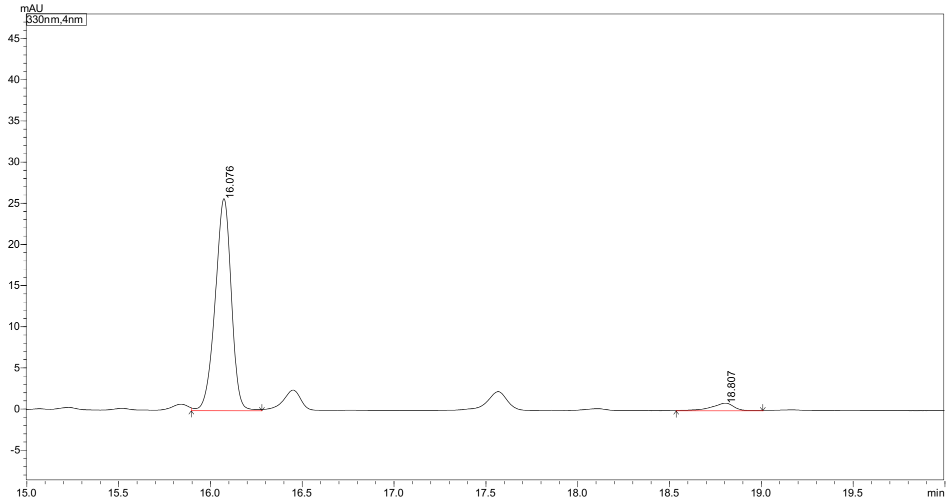

| Peak# | Peak Start | Peak End | Ret. Time | Area | Area% |
|---|---|---|---|---|---|
| 1 | 15.896 | 16.280 | 16.076 | 156425 | 95.864 |
| 2 | 18.536 | 19.008 | 18.807 | 6749 | 4.136 |
| Total | | | | 163174 | 100.000 |

Marfey's analysis of D-**5e**

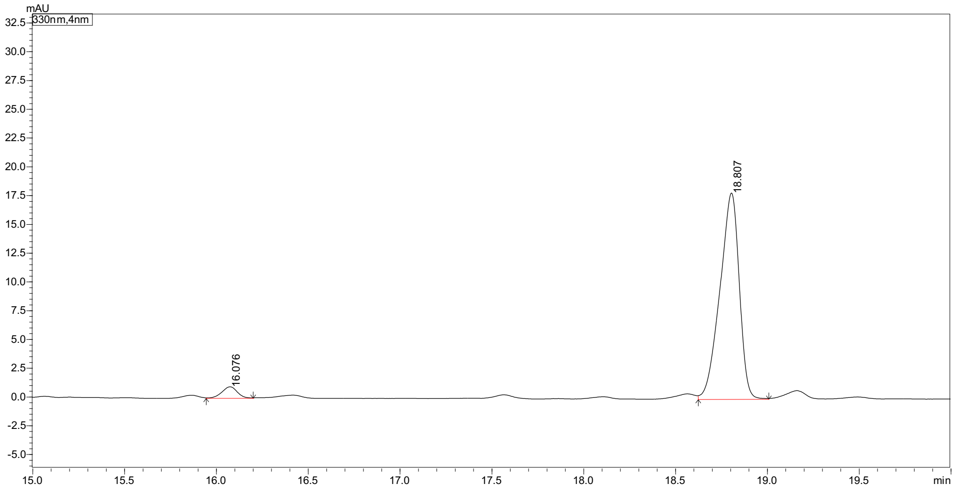

| Peak# | Peak Start | Peak End | Ret. Time | Area | Area% |
|---|---|---|---|---|---|
| 1 | 15.944 | 16.200 | 16.076 | 5287 | 3.877 |
| 2 | 18.624 | 19.008 | 18.807 | 131080 | 96.123 |
| Total | | | | 136366 | 100.000 |

#### 2-Amino-8-azidooctanoic acid (5f)

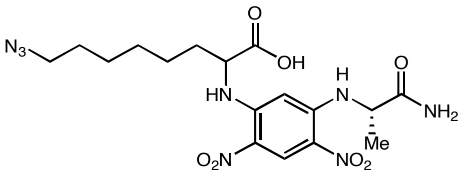

(*rac*)-**5l** derivative

##### Marfey's analysis of (*rac*)-**5f**

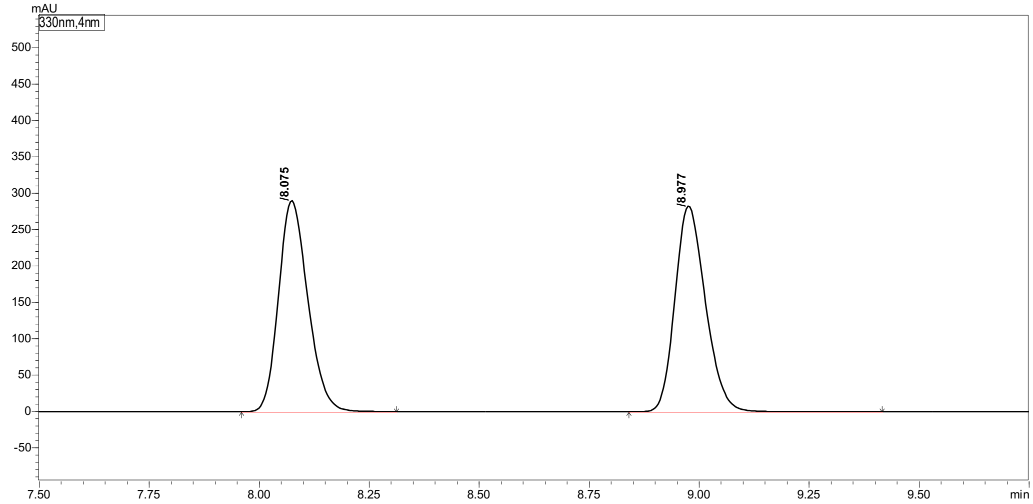

| Peak# | Ret. Time | Height | Area | Area% |
|---|---|---|---|---|
| 1 | 8.075 | 288633 | 1344908 | 49.757 |
| 2 | 8.977 | 280662 | 1358034 | 50.243 |
| Total | | 569294 | 2702942 | 100.000 |

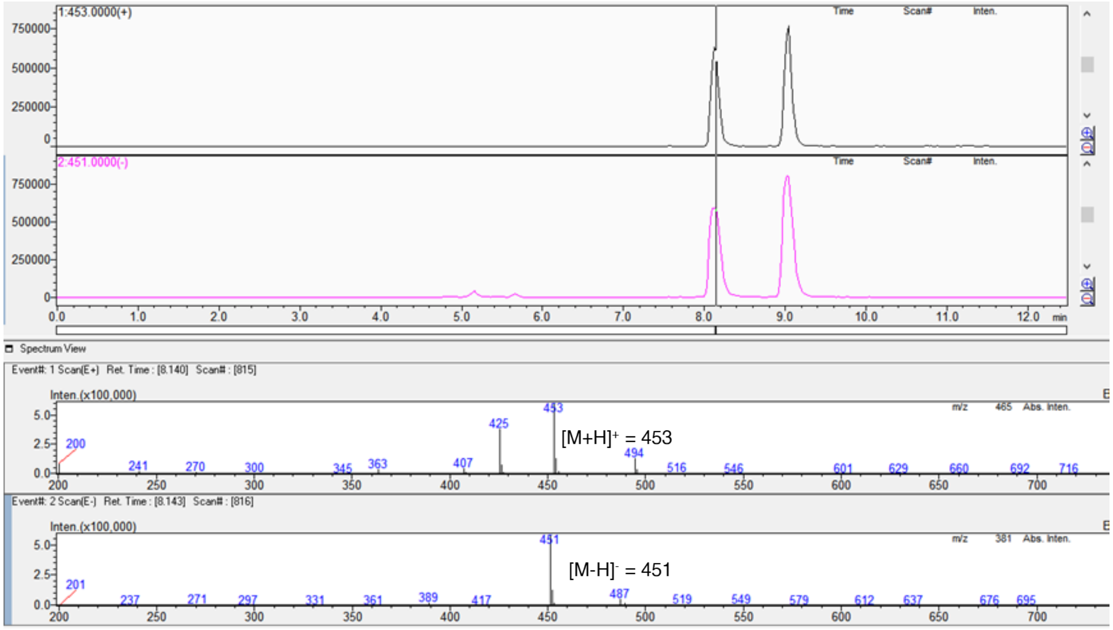

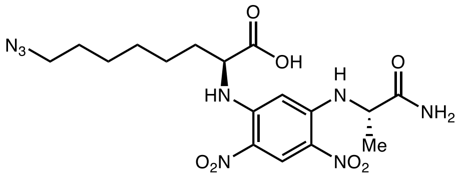

L-**5f** derivative

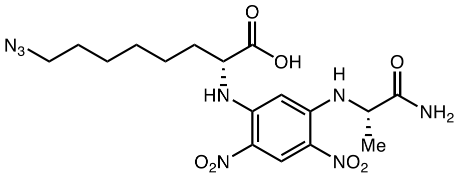

D-**5f** derivative

#### Marfey’s analysis of L-**5f**

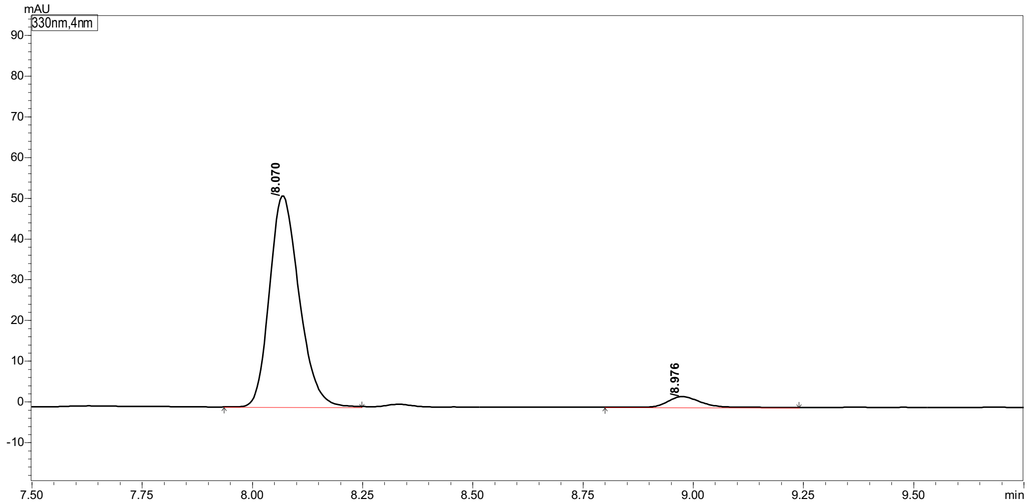

Peak Table | Compound | Group | Calibration Curve

| Peak# | Ret. Time | Height | Area | Area% |
|---|---|---|---|---|
| 1 | 8.070 | 51773 | 242446 | 94.960 |
| 2 | 8.976 | 2616 | 12869 | 5.040 |
| Total | | 54390 | 255315 | 100.000 |

#### Marfey’s analysis of D-**5f**

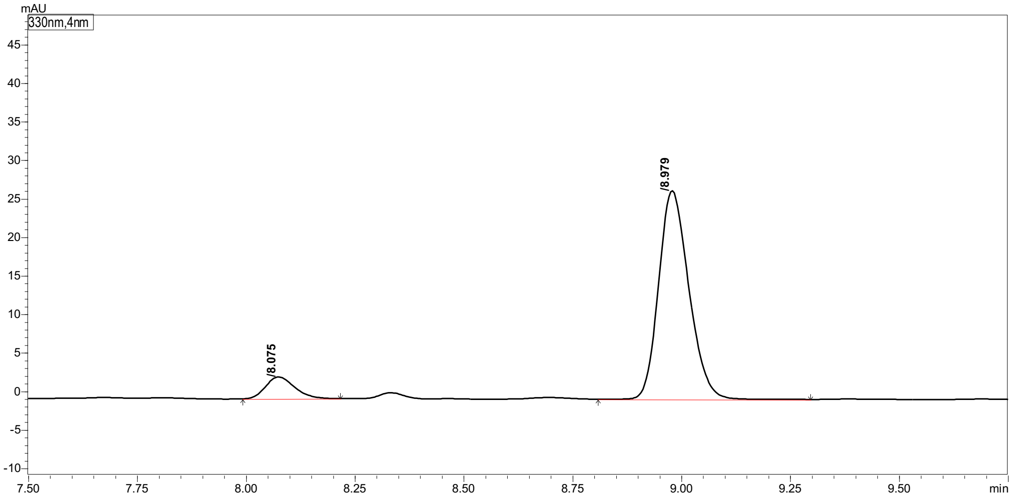

Peak Table | Compound | Group | Calibration Curve

| Peak# | Ret. Time | Height | Area | Area% |
|---|---|---|---|---|
| 1 | 8.075 | 2833 | 13219 | 9.212 |
| 2 | 8.979 | 26907 | 130283 | 90.788 |
| Total | | 29740 | 143502 | 100.000 |

#### 2-Aminooct-7-ynoic acid (5g)

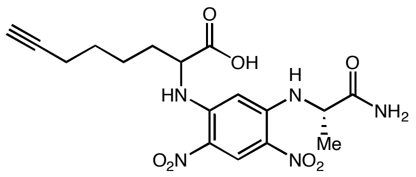

(*rac*)-**5g** derivative

#### Marfey's analysis of (*rac*)-**5g**

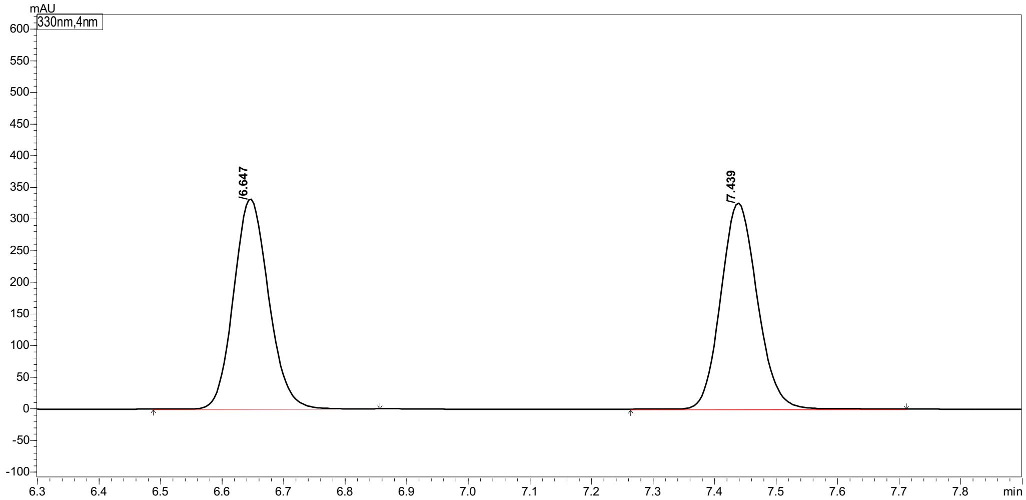

Peak Table | Compound | Group | Calibration Curve

| Peak# | Ret. Time | Height | Area | Area% |
|---|---|---|---|---|
| 1 | 6.647 | 330868 | 1307558 | 49.329 |
| 2 | 7.439 | 324501 | 1343125 | 50.671 |
| Total | | 655369 | 2650683 | 100.000 |

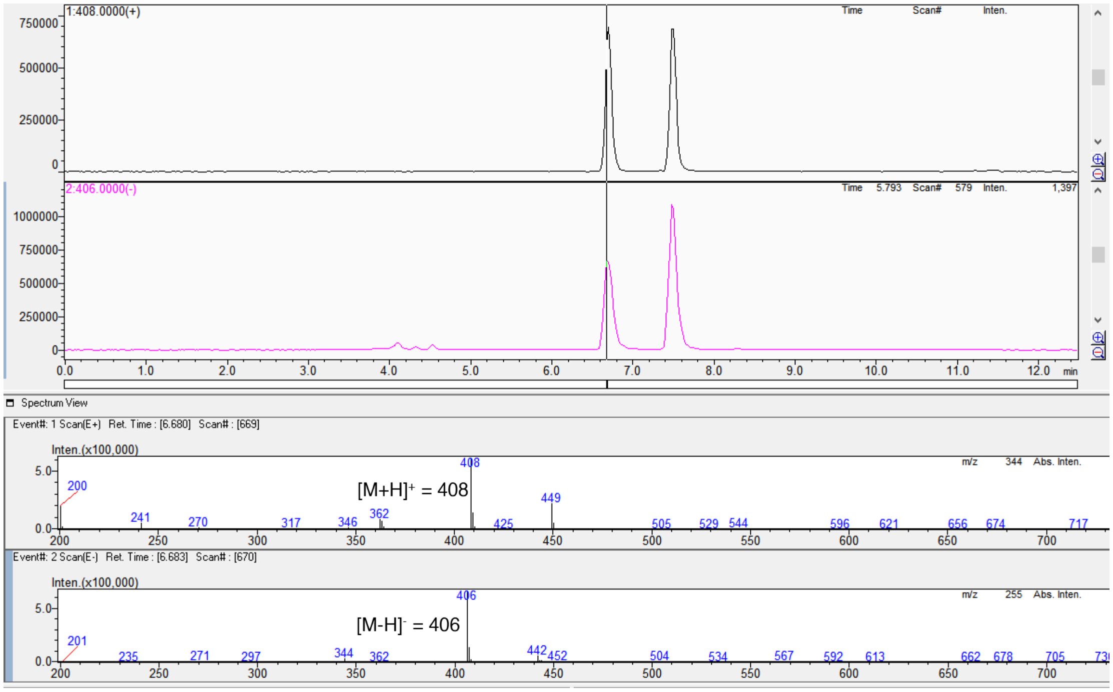

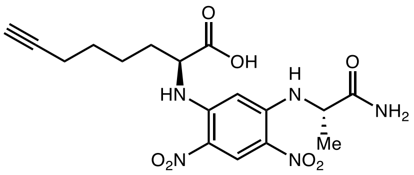

L-**5g** derivative

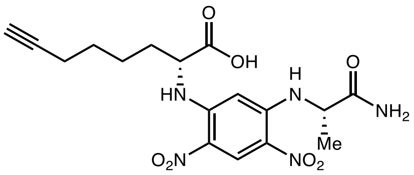

D-**5g** derivative

Marfey's analysis of L-**5g**

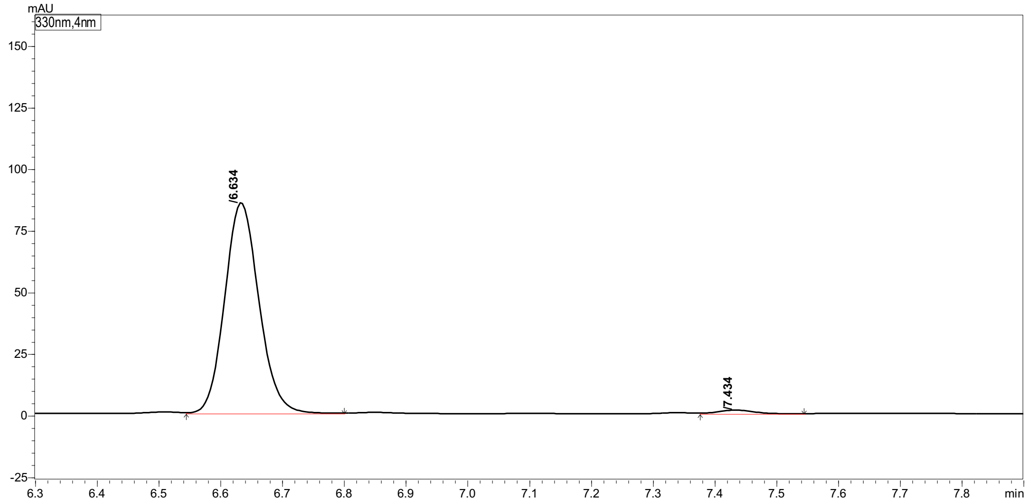

Peak Table | Compound | Group | Calibration Curve

| Peak# | Ret. Time | Height | Area | Area% |
|---|---|---|---|---|
| 1 | 6.634 | 84470 | 330141 | 98.309 |
| 2 | 7.434 | 1442 | 5680 | 1.691 |
| Total | | 85912 | 335821 | 100.000 |

Marfey's analysis of D-**5g**

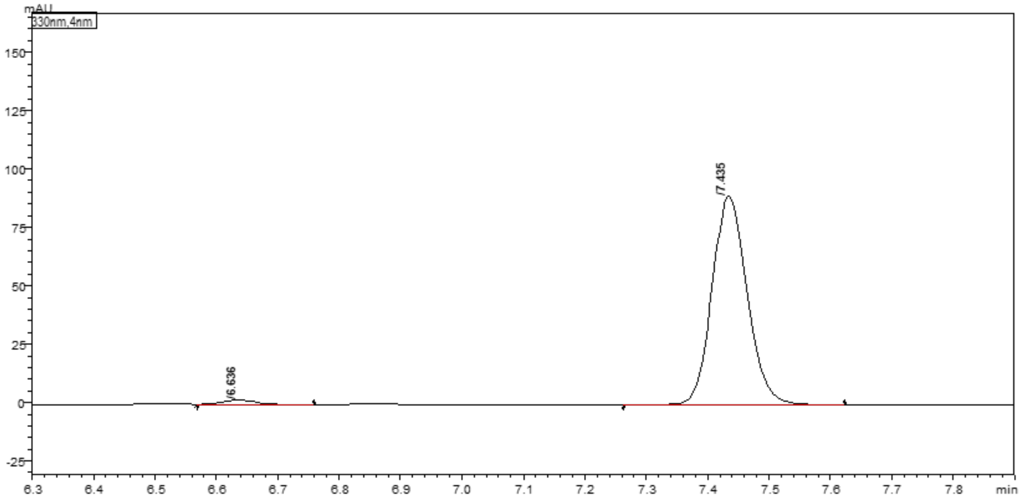

Peak Table | Compound | Group | Calibration Curve

| Peak# | Ret. Time | Height | Area | Area% |
|---|---|---|---|---|
| 1 | 6.636 | 1949 | 7371 | 2.026 |
| 2 | 7.435 | 88734 | 356460 | 97.974 |
| Total | | 90683 | 363831 | 100.000 |

#### 2-Aminohept-6-enoic acid (5h)

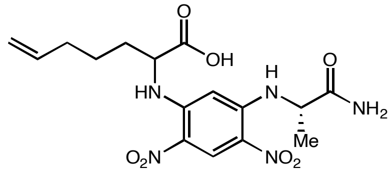

(*rac*)-**5h** derivative

Marfey's analysis of (*rac*)-**5h** (Long HPLC **method b** was used for separation)

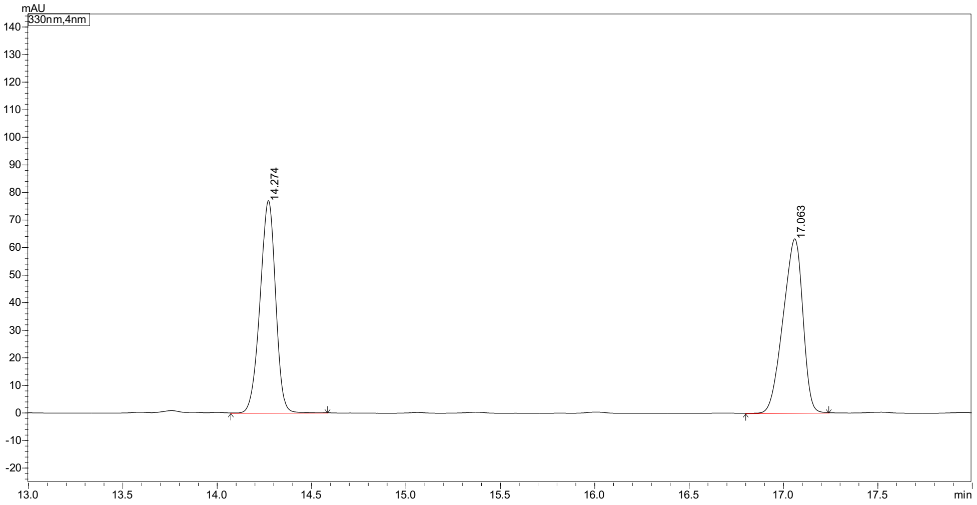

| Peak# | Peak Start | Peak End | Ret. Time | Area | Area% |
|---|---|---|---|---|---|
| 1 | 14.072 | 14.584 | 14.274 | 439847 | 49.006 |
| 2 | 16.800 | 17.240 | 17.063 | 457691 | 50.994 |
| Total | | | | 897538 | 100.000 |

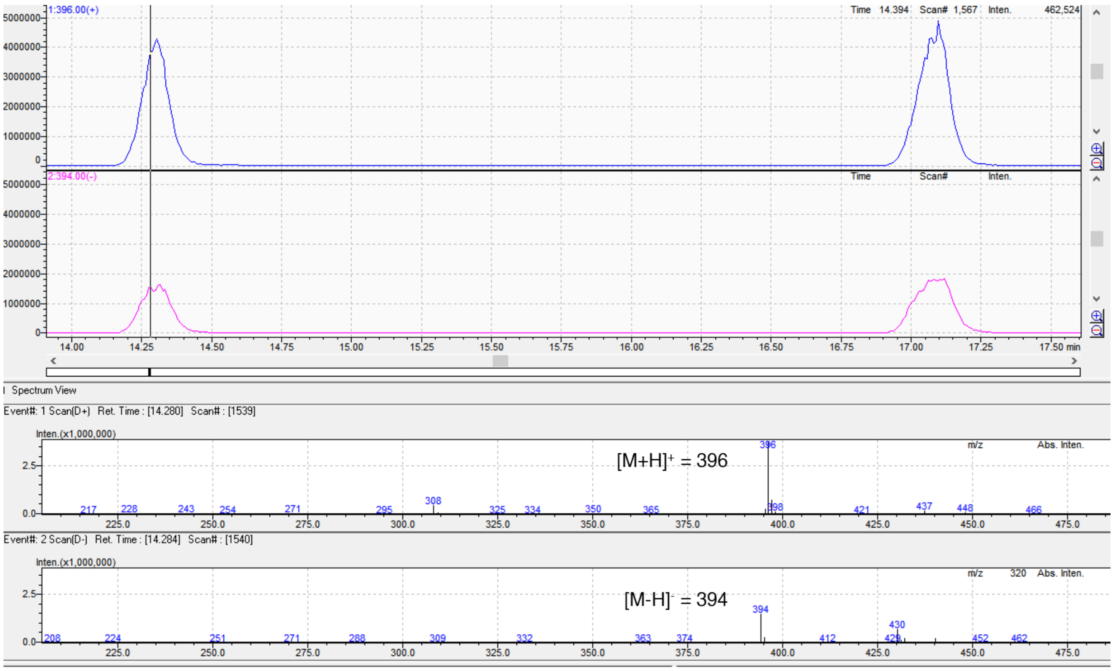

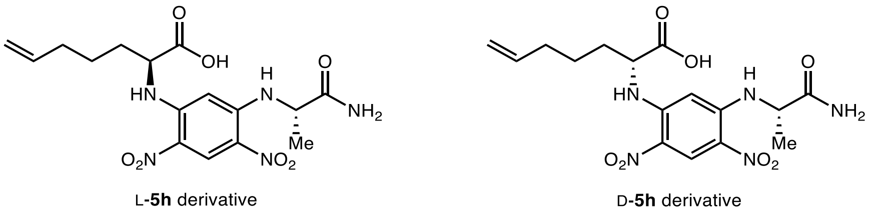

Marfey's analysis of L-**5h**

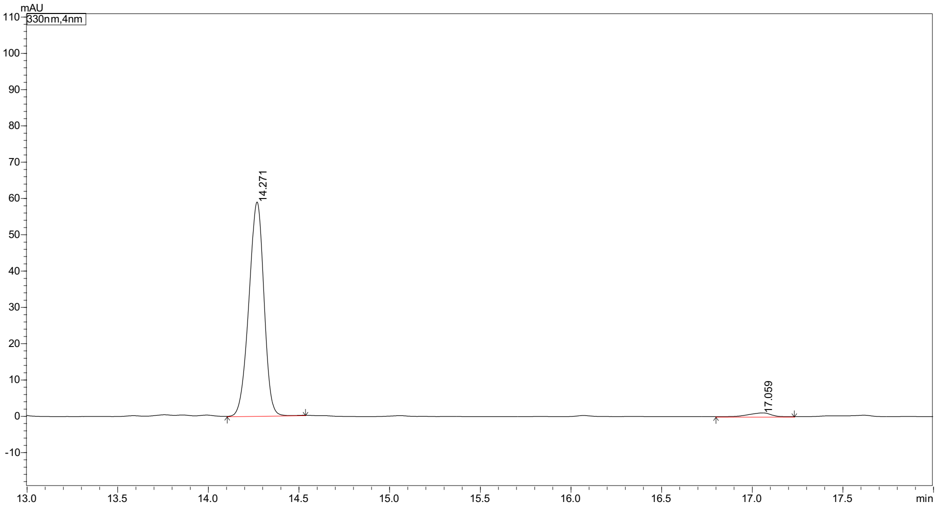

| Peak# | Peak Start | Peak End | Ret. Time | Area | Area% |
|---|---|---|---|---|---|
| 1 | 14.104 | 14.536 | 14.271 | 339417 | 97.581 |
| 2 | 16.800 | 17.232 | 17.059 | 8415 | 2.419 |
| Total | | | | 347832 | 100.000 |

Marfey's analysis of D-**5h**

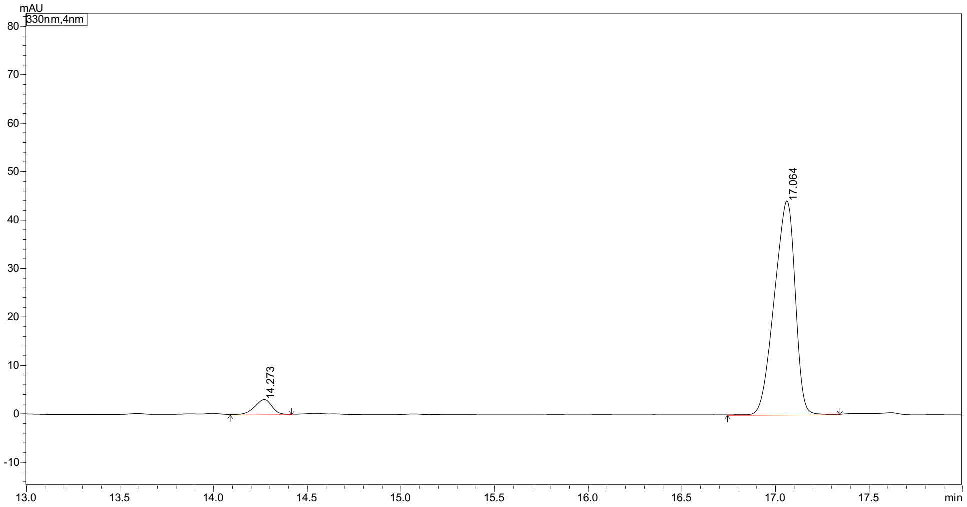

| Peak# | Peak Start | Peak End | Ret. Time | Area | Area% |
|---|---|---|---|---|---|
| 1 | 14.088 | 14.416 | 14.273 | 19005 | 5.310 |
| 2 | 16.744 | 17.344 | 17.064 | 338932 | 94.690 |
| Total | | | | 357937 | 100.000 |

#### 2-Amino-3-cyclopentylpropanoic acid (5i)

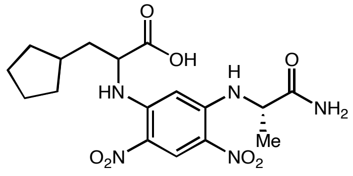

($rac$)-**5i** derivative

Marfey's analysis of ($rac$)-**5i**

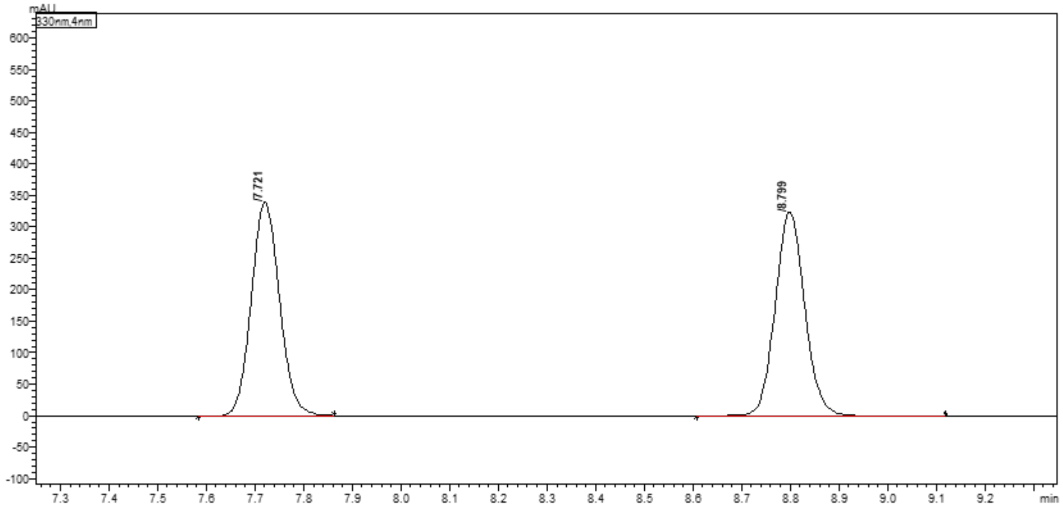

Peak Table | Compound | Group | Calibration Curve

| Peak# | Ret. Time | Height | Area | Area% |
|---|---|---|---|---|
| 1 | 7.721 | 336553 | 1358699 | 49.673 |
| 2 | 8.799 | 323929 | 1376614 | 50.327 |
| Total | | 660482 | 2735314 | 100.000 |

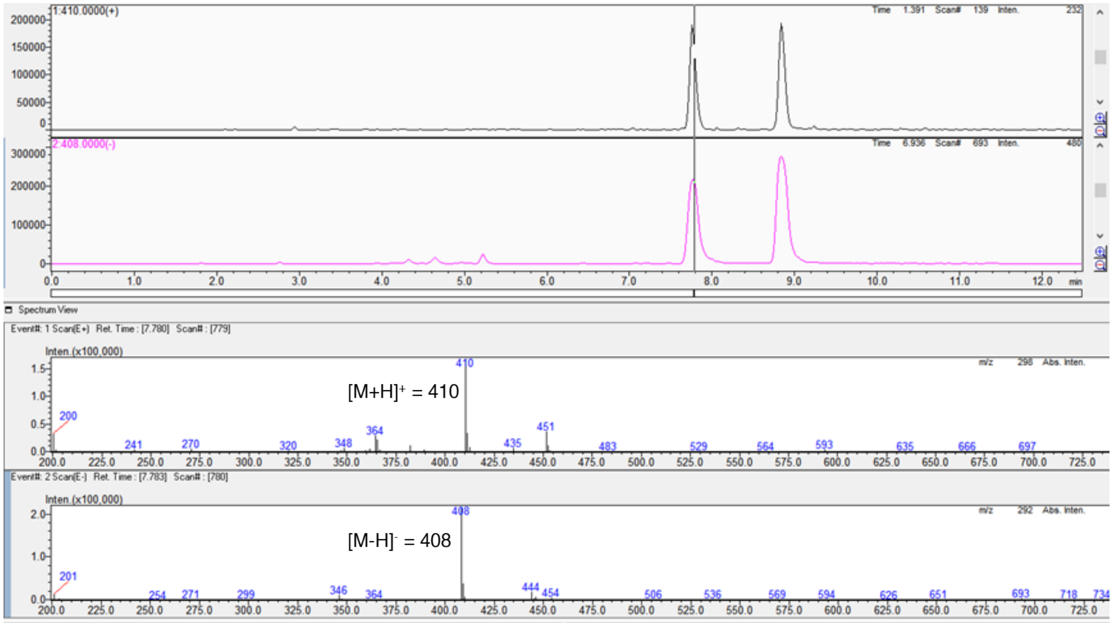

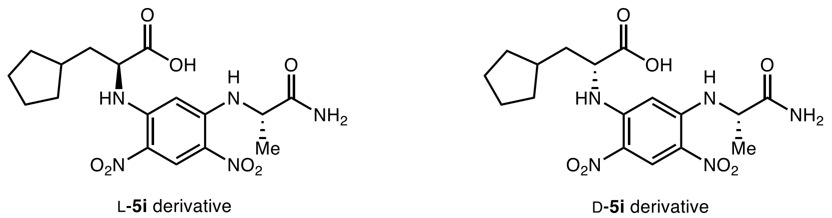

Marfey’s analysis of L-**5i**

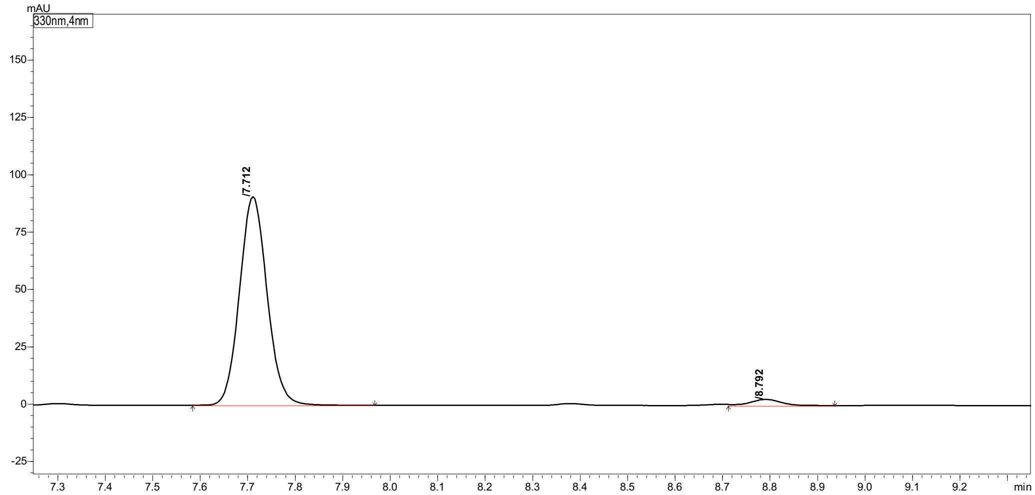

| Peak# | Ret. Time | Height | Area | Area% |
|---|---|---|---|---|
| 1 | 7.712 | 90451 | 367184 | 96.873 |
| 2 | 8.792 | 2673 | 11854 | 3.127 |
| Total | | 93124 | 379038 | 100.000 |

Marfey’s analysis of D-**5i**

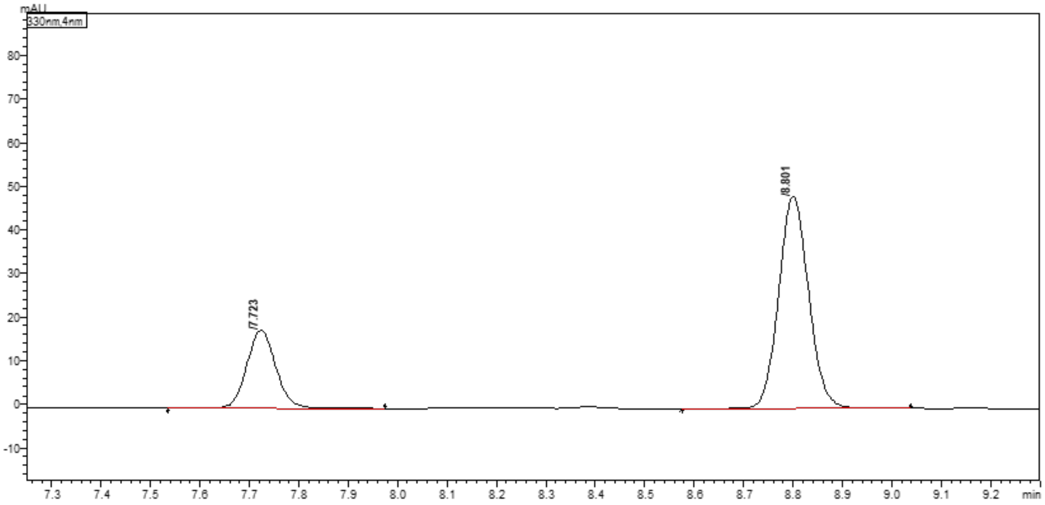

| Peak# | Ret. Time | Height | Area | Area% |
|---|---|---|---|---|
| 1 | 7.723 | 17760 | 73237 | 26.347 |
| 2 | 8.801 | 48253 | 204735 | 73.653 |
| Total | | 66012 | 277972 | 100.000 |

#### 2-Aminopentanoic acid (5k)

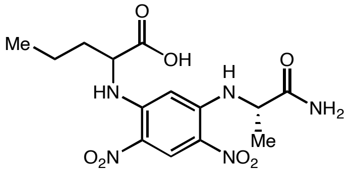

(*rac*)-**5k** derivative

#### Marfey's analysis of (*rac*)-5k

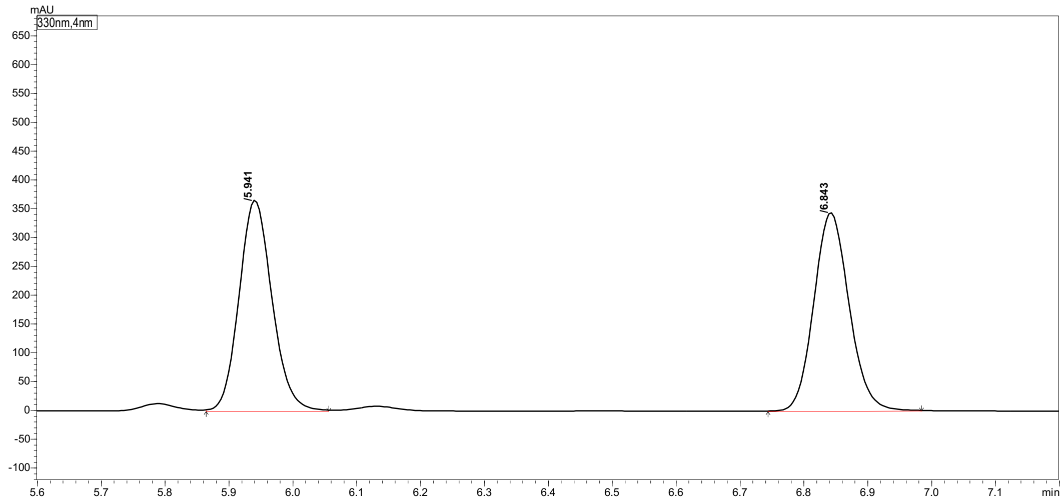

Peak Table | Compound | Group | Calibration Curve

| Peak# | Ret. Time | Height | Area | Area% |
|---|---|---|---|---|
| 1 | 5.941 | 363924 | 1311731 | 49.946 |
| 2 | 6.843 | 339629 | 1314554 | 50.054 |
| Total | | 703553 | 2626285 | 100.000 |

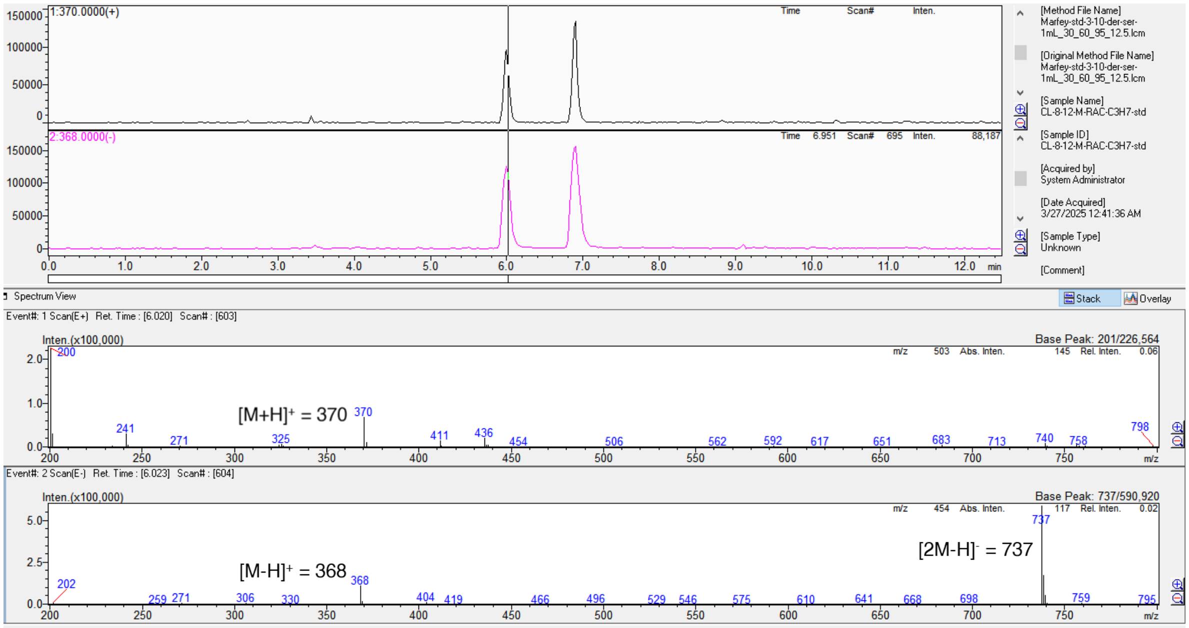

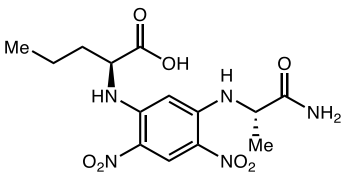

L-**5k** derivative

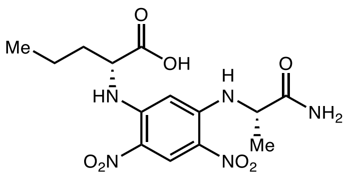

D-**5k** derivative

#### Marfey's analysis of L-**5k**

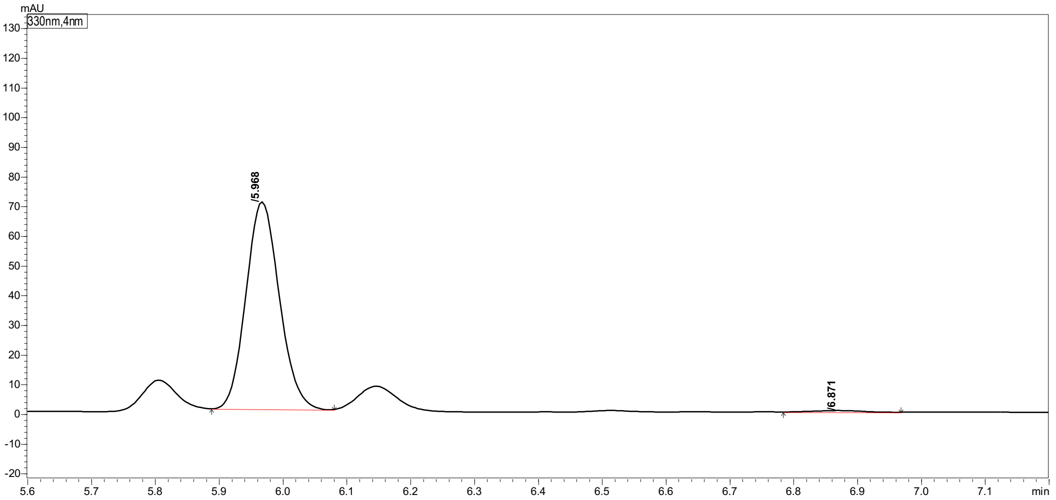

| Peak# | Ret. Time | Height | Area | Area% |
|---|---|---|---|---|
| 1 | 5.968 | 69459 | 253717 | 98.893 |
| 2 | 6.871 | 544 | 2841 | 1.107 |
| Total | | 70003 | 256558 | 100.000 |

#### Marfey's analysis of D-**5k**

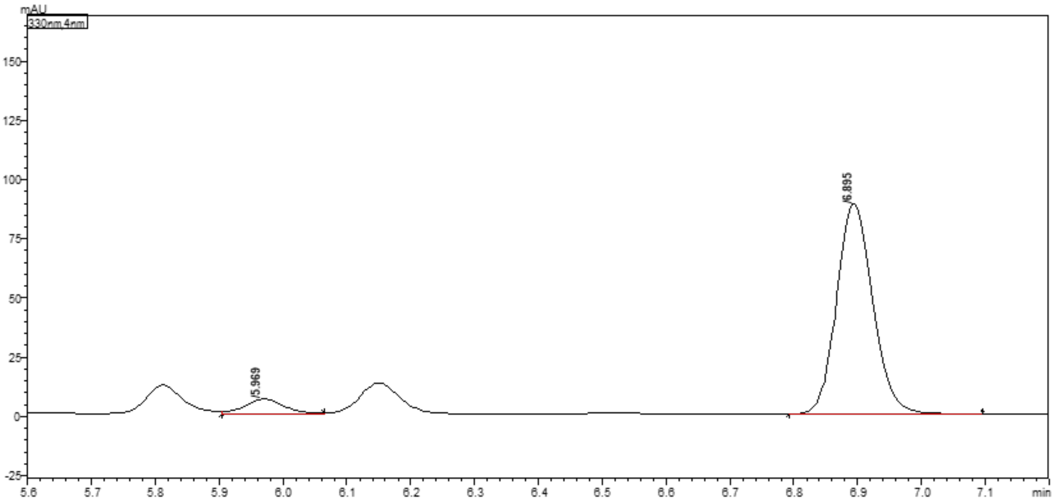

| Peak# | Ret. Time | Height | Area | Area% |
|---|---|---|---|---|
| 1 | 5.969 | 6128 | 25186 | 6.721 |
| 2 | 6.895 | 88695 | 349527 | 93.279 |
| Total | | 94822 | 374713 | 100.000 |

#### 2-Aminohexanoic acid (5l)

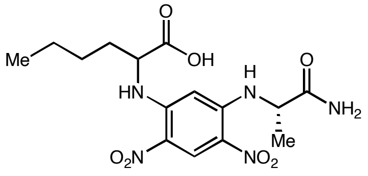

(*rac*)-**5l** derivative

#### Marfey's analysis of (*rac*)-**5l**

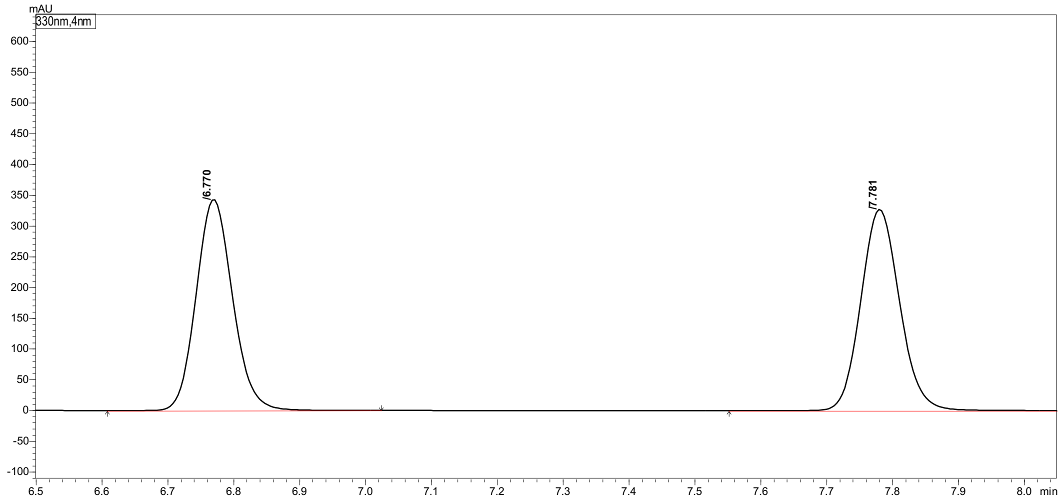

| Peak# | Ret. Time | Height | Area | Area% |
|---|---|---|---|---|
| 1 | 6.770 | 337555 | 1316960 | 49.721 |
| 2 | 7.781 | 325845 | 1331727 | 50.279 |
| Total | | 663400 | 2648687 | 100.000 |

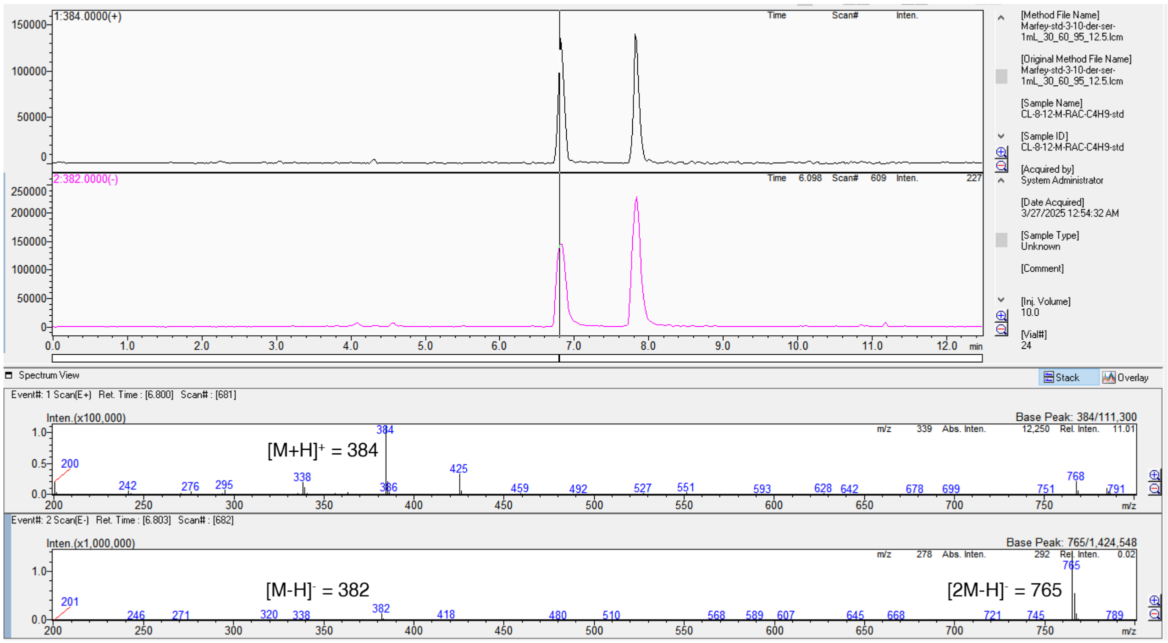

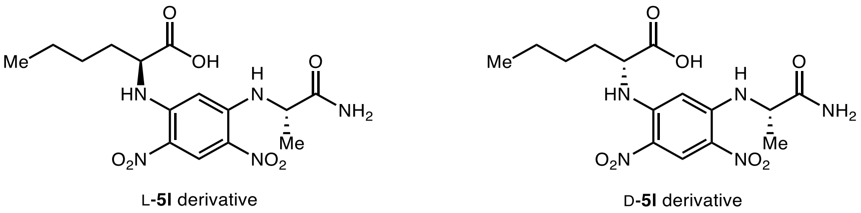

Marfey's analysis of L-**5l**

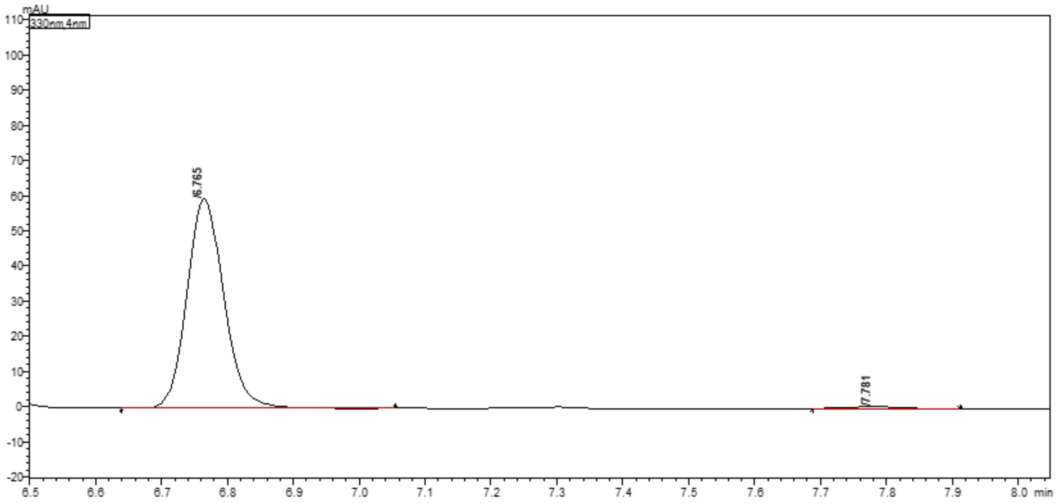

| Peak# | Ret. Time | Height | Area | Area% |
|---|---|---|---|---|
| 1 | 6.765 | 59452 | 231639 | 98.788 |
| 2 | 7.781 | 619 | 2842 | 1.212 |
| Total | | 60070 | 234480 | 100.000 |

Marfey's analysis of D-**5l**

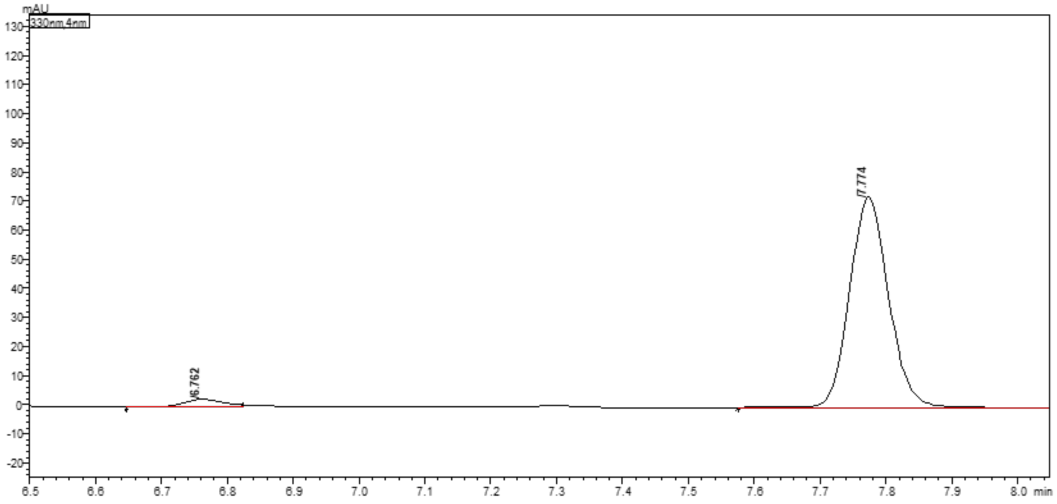

| Peak# | Ret. Time | Height | Area | Area% |
|---|---|---|---|---|
| 1 | 6.762 | 2663 | 11375 | 3.708 |
| 2 | 7.774 | 72127 | 295415 | 96.292 |
| Total | | 74790 | 306790 | 100.000 |

#### 2-Aminoheptanoic acid (5m)

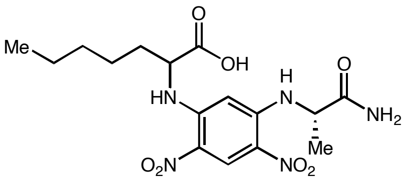

(*rac*)-**5m** derivative

#### Marfey’s analysis of (*rac*)-**5m**

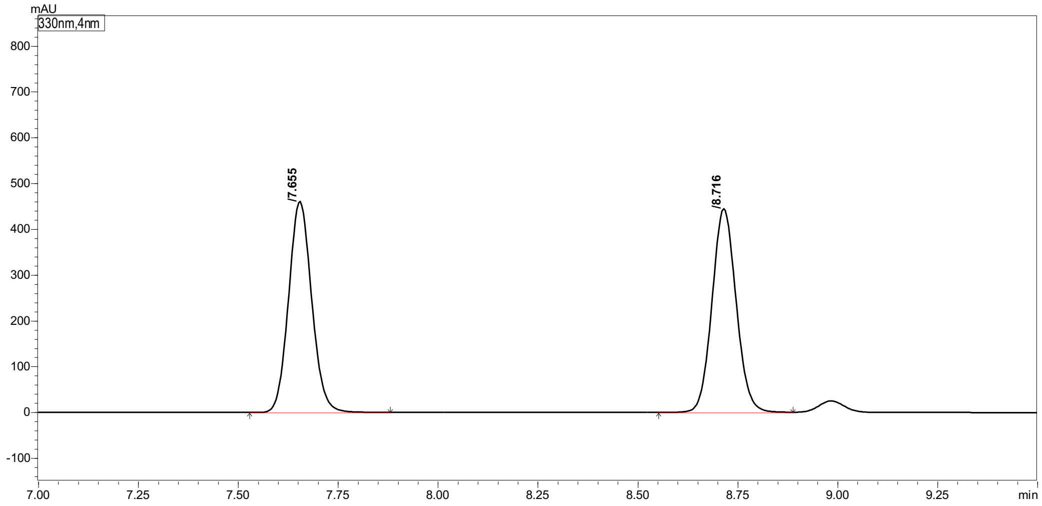

Peak Table | Compound | Group | Calibration Curve

| Peak# | Ret. Time | Height | Area | Area% |
|---|---|---|---|---|
| 1 | 7.655 | 459110 | 1849354 | 49.676 |
| 2 | 8.716 | 442834 | 1873455 | 50.324 |
| Total | | 901944 | 3722809 | 100.000 |

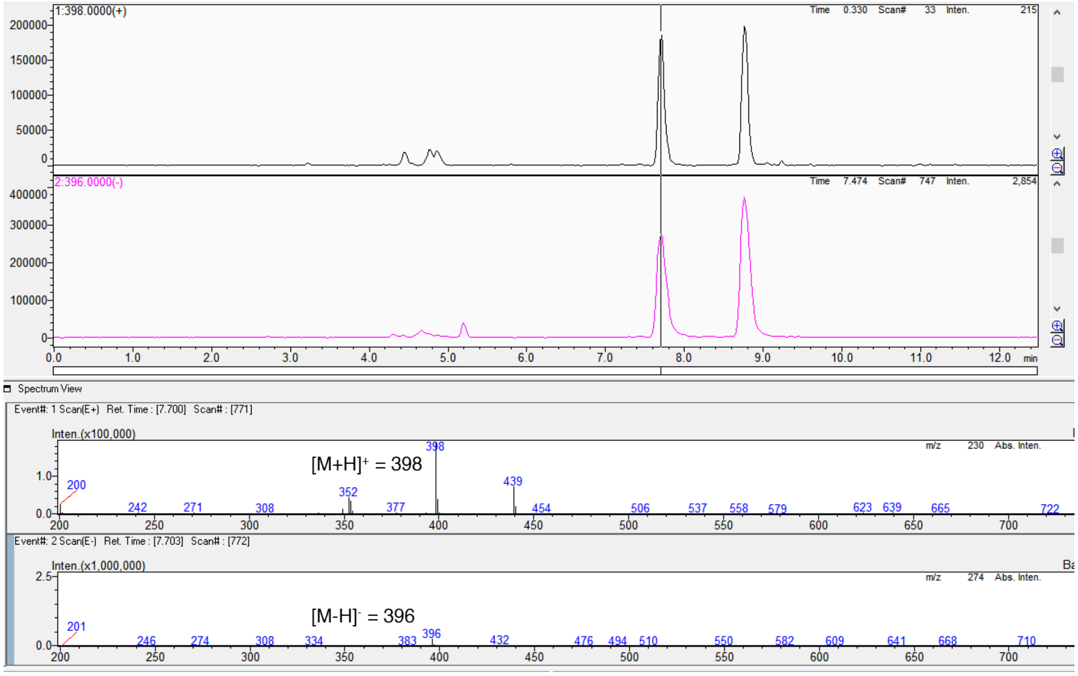

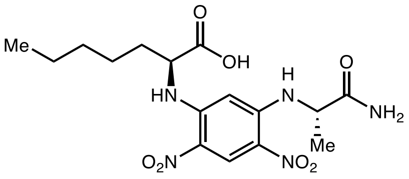

L-**5m** derivative

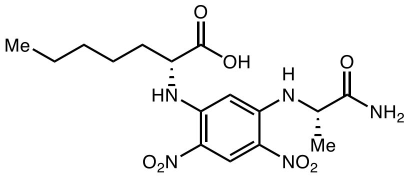

D-**5m** derivative

#### Marfey's analysis of L-**5m**

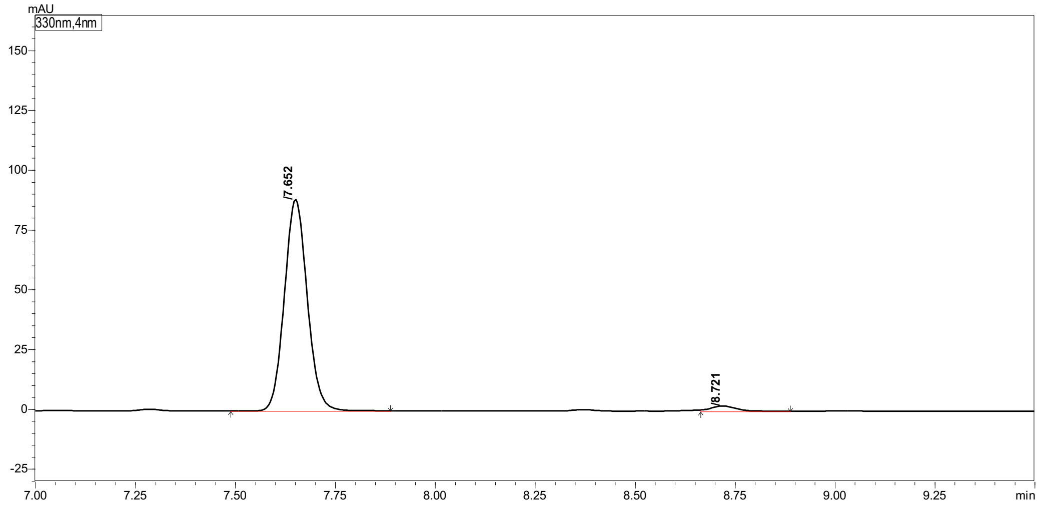

Peak Table | Compound | Group | Calibration Curve

| Peak# | Ret. Time | Height | Area | Area% |
|---|---|---|---|---|
| 1 | 7.652 | 87642 | 351333 | 97.514 |
| 2 | 8.721 | 2109 | 8956 | 2.486 |
| Total | | 89751 | 360289 | 100.000 |

#### Marfey's analysis of D-**5m**

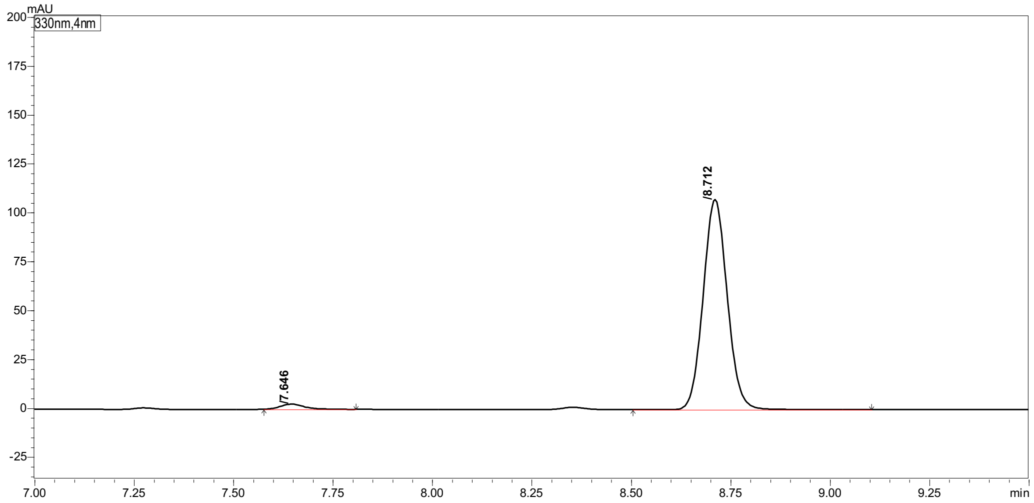

Peak Table | Compound | Group | Calibration Curve

| Peak# | Ret. Time | Height | Area | Area% |
|---|---|---|---|---|
| 1 | 7.646 | 2639 | 11120 | 2.405 |
| 2 | 8.712 | 107031 | 451328 | 97.595 |
| Total | | 109671 | 462448 | 100.000 |

#### 2-Aminononanoic acid (5n)

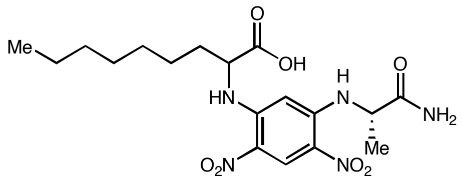

(*rac*)-**5n** derivative

#### Marfey's analysis of (*rac*)-**5n**

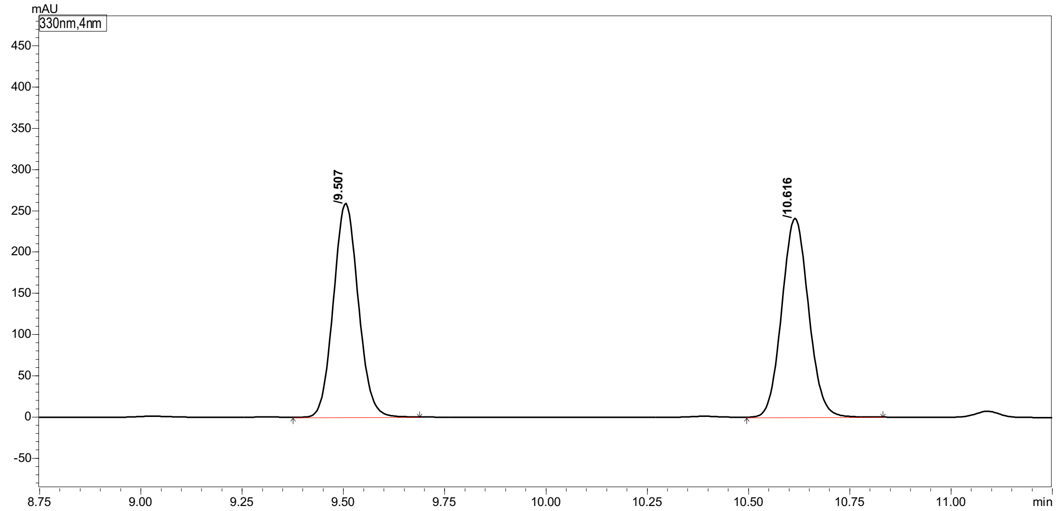

Peak Table | Compound | Group | Calibration Curve

| Peak# | Ret. Time | Height | Area | Area% |
|---|---|---|---|---|
| 1 | 9.507 | 256829 | 1104767 | 49.935 |
| 2 | 10.616 | 240253 | 1107644 | 50.065 |
| Total | | 497082 | 2212411 | 100.000 |

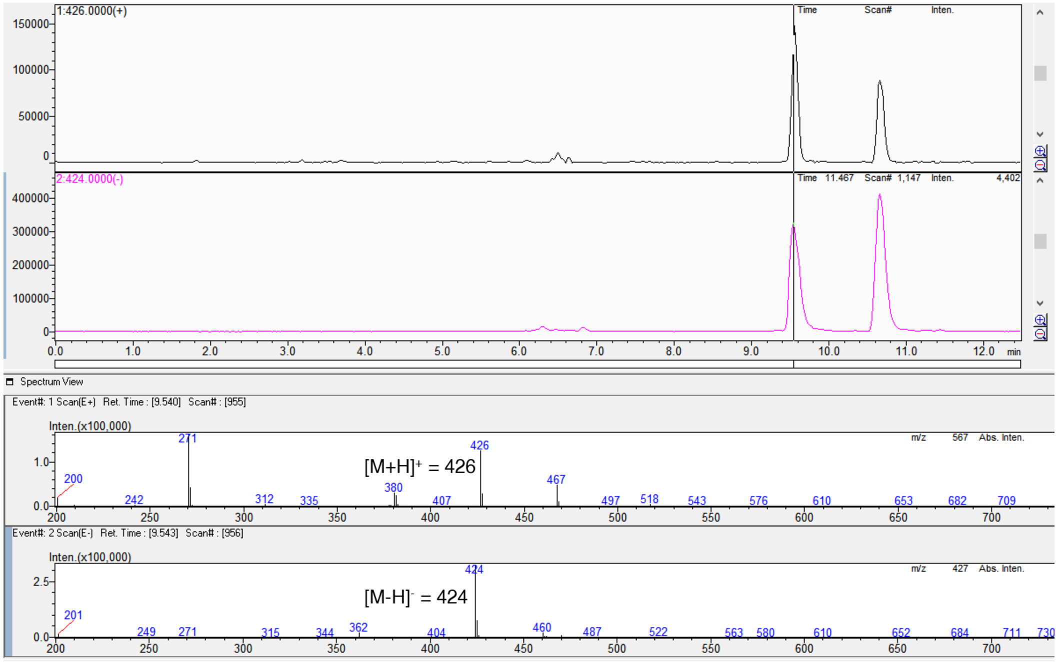

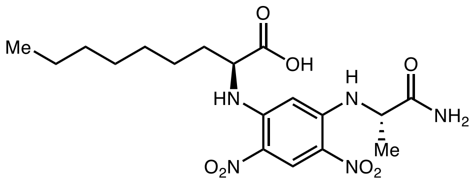

L-**5n** derivative

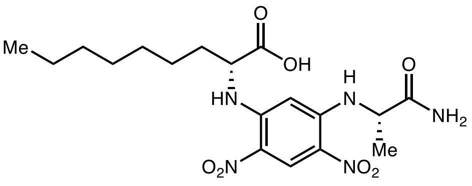

D-**5n** derivative

#### Marfey's analysis of L-**5n**

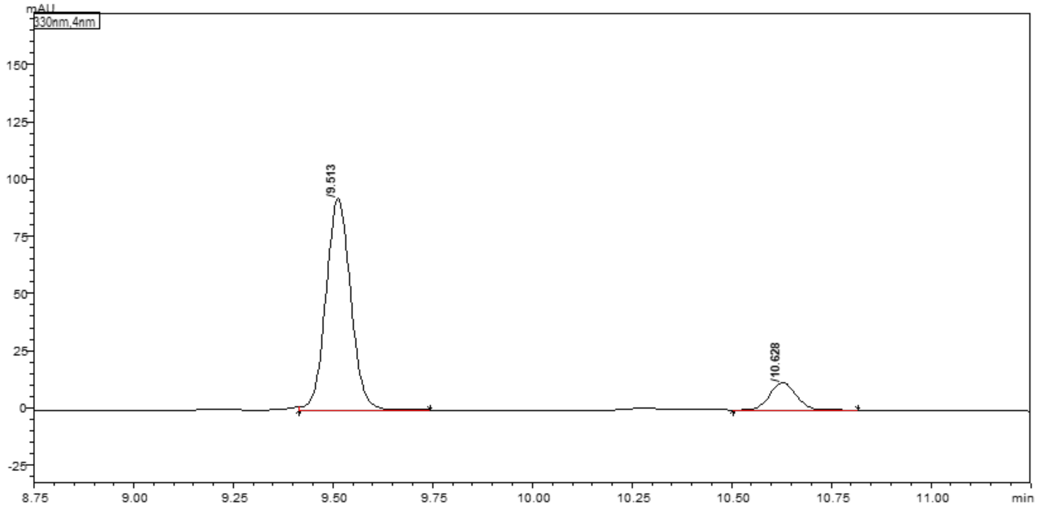

Peak Table | Compound | Group | Calibration Curve

| Peak# | Ret. Time | Height | Area | Area% |
|---|---|---|---|---|
| 1 | 9.513 | 91806 | 395478 | 87.902 |
| 2 | 10.628 | 11942 | 54430 | 12.098 |
| Total | | 103748 | 449908 | 100.000 |

#### Marfey's analysis of D-**5n**

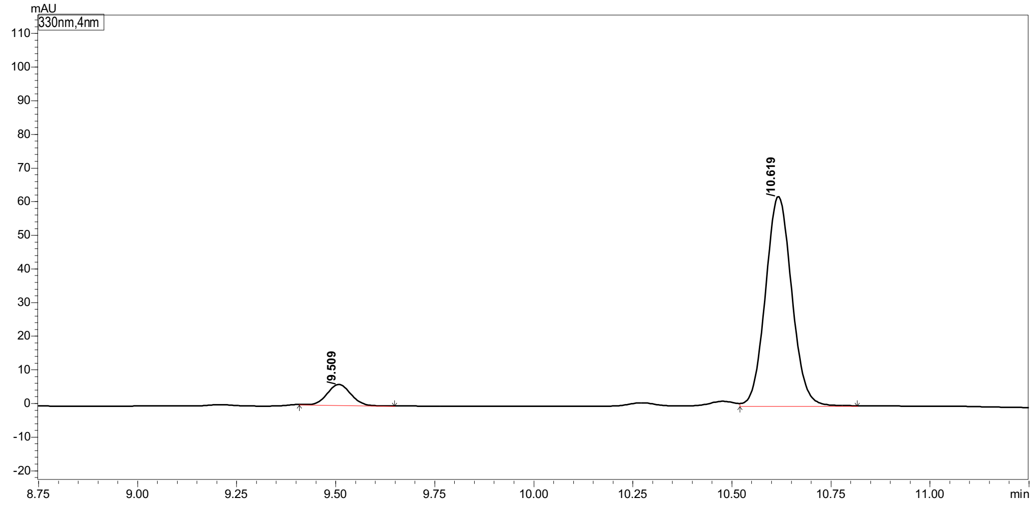

Peak Table | Compound | Group | Calibration Curve

| Peak# | Ret. Time | Height | Area | Area% |
|---|---|---|---|---|
| 1 | 9.509 | 6140 | 24912 | 8.014 |
| 2 | 10.619 | 61768 | 285957 | 91.986 |
| Total | | 67908 | 310869 | 100.000 |

#### 2-Aminoundecanoic acid (5o)

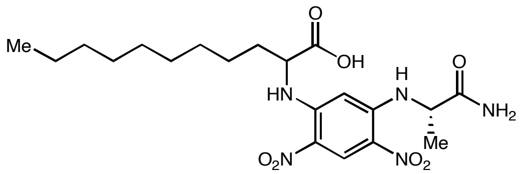

(*rac*)-**5o** derivative

Marfey's analysis of (*rac*)-**5o**

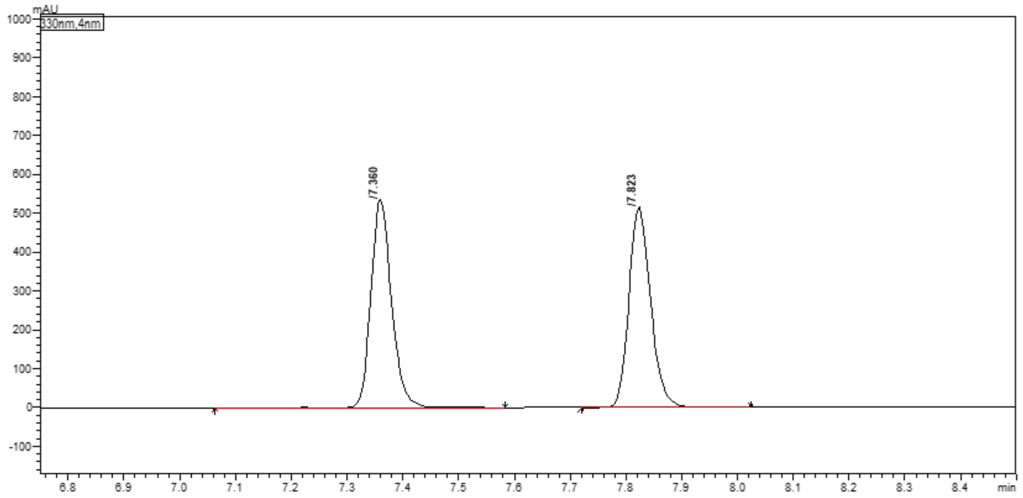

Peak Table | Compound | Group | Calibration Curve

| Peak# | Ret. Time | Height | Area | Area% |
|---|---|---|---|---|
| 1 | 7.360 | 529404 | 1369853 | 49.648 |
| 2 | 7.823 | 512487 | 1389255 | 50.352 |
| Total | | 1041891 | 2759108 | 100.000 |

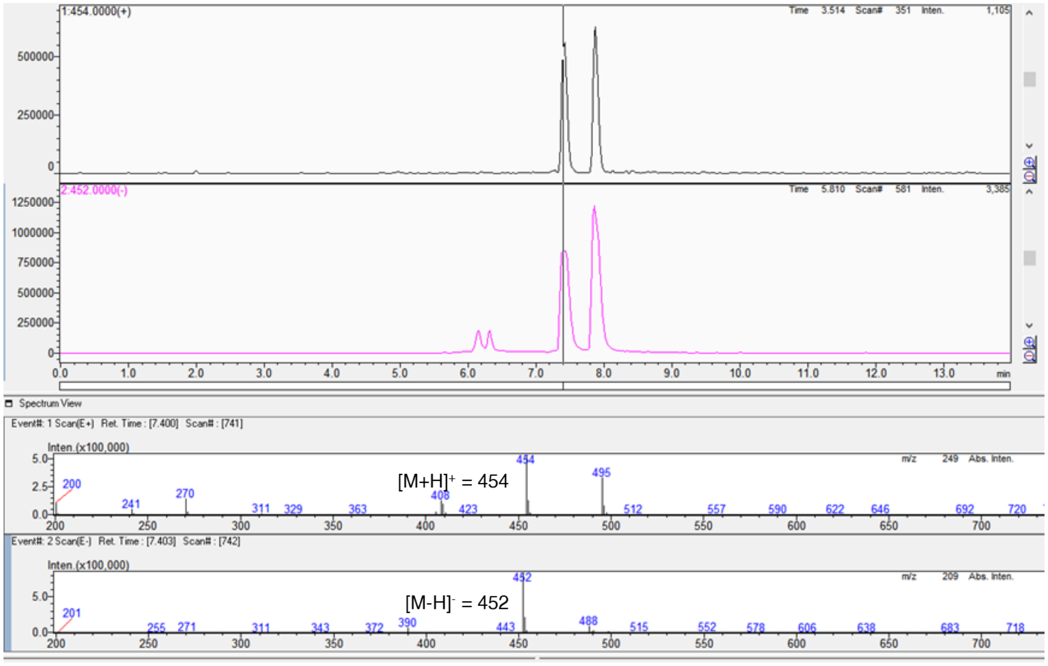

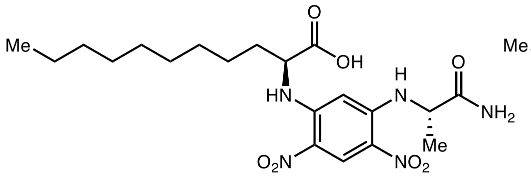

L-**5o** derivative

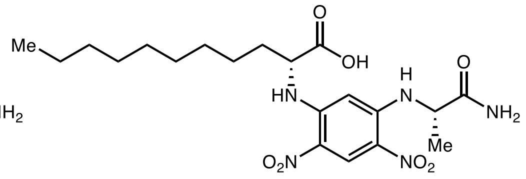

D-**5o** derivative

#### Marfey's analysis of L-**5o**

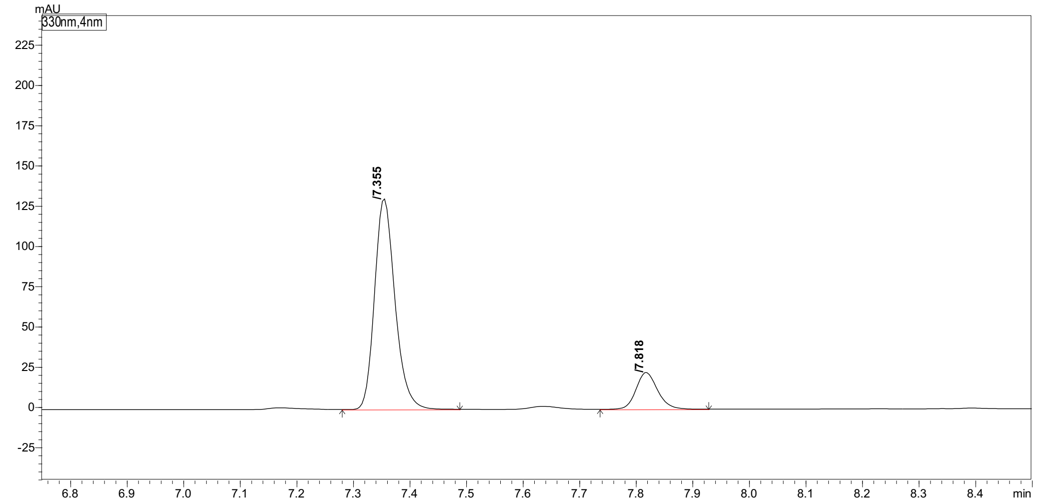

Peak Table | Compound | Group | Calibration Curve

| Peak# | Ret. Time | Height | Area | Area% |
|---|---|---|---|---|
| 1 | 7.355 | 127647 | 328629 | 84.480 |
| 2 | 7.818 | 22021 | 60371 | 15.520 |
| Total | | 149668 | 389000 | 100.000 |

#### Marfey's analysis of D-**5o**

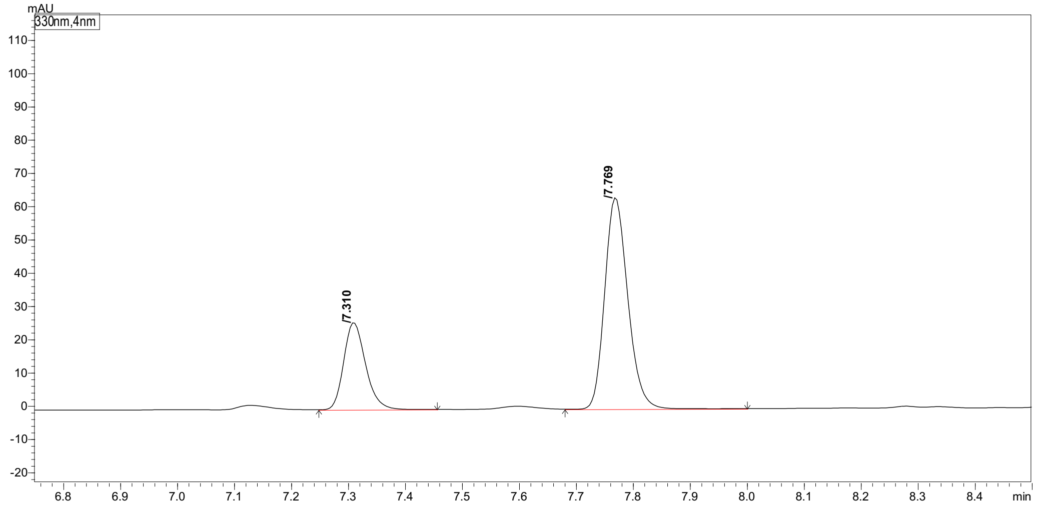

Peak Table | Compound | Group | Calibration Curve

| Peak# | Ret. Time | Height | Area | Area% |
|---|---|---|---|---|
| 1 | 7.310 | 26025 | 69475 | 27.976 |
| 2 | 7.769 | 62342 | 178862 | 72.024 |
| Total | | 88367 | 248337 | 100.000 |

#### 2-Aminotridecanoic acid (5p)

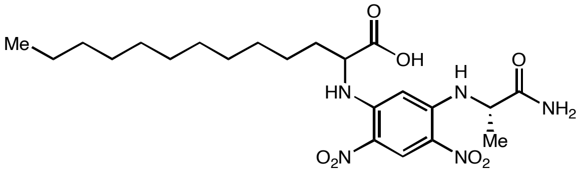

(*rac*)-**5p** derivative

Marfey's analysis of (*rac*)-**5p**

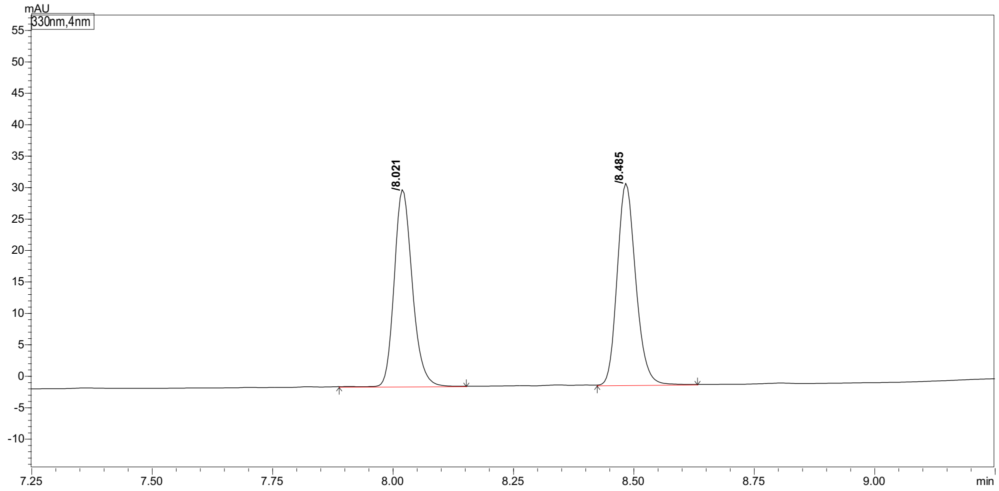

Peak Table | Compound | Group | Calibration Curve

| Peak# | Ret. Time | Height | Area | Area% |
|---|---|---|---|---|
| 1 | 8.021 | 31018 | 81272 | 48.644 |
| 2 | 8.485 | 31731 | 85801 | 51.356 |
| Total | | 62749 | 167073 | 100.000 |

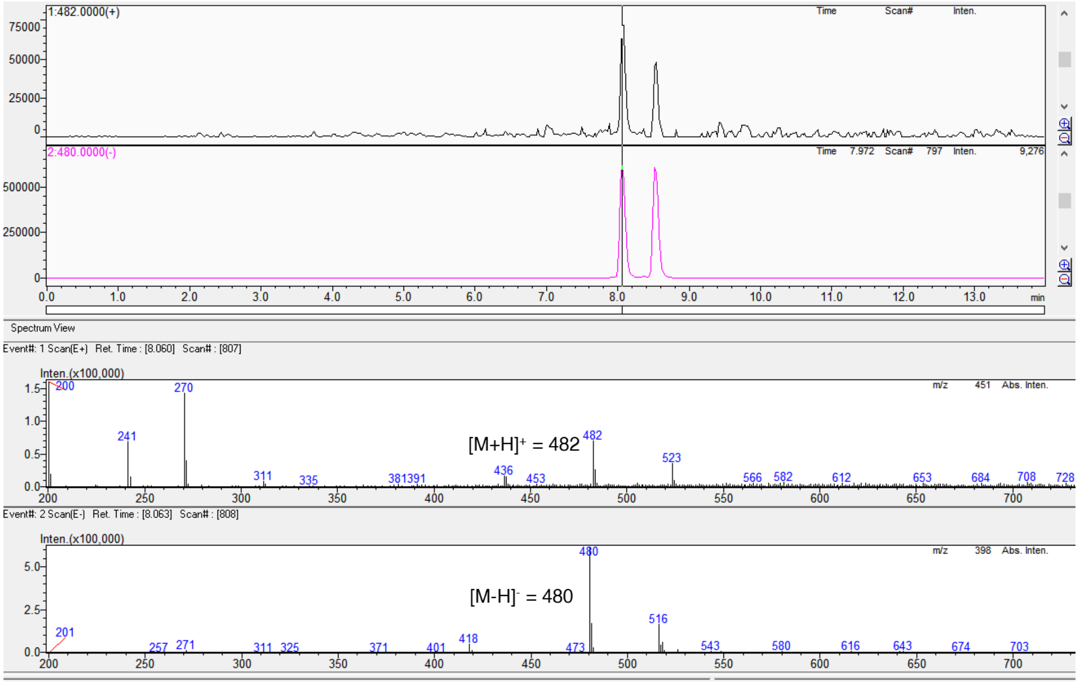

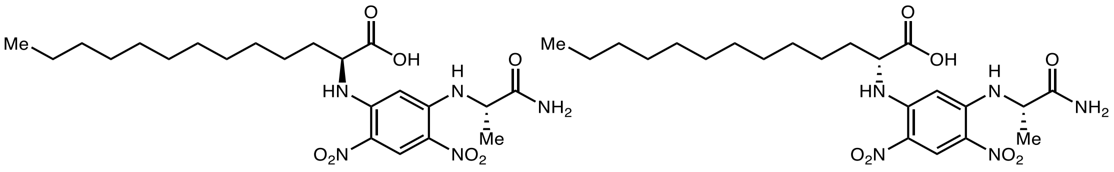

L-**5g** derivative D-**5g** derivative

Marfey's analysis of L-**5p**

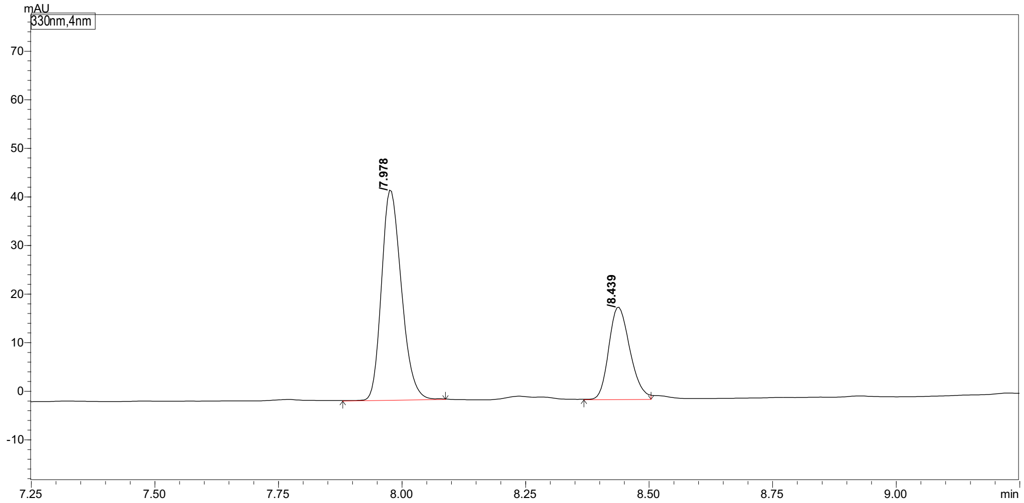

| Peak# | Ret. Time | Height | Area | Area% |
|---|---|---|---|---|
| 1 | 7.978 | 42192 | 120464 | 68.218 |
| 2 | 8.439 | 18871 | 56122 | 31.782 |
| Total | | 61063 | 176586 | 100.000 |

Marfey's analysis of D-**5p**

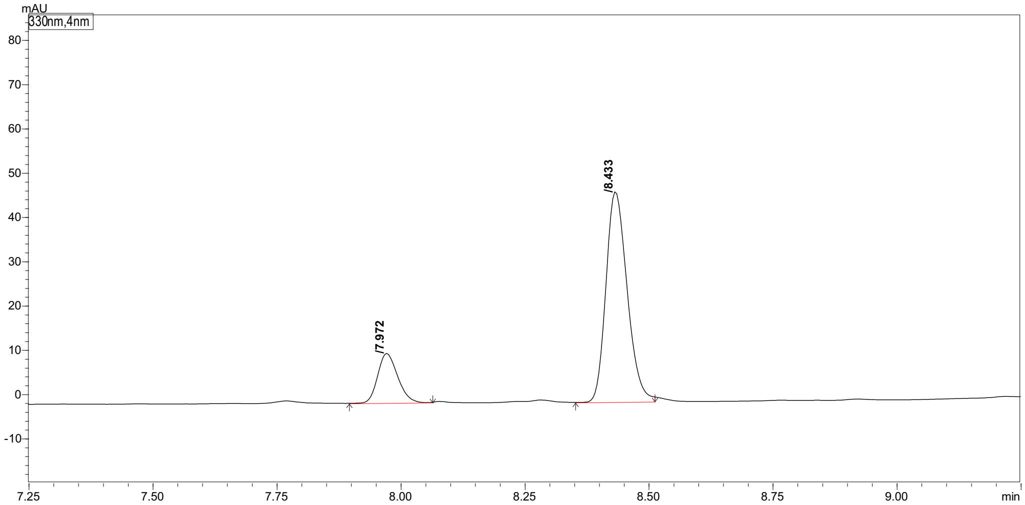

| Peak# | Ret. Time | Height | Area | Area% |
|---|---|---|---|---|
| 1 | 7.972 | 11058 | 31029 | 18.036 |
| 2 | 8.433 | 46612 | 141013 | 81.964 |
| Total | | 57670 | 172042 | 100.000 |

#### 2-Amino-4-cyclopentylbutanoic acid (5r)

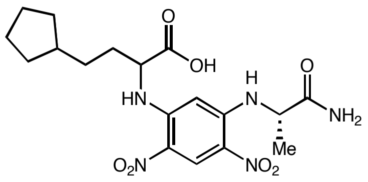

(*rac*)-**5r** derivative

Marfey's analysis of (*rac*)-**5q** (Long HPLC **method b** was used for separation)

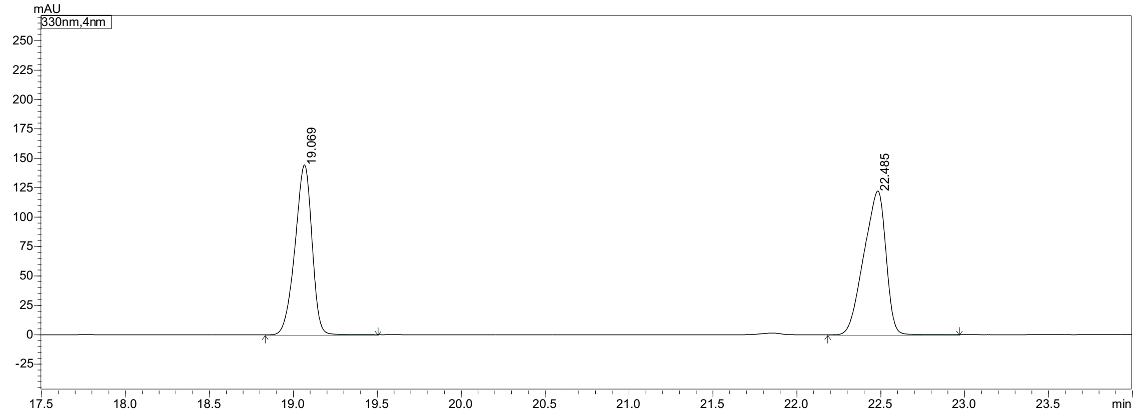

| Peak# | Peak Start | Peak End | Ret. Time | Area | Area% |
|---|---|---|---|---|---|
| 1 | 18.832 | 19.504 | 19.069 | 1011042 | 47.579 |
| 2 | 22.184 | 22.968 | 22.485 | 1113937 | 52.421 |
| Total | | | | 2124979 | 100.000 |

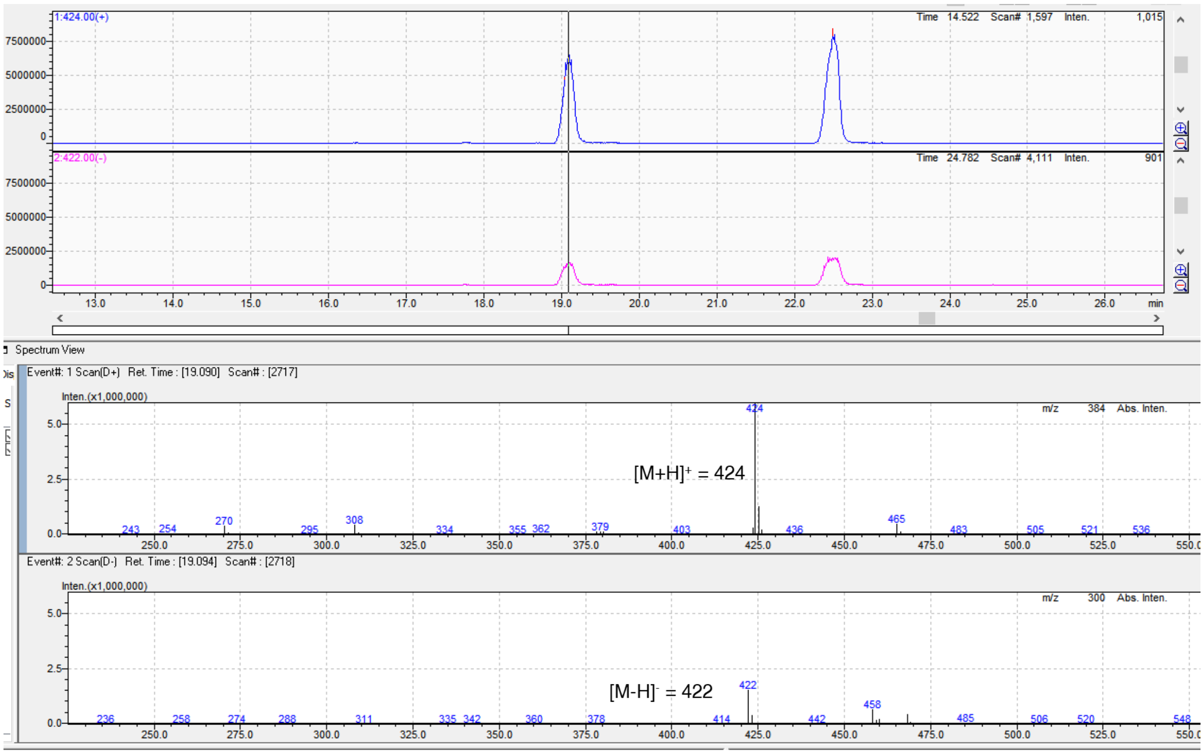

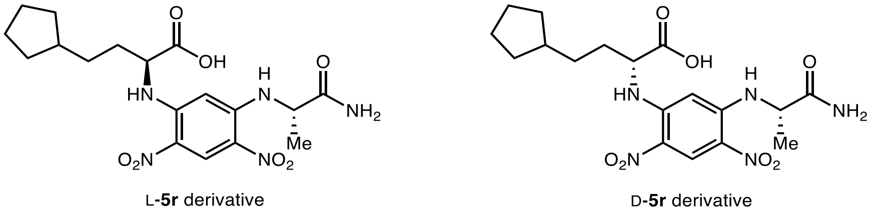

Marfey's analysis of L-**5r**

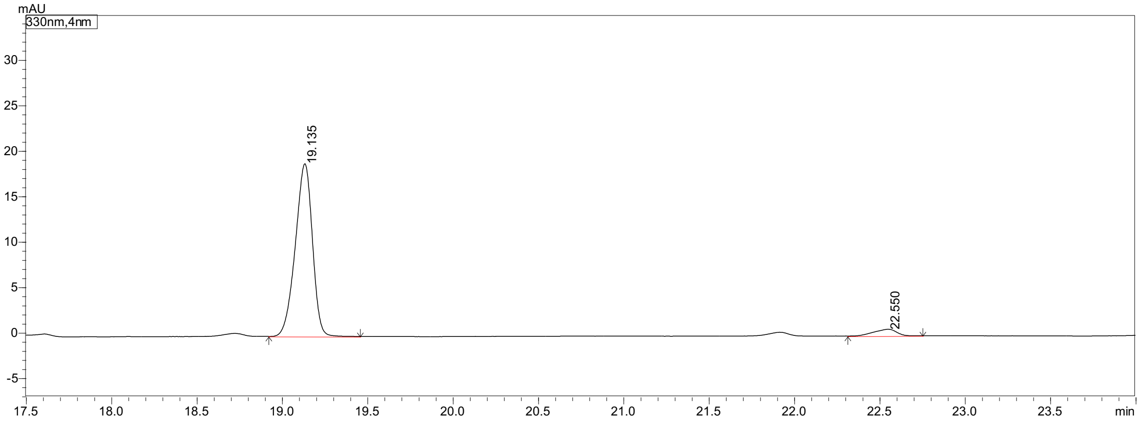

| Peak# | Peak Start | Peak End | Ret. Time | Area | Area% |
|---|---|---|---|---|---|
| 1 | 18.920 | 19.456 | 19.135 | 132015 | 95.216 |
| 2 | 22.312 | 22.752 | 22.550 | 6632 | 4.784 |
| Total | | | | 138647 | 100.000 |

Marfey's analysis of D-**5r**

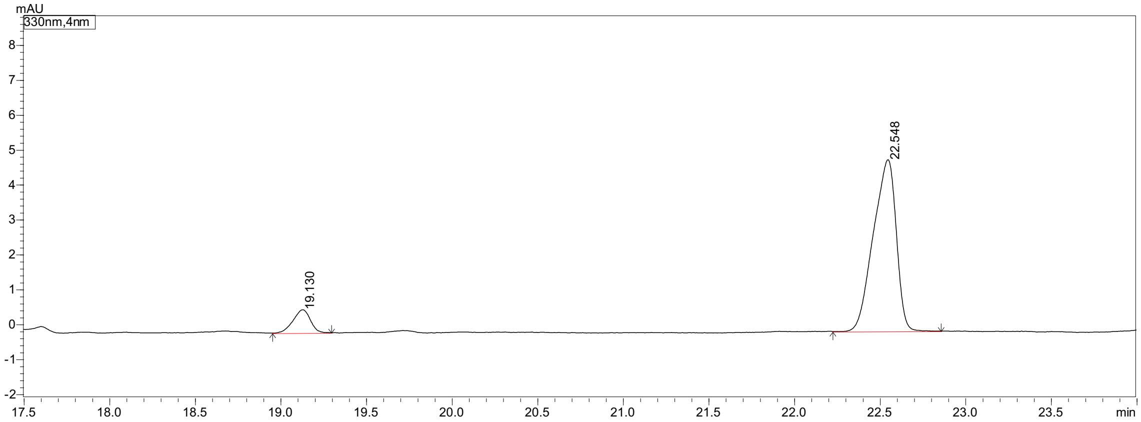

| Peak# | Peak Start | Peak End | Ret. Time | Area | Area% |
|---|---|---|---|---|---|
| 1 | 18.952 | 19.296 | 19.130 | 4631 | 9.201 |
| 2 | 22.224 | 22.856 | 22.548 | 45699 | 90.799 |
| Total | | | | 50330 | 100.000 |

#### X. Computational Details

##### Density functional theory (DFT) and local coupled-cluster (DLPNO-CCSD(T)) calculations of the lifetime of unstabilized radical intermediates in photobiocatalytic C–C coupling

To probe the lifetime of unstabilized radical intermediates in ring opening of **4q-R1** (Figure 5a), cyclization of **4r-R1** (Figure 5b), 1,2-aryl migration of **1r-R1** (Figure 5c), 1,5-HAT of **4n-R1** (Figure 5d), we performed DFT and DLPNO-CCSD(T) calculations at the DLPNO-CCSD(T)/def2-TZVPP/SMD(water)//M06-2X/def2-TZVP level of theory using the Gaussian 16[27] and ORCA 6.0.1[28] software packages. Geometries were optimized in the gas phase using the (U)M06-2X functional[29] with the def2-TZVP basis set. Vibrational frequency calculations were performed at the same level of theory as geometry optimization to confirm that the obtained transition state connects its respective minima. Grimme's quasi-harmonic approximations[30] were applied for entropy calculations using 100 $cm^{-1}$ as the frequency cutoff at the experimental temperature (323 K) using the GoodVibes software package[31]. Single-point energies were calculated with the DLPNO-CCSD(T) method[32,33,34] with the "tightPNO" keyword and the def2-TZVPP basis set. Solvation effects were considered in the single-point energy calculations using the SMD model[35] in water. The lifetimes of these radical intermediates at 323 K were calculated by converting the computed activation Gibbs free energy barriers using the Eyring–Polanyi equation[36,37]. As shown in Figure 5d, the activation barrier from the most stable linear conformer of the 1-hexyl radical (**4n-R1**) is 17.4 kcal/mol, and the corresponding rate constant at 323 K is 11.8 $s^{-1}$. However, because the bent conformer **4n-R2** with two gauche C–C bonds is expected to be more stable than the linear conformer **4n-R1** in the enzyme active site, we used the bent conformer **4n-R2** as the energy zero to calculate the 1,5-HAT barrier (15.5 kcal/mol) and the rate constant at 323 K ($2.3 \times 10^2$ $s^{-1}$).

##### Cartesian coordinates of all DFT-optimized structures

### 1r-R1

M06-2X SCF energy: -349.478668 a.u.

M06-2X enthalpy: -349.296180 a.u.

M06-2X Gibbs free energy: -349.344705 a.u.

DLPNO-CCSD(T) SCF energy in solution: -348.857680 a.u.

DLPNO-CCSD(T) enthalpy in solution: -348.675192 a.u.

DLPNO-CCSD(T) Gibbs free energy in solution: -348.723717 a.u.

Cartesian coordinates

| ATOM | X | Y | Z |
|---|---|---|---|
| C | -3.12140200 | 0.21308600 | -0.00004800 |
| H | -3.19731600 | 0.85328000 | 0.88090800 |
| H | -3.19725700 | 0.85323900 | -0.88103900 |
| H | -3.97951300 | -0.45935400 | -0.00006100 |
| C | -1.80616200 | -0.55531100 | 0.00002700 |
| H | -1.76135000 | -1.20996200 | 0.87517300 |
| H | -1.76129700 | -1.21000900 | -0.87508000 |
| C | -0.59024000 | 0.36332100 | 0.00003800 |
| H | -0.63297300 | 1.01978900 | -0.87579100 |
| H | -0.63296900 | 1.01976200 | 0.87588800 |
| C | 0.73184000 | -0.39304700 | 0.00001000 |
| H | 0.78379800 | -1.04657300 | -0.87598700 |
| H | 0.78380200 | -1.04661500 | 0.87597700 |
| C | 1.95176100 | 0.54460200 | 0.00002500 |
| H | 1.88561600 | 1.19345600 | 0.87833400 |
| H | 1.88557300 | 1.19354900 | -0.87821300 |
| C | 3.24522600 | -0.18638200 | -0.00004500 |
| H | 3.67885800 | -0.53902800 | -0.92465600 |
| H | 3.67888600 | -0.53915100 | 0.92450600 |

**1r-R2**

M06-2X SCF energy: -349.465517 a.u.

M06-2X enthalpy: -349.282609 a.u.

M06-2X Gibbs free energy: -349.329514 a.u.

DLPNO-CCSD(T) SCF energy in solution: -348.844587 a.u.

DLPNO-CCSD(T) enthalpy in solution: -348.661679 a.u.

DLPNO-CCSD(T) Gibbs free energy in solution: -348.708584 a.u.

Cartesian coordinates

| ATOM | X | Y | Z |
|---|---|---|---|
| C | 2.46368100 | 0.35171500 | -0.19519600 |
| C | 1.55906500 | 1.35529000 | 0.20414500 |
| C | 0.24078100 | 1.08379800 | 0.41010200 |
| C | -0.29852400 | -0.26562200 | 0.21989400 |
| C | 0.68480700 | -1.27811300 | -0.17548000 |
| C | 1.99345500 | -0.96569200 | -0.37362800 |
| H | 3.50576600 | 0.58720400 | -0.35597300 |
| H | 1.92050800 | 2.36483000 | 0.35642100 |
| H | -0.43020800 | 1.86967000 | 0.73170100 |
| H | 0.33269000 | -2.29452300 | -0.31210200 |
| H | 2.68636800 | -1.74272100 | -0.67172000 |
| C | -1.73107600 | -0.44428100 | -0.32916100 |
| C | -1.47164600 | -0.72426100 | 1.09954200 |
| H | -1.82157400 | -0.00106200 | 1.82530100 |
| H | -1.43705400 | -1.74746600 | 1.44777100 |
| C | -2.54758500 | 0.74311700 | -0.76381800 |
| H | -2.19021900 | 1.14214300 | -1.71414700 |
| H | -3.59270900 | 0.45414600 | -0.88662200 |
| H | -2.51416800 | 1.54443800 | -0.02664400 |
| H | -1.81714700 | -1.31236700 | -0.97238500 |

**4q-R1**

M06-2X SCF energy: -156.521525 a.u.

M06-2X enthalpy: -156.419438 a.u.

M06-2X Gibbs free energy: -156.456234 a.u.

DLPNO-CCSD(T) SCF energy in solution: -156.246600 a.u.

DLPNO-CCSD(T) enthalpy in solution: -156.144513 a.u.

DLPNO-CCSD(T) Gibbs free energy in solution: -156.181309 a.u.

Cartesian coordinates

| ATOM | X | Y | Z |
|---|---|---|---|
| C | 0.89903700 | 0.74402700 | -0.13676400 |
| C | 0.89895800 | -0.74410500 | -0.13699800 |
| C | -0.26430800 | 0.00002400 | 0.49267600 |
| H | 1.57849700 | 1.25870500 | 0.52754600 |
| H | 0.69641700 | 1.25364100 | -1.06826600 |
| H | 1.57837600 | -1.25918000 | 0.52704000 |
| H | 0.69610600 | -1.25329200 | -1.06868300 |
| H | -0.28084900 | -0.00016900 | 1.57414800 |
| C | -1.56158600 | 0.00018000 | -0.16187100 |
| H | -2.47741800 | -0.00112400 | 0.40712300 |
| H | -1.62373300 | 0.00066100 | -1.24116500 |

**4r-R1**

M06-2X SCF energy: -235.134333 a.u.

M06-2X enthalpy: -234.972414 a.u.

M06-2X Gibbs free energy: -235.018815 a.u.

DLPNO-CCSD(T) SCF energy in solution: -234.728850 a.u.

DLPNO-CCSD(T) enthalpy in solution: -234.566931 a.u.

DLPNO-CCSD(T) Gibbs free energy in solution: -234.613332 a.u.

Cartesian coordinates

| ATOM | X | Y | Z |
|---|---|---|---|
| C | -2.98432900 | -0.18459600 | 0.43719400 |

| | | | |
|---|---|---|---|
| H | -2.97445400 | 0.42687100 | 1.33271200 |
| H | -3.86827400 | -0.78267000 | 0.25838900 |
| C | -1.95768100 | -0.19276700 | -0.39954800 |
| H | -2.00289400 | -0.82125200 | -1.28604500 |
| C | -0.69401300 | 0.58767000 | -0.21500800 |
| H | -0.55028400 | 1.25737900 | -1.06987900 |
| H | -0.77810100 | 1.21847900 | 0.67384300 |
| C | 0.53269200 | -0.31791200 | -0.09256700 |
| H | 0.60355900 | -0.96412700 | -0.97226500 |
| H | 0.41300000 | -0.97834700 | 0.77001700 |
| C | 1.83773200 | 0.48352400 | 0.05204300 |
| H | 1.74853400 | 1.13065700 | 0.92948600 |
| H | 1.93588800 | 1.14022600 | -0.81749100 |
| C | 3.03564900 | -0.38575800 | 0.17604600 |
| H | 3.52215900 | -0.78277800 | -0.70326300 |
| H | 3.33057200 | -0.78540300 | 1.13553600 |

**4n-R1**

M06-2X SCF energy: -236.362376 a.u.

M06-2X enthalpy: -236.176306 a.u.

M06-2X Gibbs free energy: -236.224222 a.u.

DLPNO-CCSD(T) SCF energy in solution: -235.959917 a.u.

DLPNO-CCSD(T) enthalpy in solution: -235.773847 a.u.

DLPNO-CCSD(T) Gibbs free energy in solution: -235.821763 a.u.

Cartesian coordinates

| ATOM | X | Y | Z |
|---|---|---|---|
| C | -3.12140200 | 0.21308600 | -0.00004800 |
| H | -3.19731600 | 0.85328000 | 0.88090800 |
| H | -3.19725700 | 0.85323900 | -0.88103900 |

| | | | |
|---|---|---|---|
| H | -3.97951300 | -0.45935400 | -0.00006100 |
| C | -1.80616200 | -0.55531100 | 0.00002700 |
| H | -1.76135000 | -1.20996200 | 0.87517300 |
| H | -1.76129700 | -1.21000900 | -0.87508000 |
| C | -0.59024000 | 0.36332100 | 0.00003800 |
| H | -0.63297300 | 1.01978900 | -0.87579100 |
| H | -0.63296900 | 1.01976200 | 0.87588800 |
| C | 0.73184000 | -0.39304700 | 0.00001000 |
| H | 0.78379800 | -1.04657300 | -0.87598700 |
| H | 0.78380200 | -1.04661500 | 0.87597700 |
| C | 1.95176100 | 0.54460200 | 0.00002500 |
| H | 1.88561600 | 1.19345600 | 0.87833400 |
| H | 1.88557300 | 1.19354900 | -0.87821300 |
| C | 3.24522600 | -0.18638200 | -0.00004500 |
| H | 3.67885800 | -0.53902800 | -0.92465600 |
| H | 3.67888600 | -0.53915100 | 0.92450600 |

**4n-R2**

M06-2X SCF energy: -236.361486 a.u.

M06-2X enthalpy: -236.175573 a.u.

M06-2X Gibbs free energy: -236.222465 a.u.

DLPNO-CCSD(T) SCF energy in solution: -235.957825 a.u.

DLPNO-CCSD(T) enthalpy in solution: -235.771912 a.u.

DLPNO-CCSD(T) Gibbs free energy in solution: -235.818804 a.u.

Cartesian coordinates

| ATOM | X | Y | Z |
|---|---|---|---|
| C | 1.18015300 | 0.41621800 | -0.18069900 |
| C | 0.53842300 | -0.94110700 | 0.08800300 |
| C | -0.92700700 | -1.06790500 | -0.32410200 |

| | | | |
|---|---|---|---|
| C | -1.88012400 | -0.11369000 | 0.40674000 |
| C | -1.88966800 | 1.27177000 | -0.13302900 |
| H | 1.11481900 | -1.70883700 | -0.43825700 |
| H | 1.01621100 | 0.69258400 | -1.22696500 |
| H | -1.01963600 | -0.90312500 | -1.40279400 |
| H | -1.24762200 | -2.09587900 | -0.14043800 |
| H | -2.89610200 | -0.53065400 | 0.34479500 |
| H | -1.63627400 | -0.10016400 | 1.47392100 |
| H | -1.85596600 | 1.43351600 | -1.20185300 |
| H | -2.12984800 | 2.11668400 | 0.49551000 |
| H | 0.62969400 | -1.17239300 | 1.15518500 |
| C | 2.67276700 | 0.41208100 | 0.12648300 |
| H | 3.19649500 | -0.31942600 | -0.49218100 |
| H | 3.12310100 | 1.38835200 | -0.05474700 |
| H | 2.85274500 | 0.14945100 | 1.17105500 |
| H | 0.68511900 | 1.18568800 | 0.41639100 |

**1r-R3**

M06-2X SCF energy: -349.484008 a.u.

M06-2X enthalpy: -349.301320 a.u.

M06-2X Gibbs free energy: -349.350972 a.u.

DLPNO-CCSD(T) SCF energy in solution: -348.862627 a.u.

DLPNO-CCSD(T) enthalpy in solution: -348.679939 a.u.

DLPNO-CCSD(T) Gibbs free energy in solution: -348.729591 a.u.

Cartesian coordinates

| ATOM | X | Y | Z |
|---|---|---|---|
| C | -2.79474900 | 0.33374400 | 0.18905200 |
| C | -1.88242400 | 1.35489300 | -0.04750100 |
| C | -0.54537800 | 1.05778800 | -0.25938500 |

| | | | |
|---|---|---|---|
| C | -0.09512900 | -0.26130300 | -0.24210600 |
| C | -1.01649200 | -1.27434100 | -0.00313100 |
| C | -2.35781700 | -0.98177500 | 0.21118000 |
| H | -3.83883800 | 0.56422200 | 0.35621400 |
| H | -2.21519900 | 2.38488900 | -0.06656400 |
| H | 0.16716800 | 1.85535200 | -0.43874600 |
| H | -0.68028700 | -2.30455900 | 0.01709200 |
| H | -3.06035100 | -1.78404400 | 0.39751000 |
| C | 2.28127400 | 0.15873000 | 0.45005300 |
| C | 1.36403200 | -0.56420300 | -0.47736600 |
| H | 1.62857500 | -0.30717300 | -1.51097000 |
| H | 1.51998200 | -1.65039900 | -0.40203700 |
| C | 3.73928100 | 0.21195900 | 0.16868100 |
| H | 3.93066100 | 0.49961100 | -0.86898100 |
| H | 4.25250900 | 0.91832000 | 0.82041600 |
| H | 4.21723200 | -0.76746700 | 0.31146400 |
| H | 1.92295900 | 0.37829100 | 1.44773700 |

**4n-R3**

M06-2X SCF energy: -236.367597 a.u.

M06-2X enthalpy: -236.181708 a.u.

M06-2X Gibbs free energy: -236.229731 a.u.

DLPNO-CCSD(T) SCF energy in solution: -235.964496 a.u.

DLPNO-CCSD(T) enthalpy in solution: -235.778607 a.u.

DLPNO-CCSD(T) Gibbs free energy in solution: -235.826630 a.u.

Cartesian coordinates

| ATOM | X | Y | Z |
|---|---|---|---|
| C | 0.62441200 | -0.37094100 | -0.01577000 |
| C | 1.84601700 | 0.53741000 | 0.03494100 |

| | | | |
|---|---|---|---|
| C | 3.15518200 | -0.23517700 | -0.06100600 |
| C | -1.89477400 | -0.48753300 | 0.04012300 |
| C | -0.69409200 | 0.38914200 | 0.08105200 |
| H | 4.01839200 | 0.42978900 | -0.02347000 |
| H | 1.78675400 | 1.26502400 | -0.77989700 |
| H | 1.82574500 | 1.11536300 | 0.96368100 |
| H | 0.63730600 | -0.94978900 | -0.94473100 |
| H | 0.67967500 | -1.09897200 | 0.80045800 |
| H | -1.79130700 | -1.51615600 | 0.36392400 |
| H | -0.75345300 | 1.12789300 | -0.72878500 |
| H | -0.69310400 | 0.98568800 | 1.00828000 |
| H | 3.24604000 | -0.94886500 | 0.76014200 |
| C | -3.25427200 | 0.09814400 | -0.08985800 |
| H | -4.01069800 | -0.66252800 | -0.28036200 |
| H | -3.55440400 | 0.63491100 | 0.82123100 |
| H | -3.29212500 | 0.82966900 | -0.90249800 |
| H | 3.20634400 | -0.79829200 | -0.99486900 |

**4q-R2**

M06-2X SCF energy: -156.519864 a.u.

M06-2X enthalpy: -156.418542 a.u.

M06-2X Gibbs free energy: -156.457497 a.u.

DLPNO-CCSD(T) SCF energy in solution: -156.248386 a.u.

DLPNO-CCSD(T) enthalpy in solution: -156.147064 a.u.

DLPNO-CCSD(T) Gibbs free energy in solution: -156.186019 a.u.

Cartesian coordinates

| ATOM | X | Y | Z |
|---|---|---|---|
| C | -1.78158200 | 0.05102600 | 0.26574600 |
| H | -1.85157700 | -0.63406500 | 1.10337000 |

| | | | |
|---|---|---|---|
| C | -0.65743100 | 0.18306200 | -0.42111900 |
| C | 1.73873500 | 0.36135900 | 0.21515500 |
| C | 0.61623300 | -0.55643900 | -0.12641900 |
| H | -0.61525500 | 0.88331600 | -1.25100000 |
| H | 2.76363700 | 0.08341200 | 0.01967200 |
| H | 1.55116100 | 1.24429800 | 0.80906200 |
| H | 0.89527800 | -1.16535800 | -0.99199400 |
| H | 0.42987900 | -1.26372200 | 0.69422500 |
| H | -2.66885500 | 0.61807100 | 0.01648700 |

**4r-R2**

M06-2X SCF energy: -235.160425 a.u.

M06-2X enthalpy: -234.997784 a.u.

M06-2X Gibbs free energy: -235.039202 a.u.

DLPNO-CCSD(T) SCF energy in solution: -234.756006 a.u.

DLPNO-CCSD(T) enthalpy in solution: -234.593365 a.u.

DLPNO-CCSD(T) Gibbs free energy in solution: -234.634783 a.u.

Cartesian coordinates

| ATOM | X | Y | Z |
|---|---|---|---|
| C | 0.81638200 | -0.02670200 | -0.36526600 |
| C | -0.02004900 | -1.18476400 | 0.19452900 |
| C | -1.46120500 | -0.76501900 | -0.10173500 |
| C | -1.46963700 | 0.77653700 | 0.04011900 |
| C | 0.01151300 | 1.19990000 | 0.12769500 |
| H | 0.75216500 | -0.05474000 | -1.45846000 |
| H | 0.14850500 | -1.24524200 | 1.27463400 |
| H | 0.24821600 | -2.14878800 | -0.23795300 |
| H | -1.71708100 | -1.04751100 | -1.12466900 |
| H | -2.18372500 | -1.24972600 | 0.55415000 |

| | | | |
|---|---|---|---|
| H | -1.96213400 | 1.23963800 | -0.81480400 |
| H | -2.01657000 | 1.09317300 | 0.92793400 |
| H | 0.23958800 | 2.09975200 | -0.44288400 |
| H | 0.29042300 | 1.39472500 | 1.16623400 |
| C | 2.23725800 | -0.02788700 | 0.05092700 |
| H | 3.02419500 | 0.31008000 | -0.60663400 |
| H | 2.49084500 | -0.22375600 | 1.08484600 |

**TS1**

M06-2X SCF energy: -156.501546 a.u.

M06-2X enthalpy: -156.401321 a.u.

M06-2X Gibbs free energy: -156.437200 a.u.

DLPNO-CCSD(T) SCF energy in solution: -156.231981 a.u.

DLPNO-CCSD(T) enthalpy in solution: -156.131756 a.u.

DLPNO-CCSD(T) Gibbs free energy in solution: -156.167635 a.u.

Imaginary frequency: -712.0836 $cm^{-1}$

Cartesian coordinates

| ATOM | X | Y | Z |
|---|---|---|---|
| C | -1.58965400 | -0.12801300 | -0.18398200 |
| H | -1.67855100 | 0.05180200 | -1.24790500 |
| C | -0.43240200 | 0.16004700 | 0.48501500 |
| C | 1.19610000 | -0.69247300 | -0.10483600 |
| C | 0.79067600 | 0.72535100 | -0.15429600 |
| H | -0.39640200 | 0.03971100 | 1.56002200 |
| H | 0.98418700 | -1.34470000 | -0.93615200 |
| H | 1.74604800 | -1.08462500 | 0.73626200 |
| H | 1.36524300 | 1.40677000 | 0.46483800 |
| H | 0.61912900 | 1.12795700 | -1.14821100 |
| H | -2.42797900 | -0.58639100 | 0.31974400 |

**TS2**

M06-2X SCF energy: -235.120487 a.u.

M06-2X enthalpy: -235.959611 a.u.

M06-2X Gibbs free energy: -235.001264 a.u.

DLPNO-CCSD(T) SCF energy in solution: -234.717308 a.u.

DLPNO-CCSD(T) enthalpy in solution: -234.556432 a.u.

DLPNO-CCSD(T) Gibbs free energy in solution: -234.598085 a.u.

Imaginary frequency: -619.3958 $cm^{-1}$

Cartesian coordinates

| ATOM | X | Y | Z |
|---|---|---|---|
| C | 1.07781000 | -0.16025600 | 0.41752600 |
| C | -0.44472900 | 1.43944200 | 0.21108000 |
| C | -1.48982700 | 0.56349600 | -0.39988900 |
| C | -1.31566000 | -0.84694800 | 0.16427300 |
| C | 0.13973600 | -1.23743100 | -0.07037400 |
| H | 1.07866600 | 0.00915700 | 1.49049600 |
| H | 0.03041100 | 2.20311300 | -0.38978500 |
| H | -0.56533100 | 1.69800200 | 1.25772000 |
| H | -2.49985900 | 0.93806500 | -0.19990400 |
| H | -1.36727200 | 0.53478500 | -1.48541400 |
| H | -1.52352100 | -0.83342300 | 1.23877900 |
| H | -2.00369500 | -1.56042000 | -0.29214500 |
| H | 0.37572300 | -2.17679100 | 0.43613700 |
| H | 0.30992400 | -1.39618300 | -1.13897500 |
| C | 2.18575500 | 0.20860300 | -0.28060900 |
| H | 2.93995200 | 0.85183400 | 0.15166100 |
| H | 2.30649400 | -0.06956600 | -1.32061400 |

**TS3**

M06-2X SCF energy: -349.455569 a.u.

M06-2X enthalpy: -349.274435 a.u.

M06-2X Gibbs free energy: -349.320992 a.u.

DLPNO-CCSD(T) SCF energy in solution: -348.837569 a.u.

DLPNO-CCSD(T) enthalpy in solution: -348.656435 a.u.

DLPNO-CCSD(T) Gibbs free energy in solution: -348.702992 a.u.

Imaginary frequency: -609.4559 $cm^{-1}$

Cartesian coordinates

| ATOM | X | Y | Z |
|---|---|---|---|
| C | 2.51640300 | 0.29798000 | -0.00877400 |
| C | 1.61015100 | 1.34312200 | 0.18316500 |
| C | 0.25283700 | 1.12646000 | 0.13540000 |
| C | -0.26916100 | -0.18632700 | -0.07403300 |
| C | 0.67731900 | -1.22426800 | -0.33851600 |
| C | 2.02787800 | -0.98227500 | -0.28635400 |
| H | 3.58096900 | 0.48205800 | 0.02848300 |
| H | 1.97863700 | 2.34645700 | 0.35731800 |
| H | -0.42773100 | 1.95681300 | 0.26888500 |
| H | 0.30711000 | -2.21682800 | -0.56862200 |
| H | 2.72047900 | -1.79206100 | -0.47920400 |
| C | -1.73972800 | -0.45915800 | -0.22206200 |
| C | -1.50752400 | -0.69647600 | 1.21407900 |
| H | -1.67025900 | 0.10521000 | 1.91968100 |
| H | -1.28710600 | -1.68206300 | 1.59267900 |
| C | -2.66316700 | 0.66661700 | -0.62678300 |
| H | -2.45584200 | 0.99638100 | -1.64543100 |
| H | -3.69960000 | 0.33222500 | -0.57286900 |
| H | -2.56163700 | 1.52407100 | 0.03808700 |

| | | | |
|---|---|---|---|
| H | -1.91506400 | -1.36631400 | -0.79573900 |

**TS4**

| | |
|---|---|
| M06-2X SCF energy: | -349.457288 a.u. |
| M06-2X enthalpy: | -349.275944 a.u. |
| M06-2X Gibbs free energy: | -349.323097 a.u. |
| DLPNO-CCSD(T) SCF energy in solution: | -348.838324 a.u. |
| DLPNO-CCSD(T) enthalpy in solution: | -348.656980 a.u. |
| DLPNO-CCSD(T) Gibbs free energy in solution: | -348.704133 a.u. |

Imaginary frequency: -592.5127 $cm^{-1}$

Cartesian coordinates

| ATOM | X | Y | Z |
|---|---|---|---|
| C | 2.39268300 | 0.41038600 | -0.32491900 |
| C | 1.53016000 | 1.33106500 | 0.27580100 |
| C | 0.26859700 | 0.96340400 | 0.68080100 |
| C | -0.21206800 | -0.36289300 | 0.45416300 |
| C | 0.71646600 | -1.30699900 | -0.08547800 |
| C | 1.97202900 | -0.91516000 | -0.47909500 |
| H | 3.38348500 | 0.71036200 | -0.63601400 |
| H | 1.86558300 | 2.34653400 | 0.44699400 |
| H | -0.37403100 | 1.68096600 | 1.17567400 |
| H | 0.40461400 | -2.33949100 | -0.19451900 |
| H | 2.64938100 | -1.64805900 | -0.89951400 |
| C | -1.82799700 | -0.41754600 | -0.44372400 |
| C | -1.55349500 | -0.80684800 | 0.95069700 |
| H | -1.98780900 | -0.16874100 | 1.71482500 |
| H | -1.62558900 | -1.86144600 | 1.19666200 |
| C | -2.42044500 | 0.89568300 | -0.82901600 |
| H | -1.83267900 | 1.39506500 | -1.60313900 |

| | | | |
|---|---|---|---|
| H | -3.43190700 | 0.75788600 | -1.22263500 |
| H | -2.48894000 | 1.56800800 | 0.02579300 |
| H | -1.75768000 | -1.18763500 | -1.19950500 |

**TS5**

M06-2X SCF energy: -236.333165 a.u.

M06-2X enthalpy: -236.152315 a.u.

M06-2X Gibbs free energy: -236.196192 a.u.

DLPNO-CCSD(T) SCF energy in solution: -235.931052 a.u.

DLPNO-CCSD(T) enthalpy in solution: -235.750202 a.u.

DLPNO-CCSD(T) Gibbs free energy in solution: -235.794079 a.u.

Imaginary frequency: -1578.0134 $cm^{-1}$

Cartesian coordinates

| ATOM | X | Y | Z |
|---|---|---|---|
| C | 1.10645400 | 0.02487700 | -0.44825300 |
| C | 0.34144700 | -1.14744300 | 0.12421800 |
| C | -1.14961600 | -1.05963100 | -0.20929600 |
| C | -1.76095800 | 0.21366200 | 0.38801900 |
| C | -1.08208000 | 1.45049300 | -0.15082000 |
| H | 0.75481400 | -2.09492600 | -0.23961000 |
| H | 1.19483300 | -0.02418200 | -1.53483800 |
| H | -1.27893200 | -1.04908800 | -1.29690200 |
| H | -1.67457500 | -1.93980900 | 0.16609300 |
| H | -2.83664100 | 0.23490800 | 0.18419700 |
| H | -1.65128500 | 0.17497900 | 1.47544000 |
| H | -1.34941200 | 1.71040800 | -1.17334600 |
| H | -1.04996300 | 2.31720900 | 0.50328600 |
| H | 0.45874900 | -1.15621700 | 1.21363600 |
| H | 0.20276300 | 0.99021200 | -0.29366300 |

| | | | |
|---|---|---|---|
| C | 2.38738800 | 0.40690200 | 0.24570700 |
| H | 3.11774700 | -0.40934700 | 0.21021100 |
| H | 2.85001900 | 1.28199800 | -0.21135800 |
| H | 2.20607200 | 0.63069500 | 1.29940500 |

**XII. $^{1}$H, $^{11}$B, $^{13}$C and $^{19}$F NMR spectra of compounds**

$^{1}$H NMR (400 MHz, $CDCl_3$) spectrum of 2-(2-(furan-2-yl)ethyl)-4,4,5,5-tetramethyl-1,3,2-dioxaborolane **(1m)**

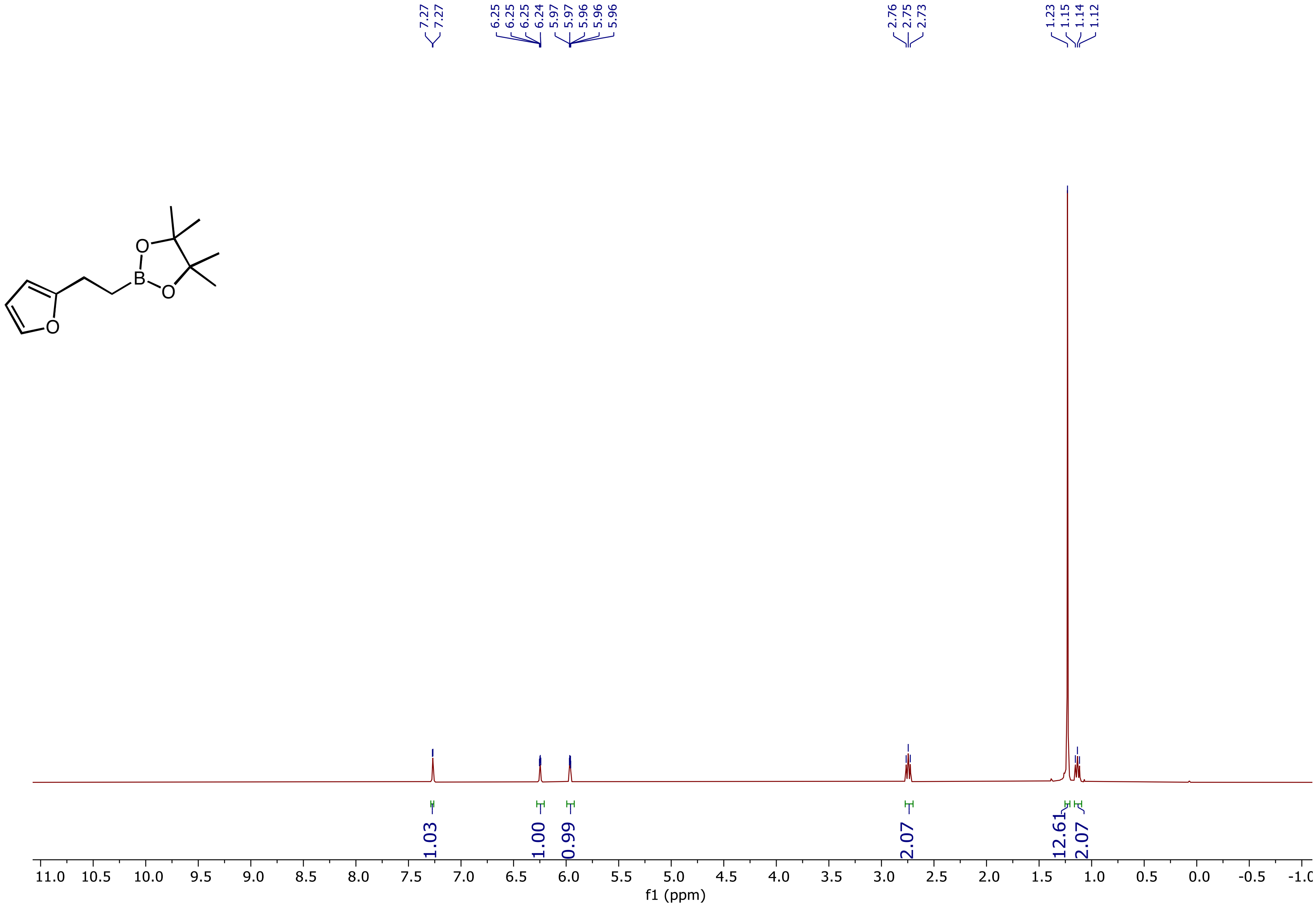

$^{13}C$ NMR (101 MHz, $CDCl_3$) spectrum of 2-(2-(furan-2-yl)ethyl)-4,4,5,5-tetramethyl-1,3,2-dioxaborolane **(1m)**

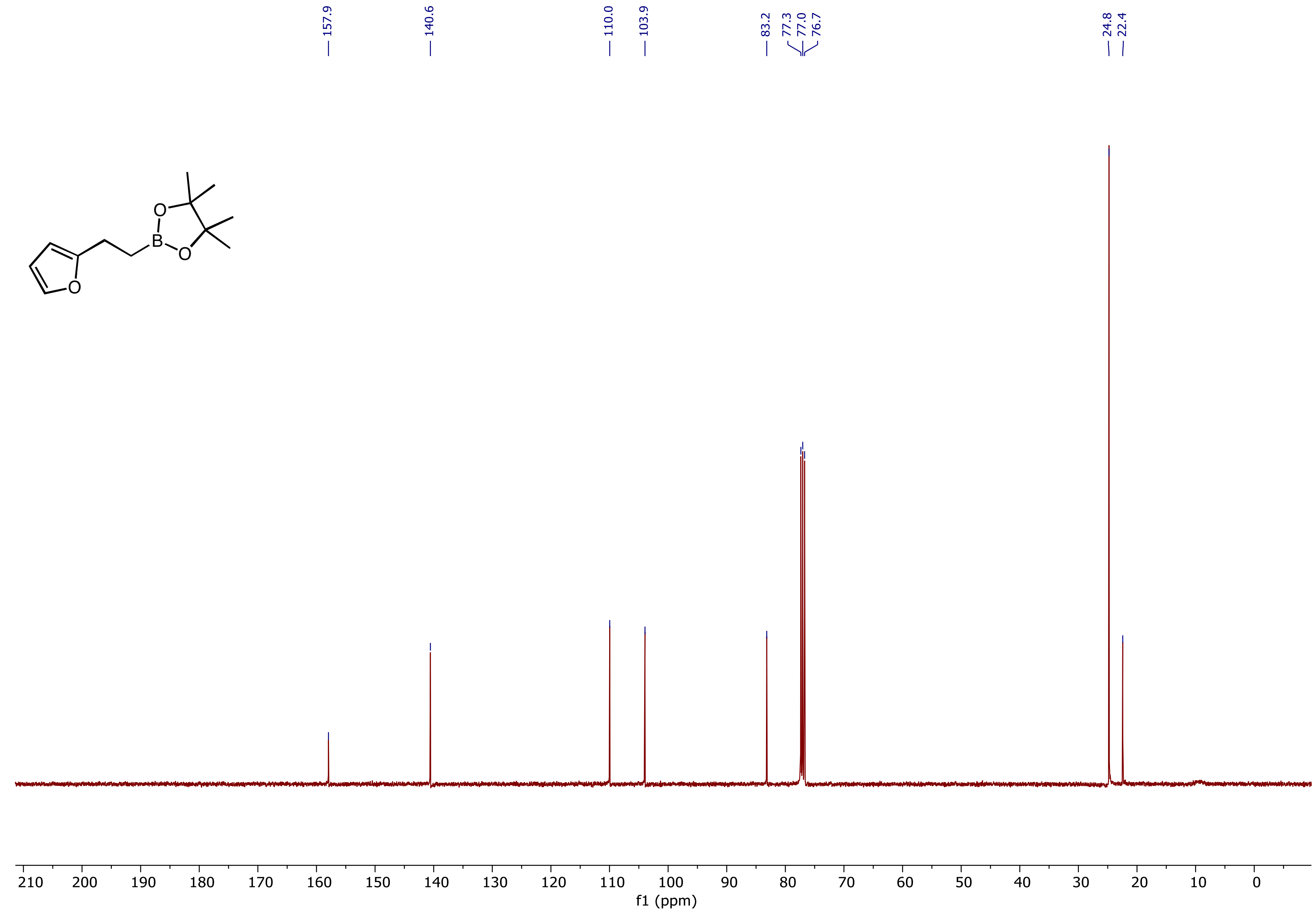

$^{11}$B NMR (160 MHz, $CDCl_3$) spectrum of 2-(2-(furan-2-yl)ethyl)-4,4,5,5-tetramethyl-1,3,2-dioxaborolane **(1m)**

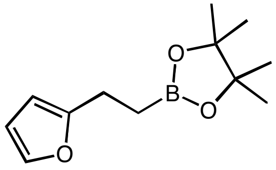

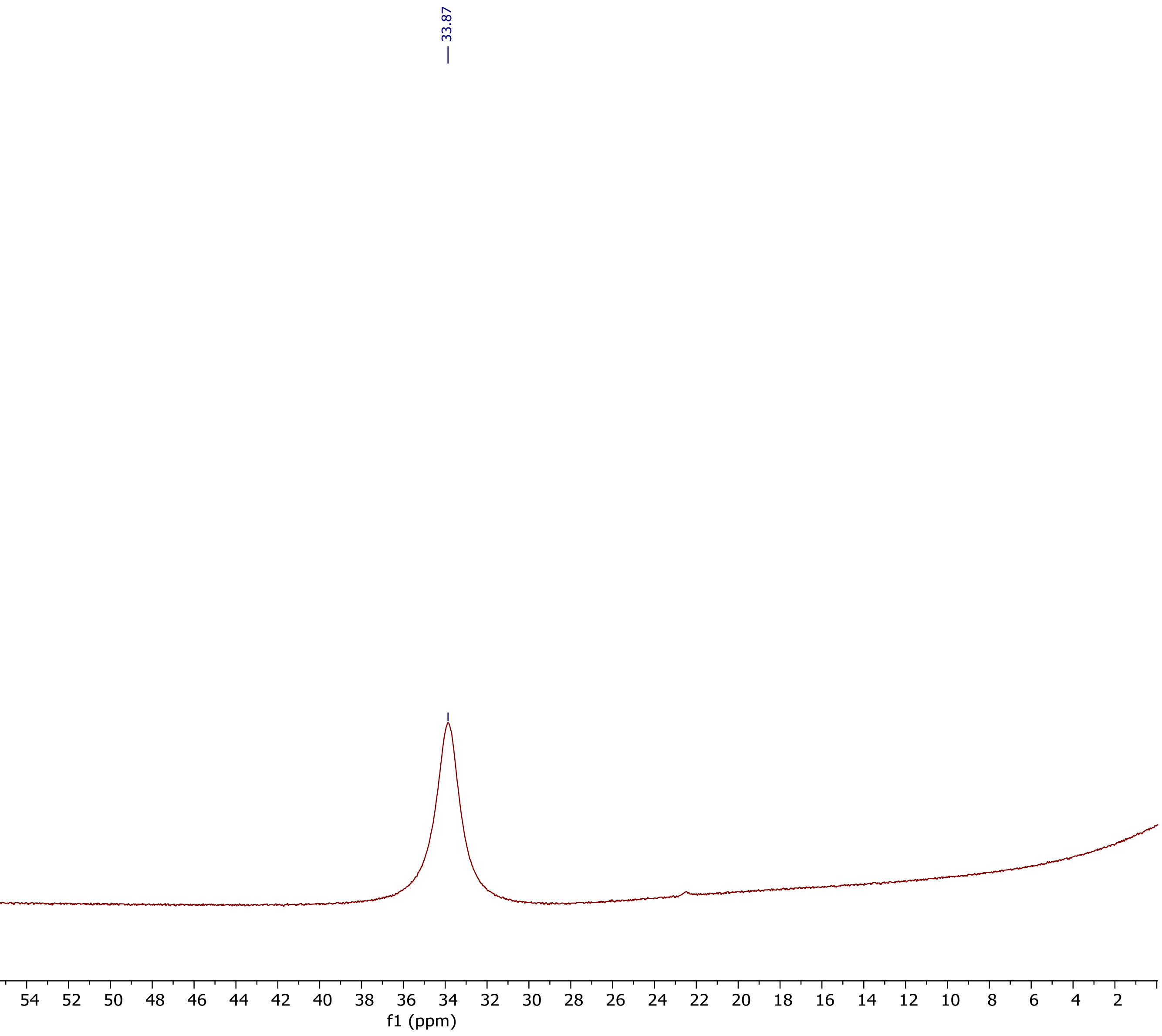

$^1$H NMR (400 MHz, $CDCl_3$) spectrum of 2-Methoxy-5-(2-(4,4,5,5-tetramethyl-1,3,2-dioxaborolan-2-yl)ethyl)pyridine **(1o)**

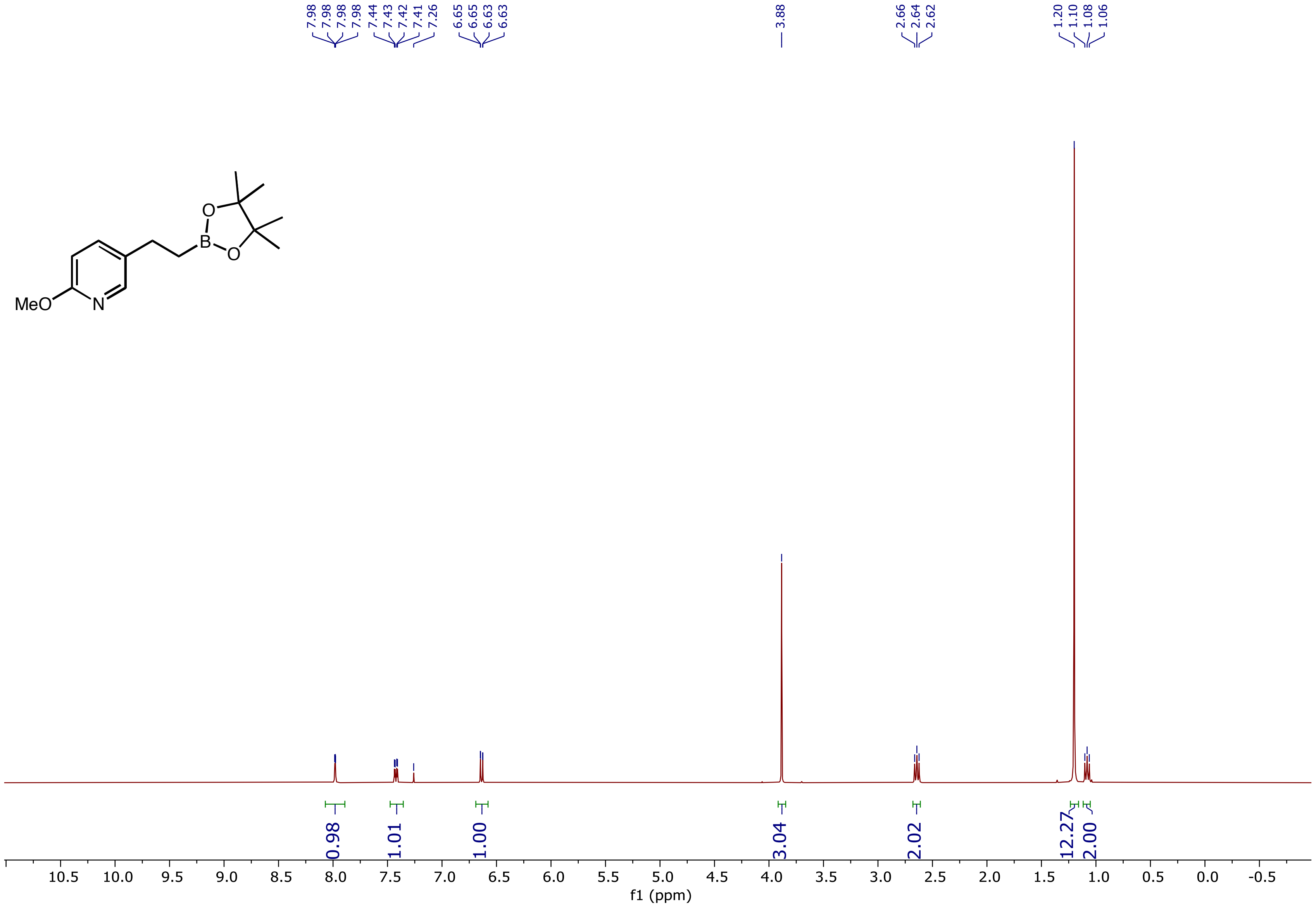

$^{13}C$ NMR (101 MHz, $CDCl_3$) spectrum of 2-Methoxy-5-(2-(4,4,5,5-tetramethyl-1,3,2-dioxaborolan-2-yl)ethyl)pyridine **(1o)**

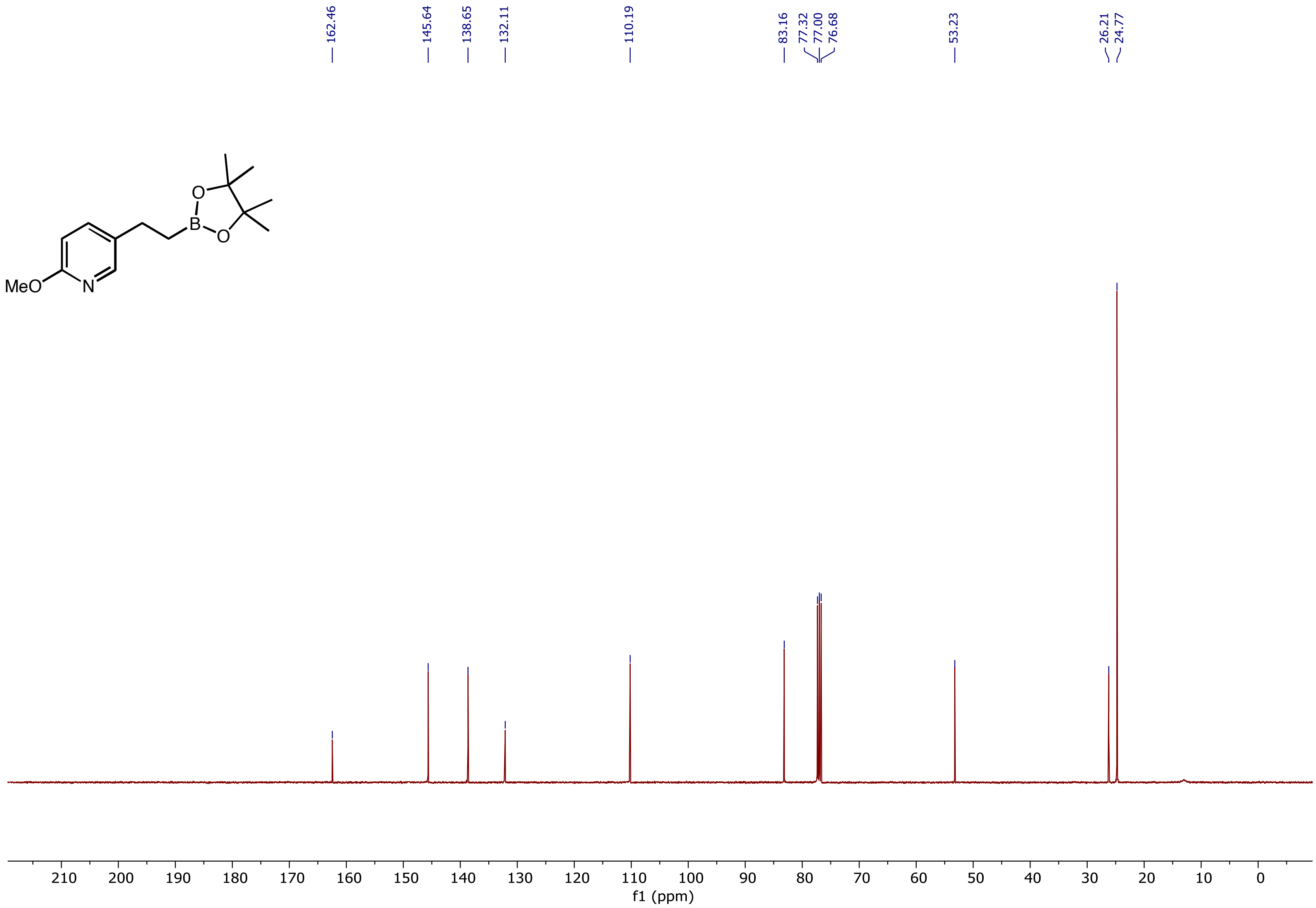

$^{11}B$ NMR (160 MHz, $CDCl_3$) spectrum of 2-Methoxy-5-(2-(4,4,5,5-tetramethyl-1,3,2-dioxaborolan-2-yl)ethyl)pyridine **(1o)**

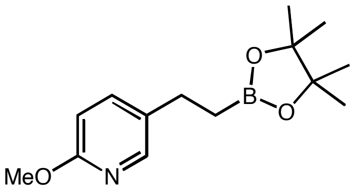

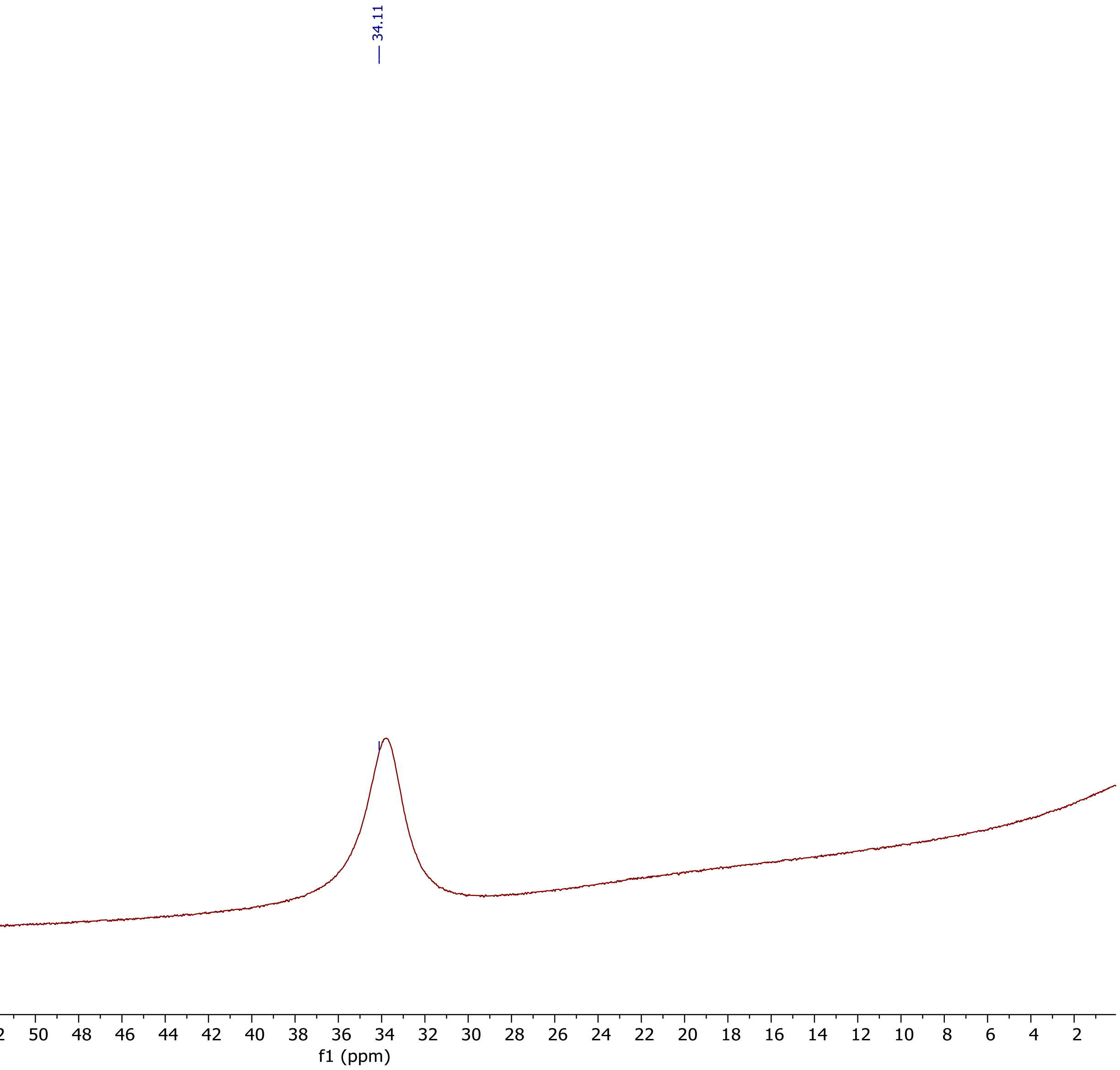

[1]H NMR (500 MHz, $CDCl_3$) spectrum of 2-(4-Fluorobutyl)-4,4,5,5-tetramethyl-1,3,2-dioxaborolane **(4b)**

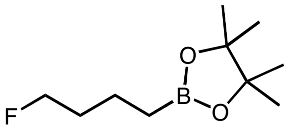

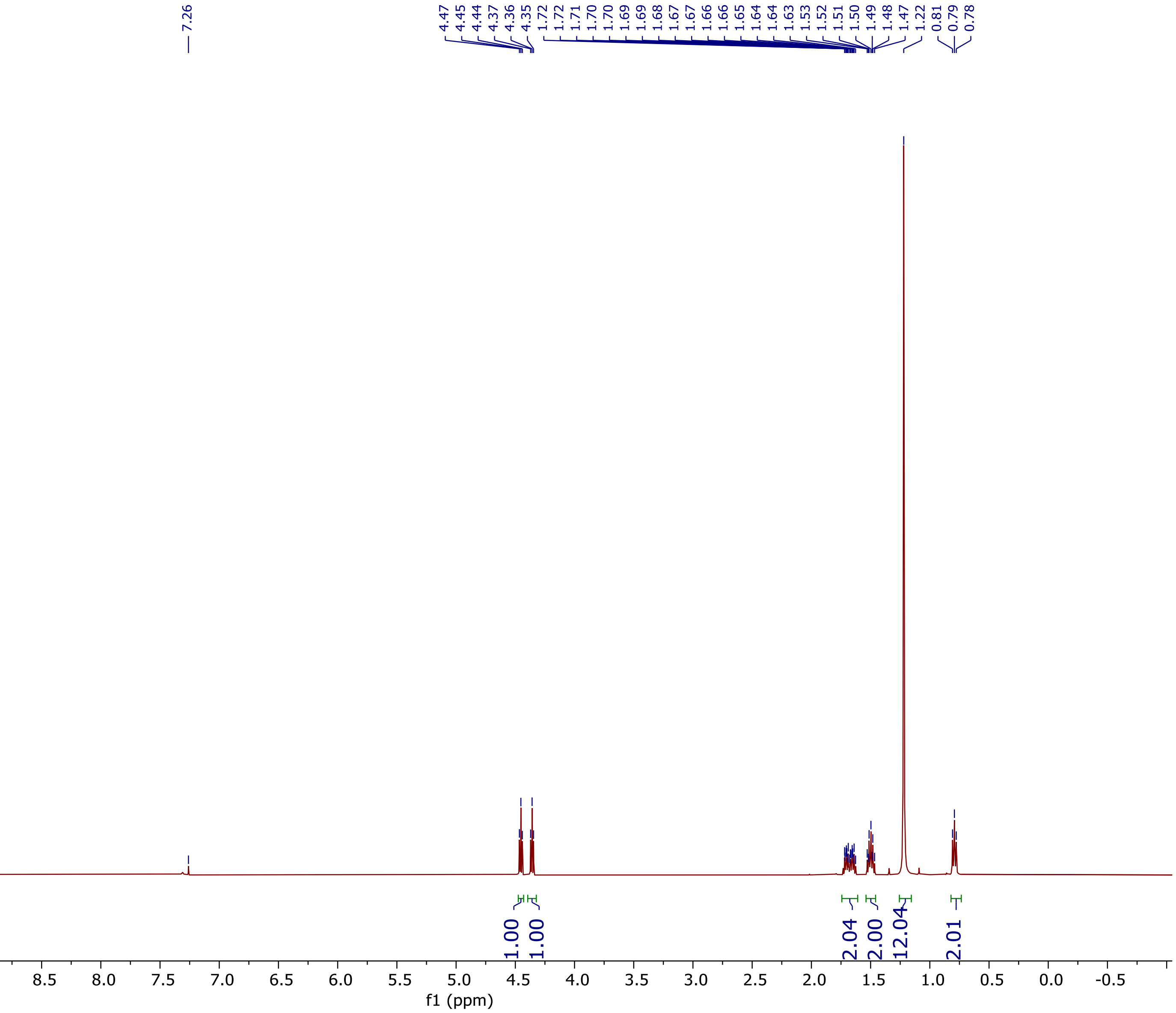

$^{13}C$ NMR (126 MHz, $CDCl_3$) spectrum of 2-(4-Fluorobutyl)-4,4,5,5-tetramethyl-1,3,2-dioxaborolane **(4b)**

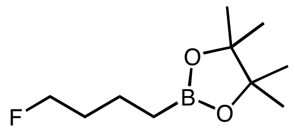

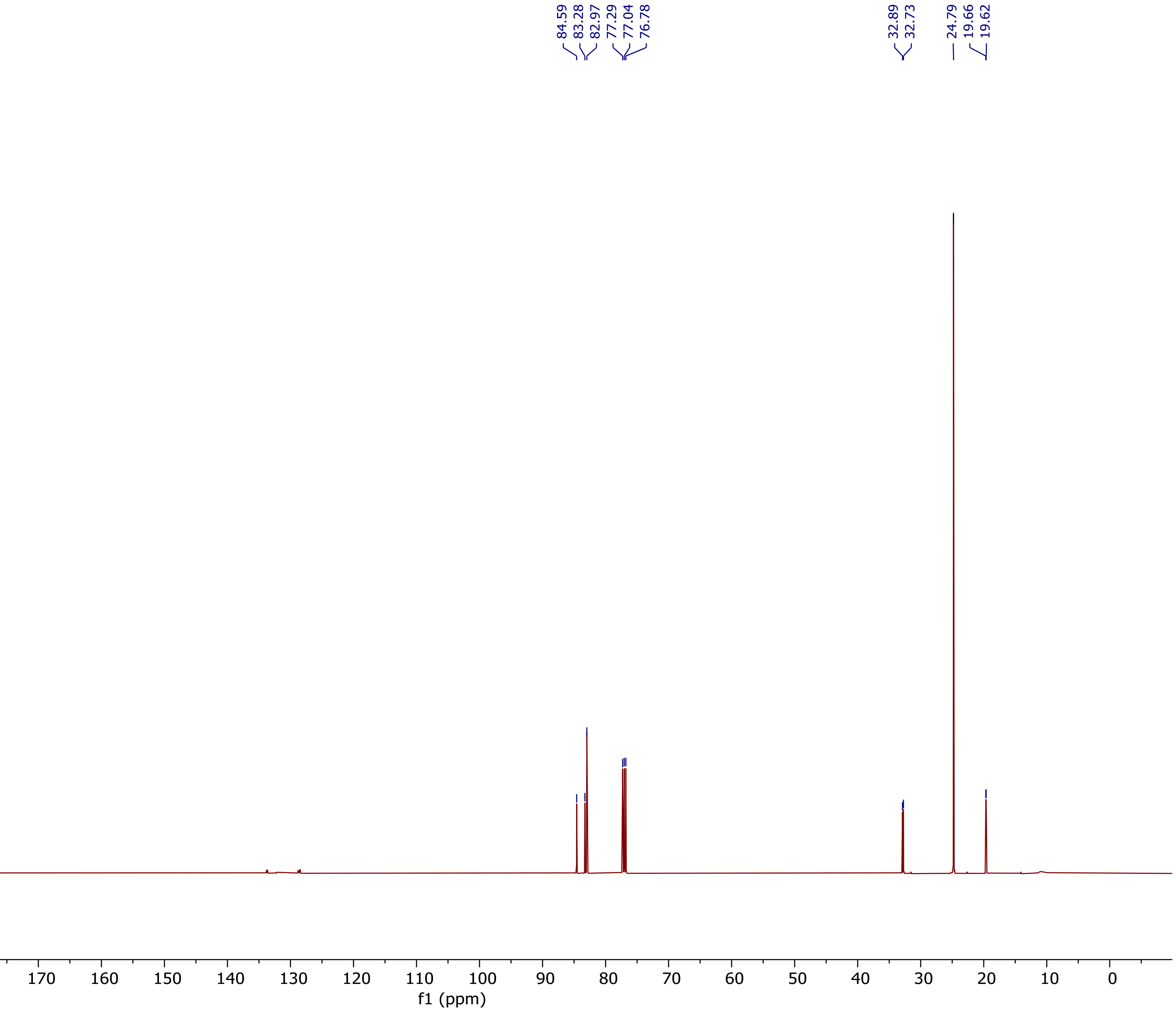

[11]B NMR (160 MHz, $CDCl_3$) spectrum of 2-(4-Fluorobutyl)-4,4,5,5-tetramethyl-1,3,2-dioxaborolane **(4b)**

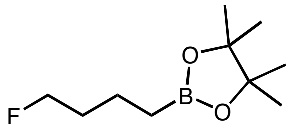

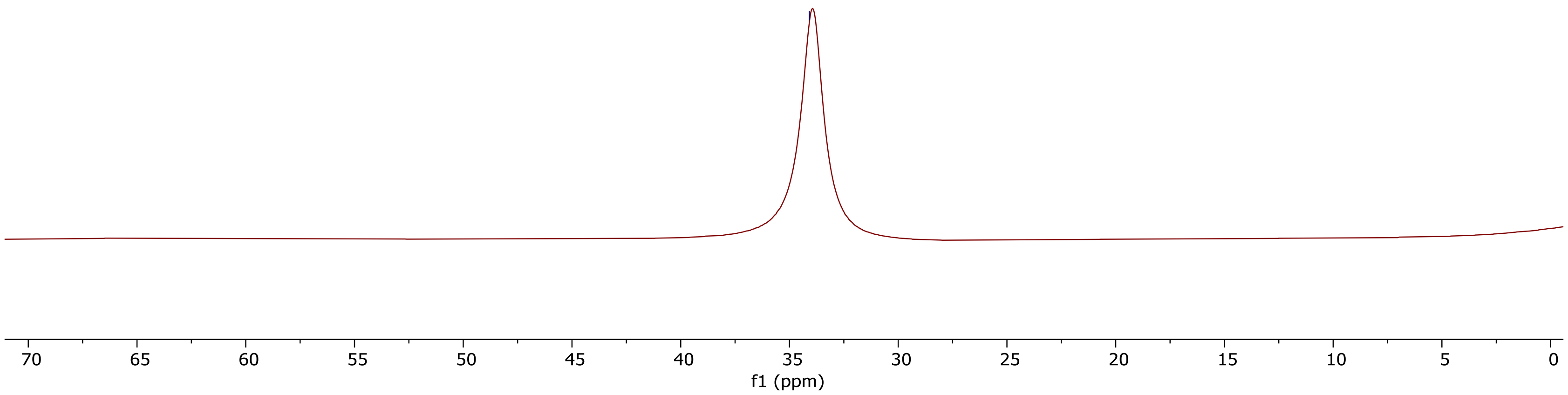

$^{19}F$ NMR (376 MHz, $CDCl_3$) spectrum of 2-(4-Fluorobutyl)-4,4,5,5-tetramethyl-1,3,2-dioxaborolane **(4b)**

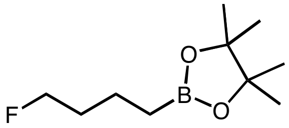

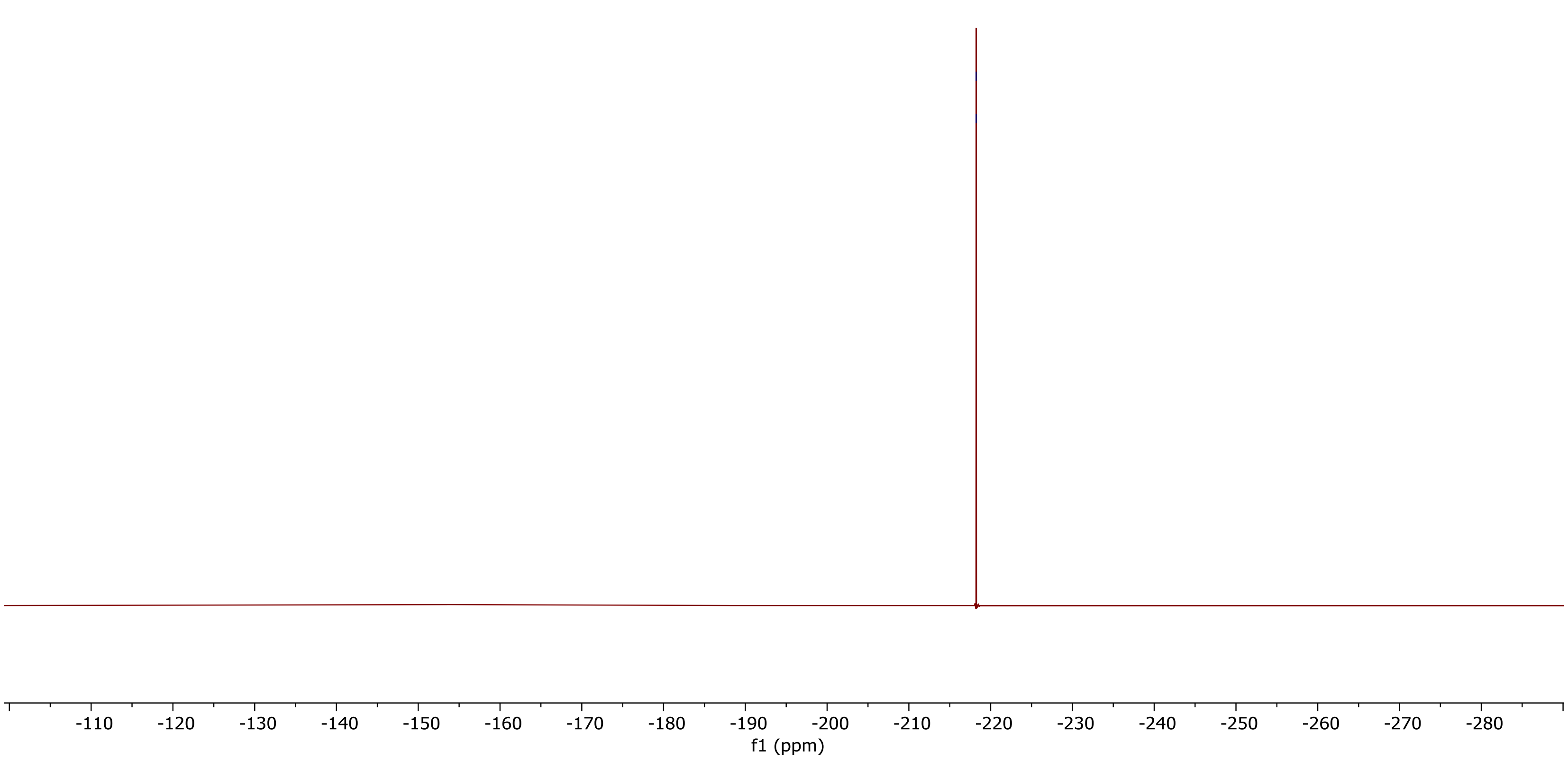

[1]H NMR (400 MHz, $CDCl_3$) spectrum of 2-(5-Azidopentyl)-4,4,5,5-tetramethyl-1,3,2-dioxaborolane **(4f)**

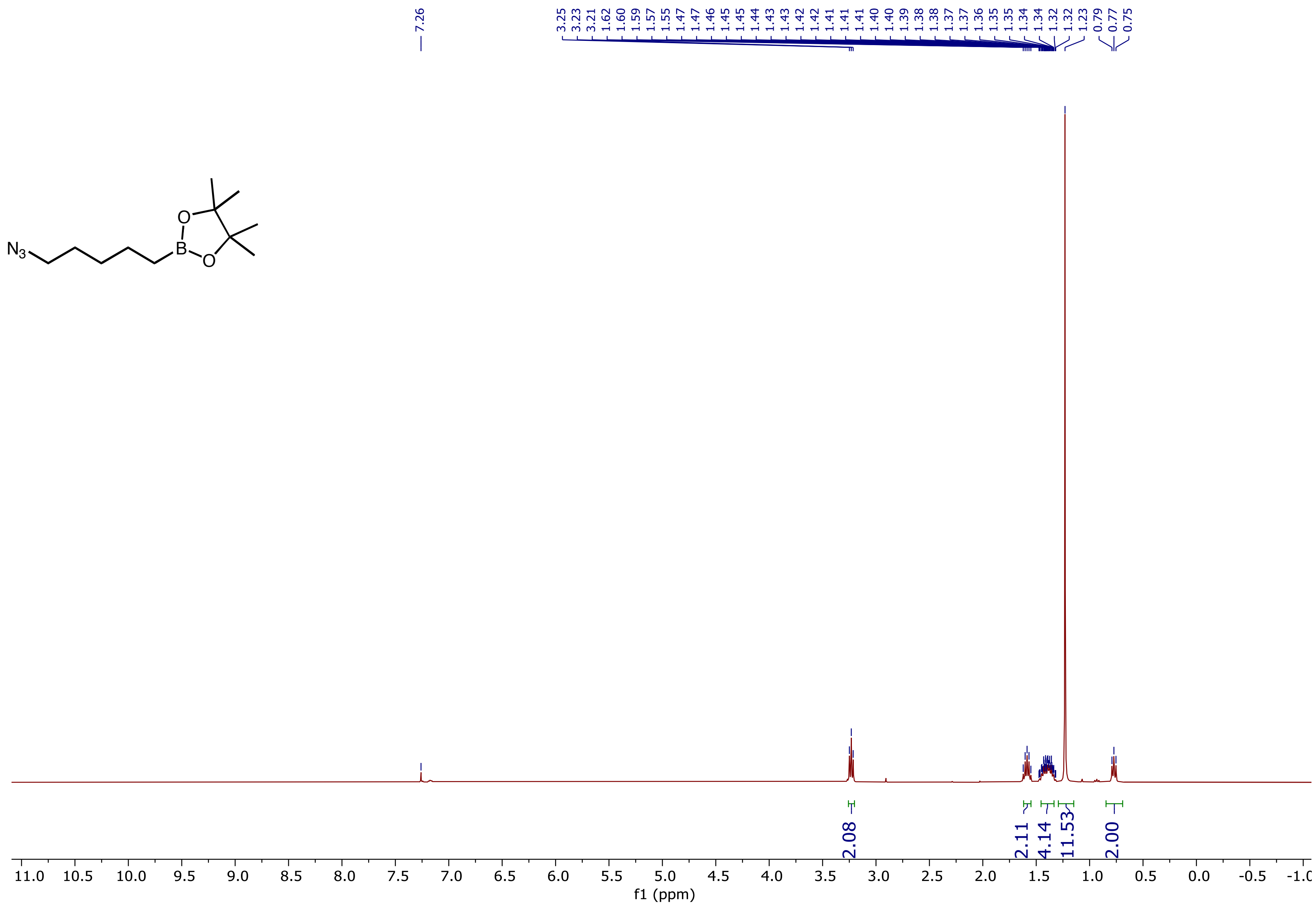

[13]C NMR (126 MHz, $CDCl_3$) spectrum of 2-(5-Azidopentyl)-4,4,5,5-tetramethyl-1,3,2-dioxaborolane **(4f)**

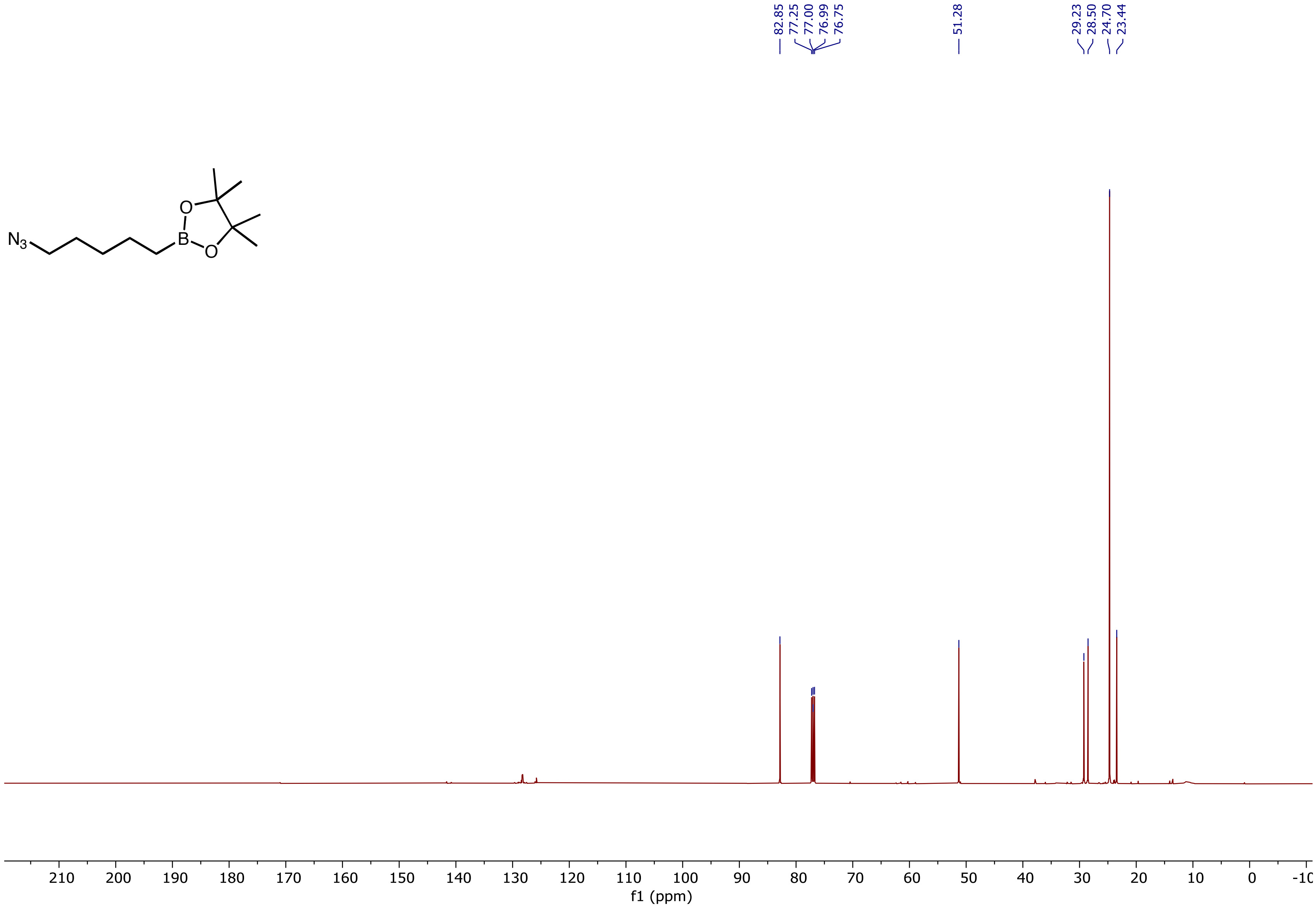

[11]B NMR (128 MHz, $CDCl_3$) spectrum of 2-(5-Azidopentyl)-4,4,5,5-tetramethyl-1,3,2-dioxaborolane **(4f)**

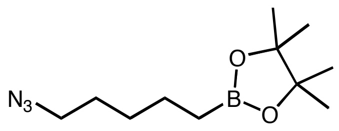

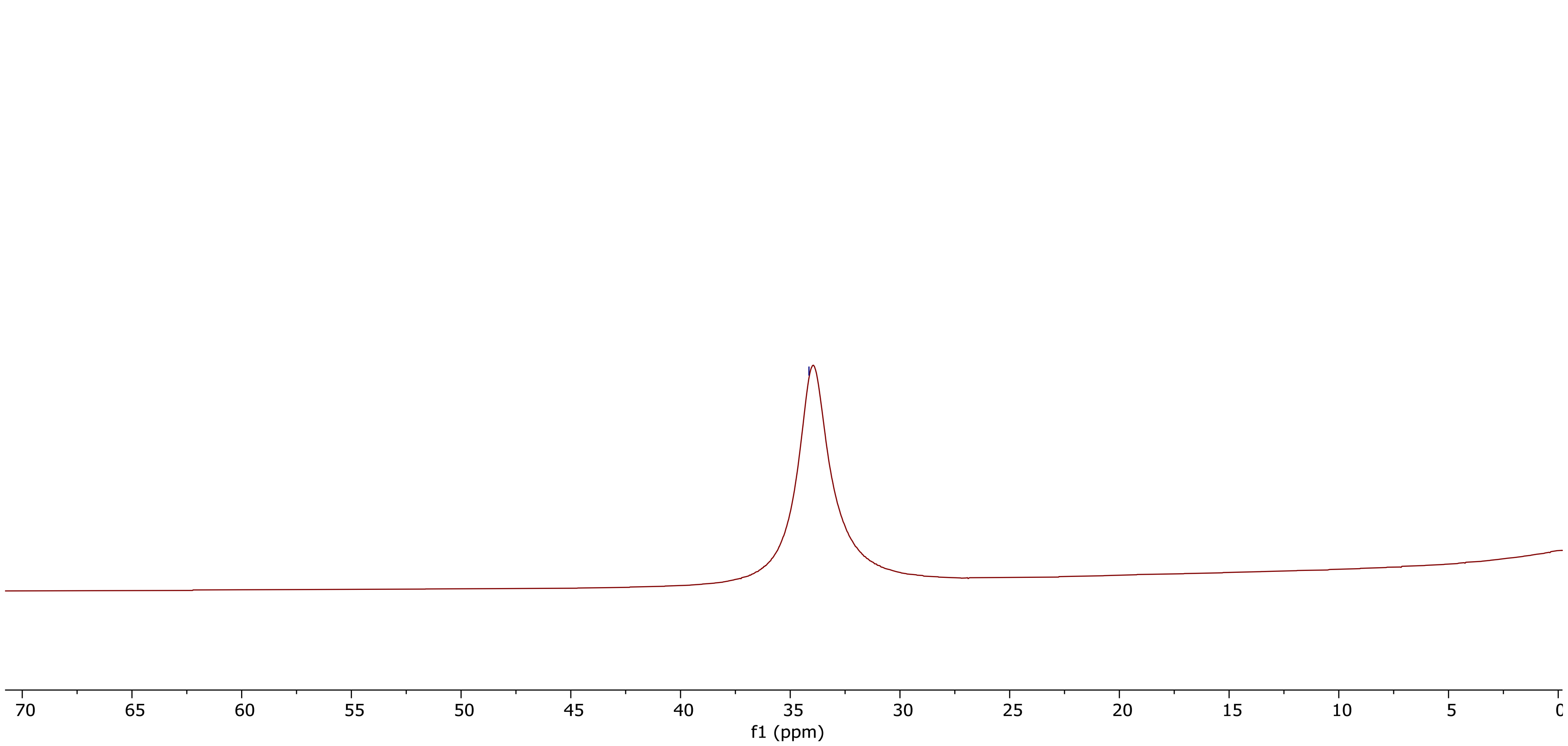

$^{1}H$ NMR (400 MHz, $D_2O$) spectrum of 2-Amino-5-phenylpentanoic acid **(3a)**

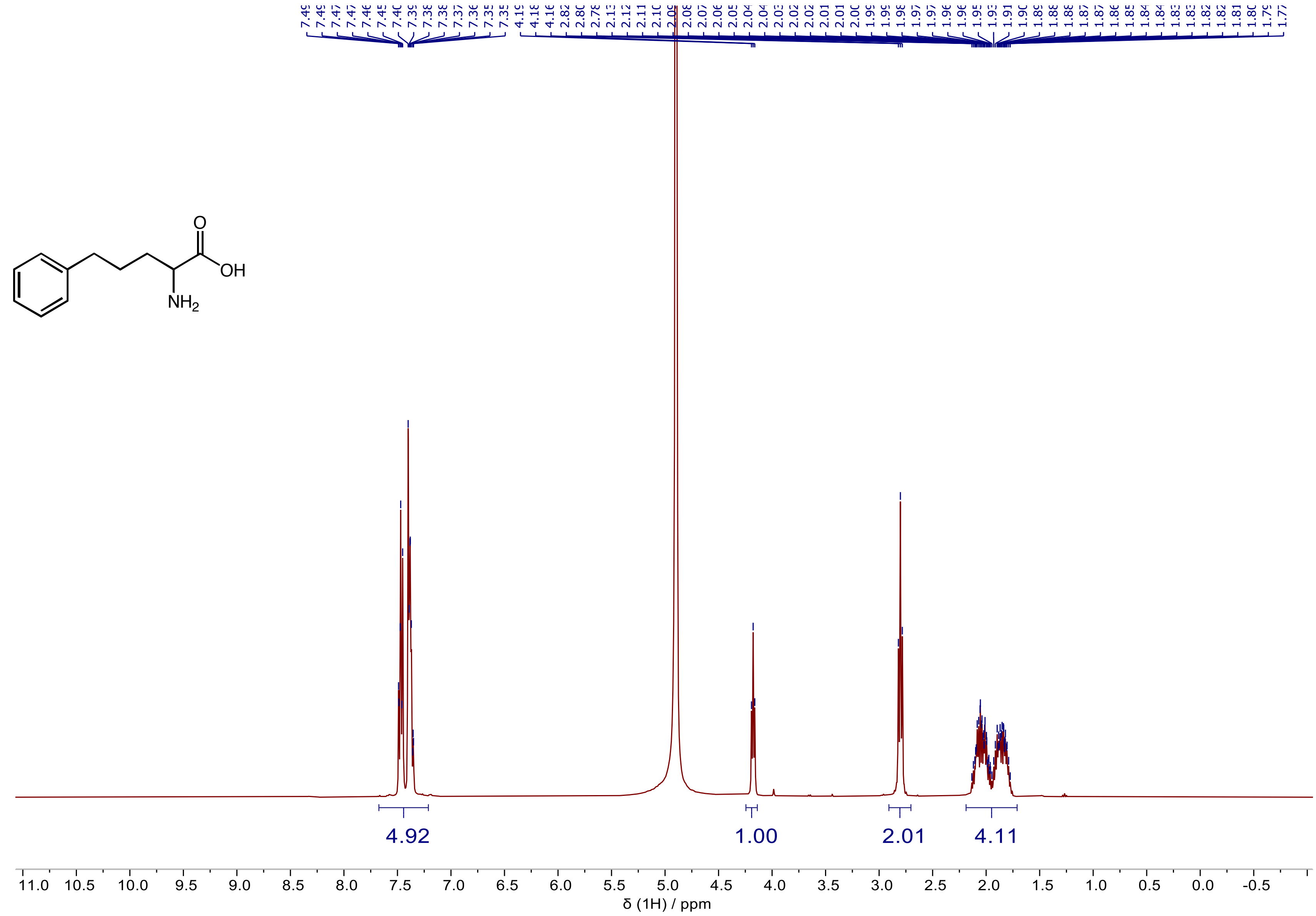

$^{13}C$ NMR (101 MHz, $D_2O$) spectrum of 2-Amino-5-phenylpentanoic acid **(3a)**

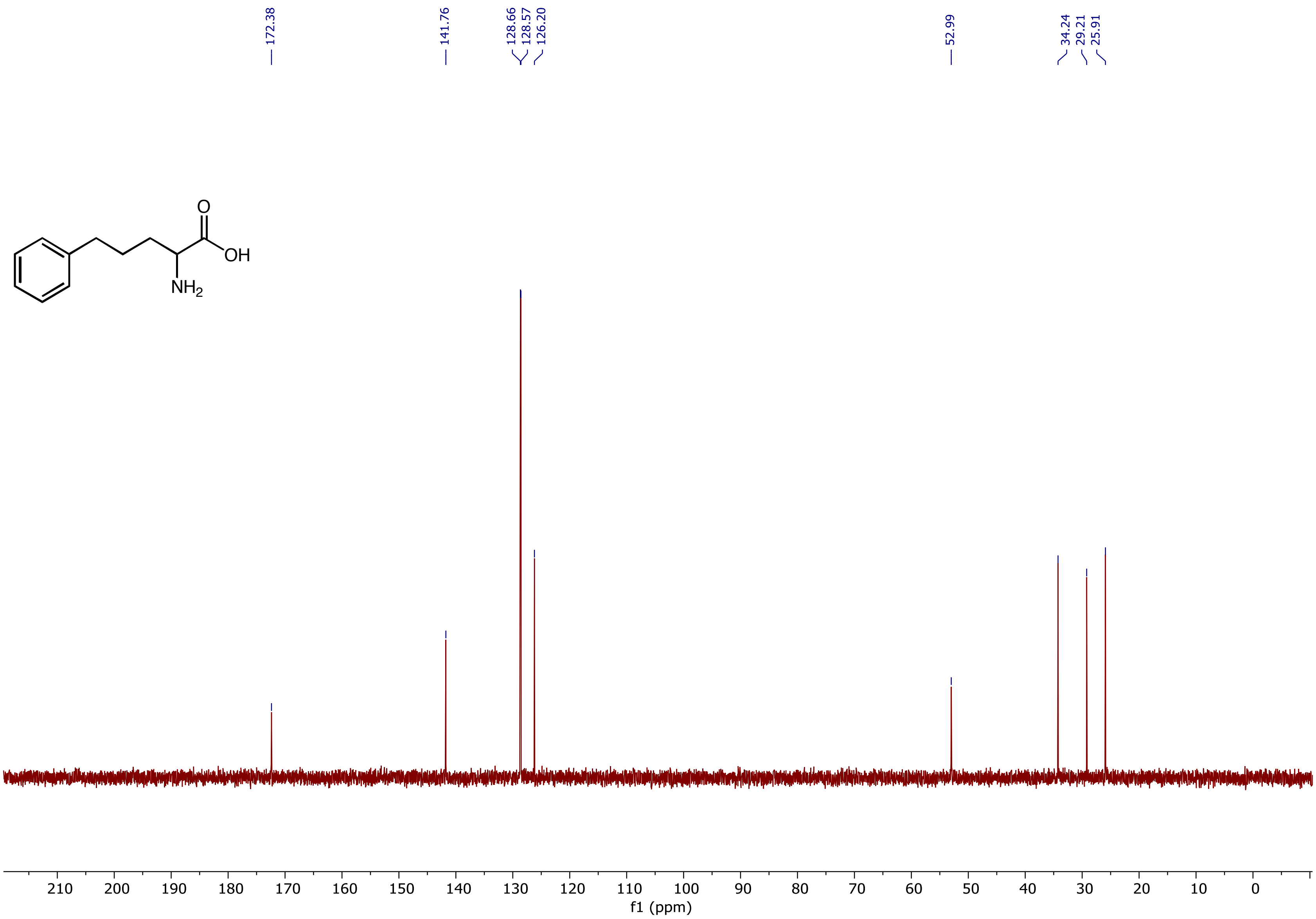

[1]H NMR (500 MHz, $D_2O$) spectrum of 2-Amino-5-(p-tolyl)pentanoic acid **(3b)**

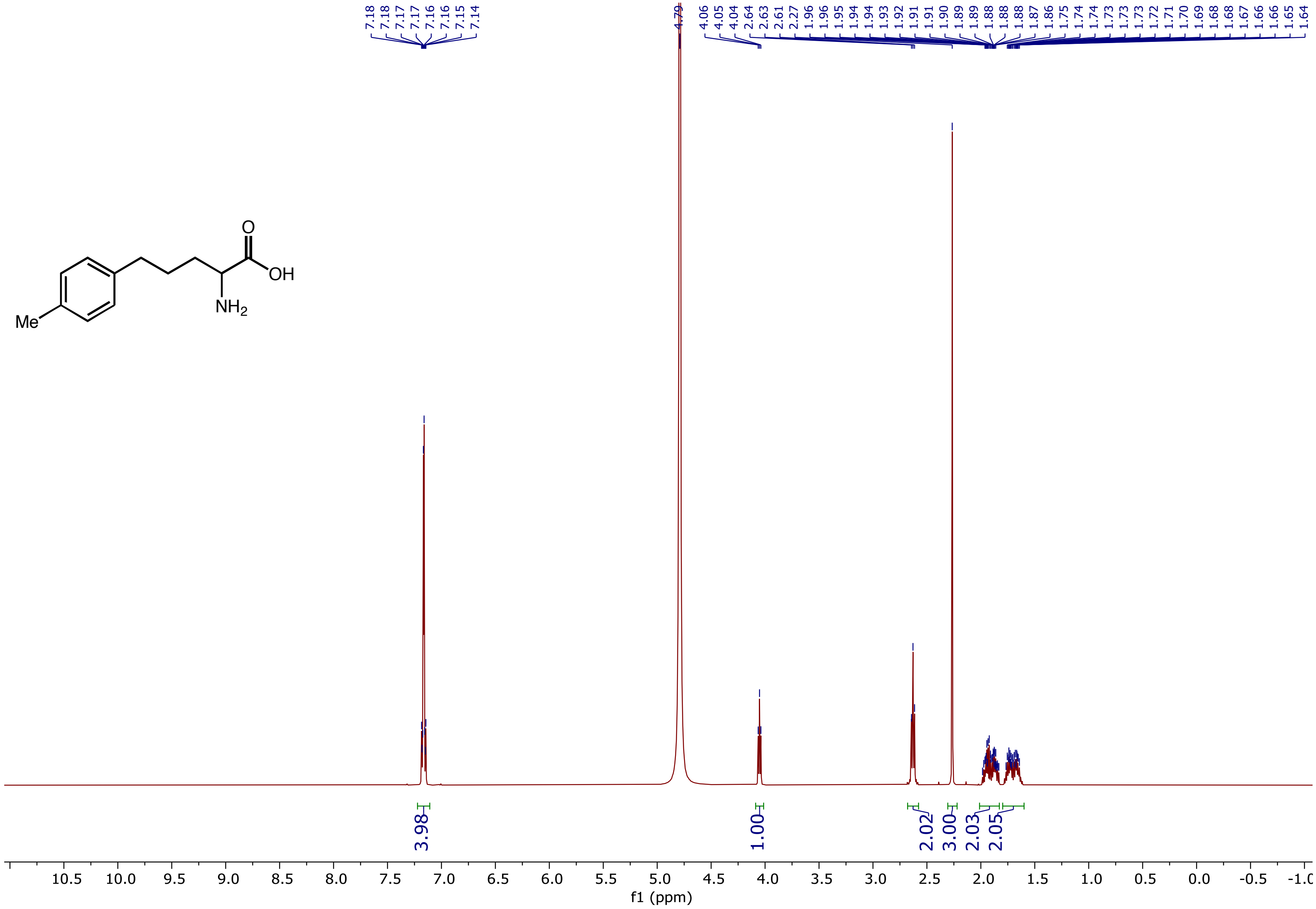

$^{13}C$ NMR (126 MHz, $D_2O$) spectrum of 2-Amino-5-(p-tolyl)pentanoic acid **(3b)**

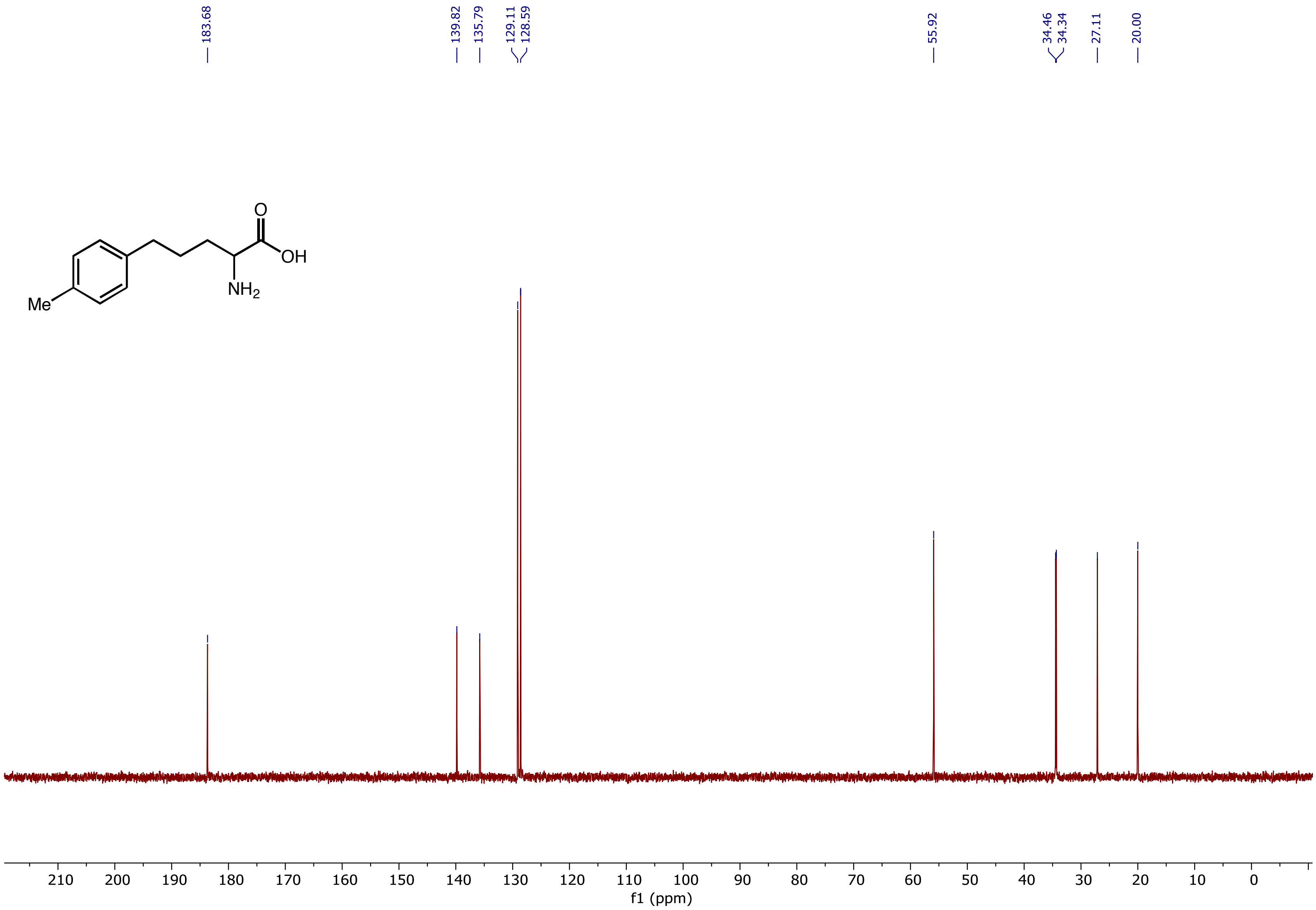

$^{1}H$ NMR (500 MHz, $D_2O$) spectrum of 2-Amino-5-(4-ethylphenyl)pentanoic acid **(3c)**

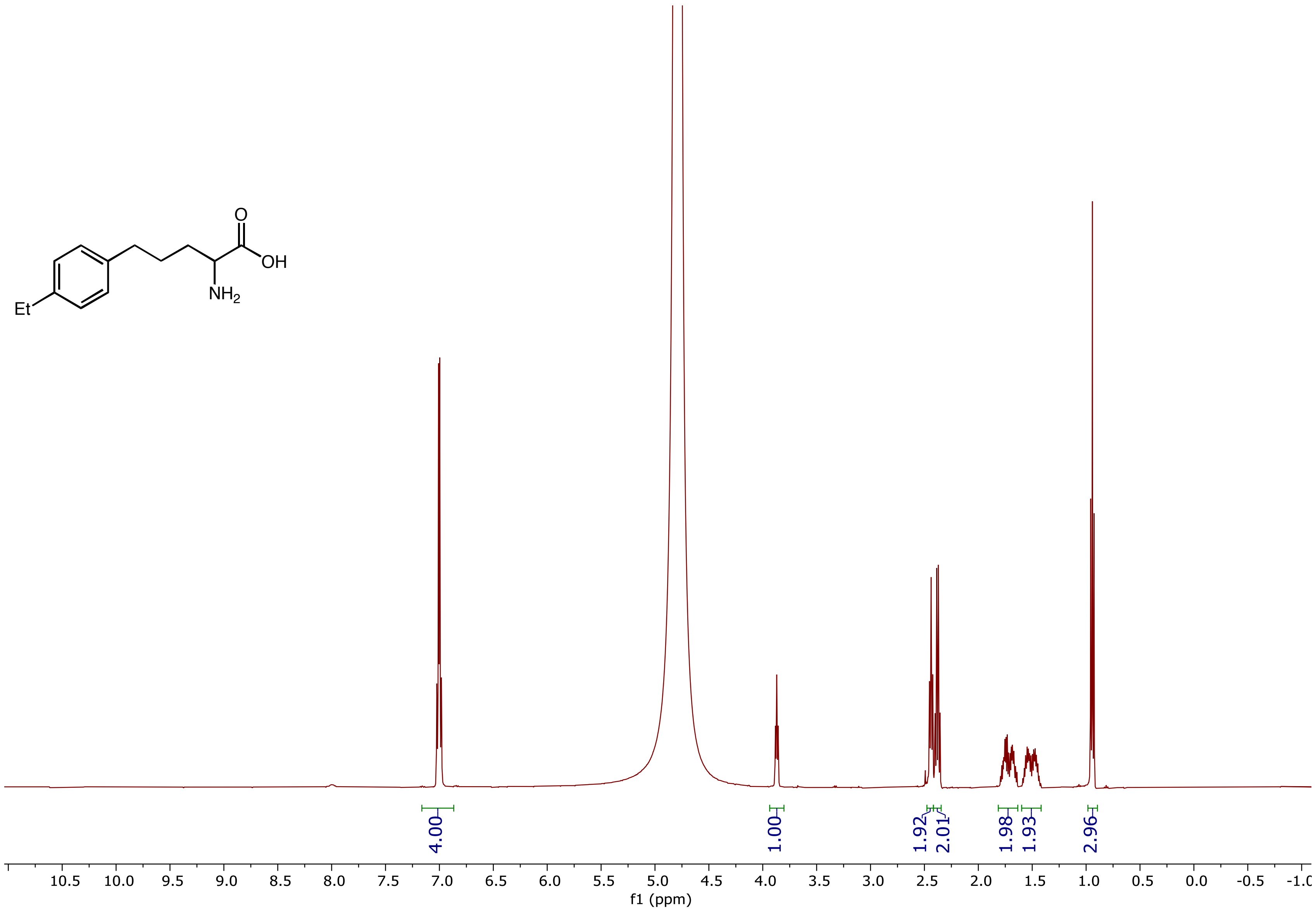

$^{13}$C NMR (126 MHz, $D_2O$) spectrum of 2-Amino-5-(4-ethylphenyl)pentanoic acid **(3c)**

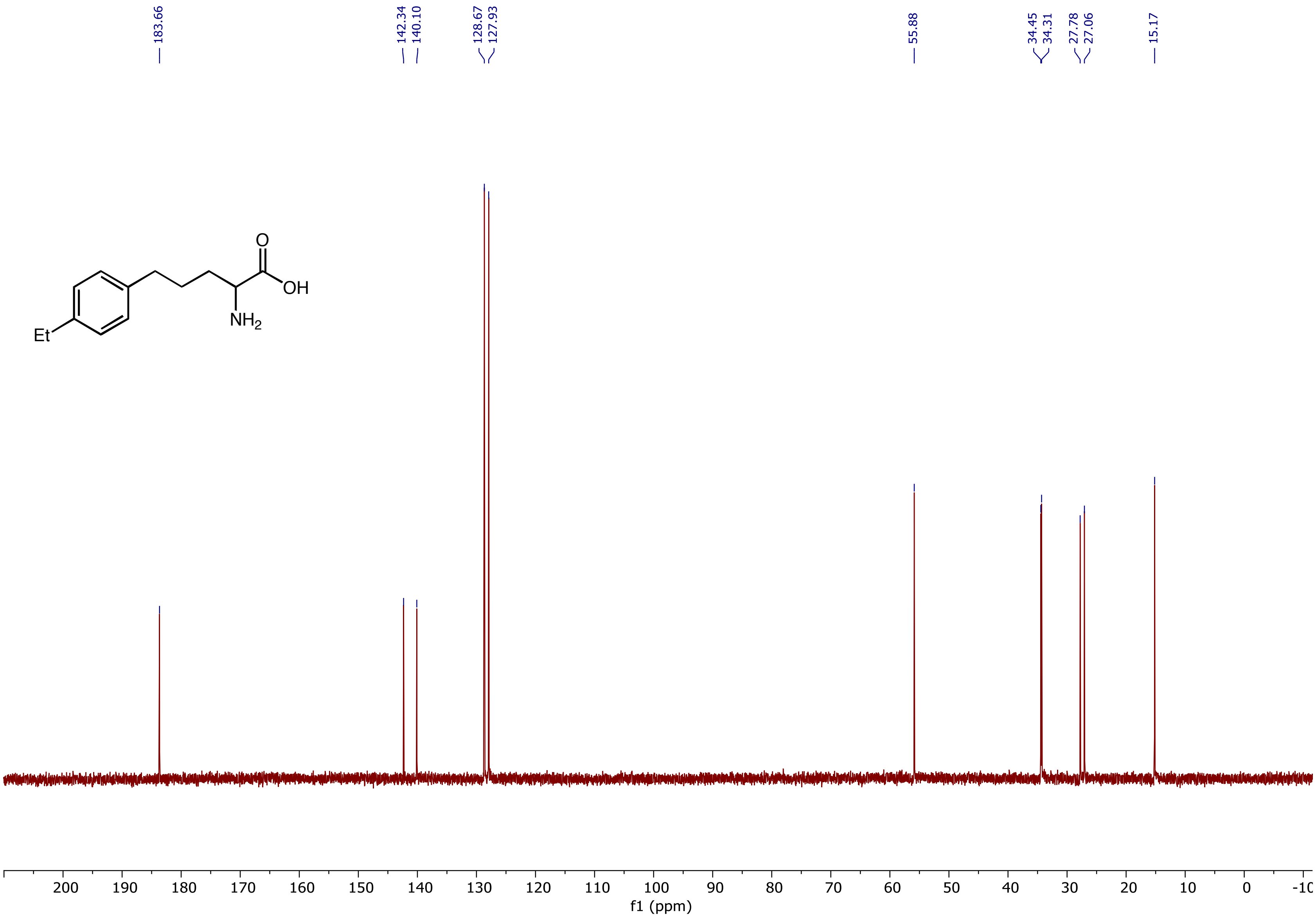

[1]H NMR (500 MHz, $D_2O$) spectrum of 2-Amino-5-(4-methoxyphenyl)pentanoic acid **(3d)**

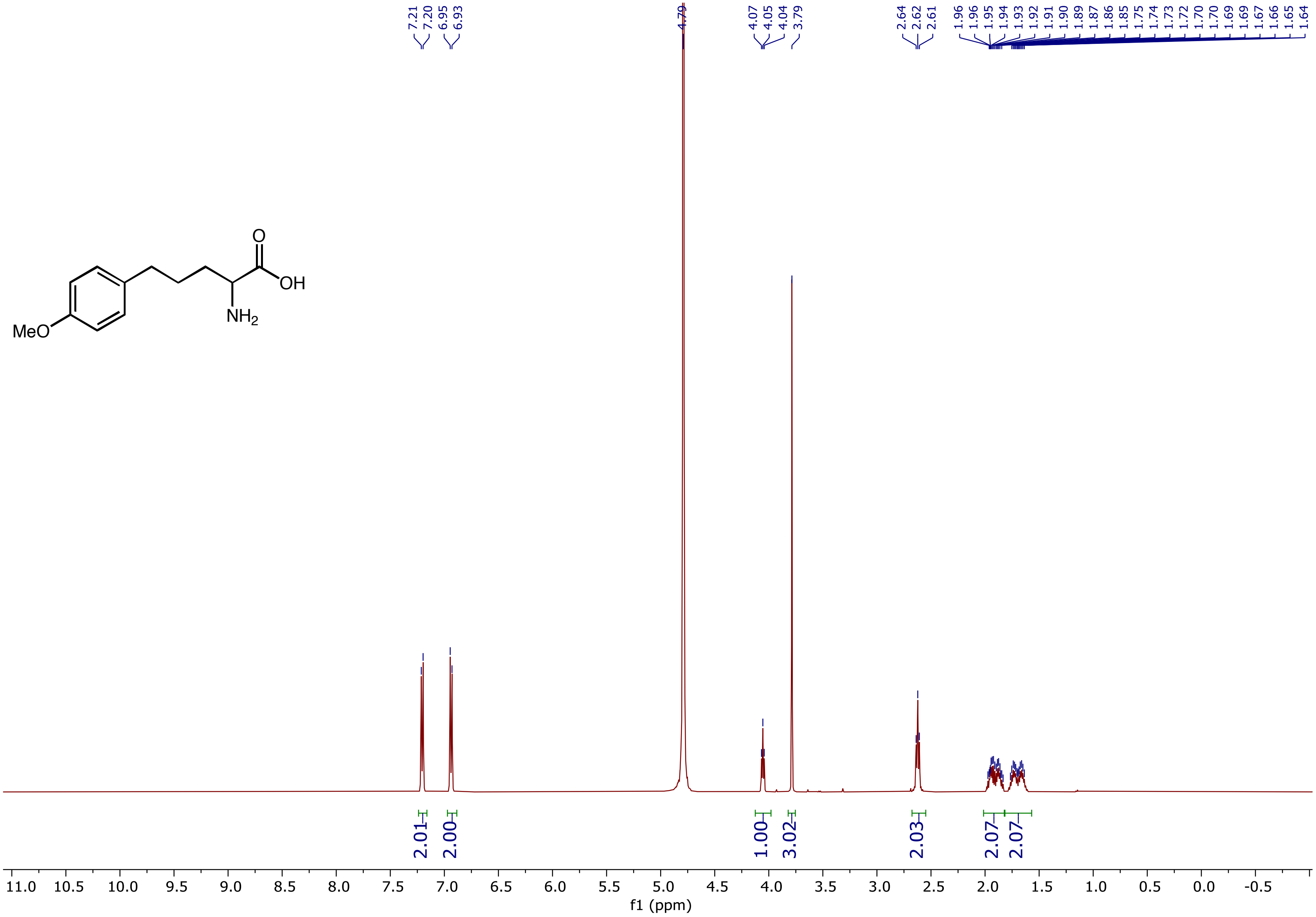

$^{13}$C NMR (126 MHz, $D_2O$) spectrum of 2-Amino-5-(4-methoxyphenyl)pentanoic acid **(3d)**

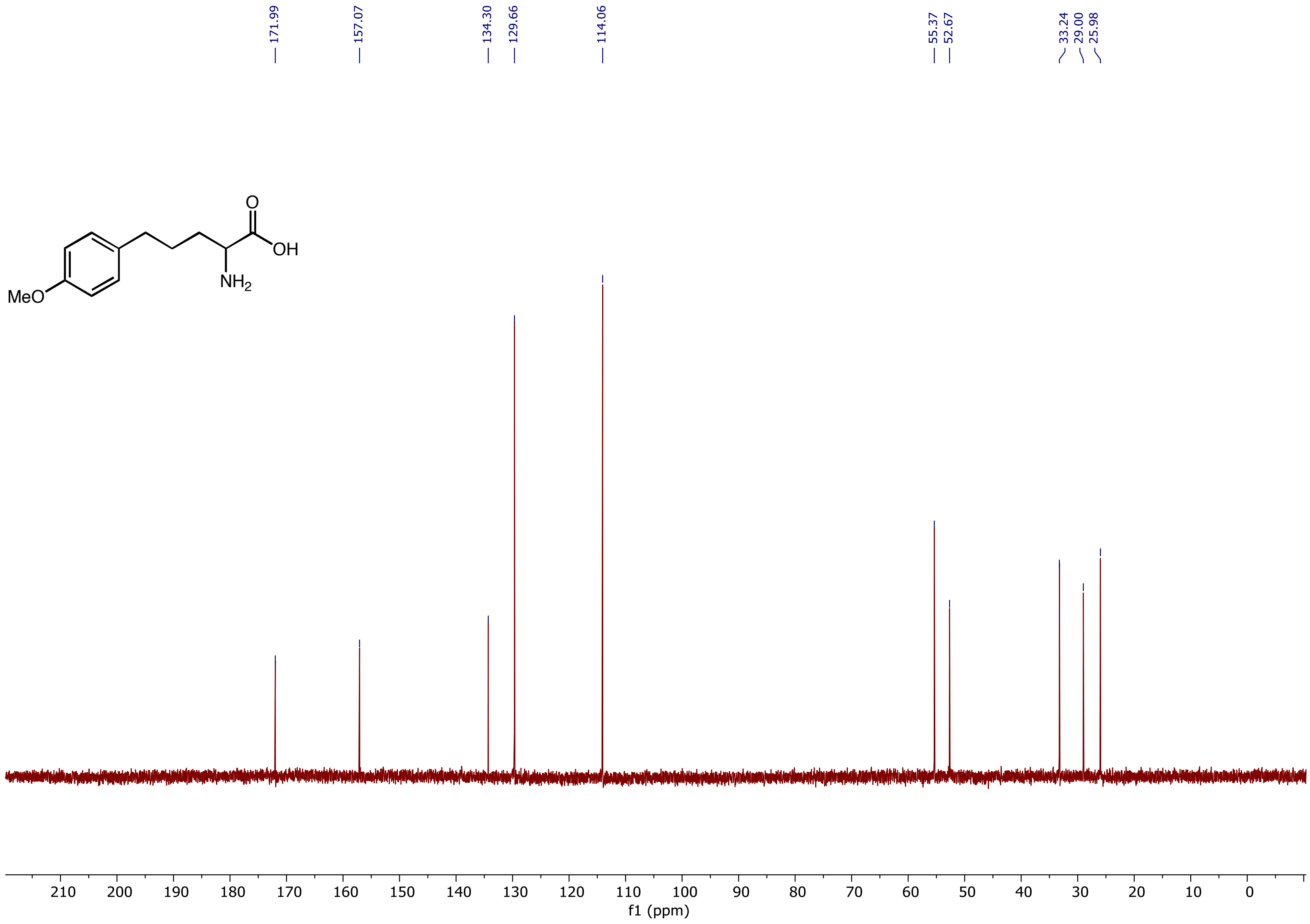

$^{1}H$ NMR (500 MHz, $D_2O$) spectrum of 2-Amino-5-(4-(methylthio)phenyl)pentanoic acid **(3e)**

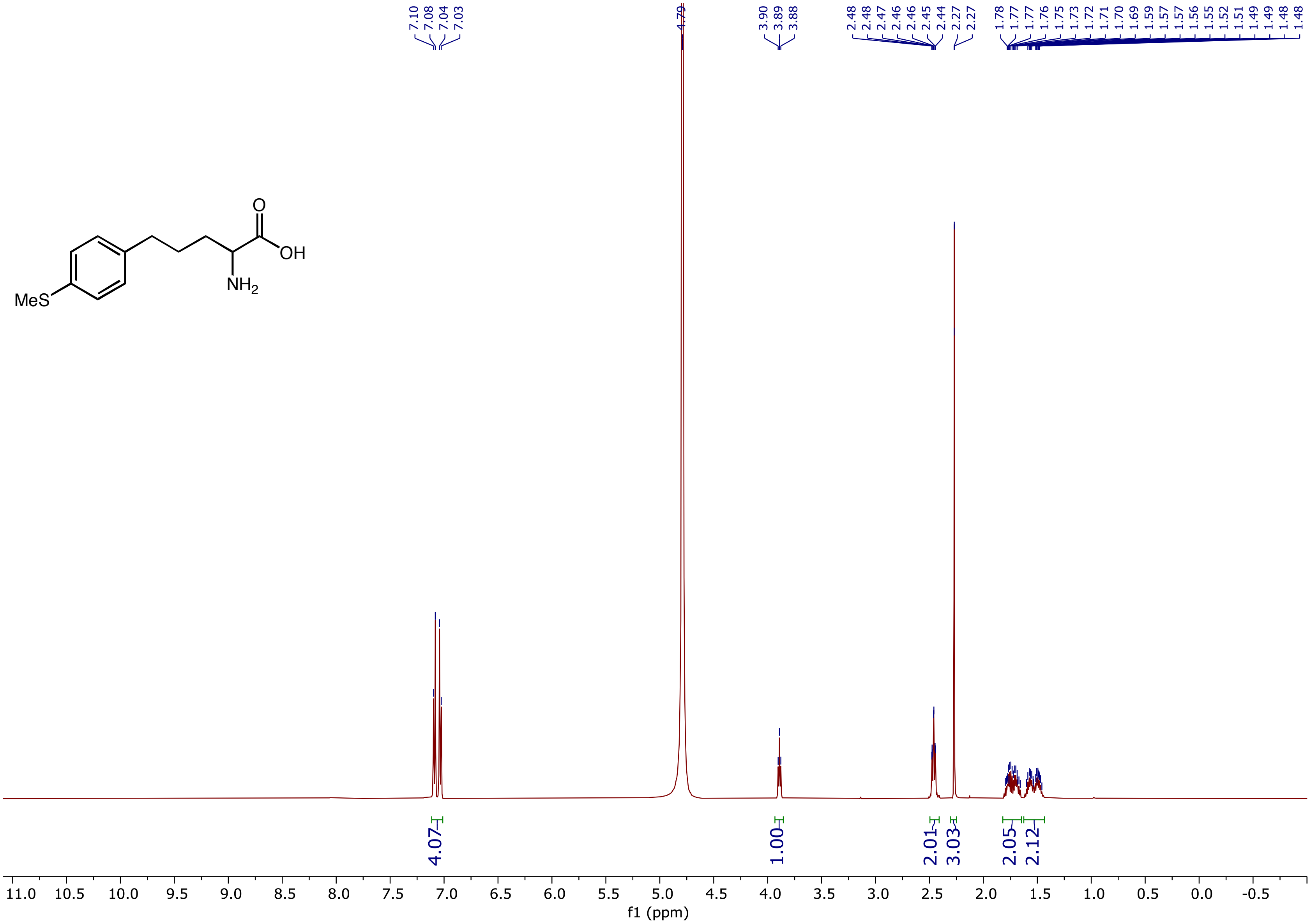

$^{13}C$ NMR (126 MHz, $D_2O$) spectrum of 2-Amino-5-(4-(methylthio)phenyl)pentanoic acid **(3e)**

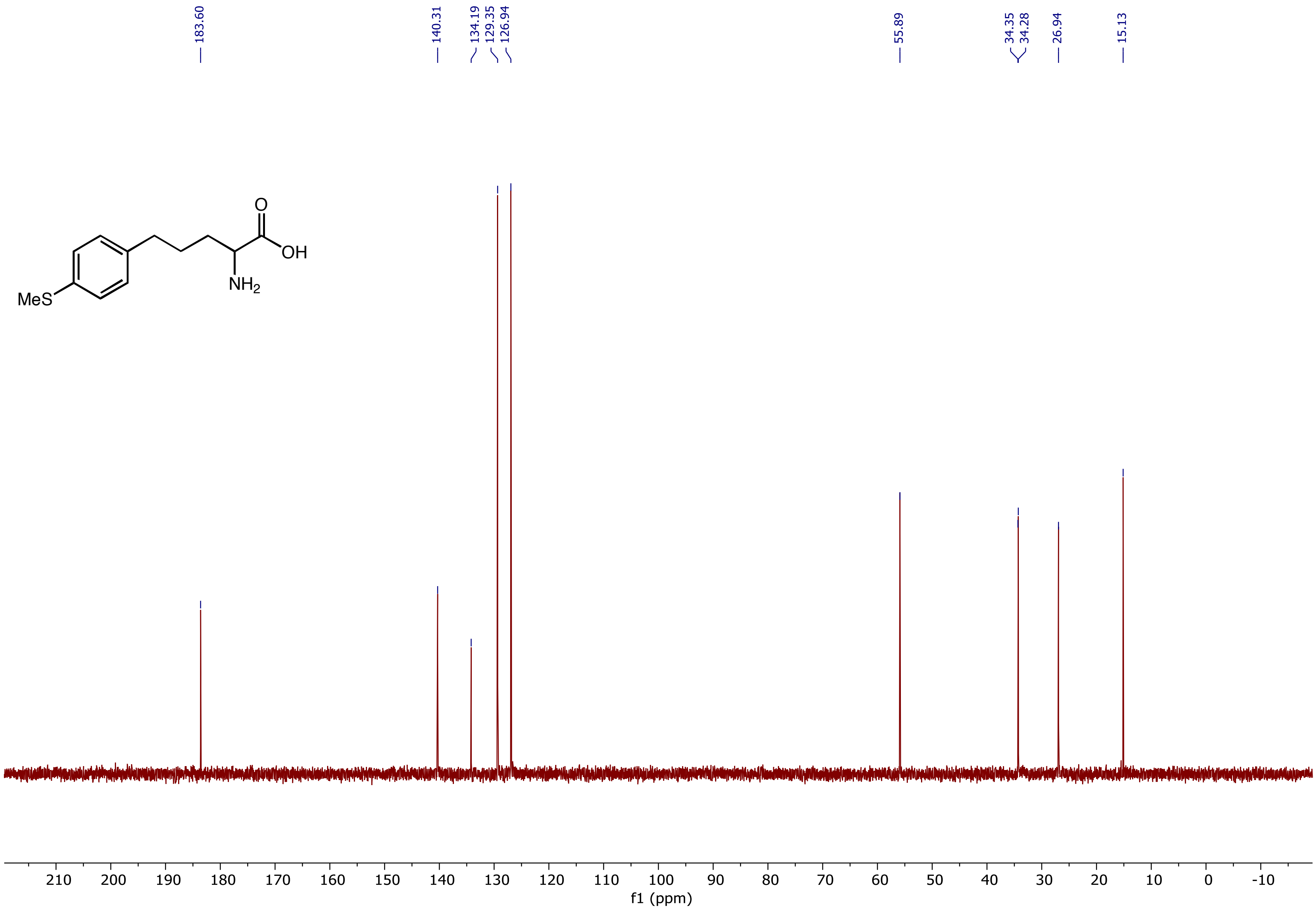

$^1H$ NMR (500 MHz, $D_2O$) spectrum of 2-Amino-5-(4-fluorophenyl)pentanoic acid **(3f)**

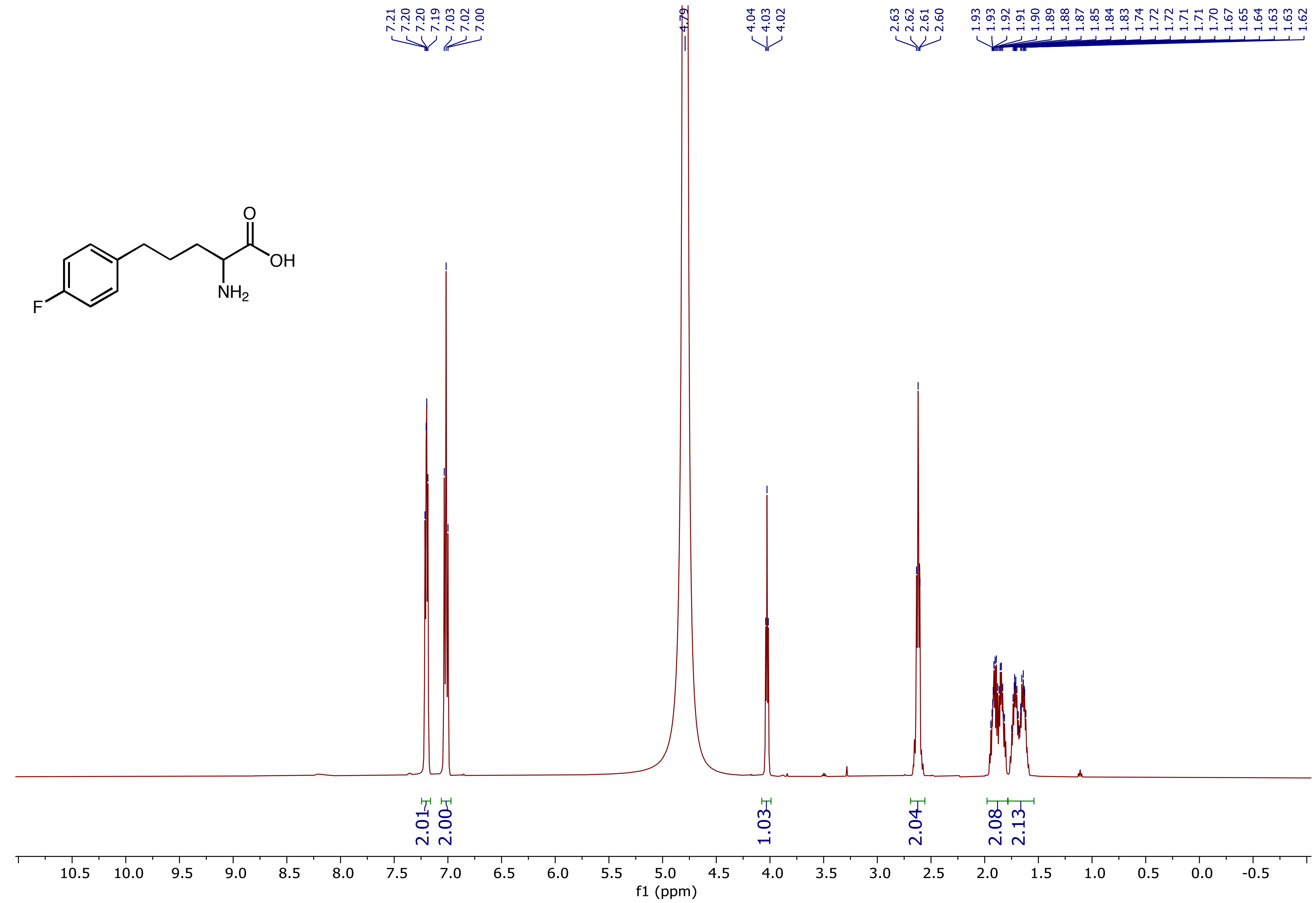

$^{13}C$ NMR (126 MHz, $D_2O$) spectrum of 2-Amino-5-(4-fluorophenyl)pentanoic acid **(3f)**

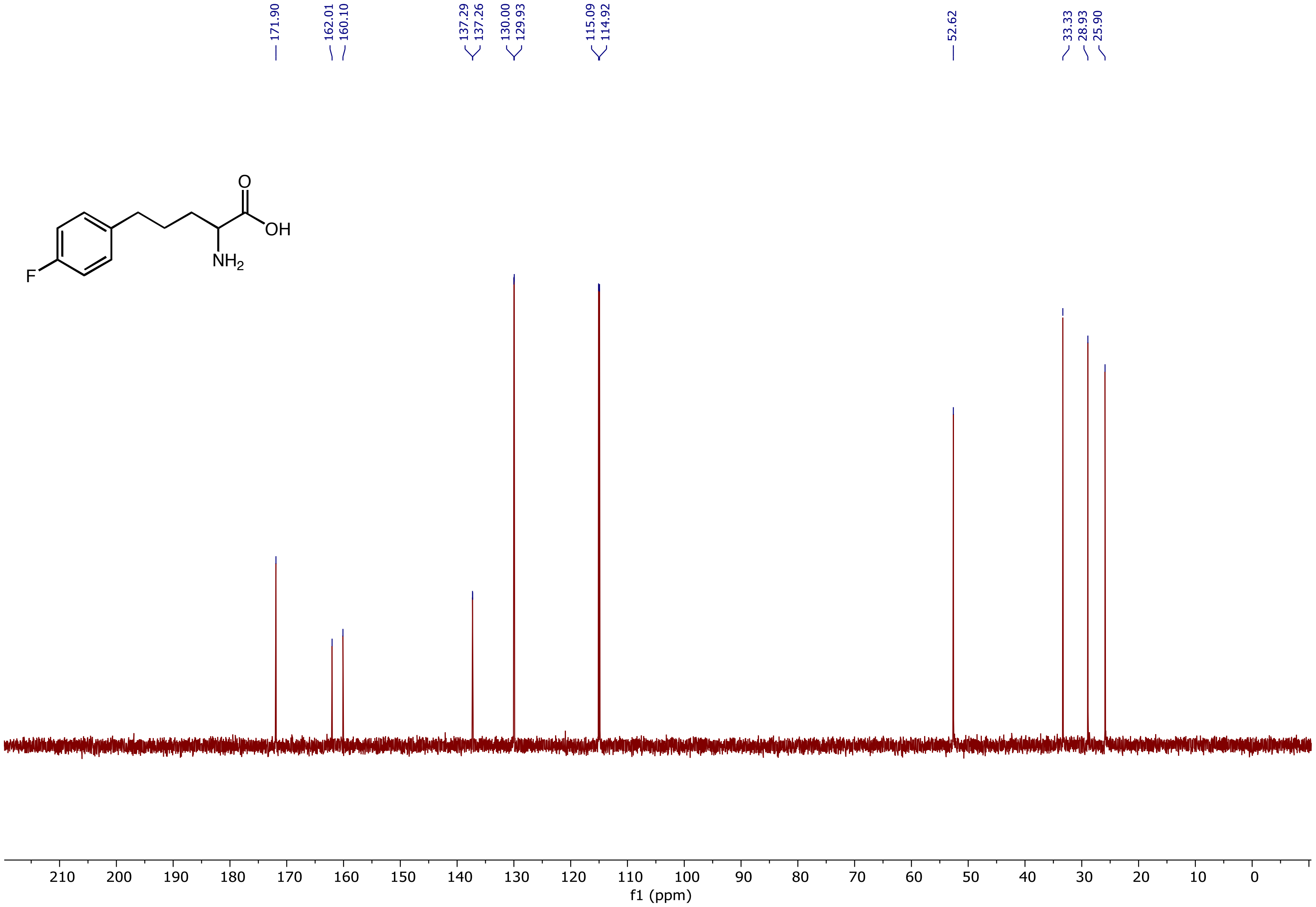

$^{19}F$ NMR (471 MHz, $D_2O$) spectrum of 2-Amino-5-(4-fluorophenyl)pentanoic acid **(3f)**

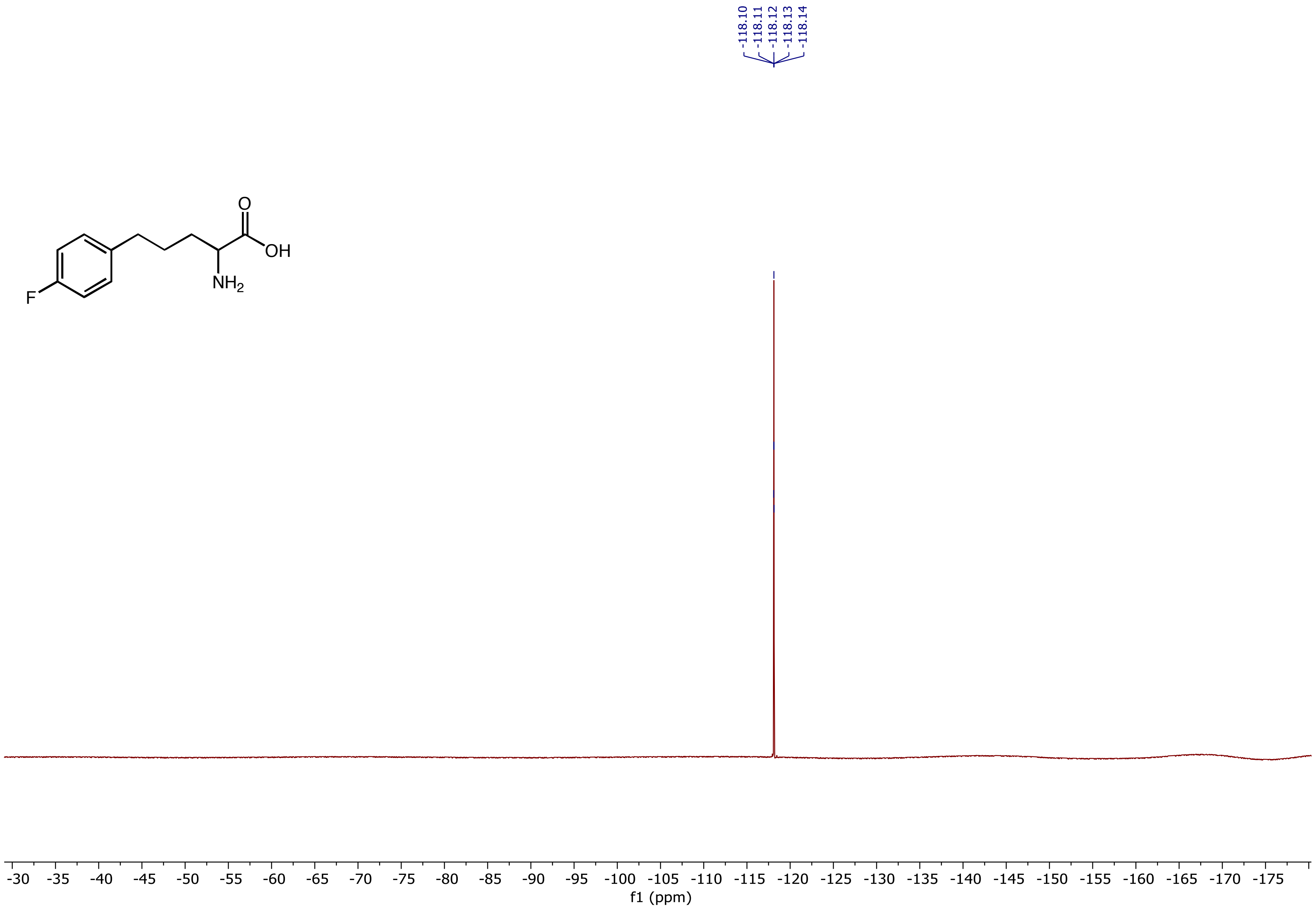

$^1H$ NMR (500 MHz, $D_2O$) spectrum of 2-Amino-5-(4-chlorophenyl)pentanoic acid **(3g)**

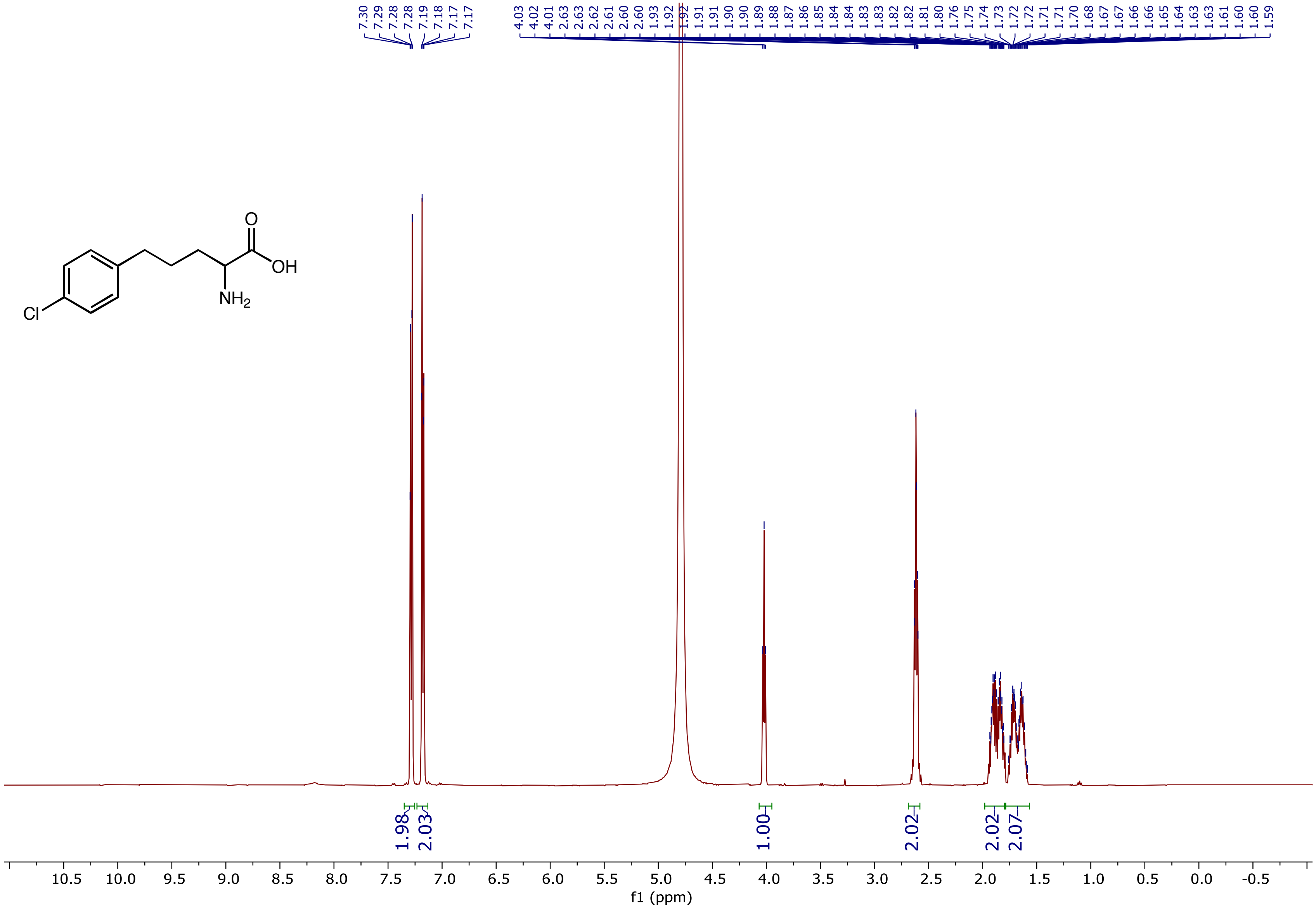

$^{13}C$ NMR (126 MHz, $D_2O$) spectrum of 2-Amino-5-(4-chlorophenyl)pentanoic acid **(3g)**

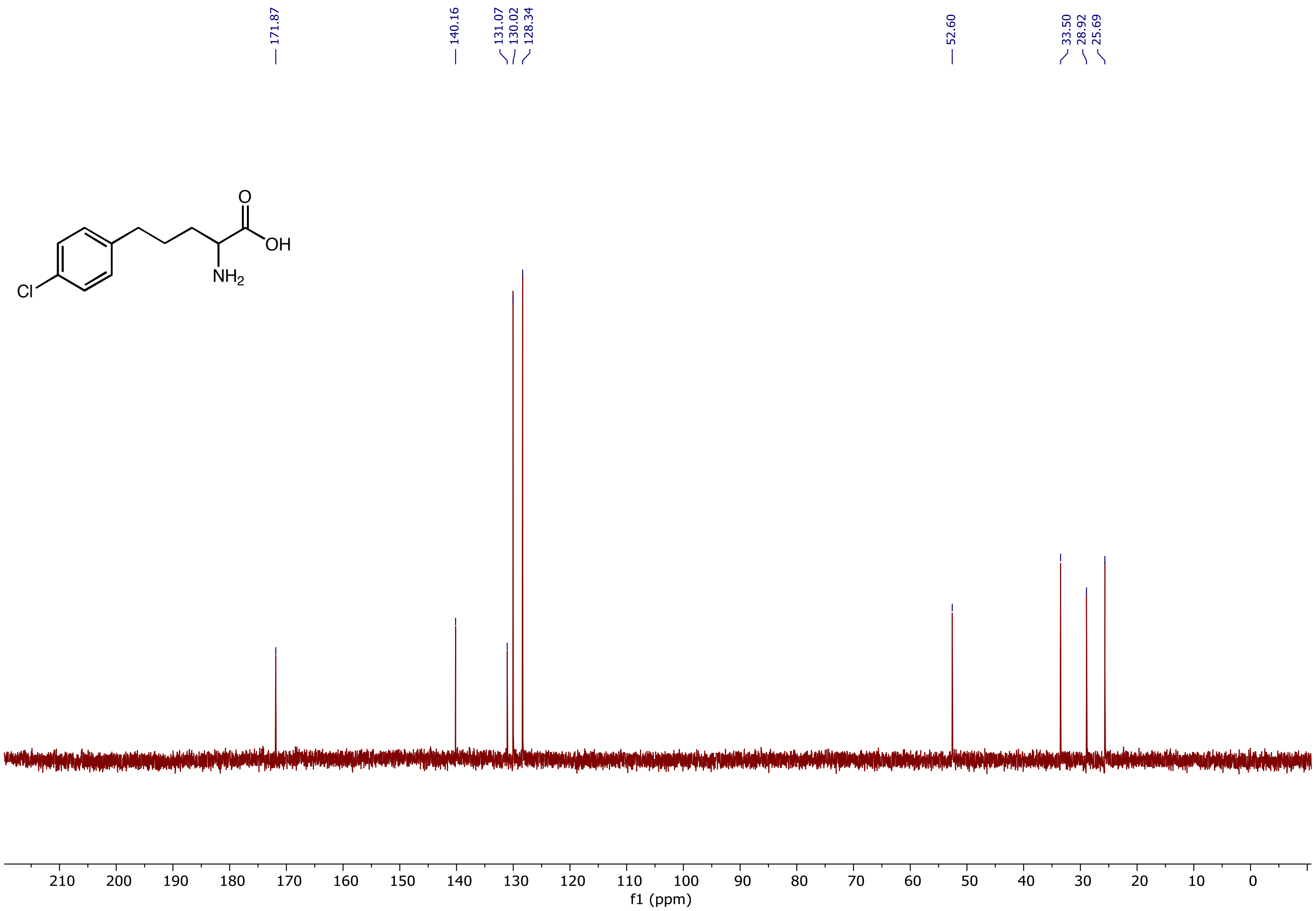

$^{1}H$ NMR (400 MHz, $D_2O$) spectrum of 2-Amino-5-(4-bromophenyl)pentanoic acid **(3h)**

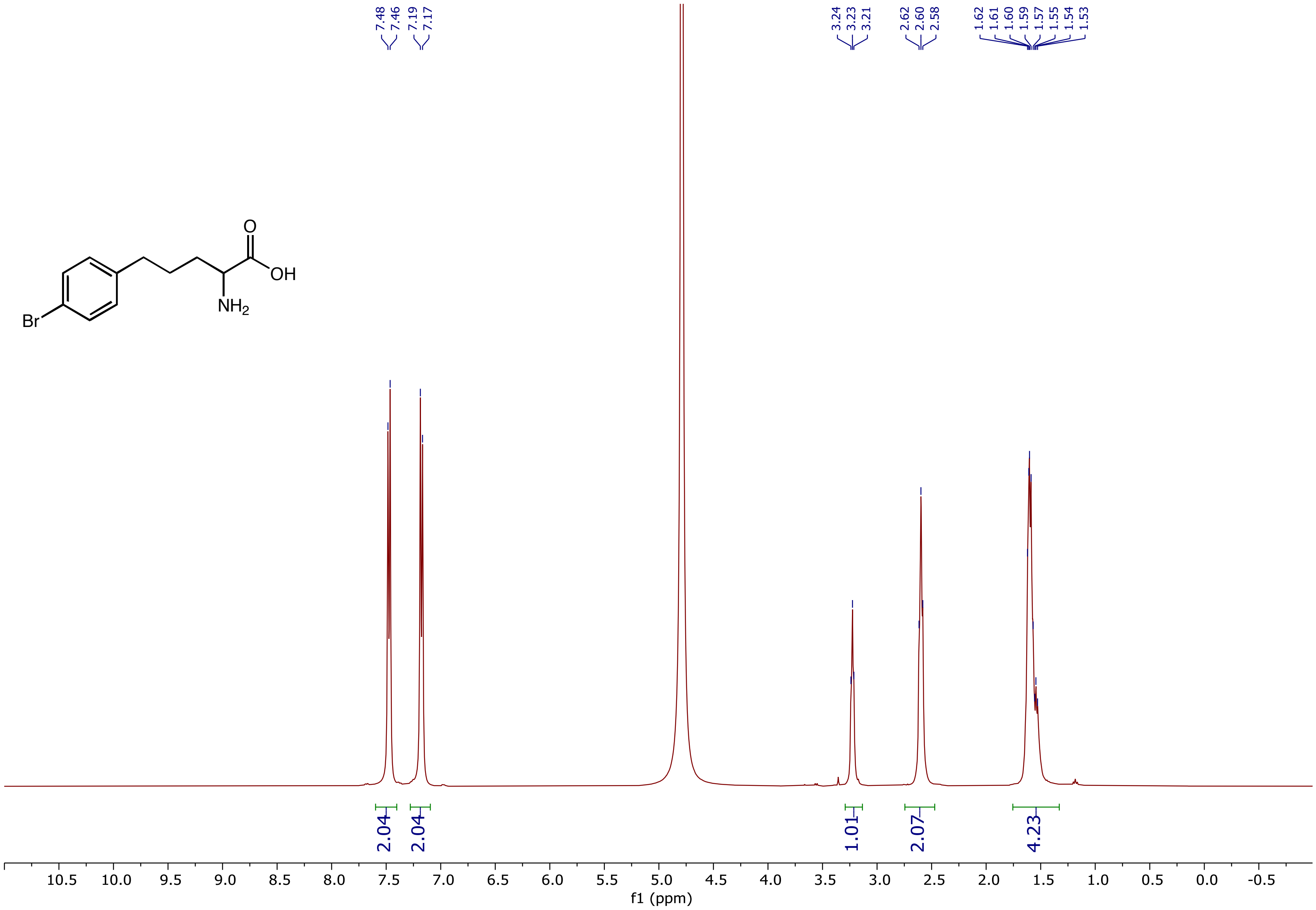

$^{13}C$ NMR (101 MHz, $D_2O$) spectrum of 2-Amino-5-(4-bromophenyl)pentanoic acid **(3h)**

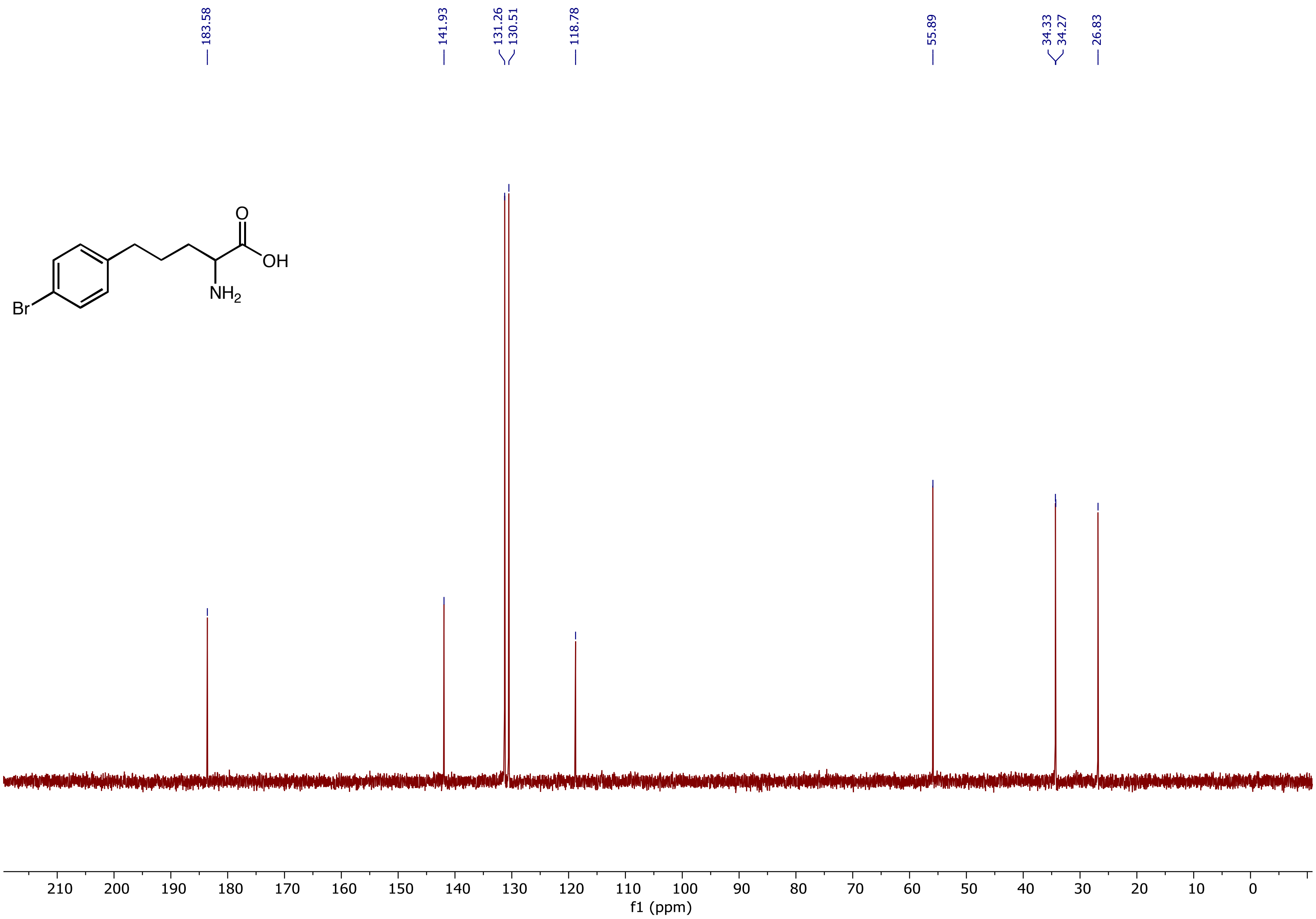

$^{1}H$ NMR (400 MHz, $D_2O$) spectrum of 2-Amino-5-(4-cyanophenyl)pentanoic acid **(3i)**

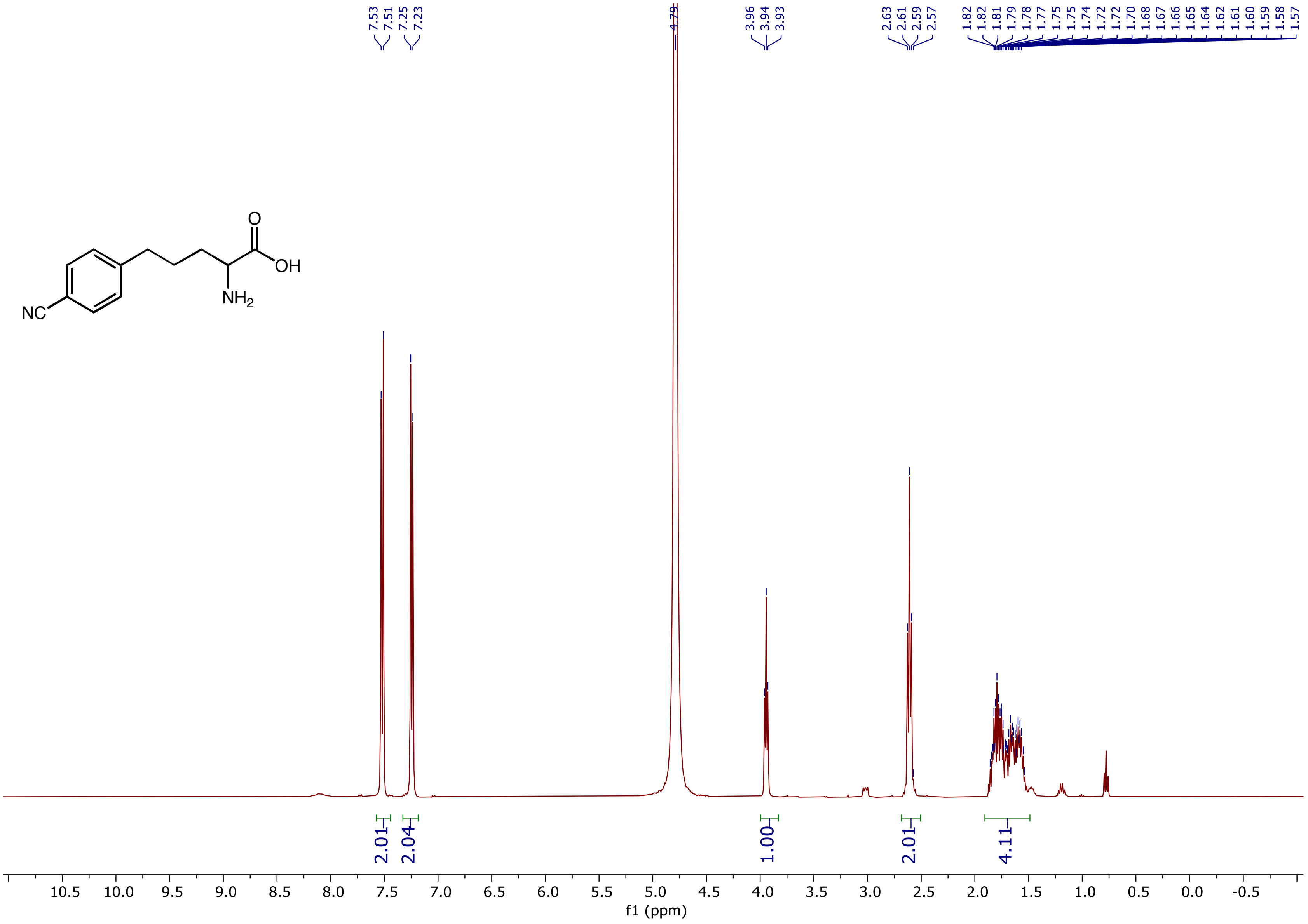

$^{13}C$ NMR (101 MHz, $D_2O$) spectrum of 2-Amino-5-(4-cyanophenyl)pentanoic acid **(3i)**

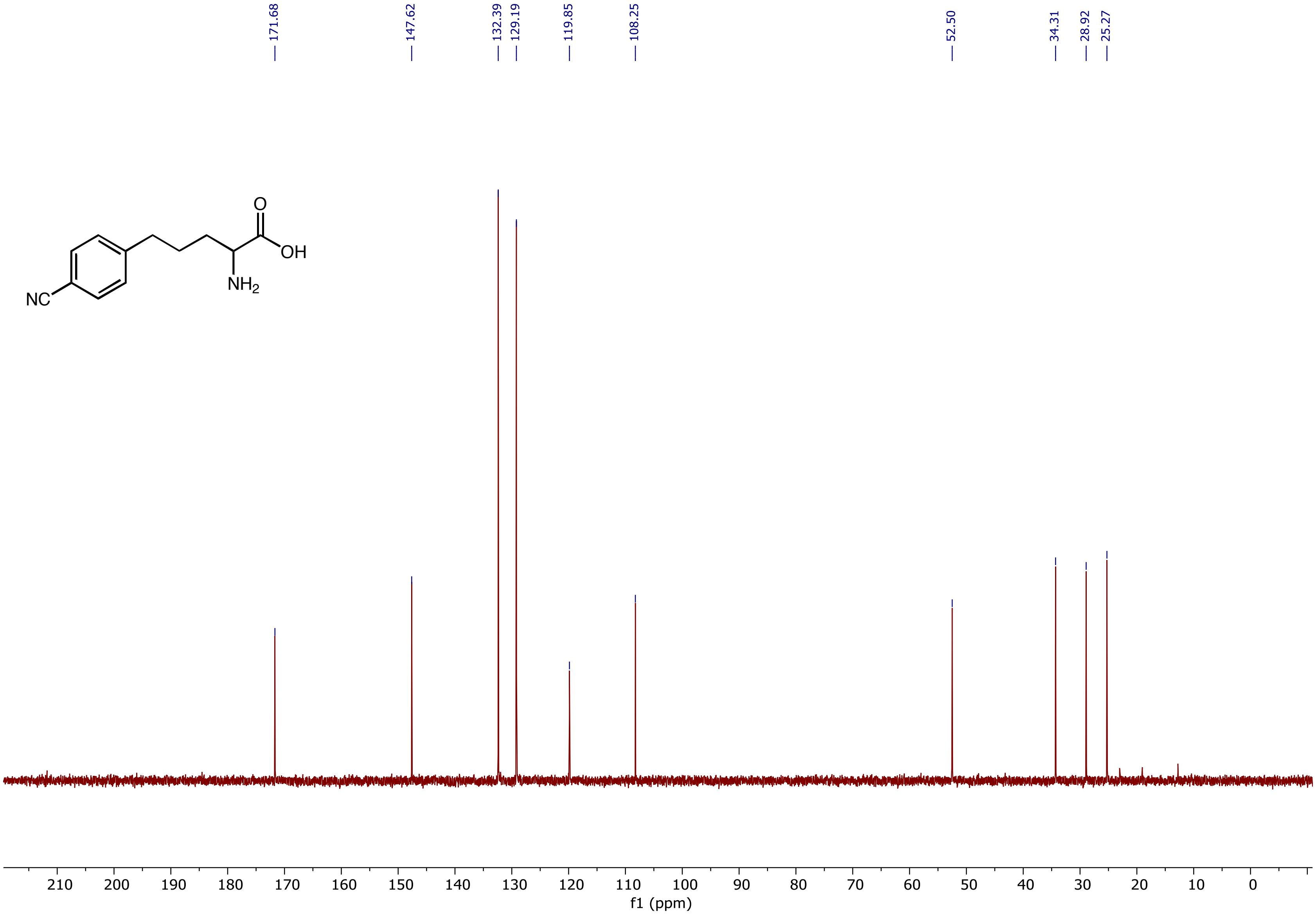

$^{1}H$ NMR (500 MHz, $D_2O$) spectrum of 2-Amino-5-(4-(trifluoromethyl)phenyl)pentanoic acid **(3j)**

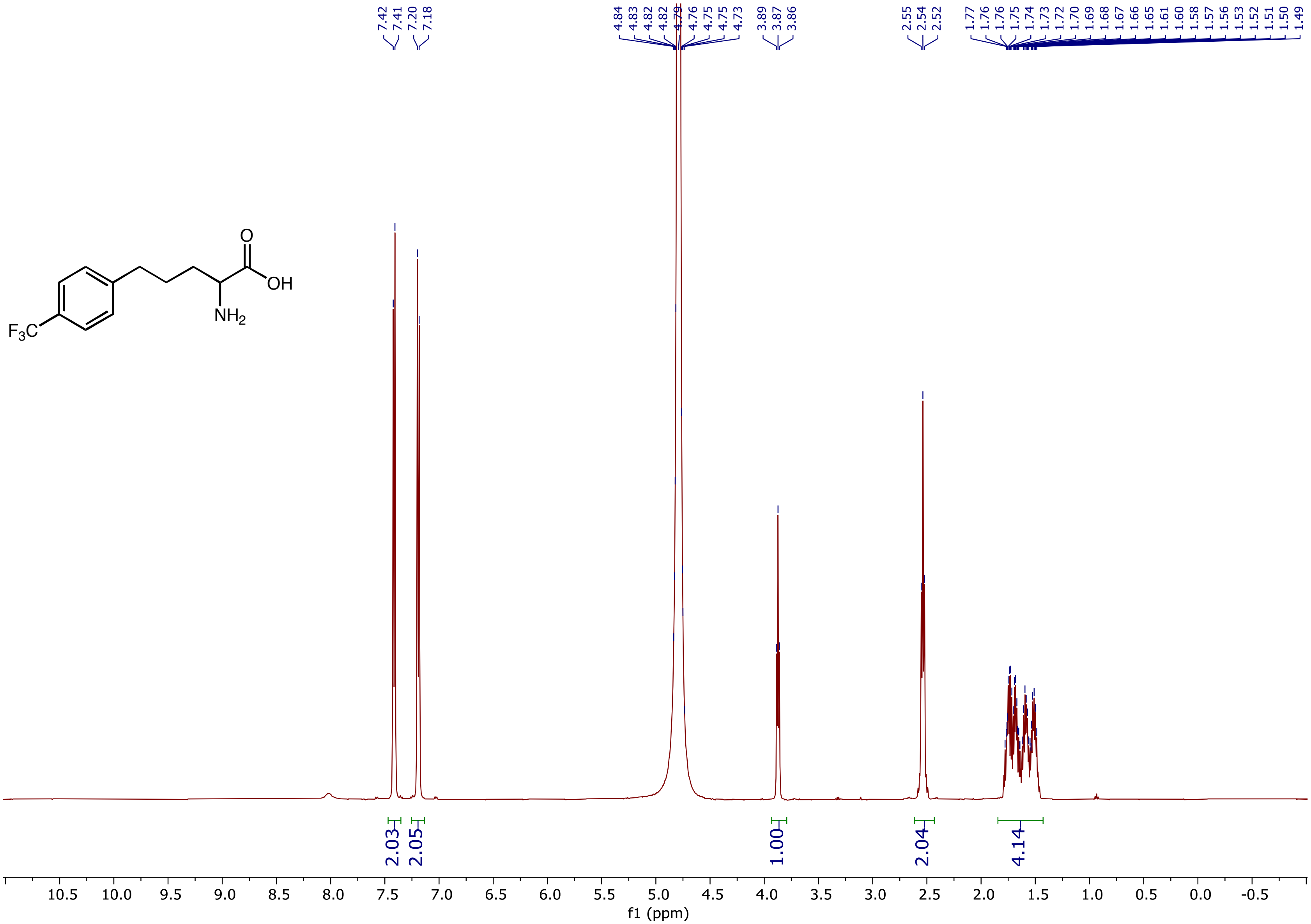

$^{13}C$ NMR (126 MHz, $D_2O$) spectrum of 2-Amino-5-(4-(trifluoromethyl)phenyl)pentanoic acid **(3j)**

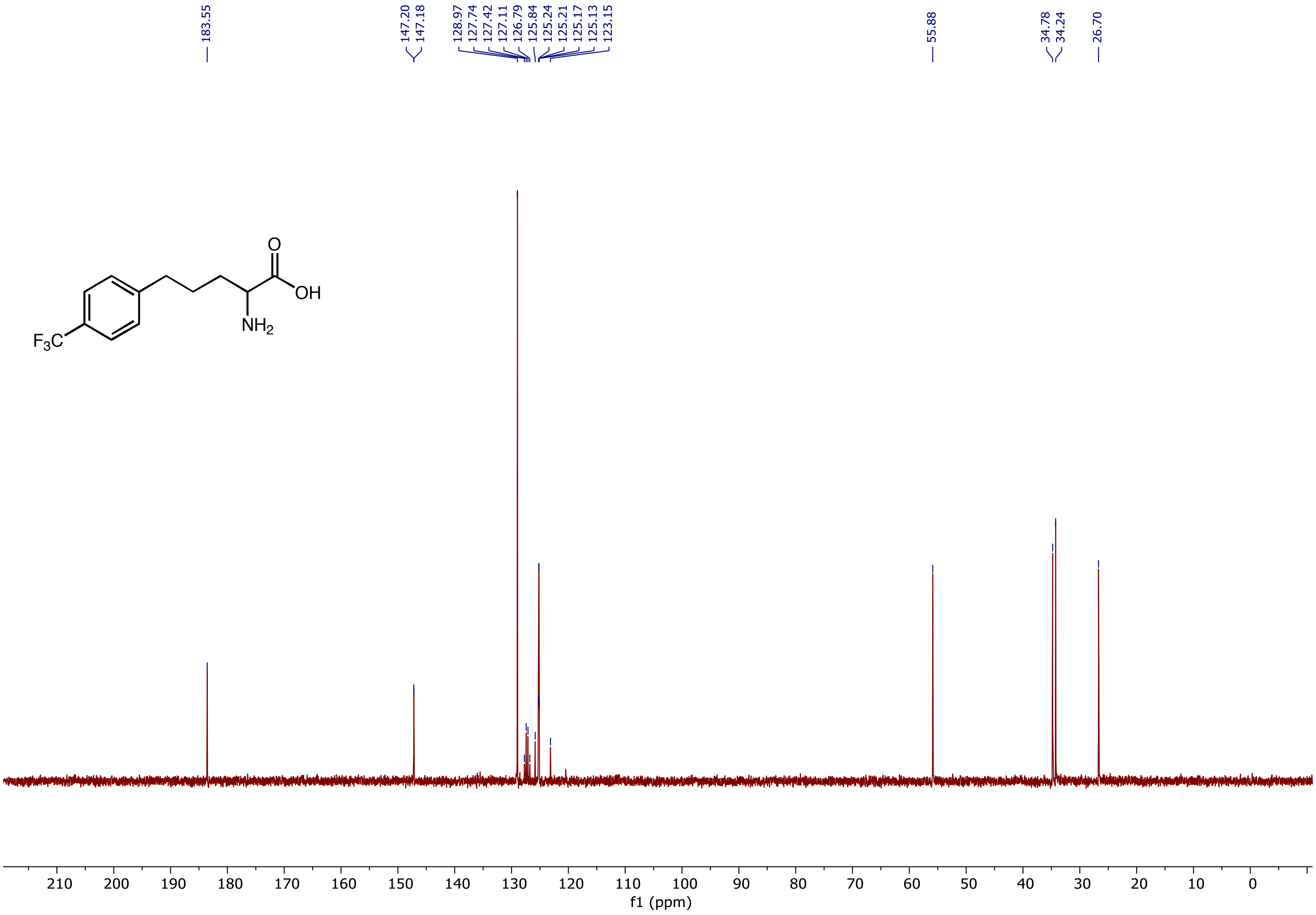

$^{19}F$ NMR (471 MHz, $D_2O$) spectrum of 2-Amino-5-(4-(trifluoromethyl)phenyl)pentanoic acid **(3j)**

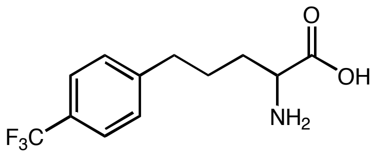

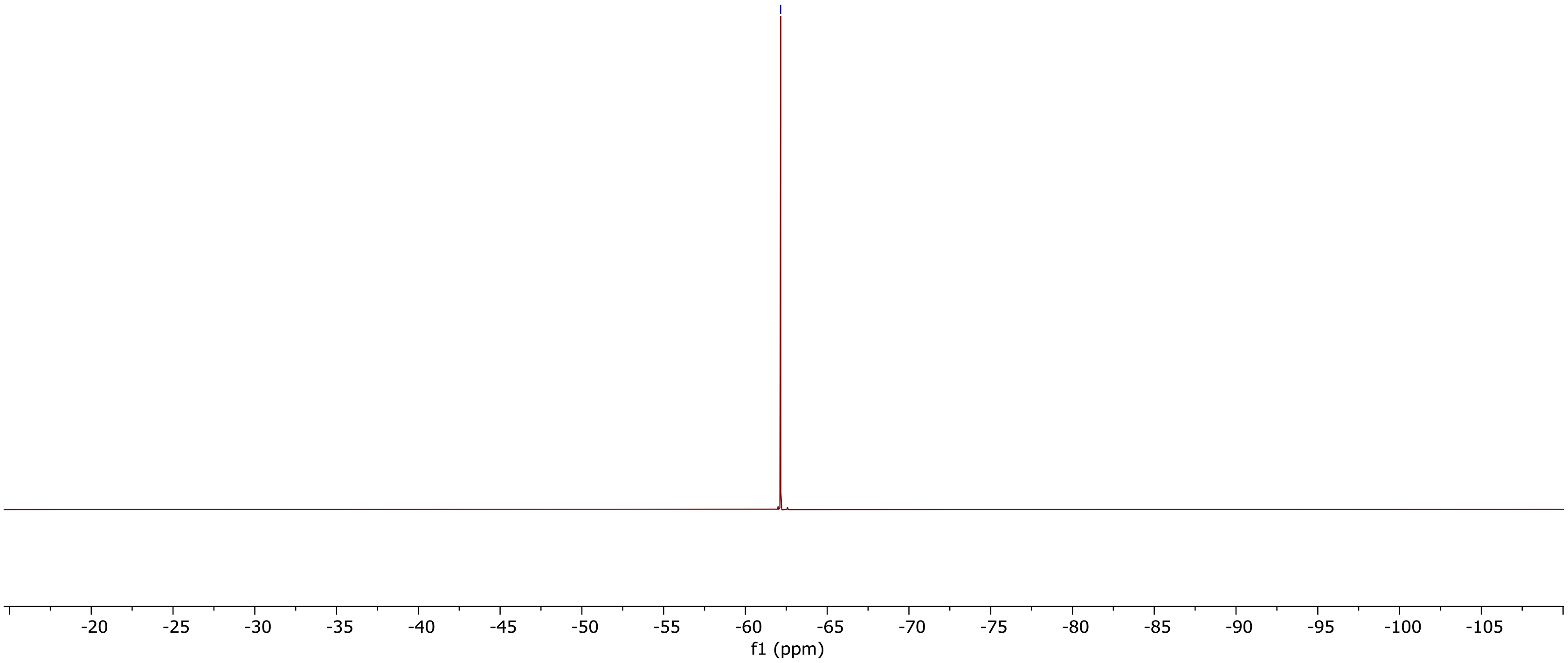

$^1H$ NMR (500 MHz, $D_2O$) spectrum of 2-Amino-5-(m-tolyl)pentanoic acid **(3k)**

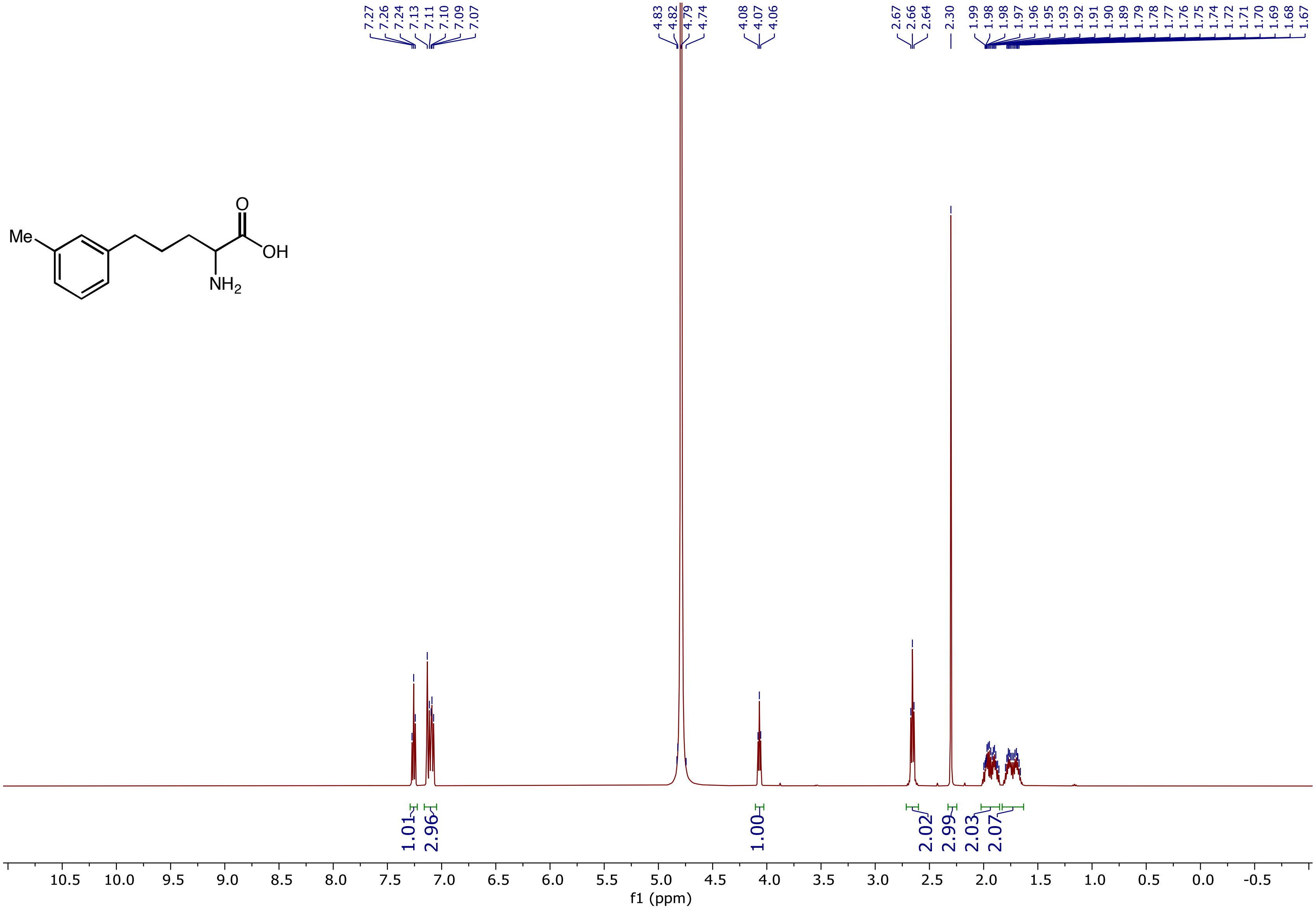

$^{13}$C NMR (126 MHz, $D_2O$) spectrum of 2-Amino-5-(m-tolyl)pentanoic acid **(3k)**

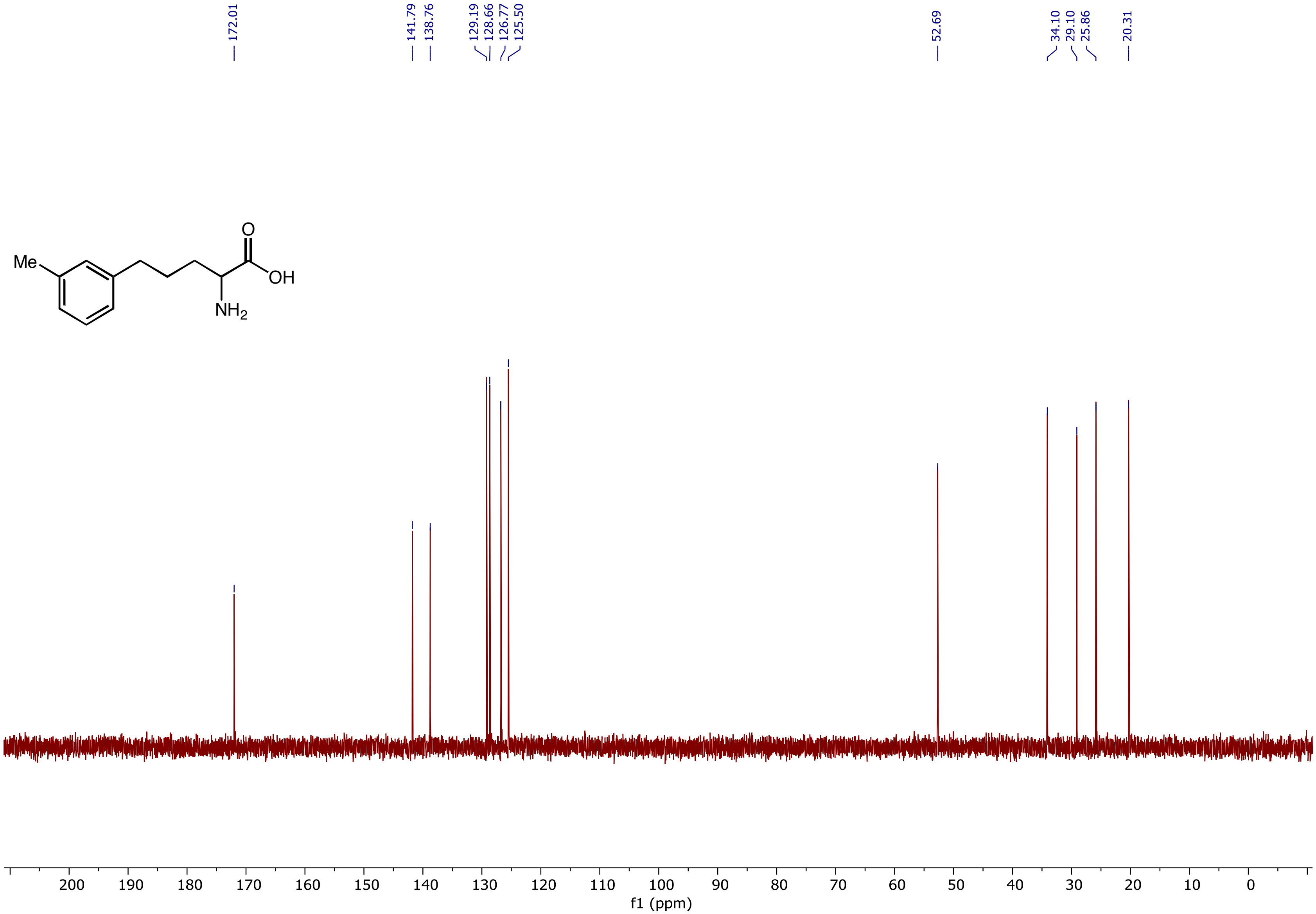

[1]H NMR (500 MHz, $D_2O$) spectrum of 2-Amino-5-(o-tolyl)pentanoic acid **(3l)**

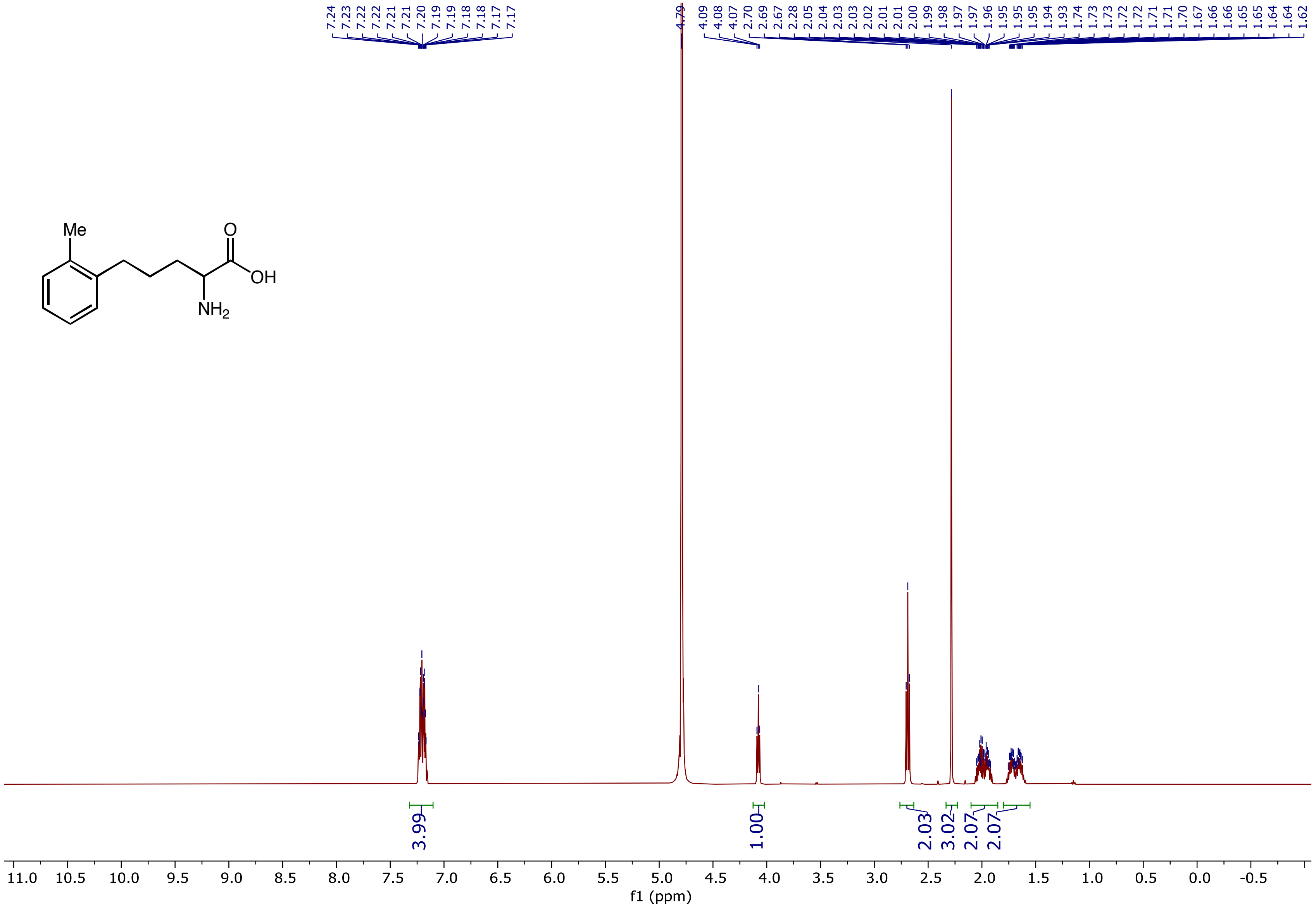

$^{13}C$ NMR (126 MHz, $D_2O$) spectrum of 2-Amino-5-(o-tolyl)pentanoic acid **(3l)**

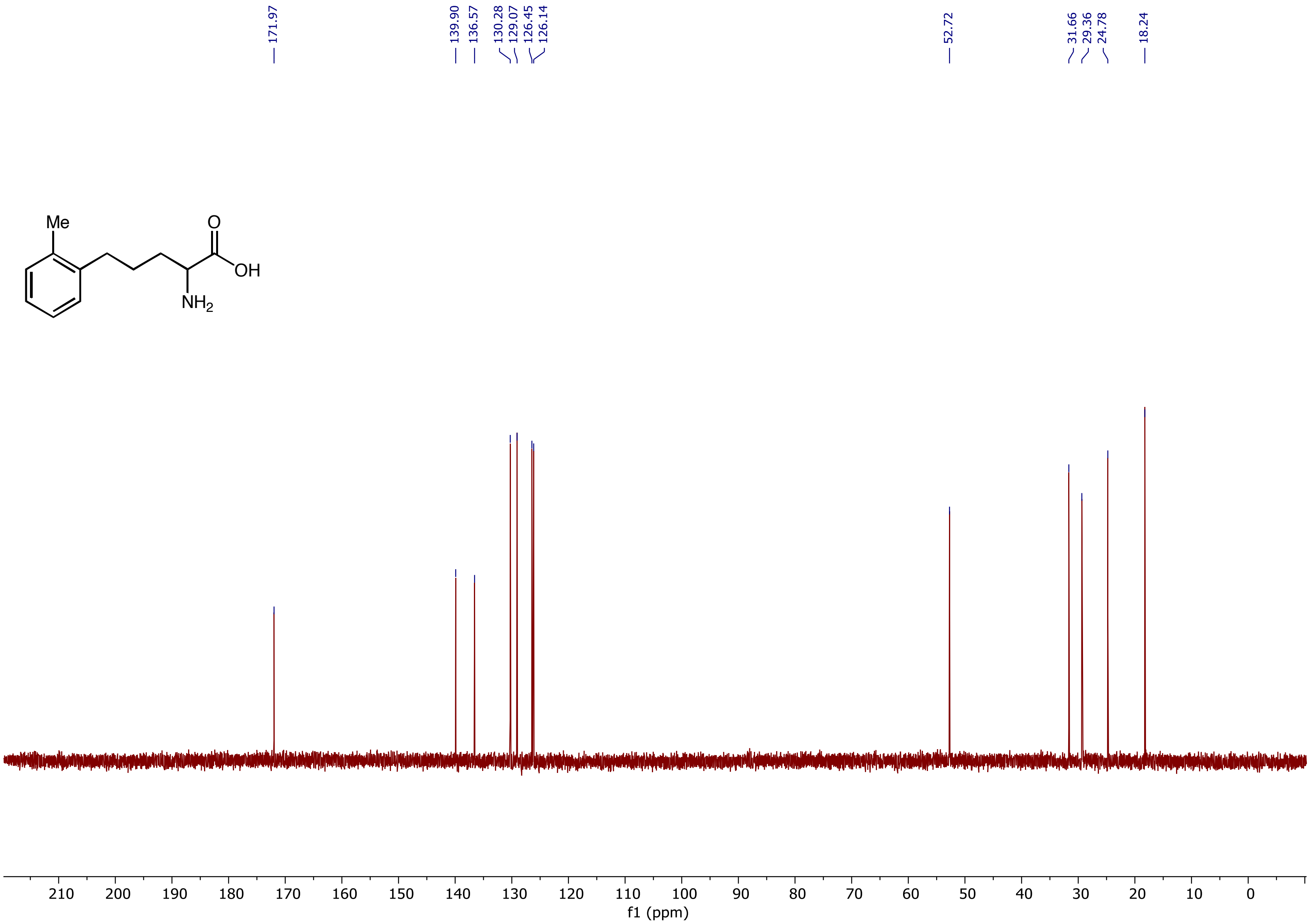

$^{1}H$ NMR (500 MHz, $D_2O$) spectrum of 2-Amino-5-(furan-2-yl)pentanoic acid **(3m)**

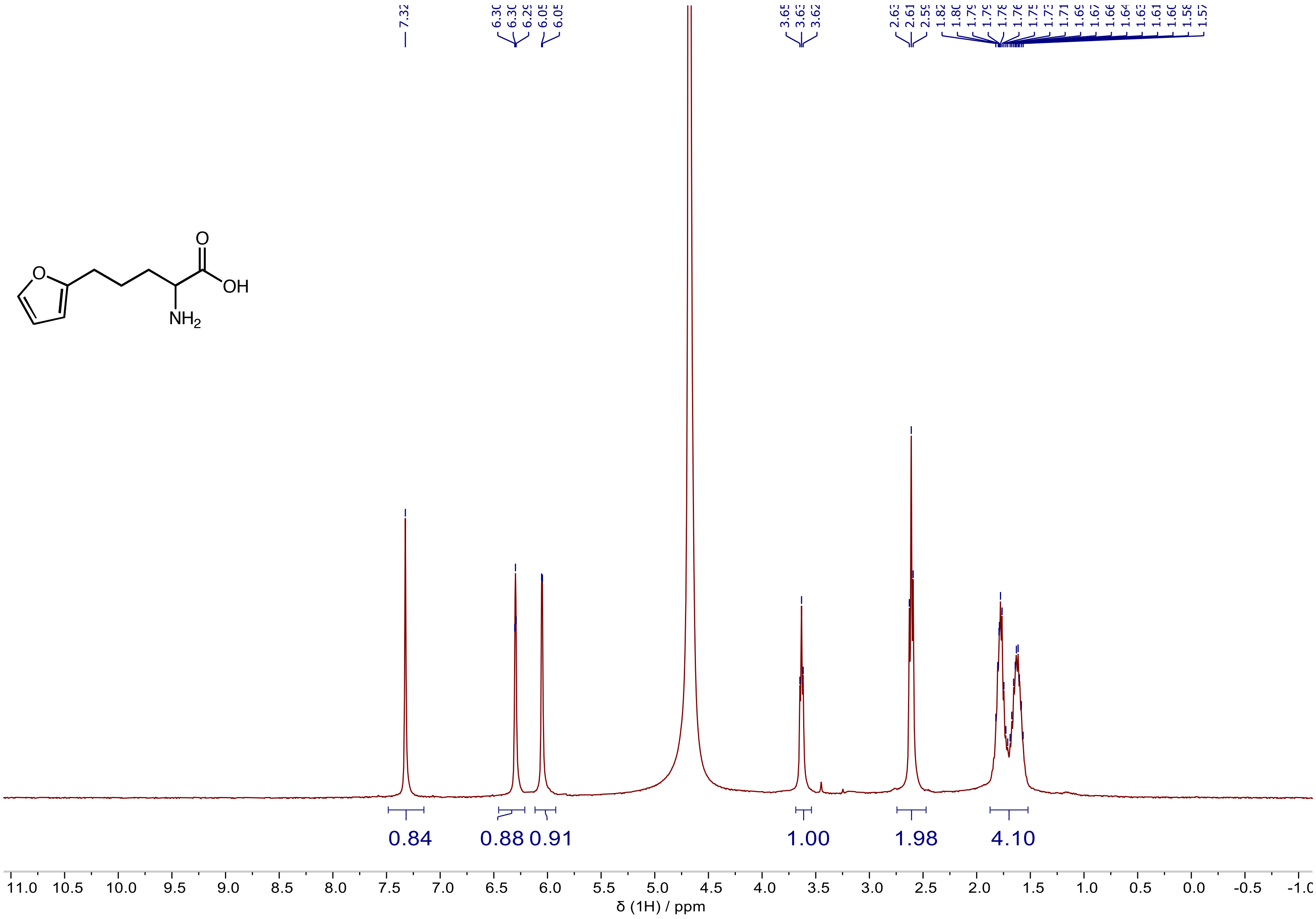

$^{13}$C NMR (101 MHz, $D_2O$) spectrum of 2-Amino-5-(furan-2-yl)pentanoic acid **(3m)**

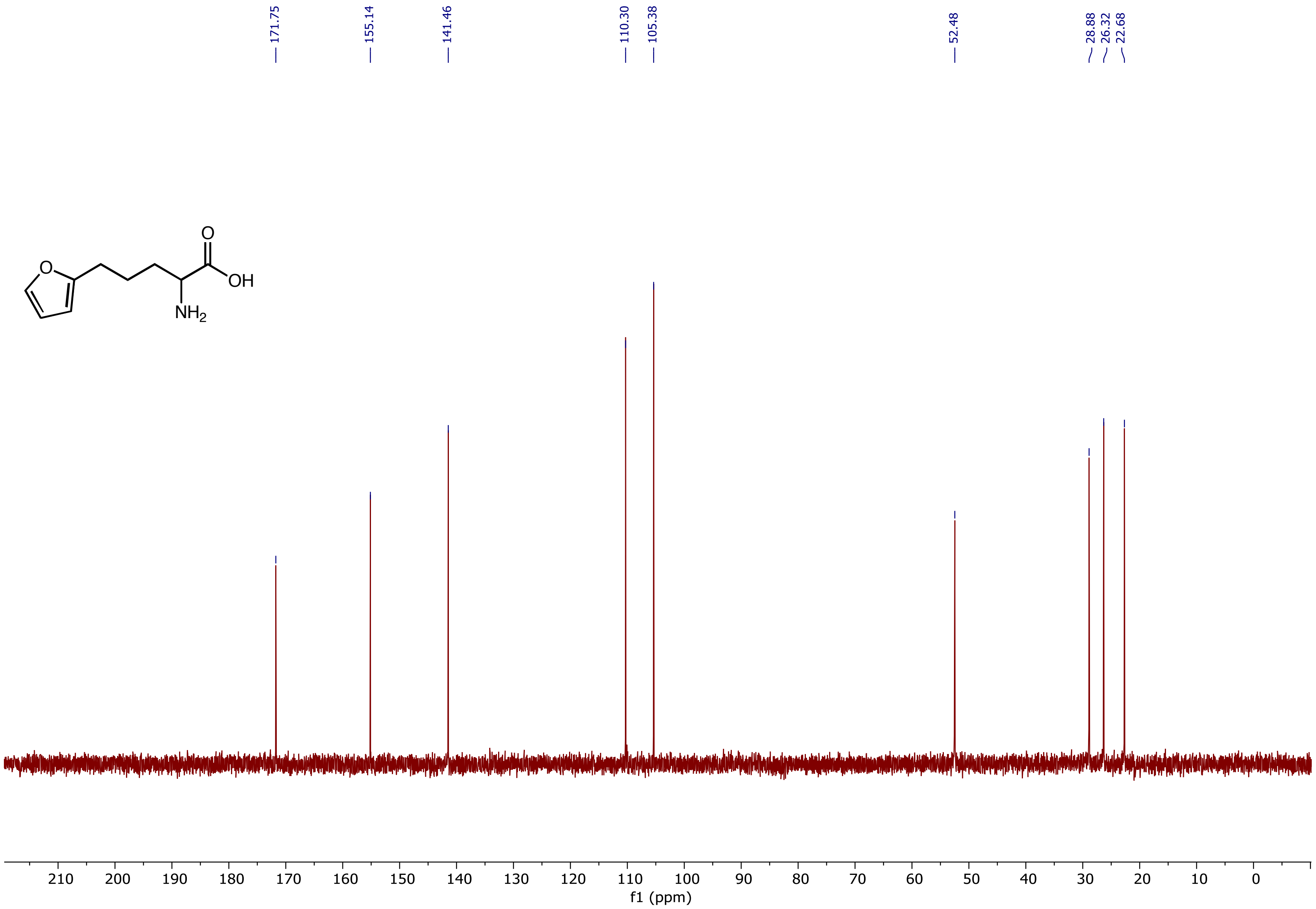

[1]H NMR (500 MHz, $D_2O$) spectrum of 2-Amino-5-(thiophen-2-yl)pentanoic acid **(3n)**

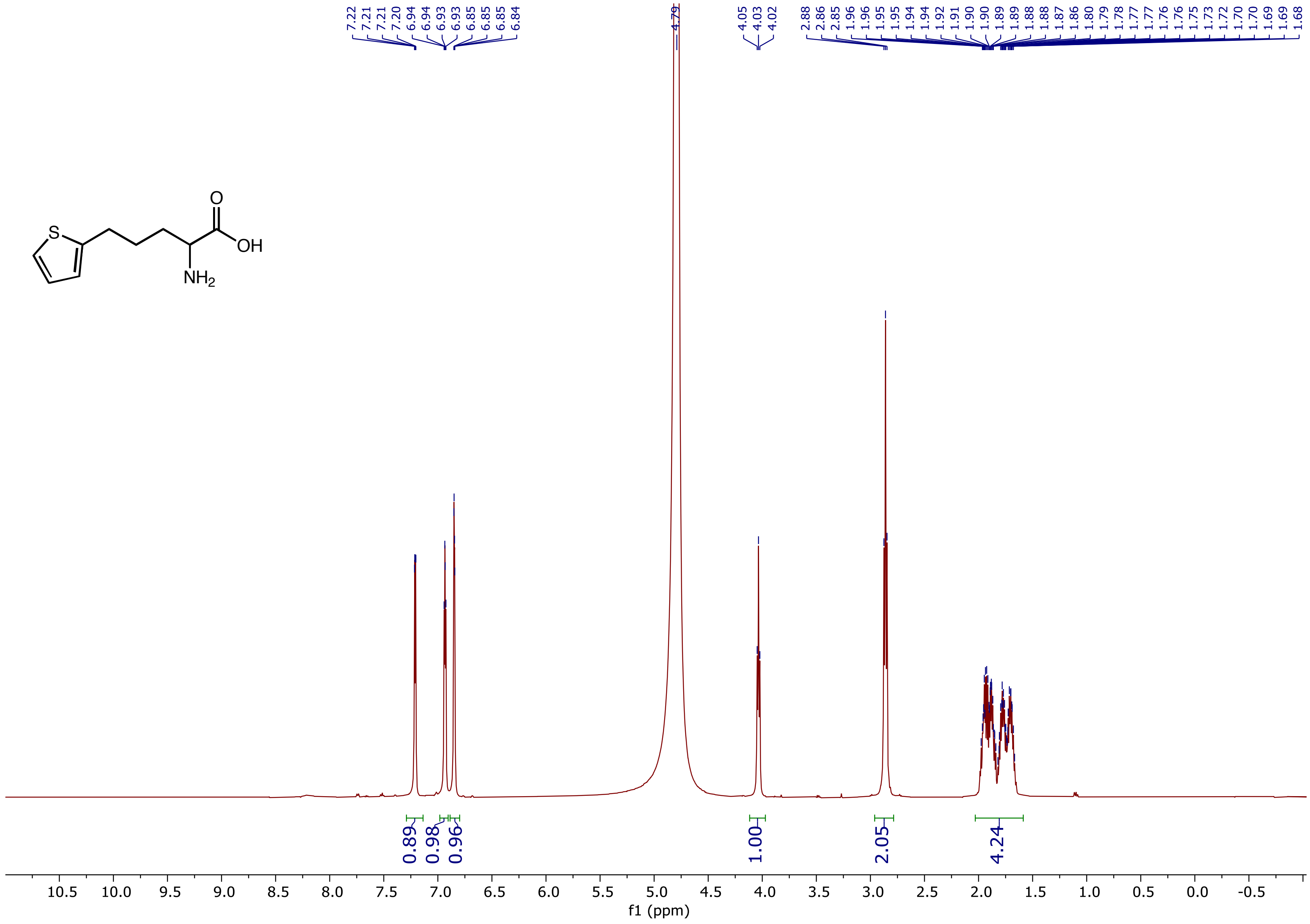

$^{13}C$ NMR (126 MHz, $D_2O$) spectrum of 2-Amino-5-(thiophen-2-yl)pentanoic acid **(3n)**

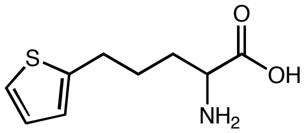

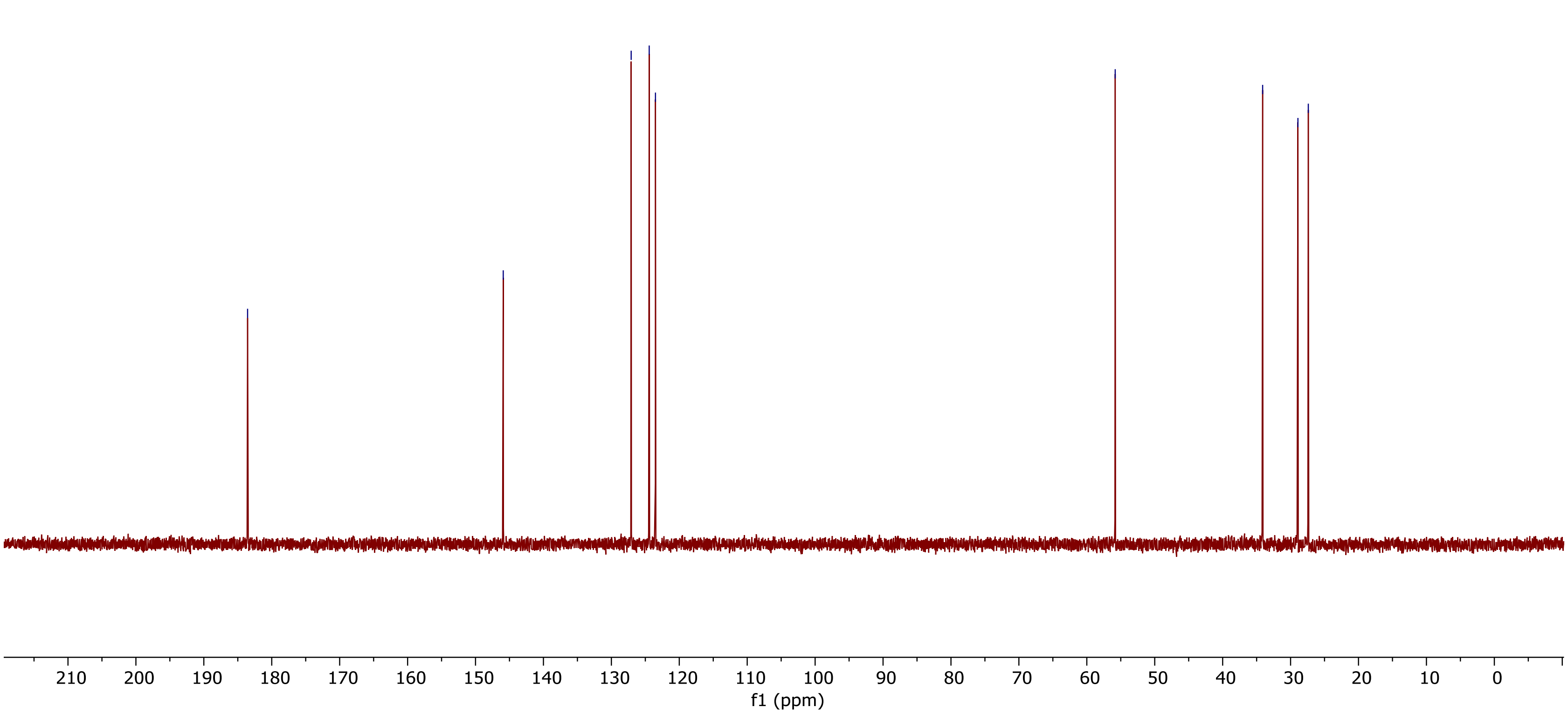

$^1H$ NMR (400 MHz, $D_2O$) spectrum of 2-Amino-5-(6-methoxypyridin-3-yl)pentanoic acid **(3o)**

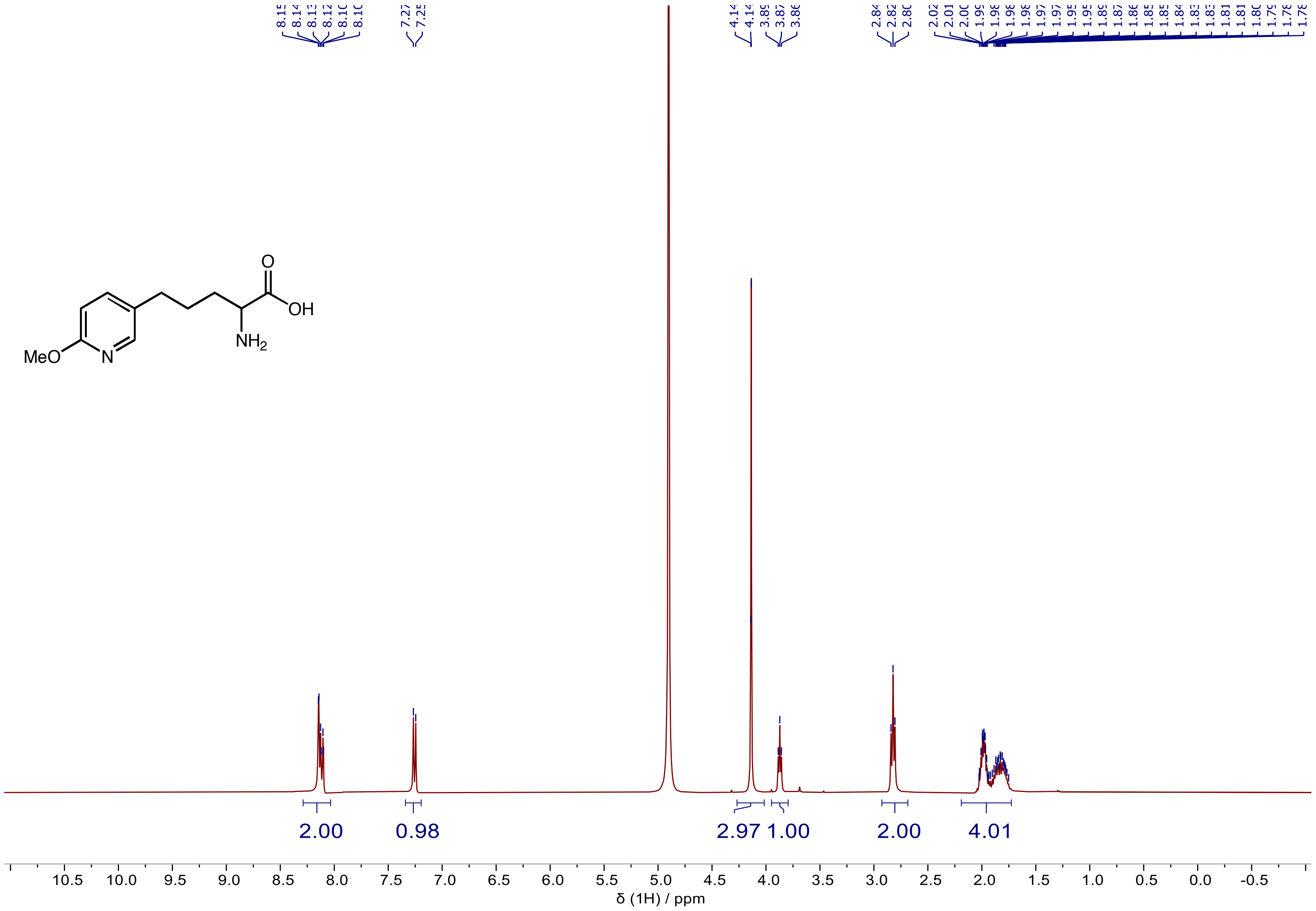

$^{13}C$ NMR (101 MHz, $D_2O$) spectrum of 2-Amino-5-(6-methoxypyridin-3-yl)pentanoic acid **(3o)**

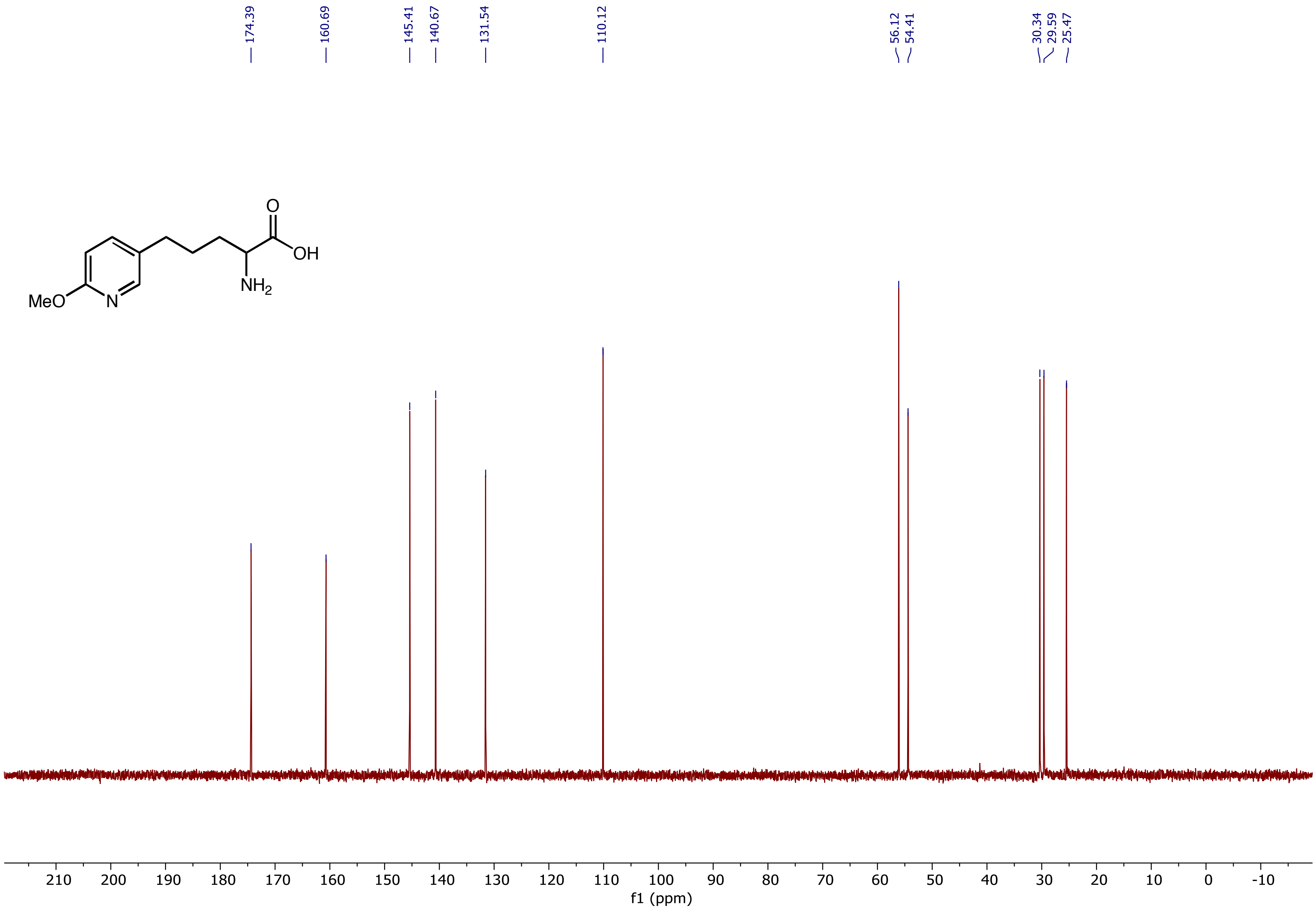

$^{1}$H NMR (400 MHz, $D_2O$) spectrum of 2-Amino-6-phenylhexanoic acid **(3p)**

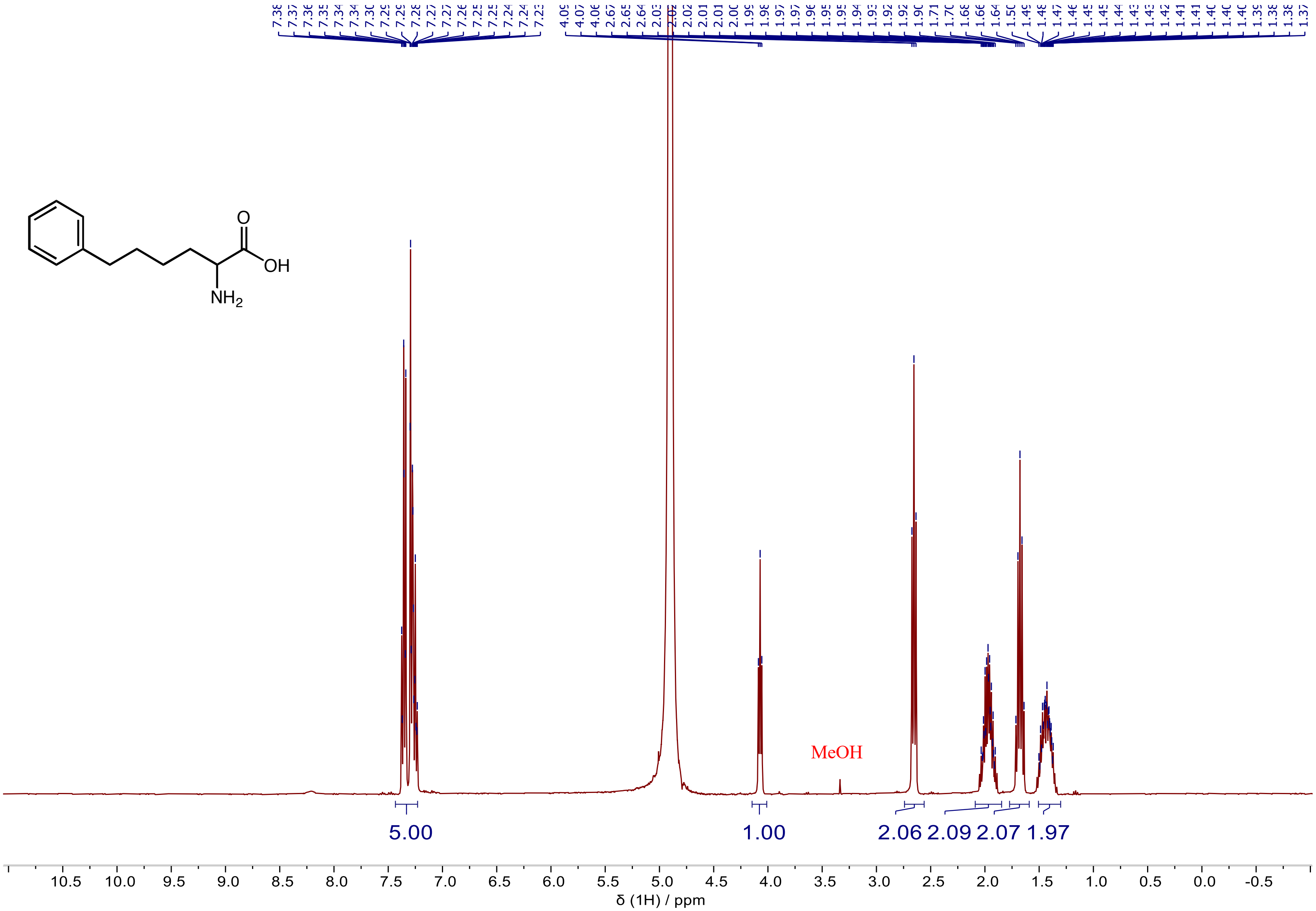

$^{1}H$ NMR (400 MHz, $D_2O$) spectrum of 2-Amino-6-phenylhexanoic acid **(3p)**

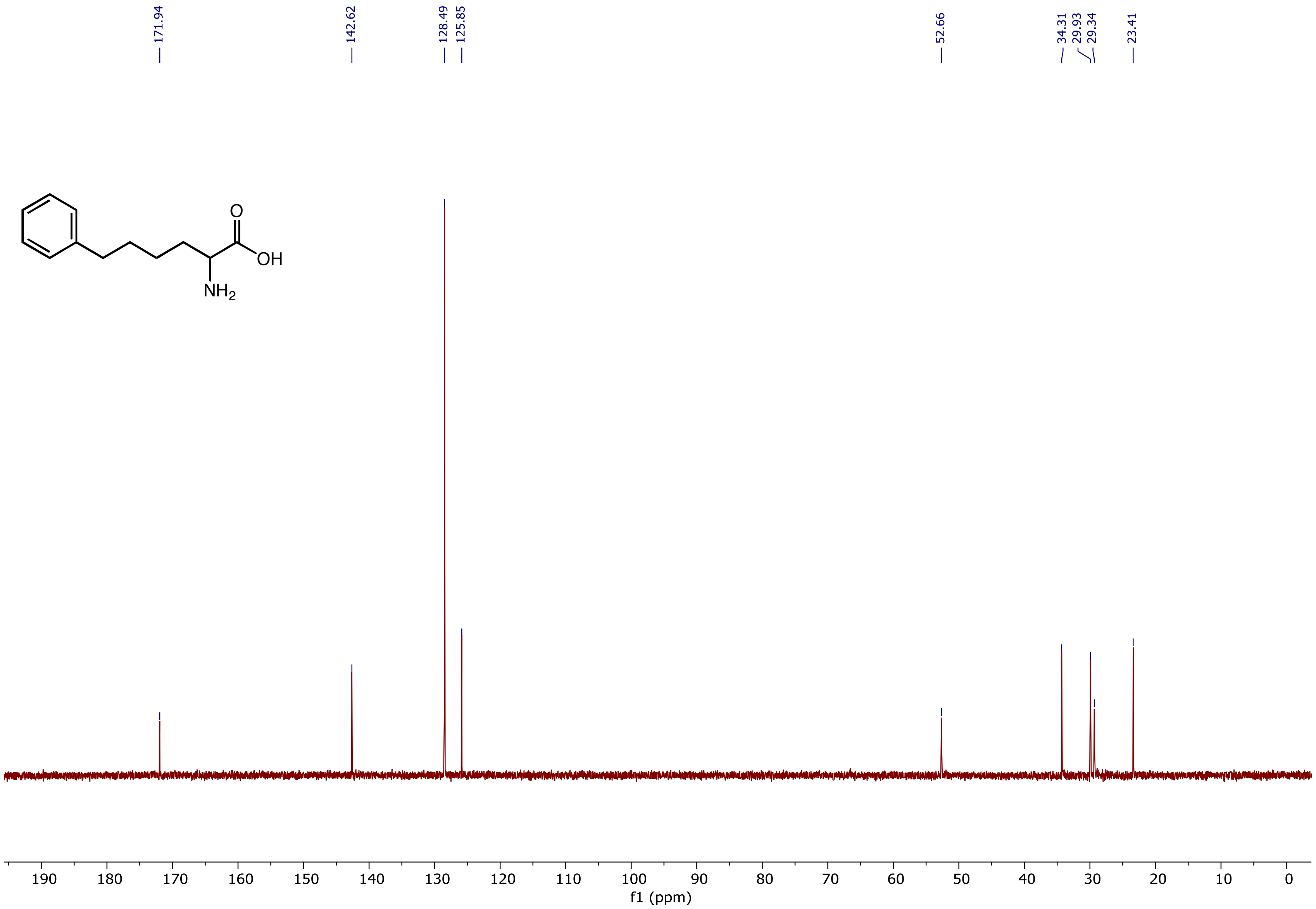

[1]H NMR (500 MHz, $D_2O$) spectrum of 2-Amino-3-(2,3-dihydro-1H-inden-2-yl)propanoic acid **(3q)**

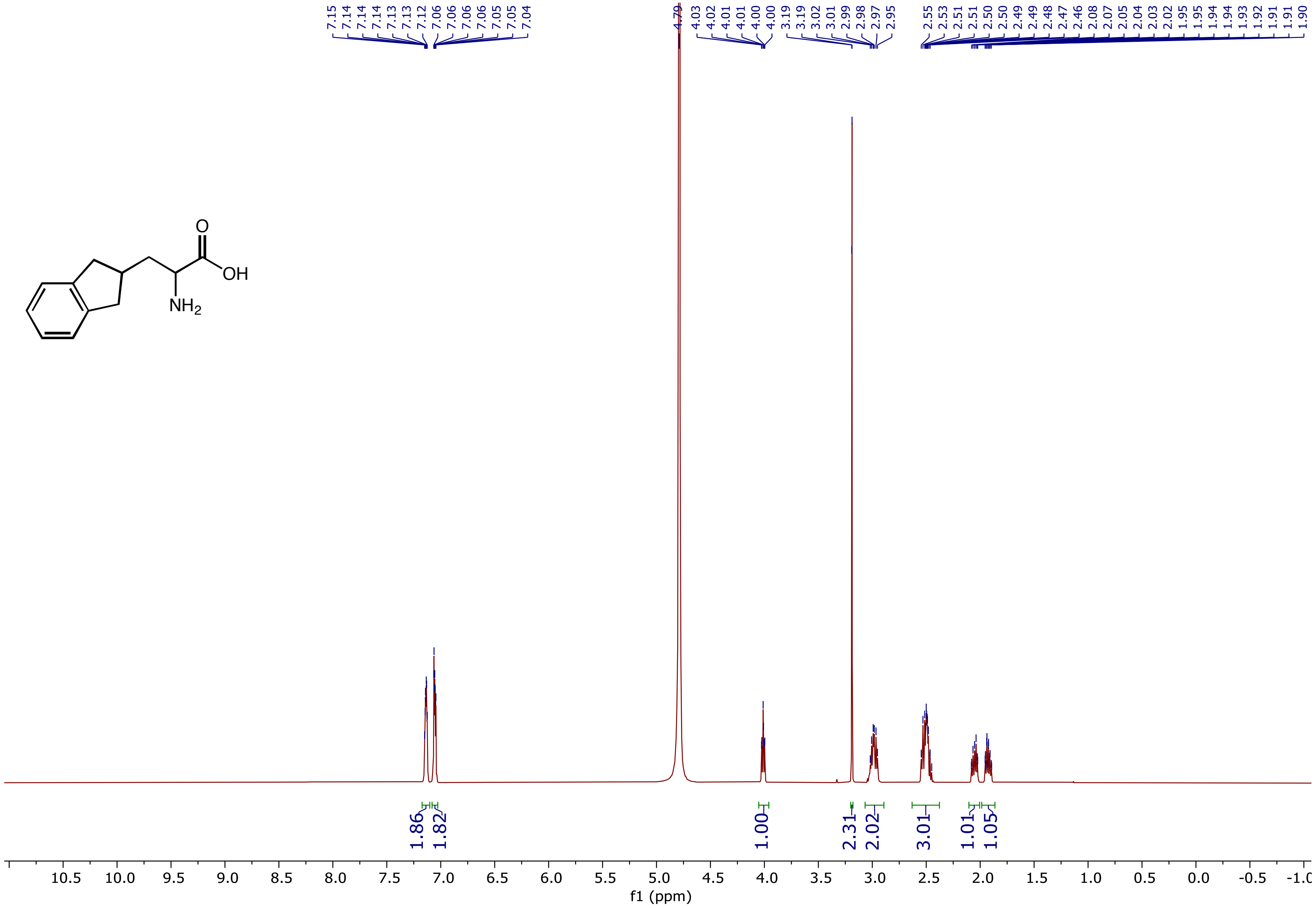

$^{13}C$ NMR (126 MHz, $D_2O$) spectrum of 2-Amino-3-(2,3-dihydro-1H-inden-2-yl)propanoic acid **(3q)**

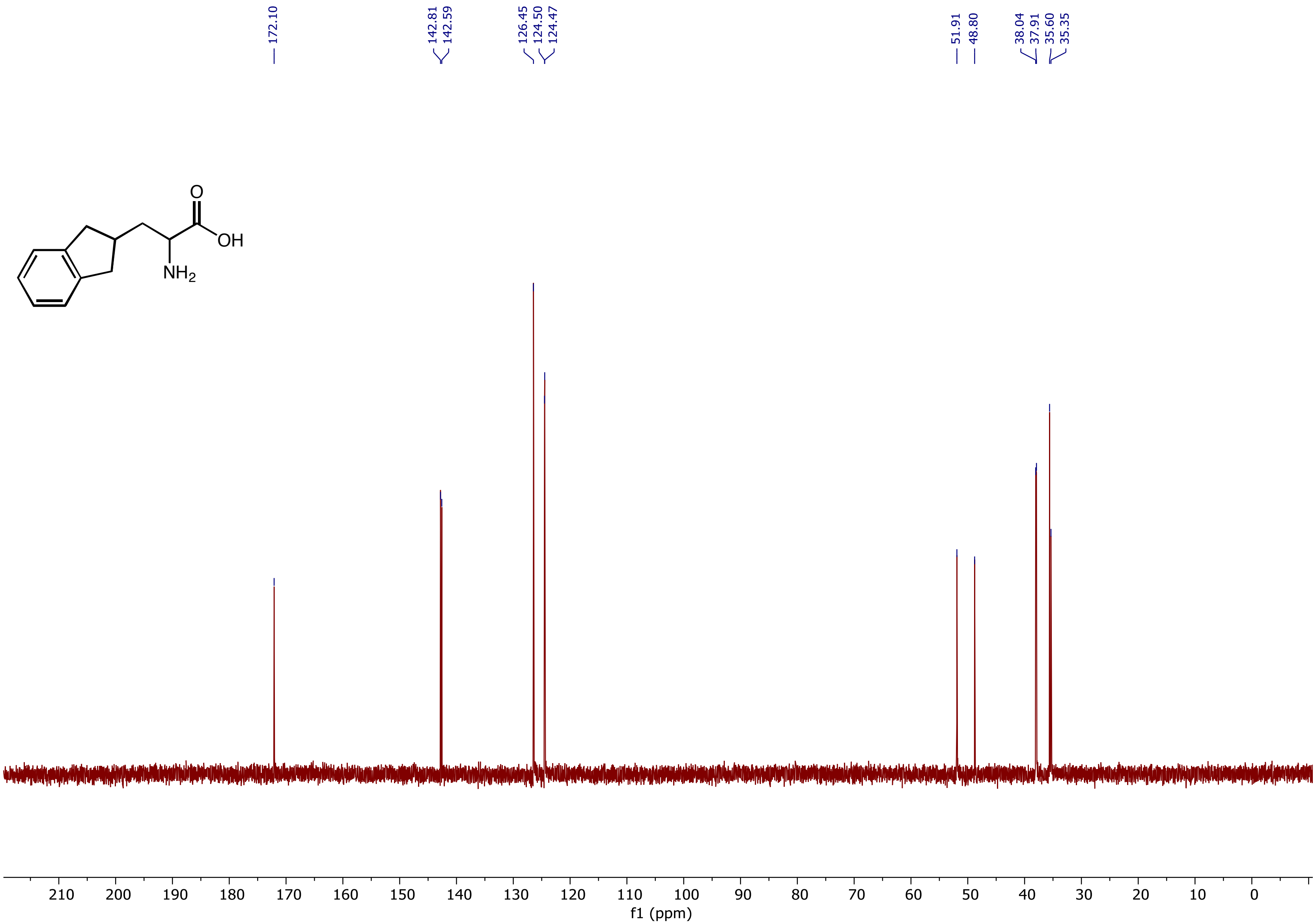

$^{1}H$ NMR (400 MHz, $D_2O$) spectrum of 2-Amino-5-phenylhexanoic acid (**3r**)

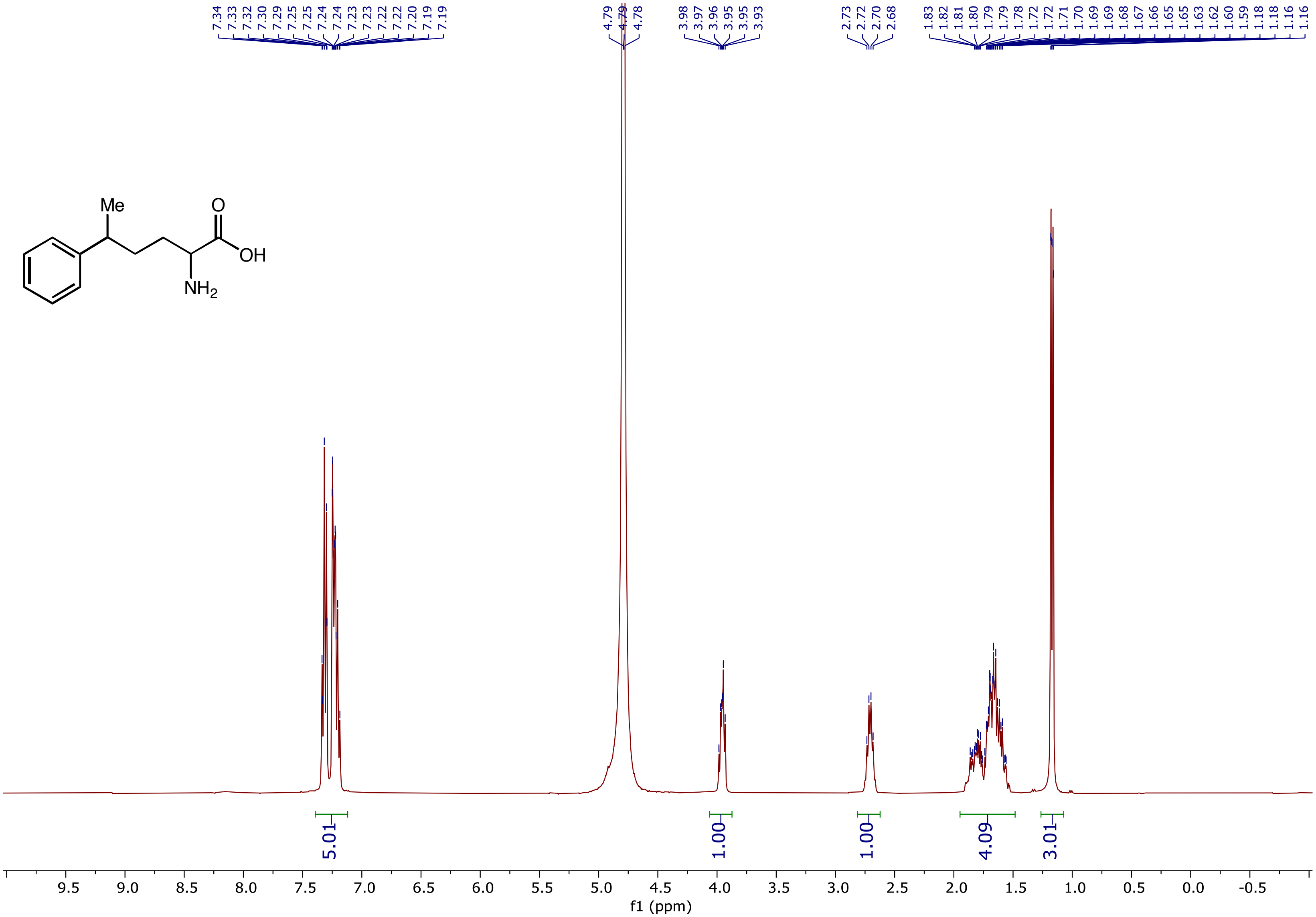

$^{13}C$ NMR (101 MHz, $D_2O$) spectrum of 2-Amino-5-phenylhexanoic acid (**3r**)

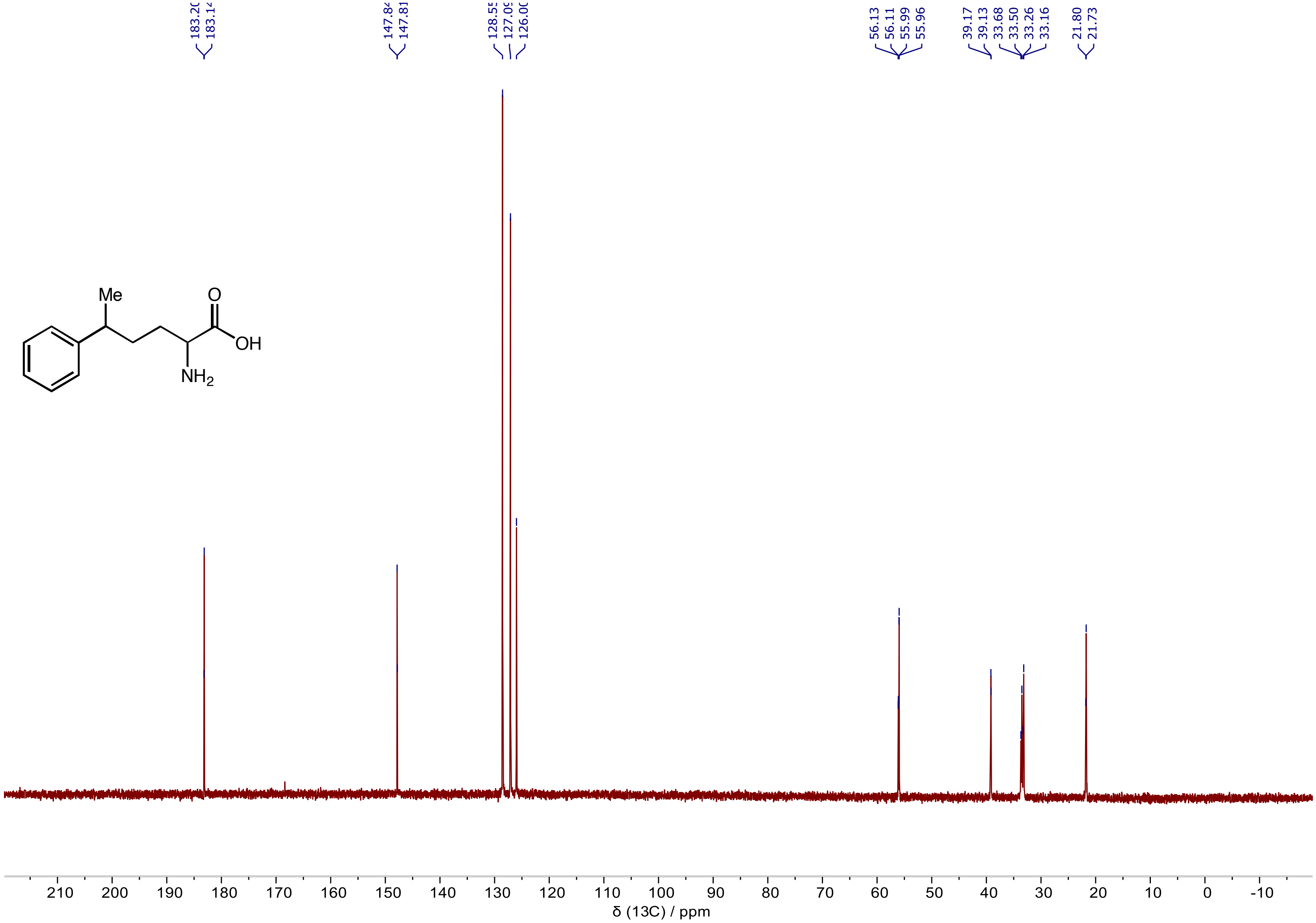

$^{1}H$ NMR (400 MHz, $D_2O$) spectrum of 2-amino-4-methyl-5-phenylpentanoic acid (**3s**)

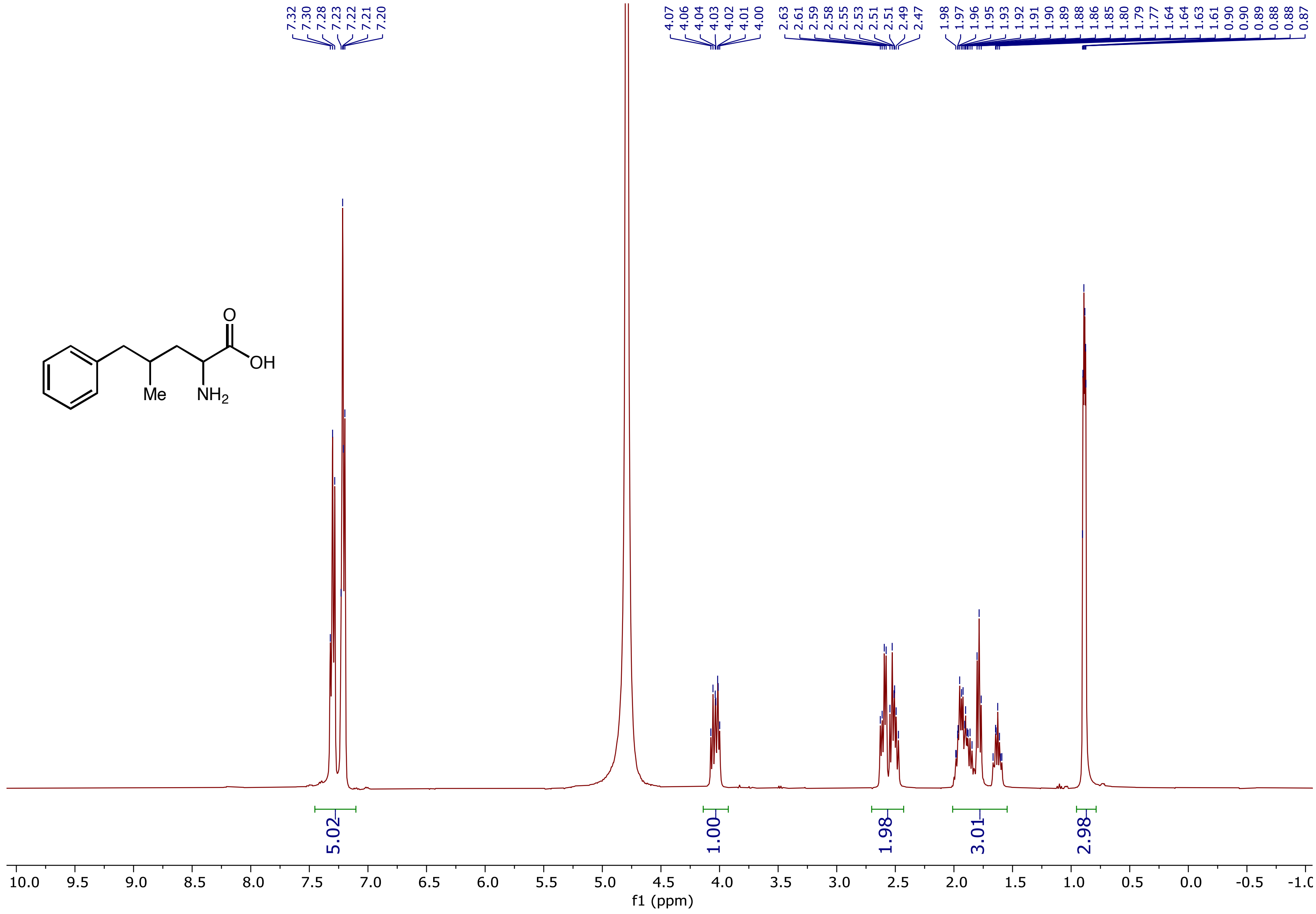

$^{13}C$ NMR (101 MHz, $D_2O$) spectrum of 2-amino-4-methyl-5-phenylpentanoic acid (**3s**)

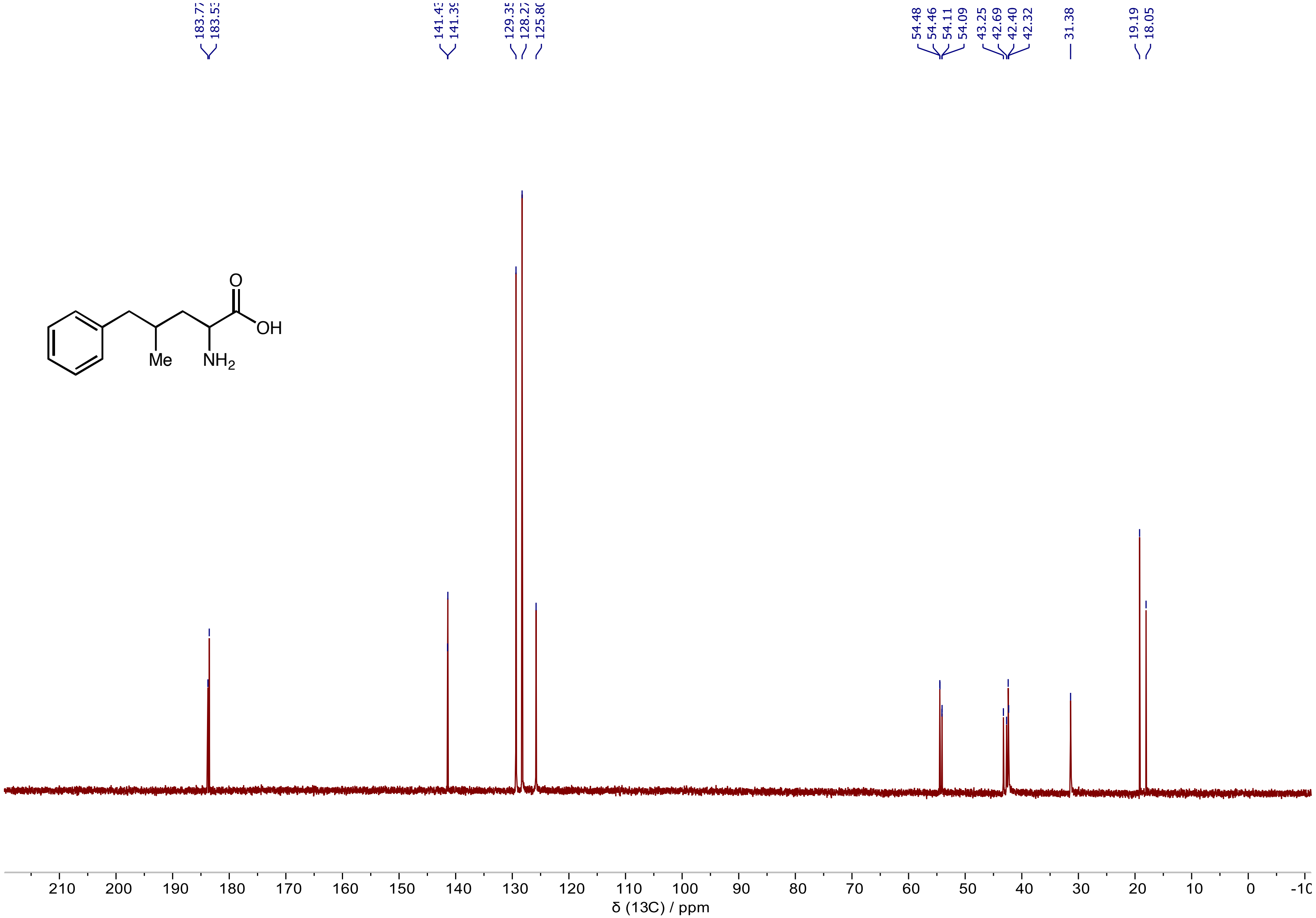

$^{1}H$ NMR (400 MHz, $D_2O$) spectrum of 2-Aminooctanoic acid **(5a)**

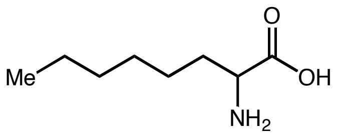

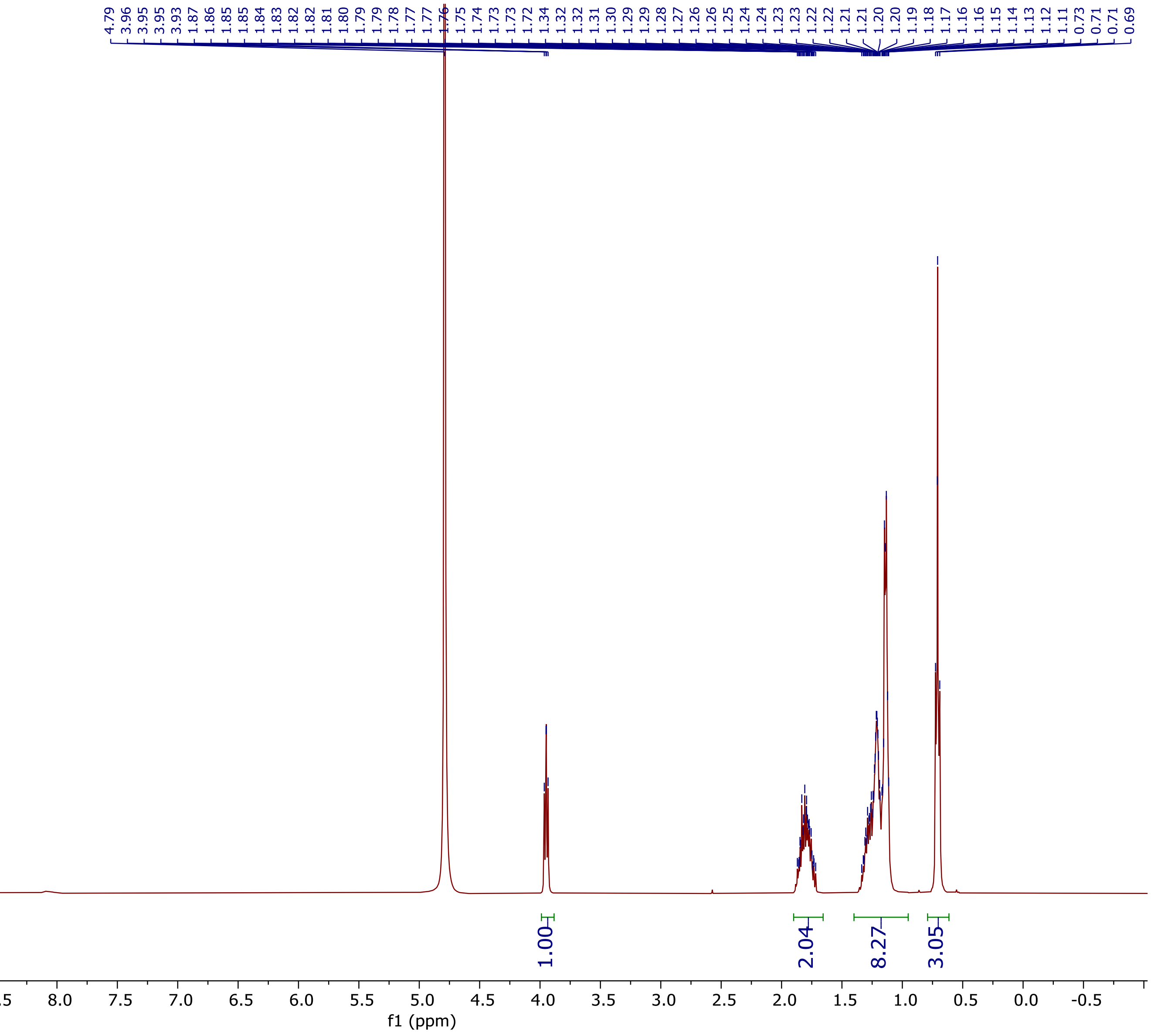

$^{13}C$ NMR (101 MHz, $D_2O$) spectrum of 2-Aminooctanoic acid **(5a)**

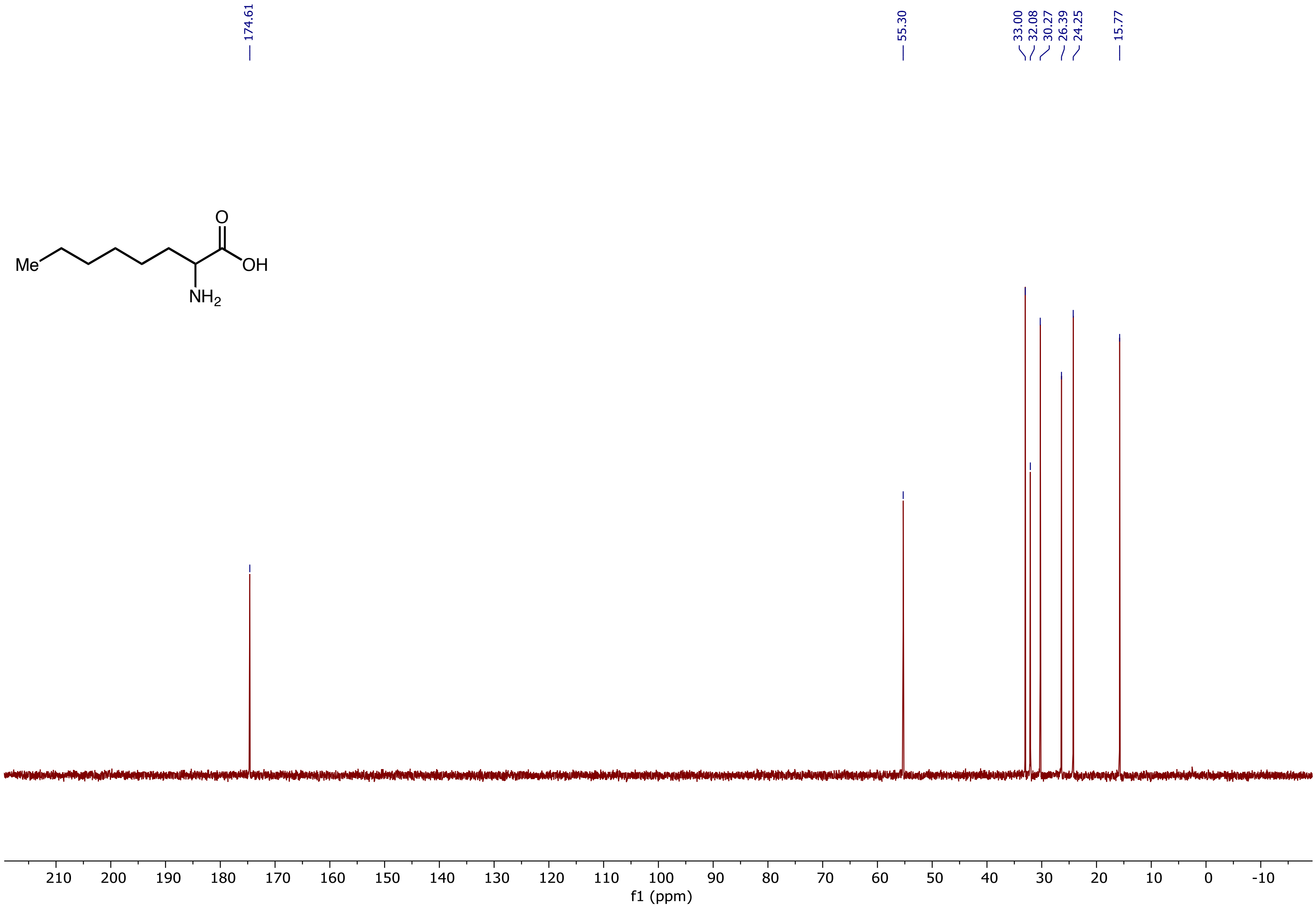

$^{1}H$ NMR (400 MHz, $D_2O$) spectrum of 2-Amino-7-fluoroheptanoic acid **(5b)**

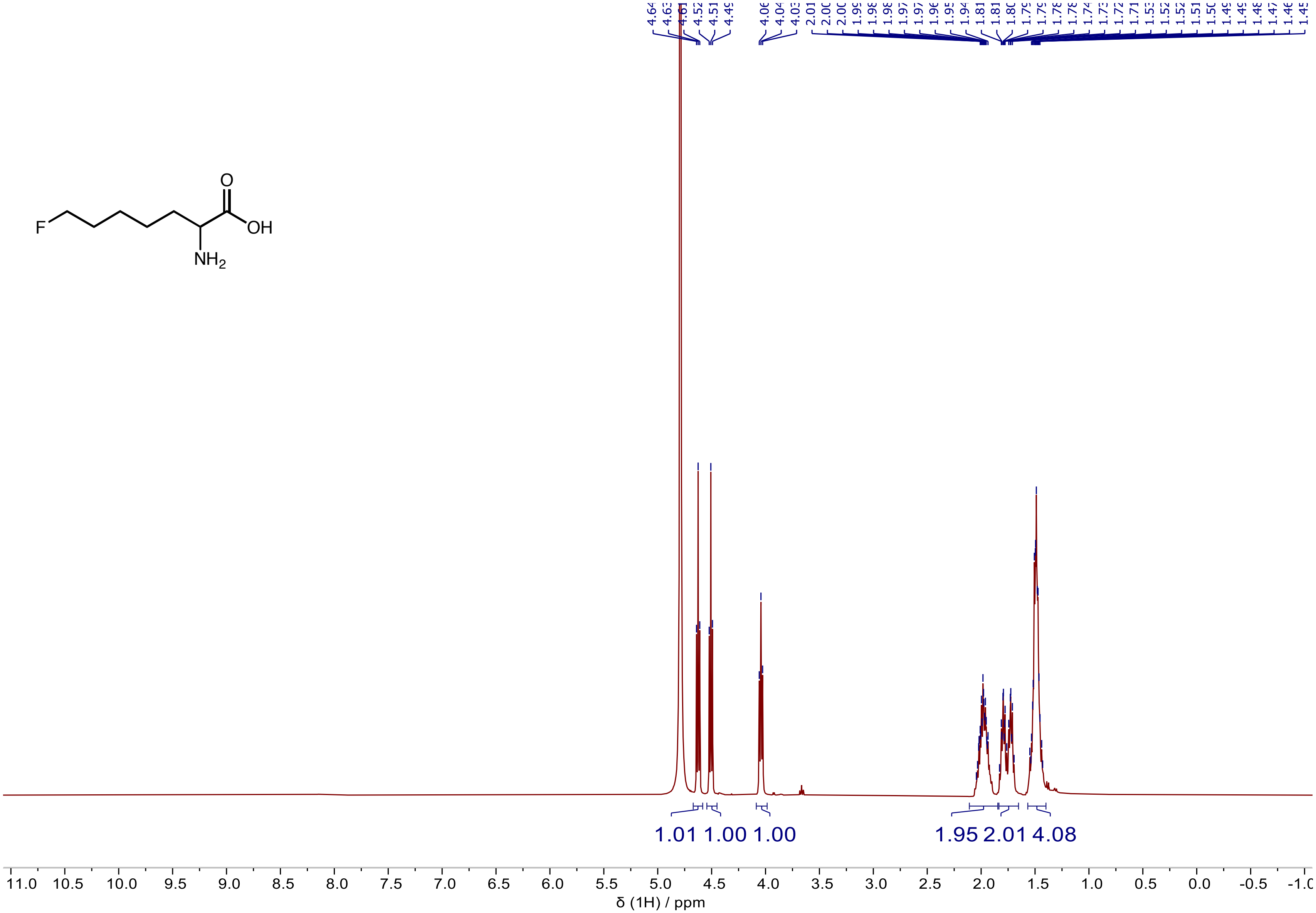

$^{13}$C NMR (101 MHz, $D_2O$) spectrum of 2-Amino-7-fluoroheptanoic acid **(5b)**

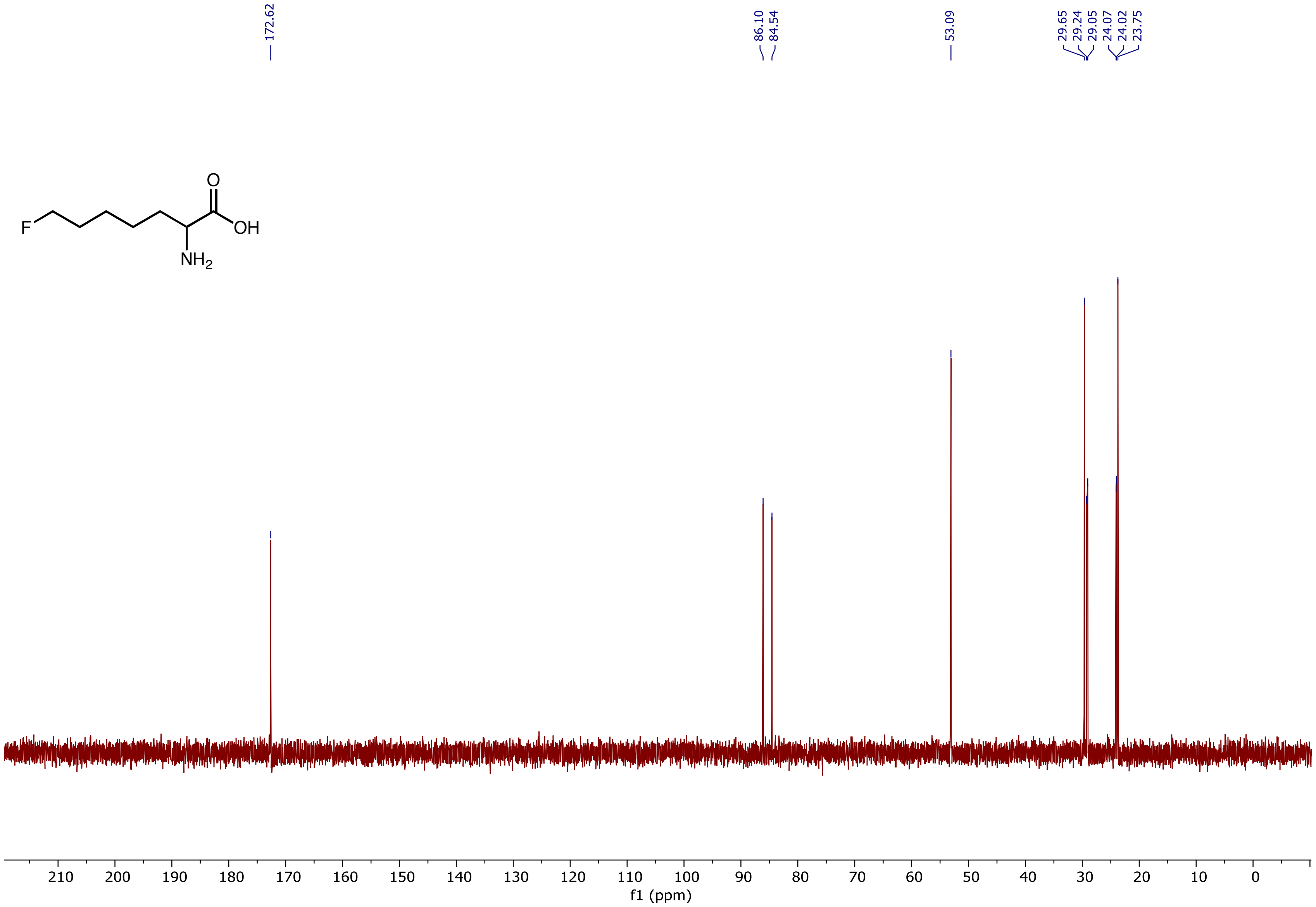

[19]F NMR (376 MHz, $D_2O$) spectrum of 2-Amino-7-fluoroheptanoic acid **(5b)**

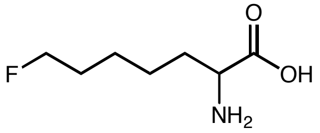

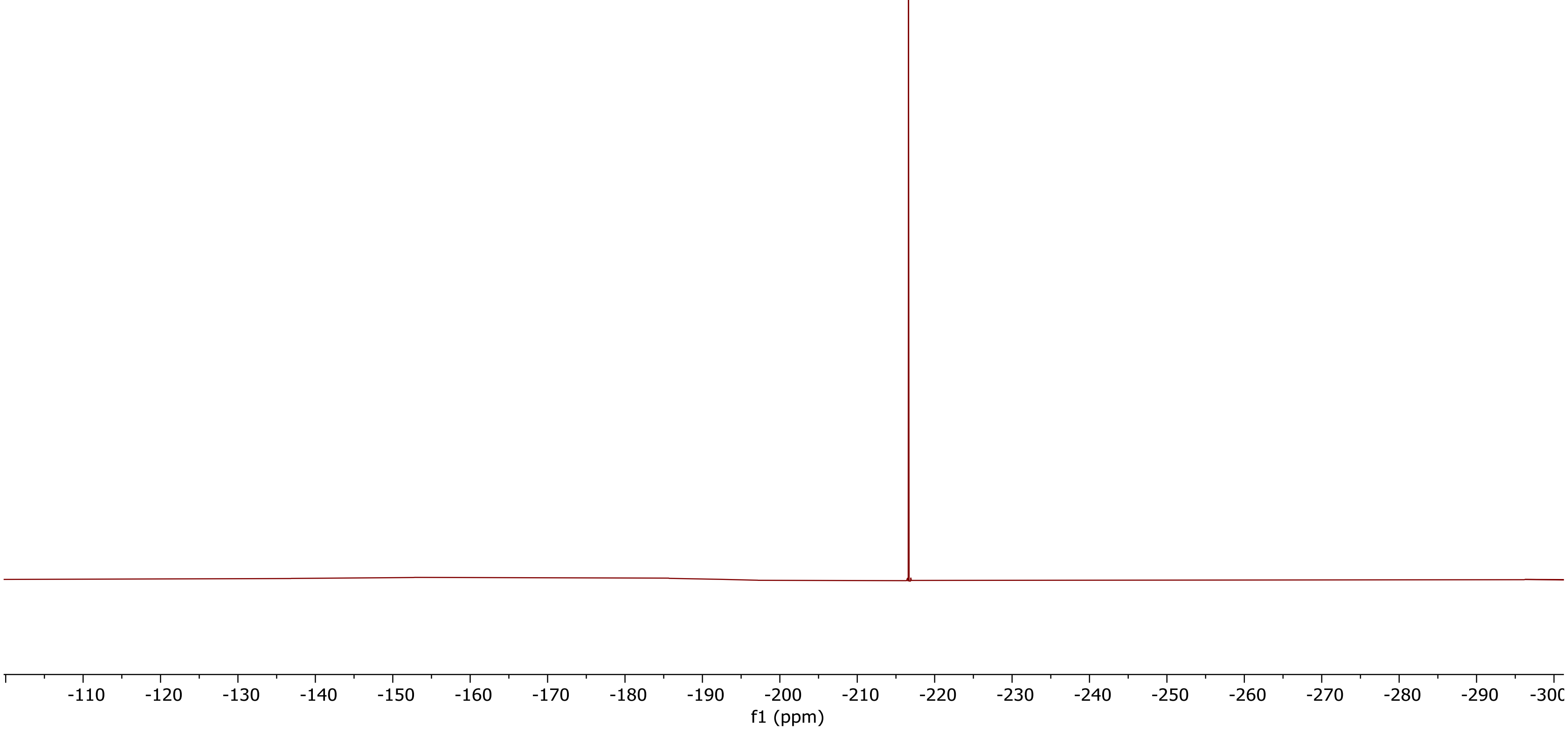

$^1H$ NMR (400 MHz, $D_2O$) spectrum of 2-Amino-8-chlorooctanoic acid **(5c)**

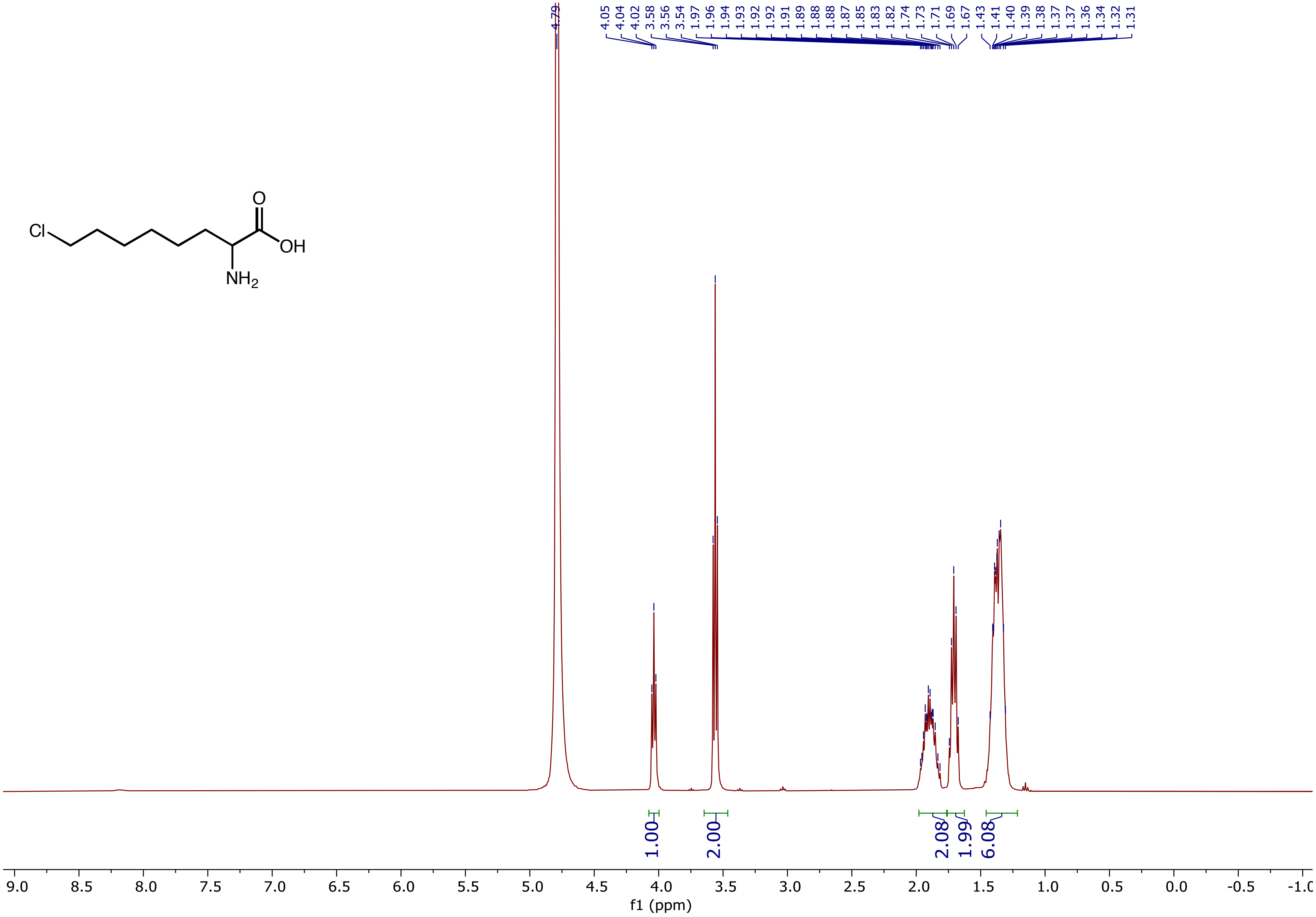

$^{13}$C NMR (101 MHz, $D_2O$) spectrum of 2-Amino-8-chlorooctanoic acid **(5c)**

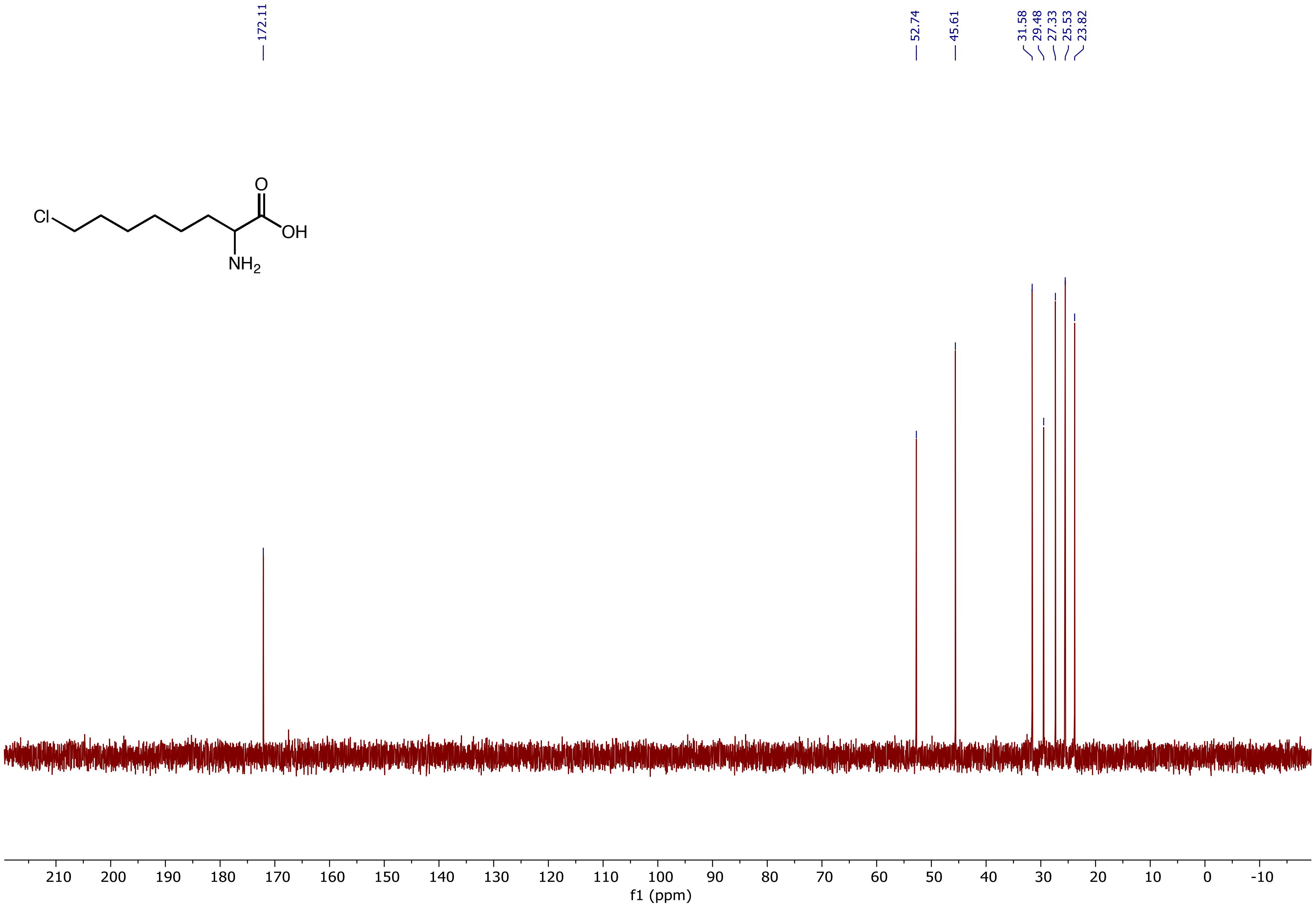

$^1$H NMR (400 MHz, $D_2O$) spectrum of 2-Amino-7-methoxyheptanoic acid **(5d)**

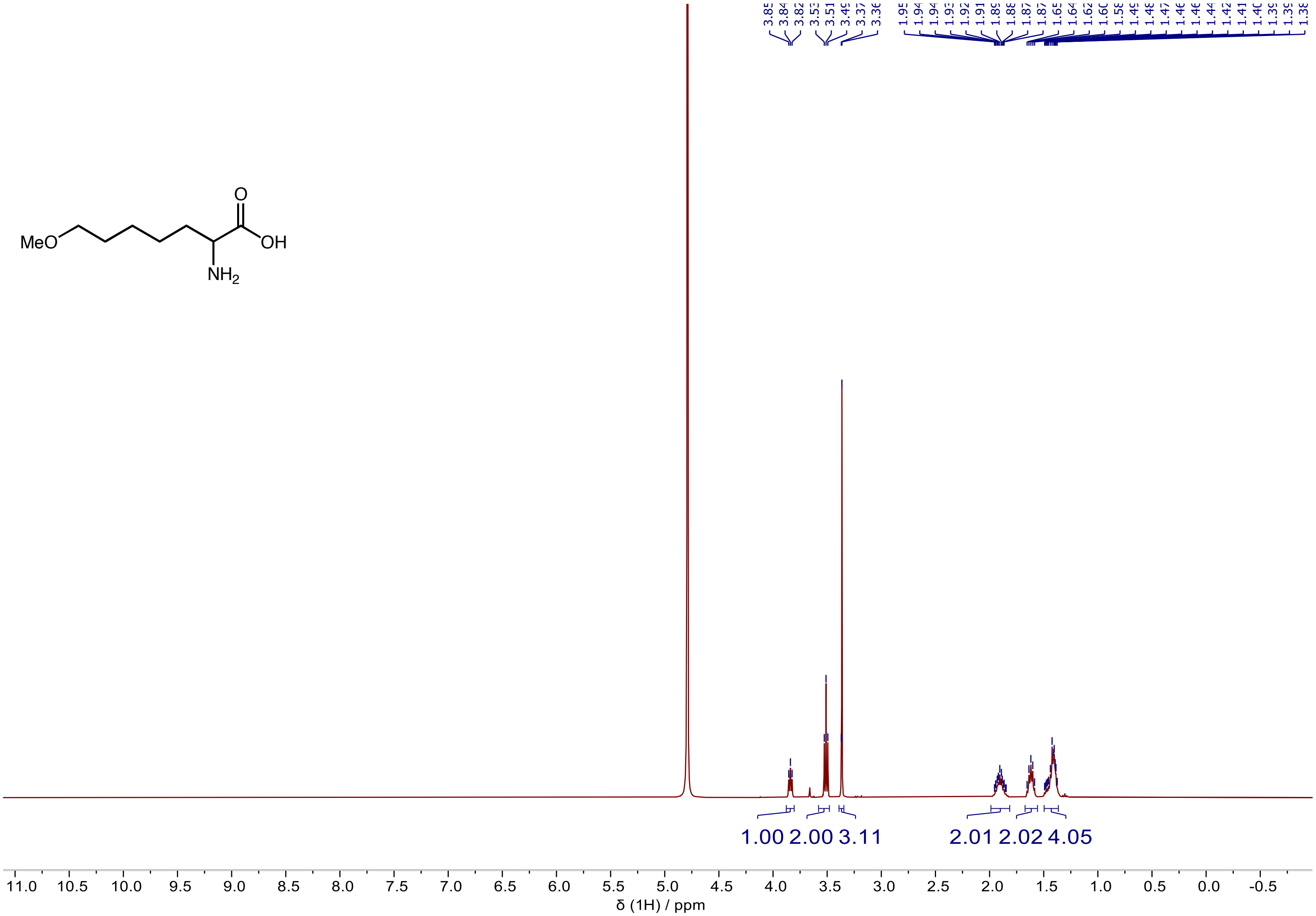

$^{13}C$ NMR (101 MHz, $D_2O$) spectrum of 2-Amino-7-methoxyheptanoic acid **(5d)**

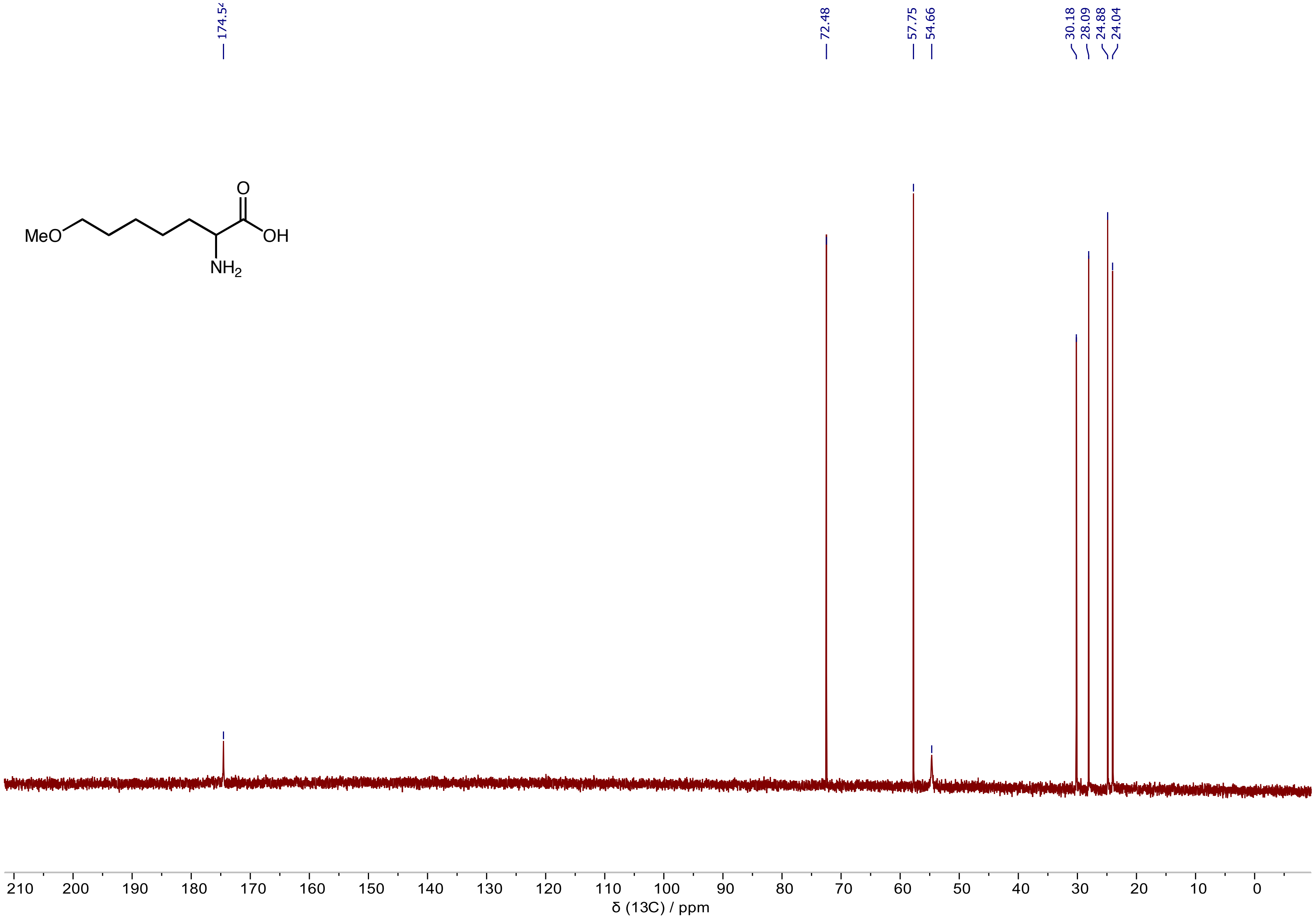

$^1H$ NMR (400 MHz, $D_2O$) spectrum of 2-Amino-7-(methylthio)heptanoic acid **(5e)**

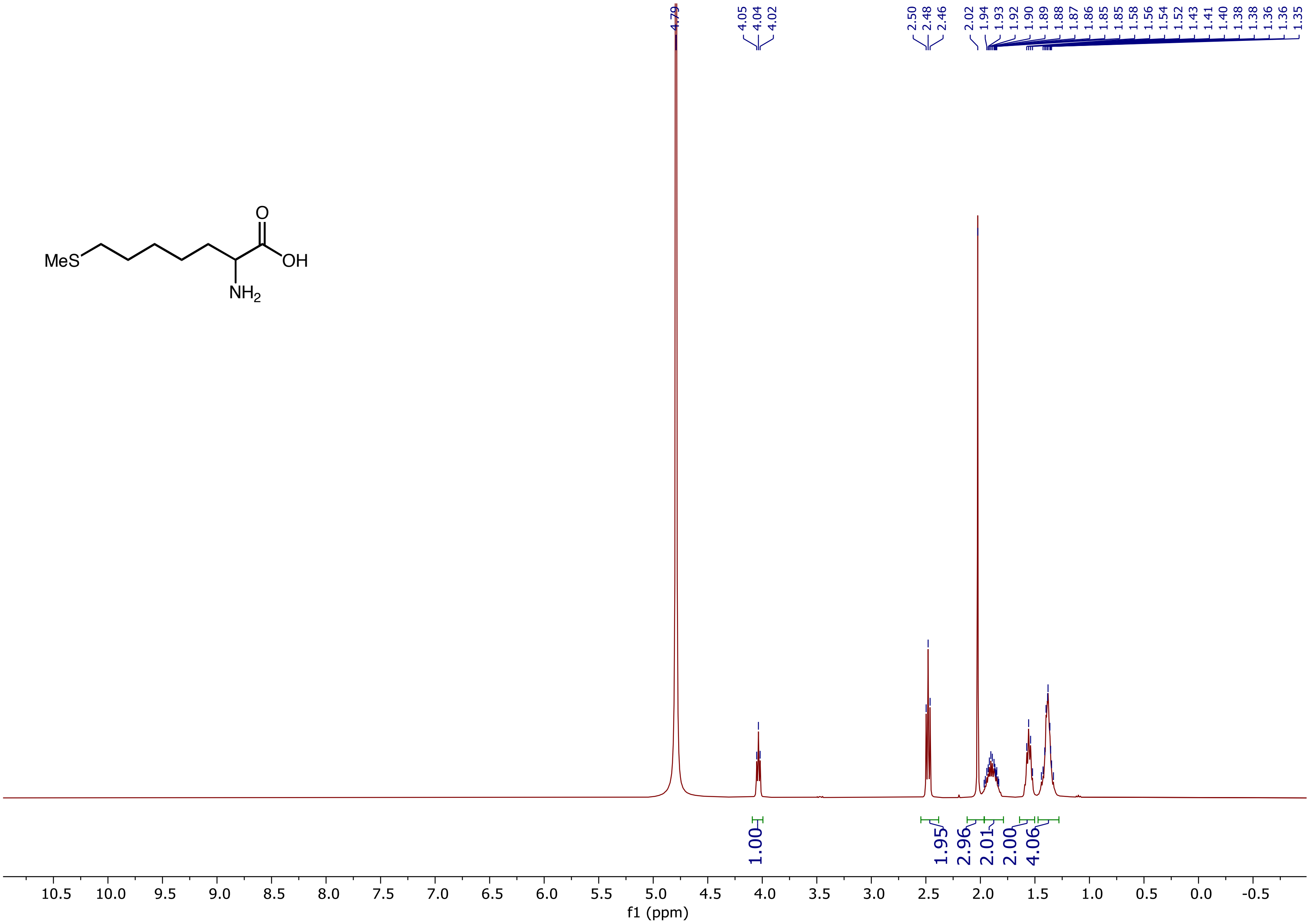

$^{13}C$ NMR (101 MHz, $D_2O$) spectrum of 2-Amino-7-(methylthio)heptanoic acid **(5e)**

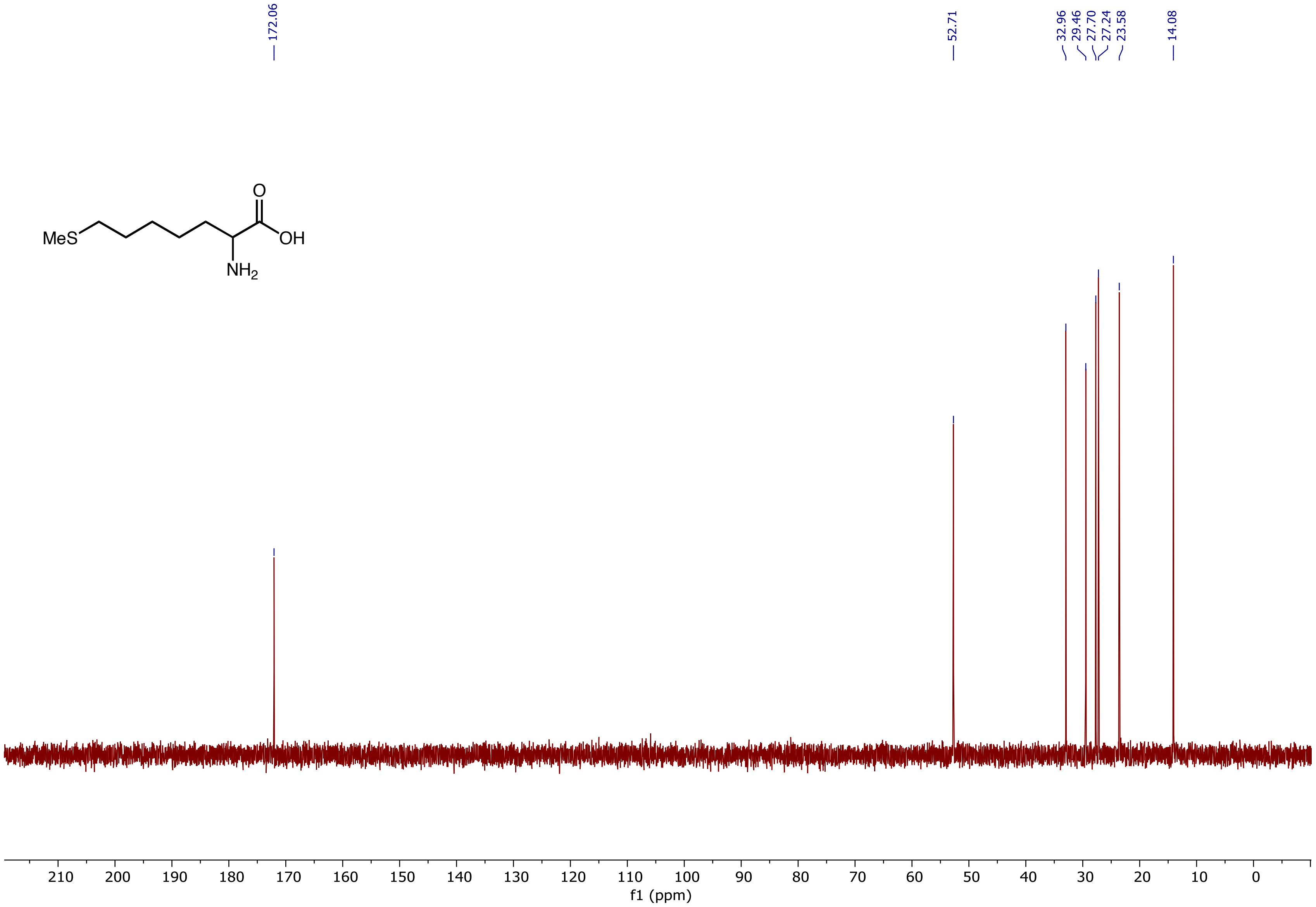

$^{1}H$ NMR (400 MHz, $D_2O$) spectrum of 2-Amino-8-azidooctanoic acid **(5f)**

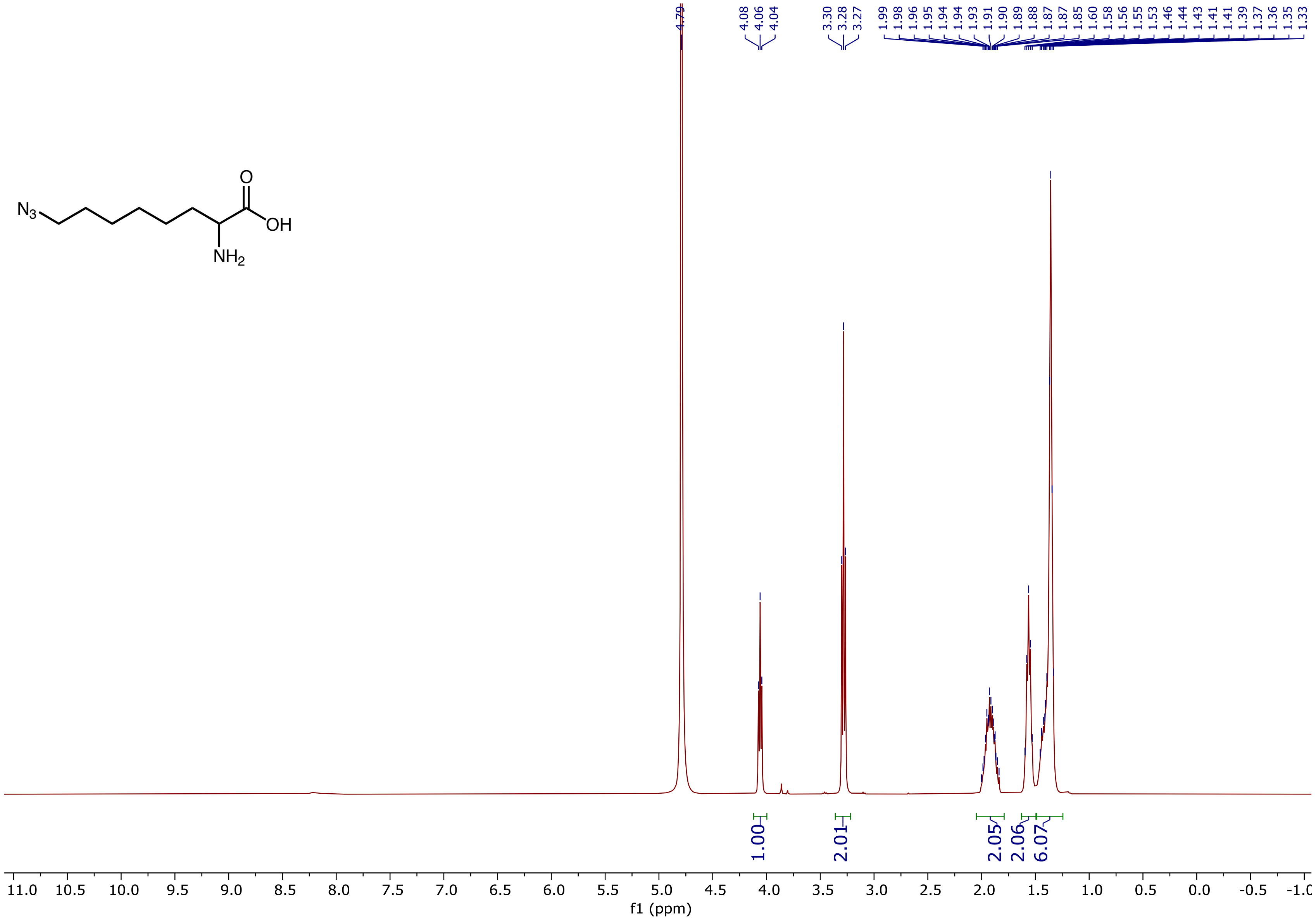

$^{13}$C NMR (101 MHz, $D_2O$) spectrum of 2-Amino-8-azidooctanoic acid **(5f)**

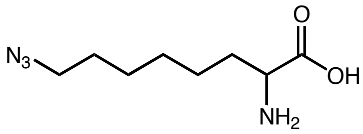

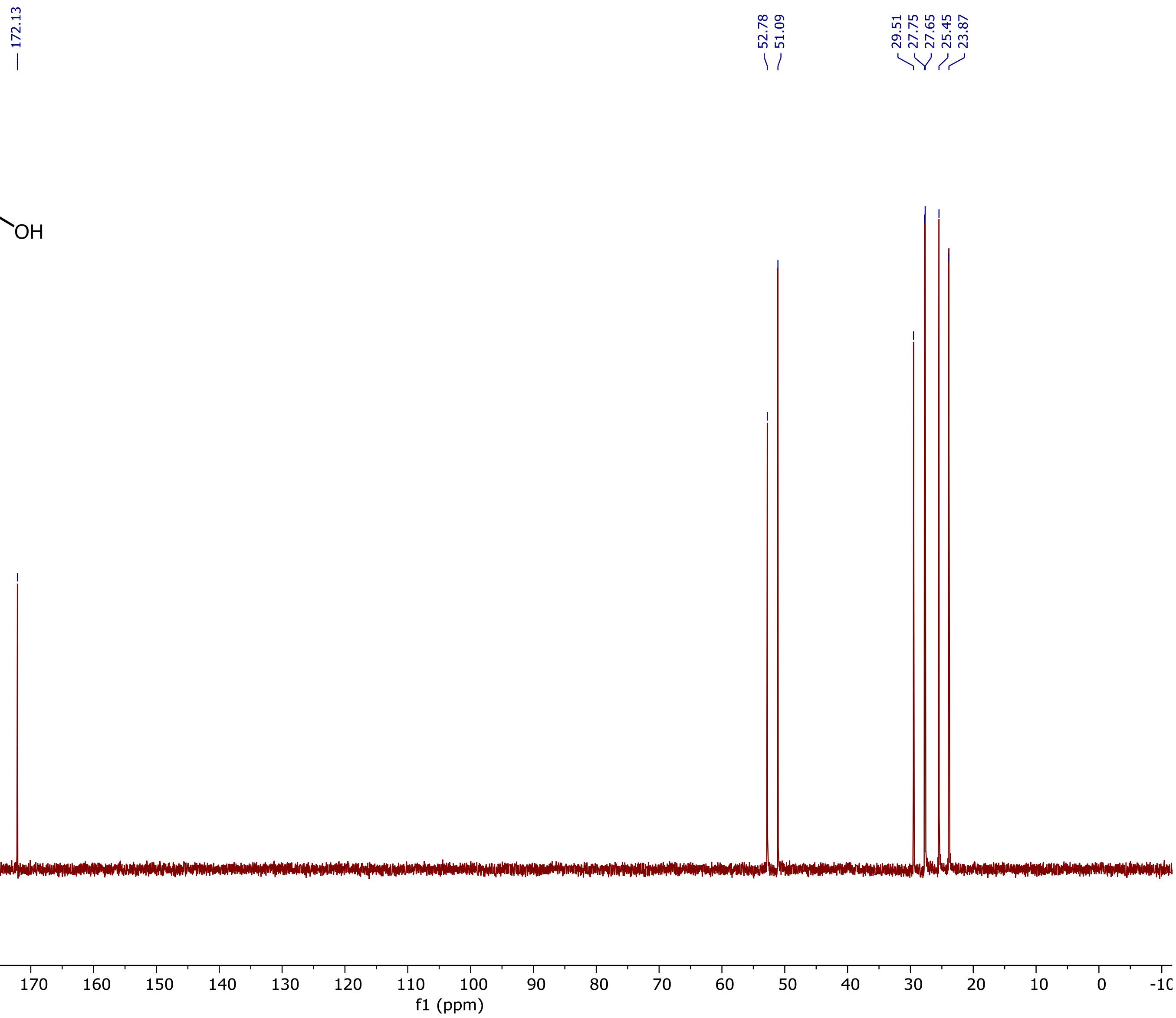

[1]H NMR (400 MHz, $D_2O$) spectrum of 2-Aminooct-7-ynoic acid **(5g)**

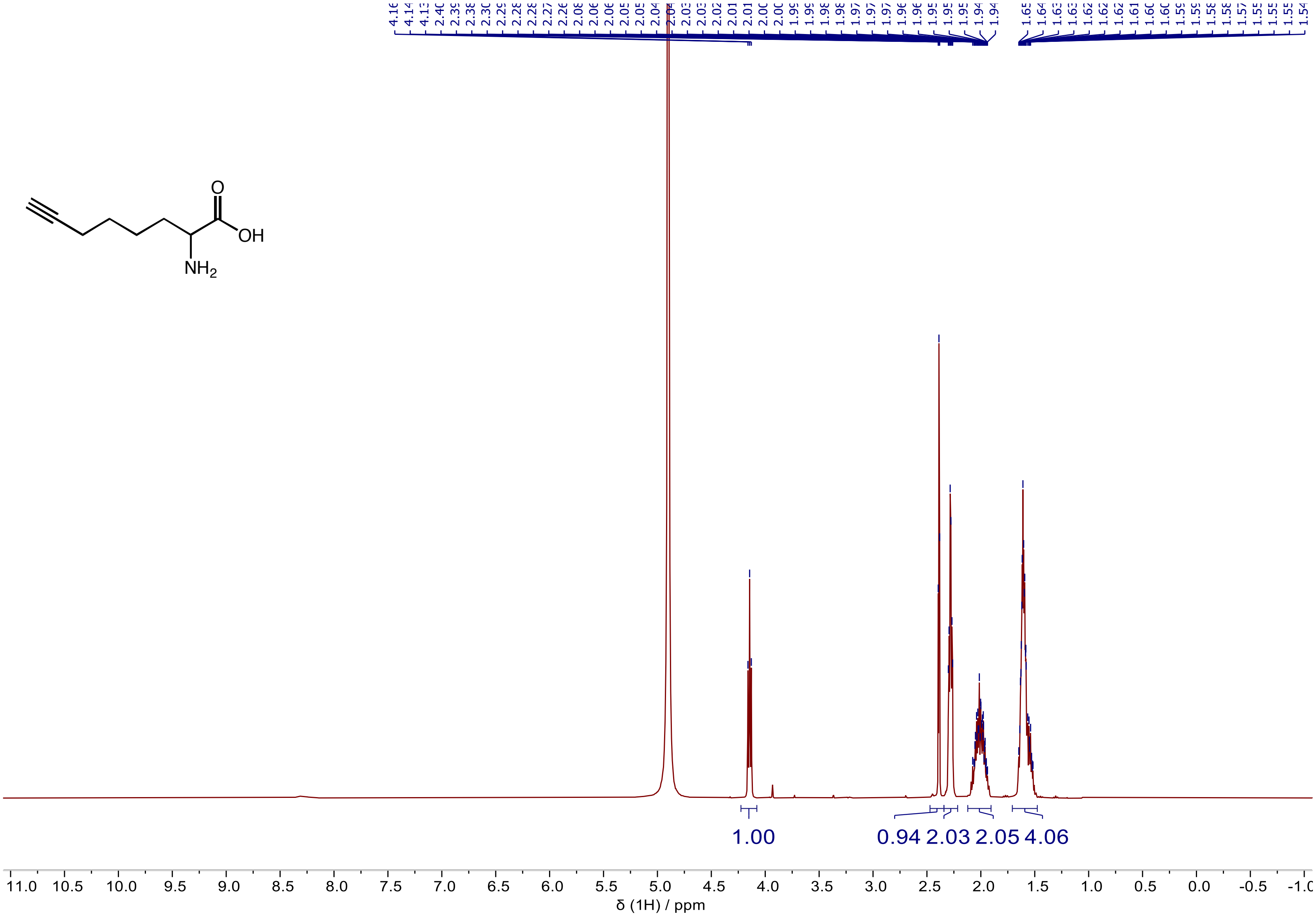

$^{13}C$ NMR (101 MHz, $D_2O$) spectrum of 2-Aminooct-7-ynoic acid **(5g)**

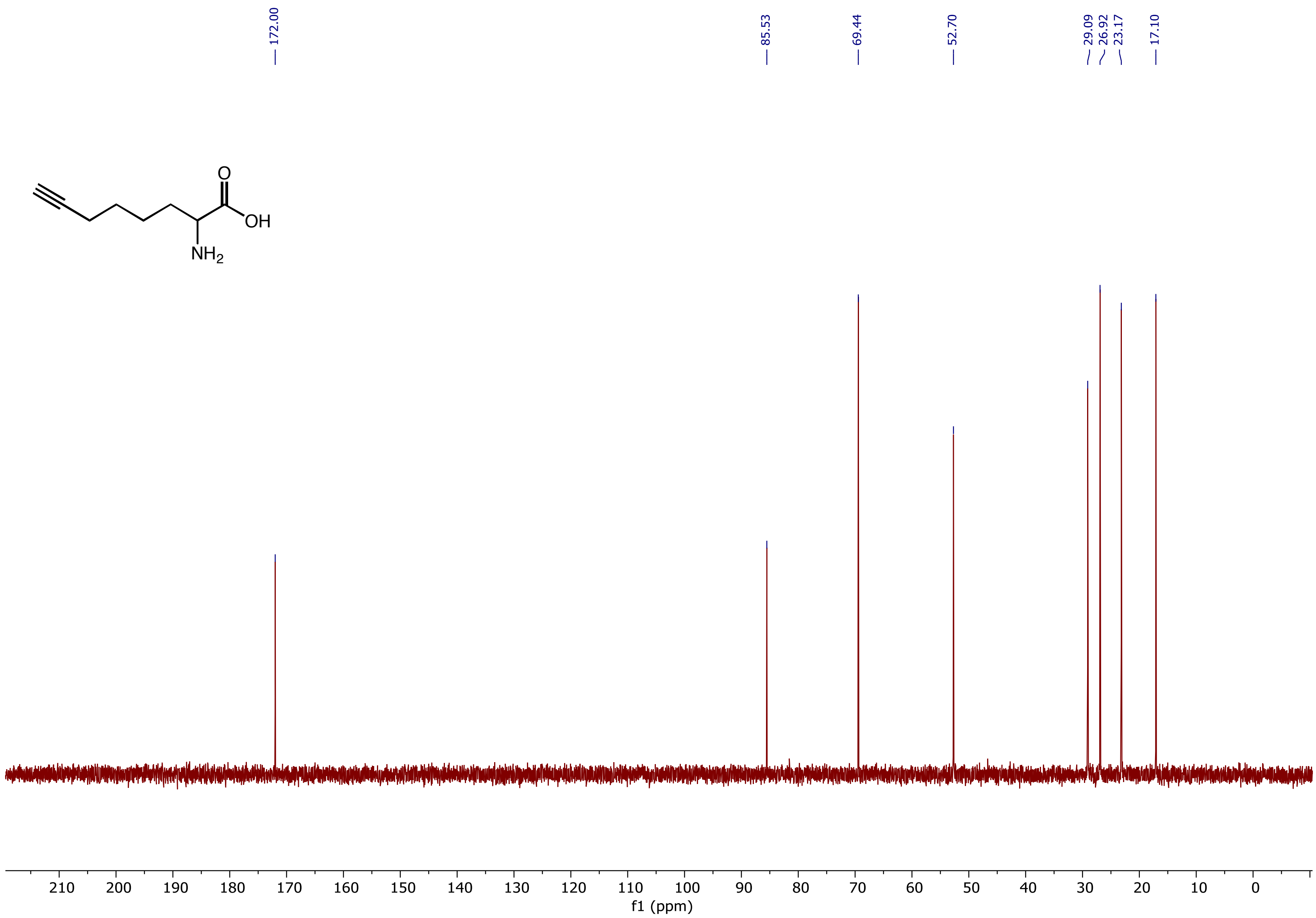

[1]H NMR (400 MHz, $D_2O$) spectrum of 2-Aminohept-6-enoic acid **(5h)**

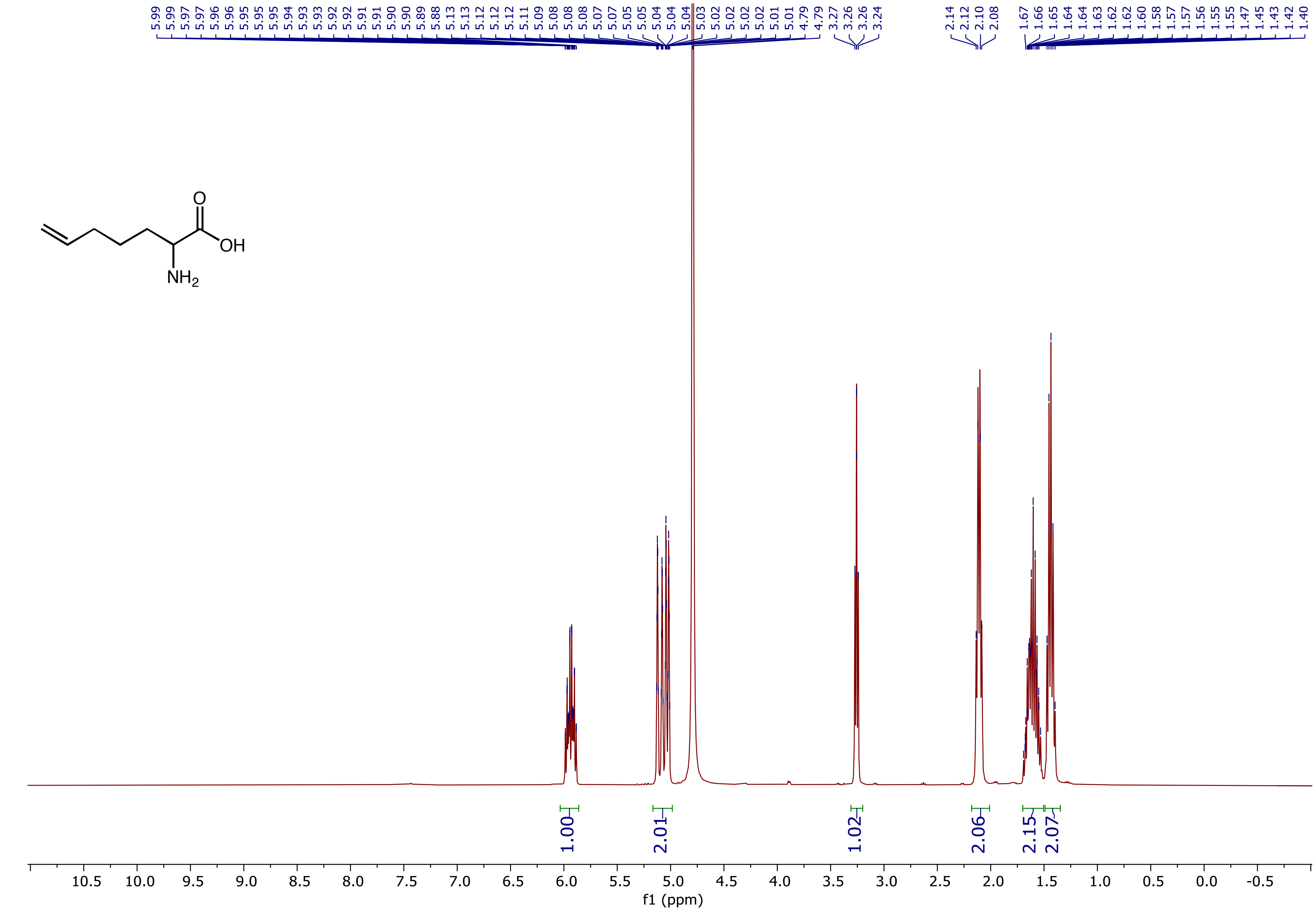

$^{13}C$ NMR (101 MHz, $D_2O$) spectrum of 2-Aminohept-6-enoic acid **(5h)**

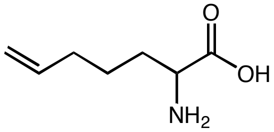

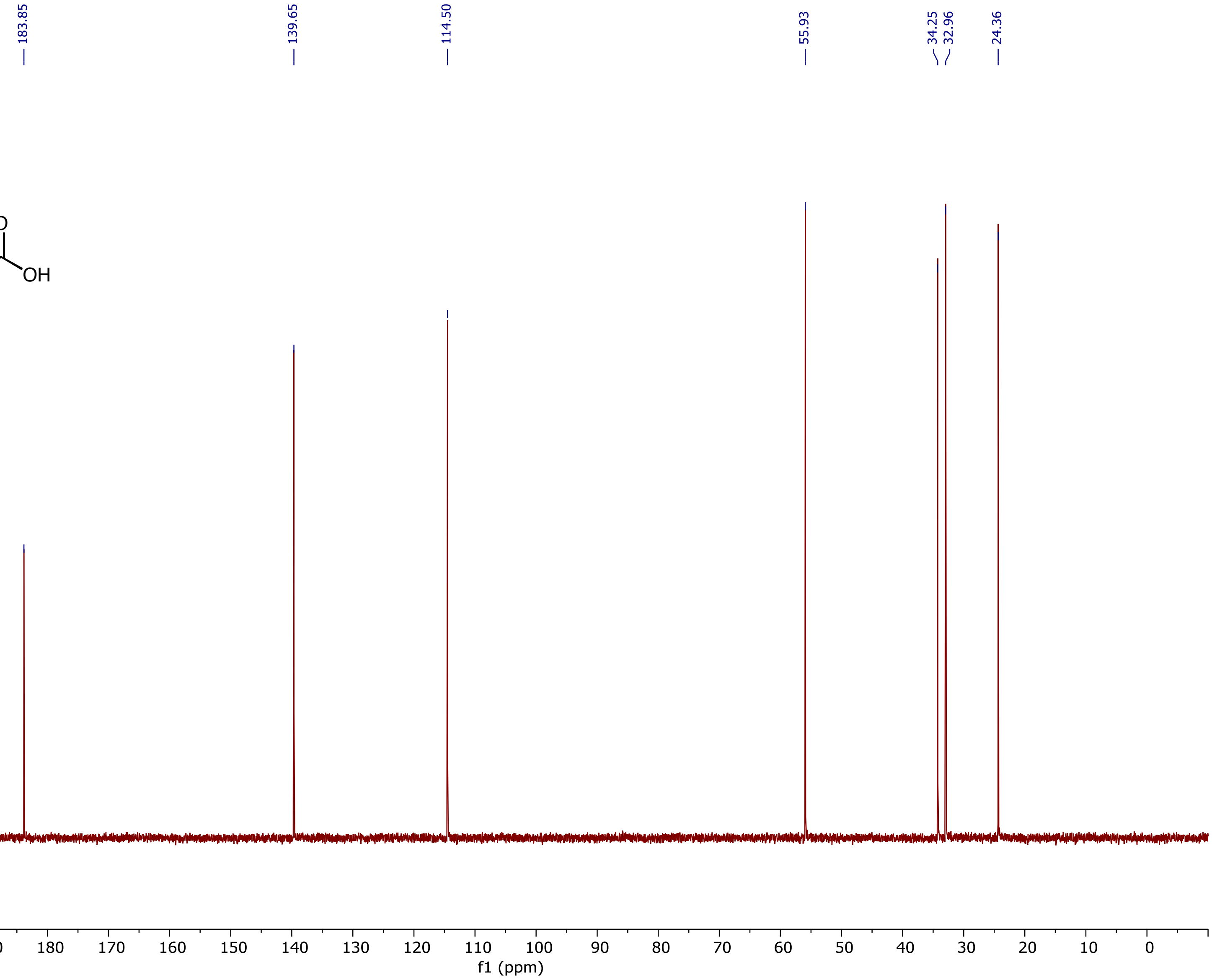

$^1$H NMR (500 MHz, MeOD) spectrum of 2-Amino-3-cyclopentylpropanoic acid **(5i)**

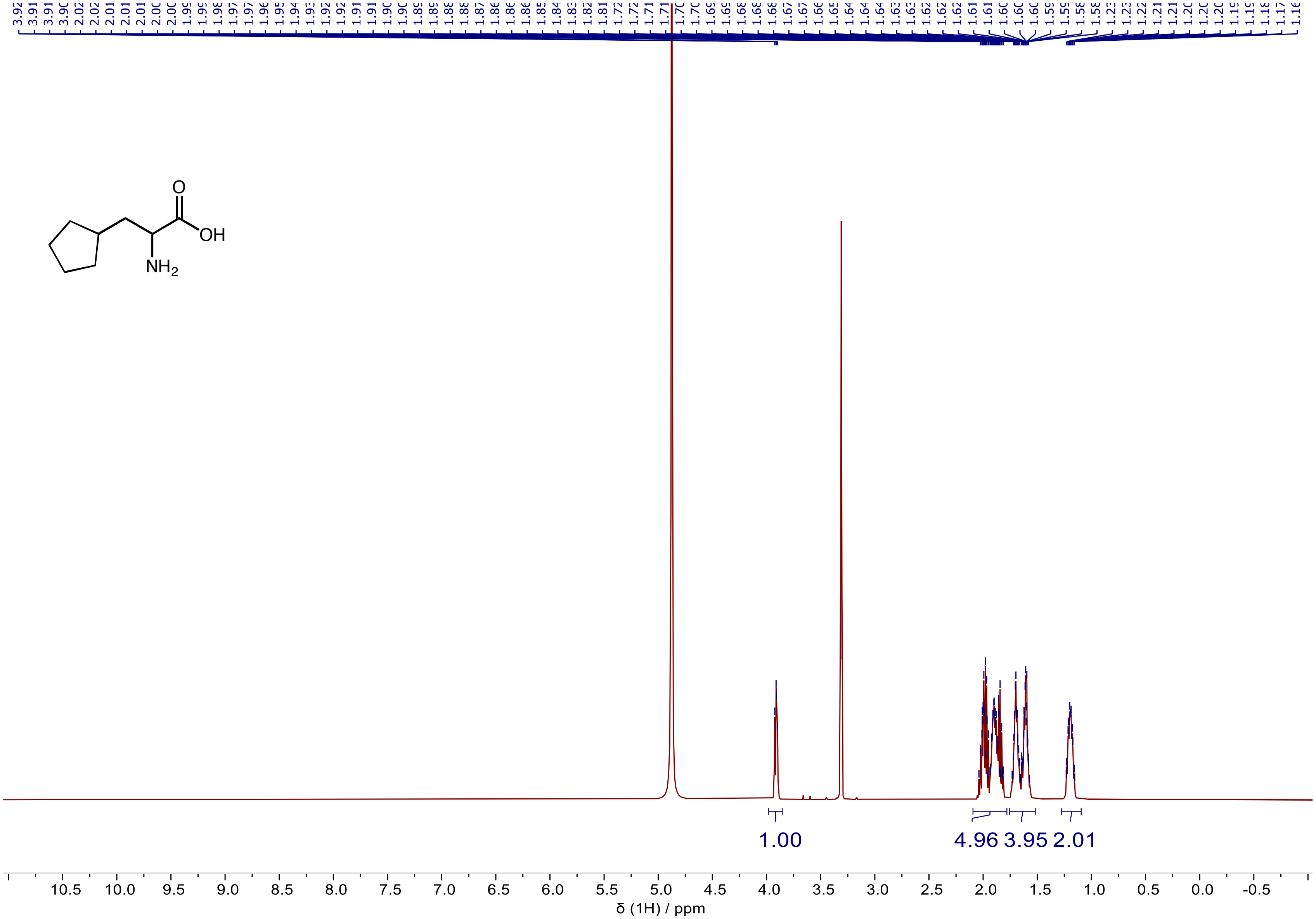

$^{13}C$ NMR (126 MHz, MeOD) spectrum of 2-Amino-3-cyclopentylpropanoic acid **(5i)**

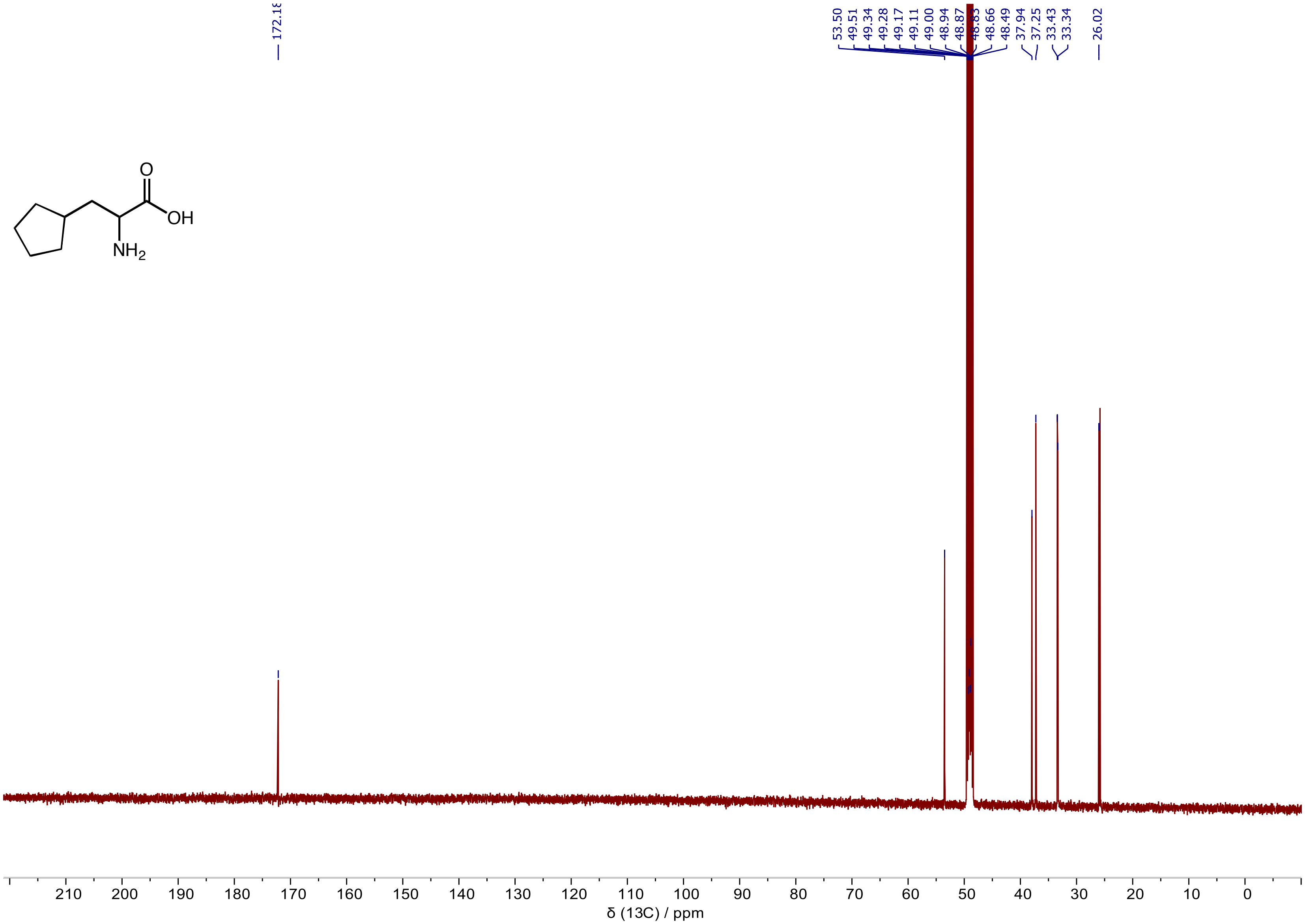

$^{1}H$ NMR (400 MHz, $D_2O$) spectrum of 2-Aminopentanoic acid **(5k)**

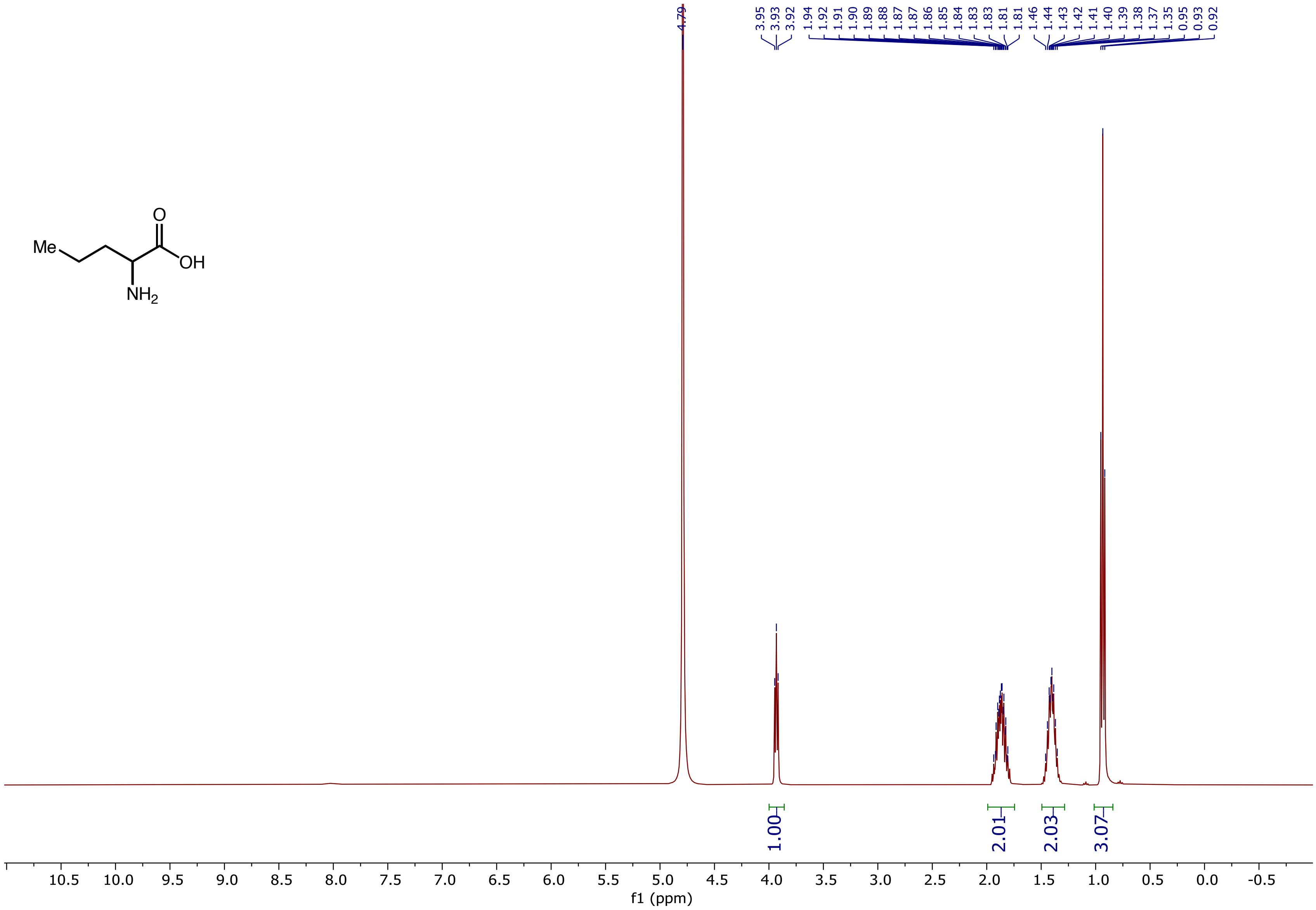

[13]C NMR (101 MHz, D2O) spectrum of 2-Aminopentanoic acid **(5k)**

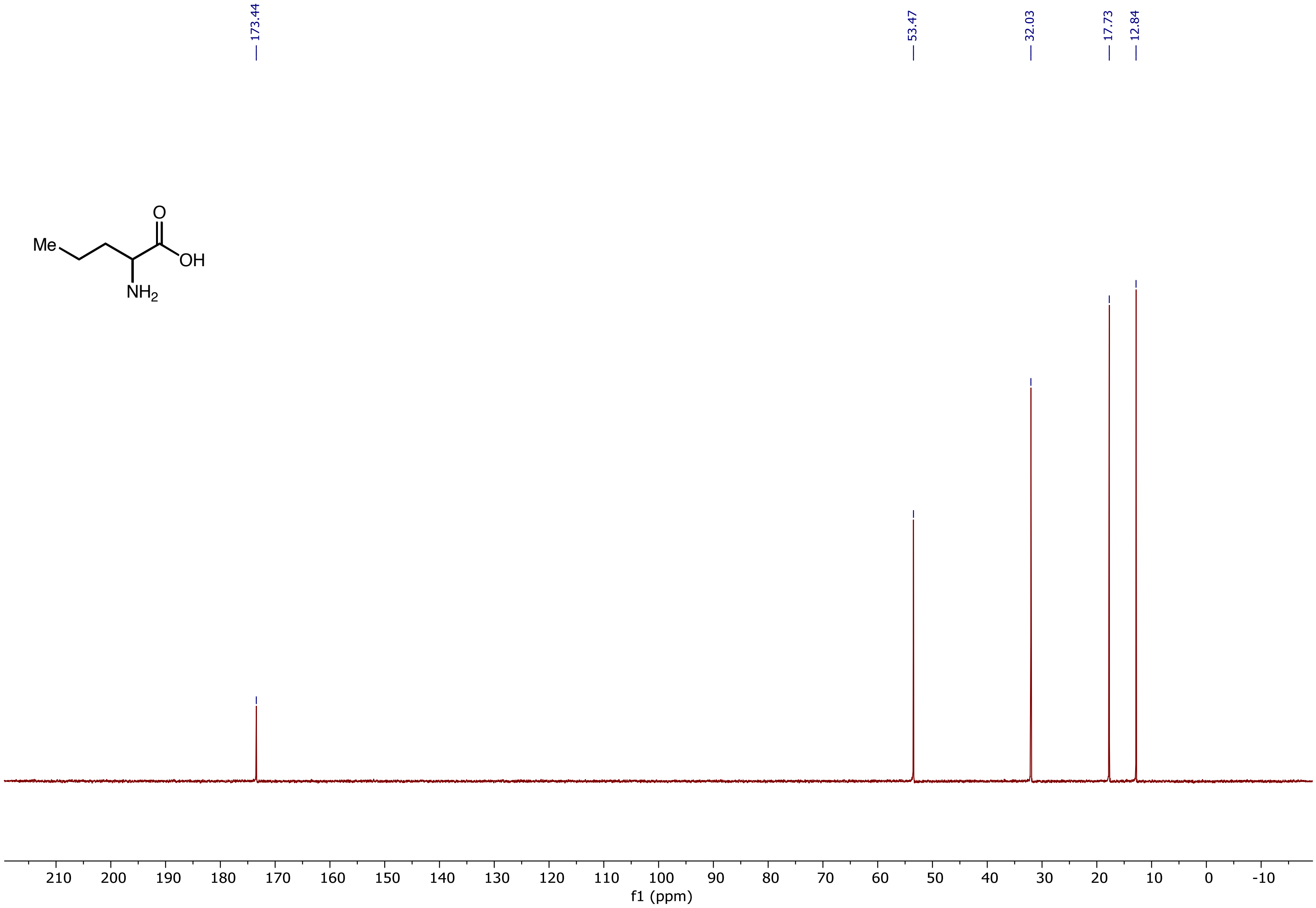

[1]H NMR (400 MHz, $D_2O$) spectrum of 2-Aminohexanoic acid **(5l)**

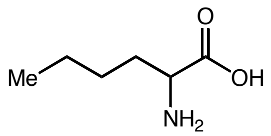

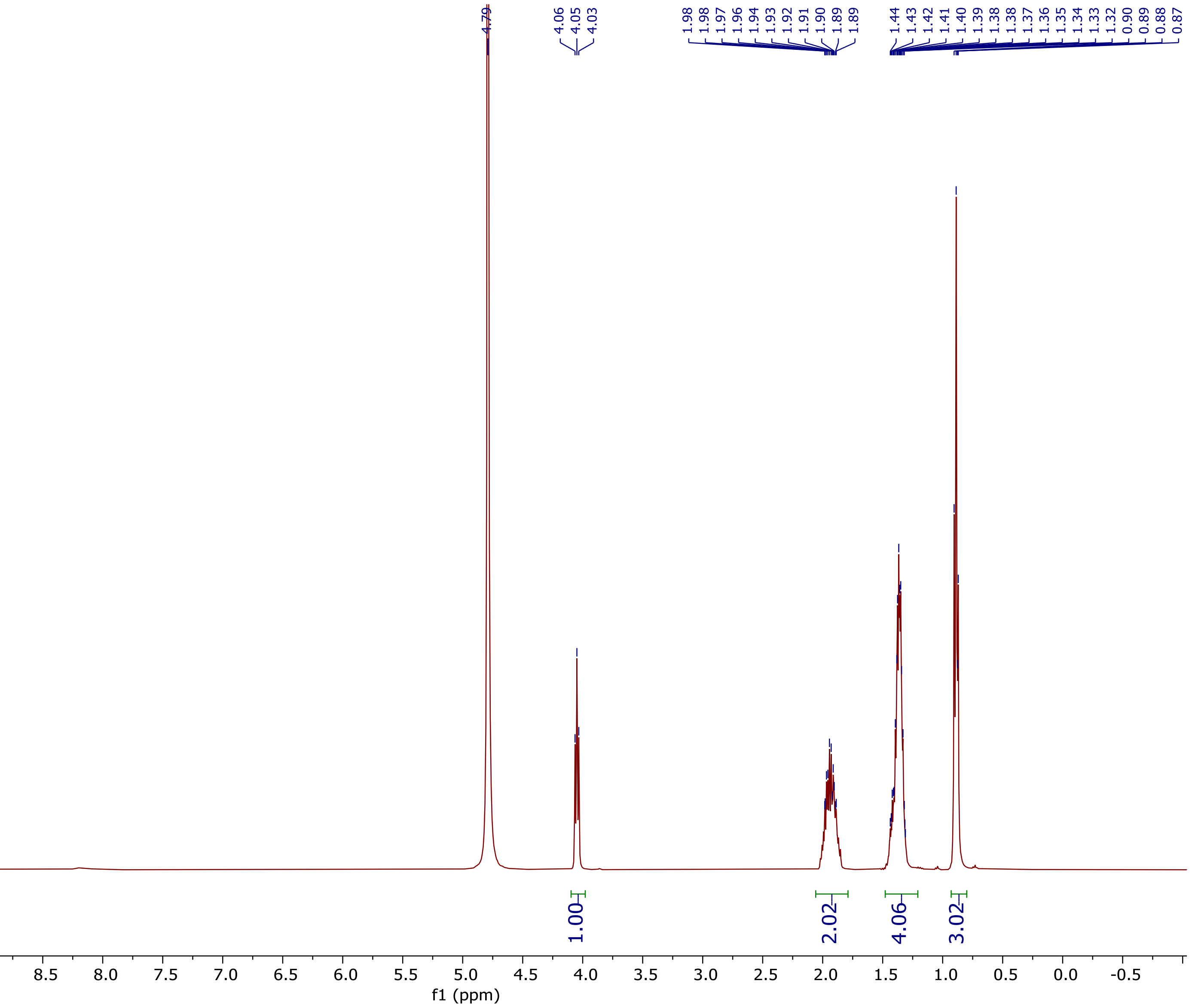

[13]C NMR (101 MHz, $D_2O$) spectrum of 2-Aminohexanoic acid **(5l)**

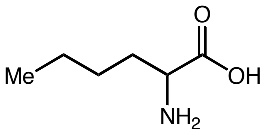

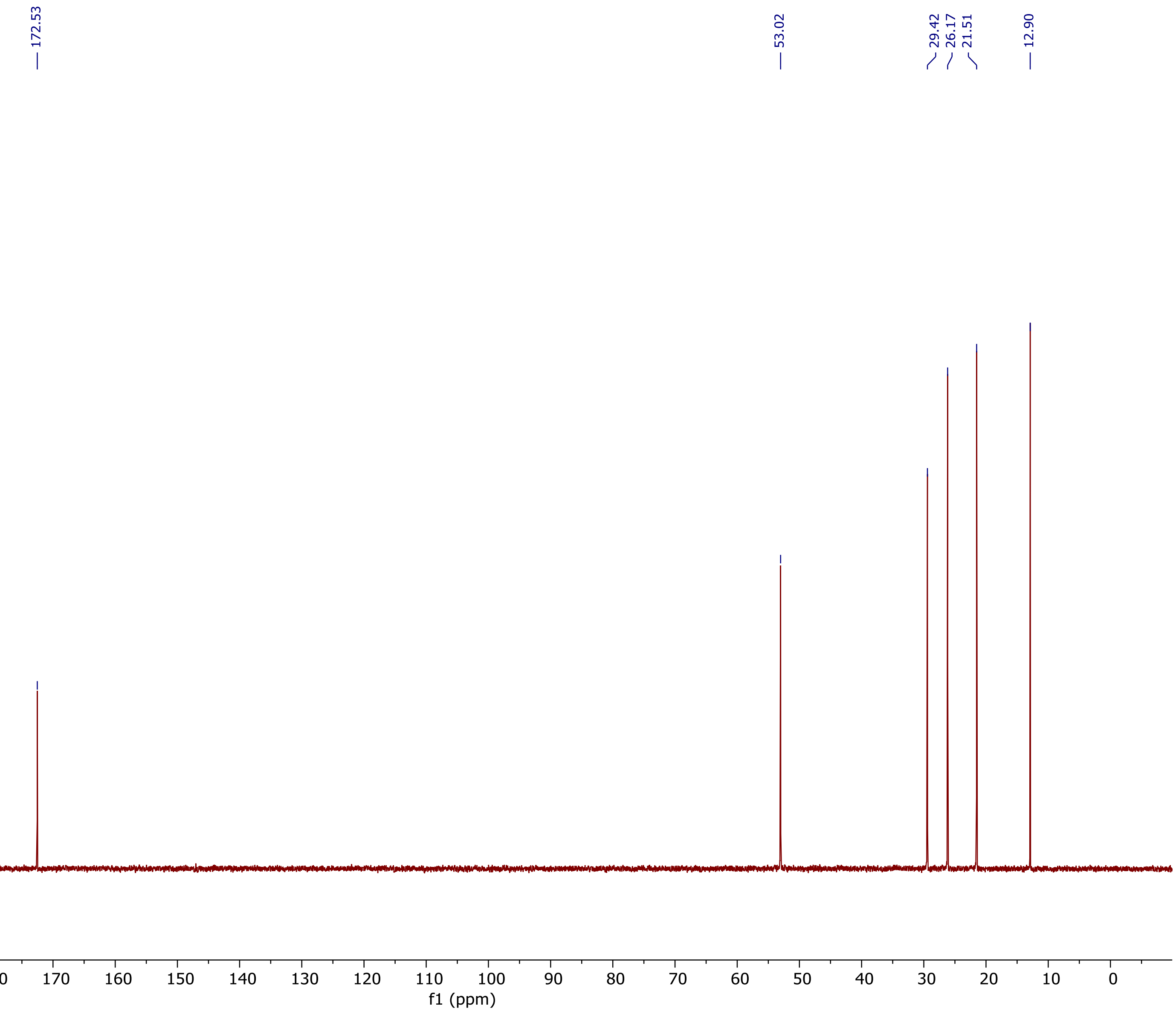

[1]H NMR (400 MHz, $D_2O$) spectrum of 2-Aminoheptanoic acid **(5m)**

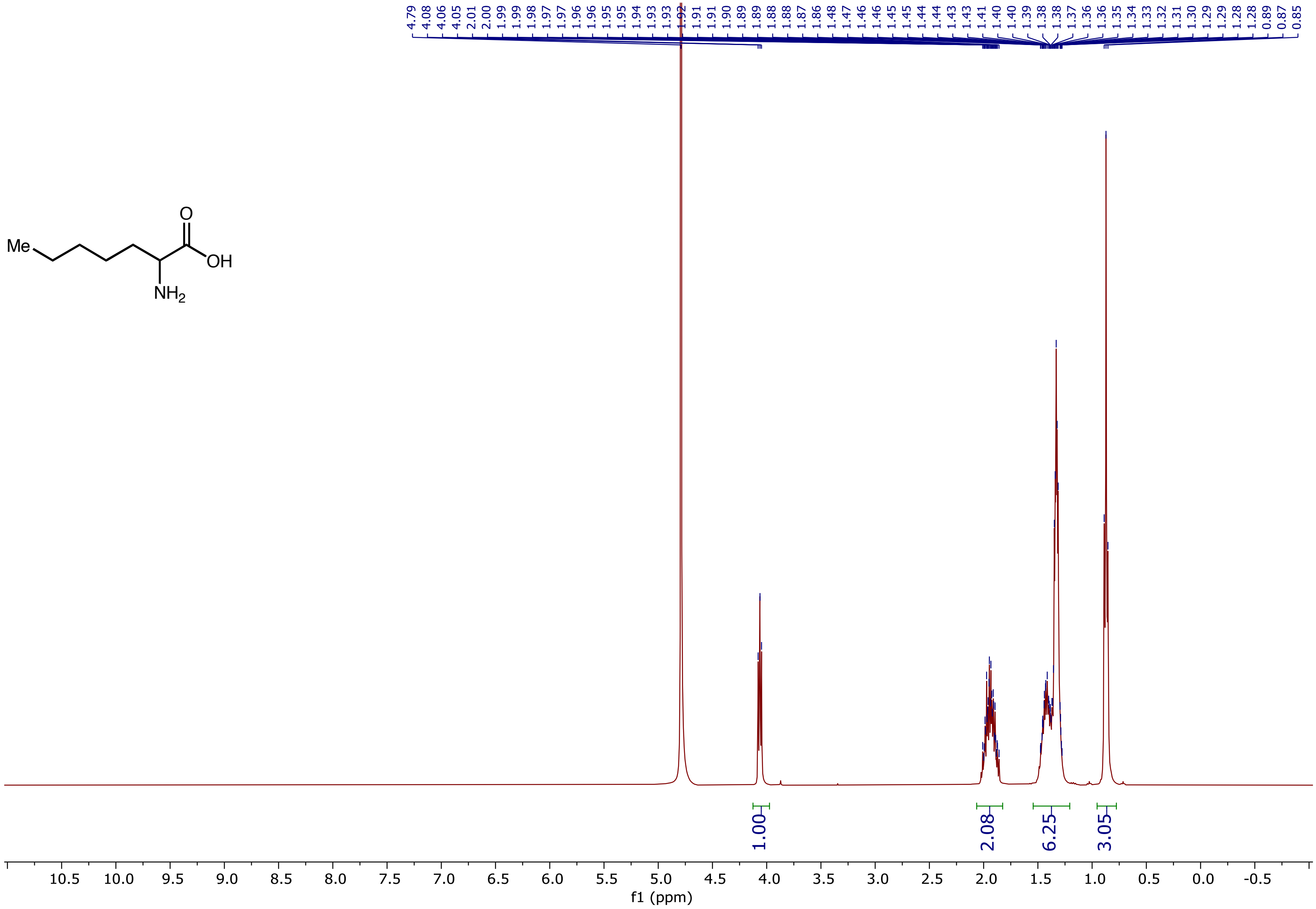

$^{13}C$ NMR (101 MHz, $D_2O$) spectrum of 2-Aminoheptanoic acid **(5m)**

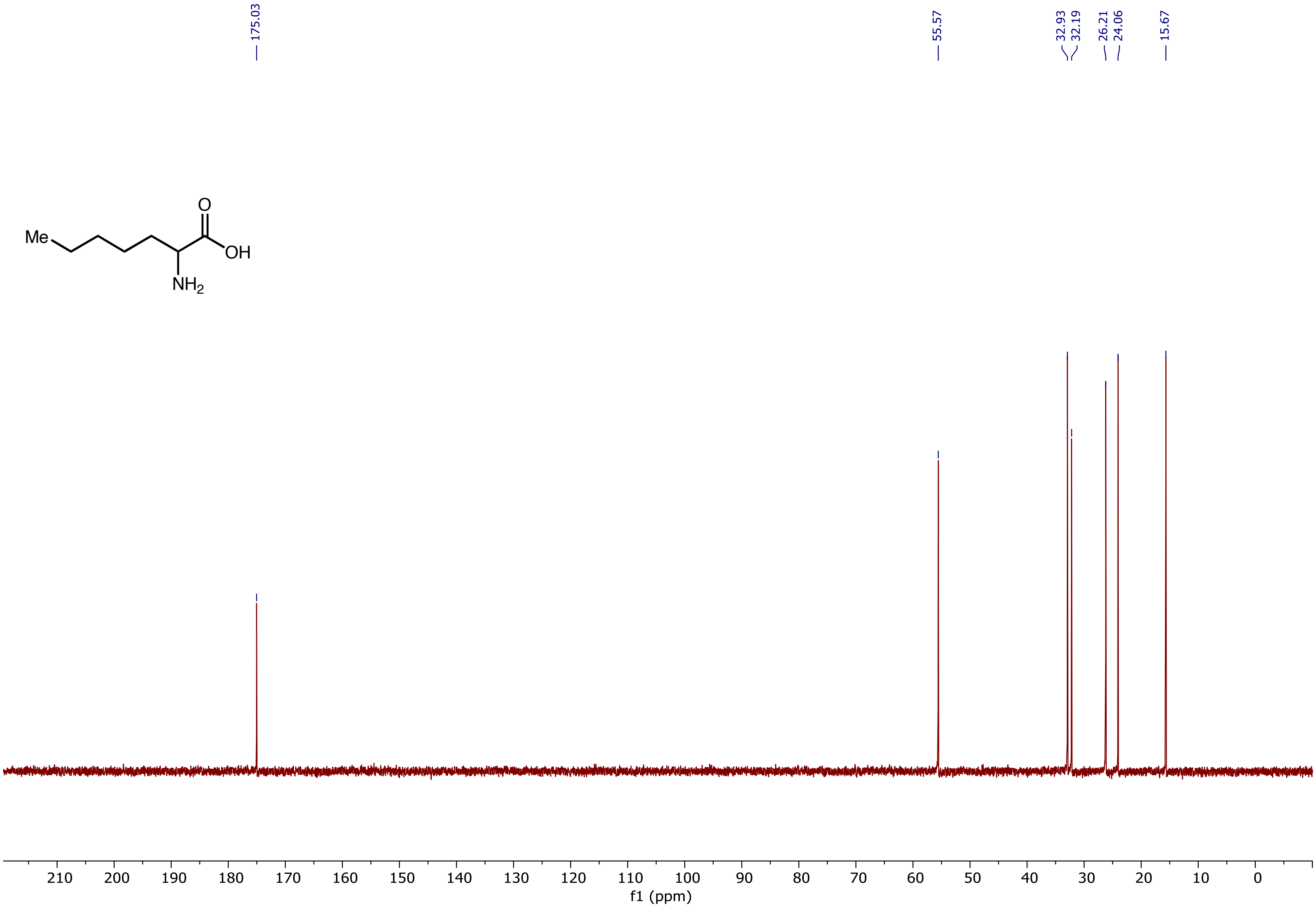

$^{1}H$ NMR (400 MHz, $D_2O$) spectrum of 2-Aminononanoic acid **(5n)**

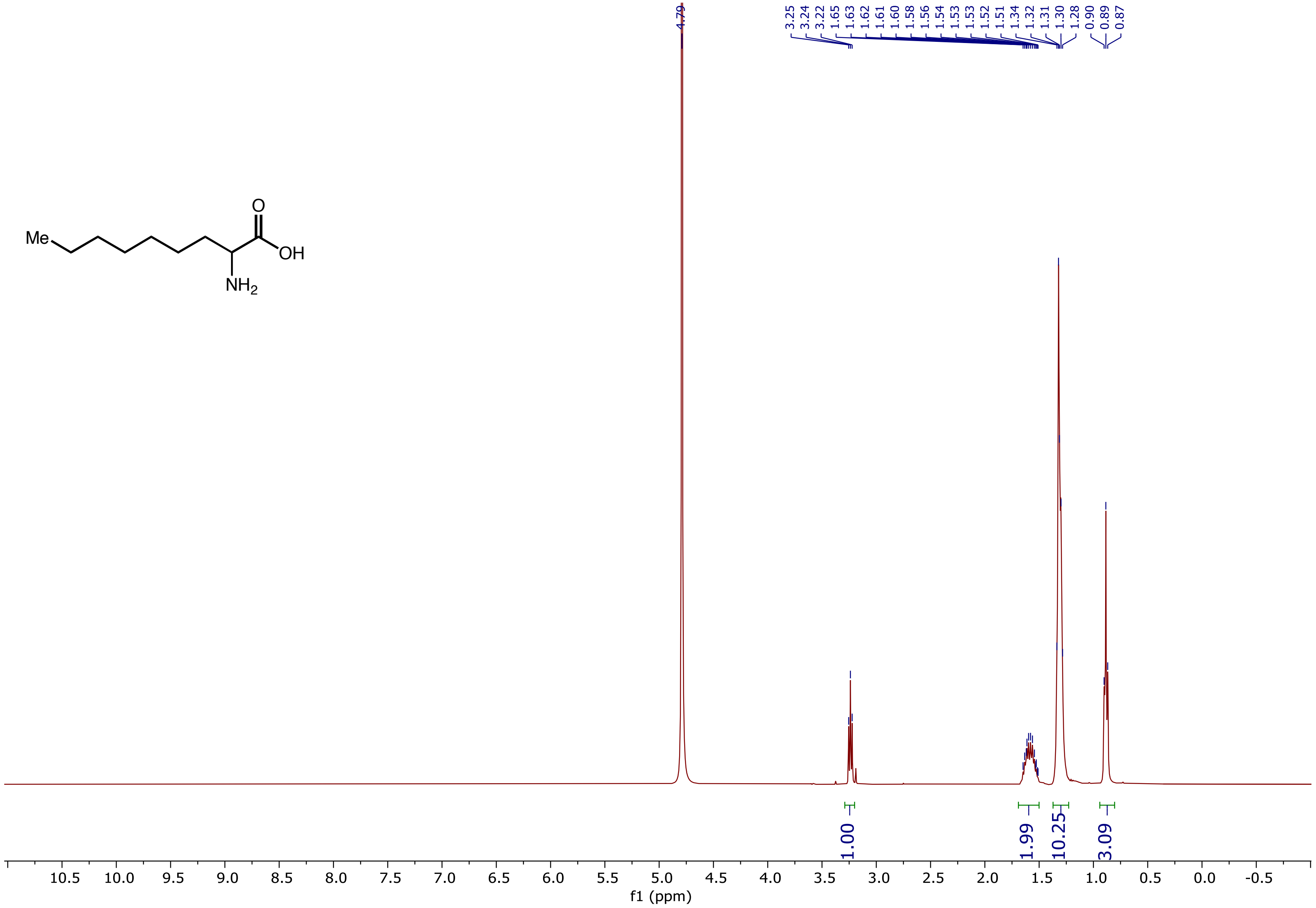

$^{13}C$ NMR (101 MHz, $D_2O$) spectrum of 2-Aminononanoic acid **(5n)**

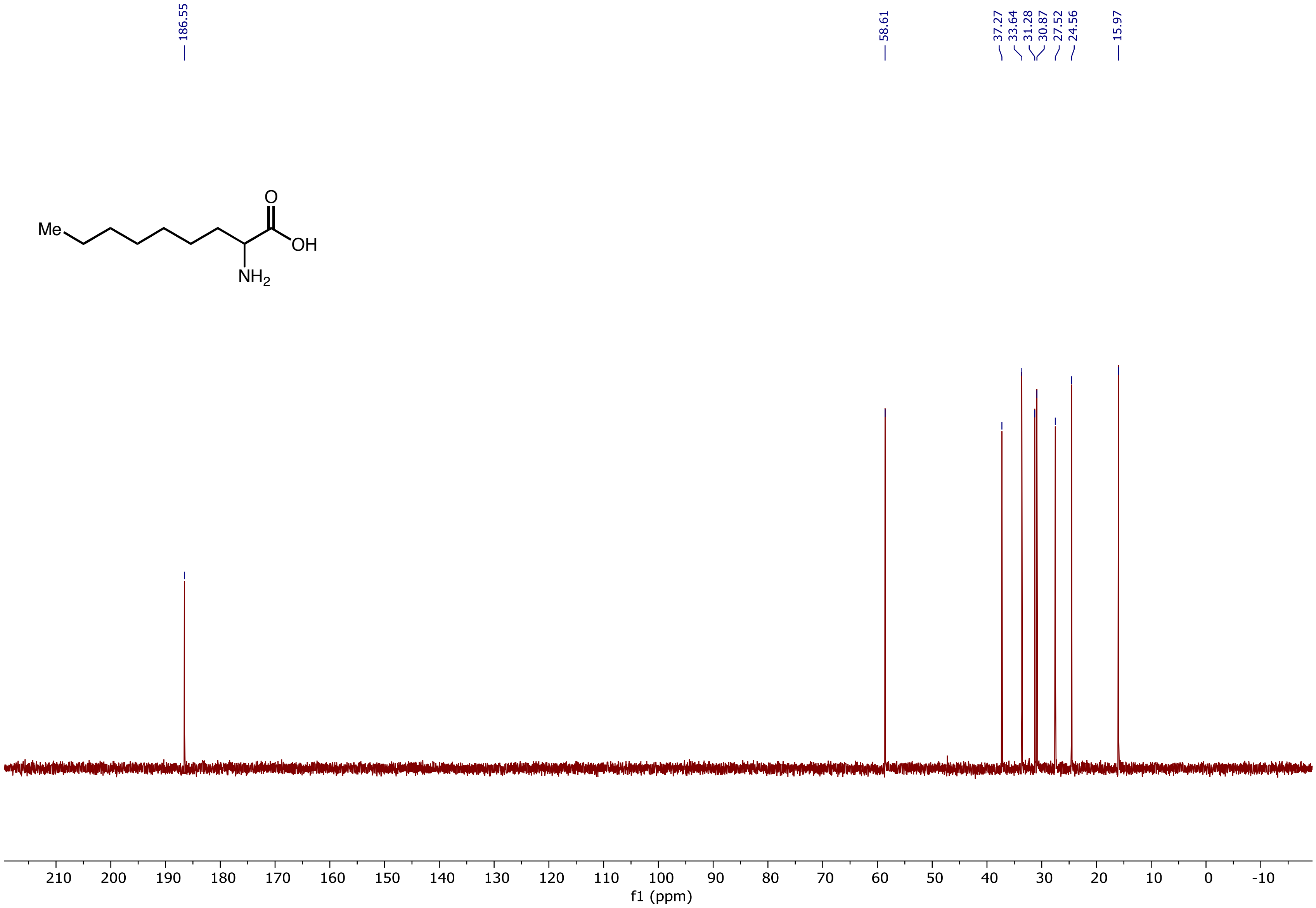

$^1H$ NMR (400 MHz, $D_2O$) spectrum of 2-amino-4-methyloctanoic acid **(5n')**

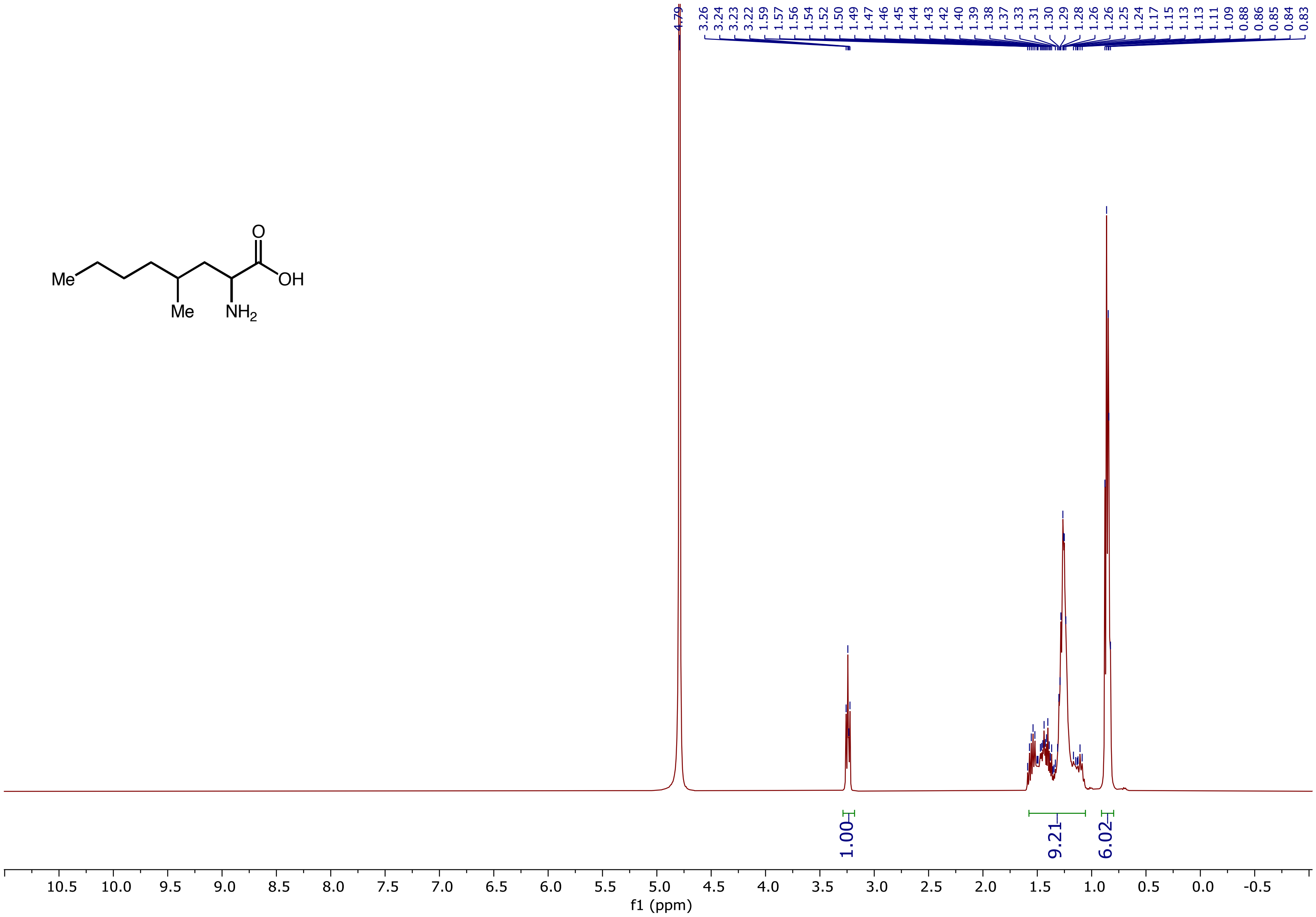

$^{13}C$ NMR (101 MHz, $D_2O$) spectrum of 2-amino-4-methyloctanoic acid **(5n')**

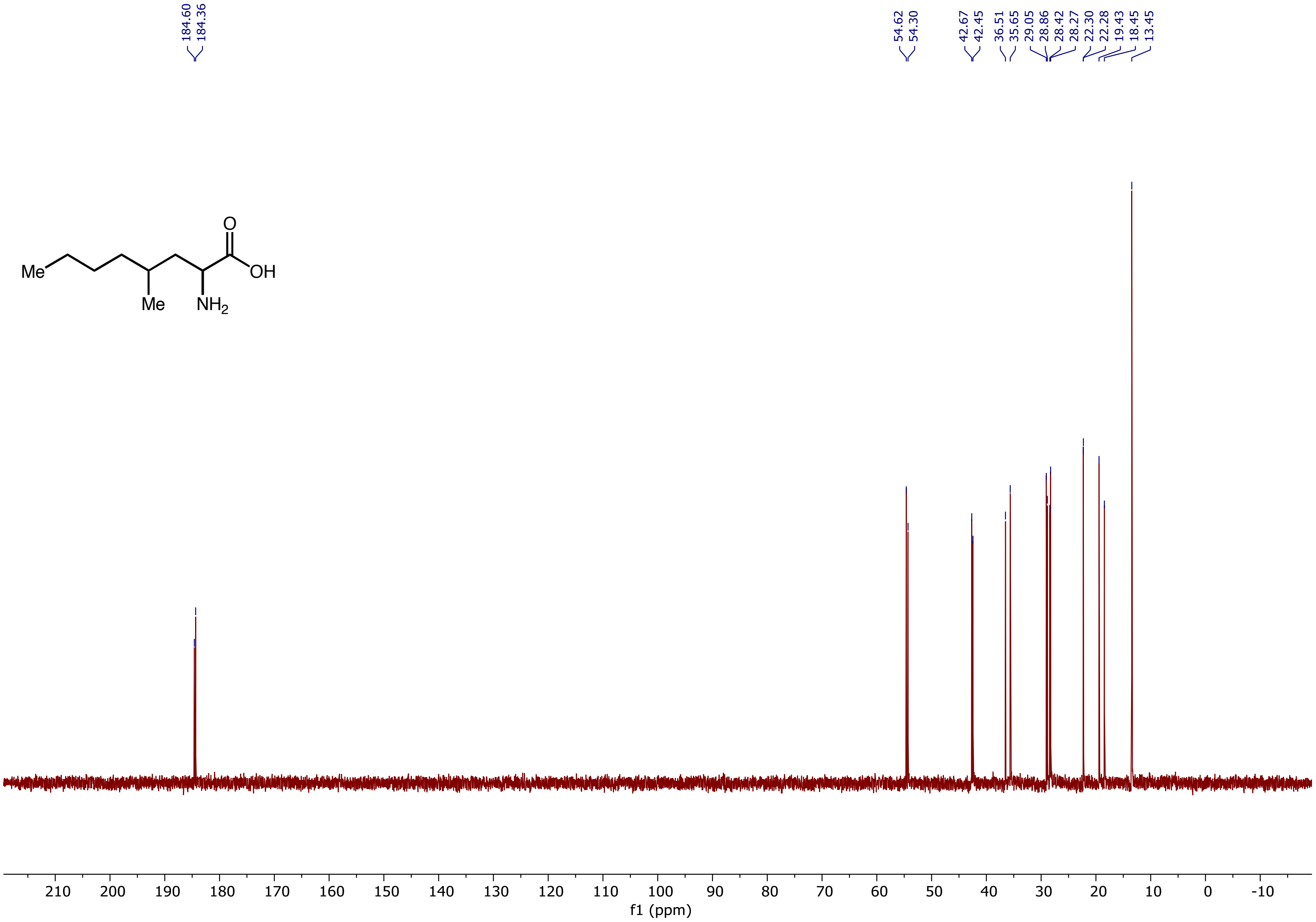

[1]H NMR (400 MHz, $D_2O$) spectrum of 2-Aminoundecanoic acid **(5o)**

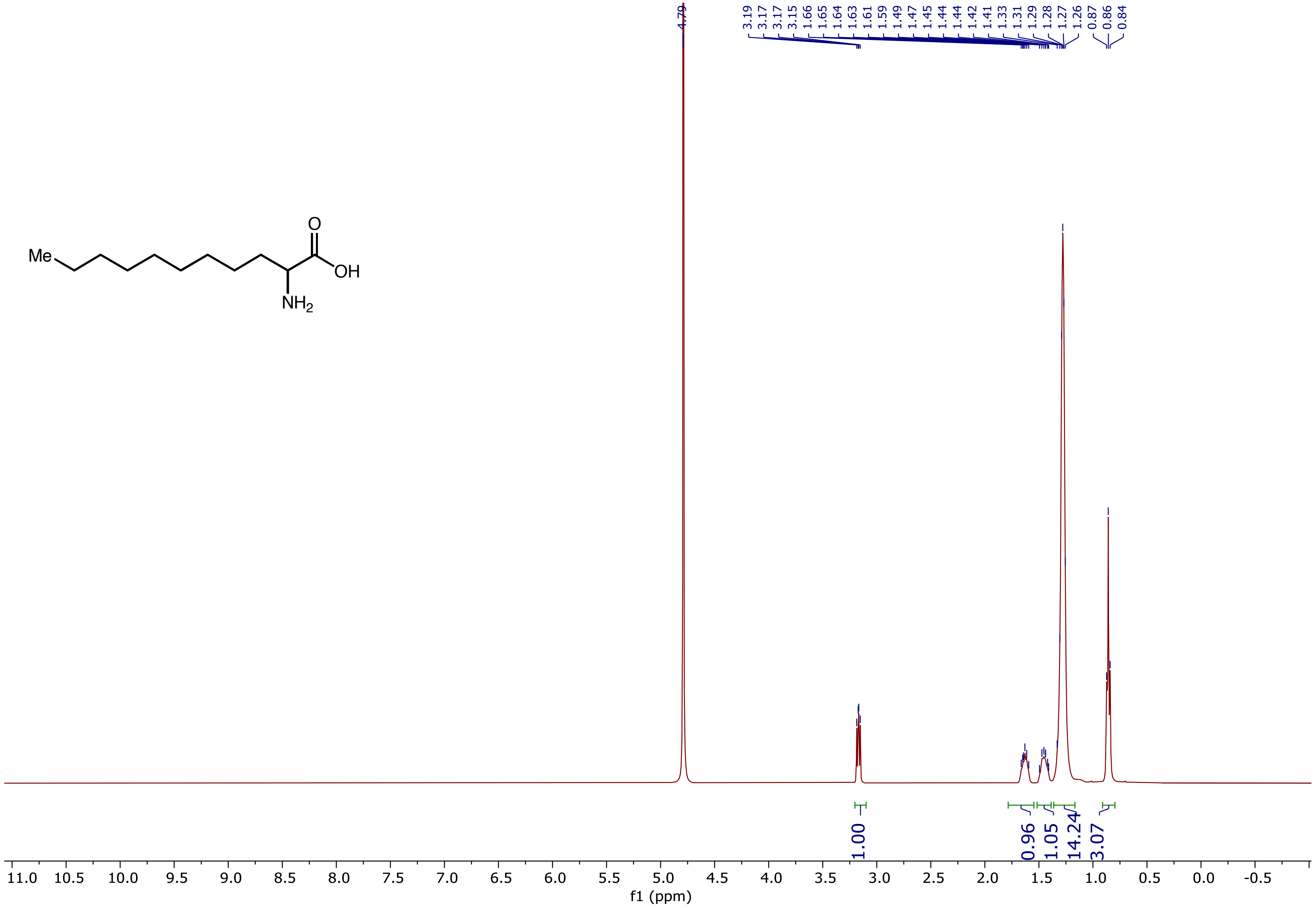

$^{13}C$ NMR (101 MHz, $D_2O$) spectrum of 2-Aminoundecanoic acid **(5o)**

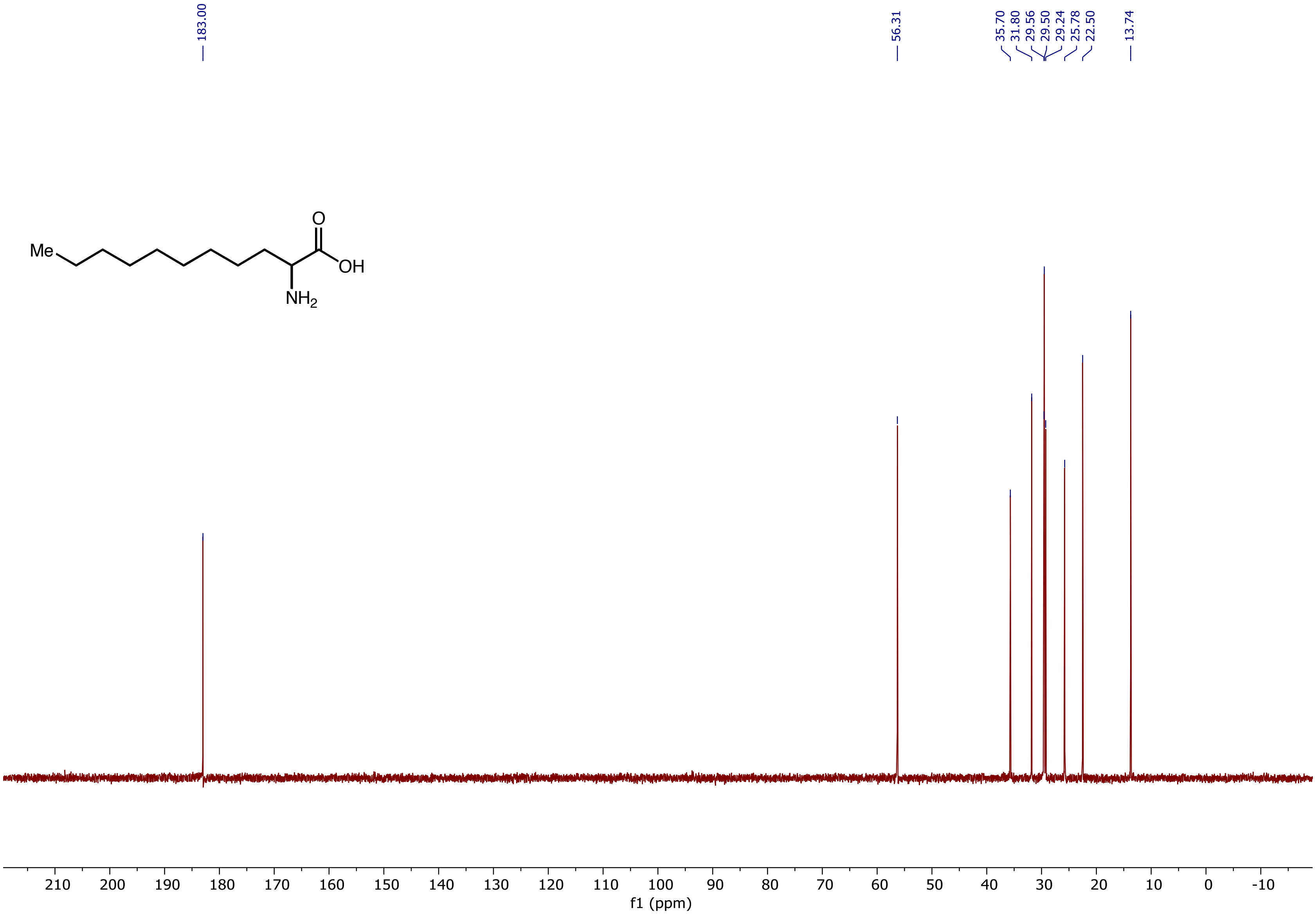

$^{1}H$ NMR (400 MHz, $D_2O$) spectrum of 2-Aminotridecanoic acid **(5p)**

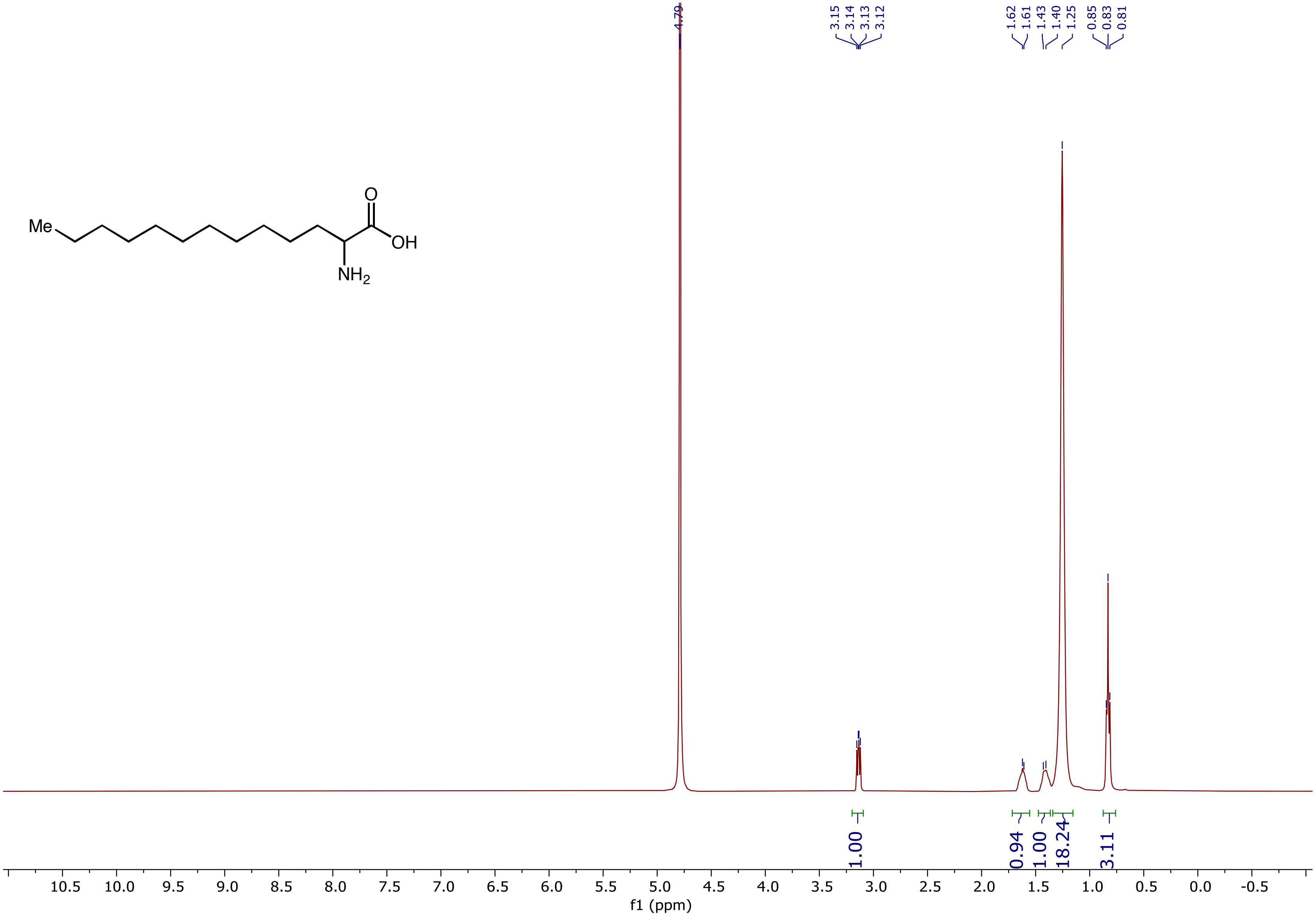

$^{13}C$ NMR (101 MHz, $D_2O$) spectrum of 2-Aminotridecanoic acid **(5p)**

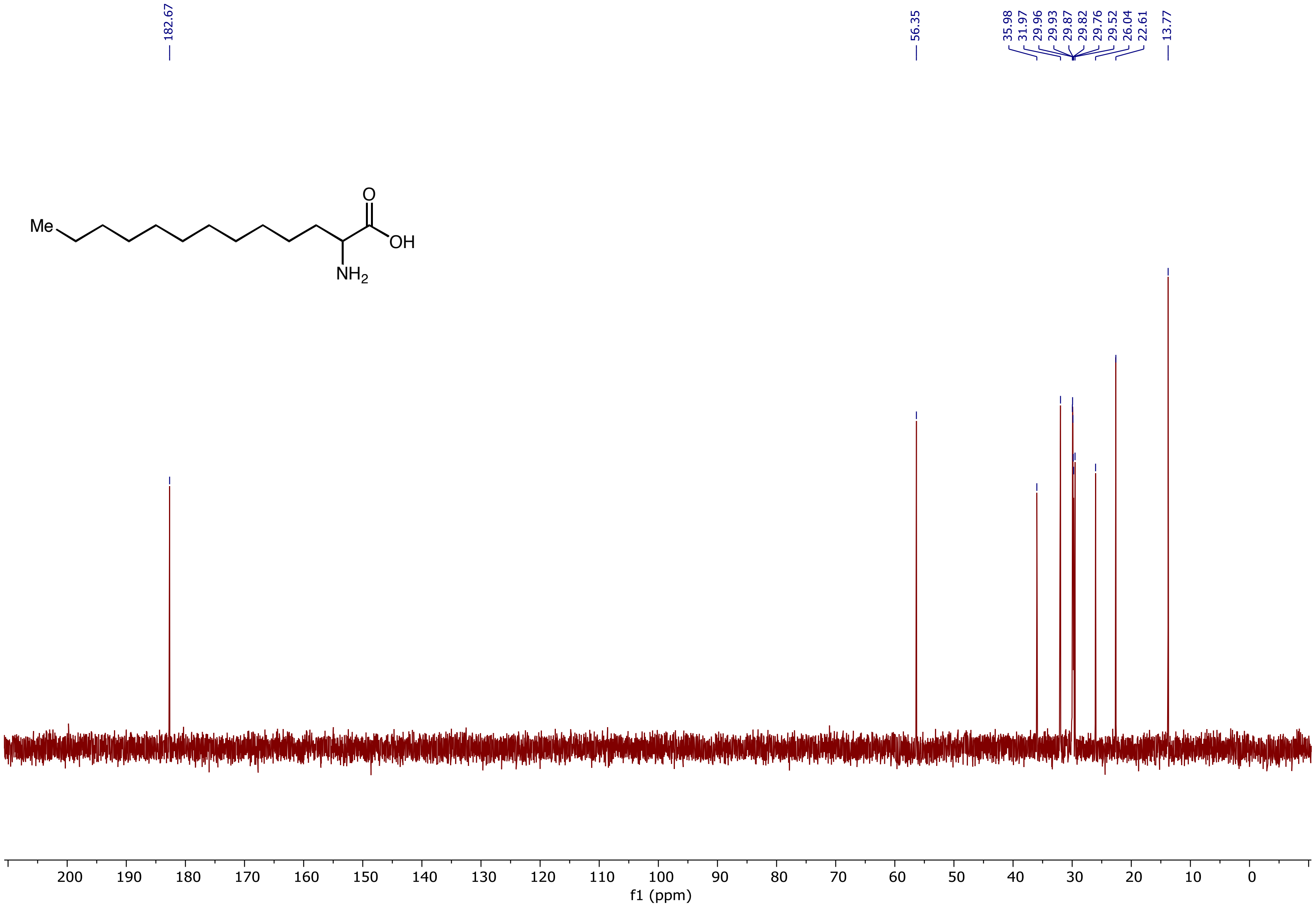

[1]H NMR (400 MHz, $D_2O$) spectrum of 2-Amino-4-cyclopropylbutanoic acid **(5q)**

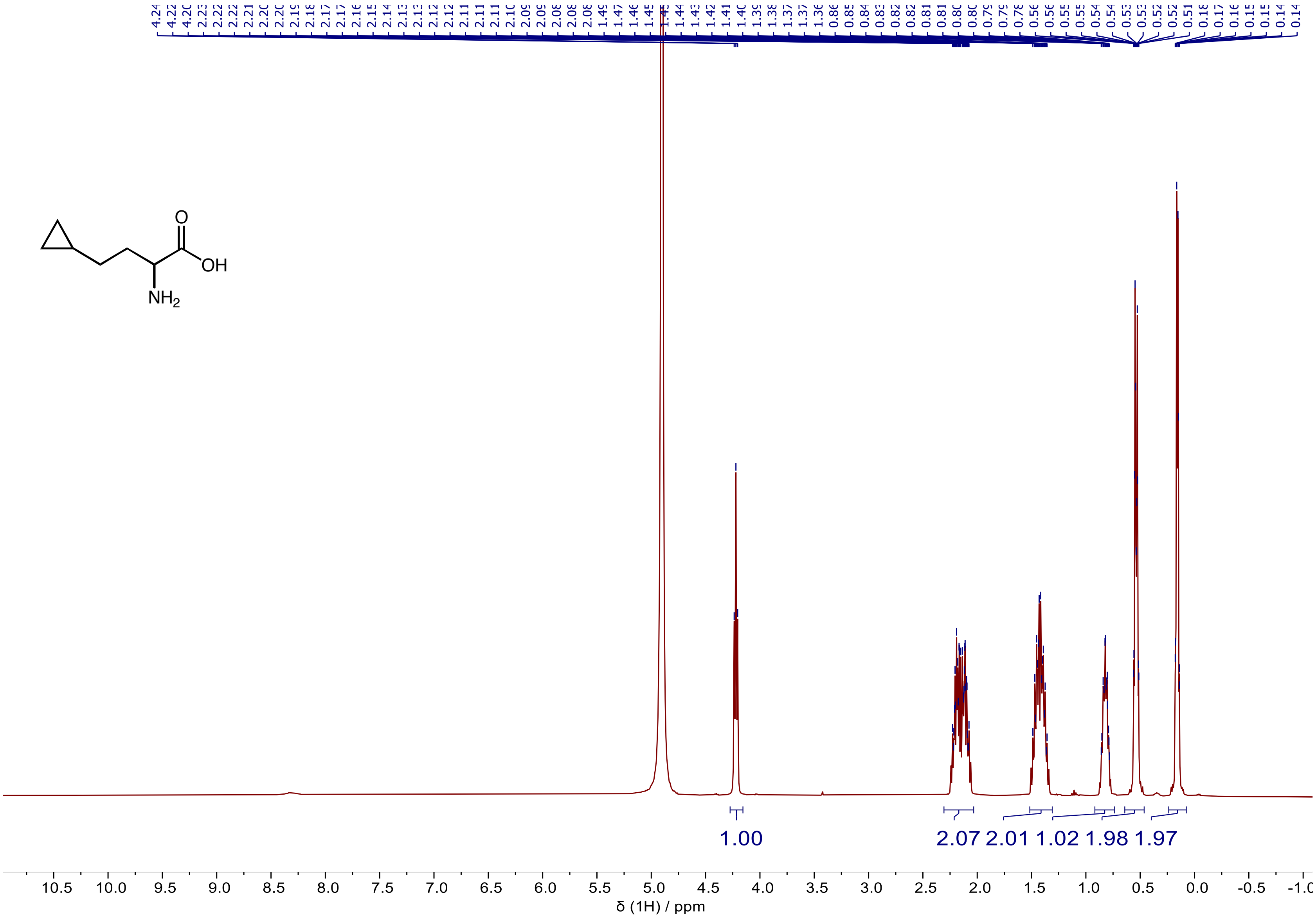

$^{13}$C NMR (101 MHz, $D_2O$) spectrum of 2-Amino-4-cyclopropylbutanoic acid **(5q)**

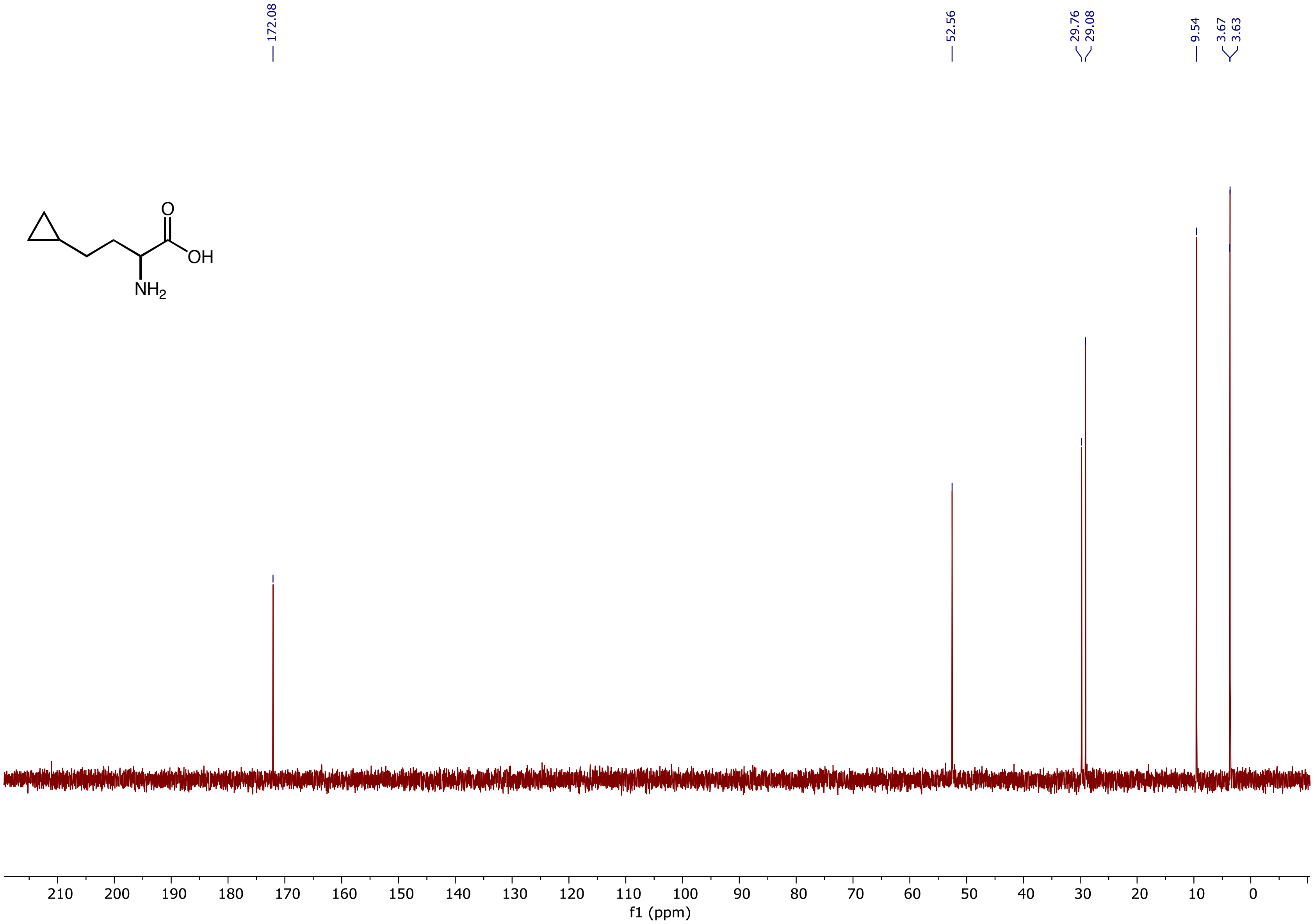

[1]H NMR (400 MHz, $D_2O$) spectrum of 2-Amino-4-cyclopentylbutanoic acid **(5r)**

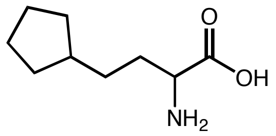

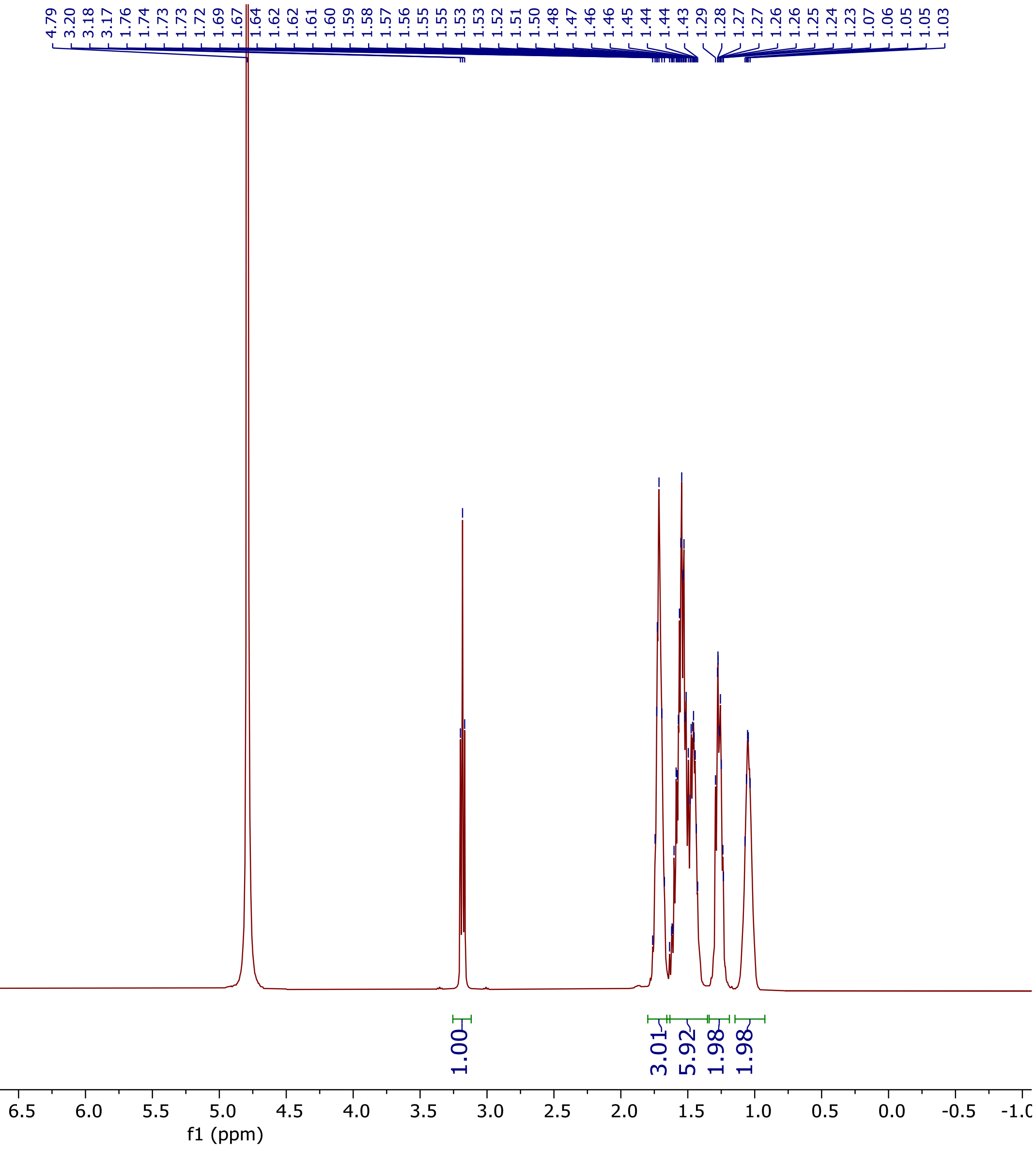

$^{13}C$ NMR (101 MHz, $D_2O$) spectrum of 2-Amino-4-cyclopentylbutanoic acid **(5r)**

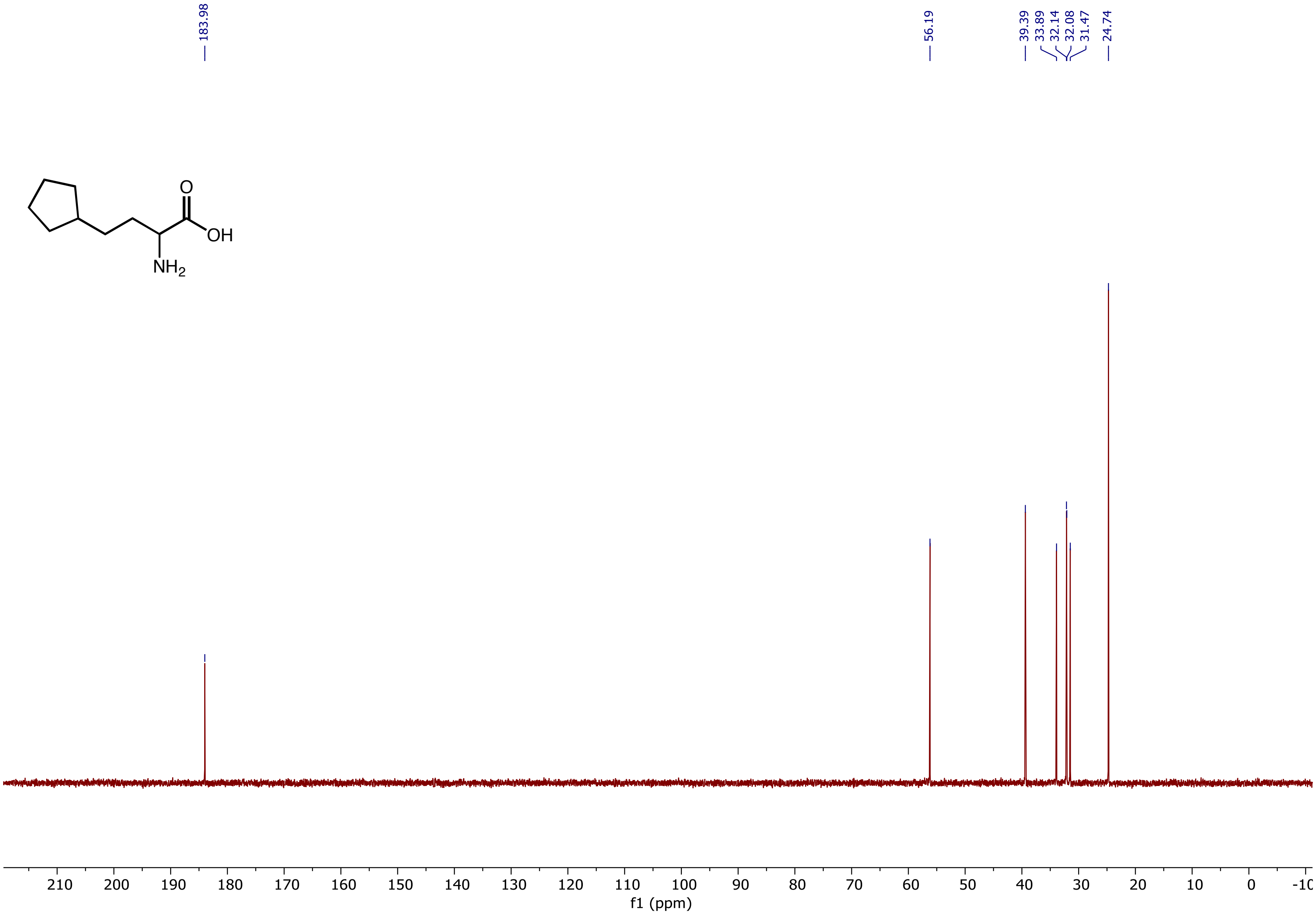

$^{1}H$ NMR (400 MHz, $D_2O$) spectrum of 2-Aminonon-8-enoic acid **(5s)**

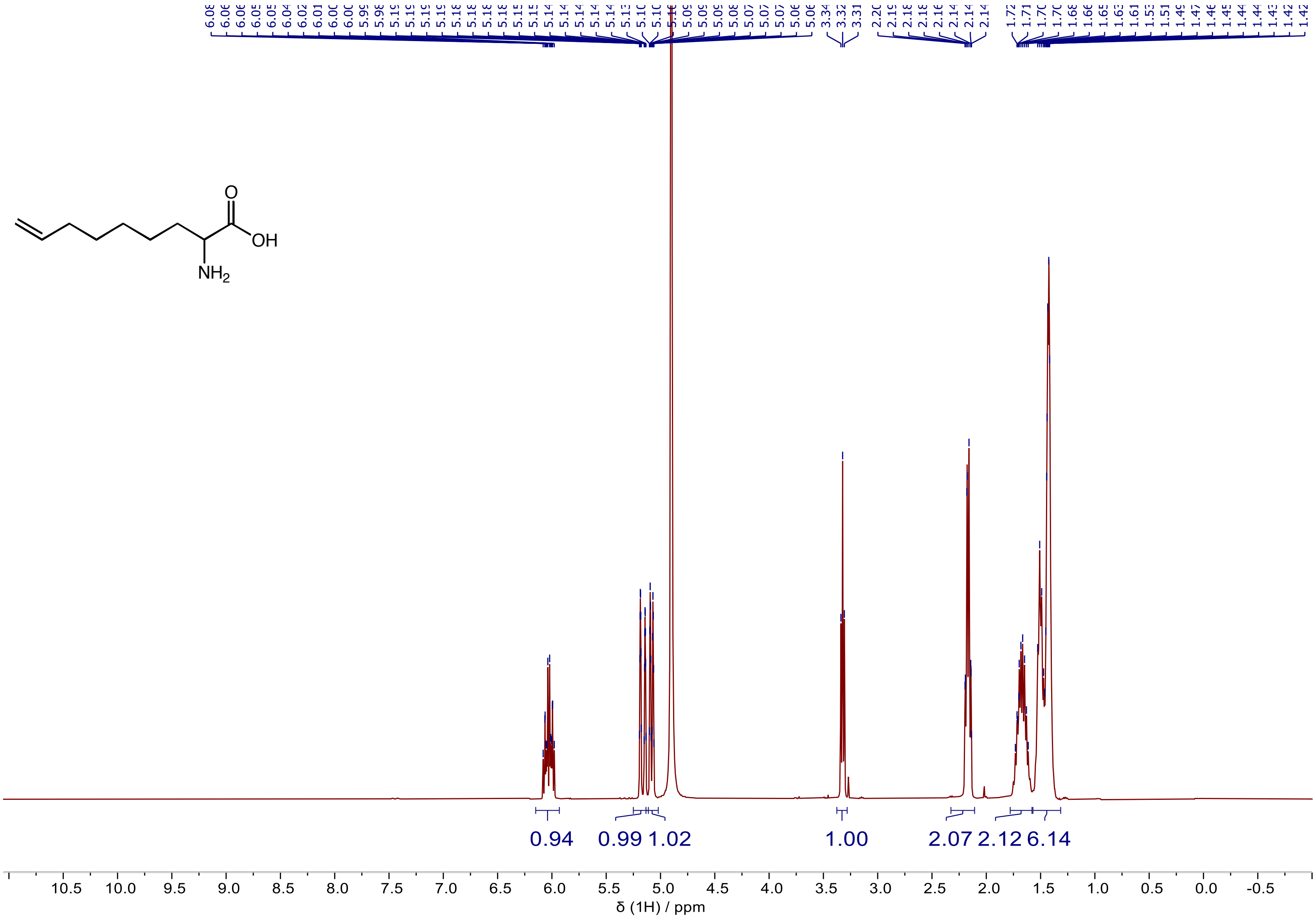

$^{13}$C NMR (101 MHz, $D_2O$) spectrum of 2-Aminonon-8-enoic acid **(5s)**

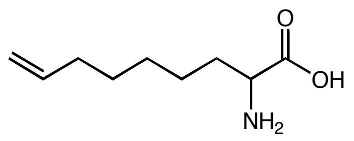

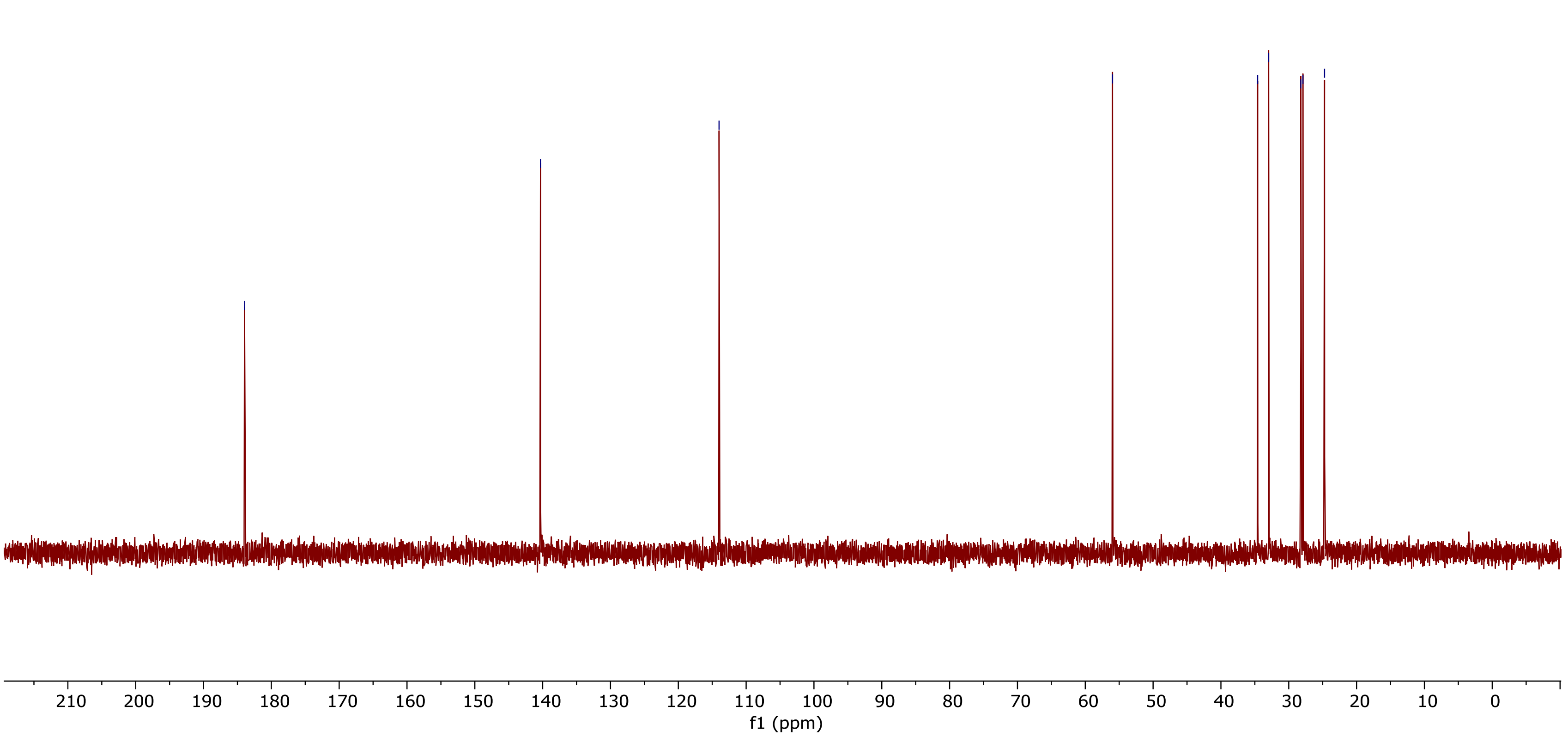
